# Novel ‘GaEl antigenic patches’ identified by ‘reverse epitomics’ approach to design multi-patch vaccines against NIPAH infection, a silent threat to global human health

**DOI:** 10.1101/2022.09.09.507124

**Authors:** Sukrit Srivastava, Michael Kolbe

**Author notes:** Correspondence should be addressed to S.S. or M.K.

## Abstract

**Background:** Nipah virus (NiV) is a zoonotic virus that causes lethal encephalitis and respiratory disease with the symptom of endothelial cell-cell fusion. Several NiV outbreaks have been reported since 1999 with nearly annual occurrences in Bangladesh. The outbreaks had high mortality rates ranging from 40 to 90%. No specific vaccine has yet been reported against NiV.

**Methodology:** Recently, several vaccine candidates and different designs of vaccines composed of epitopes against NiV were proposed. Most of the vaccines target single protein or protein complex subunits of the pathogen. The Multi-epitope vaccines proposed also cover a largely limited number of epitopes and hence their efficiency is still pending. To address the urgent need for a specific and effective vaccine against NiV infection in the present study, we have utilized the ‘Reverse Epitomics’ approach (“overlapping-epitope-clusters-to-patches” method) to identify ‘antigenic patches’ (Ag-Patches) and utilize them as immunogenic composition for Multi-Patch vaccine (MPV) design. The designed MPVs were analyzed for immunologically crucial parameters, physiochemical properties and interaction with Toll-like receptor 3 ectodomain.

**Results:** In total 30 CTL (Cytotoxic T lymphocyte) and 27 HTL (Helper T lymphocytes) antigenic patches were identified from the entire NiV proteome based on the clusters of overlapping epitopes. These identified Ag-Patches cover a total of discreet 362 CTL and 414 HTL epitopes from entire proteome of NiV. The antigenic patches were utilized as immunogenic composition for the design of two CTL and two HTL multi-patch vaccines. The 57 antigenic patches utilized here cover 776 overlapping epitopes targeting 52 different HLA class I and II alleles providing a global ethnically distributed human population coverage of 99.71%. Such large number of epitope coverage resulting in large human population coverage cannot be reached with single protein/subunit or multi-epitope based vaccines. The reported antigenic patches also provide potential immunogenic composition for early detection diagnostic kits for NiV infection. Further, all the MPVs & Toll-Like Receptor ectodomain complexes show stable nature of molecular interaction with numerous hydrogen bonds, salt bridges and non-bounded contacts formation and acceptable root mean square deviation and fluctuation. The cDNA analysis show a favorable large scale expression of the MPV constructs in human cell line.

**Conclusion:** By utilizing the novel ‘Reverse epitomics’ approach highly immunogenic novel ‘GaEl antigenic patches’ (GaEl Ag-Patches) a synonym term for ‘antigenic patches’, were identified and utilized as immunogenic composition to design four MPVs against NiV. We conclude that the novel Multi-Patch Vaccines is a potential candidate to combat NiV, with greater effectiveness, high specificity and large human population coverage worldwide.

## INTRODUCTION

Nipah virus (NiV) is a zoonotic virus of the genus Henipavirus and family Paramyxoviridae^1^. The first NiV outbreak was reported amongst pigs in Malaysia in 1999^2^. Later another NiV outbreak was reported from Meherpur, Bangladesh in 2001, this time in humans. The transmission of NiV infection in Bangladesh and India was associated with contaminated date palm sap and human-to-human contact^3^. Bats were identified as the main reservoir for NiV and they are responsible for the transmission of the infection to both humans and animals^4^. Since 2001 NiV outbreaks were reported for Bangladesh almost every year (2003-05, 2007-12). In India two NiV outbreaks were reported in the state of West Bengal in 2001 and 2007^5^. Afterwards another NiV outbreak was reported from the state of Kerala in India during the period of May to June-2018. The Kerala outbreak claimed 17 human lives leaving only two survivors out of 19 confirmed cases^6^. Hitherto, no efficient vaccine against NiV has been reported. Vaccines targeting multiple proteins of NiV might provide efficient protection and their potential needs to be explored in the future.

The essential proteins involved in NiV pathogenesis to human include C protein, Fusion glycoproteins (F), Glycoproteins (G), Matrix proteins (M), Nucleocapsid protein, Phosphoprotein, Polymerase, V protein and the W protein^7–21^. The C protein regulates the early host pro-inflammatory response as well as the pathogen virulence, the glycoprotein (G), the fusion protein (F) and the matrix protein (M) together form a cluster on the host human cell membrane and facilitate virus particle assembly formation^22,23^. The NiV Polymerase is essentially responsible for the initiation of RNA synthesis, primer extension, and transition to elongation mode. The phosphoprotein and the glycoprotein are also essentially involved viral replication while the V protein is responsible for the host interferon (IFN) signaling evasion^17–20^. The identical N-terminal region of the V and W proteins were found to be sufficient to exert the IFN-antagonist activity^21^. Hence, all the above mentioned nine NiV proteins play essential role in NiV proliferation and pathogenesis and so provide important drug and vaccine targets candidates.

In recent studies a number of B cell and T cell epitope candidates from different NiV proteins have been reported. Additionally, a number of vaccine design approaches and vaccine candidates including multi epitope vaccines were reported^24–38^. However, using a single or a small number of epitopes might be limiting the vaccine potential. The long-term adaptive immunity essentially involves the presentation of the antigen as short peptides (epitope) on the surface of Antigen Presenting Cells (APC). To achieve this presentation an intra cellular proteolytic chop down process is orchestrated by the proteasome and lysosome. The ‘Transporter associated with antigen processing’ (TAP) and further the HLA allele (human leukocyte antigen) molecules facilitate this epitope presentation^39,40^

Here, we introduce the term ‘GaEl antigenic patch’ (GaEl Ag-Patch) as a synonymous for ‘antigenic patch’. The prefix GaEl is derived from river Ganga and Elna of the inventors home countries. This is to acknowledge the inventors of ‘Reverse Epitomics’ and the prefix GaEl is derived from river Ganga and Elna of inventors home countries. The novel ‘Reverse Epitomics’ approach to identify ‘GaEl Ag-Patch’ has been introduced in our earlier study on SARS-CoV-2^41–44^. In the present study, we have utilized the said ‘reverse epitomics’ approach to identify ‘GaEl Ag-Patch’ from NiV proteins. The reverse epitomics approach applies the novel “overlapping-epitope-clusters-to-patches” method to identify the ‘GaEl Ag-Patches’. Here we first identify the overlapping epitopes which arise from a particular region of the protein. This particular region is a consensus peptide sequence of all the overlapping epitopes and we identify this region of protein as ‘GaEl Ag-Patches’. From the proteome of NiV here we report a total of 57 ‘GaEl Ag-Patches’ identified from overlapping 776 epitopes. Next, we utilized these ‘GaEl Ag-Patches’ from NiV proteome as immunogenic composition to design Multi-Patch Vaccine.

## METHODOLOGY

In the present study, we have designed two CTL and two HTL multi-patch vaccines (MPVs). These vaccines are composed of ‘GaEL antigenic patches’ (GaEl Ag-Patches) identified from proteome of NiV. The ‘GaEl Ag-Patches’ were identified by reverse epitomics approach as introduced in our earlier studies^41–44^. All nine proteins of the NiV proteome (https://www.uniprot.org/proteomes/UP000120177) were used in this study viz. C protein (gi-1859635642); glycoprotein (gi-253559848); fusion protein (gi-13559813); nucleoprotein (gi-1679387250); matrix protein (gi-13559811); phosphoprotein (gi-1802790259); W protein (gi-374256971); V protein (gi-1802790260) and RNA polymerase (gi-15487370). The full-length protein sequences were retrieved from National Center for Biotechnology Information (NCBI). Up to 96 full length protein sequences belonging to different strains and origins of NiV were retrieved. The MPVs designed with antigenic patches as immunogenic composition, also carry potential discontinuous B cell epitopes as well as IFN-γ inducing epitopes in their tertiary structure model. Hence, the designed MPVs carry the potential to elicit cell-mediated as well as humoral immune response. Furthermore, both MPVs were designed with the adjuvants human β Defensin 2 and human β Defensin 3 at N and C termini^45,46^. The β-defensins have considerable immunological adjuvant activity and generate potent humoral immune responses^47–50^. Tertiary structure models of the MPVs were generated, refined, and further analyzed for molecular interaction with the ectodomain of the human Toll-Like Receptor 3 (TLR3) by molecular docking. The TLR3 is an essential immuno-receptor and acts as sentinel to bind and process antigens causing activation of the IFN response against foreign antigen^51,52^. With these functions TLR3 plays an important role as immune-receptor during the NiV infection and hence was chosen to examine the candidate MPVs^53,54^. The ability of CTL or HTL MPVs forming a complex with the ectodomain of human TLR3 was further investigated by molecular dynamics simulation studies. The cDNA of the designed MPVs was also analyzed for its high expression potential in the Human (mammalian) host cell line. Overall, from the present study, we may put forward the design of candidate Multi-Patch Vaccines, which qualify all the significant criterions of a potential vaccine against NiV infections. Note, that the reported ‘GaEl antigenic patches’ also provide potential candidate for diagnostics of NiV infection. The corresponding workflow for the vaccine design and utility of ‘GaEl Ag-Patches’ in diagnostics is shown in Supplementary Figure S1.

### Screening of Potential Epitopes

#### T cell Epitope Prediction

##### Screening for Cytotoxic T lymphocyte (CTL) Epitope

Identification of Cytotoxic T lymphocyte epitopes was performed by IEDB (Immune Epitope Database) tool “Proteasomal cleavage/TAP transport/MHC class I combined predictor” (http://tools.iedb.org/processing/) and “MHC-I Binding Predictions” (http://tools.iedb.org/mhci/)^55,56^. The tool provides a “Total Score” which is a combined score including results from the proteasome, MHC, TAP (N-terminal interaction), and processing analysis. Combination of six different methods viz. Consensus, NN-align, SMM-align, Combinatorial library, Sturniolo and NetMHCIIpan is applied^57^. Further Immunogenicity of all the shortlisted CTL epitopes was obtained by using “MHC I Immunogenicity” tool of IEDB (http://tools.iedb.org/immunogenicity/)^57^. The tool utilizes the physiochemical properties of constituting amino acid of the peptide sequence.

##### Screening for Helper T lymphocyte (HTL) Epitopes

The IEDB tool “MHC-II Binding Predictions” (http://tools.iedb.org/mhcii/) was used to identify Helper T lymphocyte (HTL) epitopes from NiV proteins^58–61^. Three different methods viz. combinatorial library, SMM_align and Sturniolo were used, further a comparison of score of the peptide against the scores of other known five million 15-mer peptides of SWISSPROT database was performed to screen HTL epitopes^58–61^. The tool generates a percentile rank for each potential peptide. The lower value of percentile rank indicates higher immunogenic potential of the HTL epitope.

##### CTL and HTL epitope Toxicity prediction

The ToxinPred tool was used to characterize the toxic potential of shortlisted CTL and HTL epitopes. The tool facilitates the identification of the highly toxic or non-toxic peptides^62^. The “SVM (Swiss-Prot) based” (support vector machine) method was used here. The method utilizes a dataset of 1805 sequences as positive (toxic) and 3593 sequences as negative (non-toxic) peptides from Swissprot as well as 1805 positive and 12541 negative peptide sequences from TrEMBLE.

### Overlapping CTL and HTL epitope clusters to ‘GaEl antigenic patches’

#### CTL & HTL overlapping epitope clusters based GaEl antigenic patch identification

All the shortlisted high scoring epitopes from NiV proteome were aligned using their amino acid sequences with the multiple sequence alignment (MSA) tool Clustal Omega^63^. The consensus amino acid sequence of overlapping epitopes were identified as ‘GaEl Ag-Patches’. This novel approach of search and identification of ‘GaEl antigenic patches’ from a source protein is named as ‘Reverse Epitomics’ as introduced and explained in our earlier studies^41–44^. This approach is applicable to proteins/antigens of any pathogen, not only to SARS-CoV-2 or NiV.

#### Population coverage by CTL & HTL epitopes covered by the ‘GaEl Ag-Patches’

The world population coverage by the overlapping CTL and HTL epitopes which were utilized to identify the GaEl Ag-Patches, was studied by the “population coverage” tool of IEDB^64^. T cells recognize complexes of MHC molecules with a given epitope. The epitope can elicit an immune response in individuals who expresses the binding MHC molecule^61^. The MHC molecules are expressed differentially in human population in different ethically distributed population. This MHC restricted epitope binding provides an opportunity to analyse worldwide population coverage by the given epitope.

#### Conservation analysis of antigenic patches

The amino acid sequence conservation of the shortlisted CTL and HTL ‘GaEl antigenic patches’ was analyzed with the “Epitope Conservancy Analysis” tool of IEDB. The epitope conservancy was performed against up to 96 protein sequences of NiV of different strains and origins collected from NCBI^65^.

### Multi-Patch Vaccines against NiV

The identified and shortlisted ‘GaEl Ag-Patches’ from the NiV proteins were further used as immunogenic composition to design two CTL and two HTL Multi-Patch Vaccine constructs as explained in the result section.

#### Physicochemical property analysis of the designed MPVs

Two CTL and two HTL MPVs were analyzed with the ProtParam tool^66^. The tool performs an empirical investigation of various physicochemical parameters viz. amino acid length, molecular weight, theoretical pI, expected half-life (in *E. coli*, Yeast & Mammalian cell), Aliphatic index, Grand average of hydropathicity (GRAVY) and instability index score. The aliphatic index and grand average of hydropathicity (GRAVY) indicate globular and hydrophilic nature of the protein. The instability index score indicates the stable nature of the fusion protein.

#### Interferon-gamma inducing epitope prediction from the MPVs

The two CTL and two HTL MPVs were screened for potential interferon-gamma (IFN-γ) inducing epitopes (from CD4+ T cells) utilizing the “IFN epitope” tool implementing the “Motif and SVM hybrid” method i.e. MERCI: Motif-EmeRging and with Classes-Identification, and the SVM: support vector machine hybrid. The prediction is based on the IEDB database with 3705 IFN-gamma inducing and 6728 non-inducing epitopes^67,68^.

#### MPVs allergenicity and antigenicity prediction

The CTL and HTL MPVs were analyzed for allergenicity and antigenicity with the AllergenFP and VaxiJen tools respectively^69,70^. The AllergenFP prediction is based on Hydrophobicity, their size, their helix-forming propensity, relative abundance of amino acids, β-strand forming propensity. The VaxiJen tool utilizes an alignment-free approach based on physicochemical properties of given query protein sequence.

#### Tertiary structure modelling and refinement of MPVs

The tertiary structures of the MPVs were calculated by homology modelling utilizing I-TASSER. The I-TASSER utilizes the sequence-to-structure-to-function paradigm for protein structure prediction^71^.

The refinement of all calculated structural models (two CTL and two HTL MPV) was performed with the GalaxyRefine tools^73,74^. The GalaxyRefine tool refines the input tertiary structure by repeated structure perturbation as well as by subsequent structural relaxation and molecular dynamics simulation^75,76^.

#### Validation of CTL and HTL MPVs refined models

Both the refined two CTL and two HTL MPV tertiary models were further validated by Ramachandran Plot Server (https://zlab.umassmed.edu/bu/rama/index.pl)^77^. The Ramachandran plots show sterically allowed and disallowed residues along with their dihedral psi (ψ) and phi (φ) angles.

#### Linear and Discontinuous B-cell epitope prediction from the MPVs

The IEDB tool, Ellipro (ElliPro: Antibody Epitope Prediction tool) was used to screen the linear and the discontinuous B cell epitopes from all the CTL and HTL MPVs tertiary models. The farthest residue to be considered was limited to 6 Å, the residues lying outside of an ellipsoid covering 90% of the inner core residues of the protein score highest Protrusion Index (PI) of 0.9; and so on. The discontinuous epitopes predicted based on the distance “R” in Å between the center of mass of two residues lying outside of the largest possible ellipsoid. The larger the value of R indicates greater distance between the residues (residue discontinuity) of the discontinuous epitopes^78,79^.

#### Molecular interaction analysis of MPVs and immune receptor complexes

The molecular interaction of the CTL and HTL MPVs with the ectodomain of Toll-Like receptor3 (TLR3) was analyzed by molecular docking followed by a molecular dynamics simulation study. The protein-protein molecular docking was performed by GRAMM-X Protein-Protein Docking v.1.2.0 tool^80^. The GRAMM-X tool utilizes the GRAMM Fast Fourier Transformation (FFT) methodology by employing smoothed potentials, refinement stage, and knowledge-based scoring.

#### Molecular Dynamics (MD) Simulations study of MPVs-TLR3(ECD) complexes

The molecular interactions of MPVs-TLR3(ECD) complexes were further evaluated using Molecular Dynamics (MD) simulations analysis. MD simulation studies were performed for 150 nanoseconds (ns) using the GROMACS tool^81,82^. The MD simulations studies were carried out in an explicit water environment in a cubic box as the unit cell simulation box at a stabilized temperature of 300 K and a pressure of 1 atm with periodic cell boundary condition. The solvated systems were neutralized with counter Cl^-^ ions. Only the ions necessary to neutralize the net charge on the protein are added by *gmx genion*. The OPLS all-atom force field was used on the systems during MD simulation^83^. The solvated structures were energy minimized by the steepest descent. Further, the complexes were equilibrated for a period of 100 ps. After equilibration MD simulation was run for 150 ns. The RMSD and RMSF values for Cα, Backbone and all the atoms of all the CTL and HTL MPVs in complex with TLR3 were analyzed.

#### Analysis of MPVs cDNA for expression in human host cell line

Codon-optimized complementary DNA (cDNA) of all the two CTL and two HTL MPVs were generated and analyzed for favored expression in Mammalian cell line (Human) by Java Codon Adaptation Tool. The cDNA were further analyzed by GenScript Rare Codon Analysis Tool for large scale expression potential. The tool analyses GC content, Codon Adaptation Index (CAI) and Tandem rare codon frequency^84,85^. The CAI indicates the possibility of expression in the chosen human cell line expression system. The tandem rare codon frequency indicates the presence of low-frequency codons.

## RESULTS & DISCUSSION

### Screening of potential epitopes

#### T cell Epitope Prediction

##### Screening of Cytotoxic T lymphocyte (CTL) Epitope

The Cytotoxic T lymphocyte (CTL) epitopes were screened and shortlisted according to their highest “Total Score”, low IC(50) (nM) value for epitope-HLA class I allele complexes and larger number of the HLA class I allele binders. Additional epitopes were screened from the protein sequences of NiV on the basis of their percentile rank. Smaller value of percentile rank indicate higher affinity of the peptide with its respective HLA class I allele binders. A total of 1811 (579+1232) CD8+ T cell epitopes:HLA class I allele pairs were screened. These included the top scoring epitopes and the highest percentile ranking epitopes from the NiV proteome. The immunogenicity of the shortlisted CTL epitopes were also determined. All the screened and shortlisted CTL epitopes are assessed as highly immunogenic in humans (Supplementary Table S1, S2).

##### Screening of Helper T lymphocyte (HTL) epitopes

The screening of helper T lymphocyte (HTL) epitopes was performed on the basis of “Percentile rank”. The smaller the value of percentile rank indicates higher affinity of the peptide with its respective HLA allele binders. We screened in total 773 potential CD4+ T cell epitopes:allele pairs showing highest percentile rank. The epitopes with largest number of HLA class II allele binders were also included (Supplementary Table S3).

##### CTL and HTL epitope Toxicity prediction

All the screened and shortlisted CTL and HTL epitopes were evaluated as Non-Toxic (Supplementary Table S1, S2 and S3).

Further, all the 1811 CTL and 773 HTL epitopes were evaluated to be highly conserved (IEDB: “Epitope Conservancy Analysis”) with 100% amino acid sequence of epitope being present in most of the NiV strains as shown in Supplementary Table S1, S2, and S3.

### Overlapping CTL and HTL epitope clusters to antigenic patches

#### CTL & HTL overlapping epitope clusters based antigenic patch identification

A total of 30 GaEl Ag-Patches from CTL and 27 GaEl Ag-Patches from HTL high scoring 776 overlapping epitopes (362 CTL and 414 HTL epitopes) were identified. To identify the GaEl Ag-Patches, a novel ‘Reverse epitomics’ approach involving “Overlapping-epitope-clusters-to-patches” method was utilized^41–44^ (Table 1 & 2). The herewith suggested GaEl Ag-Patches are expected to result in up to 776 overlapping epitopes upon intercellular proteolytic chop down by Antigen Presenting Cells (APC). Such large numbers of epitopes cannot be accommodated by Multi-Epitope Vaccines (MEV) candidates^38,86–90^. Furthermore, lagging behind to MPVs in respect of encoding a high number of epitopes, the MEVs with short peptide of epitopes might also result in wrong or missfoled peptides output upon intracellular proteolytic chop down by APC. Structural analysis showed that all the identified GaEl Ag-Patches locate at the surface of NiV proteins providing a more accessible target for immunogenic response. Therefore, the Multi-Patch Vaccine (MPVs) candidates are expected to be superior by utilizing ‘GaEl Ag-Patches’ as immunogenic composition (Figure 1, 2 & 3).

**Figure 1.**
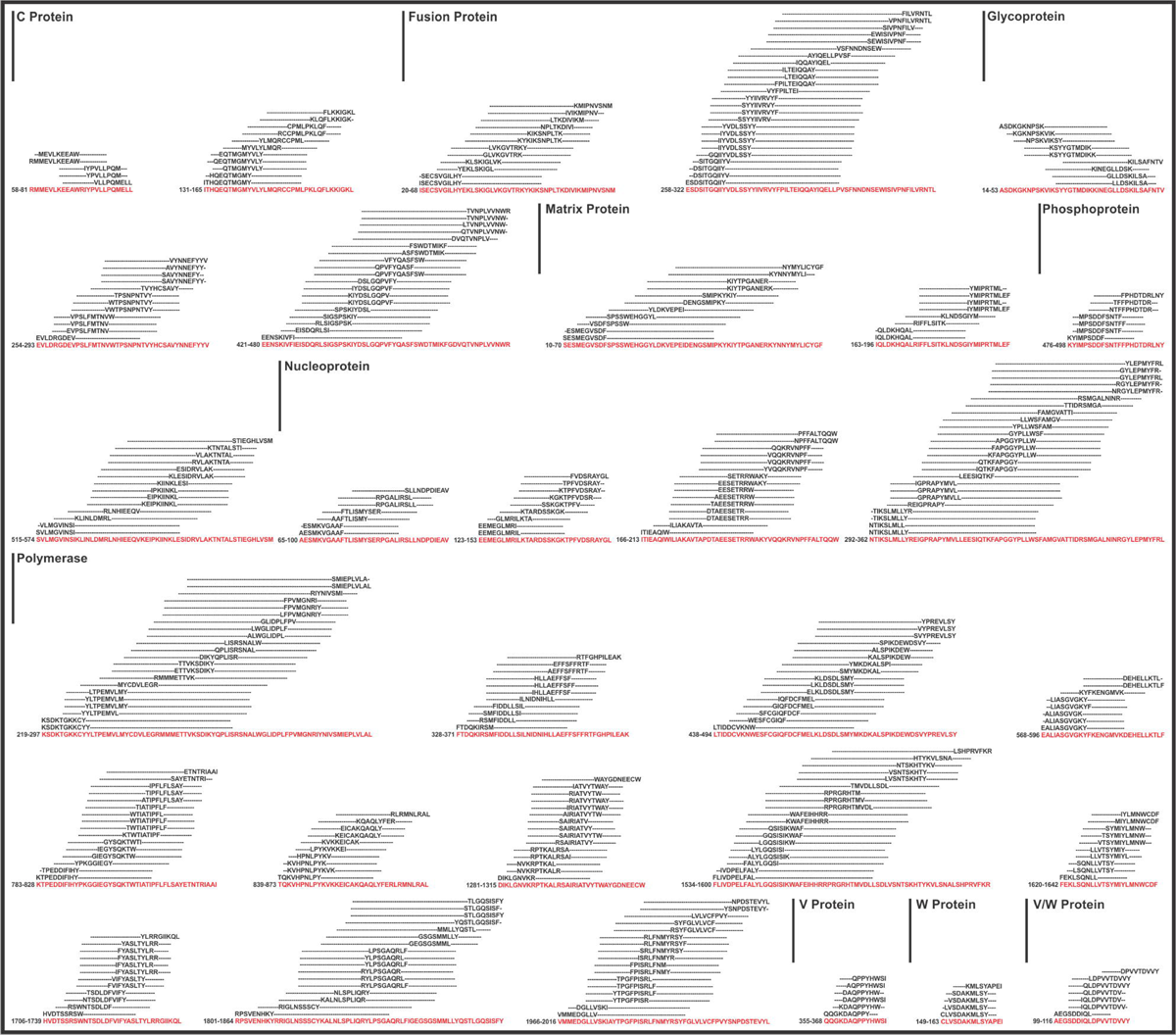
Graphical representation of the identified 30 CTL Ag-Patches from overlapping 362 CTL epitope screened from NiV proteins. Ag-Patches amino acid consensus sequences are highlighted in red.

**Figure 2.**
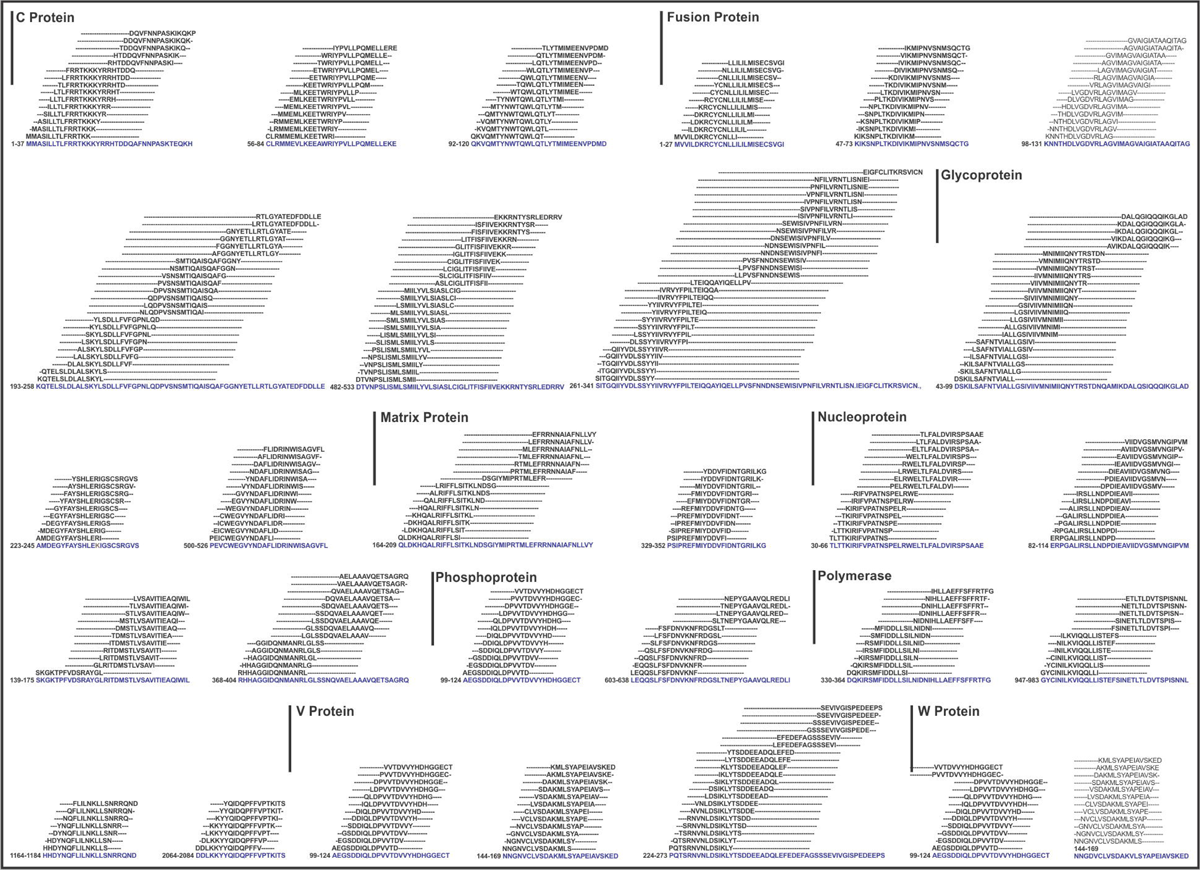
Graphical representation of the identified 27 HTL GaEl Ag-Patches from overlapping 414 HTL epitope clusters of NiV proteins. Ag-Patches are shown in blue amino acid sequences consensus.

**Figure 3.**
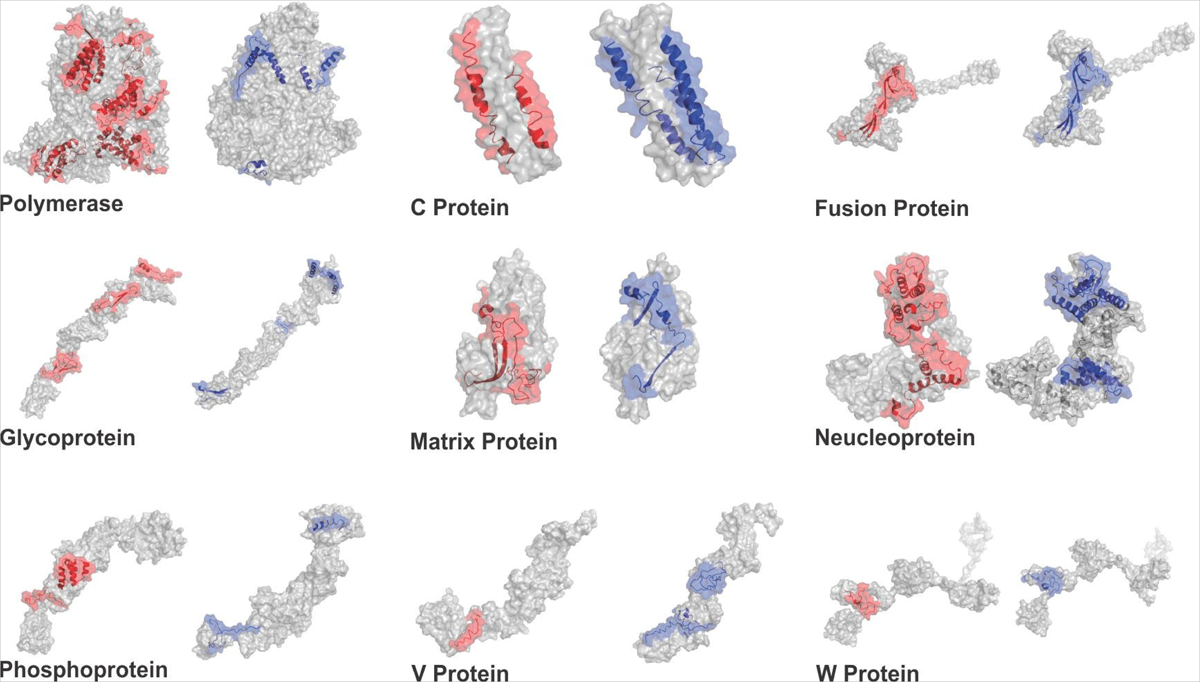
The identified 57 ‘GaEl Ag-Patches’ are shown in the tertiary structure models of the NiV proteins. The tertiary structure models of NiV proteins are generated by SwissModel and I-TASSER homology modeling^71,72^. The 30 CTL GaEl Ag-Patches are shown in red and the 27 HTL GaEl Ag-Patches are shown in blue color. Most of the GaEl Ag-Patches identified are observed to be on the exposed surface of the NiV proteins. Structural modeling details are given in Supplementary Table 4.

**Table 1:**
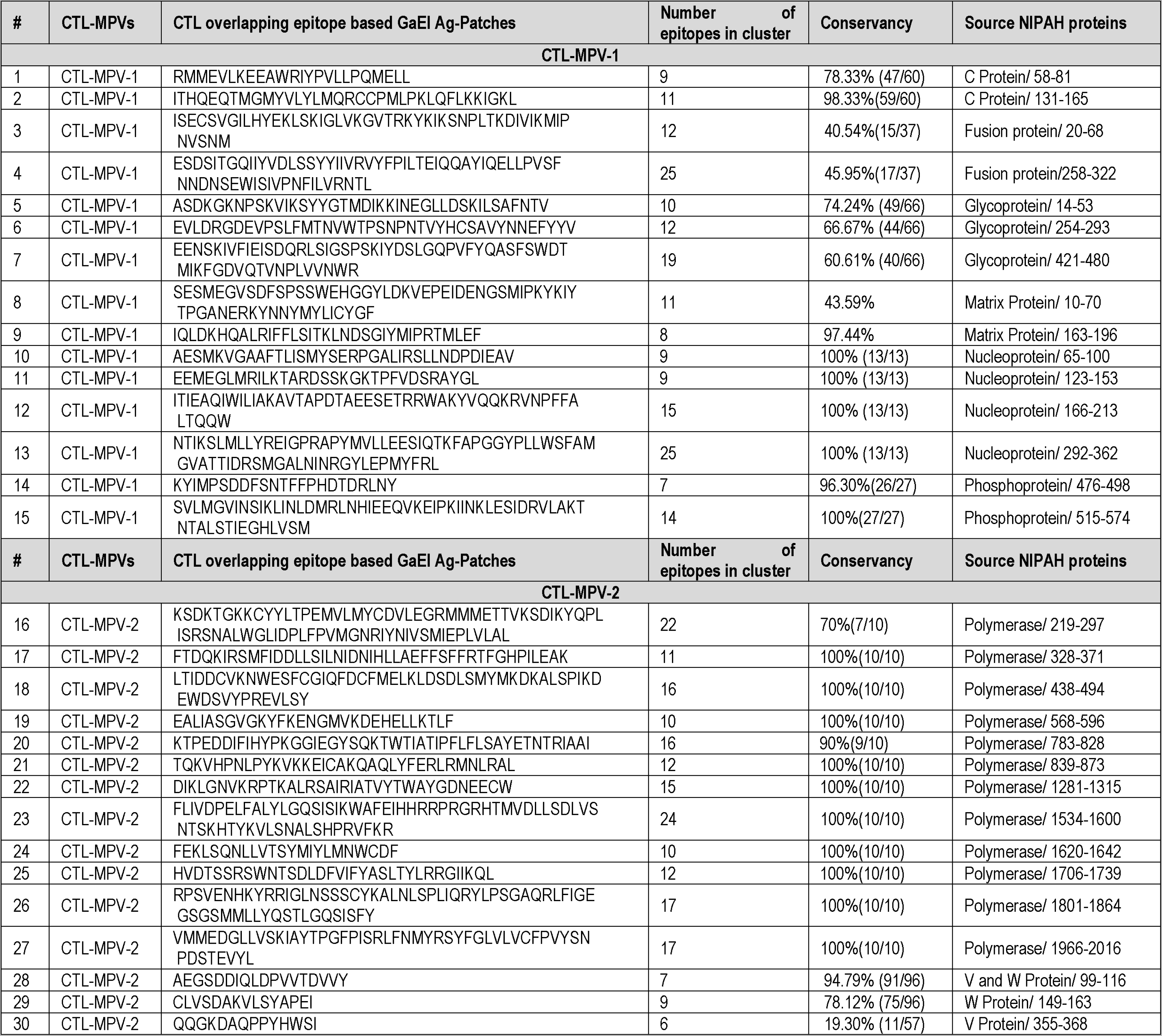
CTL ‘GaEl Ag-Patches’ from NiV proteome were identified by novel Reverse Epitomics approach involving the “overlapping-epitope-clusters-to-patches” method. The highly immunogenic 30 Ag-Patches were utilized to design two CTL (CTL-MPV-1 & CTL-MPV-2) Multi-Patch Vaccine candidates. The identified ‘GaEl Ag-Patches’ were highly conserved in nature. CTL ‗GaEl Ag-Patches’ from NIPAH Proteome: Supplementary txt file 1.

**Table 2:**
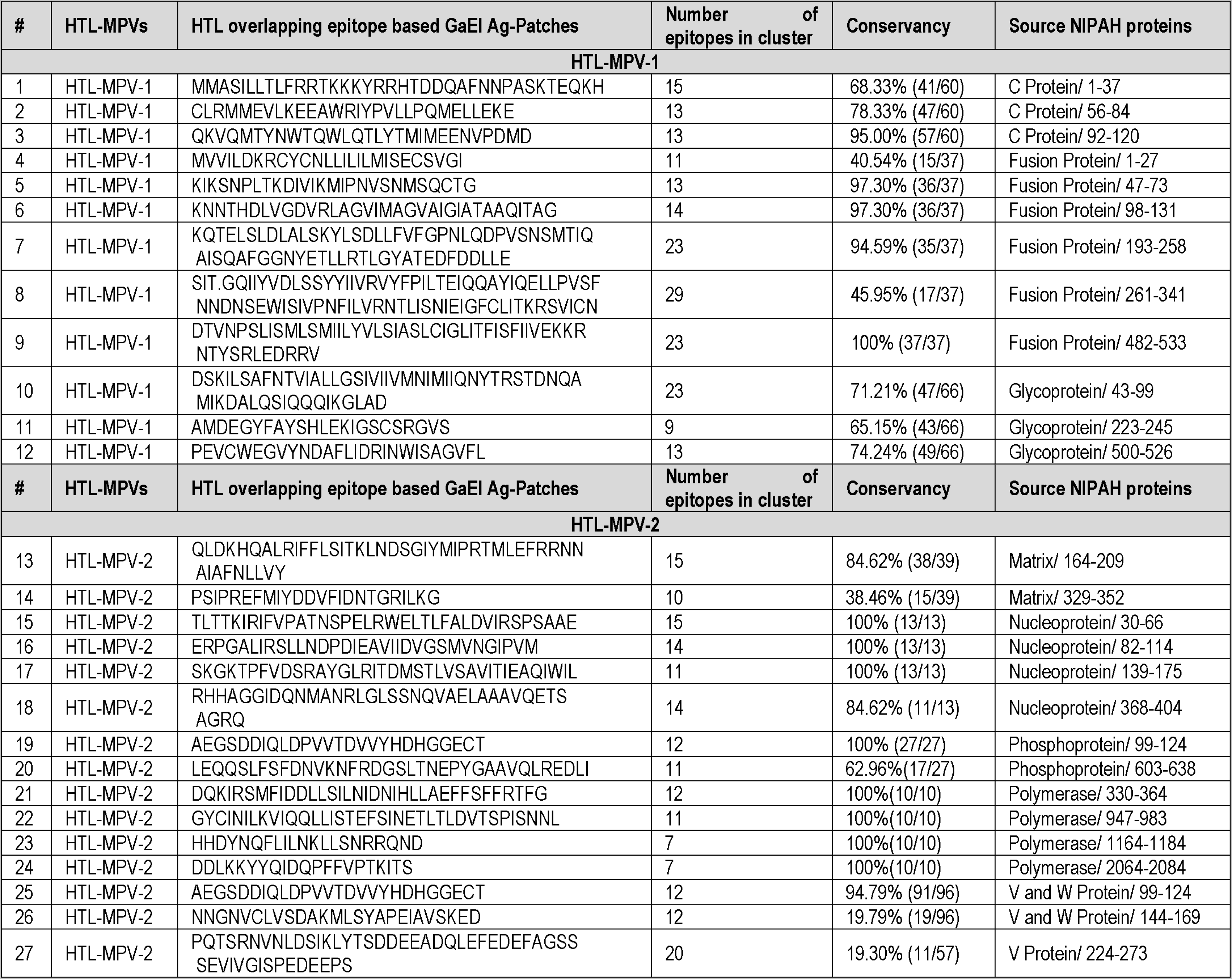
HTL GaEl Ag-Patches from NiV proteome were identified by novel Reverse Epitomics approach involving the “overlapping-epitope-clusters-to-patches” method. The highly immunogenic 27 Ag-Patches were utilized to design two HTL (HTL-MPV-1 & HTL-MPV-2) Multi-Patch Vaccine candidates. The identified GaEl Ag-Patches were highly conserved in nature. HTL ‘GaEl Ag-Patches’ from NIPAH Proteome: Supplementary txt file 2.

#### Population Coverage by antigenic patches

The population coverage by the GaEl antigenic patches was analyzed by the overlapping epitopes and their HLA allele binding pairs using the “Population Coverage” tool of IEDB. The 57 GaEL Ag-Patches constructed in this study from 362 CTL and 414 HTL overlapping epitopes, target a total of discret 27 HLA class I and 25 HLA class II alleles (Supplementary Table S5). Since HLA alleles are differentially expressed in ethnically distributed global population they provide us with the opportunity to analyse population coverage. The convincing 99.71% (average: 85.79 and standard deviation: 20.73) of the global human population is predicted to be covered by the CTL and HTL Multi-Patch Vaccine candidates proposed in this study. The countries most affected by NiV infections showed a significant coverage like India, 97.17%; Malasia, 91.87% etc. (Supplementary Table S6). The Epitope-HLA allele pairs are summarized in Supplementary txt file 3.

#### Conservation analysis of antigenic patches

The GaEl Ag-Patches were further analyzed for their amino acid sequence conservation. Up to 96 full length NiV protein sequences of different strains/origins, retrieved from NCBI protein database were analyzed with the “Epitope Conservancy Analysis” tool of IEDB. The CTL GaEl Ag-Patches were in the range of 43.59% to 100% (mostly above 90%) and HTL GaEl Ag-Patches between 40.54% and 100% conservation (mostly above 90%) (Table 1 & 2). Some of the assigned patches have also lower conservancy of around 19%, nevertheless the variation in their amino acid sequences is limited to only a few residues.

### Multi-Patch Vaccines

#### Design of Multi-Patch Vaccines

The GaEl Ag-Patches identified from the proteome of NiV were utilized as immunogenic composition for the design of two CTL and two HTL Multi-Patch vaccines resp. (Figure 4, Supplementary Table S7). Short amino acid linkers EAAAK and GGGGS provide a covalent and flexible connection between the constituting peptide, respectively. The EAAAK linker facilitates domain formation and the GGGGS linker provides conformational flexibility hence together favors stable protein folding. The EAAAK linker was used to fuse the human β defensin 2 and 3 (hBD-2 and hBD-3) at N and C terminal ends of the MPVs, respectively. The constructs include hBD-2 with sequence: GIGDPVTCLKSGAICHPVFCPRRYKQIGTCGLPGTKCCKKP and 3D structure deposited in the Protein Data Base (pdb code 1FD3) and hBD-3 with sequence: GIINTLQKYYCRVRGGRCAVLSCLPKEEQIGKCSTRGRKCCRRKK and 3D structure (pdb code 1KJ6). hBD 2 and 3 are utilized here as adjuvants to enhance the immunogenic response^45–47,49–50,91–93^. The Multi-Patch Vaccine design and the method thereof are included in the field patents 202011037939, 202011037939, PCT/IN2021/050841 and previous publications^41–44^.

**Figure 4.**
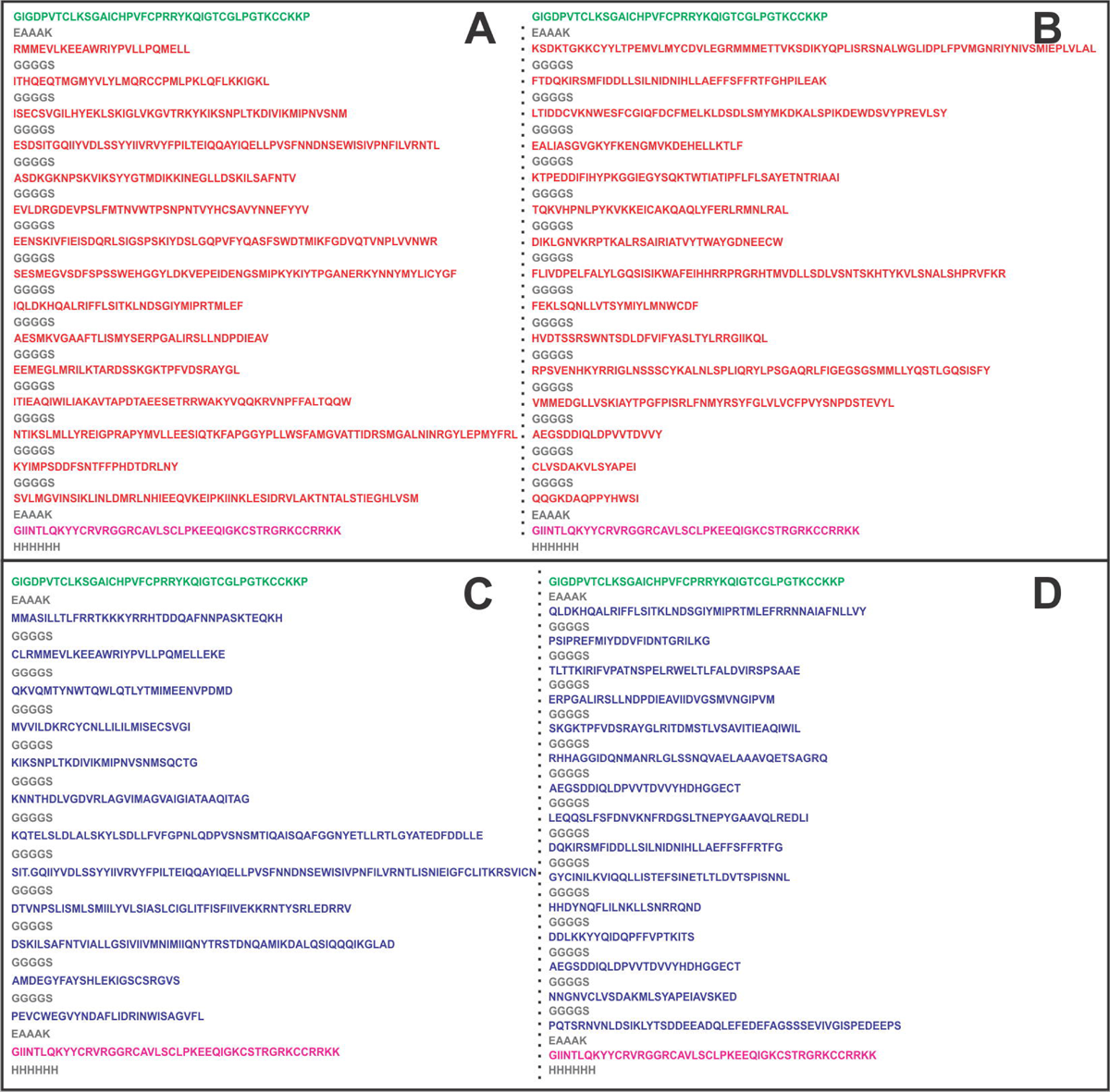
The CTL and HTL GaEl Ag-Patches are utilized as immunogenic composition to design Multi-Patch Vaccine. Short peptide linkers GGGGS and EAAK are used to fuse the GaEl Ag-Patches. The CTL MPV constructs includes (A) CTL-MPV-1 and (B) CTL-MPV-2. The HTL MPV construct includes (C) HTL-MPV-1 and (D) HTL-MPV-2 (Supplementary Table S7).

#### Physicochemical property analysis of the MPVs

ProtParam analysis was performed to analyze their physiochemical properties of all the suggested CTL and HTL MPVs. The retrieved empirical physiochemical properties of CTL and HTL MPVs are summarized in Table 3. The molecular weight of all MPVs range from 70.04 to 93.27 kDa. The expected half-life of the MPVs is predicted to be around ∼30 hours in mammalian cells suggesting stable expression products. The aliphatic index (81.80 to 96.13) and grand average of hydropathicity (GRAVY) (0.044 to -0.251) of the MPVs indicate their globular and hydrophilic nature. The instability index score of the MPVs (42.75 to 52.49) indicate stable nature of the MPV protein molecules. Taken together, the physiochemical parameters of MPV candidates indicate a favorable expression of the designed MPVs in human cells (Table 3).

**Table 3.**
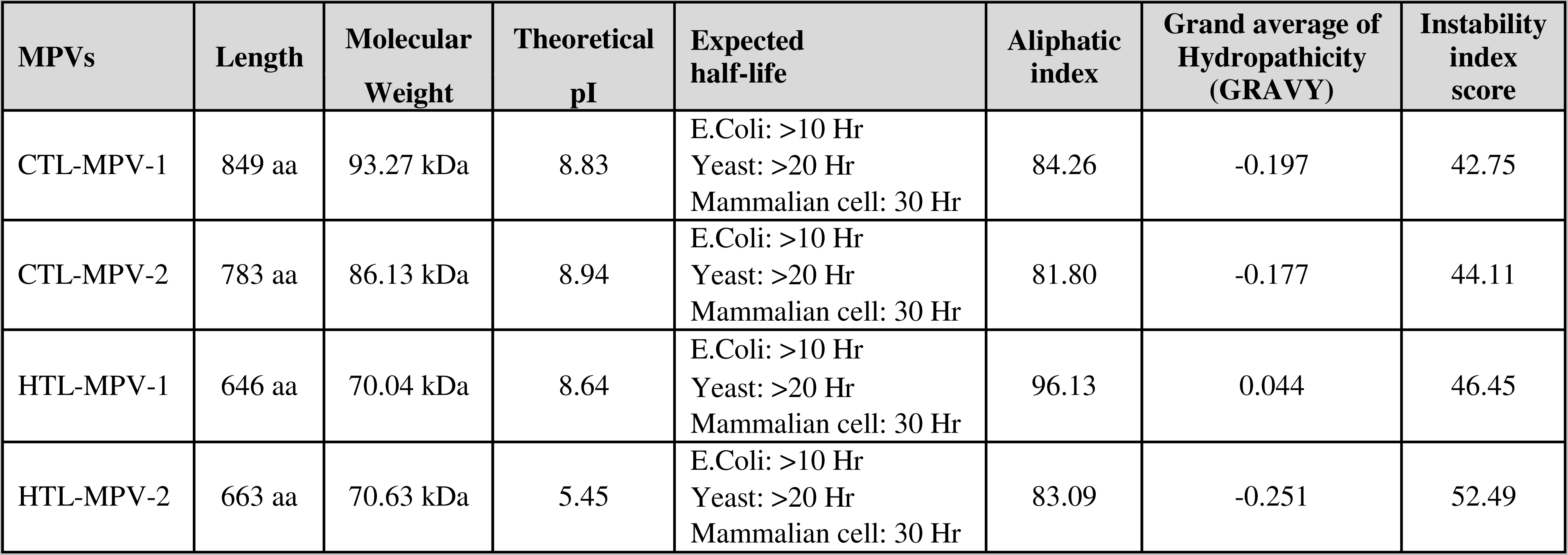
Empirical physicochemical properties of Multi-Patch Vaccine candidates.

#### Interferon-gamma inducing epitope prediction from the MPVs

Further, the MPV candidate molecules were also analyzed for the presence of potential Interferon-gamma (IFN-γ) inducing epitopes. The amino acid sequence of all the MPV candidates were screened for IFN-γ inducing 15 mer peptide epitopes by utilizing the IFNepitope tool. A total of 75 IFN-γ inducing epitopes was screened from two CTL MPVs and two HTL MPVs (Supplementary Table S8 & S9).

#### Allergenicity and antigenicity of MPV candidates

The characterization of the immunogenic properties of the suggested MPV is a crucial step in selecting appropriate vaccine candidates. We investigated both the *allergenicity and* the *antigenicity* using the bionformatic tools AllergenFP and Vaxijen resp. Note, all the MPV candidates were found to be good potential ANTIGENS and NON-ALLERGEN (Supplementary Table 10).

#### Tertiary structure modeling and refinement of MPVs

Tertiary structure homology models of MPV candidates were generated by using I-TASSER modeling tool (Figure 5). The parameters (C-score, TM-Score and RMSD) of homology modeling have shown acceptable values for all the MPV models (Supplementary Table S11). The C-score is a confidence score indicating the significance of template alignments and convergence parameters of the structure assembly simulations. C-score typically ranges from -5 to 2, with higher value indicating a model with high confidence and vice-versa. The C-score of all the four MPV models are acceptable indicating high confidence on the generated models. The TM-score indicates the structural alignment between the query structure and template structure. The RMSD (Root Mean Square Deviation) is the deviation between the residues that are structurally aligned (by TM-align) to the template structure.

**Figure 5.**
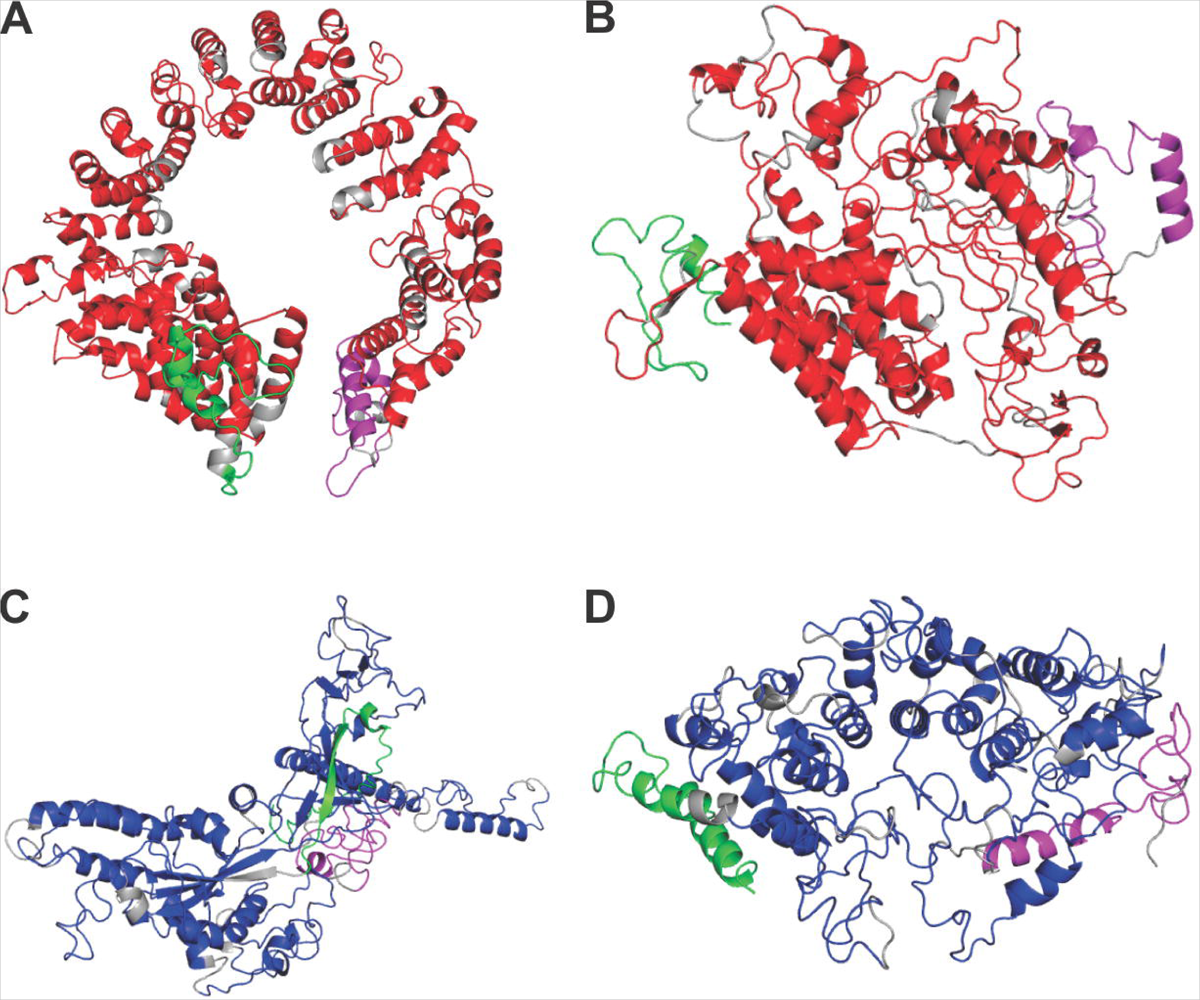
Tertiary structure modelling of Multi-Patch Vaccine candidates. Tertiary structural models of CTL and HTL MPVs are shown in cartoon (A) CTL-MPV-1, (B) CTL-MPV-2, (C) HTL-MPV-1 and (D) HTL-MPV-2. The Ag-Patches are shown in Red (CTL Ag-Patches), and blue (HTL Ag-Patches). Linkers are shown in gray, adjuvants are in green and magenta.

All the generated MPV tertiary models were further refined with the GalaxyRefine tools to improve the sterical conformation parameters of the calculated structures. We used for further analysis only the top scoring models. All selected MPV refinement output models have good Rama favoured, GDT-HA, RMSD value and MolProbity scores (Supplementary Table S12). Rama favored criteria indicates the percentage of residues which are in the favoured region of the Ramachandran plot, the GDT-HA (global distance test-High Accuracy) indicates the accuracy of the structure backbone, the RMSD (Root Mean Square Deviation) value indicated deviation from the initial model. The MolProbity score indicates log-weighted combination score of the clash score, percentage of Ramachandran not favored residues and percentage of bad side-chain rotamers.

Overall, after structural refinement, the MPV candidate had sterical parameters in acceptable range and hence all these models were carried forward for further analysis.

#### Validation of CTL and HTL MPVs refined models

We next analyzed the sterical acceptable conformation of all MPV models using Ramachandran plots. The RAMPAGE analysis tool used those plots indicated that the MPV models (CTL-MPV-1 &CTL-MPV-2) have 95.48% & 93.87% residues in favored region. Likewise, the HTL MPV models (HTL-MPV-1 & HTL-MPV-2) were found to have 90.90% & 87.3% residues in favored region (Supplementary Figure S2). Hence, all the MPV models show acceptable sterical conformations.

#### Linear and Discontinuous B-cell epitope prediction from the MPVs

The tertiary models of MPV candidates were tested for the presence of potential linear and discontinuous B Cell epitopes with the ElliPro tool of IEDB. The screening revealed that the CTL MPV carries a total of 39 linear epitopes [CTL-MPV-1 (21 epitopes) & CTL-MPV-2 (18 epitopes)]. Likewise, the HTL MPVs carries a total of 26 linear epitopes [HTL-MPV-1 (10 epitopes) & HTL-MPV-2 (16 epitopes)] (Supplementary Table S13). Further the CTL MPVs also carry 14 discontinuous B-cell epitopes [CTL-MPV-1 (3 epitopes) & CTL-MPV-2 (11 epitopes)]. Likewise the HTL MPVs carry 15 potential discontinuous epitopes [HTL-MPV-1 (7 epitopes) & HTL-MPV-2 (8 epitopes)] (Supplementary Table S14). The ElliPro associates its prediction with a PI (Protrusion Index) value. The high PI value of the linear and discontinuous epitopes suggest the MPVs to have high potential to cause humoral immune response.

#### Molecular interaction analysis of MPVs with immune receptor

The innate immune system Toll-Like Receptor 3 (TLR3) acts as sentinel against foreign antigens. For those reasons the molecular interaction of MPVs with the ectodomain (ECD) of the human TLR3 is an important aspect to study for a potential vaccine candidate. Therefore, the MPV candidates were docked with the ectodomain of human TLR3. Molecular docking study for the MPV models using a TLR3-ECD crystal structure (PDB ID: 2A0Z) was performed with the GRAMM-X Protein-Protein Docking tool of the Vakser Lab, Center for Bioinformatics at KU (Figure 6).

**Figure 6.**
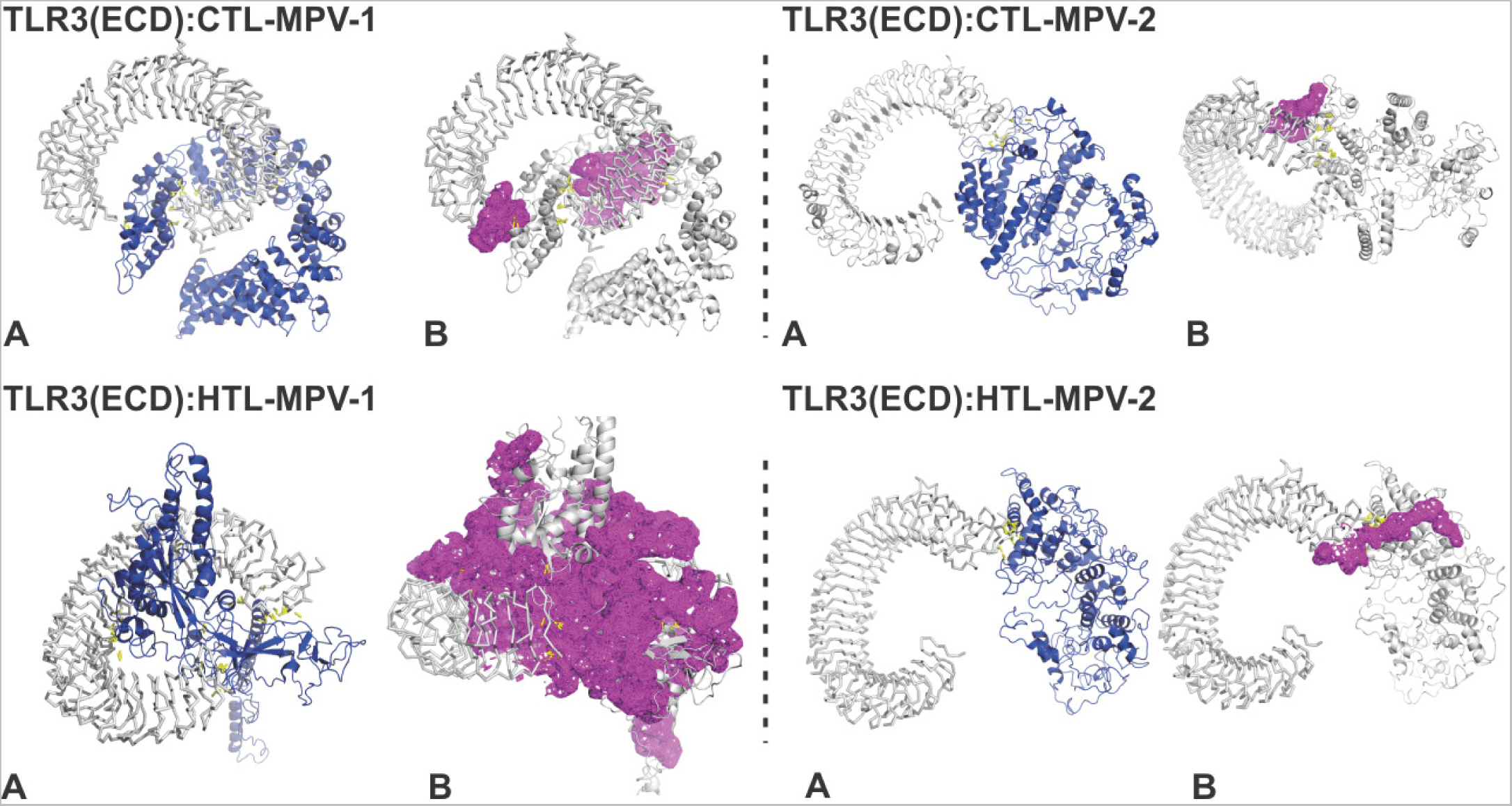
Molecular Docking Study of MPVs with TLR3-ECD. Complex formation by the MPVs (CTL-MPV-1, CTL-MPV-2, HTL-MPV-1, HTL-MPV-2) and TLR3-ECD are shown in different panels. **A:** The B-factor analysis for the MPVs in complex with TLR3-ECD is shown in VIBGYOR color, with violet showing low B-factor and red showing high B-factor. Here most of the MPV regions are in blue indicating low B-factor and hence suggesting a stable complex formation. The hydrogen bonds and the polar contacts are shown by yellow dots. **B:** Binding site clefts, generated by PDB*sum,* between TLR3-ECD:MPVs are shown by magenta surface. Molecular interaction by different bond formation is shown in Supplementary Figure S3.

The B-factor analysis of the MPV & TLR3-ECD complexes indicated the displacement of the atomic positions from an average (mean) value of the structure (Figure 6). The B-factor analysis of the MPV and TLR3-ECD complexes show most of the antigenic regions bound to TLR3-ECD is stable. The B-Factor analysis was performed using the VIBGYOR color presentation with violet representing low B-factor and red representing high B-factor. Most of the regions of MPVs & TLR3-ECD complex is in blue suggesting stable complex formation.

The docking studies of MPVs with TLR3-ECD indicated multiple binding sites as represented by the different complex structure in Figure 6 & Supplementary Figure S3. These complexes forms multiple hydrogen bonds in interaction interface (Figure 6). The residues forming hydrogen bonds, salt bridges, disulphide bonds and non-bonded contacts are shown in Supplementary Figure S3.

#### Molecular Dynamics (MD) Simulations study of MPV and TLR3 ectodomain complexes

The MPV/TLR3-ECD complexes were subjected to molecular dynamics (MD) simulation analysis with the program GROMACS version 2022.1 investigating their stability. The complexes have a very convincing Root Mean Square Deviation (RMSD) values for Backbone which gradually stabilizes towards the end of 150 nanoseconds of MD simulation [(CTL-MPV-1:TLR3-ECD, RMSD: ∼1Å) (CTL-MPV-2:TLR3-ECD, RMSD: ∼1Å) (HTL-MPV-1:TLR3-ECD, RMSD: ∼0.4 to ∼1.2 Å) (HTL-MPV-2:TLR3-ECD, RMSD: ∼0.4 to ∼1.4 Å)] (Supplementary Figure S4). In summary, all the four simulated complexes were found forming very stably interactions with TLR3-ECD at constant temperature (∼300 K) and pressure (∼1 atm). Further, the variation in radius of gyration (Rg) for all the MPV and TLR3-ECD complexes remain in acceptable range throughout the MD simulation, effectively: (CTL-MPV-1:TLR3-ECD, Rg: ∼4.5 to ∼7.5 Å) (CTL-MPV-2:TLR3-ECD, Rg: ∼4.5 to ∼7.5 Å) (HTL-MPV-1:TLR3-ECD, Rg: ∼3.55 to ∼3.85 Å) (HTL-MPV-2:TLR3-ECD, Rg: ∼4.0 to ∼4.8 Å)] (Supplementary Figure S4). The stable Rg indicate presence of structurally compact complexes throughout the MD simulation. Furthermore, the amino acid residues of the complexes show acceptable Root Mean Square Fluctuation (RMSF) from their initial confirmation [(CTL-MPV-1:TLR3-ECD, RMSF: ∼2 to ∼14 Å) (CTL-MPV-2:TLR3-ECD, RMSF: ∼1 to ∼6 Å) (HTL-MPV-1:TLR3-ECD, RMSF: ∼0.25 to ∼2.25 Å) (HTL-MPV-2:TLR3-ECD, RMSF: ∼0.2 to ∼1.2 Å)] (Supplementary Figure S4). Overall, the molecular docking and molecular dynamics simulation study of all the MPV & TLR3-ECD complexes indicate stable complex formed.

#### Analysis of complementary DNA of MPV candidates

The MPV construct were analyzed for expression feasibility in human cell line. Codon-optimized *complementary* DNA (cDNA) of MPVs were generated for expression in human cell line by the Java Codon Adaptation Tool. The optimized cDNA was further analyzed for its expression feasibility by the GenScript Rare Codon Analysis Tool. The GC content, the CAI score and the tandem rare codon frequency optimized MPV cDNA are observed to be in acceptable rage (Supplementary Table S15). The CAI score indicates a high propensity of cDNA expression in human cell expression system and the tandem rare codon frequency indicates the presence of low-frequency codons in cDNA. Briefly, the tandem rare codons which may hinder the proper expression of the cDNA in the chosen expression system was observed to be 0% in all the MPVs. Hence, as per the GenScript Rare Codon analysis, the cDNA of all the MPVs are predicted to have high potential for large-scale expression in the human cell line.

## DISCUSSION

Majority of the vaccine design against NiV are focused on the single protein like G and F proteins, protein subunits or the epitopes from NiV proteins^38,86–90^. The recent strategies for the design and development of vaccines to combat NiV involve subunit vaccines or multi-epitope vaccines. The subunit vaccine involves the use of single protein or a multiple subunits of NiV proteins. The major limitation with those vaccine candidates is with their limited efficiency being composed of single/limited number of protein/subunits. The more recent Multi-Epitope Vaccine approach provides opportunity to target multiple proteins of NiV. However, the low probability of MEV epitopes to be successfully presented by APC after intercellular chop down process, limits the efficacy of the MEV candidates. Moreover, the Multi-epitope vaccines can accommodate only a limited number of epitopes which puts it to face the challenge of frequent mutations in the proteome of the NiV.

In the present study, we have utilized the novel ‘reverse epitomics’ approach to design Multi-Patch Vaccine^41–44,94^. The consensus amino acid sequence of overlapping epitopes is identified as GaEl Ag-Patch. This method is termed as “Overlapping-epitope-clusters-to-patches” method. We have identified potential GaEl Ag-Patches from the entire proteome of NiV. All the identified GaEl Ag-Patches are observed to arise from surface of the NiV proteins providing a potential target for immunogenic response. Further, we utilized the GaEl Ag-Patch as immunogenic composition of the Multi-Patch Vaccine against NiV. Hence these GaEl Ag-Patches cover large number of overlapping epitopes which they could produce upon proteolytic intracellular chop down process by APC. These shortlisted GaEl Ag-Patches originate from all the proteins encoded by NiV, thus targeting its entire proteome^94^. These advantages of the Multi-Patch strategy would enhance the vaccine efficiency and lead to the vaccines that are more targeted/specific and effective. Since the GaEl Ag-Patches cover large number of epitopes which in turn target large number of different number of HLA alleles, the MPV would also provide larger ethnically distributed human population coverage. Here, the designed MPVs cover 99.71% of the world population targeting 52 different HLA class I and II alleles, a coverage that is much higher in comparison to the vaccine composed of single/limited number of protein/subunits/epitopes. In addition to potential immunogenic composition for vaccine candidates, the reported GaEl Ag-Patches also provides potential immunogenic composition for early detection diagnostic kits against NiV infection. Further the MPVs show stable binding with TLR3-ECD which is one of the essential criteria for an antigen to be recognized and processed by the human immune system. The physicochemical properties and cDNA analysis of MPVs favor their over expression in human cell lines.

## CONCLUSION

In the present study, we have identified highly immunogenic novel GaEl antigenic patches (GaEl Ag-Patches) (30 CTL and 27 HTL) from the entire proteome of NiV. We have utilized the novel ‘reverse epitomics’ approach involving “overlapping-epitope-clusters-to-patches” method. These GaEl Ag-Patches are highly evolutionary conserved in nature. Further, for the first time we have identified the GaEl Ag-Patches and used them as immunogenic composition to design a Multi-Patch Vaccine (MPV) against NiV. The MPVs designed against NiV in our study have potential to give rise to a total of 776 discreet epitopes (362 CTL and 414 HTL epitopes) targeting a total of discreet 27 HLA class I and 25 HLA class II alleles and hence covering the convincing 99.71% of world human population. Such large number of epitopes is not possible to accommodate in multi-epitope vaccines. We conclude that the Multi-Patch vaccine designed by novel ‘reverse epitomics’ approach, utilizing GaEl Ag-Patches as immunogenic composition could be highly potent with greater effectiveness, high specificity and large human population coverage worldwide and protect human population against NiV infection in an effective manner. The reported GaEl Ag-Patches also provide potential candidates for early detection diagnostic kit for NiV infection.

## Supporting information

Supplementary txt file 1

Supplementary txt file 2

Supplementary txt file 3

## AUTHOR’S CONTRIBUTION

Idea conceived, Methodology designed and performed: SS; Scientific reporting and revising the manuscript: S.S. and M.K.

## ACKNOWLEDGEMENTS

We acknowledge the Indian Foundation for Fundamental Research Trust (IFFR Trust) and Helmholtz Center for Infection Research (HZI) for providing resources and funding. We also acknowledge kind help of Michele Lunelli for running the MD simulation.

## ADDITIONAL INFORMATION

Authors declare to have no competing interests.

## DICLARATION

Novel term “GaEl antigenic patch” is introduced. Three patents are used with due permissions. The study is part of patent proceedings. All rights reserved.

---

**Supplementary Figure S1.**
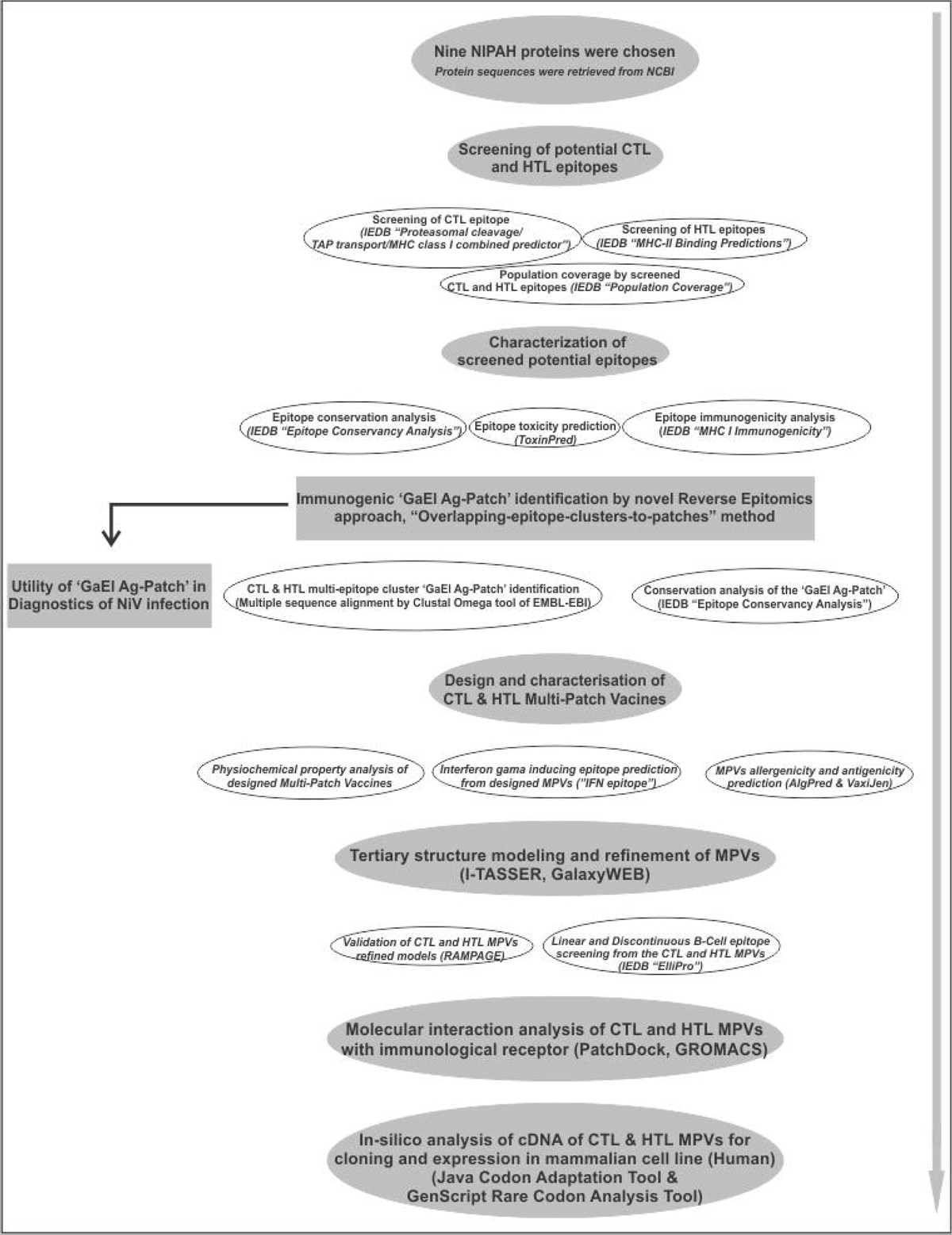
Schematic representation of workflow and methodology.

**Supplementary Figure S2.**
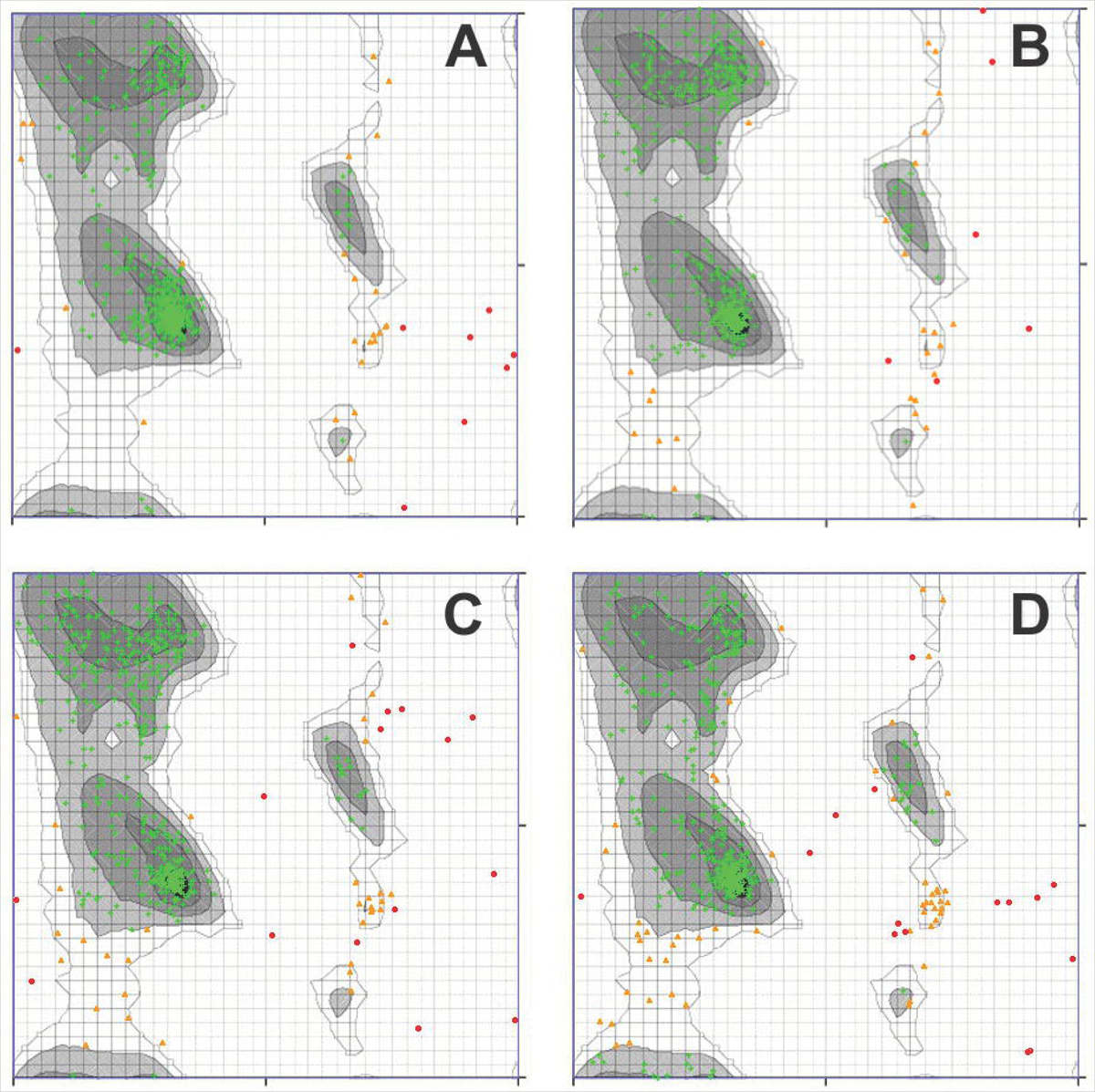
RAMPAGE analysis for all the MPVs **(A)** CTL-MPV-1, **(B)** CTL-MPV-2, **(C)** HTL-MPV-1, **(D)** HTL-MPV-2.

**Supplementary Figure S3.**
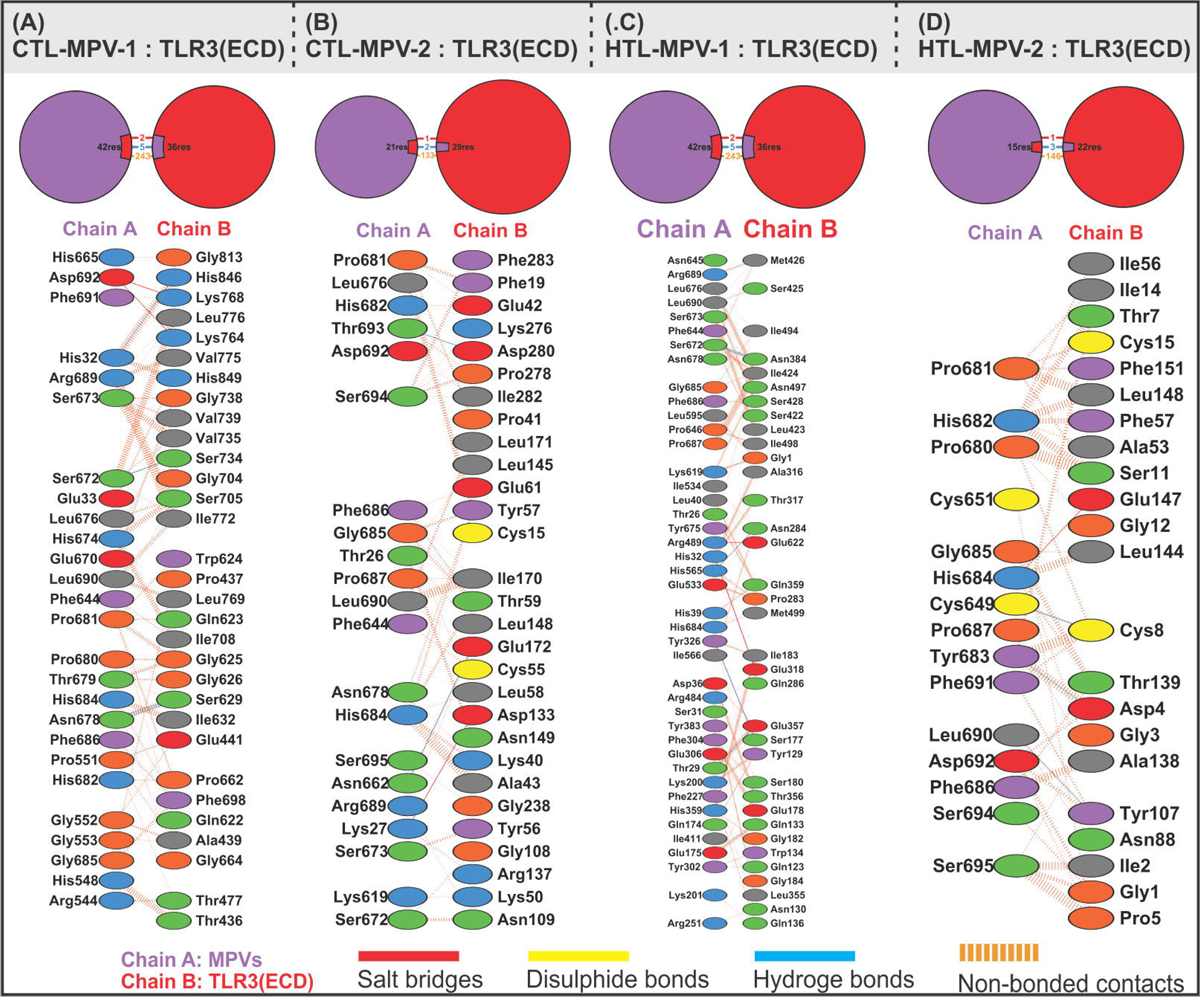
Molecular interaction between MPVs and TLR3 Ectodomain **(A)** CTL-MPV-1, **(B)** CTL-MPV-2, **(C)** HTL-MPV-1 and **(D)** HTL-MPV-2.

**Supplementary Figure S4.**
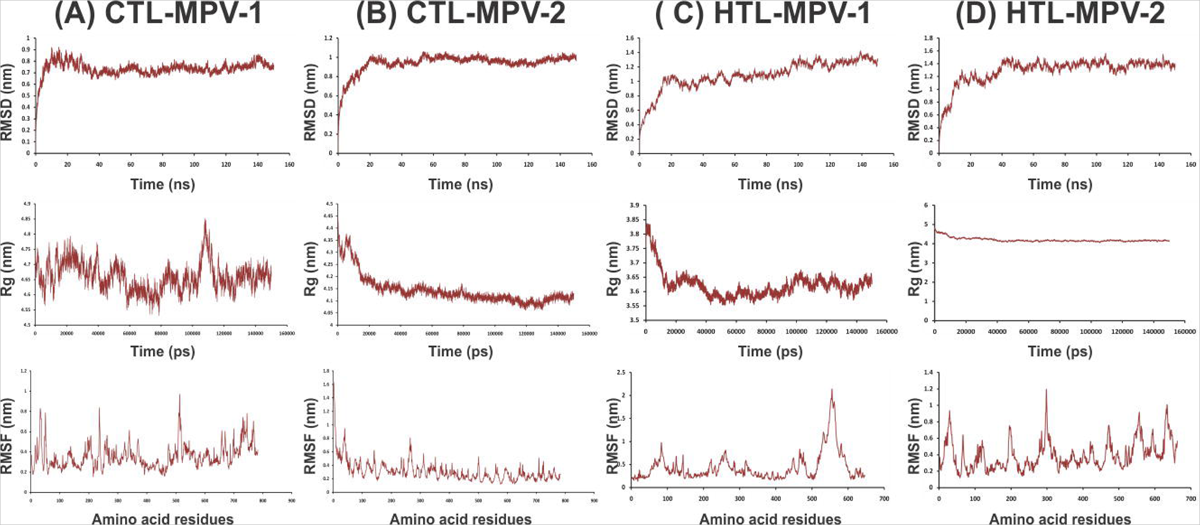
Molecular Dynamics simulation study of the MPVs and TLR3-ECD complexes. The Root Mean Square Deviation (RMSD) for Cα, Backbone, the Radius of gyration for all the MPVs and TLR3-ECD complexes and the Root Mean Square Fluctuation (RMSF) in the conformation of residues of the MPVs in complex with TLR3-ECD are shown.

## SUPPLEMENTARY TABLES

**Supplementary Table S1:** High Percentile Ranking CTL epitopes-HLA allele pairs screened from entire proteome of SARS-CoV-2 by the “MHC-I Processing Predictions” tool of IEDB. These epitopes were further utilized to identify the potentially immunogenic multiple epitope cluster based CTL Ag-Patches from the entire proteome of the SRAS-CoV-2. The screened epitopes are in consensus with the previous studies^38^.

**Supplementary Table S2:** High Scoring CTL epitopes-HLA allele pairs screened from entire proteome of SARS-CoV-2 by the “MHC-I Binding Predictions” tool of IEDB. These epitopes were further utilized to identify the potentially immunogenic multiple epitope cluster based CTL Ag-Patches from the entire proteome of the SRAS-CoV-2. The screened epitopes are in consensus with the previous studies^38^.

**Supplementary Table S3:** High Percentile Ranking HTL epitopes-HLA allele pairs screened from entire proteome of SARS-CoV-2 by the “MHC-II Binding Predictions” tool of IEDB. These epitopes were further utilized to identify the potentially immunogenic multiple epitope cluster based HTL Ag-Patches from the entire proteome of the SRAS-CoV-2. The screened epitopes are in consensus with the previous studies^38^.

**Supplementary Table S4.** Homology models of full length protein tertiary structure of NIPAH virus. Nipah protein sequences were retrieved from NCBI. The homology modeling tools SwissModel and I-Tasser were used.

**Supplementary Table S5:** HLA alleles covered by the overlapping CTL and HTL epitopes.

**Supplementary Table S6:** Population coverage by all the overlapping CTL and HTL epitopes forming epitope clusters.

**Supplementary Table S7:** Construct of CTL-MPV-1, CTL-MPV-2, CTL-MPV-3, HTL-MPV-1, and HTL-MPV-2. Physicochemical property analysis based on the amino acid sequences of all the designed three CTL and two HTL Multi-Patch Vaccines.

**Supplementary Table S8:** INF-γ inducing POSITIVE epitopes with a score of 1 or more than 1, screened from the CTL MPVs.

**Supplementary Table S9:** INF-γ inducing POSITIVE epitopes with a score of 1 or more than 1, screened from the HTL MPVs.

**Supplementary Table S10:** AllergenFP and Vaxijen analysis of MPVs. For the Vaxijen the default threshold is 0.4, and here all the MPVs have scored above 0.4, indicating potential antigenic nature.

**Supplementary Table S11:** Parameters for the tertiary structure homology modeling of all the CTL and HTL MPVs by the I-TASSER tool.

**Supplementary Table S12:** Refinement parameter values for CTL and HTL MPV models after refinement by GalaxyRefine tool. RMSD value in Å indicated deviation from initial model. GDT-HA (global distance test-High Accuracy): backbone structure accuracy measured by GDT-HA.

**Supplementary Table S13:** B cell linear epitopes screened from CTL & HTL MPVs.

**Supplementary Table S14:** B cell Discontinuous epitopes screened from CTL MPVs.

**Supplementary Table S15:** Analysis of codon-optimized cDNA of all the MPVs.

## REFERENCES

(1) Angeletti, S., Presti, A.L., Cella, E. and Ciccozzi, M., 2016. Molecular epidemiology and phylogeny of nipah virus infection: a mini review. Asian Pacific journal of tropical medicine, 9(7), pp.630–634.

(2) Aguilar, H.C., Henderson, B.A., Zamora, J.L. and Johnston, G.P., 2016. Paramyxovirus glycoproteins and the membrane fusion process. Current clinical microbiology reports, 3(3), pp.142–154.

(3) Ang, B.S., Lim, T.C. and Wang, L., 2018. Nipah Virus Infection. Journal of clinical microbiology, pp.JCM-01875.

(4) WHO Report, Surveillance and outbreak alert, Nipah virus; https://www.who.int/health-topics/nipah-virus-infection#tab=tab_1

(5) Plowright RK, Becker DJ, Crowley DE, Washburne AD, Huang T, Nameer PO, Gurley ES, Han BA. Prioritizing surveillance of Nipah virus in India. PLoS Negl Trop Dis. 2019 Jun 27;13(6):e0007393. doi: 10.1371/journal.pntd.0007393. PMID: 31246966; PMCID: PMC6597033.

(6) Thomas, B., Chandran, P., Lilabi, M. P., George, B., Sivakumar, C. P., Jayadev, V. K., Bindu, V., Rajasi, R. S., Vijayan, B., Mohandas, A., & Hafeez, N. (2019). Nipah Virus Infection in Kozhikode, Kerala, South India, in 2018: Epidemiology of an Outbreak of an Emerging Disease. Indian journal of community medicine : official publication of Indian Association of Preventive & Social Medicine, 44(4), 383–387. https://doi.org/10.4103/ijcm.IJCM_198_19

(7) Mathieu, C., Guillaume, V., Volchkova, V.A., Pohl, C., Jacquot, F., Looi, R.Y., Wong, K.T., Legras-Lachuer, C., Volchkov, V.E., Lachuer, J. and Horvat, B., 2012. Nonstructural Nipah virus C protein regulates both the early host proinflammatory response and viral virulence. Journal of virology, pp.JVI-01203.

(8) Liu, Q., Chen, L., Aguilar, H.C. and Chou, K.C., 2018. A stochastic assembly model for Nipah virus revealed by super-resolution microscopy. Nature communications, 9(1), p.3050.

(9) Johnston, G.P., Contreras, E.M., Dabundo, J., Henderson, B.A., Matz, K.M., Ortega, V., Ramirez, A., Park, A. and Aguilar, H.C., 2017. Cytoplasmic motifs in the nipah virus fusion protein modulate virus particle assembly and egress. Journal of virology, pp.JVI–02150.

(10) Satterfield, B.A., Cross, R.W., Fenton, K.A., Borisevich, V., Agans, K.N., Deer, D.J., Graber, J., Basler, C.F., Geisbert, T.W. and Mire, C.E., 2016. The Nipah virus C and W proteins contribute to respiratory disease in ferrets. Journal of virology, pp.JVI–00215.

(11) Lamp, B., Dietzel, E., Kolesnikova, L., Sauerhering, L., Erbar, S., Weingartl, H. and Maisner, A., 2013. Nipah virus entry and egress from polarized epithelial cells. Journal of virology, pp.JVI–02696.

(12) Weise, C., Erbar, S., Lamp, B., Vogt, C., Diederich, S. and Maisner, A., 2010. Tyrosine residues in the cytoplasmic domains affect sorting and fusion activity of the Nipah virus glycoproteins in polarized epithelial cells. Journal of virology, 84(15), pp.7634–7641.

(13) Ciancanelli, M.J. and Basler, C.F., 2006. Mutation of YMYL in the Nipah virus matrix protein abrogates budding and alters subcellular localization. Journal of virology, 80(24), pp.12070–12078.

(14) Patch, J.R., Crameri, G., Wang, L.F., Eaton, B.T. and Broder, C.C., 2007. Quantitative analysis of Nipah virus proteins released as virus-like particles reveals central role for the matrix protein. Virology journal, 4(1), p.1.

(15) Patch, J.R., Han, Z., McCarthy, S.E., Yan, L., Wang, L.F., Harty, R.N. and Broder, C.C., 2008. The YPLGVG sequence of the Nipah virus matrix protein is required for budding. Virology journal, 5(1), p.137.

(16) Jordan, P.C., Liu, C., Raynaud, P., Lo, M.K., Spiropoulou, C.F., Symons, J.A., Beigelman, L. and Deval, J., 2018. Initiation, extension, and termination of RNA synthesis by a paramyxovirus polymerase. PLoS pathogens, 14(2), p.e1006889.

(17) Ranadheera, C., Proulx, R., Chaiyakul, M., Jones, S., Grolla, A., Leung, A., Rutherford, J., Kobasa, D., Carpenter, M. and Czub, M., 2018. The interaction between the Nipah virus nucleocapsid protein and phosphoprotein regulates virus replication. Scientific reports, 8(1), p.15994.

(18) Baronti, L., Erales, J., Habchi, J., Felli, I.C., Pierattelli, R. and Longhi, S., 2015. Dynamics of the intrinsically disordered C-terminal domain of the Nipah virus nucleoprotein and interaction with the X domain of the phosphoprotein as unveiled by NMR spectroscopy. ChemBioChem, 16(2), pp.268–276.

(19) Uchida, S., Horie, R., Sato, H., Kai, C. and Yoneda, M., 2018. Possible role of the Nipah virus V protein in the regulation of the interferon beta induction by interacting with UBX domain-containing protein1. Scientific reports, 8(1), p.7682.

(20) Ludlow, L.E., Lo, M.K., Rodriguez, J.J., Rota, P.A. and Horvath, C.M., 2008. Henipavirus V protein association with Polo-like kinase reveals functional overlap with STAT1 binding and interferon evasion. Journal of virology, 82(13), pp.6259–6271.

(21) Park, M.S., Shaw, M.L., Munoz-Jordan, J., Cros, J.F., Nakaya, T., Bouvier, N., Palese, P., García-Sastre, A. and Basler, C.F., 2003. Newcastle disease virus (NDV)-based assay demonstrates interferon-antagonist activity for the NDV V protein and the Nipah virus V, W, and C proteins. Journal of virology, 77(2), pp.1501–1511.

(22) Sakib, M.S., Islam, M., Hasan, A.K.M. and Nabi, A.H.M., 2014. Prediction of epitope-based peptides for the utility of vaccine development from fusion and glycoprotein of nipah virus using in silico approach. Advances in bioinformatics, 2014.

(23) Guillaume, V., Contamin, H., Loth, P., Georges-Courbot, M.C., Lefeuvre, A., Marianneau, P., Chua, K.B., Lam, S.K., Buckland, R., Deubel, V. and Wild, T.F., 2004. Nipah virus: vaccination and passive protection studies in a hamster model. Journal of virology, 78(2), pp.834–840.

(24) Kong, D., Wen, Z., Su, H., Ge, J., Chen, W., Wang, X., Wu, C., Yang, C., Chen, H. and Bu, Z., 2012. Newcastle disease virus-vectored Nipah encephalitis vaccines induce B and T cell responses in mice and long-lasting neutralizing antibodies in pigs. Virology, 432(2), pp.327–335.

(25) Kamthania, M. and Sharma, D.K., 2015. Screening and structure-based modeling of T-cell epitopes of Nipah virus proteome: an immunoinformatic approach for designing peptide-based vaccine. 3 Biotech, 5(6), pp.877-882.

(26) Kamthania, M. and Sharma, D.K., 2016. Epitope-based peptides prediction from proteome of nipah virus. International Journal of Peptide Research and Therapeutics, 22(4), pp.465–470.

(27) Ali, M.T., Morshed, M.M. and Hassan, F., 2015. A computational approach for designing a universal epitope-based peptide vaccine against Nipah virus. Interdisciplinary Sciences: Computational Life Sciences, 7(2), pp.177–185.

(28) kumar Sharma, S., Srivastava, S., Kumar, A. and Srivastava, V., 2021. Anticipation of Antigenic Sites for the Goal of Vaccine Designing Against Nipah Virus: An Immunoinformatics Inquisitive Quest. International Journal of Peptide Research and Therapeutics, pp.1–13.

(29) Dey, S., Roy, P., Dutta, T., Nandy, A. and Basak, S.C., 2018. Rational Design of Peptide Vaccines for the Highly Lethal Nipah and Hendra Viruses. bioRxiv, p.425819.

(30) Krishnamoorthy, P.K., Subasree, S., Arthi, U., Mobashir, M., Gowda, C. and Revanasiddappa, P.D., 2020. T-cell Epitope-based Vaccine Design for Nipah Virus by Reverse Vaccinology Approach. Combinatorial chemistry & high throughput screening, 23(8), pp.788–796.

(31) Sakib, M.S., Islam, M., Hasan, A.K.M. and Nabi, A.H.M., 2014. Prediction of epitope-based peptides for the utility of vaccine development from fusion and glycoprotein of nipah virus using in silico approach. Advances in bioinformatics, 2014.

(32) Eshaghi, M., Tan, W.S. and Yusoff, K., 2005. Identification of epitopes in the nucleocapsid protein of Nipah virus using a linear phage-displayed random peptide library. Journal of medical virology, 75(1), pp.147–152.

(33) Mohammed, A.A., Shantier, S.W., Mustafa, M.I., Osman, H.K., Elmansi, H.E., Osman, I.A.A., Mohammed, R.A., Abdelrhman, F.A., Elnnewery, M.E., Yousif, E.M. and Mustafa, M.M., 2020. Epitope-based peptide vaccine against glycoprotein G of Nipah henipavirus using immunoinformatics approaches. Journal of immunology research, 2020.

(34) Gupta, A.K., Kumar, A., Rajput, A., Kaur, K., Dar, S.A., Thakur, A., Megha, K. and Kumar, M., 2020. NipahVR: a resource of multi-targeted putative therapeutics and epitopes for the Nipah virus. Database, 2020.

(35) Habib, P.T., 2021. Learning from COVID-19 Pandemic: In Silico Vaccine and Cloning Design Against Nipah Virus by Studying and Analyzing the Whole Nipah Virus Proteome.

(36) Singh, R.K., Dhama, K., Chakraborty, S., Tiwari, R., Natesan, S., Khandia, R., Munjal, A., Vora, K.S., Latheef, S.K., Karthik, K. and Singh Malik, Y., 2019. Nipah virus: epidemiology, pathology, immunobiology and advances in diagnosis, vaccine designing and control strategies–a comprehensive review. Veterinary Quarterly, 39(1), pp.26–55.

(37) Majee, P., Jain, N. and Kumar, A., 2021. Designing of a multi-epitope vaccine candidate against Nipah virus by in silico approach: a putative prophylactic solution for the deadly virus. Journal of Biomolecular Structure and Dynamics, 39(4), pp.1461–1480.

(38) Srivastava, S., Saxena, A.K. and Kolbe, M., 2021. Exploring the structural basis to develop efficient multi-epitope vaccines displaying interaction with HLA and TAP and TLR3 molecules to prevent NIPAH infection, a global threat to human health. bioRxiv.

(39) Tenzer S, Peters B, Bulik S, Schoor O, Lemmel C, Schatz MM, Kloetzel PM, Rammensee HG, Schild H, Holzhutter HG. 2005. Modeling the MHC class I pathway by combining predictions of proteasomal cleavage, TAP transport and MHC class I binding. Cell Mol Life Sci 62:1025–1037.

(40) Antoniou, A.N., Powis, S.J. and Elliott, T., 2003. Assembly and export of MHC class I peptide ligands. Current opinion in immunology, 15(1), pp.75–81.

(41) Srivastava S, Verma S, Kamthania M et al (2020) Computationally validated SARS-CoV-2 CTL and HTL multi-patch vaccines, designed by reverse epitomics approach, show potential to cover large ethnically distributed human population worldwide. J Biomol Struct Dyn:1–20. https://doi.org/10.1080/07391102.2020.1838329

(42) Srivastava S (2020) A novel method for designing multi-patch vaccine and diagnostic kit. Patent No. 202011037585

(43) Srivastava S (2020) A novel multi-patch vaccine design to combat SARS-CoV-2 and a method to prepare thereof. Patent No. 202011037939

(44) Srivastava S, Kolbe M (2021) Method for producing immunogenic compositions, vaccines as produced, and uses thereof. Patent No. PCT/IN2021/050841

(45) Wilson, S.S., Wiens, M.E. and Smith, J.G., 2013. Antiviral mechanisms of human defensins. Journal of molecular biology, 425(24), pp.4965–4980.

(46) Duits, L.A., Nibbering, P.H., Strijen, E., Vos, J.B., Mannesse-Lazeroms, S.P., Sterkenburg, M.A. and Hiemstra, P.S., 2003. Rhinovirus increases human β-defensin-2 and-3 mRNA expression in cultured bronchial epithelial cells. Pathogens and Disease, 38(1), pp.59–64.

(47) Yang, D., Biragyn, A., Kwak, L.W. and Oppenheim, J.J., 2002. Mammalian defensins in immunity: more than just microbicidal. Trends in immunology, 23(6), pp.291–296.

(48) Biragyn, A., Surenhu, M., Yang, D., Ruffini, P.A., Haines, B.A., Klyushnenkova, E., Oppenheim, J.J. and Kwak, L.W., 2001. Mediators of innate immunity that target immature, but not mature, dendritic cells induce antitumor immunity when genetically fused with nonimmunogenic tumor antigens. The Journal of Immunology, 167(11), pp.6644–6653.

(49) Duits, L.A., Nibbering, P.H., van Strijen, E., Vos, J.B., Mannesse-Lazeroms, S.P., van Sterkenburg, M.A. and Hiemstra, P.S., 2003. Rhinovirus increases human β-defensin-2 and-3 mRNA expression in cultured bronchial epithelial cells. FEMS Immunology & Medical Microbiology, 38(1), pp.59–64.

(50) Kohlgraf, K.G., Pingel, L.C., Dietrich, D.E. and Brogden, K.A., 2010. Defensins as anti-inflammatory compounds and mucosal adjuvants. Future microbiology, 5(1), pp.99–113.

(51) Delneste, Y., Beauvillain, C. and Jeannin, P., 2007. Innate immunity: structure and function of TLRs. Medecine sciences: M/S, 23(1), pp.67–73.

(52) Totura, A.L., Whitmore, A., Agnihothram, S., Schäfer, A., Katze, M.G., Heise, M.T. and Baric, R.S., 2015. Toll-like receptor 3 signaling via TRIF contributes to a protective innate immune response to severe acute respiratory syndrome coronavirus infection. MBio, 6(3), pp.e00638–15.

(53) Shaw, M.L., Cardenas, W.B., Zamarin, D., Palese, P. and Basler, C.F., 2005. Nuclear localization of the Nipah virus W protein allows for inhibition of both virus-and toll-like receptor 3-triggered signaling pathways. Journal of virology, 79(10), pp.6078–6088.

(54) Farina, C., Krumbholz, M., Giese, T., Hartmann, G., Aloisi, F. and Meinl, E., 2005. Preferential expression and function of Toll-like receptor 3 in human astrocytes. Journal of neuroimmunology, 159(1-2), pp.12–19.

(55) Peters B, Bulik S, Tampe R, Van Endert PM, Holzhutter HG. 2003. Identifying MHC class I epitopes by predicting the TAP transport efficiency of epitope precursors. J Immunol 171:1741–1749.

(56) Hoof, I., Peters, B., Sidney, J., Pedersen, L.E., Sette, A., Lund, O., Buus, S. and Nielsen, M., 2009. NetMHCpan, a method for MHC class I binding prediction beyond humans. Immunogenetics, 61(1), p.1.

(57) Calis JJA, Maybeno M, Greenbaum JA, Weiskopf D, De Silva AD, Sette A, Kesmir C, Peters B. 2013. Properties of MHC class I presented peptides that enhance immunogenicity. PloS Comp. Biol. 8(1):361.

(58) Wang, P., Sidney, J., Kim, Y., Sette, A., Lund, O., Nielsen, M. and Peters, B., 2010. Peptide binding predictions for HLA DR, DP and DQ molecules. BMC bioinformatics, 11(1), p.568.

(59) Sidney, J., Assarsson, E., Moore, C., Ngo, S., Pinilla, C., Sette, A. and Peters, B., 2008. Quantitative peptide binding motifs for 19 human and mouse MHC class I molecules derived using positional scanning combinatorial peptide libraries. Immunome research, 4(1), p.2.

(60) Nielsen, M., Lundegaard, C. and Lund, O., 2007. Prediction of MHC class II binding affinity using SMM-align, a novel stabilization matrix alignment method. BMC bioinformatics, 8(1), p.238.

(61) Sturniolo, T., Bono, E., Ding, J., Raddrizzani, L., Tuereci, O., Sahin, U., Braxenthaler, M., Gallazzi, F., Protti, M.P., Sinigaglia, F. and Hammer, J., 1999. Generation of tissue-specific and promiscuous HLA ligand databases using DNA microarrays and virtual HLA class II matrices. Nature biotechnology, 17(6), p.555.

(62) Gupta, S., Kapoor, P., Chaudhary, K., Gautam, A., Kumar, R., Raghava, G.P. and Open Source Drug Discovery Consortium, 2013. In silico approach for predicting toxicity of peptides and proteins. PLoS One, 8(9), p.e73957.

(63) Sievers, F., Wilm, A., Dineen, D., Gibson, T.J., Karplus, K., Li, W., Lopez, R., McWilliam, H., Remmert, M., Söding, J. and Thompson, J.D., and Higgins D.G., 2011. Fast, scalable generation of high-quality protein multiple sequence alignments using Clustal Omega. Molecular systems biology, 7(1), p.539.

(64) Bui H. H, Sidney J, Dinh K, Southwood S, Newman M. J, Sette A. 2006. Predicting population coverage of T-cell epitope-based diagnostics and vaccines. BMC Bioinformatics 17:153.

(65) Bui HH, Sidney J, Li W, Fusseder N, Sette A. 2007. Development of an epitope conservancy analysis tool to facilitate the design of epitope-based diagnostics and vaccines. BMC Bioinformatics 8:361.

(66) Gasteiger, E., Hoogland, C., Gattiker, A., Duvaud, S.E., Wilkins, M.R., Appel, R.D. and Bairoch, A., 2005. Protein identification and analysis tools on the ExPASy server (pp. 571–607). Humana Press.

(67) Nagpal, G., Gupta, S., Chaudhary, K., Dhanda, S.K., Prakash, S. and Raghava, G.P., 2015. VaccineDA: Prediction, design and genome-wide screening of oligodeoxynucleotide-based vaccine adjuvants. Scientific reports, 5, p.12478.

(68) Dhanda, S. K., Vir, P. & Raghava, G. P. Designing of interferon-gamma inducing MHC class-II binders. Biol. Direct. 8, 30 (2013).

(69) Dimitrov I, Naneva L, Doytchinova I, Bangov I. AllergenFP: allergenicity prediction by descriptor fingerprints. Bioinformatics. 2014 Mar 15;30(6):846–51. doi: 10.1093/bioinformatics/btt619. Epub 2013 Oct 27. PMID: 24167156. https://ddg-pharmfac.net/AllergenFP/

(70) Irini A Doytchinova and Darren R Flower. VaxiJen: a server for prediction of protective antigens, tumour antigens and subunit vaccines. BMC Bioinformatics. 2007 8:4. http://www.ddg-pharmfac.net/vaxijen/VaxiJen/VaxiJen.html

(71) W Zheng, C Zhang, Y Li, R Pearce, EW Bell, Y Zhang. Folding non-homology proteins by coupling deep-learning contact maps with I-TASSER assembly simulations. Cell Reports Methods, 1: 100014 (2021). https://zhanggroup.org/I-TASSER/

(72) Arnold K, Bordoli L, Kopp J, and Schwede T (2006). The SWISS-MODEL Workspace: A web-based environment for protein structure homology modelling. Bioinformatics.,22,195–201.

(73) Dong Xu and Yang Zhang. Improving the Physical Realism and Structural Accuracy of Protein Models by a Two-step Atomic-level Energy Minimization. Biophysical Journal, vol 101, 2525–2534 (2011).

(74) J. Ko, H. Park, L. Heo, and C. Seok, GalaxyWEB server for protein structure prediction and refinement, Nucleic Acids Res. 40 (W1), W294–W297 (2012).

(75) Wang, Z. and Xu, J., 2013. Predicting protein contact map using evolutionary and physical constraints by integer programming. Bioinformatics, 29(13), pp.i266–i273.

(76) Shin, W.H., Lee, G.R., Heo, L., Lee, H. and Seok, C., 2014. Prediction of protein structure and interaction by GALAXY protein modeling programs. Bio Design, 2(1), pp.1–11.

(77) Ramakrishnan, C. and Ramachandran, G.N., 1965. Stereochemical criteria for polypeptide and protein chain conformations: II. Allowed conformations for a pair of peptide units. Biophysical journal, 5(6), pp.909–933.

(78) J. V. Kringelum, C. Lundegaard, O. Lund, M. Nielsen. 2012. Reliable B cell epitope predictions: impacts of method development and improved benchmarking. PLoS Comput Biol. 8:(12):e1002829.

(79) Ponomarenko JV, Bui H, Li W, Fusseder N, Bourne PE, Sette A, Peters B. 2008. ElliPro: a new structure-based tool for the prediction of antibody epitopes. BMC Bioinformatics 9:514.

(80) Tovchigrechko A, Vakser IA. GRAMM-X public web server for protein-protein docking. Nucleic Acids Res. 2006; 34:W310–4.

(81) H. Bekker, H.J.C. Berendsen, E.J. Dijkstra, S. Achterop, R. van Drunen, D. van der Spoel, A. Sijbers, and H. Keegstra et al., “Gromacs: A parallel computer for molecular dynamics simulations”; pp. 252–256 in Physics computing 92. Edited by R.A. de Groot and J. Nadrchal. World Scientific, Singapore, 1993.

(82) Abraham, M. J., Murtola, T., Schulz, R., Pa ll, S., Smith, J. C., Hess, B., Lindahl, E. GROMACS: High performance molecular simulations through multi-level parallelism from lap-tops to supercomputers. SoftwareX 1–2:19–25, 2015.

(83) Jorgensen, W.L., Maxwell, D.S. and Tirado-Rives, J., 1996. Development and testing of the OPLS all-atom force field on conformational energetics and properties of organic liquids. Journal of the American Chemical Society, 118(45), pp.11225–11236.

(84) Morla, S., Makhija, A. & Kumar, S. Synonymous codon usage pattern in glycoprotein gene of rabies virus. Gene. 584, 1–6 (2016).

(85) Wu, X., Wu, S., Li, D., Zhang, J., Hou, L., Ma, J., Liu, W., Ren, D., Zhu, Y. and He, F., 2010. Computational identification of rare codons of Escherichia coli based on codon pairs preference. Bmc Bioinformatics, 11(1), p.61.

(86) Ojha, R., Pareek, A., Pandey, R.K., Prusty, D. and Prajapati, V.K., 2019. Strategic development of a next-generation multi-epitope vaccine to prevent Nipah virus zoonotic infection. ACS omega, 4(8), pp.13069–13079.

(87) Majee, P., Jain, N. and Kumar, A., 2021. Designing of a multi-epitope vaccine candidate against Nipah virus by in silico approach: a putative prophylactic solution for the deadly virus. Journal of Biomolecular Structure and Dynamics, 39(4), pp.1461–1480.

(88) Mohammed, A.A., Shantier, S.W., Mustafa, M.I., Osman, H.K., Elmansi, H.E., Osman, I.A.A., Mohammed, R.A., Abdelrhman, F.A., Elnnewery, M.E., Yousif, E.M. and Mustafa, M.M., 2020. Epitope-based peptide vaccine against glycoprotein G of Nipah henipavirus using immunoinformatics approaches. Journal of immunology research, 2020.

(89) Ali, M.T., Morshed, M.M. and Hassan, F., 2015. A computational approach for designing a universal epitope-based peptide vaccine against Nipah virus. Interdisciplinary Sciences: Computational Life Sciences, 7(2), pp.177–185.

(90) Soltan, M.A., Eldeen, M.A., Elbassiouny, N., Mohamed, I., El-Damasy, D.A., Fayad, E., Abu Ali, O.A., Raafat, N., Eid, R.A. and Al-Karmalawy, A.A., 2021. Proteome Based Approach Defines Candidates for Designing a Multitope Vaccine against the Nipah Virus. International Journal of Molecular Sciences, 22(17), p.9330.

(91) Hu W, Li F, Yang X, Li Z, Xia H, Li G, Wang Y, Zhang Z. A flexible peptide linker enhances the immunoreactivity of two copies HBsAg preS1 (21-47) fusion protein. J Biotechnol. 2004;107:83–90.

(92) Chen, X., Zaro, J.L. and Shen, W.C., 2013. Fusion protein linkers: property, design and functionality. Advanced drug delivery reviews, 65(10), pp.1357–1369.

(93) Hoover, D.M., Rajashankar, K.R., Blumenthal, R., Puri, A., Oppenheim, J.J., Chertov, O. and Lubkowski, J., 2000. The structure of human β-defensin-2 shows evidence of higher order oligomerization. Journal of Biological Chemistry, 275(42), pp.32911–32918. PDB ID: 1FD3.

(94) Srivastava, S., Chatziefthymiou, S.D. and Kolbe, M., 2022. Vaccines Targeting Numerous Coronavirus Antigens, Ensuring Broader Global Population Coverage: Multi-epitope and Multi-patch Vaccines. In Vaccine Design (pp. 149–175). Humana, New York, NY.

