## Supplementary for "Novel ‘GaEl antigenic patches’ identified by ‘reverse epitomics’ approach to design multi-patch vaccines against NIPAH infection, a silent threat to global human health"

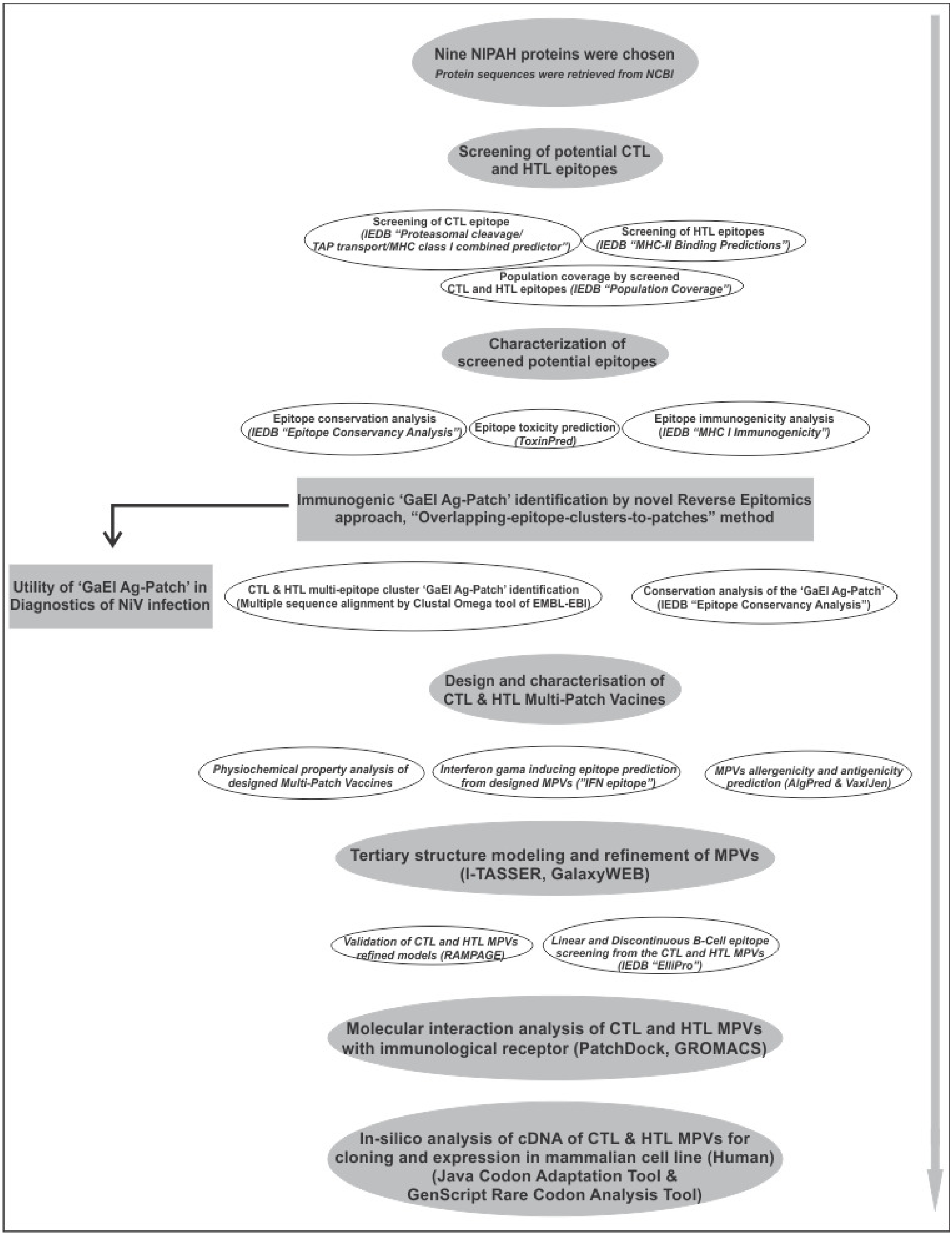

**Supplementary Figure S1.** Schematic representation of workflow and methodology.

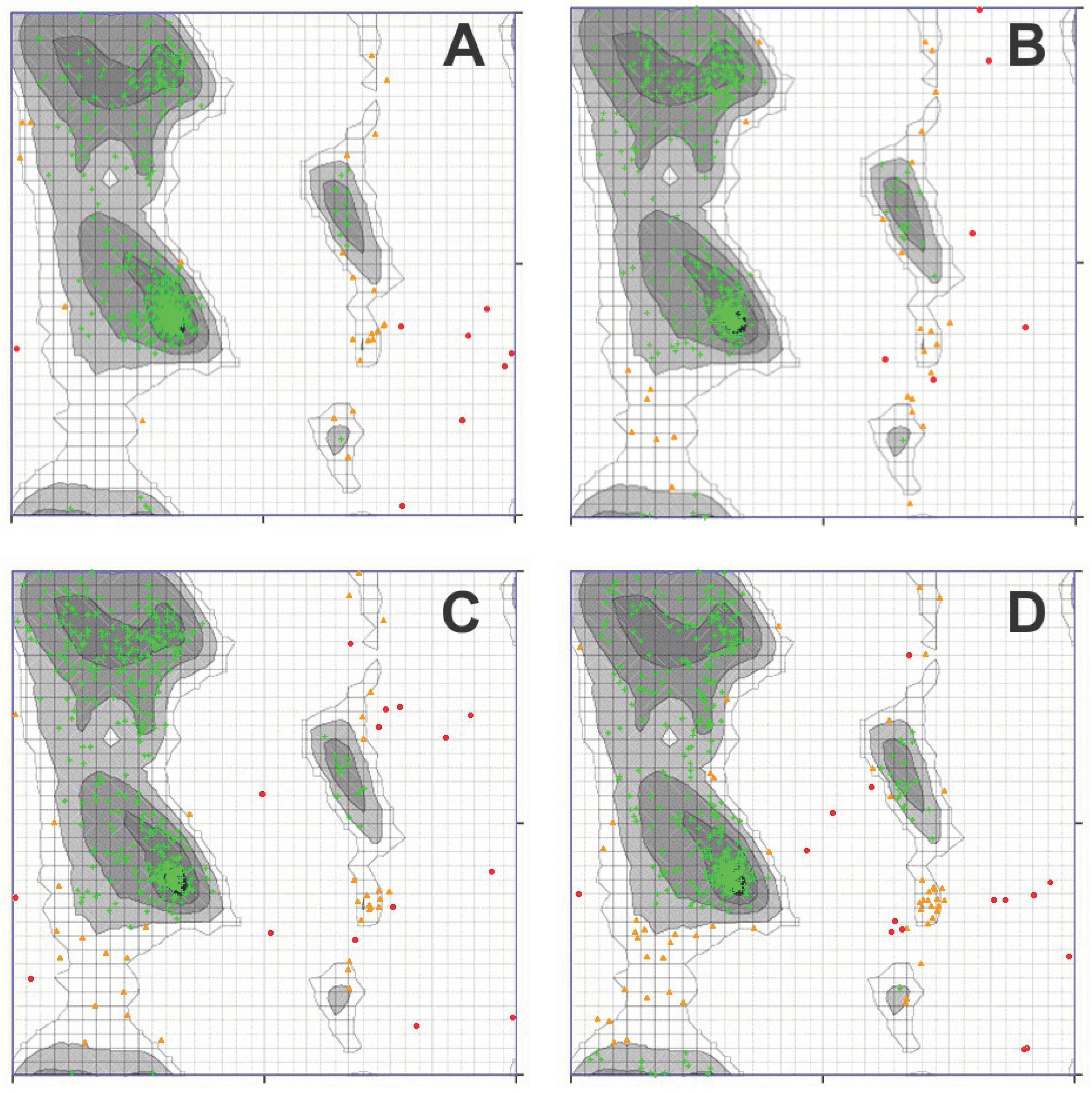

| # | Ramachandran Plot | (A) CTL-MPV-1 | (B) CTL-MPV-2 | (C) HTL-MPV-1 | (D) HTL-MPV-2 |
|---|---|---|---|---|---|
| 1 | Highly Preferred (GREEN) | 676 (95.480%) | 620 (94.946%) | 500 (90.909%) | 476 (87.339%) |
| 2 | Preferred (BROWN) | 24 (3.390%) | 27 (4.135%) | 35 (6.364%) | 54 (9.908%) |
| 3 | Questionable (RED) | 8 (1.130%) | 6 (0.919%) | 15 (2.727%) | 15 (2.752%) |

**Supplementary Figure S2.** RAMPAGE analysis for all the MPVs **(A)** CTL-MPV-1, **(B)** CTL-MPV-2, **(C)** HTL-MPV-1, **(D)** HTL-MPV-2.

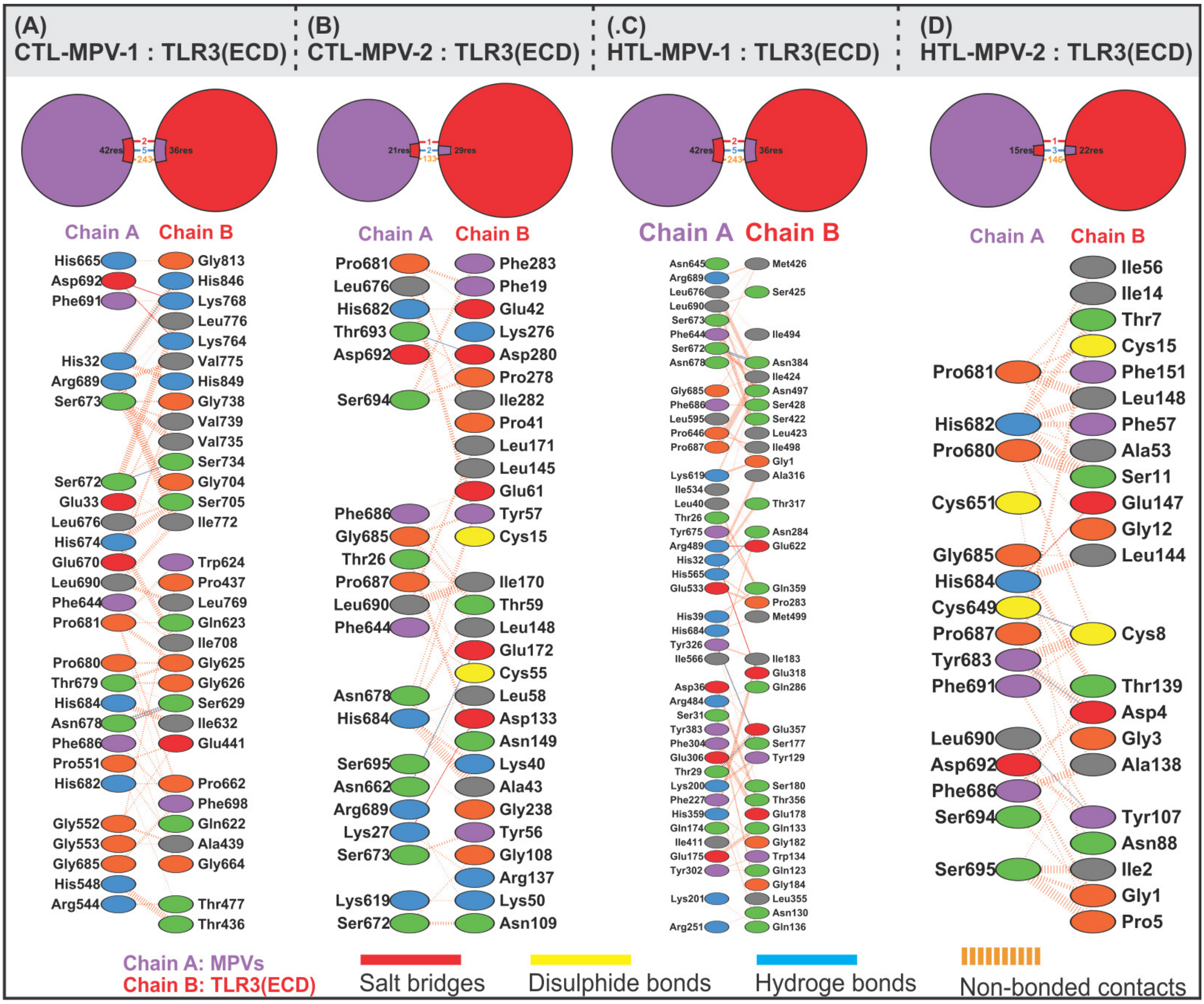

| # | Ramachandran Plot | (A) CTL-MPV-1 | (B) CTL-MPV-2 | (C) HTL-MPV-1 | (D) HTL-MPV-2 |
|---|---|---|---|---|---|
| 1 | Hydrogen Bonds | 5 | 2 | 5 | 3 |
| 2 | Salt Bridge | 2 | 1 | 2 | 1 |
| 3 | Disulphide bonds | 0 | 0 | 0 | 0 |
| 4 | Non-Bonded contacts | 243 | 133 | 243 | 146 |

**Supplementary Figure S3.** Molecular interaction between MPVs and TLR3 Ectodomain **(A)** CTL-MPV-1, **(B)** CTL-MPV-2, **(C)** HTL-MPV-1 and **(D)** HTL-MPV-2.

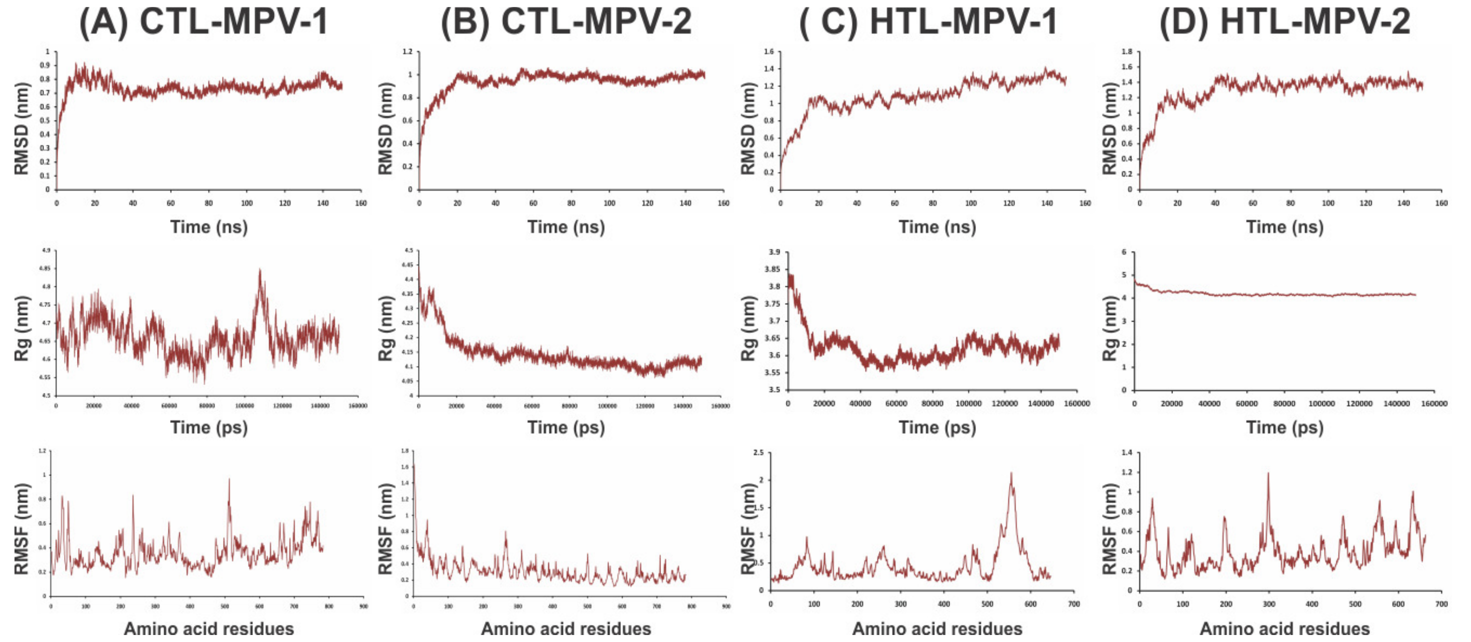

**Supplementary Figure S4.** Molecular Dynamics simulation study of the MPVs and TLR3-ECD complexes. The Root Mean Square Deviation (RMSD) for Cα, Backbone, the Radius of gyration for all the MPVs and TLR3-ECD complexes and the Root Mean Square Fluctuation (RMSF) in the conformation of residues of the MPVs in complex with TLR3-ECD are shown.

**Supplementary Table S1:** High Percentile Ranking CTL epitopes-HLA allele pairs screened from entire proteome of SARS-CoV-2 by the "MHC-I Processing Predictions" tool of IEDB. These epitopes were further utilized to identify the potentially immunogenic multiple epitope cluster based CTL Ag-Patches from the entire proteome of the SRAS-CoV-2. The screened epitopes are in consensus with the previous studies[38].

| S.No | Protein | Peptide | Allele | Peptide Length | Immunogenicity | Conservancy | Toxicity | Proteasome Score | TAP Score | MHC Score | Processing Score | Total Score | MHC IC50[nM] |
|---|---|---|---|---|---|---|---|---|---|---|---|---|---|
| 1 | c protein | ASILLTLFR | HLA-A*11:01 | 9 | 0.11682 | 100.00% (60/60) | Non-Toxin | 1.27 | 0.77 | -1.33 | 2.04 | 0.71 | 21.4 |
| 2 | c protein | ASILLTLFR | HLA-A*68:01 | 9 | 0.11682 | 100.00% (60/60) | Non-Toxin | 1.27 | 0.77 | -1.48 | 2.04 | 0.55 | 30.5 |
| 3 | c protein | ASILLTLFR | HLA-A*31:01 | 9 | 0.11682 | 100.00% (60/60) | Non-Toxin | 1.27 | 0.77 | -1.5 | 2.04 | 0.53 | 31.8 |
| 4 | c protein | EQTMGMYVLY | HLA-A*26:01 | 10 | -0.27722 | 100.00% (60/60) | Non-Toxin | 1.3 | 1.18 | -1.34 | 2.47 | 1.14 | 21.8 |
| 5 | c protein | EQTMGMYVLY | HLA-A*68:01 | 10 | -0.27722 | 100.00% (60/60) | Non-Toxin | 1.3 | 1.18 | -1.74 | 2.47 | 0.74 | 54.6 |
| 6 | c protein | MASILLTLF | HLA-B*58:01 | 9 | 0.08526 | 100.00% (60/60) | Non-Toxin | 1.2 | 1.16 | -1.04 | 2.35 | 1.31 | 11 |
| 7 | c protein | MASILLTLF | HLA-B*35:01 | 9 | 0.08526 | 100.00% (60/60) | Non-Toxin | 1.2 | 1.16 | -1.26 | 2.35 | 1.1 | 18.1 |
| 8 | c protein | MASILLTLF | HLA-B*53:01 | 9 | 0.08526 | 100.00% (60/60) | Non-Toxin | 1.2 | 1.16 | -1.56 | 2.35 | 0.8 | 35.9 |
| 9 | c protein | MASILLTLF | HLA-B*15:01 | 9 | 0.08526 | 100.00% (60/60) | Non-Toxin | 1.2 | 1.16 | -1.78 | 2.35 | 0.57 | 60.9 |
| 10 | c protein | MASILLTLFR | HLA-A*68:01 | 10 | 0.1542 | 100.00% (60/60) | Non-Toxin | 1.27 | 0.7 | -0.72 | 1.97 | 1.25 | 5.2 |
| 11 | c protein | MASILLTLFR | HLA-A*31:01 | 10 | 0.1542 | 100.00% (60/60) | Non-Toxin | 1.27 | 0.7 | -1.4 | 1.97 | 0.57 | 25.1 |
| 12 | c protein | MEMLKEETW | HLA-B*44:02 | 9 | -0.07673 | 21.67% (13/60) | Non-Toxin | 1.82 | 0.39 | -1.4 | 2.21 | 0.8 | 25.2 |
| 13 | c protein | MEMLKEETW | HLA-B*44:03 | 9 | -0.07673 | 21.67% (13/60) | Non-Toxin | 1.82 | 0.39 | -1.54 | 2.21 | 0.67 | 34.7 |
| 14 | c protein | MLKEETWRIY | HLA-B*15:01 | 10 | 0.496 | 21.67% (13/60) | Non-Toxin | 1.29 | 1.23 | -1.72 | 2.51 | 0.79 | 52.4 |
| 15 | c protein | MMASILLTL | HLA-A*02:03 | 9 | -0.02129 | 100.00% (60/60) | Non-Toxin | 1.45 | 0.45 | -0.61 | 1.9 | 1.29 | 4.1 |
| 16 | c protein | MMASILLTL | HLA-A*02:01 | 9 | -0.02129 | 100.00% (60/60) | Non-Toxin | 1.45 | 0.45 | -0.75 | 1.9 | 1.15 | 5.6 |
| 17 | c protein | MMASILLTL | HLA-A*02:06 | 9 | -0.02129 | 100.00% (60/60) | Non-Toxin | 1.45 | 0.45 | -0.9 | 1.9 | 1 | 7.9 |
| 18 | c protein | MMASILLTL | HLA-A*32:01 | 9 | -0.02129 | 100.00% (60/60) | Non-Toxin | 1.45 | 0.45 | -1.11 | 1.9 | 0.79 | 12.9 |
| 19 | c protein | MMASILLTL | HLA-B*15:01 | 9 | -0.02129 | 100.00% (60/60) | Non-Toxin | 1.45 | 0.45 | -1.38 | 1.9 | 0.52 | 23.9 |
| 20 | c protein | MMASILLTLF | HLA-B*15:01 | 10 | -0.01913 | 100.00% (60/60) | Non-Toxin | 1.2 | 1.13 | -0.76 | 2.33 | 1.57 | 5.8 |
| 21 | c protein | MMASILLTLF | HLA-A*02:03 | 10 | -0.01913 | 100.00% (60/60) | Non-Toxin | 1.2 | 1.13 | -1.37 | 2.33 | 0.96 | 23.6 |
| 22 | c protein | MMASILLTLF | HLA-B*58:01 | 10 | -0.01913 | 100.00% (60/60) | Non-Toxin | 1.2 | 1.13 | -1.53 | 2.33 | 0.8 | 33.7 |
| 23 | c protein | MMASILLTLF | HLA-A*02:01 | 10 | -0.01913 | 100.00% (60/60) | Non-Toxin | 1.2 | 1.13 | -1.71 | 2.33 | 0.62 | 51.5 |
| 24 | c protein | MMASILLTLF | HLA-A*32:01 | 10 | -0.01913 | 100.00% (60/60) | Non-Toxin | 1.2 | 1.13 | -1.77 | 2.33 | 0.56 | 59.5 |
| 25 | c protein | MMASILLTLF | HLA-A*23:01 | 10 | -0.01913 | 100.00% (60/60) | Non-Toxin | 1.2 | 1.13 | -1.8 | 2.33 | 0.53 | 62.6 |
| 26 | c protein | MMASILLTLFR | HLA-A*68:01 | 11 | 0.04981 | 100.00% (60/60) | Non-Toxin | 1.27 | 0.68 | -1.25 | 1.95 | 0.7 | 17.6 |
| 27 | c protein | MMEMLKEETW | HLA-B*58:01 | 10 | -0.16357 | 21.67% (13/60) | Non-Toxin | 1.82 | 0.45 | -1.73 | 2.27 | 0.53 | 54.1 |
| 28 | c protein | MTYNWTQW | HLA-B*58:01 | 8 | 0.14677 | 95.00% (57/60) | Non-Toxin | 1.24 | 0.45 | -1.17 | 1.7 | 0.53 | 14.8 |
| 29 | c protein | MTYNWTQWL | HLA-A*68:02 | 9 | 0.27619 | 95.00% (57/60) | Non-Toxin | 1.5 | 0.49 | -1.16 | 1.99 | 0.83 | 14.4 |
| 30 | c protein | MTYNWTQWL | HLA-A*32:01 | 9 | 0.27619 | 95.00% (57/60) | Non-Toxin | 1.5 | 0.49 | -1.24 | 1.99 | 0.75 | 17.5 |
| 31 | c protein | MYVLYLMQR | HLA-A*33:01 | 9 | -0.22768 | 100.00% (60/60) | Non-Toxin | 1 | 0.8 | -1.08 | 1.8 | 0.72 | 12 |
| 32 | c protein | NWTQWLQTLY | HLA-A*01:01 | 10 | 0.01818 | 100.00% (60/60) | Non-Toxin | 1.29 | 1.35 | -2.15 | 2.64 | 0.5 | 139.8 |
| 33 | c protein | QEQTMGMYVLY | HLA-B*44:03 | 11 | -0.28266 | 100.00% (60/60) | Non-Toxin | 1.3 | 1.19 | -1.95 | 2.49 | 0.54 | 88.5 |
| 34 | c protein | QTMGMYVLY | HLA-A*11:01 | 9 | -0.16902 | 100.00% (60/60) | Non-Toxin | 1.3 | 1.22 | -1.4 | 2.52 | 1.12 | 25 |
| 35 | c protein | QTMGMYVLY | HLA-A*68:01 | 9 | -0.16902 | 100.00% (60/60) | Non-Toxin | 1.3 | 1.22 | -1.42 | 2.52 | 1.1 | 26.4 |
| 36 | c protein | QTMGMYVLY | HLA-A*30:02 | 9 | -0.16902 | 100.00% (60/60) | Non-Toxin | 1.3 | 1.22 | -1.45 | 2.52 | 1.06 | 28.5 |
| 37 | c protein | QTMGMYVLY | HLA-A*26:01 | 9 | -0.16902 | 100.00% (60/60) | Non-Toxin | 1.3 | 1.22 | -1.52 | 2.52 | 1 | 32.9 |
| 38 | c protein | QTMGMYVLY | HLA-A*32:01 | 9 | -0.16902 | 100.00% (60/60) | Non-Toxin | 1.3 | 1.22 | -1.76 | 2.52 | 0.76 | 57.2 |
| 39 | c protein | RMMEMLKEETW | HLA-B*58:01 | 11 | -0.14662 | 21.67% (13/60) | Non-Toxin | 1.82 | 0.59 | -1.46 | 2.41 | 0.95 | 28.7 |
| 40 | c protein | WTQWLQTLY | HLA-A*01:01 | 9 | 0.09173 | 100.00% (60/60) | Non-Toxin | 1.29 | 1.3 | -1.24 | 2.59 | 1.34 | 17.5 |
| 41 | c protein | WTQWLQTLY | HLA-A*30:02 | 9 | 0.09173 | 100.00% (60/60) | Non-Toxin | 1.29 | 1.3 | -1.61 | 2.59 | 0.98 | 40.3 |

| | | | | | | | | | | | | | |
|---|---|---|---|---|---|---|---|---|---|---|---|---|---|
| 42 | c protein | YLMQRCCPML | HLA-B*08:01 | 10 | -0.33837 | 100.00% (60/60) | Non-Toxin | 1.41 | 0.47 | -1.1 | 1.87 | 0.77 | 12.6 |
| 43 | c protein | YLMQRCCPML | HLA-A*02:03 | 10 | -0.33837 | 100.00% (60/60) | Non-Toxin | 1.41 | 0.47 | -1.23 | 1.87 | 0.65 | 16.9 |
| 44 | c protein | YLMQRCCPML | HLA-A*02:01 | 10 | -0.33837 | 100.00% (60/60) | Non-Toxin | 1.41 | 0.47 | -1.29 | 1.87 | 0.59 | 19.3 |
| | **Protein** | Peptide | **Allele** | **Peptide Length** | **Immunogenicity** | **Conservancy** | **Toxicity** | **Proteasome Score** | **TAP Score** | **MHC Score** | **Processing Score** | **Total Score** | **MHC IC50[nM]** |
| 45 | fusion protein | AISQAFGGNY | HLA-A*30:02 | 10 | 0.03856 | 97.30% (36/37) | Non-Toxin | 1.35 | 1.37 | -1.72 | 2.71 | 0.99 | 52.6 |
| 46 | fusion protein | AQITAGVALY | HLA-B*15:01 | 10 | 0.21876 | 97.30% (36/37) | Non-Toxin | 1.18 | 1.46 | -1 | 2.64 | 1.64 | 10 |
| 47 | fusion protein | ETLLRTLGY | HLA-A*26:01 | 9 | 0.08262 | 97.30% (36/37) | Non-Toxin | 1.61 | 1.16 | -1.56 | 2.76 | 1.2 | 36.7 |
| 48 | fusion protein | FALSNGVLF | HLA-B*35:01 | 9 | -0.11611 | 100.00% (37/37) | Non-Toxin | 1.49 | 1.15 | -0.77 | 2.64 | 1.86 | 5.9 |
| 49 | fusion protein | FALSNGVLF | HLA-B*53:01 | 9 | -0.11611 | 100.00% (37/37) | Non-Toxin | 1.49 | 1.15 | -1.53 | 2.64 | 1.11 | 33.9 |
| 50 | fusion protein | FPILTEIQQAY | HLA-B*35:01 | 11 | 0.113 | 100.00% (37/37) | Non-Toxin | 1.17 | 1.1 | -0.96 | 2.28 | 1.32 | 9.1 |
| 51 | fusion protein | IIYVDLSSY | HLA-B*15:01 | 9 | -0.18478 | 45.95% (17/37) | Non-Toxin | 1.41 | 1.4 | -1.63 | 2.8 | 1.17 | 42.8 |
| 52 | fusion protein | IIYVDLSSY | HLA-A*30:02 | 9 | -0.18478 | 45.95% (17/37) | Non-Toxin | 1.41 | 1.4 | -1.8 | 2.8 | 1.01 | 62.8 |
| 53 | fusion protein | IIYVDLSSYY | HLA-A*30:02 | 10 | -0.24637 | 45.95% (17/37) | Non-Toxin | 1.24 | 1.4 | -1.61 | 2.64 | 1.03 | 40.3 |
| 54 | fusion protein | ISQAFGGNY | HLA-A*30:02 | 9 | 0.17249 | 97.30% (36/37) | Non-Toxin | 1.35 | 1.35 | -1.44 | 2.7 | 1.25 | 27.7 |
| 55 | fusion protein | KYLSDLLFVF | HLA-A*23:01 | 10 | -0.04679 | 97.30% (36/37) | Non-Toxin | 1.55 | 1.29 | -0.74 | 2.84 | 2.1 | 5.5 |
| 56 | fusion protein | KYLSDLLFVF | HLA-A*24:02 | 10 | -0.04679 | 97.30% (36/37) | Non-Toxin | 1.55 | 1.29 | -0.98 | 2.84 | 1.86 | 9.5 |
| 57 | fusion protein | LVVSSHVPRF | HLA-B*15:01 | 10 | -0.22293 | 100.00% (37/37) | Non-Toxin | 1.49 | 1.12 | -1.35 | 2.62 | 1.26 | 22.6 |
| 58 | fusion protein | MTIQAISQAF | HLA-B*15:01 | 10 | -0.13629 | 97.30% (36/37) | Non-Toxin | 1.4 | 1.16 | -0.92 | 2.57 | 1.64 | 8.4 |
| 59 | fusion protein | MTIQAISQAF | HLA-A*26:01 | 10 | -0.13629 | 97.30% (36/37) | Non-Toxin | 1.4 | 1.16 | -1.55 | 2.57 | 1.01 | 35.7 |
| 60 | fusion protein | RFALSNGVLF | HLA-A*23:01 | 10 | -0.1056 | 100.00% (37/37) | Non-Toxin | 1.49 | 1.22 | -1.29 | 2.71 | 1.43 | 19.3 |
| 61 | fusion protein | RFALSNGVLF | HLA-A*24:02 | 10 | -0.1056 | 100.00% (37/37) | Non-Toxin | 1.49 | 1.22 | -1.41 | 2.71 | 1.3 | 25.9 |
| 62 | fusion protein | RVRPTSSGDLY | HLA-A*30:02 | 11 | -0.23462 | 100.00% (37/37) | Non-Toxin | 1.42 | 1.51 | -1.67 | 2.94 | 1.26 | 47 |
| 63 | fusion protein | SEWISIVPNF | HLA-B*44:03 | 10 | 0.20004 | 97.30% (36/37) | Non-Toxin | 1.25 | 1.2 | -1.23 | 2.45 | 1.22 | 17 |
| 64 | fusion protein | SEWISIVPNF | HLA-A*23:01 | 10 | 0.20004 | 97.30% (36/37) | Non-Toxin | 1.25 | 1.2 | -1.29 | 2.45 | 1.15 | 19.6 |
| 65 | fusion protein | SEWISIVPNF | HLA-B*44:02 | 10 | 0.20004 | 97.30% (36/37) | Non-Toxin | 1.25 | 1.2 | -1.44 | 2.45 | 1 | 27.7 |
| 66 | fusion protein | SKYLSDLLFVF | HLA-A*23:01 | 11 | -0.05018 | 97.30% (36/37) | Non-Toxin | 1.55 | 1.25 | -1.79 | 2.79 | 1 | 61.9 |
| 67 | fusion protein | SYYIIVRVYF | HLA-A*23:01 | 10 | 0.38392 | 45.95% (17/37) | Non-Toxin | 1.46 | 1.34 | -1.45 | 2.8 | 1.35 | 28.2 |
| 68 | fusion protein | YLSDLLFVF | HLA-A*02:01 | 9 | 0.0703 | 97.30% (36/37) | Non-Toxin | 1.55 | 1.14 | -1.52 | 2.69 | 1.16 | 33.3 |
| 69 | fusion protein | YLSDLLFVF | HLA-A*02:06 | 9 | 0.0703 | 97.30% (36/37) | Non-Toxin | 1.55 | 1.14 | -1.54 | 2.69 | 1.15 | 34.6 |
| 70 | fusion protein | YLSDLLFVF | HLA-B*15:01 | 9 | 0.0703 | 97.30% (36/37) | Non-Toxin | 1.55 | 1.14 | -1.62 | 2.69 | 1.07 | 41.6 |
| 71 | fusion protein | YYIIVRVYF | HLA-A*23:01 | 9 | 0.29872 | 100.00% (37/37) | Non-Toxin | 1.46 | 1.26 | -1.63 | 2.72 | 1.08 | 43 |
| | **Protein** | Peptide | **Allele** | **Peptide Length** | **Immunogenicity** | **Conservancy** | **Toxicity** | **Proteasome Score** | **TAP Score** | **MHC Score** | **Processing Score** | **Total Score** | **MHC IC50[nM]** |
| 72 | Glycoprotein | AENPVFTVF | HLA-B*44:03 | 9 | 0.19402 | 100.00% (66/66) | Non-Toxin | 1.71 | 1.07 | -1.09 | 2.77 | 1.69 | 12.3 |
| 73 | Glycoprotein | AENPVFTVF | HLA-B*44:02 | 9 | 0.19402 | 100.00% (66/66) | Non-Toxin | 1.71 | 1.07 | -1.21 | 2.77 | 1.56 | 16.3 |
| 74 | Glycoprotein | AENPVFTVF | HLA-B*40:01 | 9 | 0.19402 | 100.00% (66/66) | Non-Toxin | 1.71 | 1.07 | -1.31 | 2.77 | 1.46 | 20.6 |
| 75 | Glycoprotein | AMDEGYFAY | HLA-A*01:01 | 9 | 0.25913 | 90.91% (60/66) | Non-Toxin | 1.43 | 1.33 | -1.61 | 2.76 | 1.15 | 40.7 |
| 76 | Glycoprotein | AVYNNEFYY | HLA-A*11:01 | 9 | 0.17688 | 90.91% (60/66) | Non-Toxin | 1.21 | 1.5 | -1.07 | 2.7 | 1.63 | 11.8 |
| 77 | Glycoprotein | AVYNNEFYY | HLA-A*30:02 | 9 | 0.17688 | 90.91% (60/66) | Non-Toxin | 1.21 | 1.5 | -1.11 | 2.7 | 1.59 | 13 |
| 78 | Glycoprotein | FLIDRINWI | HLA-A*02:03 | 9 | 0.36516 | 100.00% (66/66) | Non-Toxin | 1.3 | 0.3 | -0.28 | 1.6 | 1.32 | 1.9 |
| 79 | Glycoprotein | FLIDRINWI | HLA-A*02:01 | 9 | 0.36516 | 100.00% (66/66) | Non-Toxin | 1.3 | 0.3 | -0.48 | 1.6 | 1.12 | 3 |
| 80 | Glycoprotein | FLIDRINWI | HLA-A*02:06 | 9 | 0.36516 | 100.00% (66/66) | Non-Toxin | 1.3 | 0.3 | -0.49 | 1.6 | 1.1 | 3.1 |
| 81 | Glycoprotein | ILKPKLISY | HLA-B*15:01 | 9 | -0.28594 | 96.97% (64/66) | Non-Toxin | 1.43 | 1.27 | -1.52 | 2.7 | 1.18 | 33.4 |
| 82 | Glycoprotein | ILRSGLLKY | HLA-B*15:01 | 9 | -0.26247 | 98.48% (65/66) | Non-Toxin | 1.44 | 1.29 | -1.56 | 2.73 | 1.17 | 36.2 |
| 83 | Glycoprotein | IPSKVIKSY | HLA-B*35:01 | 9 | -0.38388 | 18.18% (12/66) | Non-Toxin | 1.44 | 1.19 | -1.34 | 2.63 | 1.29 | 21.7 |
| 84 | Glycoprotein | LAMDEGYFAY | HLA-B*35:01 | 10 | 0.214 | 90.91% (60/66) | Non-Toxin | 1.43 | 1.34 | -0.85 | 2.77 | 1.93 | 7 |
| 85 | Glycoprotein | LAMDEGYFAY | HLA-B*15:01 | 10 | 0.214 | 90.91% (60/66) | Non-Toxin | 1.43 | 1.34 | -1.55 | 2.77 | 1.22 | 35.7 |
| 86 | Glycoprotein | LAMDEGYFAY | HLA-A*01:01 | 10 | 0.214 | 90.91% (60/66) | Non-Toxin | 1.43 | 1.34 | -1.72 | 2.77 | 1.05 | 52.9 |

| | | | | | | | | | | | | | |
|---|---|---|---|---|---|---|---|---|---|---|---|---|---|
| 87 | Glycoprotein | LTVNPLVVNW | HLA-B*58:01 | 10 | 0.05521 | 19.70% (13/66) | Non-Toxin | 1.73 | 0.43 | -0.9 | 2.16 | 1.26 | 8 |
| 88 | Glycoprotein | NIMIIQNYTR | HLA-A*68:01 | 10 | 0.10719 | 98.48% (65/66) | Non-Toxin | 1.34 | 0.76 | -1 | 2.1 | 1.09 | 10.1 |
| 89 | Glycoprotein | SAVYNNEFY | HLA-B*35:01 | 9 | 0.15019 | 90.91% (60/66) | Non-Toxin | 1.49 | 1.35 | -1.68 | 2.84 | 1.16 | 47.7 |
| 90 | Glycoprotein | SAVYNNEFYY | HLA-A*11:01 | 10 | 0.18797 | 90.91% (60/66) | Non-Toxin | 1.21 | 1.35 | -1.51 | 2.55 | 1.04 | 32.5 |
| 91 | Glycoprotein | SAVYNNEFYY | HLA-A*30:02 | 10 | 0.18797 | 90.91% (60/66) | Non-Toxin | 1.21 | 1.35 | -1.56 | 2.55 | 0.99 | 36.3 |
| 92 | Glycoprotein | SMGIRPNSHY | HLA-B*15:01 | 10 | 0.05771 | 96.97% (64/66) | Non-Toxin | 1.18 | 1.3 | -1.47 | 2.47 | 1 | 29.6 |
| 93 | Glycoprotein | TVYHCSAVY | HLA-A*30:02 | 9 | -0.11974 | 100.00% (66/66) | Non-Toxin | 1.57 | 1.46 | -1.51 | 3.03 | 1.52 | 32.6 |
| 94 | Glycoprotein | TVYHCSAVY | HLA-B*35:01 | 9 | -0.11974 | 100.00% (66/66) | Non-Toxin | 1.57 | 1.46 | -1.53 | 3.03 | 1.5 | 34.2 |
| 95 | Glycoprotein | TVYHCSAVY | HLA-B*15:01 | 9 | -0.11974 | 100.00% (66/66) | Non-Toxin | 1.57 | 1.46 | -1.56 | 3.03 | 1.47 | 36.2 |
| | **Protein** | Peptide | **Allele** | **Peptide Length** | **Immunogenicity** | **Conservancy** | **Toxicity** | **Proteasome Score** | **TAP Score** | **MHC Score** | **Processing Score** | **Total Score** | **MHC IC50[nM]** |
| 96 | Matrix protein | FSLMDINPWL | HLA-A*02:01 | 10 | 0.08487 | 94.87% (37/39) | Non-Toxin | 1.53 | 0.4 | -0.6 | 1.92 | 1.32 | 4 |
| 97 | Matrix protein | FSLMDINPWL | HLA-A*02:06 | 10 | 0.08487 | 94.87% (37/39) | Non-Toxin | 1.53 | 0.4 | -0.78 | 1.92 | 1.14 | 6 |
| 98 | Matrix protein | IYMIPRTMLEF | HLA-A*23:01 | 11 | 0.03816 | 97.44% (38/39) | Non-Toxin | 1.4 | 1.24 | -1.34 | 2.64 | 1.29 | 22.1 |
| 99 | Matrix protein | IYMIPRTMLEF | HLA-A*24:02 | 11 | 0.03816 | 97.44% (38/39) | Non-Toxin | 1.4 | 1.24 | -1.55 | 2.64 | 1.09 | 35.5 |
| 100 | Matrix protein | KYYSVDYCR | HLA-A*31:01 | 9 | -0.14121 | 97.44% (38/39) | Non-Toxin | 1.09 | 0.87 | -0.93 | 1.96 | 1.03 | 8.6 |
| 101 | Matrix protein | KYYSVDYCRR | HLA-A*31:01 | 10 | -0.12461 | 97.44% (38/39) | Non-Toxin | 1.21 | 0.87 | -1.1 | 2.09 | 0.99 | 12.6 |
| 102 | Matrix protein | NAIAFNLLVY | HLA-B*35:01 | 10 | 0.19459 | 84.62% (33/39) | Non-Toxin | 1.25 | 1.4 | -1.55 | 2.65 | 1.1 | 35.3 |
| 103 | Matrix protein | NYMYLICYGF | HLA-A*23:01 | 10 | 0.02401 | 97.44% (38/39) | Non-Toxin | 1.29 | 1.32 | -1.09 | 2.61 | 1.52 | 12.2 |
| 104 | Matrix protein | NYMYLICYGF | HLA-A*24:02 | 10 | 0.02401 | 97.44% (38/39) | Non-Toxin | 1.29 | 1.32 | -1.34 | 2.61 | 1.27 | 21.7 |
| 105 | Matrix protein | SLMDINPWL | HLA-A*02:01 | 9 | 0.20889 | 94.87% (37/39) | Non-Toxin | 1.53 | 0.43 | -0.53 | 1.96 | 1.42 | 3.4 |
| 106 | Matrix protein | SLMDINPWL | HLA-A*02:06 | 9 | 0.20889 | 94.87% (37/39) | Non-Toxin | 1.53 | 0.43 | -0.66 | 1.96 | 1.29 | 4.6 |
| 107 | Matrix protein | SLMDINPWL | HLA-A*02:03 | 9 | 0.20889 | 94.87% (37/39) | Non-Toxin | 1.53 | 0.43 | -0.91 | 1.96 | 1.04 | 8.2 |
| 108 | Matrix protein | SMIPKYKIY | HLA-B*15:01 | 9 | -0.28568 | 97.44% (38/39) | Non-Toxin | 1.53 | 1.36 | -1.57 | 2.89 | 1.33 | 36.9 |
| 109 | Matrix protein | YMIPRTMLEF | HLA-B*15:01 | 10 | 0.00408 | 97.44% (38/39) | Non-Toxin | 1.4 | 1.18 | -0.79 | 2.57 | 1.79 | 6.1 |
| 110 | Matrix protein | YMIPRTMLEF | HLA-A*23:01 | 10 | 0.00408 | 97.44% (38/39) | Non-Toxin | 1.4 | 1.18 | -1.39 | 2.57 | 1.18 | 24.7 |
| | **Protein** | Peptide | **Allele** | **Peptide Length** | **Immunogenicity** | **Conservancy** | **Toxicity** | **Proteasome Score** | **TAP Score** | **MHC Score** | **Processing Score** | **Total Score** | **MHC IC50[nM]** |
| 111 | nucleocapsid | AAFTLISMY | HLA-A*30:02 | 9 | -0.05068 | 100.00% (13/13) | Non-Toxin | 1.32 | 1.39 | -1.7 | 2.72 | 1.01 | 50.5 |
| 112 | nucleocapsid | AAFTLISMY | HLA-B*15:01 | 9 | -0.05068 | 100.00% (13/13) | Non-Toxin | 1.32 | 1.39 | -2.02 | 2.72 | 0.69 | 105.9 |
| 113 | nucleocapsid | AAFTLISMY | HLA-B*35:01 | 9 | -0.05068 | 100.00% (13/13) | Non-Toxin | 1.32 | 1.39 | -2.15 | 2.72 | 0.56 | 141.5 |
| 114 | nucleocapsid | AESMKVGAAF | HLA-B*44:02 | 10 | -0.31242 | 100.00% (13/13) | Non-Toxin | 1.23 | 1.17 | -1.06 | 2.4 | 1.34 | 11.4 |
| 115 | nucleocapsid | AESMKVGAAF | HLA-B*44:03 | 10 | -0.31242 | 100.00% (13/13) | Non-Toxin | 1.23 | 1.17 | -1.18 | 2.4 | 1.22 | 15.1 |
| 116 | nucleocapsid | AESMKVGAAF | HLA-B*15:01 | 10 | -0.31242 | 100.00% (13/13) | Non-Toxin | 1.23 | 1.17 | -1.41 | 2.4 | 0.99 | 25.6 |
| 117 | nucleocapsid | ATNSPELRW | HLA-B*58:01 | 9 | -0.06424 | 100.00% (13/13) | Non-Toxin | 1.42 | 0.33 | -1.21 | 1.76 | 0.55 | 16.2 |
| 118 | nucleocapsid | DIFEEAASF | HLA-B*35:01 | 9 | 0.20944 | 100.00% (13/13) | Non-Toxin | 1.29 | 1.09 | -1.87 | 2.38 | 0.52 | 73.7 |
| 119 | nucleocapsid | DIFEEAASFR | HLA-A*68:01 | 10 | 0.23996 | 100.00% (13/13) | Non-Toxin | 0.98 | 0.63 | -1.07 | 1.62 | 0.55 | 11.8 |
| 120 | nucleocapsid | EEAASFRSY | HLA-B*44:03 | 9 | -0.05181 | 100.00% (13/13) | Non-Toxin | 1.17 | 1.08 | -1.35 | 2.26 | 0.91 | 22.3 |
| 121 | nucleocapsid | EEAASFRSY | HLA-B*44:02 | 9 | -0.05181 | 100.00% (13/13) | Non-Toxin | 1.17 | 1.08 | -1.37 | 2.26 | 0.89 | 23.5 |
| 122 | nucleocapsid | EIISDIGNY | HLA-A*26:01 | 9 | 0.04843 | 100.00% (13/13) | Non-Toxin | 1.19 | 1.29 | -0.6 | 2.47 | 1.87 | 4 |
| 123 | nucleocapsid | ESMKVGAAF | HLA-B*35:01 | 9 | -0.14602 | 100.00% (13/13) | Non-Toxin | 1.23 | 1.12 | -1.55 | 2.35 | 0.8 | 35.4 |
| 124 | nucleocapsid | ESMKVGAAF | HLA-A*26:01 | 9 | -0.14602 | 100.00% (13/13) | Non-Toxin | 1.23 | 1.12 | -1.7 | 2.35 | 0.65 | 50.2 |
| 125 | nucleocapsid | FAPGGYPLLW | HLA-B*53:01 | 10 | 0.03362 | 100.00% (13/13) | Non-Toxin | 1.48 | 0.4 | -1.21 | 1.89 | 0.68 | 16.1 |
| 126 | nucleocapsid | FAPGGYPLLW | HLA-B*58:01 | 10 | 0.03362 | 100.00% (13/13) | Non-Toxin | 1.48 | 0.4 | -1.27 | 1.89 | 0.61 | 18.8 |
| 127 | nucleocapsid | FMVEILIEV | HLA-A*02:06 | 9 | 0.40413 | 100.00% (13/13) | Non-Toxin | 0.99 | 0.22 | -0.32 | 1.22 | 0.9 | 2.1 |
| 128 | nucleocapsid | FMVEILIEV | HLA-A*02:01 | 9 | 0.40413 | 100.00% (13/13) | Non-Toxin | 0.99 | 0.22 | -0.34 | 1.22 | 0.88 | 2.2 |
| 129 | nucleocapsid | FMVEILIEV | HLA-A*02:03 | 9 | 0.40413 | 100.00% (13/13) | Non-Toxin | 0.99 | 0.22 | -0.43 | 1.22 | 0.79 | 2.7 |
| 130 | nucleocapsid | FTLISMYSER | HLA-A*68:01 | 10 | -0.28638 | 100.00% (13/13) | Non-Toxin | 0.84 | 0.67 | -0.88 | 1.51 | 0.63 | 7.5 |
| 131 | nucleocapsid | GLSSDQVAEL | HLA-A*02:03 | 10 | -0.18099 | 15.38% (2/13) | Non-Toxin | 1.68 | 0.37 | -1.46 | 2.05 | 0.58 | 29.1 |

| | | | | | | | | | | | | | |
|---|---|---|---|---|---|---|---|---|---|---|---|---|---|
| 132 | nucleocapsid | GMAGFFATI | HLA-A*02:03 | 9 | 0.3267 | 100.00% (13/13) | Non-Toxin | 1.35 | 0.22 | -0.63 | 1.57 | 0.93 | 4.3 |
| 133 | nucleocapsid | GPRAPYMVL | HLA-B*07:02 | 9 | -0.08219 | 100.00% (13/13) | Non-Toxin | 1.43 | 0.23 | -0.76 | 1.66 | 0.9 | 5.8 |
| 134 | nucleocapsid | GYLEPMYFRL | HLA-A*23:01 | 10 | 0.04091 | 100.00% (13/13) | Non-Toxin | 1.46 | 0.52 | -1.44 | 1.98 | 0.54 | 27.6 |
| 135 | nucleocapsid | GYPLLWSF | HLA-A*23:01 | 8 | 0.04333 | 100.00% (13/13) | Non-Toxin | 1.58 | 1.12 | -1.84 | 2.7 | 0.85 | 69.9 |
| 136 | nucleocapsid | GYPLLWSF | HLA-A*24:02 | 8 | 0.04333 | 100.00% (13/13) | Non-Toxin | 1.58 | 1.12 | -2.08 | 2.7 | 0.62 | 120.1 |
| 137 | nucleocapsid | IQTKFAPGGY | HLA-A*30:02 | 10 | -0.01434 | 100.00% (13/13) | Non-Toxin | 1.19 | 1.32 | -1.97 | 2.51 | 0.54 | 92.8 |
| 138 | nucleocapsid | KFMVEILIEV | HLA-A*02:03 | 10 | 0.37202 | 100.00% (13/13) | Non-Toxin | 0.99 | 0.34 | -0.8 | 1.33 | 0.53 | 6.3 |
| 139 | nucleocapsid | KFMVEILIEV | HLA-A*02:01 | 10 | 0.37202 | 100.00% (13/13) | Non-Toxin | 0.99 | 0.34 | -0.81 | 1.33 | 0.52 | 6.5 |
| 140 | nucleocapsid | KTPFVDSRAY | HLA-A*30:02 | 10 | 0.08681 | 100.00% (13/13) | Non-Toxin | 1.33 | 1.29 | -1.7 | 2.62 | 0.92 | 50.2 |
| 141 | nucleocapsid | LLSQSLSVRKF | HLA-B*15:01 | 11 | -0.54672 | 100.00% (13/13) | Non-Toxin | 1.46 | 1.17 | -2.03 | 2.63 | 0.6 | 107.7 |
| 142 | nucleocapsid | LLWSFAMGV | HLA-A*02:01 | 9 | -0.07214 | 100.00% (13/13) | Non-Toxin | 1.15 | 0.16 | -0.63 | 1.31 | 0.67 | 4.3 |
| 143 | nucleocapsid | LTNSLLNLR | HLA-A*68:01 | 9 | -0.20175 | 100.00% (13/13) | Non-Toxin | 0.98 | 0.65 | -1.1 | 1.63 | 0.53 | 12.7 |
| 144 | nucleocapsid | MAGFFATIR | HLA-A*68:01 | 9 | 0.39015 | 100.00% (13/13) | Non-Toxin | 1.08 | 0.6 | -0.99 | 1.69 | 0.7 | 9.7 |
| 145 | nucleocapsid | MAGFFATIRF | HLA-B*58:01 | 10 | 0.46 | 100.00% (13/13) | Non-Toxin | 1.45 | 1.06 | -1.67 | 2.51 | 0.84 | 47.1 |
| 146 | nucleocapsid | MAGFFATIRF | HLA-B*15:01 | 10 | 0.46 | 100.00% (13/13) | Non-Toxin | 1.45 | 1.06 | -1.95 | 2.51 | 0.56 | 89 |
| 147 | nucleocapsid | MAGFFATIRF | HLA-B*35:01 | 10 | 0.46 | 100.00% (13/13) | Non-Toxin | 1.45 | 1.06 | -2 | 2.51 | 0.51 | 100.8 |
| 148 | nucleocapsid | MVNGIPVMER | HLA-A*68:01 | 10 | 0.09996 | 100.00% (13/13) | Non-Toxin | 1.09 | 0.71 | -0.84 | 1.8 | 0.96 | 6.9 |
| 149 | nucleocapsid | MVNGIPVMERR | HLA-A*68:01 | 11 | 0.14044 | 100.00% (13/13) | Non-Toxin | 1.28 | 0.71 | -1.36 | 1.98 | 0.62 | 23.1 |
| 150 | nucleocapsid | NTIKSLMLLY | HLA-A*26:01 | 10 | -0.52684 | 100.00% (13/13) | Non-Toxin | 1.34 | 1.33 | -1.74 | 2.67 | 0.94 | 54.4 |
| 151 | nucleocapsid | NTIKSLMLLY | HLA-A*68:01 | 10 | -0.52684 | 100.00% (13/13) | Non-Toxin | 1.34 | 1.33 | -2.05 | 2.67 | 0.62 | 112.5 |
| 152 | nucleocapsid | NTIKSLMLLY | HLA-A*11:01 | 10 | -0.52684 | 100.00% (13/13) | Non-Toxin | 1.34 | 1.33 | -2.1 | 2.67 | 0.58 | 124.5 |
| 153 | nucleocapsid | QQKRVNPFF | HLA-B*15:01 | 9 | 0.07523 | 100.00% (13/13) | Non-Toxin | 1.54 | 1.16 | -1.95 | 2.69 | 0.74 | 90.1 |
| 154 | nucleocapsid | QTKFAPGGY | HLA-A*30:02 | 9 | 0.12386 | 100.00% (13/13) | Non-Toxin | 1.19 | 1.27 | -1.89 | 2.46 | 0.57 | 77.2 |
| 155 | nucleocapsid | RGYLEPMYFR | HLA-A*31:01 | 10 | -0.02568 | 100.00% (13/13) | Non-Toxin | 0.94 | 0.78 | -1.02 | 1.72 | 0.7 | 10.4 |
| 156 | nucleocapsid | RPGALIRSL | HLA-B*07:02 | 9 | 0.11187 | 100.00% (13/13) | Non-Toxin | 1.37 | 0.25 | -0.88 | 1.61 | 0.74 | 7.5 |
| 157 | nucleocapsid | RPGALIRSLL | HLA-B*07:02 | 10 | 0.07179 | 100.00% (13/13) | Non-Toxin | 1.57 | 0.25 | -0.88 | 1.81 | 0.94 | 7.5 |
| 158 | nucleocapsid | RSMGALNINR | HLA-A*31:01 | 10 | 0.10685 | 100.00% (13/13) | Non-Toxin | 1.16 | 0.74 | -1.07 | 1.89 | 0.82 | 11.7 |
| 159 | nucleocapsid | RSYQSKLGR | HLA-A*31:01 | 9 | -0.47142 | 100.00% (13/13) | Non-Toxin | 0.88 | 0.8 | -1.1 | 1.68 | 0.58 | 12.7 |
| 160 | nucleocapsid | RYPALALNEF | HLA-A*24:02 | 10 | 0.10567 | 100.00% (13/13) | Non-Toxin | 1.25 | 1.32 | -1.09 | 2.57 | 1.47 | 12.4 |
| 161 | nucleocapsid | RYPALALNEF | HLA-A*23:01 | 10 | 0.10567 | 100.00% (13/13) | Non-Toxin | 1.25 | 1.32 | -1.21 | 2.57 | 1.35 | 16.4 |
| 162 | nucleocapsid | RYPALALNEF | HLA-B*07:02 | 10 | 0.10567 | 100.00% (13/13) | Non-Toxin | 1.25 | 1.32 | -1.89 | 2.57 | 0.68 | 77.6 |
| 163 | nucleocapsid | RYPALALNEF | HLA-B*35:01 | 10 | 0.10567 | 100.00% (13/13) | Non-Toxin | 1.25 | 1.32 | -2.02 | 2.57 | 0.54 | 105.5 |
| 164 | nucleocapsid | SQSLSVRKF | HLA-B*15:01 | 9 | -0.26942 | 100.00% (13/13) | Non-Toxin | 1.46 | 1.17 | -1.84 | 2.63 | 0.79 | 69.4 |
| 165 | nucleocapsid | TIKSLMLLY | HLA-A*30:02 | 9 | -0.42841 | 100.00% (13/13) | Non-Toxin | 1.34 | 1.38 | -1.81 | 2.72 | 0.91 | 64.8 |
| 166 | nucleocapsid | TIKSLMLLYR | HLA-A*31:01 | 10 | -0.44023 | 100.00% (13/13) | Non-Toxin | 1.17 | 0.75 | -1.34 | 1.92 | 0.58 | 22 |
| 167 | nucleocapsid | TLFALDVIR | HLA-A*68:01 | 9 | 0.20005 | 100.00% (13/13) | Non-Toxin | 1.17 | 0.71 | -1.18 | 1.88 | 0.7 | 15.2 |
| 168 | nucleocapsid | TPFVDSRAY | HLA-B*35:01 | 9 | 0.01195 | 100.00% (13/13) | Non-Toxin | 1.33 | 1.22 | -0.79 | 2.56 | 1.77 | 6.1 |
| 169 | nucleocapsid | VEIISDIGNY | HLA-A*26:01 | 10 | 0.18772 | 100.00% (13/13) | Non-Toxin | 1.19 | 1.33 | -1.82 | 2.52 | 0.7 | 66.8 |
| 170 | nucleocapsid | VQQKRVNPF | HLA-B*15:01 | 9 | -0.17728 | 100.00% (13/13) | Non-Toxin | 1.01 | 1.12 | -1.64 | 2.13 | 0.49 | 43.4 |
| 171 | nucleocapsid | VQQKRVNPFF | HLA-B*15:01 | 10 | -0.11105 | 100.00% (13/13) | Non-Toxin | 1.54 | 1.12 | -1.98 | 2.66 | 0.68 | 95.6 |
| 172 | nucleocapsid | YLEPMYFRL | HLA-A*02:01 | 9 | -0.0241 | 100.00% (13/13) | Non-Toxin | 1.46 | 0.34 | -0.85 | 1.8 | 0.96 | 7 |
| 173 | nucleocapsid | YLEPMYFRL | HLA-A*02:06 | 9 | -0.0241 | 100.00% (13/13) | Non-Toxin | 1.46 | 0.34 | -1.13 | 1.8 | 0.67 | 13.4 |
| 174 | nucleocapsid | YPALALNEF | HLA-B*35:01 | 9 | 0.08224 | 100.00% (13/13) | Non-Toxin | 1.25 | 1.02 | -0.61 | 2.27 | 1.66 | 4.1 |
| 175 | nucleocapsid | YPALALNEF | HLA-B*53:01 | 9 | 0.08224 | 100.00% (13/13) | Non-Toxin | 1.25 | 1.02 | -1.33 | 2.27 | 0.94 | 21.6 |
| 176 | nucleocapsid | YPLLWSFAM | HLA-B*35:01 | 9 | 0.16687 | 100.00% (13/13) | Non-Toxin | 1.01 | -0.01 | -0.32 | 1 | 0.68 | 2.1 |
| 177 | nucleocapsid | YVQQKRVNPF | HLA-B*15:01 | 10 | -0.28684 | 100.00% (13/13) | Non-Toxin | 1.01 | 1.18 | -1.59 | 2.19 | 0.6 | 38.6 |
| | **Protein** | Peptide | **Allele** | **Peptide Length** | **Immunogenicity** | **Conservancy** | **Toxicity** | **Proteasome Score** | **TAP Score** | **MHC Score** | **Processing Score** | **Total Score** | **MHC IC50[nM]** |
| 178 | phosphoprotein | CLVSDAKMLSY | HLA-A*01:01 | 11 | -0.57469 | 59.26% (16/27) | Non-Toxin | 1.48 | 1.31 | -1.82 | 2.8 | 0.98 | 65.8 |

| | | | | | | | | | | | | | |
|---|---|---|---|---|---|---|---|---|---|---|---|---|---|
| 179 | phosphoprotein | DPVVTDVVY | HLA-B*35:01 | 9 | 0.17258 | 100.00% (27/27) | Non-Toxin | 1.74 | 1.05 | -1.4 | 2.8 | 1.4 | 25.1 |
| 180 | phosphoprotein | ELRSELIGY | HLA-A*26:01 | 9 | 0.06951 | 96.30% (26/27) | Non-Toxin | 1.2 | 1.27 | -1.92 | 2.46 | 0.55 | 82.8 |
| 181 | phosphoprotein | FPHDTDRLNY | HLA-B*35:01 | 10 | 0.1278 | 100.00% (27/27) | Non-Toxin | 1.41 | 1.06 | -1.06 | 2.47 | 1.41 | 11.4 |
| 182 | phosphoprotein | FPHDTDRLNY | HLA-B*53:01 | 10 | 0.1278 | 100.00% (27/27) | Non-Toxin | 1.41 | 1.06 | -1.9 | 2.47 | 0.57 | 79.2 |
| 183 | phosphoprotein | FSFDNVKNF | HLA-B*35:01 | 9 | -0.0929 | 100.00% (27/27) | Non-Toxin | 1.81 | 1.15 | -2.14 | 2.96 | 0.82 | 137 |
| 184 | phosphoprotein | FSFDNVKNF | HLA-B*15:01 | 9 | -0.0929 | 100.00% (27/27) | Non-Toxin | 1.81 | 1.15 | -2.25 | 2.96 | 0.71 | 178 |
| 185 | phosphoprotein | FSFDNVKNFR | HLA-A*68:01 | 10 | -0.04584 | 100.00% (27/27) | Non-Toxin | 0.97 | 0.69 | -0.58 | 1.66 | 1.08 | 3.8 |
| 186 | phosphoprotein | FSFDNVKNFR | HLA-A*31:01 | 10 | -0.04584 | 100.00% (27/27) | Non-Toxin | 0.97 | 0.69 | -1.15 | 1.66 | 0.51 | 14.1 |
| 187 | phosphoprotein | GLNPTAVPF | HLA-B*15:01 | 9 | 0.08973 | 62.96% (17/27) | Non-Toxin | 1.12 | 0.99 | -1.4 | 2.12 | 0.72 | 24.9 |
| 188 | phosphoprotein | IMPSDDFSNTF | HLA-B*35:01 | 11 | -0.15138 | 96.30% (26/27) | Non-Toxin | 1.53 | 1.13 | -2.14 | 2.66 | 0.52 | 139.2 |
| 189 | phosphoprotein | KLINLDMRL | HLA-A*02:01 | 9 | -0.07119 | 100.00% (27/27) | Non-Toxin | 1.61 | 0.51 | -1.21 | 2.11 | 0.91 | 16.1 |
| 190 | phosphoprotein | KLINLDMRL | HLA-A*02:03 | 9 | -0.07119 | 100.00% (27/27) | Non-Toxin | 1.61 | 0.51 | -1.54 | 2.11 | 0.57 | 34.9 |
| 191 | phosphoprotein | KMLSYAPEI | HLA-A*02:01 | 9 | -0.0877 | 59.26% (16/27) | Non-Toxin | 1.24 | 0.4 | -0.68 | 1.65 | 0.97 | 4.8 |
| 192 | phosphoprotein | KMLSYAPEI | HLA-A*02:06 | 9 | -0.0877 | 59.26% (16/27) | Non-Toxin | 1.24 | 0.4 | -0.82 | 1.65 | 0.83 | 6.6 |
| 193 | phosphoprotein | KMLSYAPEI | HLA-A*32:01 | 9 | -0.0877 | 59.26% (16/27) | Non-Toxin | 1.24 | 0.4 | -1.13 | 1.65 | 0.52 | 13.4 |
| 194 | phosphoprotein | KSRGIPIKK | HLA-A*30:01 | 9 | 0.15638 | 100.00% (27/27) | Non-Toxin | 1.09 | 0.22 | -0.73 | 1.31 | 0.58 | 5.4 |
| 195 | phosphoprotein | LDPVVTDVVY | HLA-B*35:01 | 10 | 0.19578 | 100.00% (27/27) | Non-Toxin | 1.74 | 1.11 | -1.34 | 2.85 | 1.51 | 22.1 |
| 196 | phosphoprotein | LFSFDNVKNF | HLA-A*23:01 | 10 | -0.06752 | 100.00% (27/27) | Non-Toxin | 1.81 | 1.2 | -2.34 | 3.01 | 0.67 | 216.4 |
| 197 | phosphoprotein | LVSDAKMLSY | HLA-A*01:01 | 10 | -0.4746 | 59.26% (16/27) | Non-Toxin | 1.48 | 1.33 | -1.26 | 2.81 | 1.55 | 18 |
| 198 | phosphoprotein | LVSDAKMLSY | HLA-B*15:01 | 10 | -0.4746 | 59.26% (16/27) | Non-Toxin | 1.48 | 1.33 | -1.94 | 2.81 | 0.87 | 87.9 |
| 199 | phosphoprotein | MPSDDFSNTF | HLA-B*35:01 | 10 | -0.03429 | 96.30% (26/27) | Non-Toxin | 1.53 | 0.99 | -0.88 | 2.52 | 1.64 | 7.6 |
| 200 | phosphoprotein | MPSDDFSNTF | HLA-B*53:01 | 10 | -0.03429 | 96.30% (26/27) | Non-Toxin | 1.53 | 0.99 | -1.15 | 2.52 | 1.37 | 14.2 |
| 201 | phosphoprotein | MPSDDFSNTFF | HLA-B*53:01 | 11 | 0.03819 | 96.30% (26/27) | Non-Toxin | 1.28 | 0.99 | -1.76 | 2.27 | 0.52 | 56.9 |
| 202 | phosphoprotein | NPTAVPFTLR | HLA-A*68:01 | 10 | 0.21785 | 51.85% (14/27) | Non-Toxin | 1.01 | 0.53 | -1.03 | 1.54 | 0.51 | 10.7 |
| 203 | phosphoprotein | NTFFPHDTDR | HLA-A*68:01 | 10 | 0.2431 | 100.00% (27/27) | Non-Toxin | 0.69 | 0.72 | -0.81 | 1.42 | 0.6 | 6.5 |
| 204 | phosphoprotein | REDLILPEL | HLA-B*40:01 | 9 | 0.16434 | 62.96% (17/27) | Non-Toxin | 1.61 | 0.41 | -1 | 2.02 | 1.02 | 9.9 |
| 205 | phosphoprotein | SPVIAEHYY | HLA-B*35:01 | 9 | 0.30481 | 96.30% (26/27) | Non-Toxin | 1.36 | 1.13 | -1.31 | 2.5 | 1.18 | 20.6 |
| 206 | phosphoprotein | SVLMGVINSI | HLA-A*02:03 | 10 | -0.08394 | 100.00% (27/27) | Non-Toxin | 1.25 | 0.25 | -0.86 | 1.5 | 0.64 | 7.2 |
| 207 | phosphoprotein | VLAKTNTAL | HLA-A*02:03 | 9 | -0.11697 | 100.00% (27/27) | Non-Toxin | 1.54 | 0.52 | -1.55 | 2.05 | 0.5 | 35.8 |
| 208 | phosphoprotein | VLMGVINSI | HLA-A*02:03 | 9 | 0.04046 | 100.00% (27/27) | Non-Toxin | 1.25 | 0.32 | -0.61 | 1.57 | 0.96 | 4.1 |
| 209 | phosphoprotein | VLMGVINSI | HLA-A*02:01 | 9 | 0.04046 | 100.00% (27/27) | Non-Toxin | 1.25 | 0.32 | -0.96 | 1.57 | 0.61 | 9.2 |
| 210 | phosphoprotein | VLMGVINSI | HLA-A*02:06 | 9 | 0.04046 | 100.00% (27/27) | Non-Toxin | 1.25 | 0.32 | -1.03 | 1.57 | 0.54 | 10.7 |
| 211 | phosphoprotein | VSDAKMLSY | HLA-A*01:01 | 9 | -0.43475 | 59.26% (16/27) | Non-Toxin | 1.48 | 1.26 | -0.94 | 2.74 | 1.8 | 8.7 |

| | **Protein** | Peptide | **Allele** | **Peptide Length** | **Immunogenicity** | **Conservancy** | **Toxicity** | **Proteasome Score** | **TAP Score** | **MHC Score** | **Processing Score** | **Total Score** | **MHC IC50[nM]** |
|---|---|---|---|---|---|---|---|---|---|---|---|---|---|
| 212 | polymerase | AEFFSFFRTF | HLA-B*44:03 | 10 | 0.28526 | 100.00% (10/10) | Non-Toxin | 1.49 | 1.17 | -1.05 | 2.66 | 1.61 | 11.3 |
| 213 | polymerase | AEFFSFFRTF | HLA-B*44:02 | 10 | 0.28526 | 100.00% (10/10) | Non-Toxin | 1.49 | 1.17 | -1.22 | 2.66 | 1.44 | 16.5 |
| 214 | polymerase | AEFFSFFRTF | HLA-A*23:01 | 10 | 0.28526 | 100.00% (10/10) | Non-Toxin | 1.49 | 1.17 | -1.44 | 2.66 | 1.22 | 27.7 |
| 215 | polymerase | AEFFSFFRTF | HLA-B*40:01 | 10 | 0.28526 | 100.00% (10/10) | Non-Toxin | 1.49 | 1.17 | -1.77 | 2.66 | 0.89 | 59.1 |
| 216 | polymerase | AEFFSFFRTF | HLA-A*24:02 | 10 | 0.28526 | 100.00% (10/10) | Non-Toxin | 1.49 | 1.17 | -2.06 | 2.66 | 0.6 | 115.6 |
| 217 | polymerase | AIRIATVYTW | HLA-B*58:01 | 10 | 0.28504 | 100.00% (10/10) | Non-Toxin | 1.73 | 0.55 | -1.77 | 2.28 | 0.51 | 59.2 |
| 218 | polymerase | ALIASGVGKY | HLA-B*15:01 | 10 | -0.10407 | 100.00% (10/10) | Non-Toxin | 1.13 | 1.3 | -1.6 | 2.42 | 0.83 | 39.7 |
| 219 | polymerase | ALIPAPIGGF | HLA-B*15:01 | 10 | 0.23302 | 100.00% (10/10) | Non-Toxin | 1.32 | 1.27 | -1.69 | 2.59 | 0.9 | 48.9 |
| 220 | polymerase | ALKSMSRCCF | HLA-B*15:01 | 10 | -0.59685 | 90.00% (9/10) | Non-Toxin | 1.34 | 1.15 | -1.72 | 2.49 | 0.77 | 52.9 |
| 221 | polymerase | ALWGLIDPL | HLA-A*02:01 | 9 | 0.23272 | 100.00% (10/10) | Non-Toxin | 1.07 | 0.62 | -0.92 | 1.7 | 0.78 | 8.3 |
| 222 | polymerase | ALWGLIDPL | HLA-A*02:06 | 9 | 0.23272 | 100.00% (10/10) | Non-Toxin | 1.07 | 0.62 | -1.15 | 1.7 | 0.55 | 14 |
| 223 | polymerase | APIGGFNYL | HLA-B*07:02 | 9 | 0.21288 | 100.00% (10/10) | Non-Toxin | 1.51 | 0.31 | -1.28 | 1.83 | 0.54 | 19.2 |
| 224 | polymerase | APIMKAHAIF | HLA-B*07:02 | 10 | -0.16417 | 100.00% (10/10) | Non-Toxin | 1.24 | 1.04 | -1.55 | 2.27 | 0.73 | 35.1 |
| 225 | polymerase | APIMKAHAIF | HLA-B*15:01 | 10 | -0.16417 | 100.00% (10/10) | Non-Toxin | 1.24 | 1.04 | -1.72 | 2.27 | 0.55 | 52.4 |

| | | | | | | | | | | | | | |
|---|---|---|---|---|---|---|---|---|---|---|---|---|---|
| 226 | polymerase | ATIPFLFLSAY | HLA-A*11:01 | 11 | 0.12204 | 100.00% (10/10) | Non-Toxin | 1.39 | 1.36 | -2.05 | 2.75 | 0.7 | 112.7 |
| 227 | polymerase | ATIPFLFLSAY | HLA-A*30:02 | 11 | 0.12204 | 100.00% (10/10) | Non-Toxin | 1.39 | 1.36 | -2.11 | 2.75 | 0.64 | 128.7 |
| 228 | polymerase | AYPECNNILF | HLA-A*24:02 | 10 | 0.1381 | 90.00% (9/10) | Non-Toxin | 1.47 | 1.27 | -1.97 | 2.75 | 0.78 | 92.6 |
| 229 | polymerase | AYPECNNILF | HLA-A*23:01 | 10 | 0.1381 | 90.00% (9/10) | Non-Toxin | 1.47 | 1.27 | -2.02 | 2.75 | 0.72 | 105.5 |
| 230 | polymerase | CSVDLARALR | HLA-A*68:01 | 10 | 0.13828 | 100.00% (10/10) | Non-Toxin | 1.06 | 0.68 | -1.24 | 1.74 | 0.5 | 17.4 |
| 231 | polymerase | DSYEYIINR | HLA-A*68:01 | 9 | 0.32977 | 100.00% (10/10) | Non-Toxin | 1.14 | 0.63 | -1.02 | 1.76 | 0.74 | 10.5 |
| 232 | polymerase | EAWYLASQR | HLA-A*68:01 | 9 | -0.11309 | 100.00% (10/10) | Non-Toxin | 0.98 | 0.63 | -1.05 | 1.61 | 0.56 | 11.3 |
| 233 | polymerase | EFFSFFRTF | HLA-A*23:01 | 9 | 0.16209 | 100.00% (10/10) | Non-Toxin | 1.49 | 1.2 | -1.34 | 2.69 | 1.35 | 22 |
| 234 | polymerase | EFFSFFRTF | HLA-A*24:02 | 9 | 0.16209 | 100.00% (10/10) | Non-Toxin | 1.49 | 1.2 | -1.96 | 2.69 | 0.73 | 90.6 |
| 235 | polymerase | EIINIHECR | HLA-A*68:01 | 9 | 0.24974 | 100.00% (10/10) | Non-Toxin | 1.31 | 0.58 | -0.77 | 1.89 | 1.12 | 5.9 |
| 236 | polymerase | EIINIHECR | HLA-A*33:01 | 9 | 0.24974 | 100.00% (10/10) | Non-Toxin | 1.31 | 0.58 | -1.2 | 1.89 | 0.68 | 16 |
| 237 | polymerase | EILKSEISY | HLA-B*35:01 | 9 | -0.27179 | 100.00% (10/10) | Non-Toxin | 1.47 | 1.18 | -2 | 2.65 | 0.65 | 100.9 |
| 238 | polymerase | EIYGLPGFF | HLA-A*26:01 | 9 | 0.10866 | 100.00% (10/10) | Non-Toxin | 1.3 | 1.09 | -1.88 | 2.39 | 0.51 | 75.8 |
| 239 | polymerase | ESMAIFAER | HLA-A*68:01 | 9 | 0.31369 | 100.00% (10/10) | Non-Toxin | 1 | 0.58 | -0.48 | 1.58 | 1.1 | 3 |
| 240 | polymerase | ESMAIFAER | HLA-A*33:01 | 9 | 0.31369 | 100.00% (10/10) | Non-Toxin | 1 | 0.58 | -0.72 | 1.58 | 0.85 | 5.3 |
| 241 | polymerase | ESMAIFAERL | HLA-A*68:02 | 10 | 0.37754 | 100.00% (10/10) | Non-Toxin | 1.47 | 0.35 | -1.17 | 1.82 | 0.65 | 14.7 |
| 242 | polymerase | ETDDYNGIY | HLA-A*01:01 | 9 | 0.12619 | 100.00% (10/10) | Non-Toxin | 1.41 | 1.12 | -0.91 | 2.53 | 1.62 | 8.1 |
| 243 | polymerase | ETIISTHLF | HLA-A*26:01 | 9 | 0.07338 | 100.00% (10/10) | Non-Toxin | 1.09 | 1.04 | -1.29 | 2.12 | 0.83 | 19.6 |
| 244 | polymerase | ETTWIGNLDSY | HLA-A*26:01 | 11 | 0.30341 | 100.00% (10/10) | Non-Toxin | 1.29 | 1.09 | -1.78 | 2.38 | 0.61 | 59.8 |
| 245 | polymerase | EVDNNHLIY | HLA-A*01:01 | 9 | 0.09324 | 100.00% (10/10) | Non-Toxin | 1.3 | 1.17 | -1.25 | 2.47 | 1.22 | 17.9 |
| 246 | polymerase | EVDNNHLIY | HLA-B*35:01 | 9 | 0.09324 | 100.00% (10/10) | Non-Toxin | 1.3 | 1.17 | -1.9 | 2.47 | 0.58 | 78.6 |
| 247 | polymerase | EVIDTTTML | HLA-A*68:02 | 9 | 0.07002 | 70.00% (7/10) | Non-Toxin | 1.88 | 0.42 | -1.18 | 2.31 | 1.12 | 15.2 |
| 248 | polymerase | EVIDTTTML | HLA-A*26:01 | 9 | 0.07002 | 70.00% (7/10) | Non-Toxin | 1.88 | 0.42 | -1.4 | 2.31 | 0.9 | 25.3 |
| 249 | polymerase | EVIDTTTMLR | HLA-A*68:01 | 10 | 0.02298 | 70.00% (7/10) | Non-Toxin | 0.9 | 0.65 | -0.8 | 1.55 | 0.75 | 6.3 |
| 250 | polymerase | EYIINRTAGR | HLA-A*33:01 | 10 | 0.31058 | 100.00% (10/10) | Non-Toxin | 1.16 | 0.68 | -1.25 | 1.84 | 0.59 | 17.8 |
| 251 | polymerase | FAERLDEIY | HLA-B*35:01 | 9 | 0.25692 | 100.00% (10/10) | Non-Toxin | 1.41 | 1.19 | -1.5 | 2.6 | 1.1 | 31.9 |
| 252 | polymerase | FIDDLLSIL | HLA-A*02:06 | 9 | -0.05358 | 100.00% (10/10) | Non-Toxin | 1.7 | 0.37 | -1.23 | 2.07 | 0.85 | 16.8 |
| 253 | polymerase | FIDDLLSIL | HLA-A*02:01 | 9 | -0.05358 | 100.00% (10/10) | Non-Toxin | 1.7 | 0.37 | -1.32 | 2.07 | 0.75 | 21.1 |
| 254 | polymerase | FIYSKKIHY | HLA-B*15:01 | 9 | -0.44945 | 90.00% (9/10) | Non-Toxin | 1.38 | 1.39 | -1.83 | 2.77 | 0.94 | 67.5 |
| 255 | polymerase | FLDWASDPY | HLA-B*35:01 | 9 | 0.1247 | 100.00% (10/10) | Non-Toxin | 1.08 | 1.18 | -1.15 | 2.26 | 1.11 | 14.2 |
| 256 | polymerase | FLDWASDPY | HLA-A*01:01 | 9 | 0.1247 | 100.00% (10/10) | Non-Toxin | 1.08 | 1.18 | -1.3 | 2.26 | 0.96 | 20 |
| 257 | polymerase | FLIVDPELFAL | HLA-A*02:01 | 11 | 0.30426 | 100.00% (10/10) | Non-Toxin | 1.59 | 0.34 | -1.23 | 1.93 | 0.71 | 16.8 |
| 258 | polymerase | FLIVDPELFAL | HLA-A*02:06 | 11 | 0.30426 | 100.00% (10/10) | Non-Toxin | 1.59 | 0.34 | -1.28 | 1.93 | 0.65 | 18.9 |
| 259 | polymerase | FLMDRRVIL | HLA-B*08:01 | 9 | 0.17704 | 100.00% (10/10) | Non-Toxin | 1.55 | 0.44 | -0.9 | 1.99 | 1.09 | 7.9 |
| 260 | polymerase | FLMDRRVIL | HLA-A*02:03 | 9 | 0.17704 | 100.00% (10/10) | Non-Toxin | 1.55 | 0.44 | -1 | 1.99 | 0.99 | 10 |
| 261 | polymerase | FLMDRRVIL | HLA-A*02:01 | 9 | 0.17704 | 100.00% (10/10) | Non-Toxin | 1.55 | 0.44 | -1.12 | 1.99 | 0.86 | 13.3 |
| 262 | polymerase | FNYLNLSRIF | HLA-A*23:01 | 10 | -0.06375 | 100.00% (10/10) | Non-Toxin | 1.29 | 1.06 | -1.61 | 2.35 | 0.74 | 40.9 |
| 263 | polymerase | FPISRLFNMY | HLA-B*35:01 | 10 | -0.08153 | 100.00% (10/10) | Non-Toxin | 1.46 | 1.11 | -0.98 | 2.57 | 1.59 | 9.6 |
| 264 | polymerase | FPISRLFNMY | HLA-B*53:01 | 10 | -0.08153 | 100.00% (10/10) | Non-Toxin | 1.46 | 1.11 | -1.6 | 2.57 | 0.97 | 39.9 |
| 265 | polymerase | FPLWSTEEL | HLA-B*35:01 | 9 | 0.23773 | 100.00% (10/10) | Non-Toxin | 1.65 | 0.19 | -0.91 | 1.84 | 0.93 | 8.1 |
| 266 | polymerase | FPLWSTEEL | HLA-B*53:01 | 9 | 0.23773 | 100.00% (10/10) | Non-Toxin | 1.65 | 0.19 | -1.29 | 1.84 | 0.56 | 19.3 |
| 267 | polymerase | FPVMGNRIY | HLA-B*35:01 | 9 | -0.01495 | 100.00% (10/10) | Non-Toxin | 1.57 | 1.15 | -0.51 | 2.72 | 2.22 | 3.2 |
| 268 | polymerase | FPVMGNRIY | HLA-B*53:01 | 9 | -0.01495 | 100.00% (10/10) | Non-Toxin | 1.57 | 1.15 | -1.75 | 2.72 | 0.97 | 56.4 |
| 269 | polymerase | FVIFYASLTY | HLA-B*15:01 | 10 | 0.05309 | 100.00% (10/10) | Non-Toxin | 1.22 | 1.26 | -1.28 | 2.48 | 1.2 | 19 |
| 270 | polymerase | FVIFYASLTY | HLA-B*35:01 | 10 | 0.05309 | 100.00% (10/10) | Non-Toxin | 1.22 | 1.26 | -1.66 | 2.48 | 0.83 | 45.2 |
| 271 | polymerase | FVNDENFDPY | HLA-B*35:01 | 10 | 0.23386 | 100.00% (10/10) | Non-Toxin | 1.05 | 1.27 | -1.3 | 2.31 | 1.01 | 20.1 |
| 272 | polymerase | FVNDENFDPY | HLA-A*01:01 | 10 | 0.23386 | 100.00% (10/10) | Non-Toxin | 1.05 | 1.27 | -1.78 | 2.31 | 0.53 | 60.4 |
| 273 | polymerase | FYASLTYLR | HLA-A*33:01 | 9 | -0.13763 | 100.00% (10/10) | Non-Toxin | 1.06 | 0.73 | -0.95 | 1.79 | 0.84 | 8.9 |

| | | | | | | | | | | | | | |
|---|---|---|---|---|---|---|---|---|---|---|---|---|---|
| 274 | polymerase | FYASLTYLRR | HLA-A*33:01 | 10 | -0.10937 | 100.00% (10/10) | Non-Toxin | 1.29 | 0.73 | -1.32 | 2.02 | 0.7 | 21 |
| 275 | polymerase | GIQFDCFMEL | HLA-A*02:06 | 10 | 0.0698 | 100.00% (10/10) | Non-Toxin | 1.55 | 0.38 | -1.32 | 1.93 | 0.61 | 20.7 |
| 276 | polymerase | GLIDPLFPV | HLA-A*02:06 | 9 | 0.1366 | 100.00% (10/10) | Non-Toxin | 0.88 | 0.07 | -0.26 | 0.95 | 0.69 | 1.8 |
| 277 | polymerase | GLIDPLFPV | HLA-A*02:01 | 9 | 0.1366 | 100.00% (10/10) | Non-Toxin | 0.88 | 0.07 | -0.28 | 0.95 | 0.67 | 1.9 |
| 278 | polymerase | GLIDPLFPV | HLA-A*02:03 | 9 | 0.1366 | 100.00% (10/10) | Non-Toxin | 0.88 | 0.07 | -0.36 | 0.95 | 0.59 | 2.3 |
| 279 | polymerase | GQSISIKWAF | HLA-B*15:01 | 10 | 0.05552 | 100.00% (10/10) | Non-Toxin | 1.4 | 1.07 | -1.78 | 2.47 | 0.7 | 59.9 |
| 280 | polymerase | GSGSMMLLY | HLA-A*30:02 | 9 | -0.50761 | 100.00% (10/10) | Non-Toxin | 1.14 | 1.07 | -1.6 | 2.21 | 0.61 | 39.9 |
| 281 | polymerase | GTKFDTVSAF | HLA-B*15:01 | 10 | 0.0293 | 100.00% (10/10) | Non-Toxin | 1.24 | 1.02 | -1.75 | 2.25 | 0.51 | 55.6 |
| 282 | polymerase | HLLAEFFSF | HLA-A*23:01 | 9 | 0.24561 | 100.00% (10/10) | Non-Toxin | 1.14 | 1.08 | -1.41 | 2.22 | 0.8 | 25.9 |
| 283 | polymerase | HLLAEFFSF | HLA-A*32:01 | 9 | 0.24561 | 100.00% (10/10) | Non-Toxin | 1.14 | 1.08 | -1.47 | 2.22 | 0.74 | 29.7 |
| 284 | polymerase | HLLAEFFSF | HLA-B*15:01 | 9 | 0.24561 | 100.00% (10/10) | Non-Toxin | 1.14 | 1.08 | -1.64 | 2.22 | 0.58 | 43.4 |
| 285 | polymerase | HLLAEFFSFF | HLA-B*15:01 | 10 | 0.28625 | 100.00% (10/10) | Non-Toxin | 1.37 | 1.08 | -1.58 | 2.45 | 0.87 | 37.8 |
| 286 | polymerase | HMLADKVLEY | HLA-B*15:01 | 10 | -0.06463 | 100.00% (10/10) | Non-Toxin | 1.56 | 1.22 | -1.74 | 2.78 | 1.04 | 54.5 |
| 287 | polymerase | HMLADKVLEY | HLA-A*30:02 | 10 | -0.06463 | 100.00% (10/10) | Non-Toxin | 1.56 | 1.22 | -2.26 | 2.78 | 0.52 | 183.1 |
| 288 | polymerase | HTEFNPHNHY | HLA-A*30:02 | 10 | 0.17709 | 100.00% (10/10) | Non-Toxin | 1.36 | 1.14 | -1.7 | 2.5 | 0.8 | 50.3 |
| 289 | polymerase | IATVYTWAY | HLA-B*35:01 | 9 | 0.29688 | 100.00% (10/10) | Non-Toxin | 1.56 | 1.34 | -0.93 | 2.9 | 1.97 | 8.6 |
| 290 | polymerase | IATVYTWAY | HLA-B*58:01 | 9 | 0.29688 | 100.00% (10/10) | Non-Toxin | 1.56 | 1.34 | -2.04 | 2.9 | 0.87 | 109 |
| 291 | polymerase | IATVYTWAY | HLA-B*53:01 | 9 | 0.29688 | 100.00% (10/10) | Non-Toxin | 1.56 | 1.34 | -2.21 | 2.9 | 0.69 | 162.7 |
| 292 | polymerase | IATVYTWAY | HLA-A*30:02 | 9 | 0.29688 | 100.00% (10/10) | Non-Toxin | 1.56 | 1.34 | -2.23 | 2.9 | 0.68 | 168.5 |
| 293 | polymerase | IFYASLTYLR | HLA-A*31:01 | 10 | -0.10679 | 100.00% (10/10) | Non-Toxin | 1.06 | 0.83 | -0.97 | 1.89 | 0.92 | 9.4 |
| 294 | polymerase | IFYASLTYLR | HLA-A*33:01 | 10 | -0.10679 | 100.00% (10/10) | Non-Toxin | 1.06 | 0.83 | -1.18 | 1.89 | 0.71 | 15.3 |
| 295 | polymerase | IFYASLTYLRR | HLA-A*31:01 | 11 | -0.07853 | 100.00% (10/10) | Non-Toxin | 1.29 | 0.83 | -1.59 | 2.12 | 0.53 | 38.9 |
| 296 | polymerase | IHLLAEFFSF | HLA-A*23:01 | 10 | 0.23318 | 100.00% (10/10) | Non-Toxin | 1.14 | 1.18 | -1.79 | 2.32 | 0.52 | 62 |
| 297 | polymerase | IIMLNEAMNY | HLA-B*15:01 | 10 | -0.09211 | 100.00% (10/10) | Non-Toxin | 1.48 | 1.37 | -2.15 | 2.85 | 0.7 | 141.9 |
| 298 | polymerase | IIMLNEAMNY | HLA-A*30:02 | 10 | -0.09211 | 100.00% (10/10) | Non-Toxin | 1.48 | 1.37 | -2.32 | 2.85 | 0.53 | 210.3 |
| 299 | polymerase | IISTHLFIY | HLA-A*30:02 | 9 | 0.18298 | 90.00% (9/10) | Non-Toxin | 1.26 | 1.34 | -1.85 | 2.6 | 0.74 | 71 |
| 300 | polymerase | ILNIDNIHLL | HLA-A*02:03 | 10 | 0.29322 | 100.00% (10/10) | Non-Toxin | 1.57 | 0.42 | -1.36 | 1.99 | 0.63 | 22.7 |
| 301 | polymerase | ILNIDNIHLL | HLA-A*02:01 | 10 | 0.29322 | 100.00% (10/10) | Non-Toxin | 1.57 | 0.42 | -1.4 | 1.99 | 0.59 | 25.1 |
| 302 | polymerase | ILNPNLICIF | HLA-B*15:01 | 10 | 0.12718 | 100.00% (10/10) | Non-Toxin | 1.35 | 1.08 | -1.62 | 2.43 | 0.81 | 41.6 |
| 303 | polymerase | ILVEHSHLI | HLA-A*02:03 | 9 | 0.01074 | 100.00% (10/10) | Non-Toxin | 1.21 | 0.21 | -0.9 | 1.42 | 0.52 | 8 |
| 304 | polymerase | IMKKSFKAY | HLA-B*15:01 | 9 | -0.49704 | 100.00% (10/10) | Non-Toxin | 1.59 | 1.39 | -1.07 | 2.98 | 1.92 | 11.7 |
| 305 | polymerase | IMKKSFKAY | HLA-A*30:02 | 9 | -0.49704 | 100.00% (10/10) | Non-Toxin | 1.59 | 1.39 | -1.83 | 2.98 | 1.15 | 67.8 |
| 306 | polymerase | IMLNEAMNY | HLA-A*30:02 | 9 | -0.02776 | 100.00% (10/10) | Non-Toxin | 1.48 | 1.34 | -1.81 | 2.83 | 1.01 | 64.9 |
| 307 | polymerase | IMLNEAMNY | HLA-B*15:01 | 9 | -0.02776 | 100.00% (10/10) | Non-Toxin | 1.48 | 1.34 | -1.81 | 2.83 | 1.01 | 65.3 |
| 308 | polymerase | IMLNEAMNY | HLA-A*03:01 | 9 | -0.02776 | 100.00% (10/10) | Non-Toxin | 1.48 | 1.34 | -2.15 | 2.83 | 0.68 | 141.1 |
| 309 | polymerase | IMLNEAMNYF | HLA-B*15:01 | 10 | -0.04743 | 100.00% (10/10) | Non-Toxin | 1.38 | 1.17 | -1.38 | 2.56 | 1.18 | 23.9 |
| 310 | polymerase | IPAPIGGFNY | HLA-B*35:01 | 10 | 0.29106 | 100.00% (10/10) | Non-Toxin | 1.26 | 1.18 | -1.49 | 2.43 | 0.94 | 30.9 |
| 311 | polymerase | IPFLFLSAY | HLA-B*35:01 | 9 | 0.01364 | 100.00% (10/10) | Non-Toxin | 1.39 | 1.2 | -0.67 | 2.59 | 1.92 | 4.7 |
| 312 | polymerase | IPFLFLSAY | HLA-B*53:01 | 9 | 0.01364 | 100.00% (10/10) | Non-Toxin | 1.39 | 1.2 | -2.07 | 2.59 | 0.52 | 118.6 |
| 313 | polymerase | IQFDCFMEL | HLA-A*02:06 | 9 | 0.02832 | 100.00% (10/10) | Non-Toxin | 1.55 | 0.49 | -0.81 | 2.05 | 1.23 | 6.5 |
| 314 | polymerase | IQFDCFMEL | HLA-A*02:01 | 9 | 0.02832 | 100.00% (10/10) | Non-Toxin | 1.55 | 0.49 | -1.27 | 2.05 | 0.77 | 18.8 |
| 315 | polymerase | IQQLLISTEF | HLA-B*15:01 | 10 | 0.00557 | 100.00% (10/10) | Non-Toxin | 1.56 | 1.15 | -1.73 | 2.71 | 0.98 | 53.3 |
| 316 | polymerase | IRIATVYTWAY | HLA-A*30:02 | 11 | 0.40331 | 100.00% (10/10) | Non-Toxin | 1.56 | 1.47 | -2.43 | 3.03 | 0.6 | 267.3 |
| 317 | polymerase | ISRLFNMYR | HLA-A*31:01 | 9 | -0.03681 | 100.00% (10/10) | Non-Toxin | 0.98 | 0.63 | -0.69 | 1.61 | 0.92 | 4.9 |
| 318 | polymerase | IVDPELFALY | HLA-A*01:01 | 10 | 0.21948 | 100.00% (10/10) | Non-Toxin | 1.03 | 1.28 | -1.34 | 2.32 | 0.98 | 21.8 |
| 319 | polymerase | IYGLPGFFNW | HLA-A*23:01 | 10 | 0.22726 | 100.00% (10/10) | Non-Toxin | 1.52 | 0.36 | -0.94 | 1.89 | 0.94 | 8.8 |
| 320 | polymerase | IYGLPGFFNW | HLA-A*24:02 | 10 | 0.22726 | 100.00% (10/10) | Non-Toxin | 1.52 | 0.36 | -1.25 | 1.89 | 0.64 | 17.6 |
| 321 | polymerase | IYLMNWCDF | HLA-A*23:01 | 9 | -0.01063 | 100.00% (10/10) | Non-Toxin | 0.96 | 1.25 | -1.09 | 2.21 | 1.12 | 12.3 |

| | | | | | | | | | | | | | |
|---|---|---|---|---|---|---|---|---|---|---|---|---|---|
| 322 | polymerase | IYLMNWCDF | HLA-A*24:02 | 9 | -0.01063 | 100.00% (10/10) | Non-Toxin | 0.96 | 1.25 | -1.24 | 2.21 | 0.97 | 17.5 |
| 323 | polymerase | KALSPIKDEW | HLA-B*58:01 | 10 | -0.17705 | 100.00% (10/10) | Non-Toxin | 1.25 | 0.4 | -0.89 | 1.65 | 0.76 | 7.8 |
| 324 | polymerase | KEICAKQAQLY | HLA-B*44:03 | 11 | -0.36316 | 100.00% (10/10) | Non-Toxin | 1.18 | 1.33 | -2 | 2.51 | 0.51 | 100.5 |
| 325 | polymerase | KIWWKIIGY | HLA-A*32:01 | 9 | 0.34219 | 100.00% (10/10) | Non-Toxin | 1.24 | 1.47 | -2.12 | 2.7 | 0.58 | 131.2 |
| 326 | polymerase | KKYYQIDQPF | HLA-A*23:01 | 10 | -0.07148 | 100.00% (10/10) | Non-Toxin | 0.86 | 1.27 | -1.27 | 2.12 | 0.85 | 18.7 |
| 327 | polymerase | KLFPSEYSI | HLA-A*32:01 | 9 | -0.13979 | 100.00% (10/10) | Non-Toxin | 1.34 | 0.36 | -0.9 | 1.7 | 0.79 | 8 |
| 328 | polymerase | KLFPSEYSI | HLA-A*02:01 | 9 | -0.13979 | 100.00% (10/10) | Non-Toxin | 1.34 | 0.36 | -1.16 | 1.7 | 0.54 | 14.5 |
| 329 | polymerase | KLKGLVVPL | HLA-A*02:03 | 9 | 0.02052 | 100.00% (10/10) | Non-Toxin | 1.33 | 0.46 | -1.2 | 1.79 | 0.59 | 15.9 |
| 330 | polymerase | KLKGLVVPLF | HLA-B*15:01 | 10 | 0.01652 | 100.00% (10/10) | Non-Toxin | 1.37 | 1.14 | -1.65 | 2.51 | 0.87 | 44.6 |
| 331 | polymerase | KLKGLVVPLF | HLA-A*32:01 | 10 | 0.01652 | 100.00% (10/10) | Non-Toxin | 1.37 | 1.14 | -2.01 | 2.51 | 0.5 | 103 |
| 332 | polymerase | KQFHDDLKKY | HLA-A*30:02 | 10 | -0.20469 | 100.00% (10/10) | Non-Toxin | 1.58 | 1.42 | -2.4 | 3 | 0.61 | 248.6 |
| 333 | polymerase | KQFHDDLKKY | HLA-B*15:01 | 10 | -0.20469 | 100.00% (10/10) | Non-Toxin | 1.58 | 1.42 | -2.42 | 3 | 0.59 | 260.2 |
| 334 | polymerase | KSFLDYHTEF | HLA-B*58:01 | 10 | 0.16655 | 100.00% (10/10) | Non-Toxin | 1.46 | 1.24 | -1.51 | 2.69 | 1.19 | 32.2 |
| 335 | polymerase | KSFLDYHTEF | HLA-B*57:01 | 10 | 0.16655 | 100.00% (10/10) | Non-Toxin | 1.46 | 1.24 | -1.71 | 2.69 | 0.99 | 50.9 |
| 336 | polymerase | KSFLDYHTEF | HLA-A*32:01 | 10 | 0.16655 | 100.00% (10/10) | Non-Toxin | 1.46 | 1.24 | -1.92 | 2.69 | 0.77 | 83.2 |
| 337 | polymerase | KSFLDYHTEF | HLA-A*23:01 | 10 | 0.16655 | 100.00% (10/10) | Non-Toxin | 1.46 | 1.24 | -2.01 | 2.69 | 0.68 | 102.4 |
| 338 | polymerase | KSMSRCCFW | HLA-B*58:01 | 9 | -0.20092 | 90.00% (9/10) | Non-Toxin | 1.46 | 0.5 | -0.69 | 1.96 | 1.27 | 4.9 |
| 339 | polymerase | KSMSRCCFW | HLA-B*57:01 | 9 | -0.20092 | 90.00% (9/10) | Non-Toxin | 1.46 | 0.5 | -0.95 | 1.96 | 1 | 9 |
| 340 | polymerase | KSRELDFPLW | HLA-B*58:01 | 10 | 0.22271 | 100.00% (10/10) | Non-Toxin | 1.56 | 0.49 | -0.74 | 2.05 | 1.31 | 5.5 |
| 341 | polymerase | KSRELDFPLW | HLA-B*57:01 | 10 | 0.22271 | 100.00% (10/10) | Non-Toxin | 1.56 | 0.49 | -0.83 | 2.05 | 1.22 | 6.7 |
| 342 | polymerase | KTIKNITAR | HLA-A*31:01 | 9 | 0.0008 | 100.00% (10/10) | Non-Toxin | 1.13 | 0.72 | -1.13 | 1.85 | 0.72 | 13.4 |
| 343 | polymerase | KTWTIATIPF | HLA-A*32:01 | 10 | 0.42104 | 100.00% (10/10) | Non-Toxin | 0.97 | 1.19 | -1.03 | 2.16 | 1.13 | 10.7 |
| 344 | polymerase | KTWTIATIPF | HLA-B*58:01 | 10 | 0.42104 | 100.00% (10/10) | Non-Toxin | 0.97 | 1.19 | -1.45 | 2.16 | 0.71 | 28.2 |
| 345 | polymerase | KVIVYSLIKF | HLA-A*32:01 | 10 | -0.10408 | 100.00% (10/10) | Non-Toxin | 1.56 | 1.28 | -2.22 | 2.84 | 0.62 | 164.9 |
| 346 | polymerase | KVIVYSLIKF | HLA-B*15:01 | 10 | -0.10408 | 100.00% (10/10) | Non-Toxin | 1.56 | 1.28 | -2.26 | 2.84 | 0.58 | 181.5 |
| 347 | polymerase | KWYECFLF | HLA-A*23:01 | 8 | 0.14789 | 100.00% (10/10) | Non-Toxin | 1.15 | 1.36 | -1.12 | 2.51 | 1.38 | 13.3 |
| 348 | polymerase | KWYECFLF | HLA-A*24:02 | 8 | 0.14789 | 100.00% (10/10) | Non-Toxin | 1.15 | 1.36 | -1.55 | 2.51 | 0.96 | 35.1 |
| 349 | polymerase | KWYECFLFW | HLA-A*23:01 | 9 | 0.21629 | 100.00% (10/10) | Non-Toxin | 1.52 | 0.64 | -1 | 2.16 | 1.16 | 10.1 |
| 350 | polymerase | KWYECFLFW | HLA-A*24:02 | 9 | 0.21629 | 100.00% (10/10) | Non-Toxin | 1.52 | 0.64 | -1.5 | 2.16 | 0.66 | 31.6 |
| 351 | polymerase | KWYECFLFWF | HLA-A*23:01 | 10 | 0.37883 | 100.00% (10/10) | Non-Toxin | 1.24 | 1.36 | -1 | 2.6 | 1.6 | 10.1 |
| 352 | polymerase | KWYECFLFWF | HLA-A*24:02 | 10 | 0.37883 | 100.00% (10/10) | Non-Toxin | 1.24 | 1.36 | -1.5 | 2.6 | 1.1 | 31.4 |
| 353 | polymerase | KYYQIDQPF | HLA-A*23:01 | 9 | -0.07152 | 100.00% (10/10) | Non-Toxin | 0.86 | 1.39 | -0.95 | 2.24 | 1.29 | 9 |
| 354 | polymerase | KYYQIDQPF | HLA-A*24:02 | 9 | -0.07152 | 100.00% (10/10) | Non-Toxin | 0.86 | 1.39 | -1.24 | 2.24 | 1 | 17.3 |
| 355 | polymerase | KYYQIDQPFF | HLA-A*23:01 | 10 | -0.01656 | 100.00% (10/10) | Non-Toxin | 1.33 | 1.39 | -0.98 | 2.72 | 1.73 | 9.6 |
| 356 | polymerase | KYYQIDQPFF | HLA-A*24:02 | 10 | -0.01656 | 100.00% (10/10) | Non-Toxin | 1.33 | 1.39 | -1.3 | 2.72 | 1.42 | 19.8 |
| 357 | polymerase | LAEFFSFFR | HLA-A*68:01 | 9 | 0.27577 | 100.00% (10/10) | Non-Toxin | 1.02 | 0.63 | -0.89 | 1.64 | 0.75 | 7.8 |
| 358 | polymerase | LARALRSHMW | HLA-B*58:01 | 10 | -0.13526 | 100.00% (10/10) | Non-Toxin | 1.6 | 0.37 | -1.14 | 1.96 | 0.82 | 13.9 |
| 359 | polymerase | LARALRSHMW | HLA-B*57:01 | 10 | -0.13526 | 100.00% (10/10) | Non-Toxin | 1.6 | 0.37 | -1.31 | 1.96 | 0.65 | 20.5 |
| 360 | polymerase | LETDDYNGIY | HLA-A*01:01 | 10 | 0.15319 | 100.00% (10/10) | Non-Toxin | 1.41 | 1.25 | -0.93 | 2.66 | 1.73 | 8.6 |
| 361 | polymerase | LFPVMGNRIY | HLA-B*35:01 | 10 | 0.01529 | 100.00% (10/10) | Non-Toxin | 1.57 | 1.28 | -1.68 | 2.85 | 1.16 | 48.4 |
| 362 | polymerase | LIASGVGKY | HLA-A*30:02 | 9 | -0.17931 | 100.00% (10/10) | Non-Toxin | 1.13 | 1.4 | -1.68 | 2.53 | 0.85 | 47.7 |
| 363 | polymerase | LIASGVGKY | HLA-B*15:01 | 9 | -0.17931 | 100.00% (10/10) | Non-Toxin | 1.13 | 1.4 | -1.8 | 2.53 | 0.73 | 63 |
| 364 | polymerase | LIASGVGKYF | HLA-B*15:01 | 10 | -0.23283 | 100.00% (10/10) | Non-Toxin | 1.41 | 1.23 | -1.63 | 2.64 | 1.01 | 42.4 |
| 365 | polymerase | LISRSNALW | HLA-B*58:01 | 9 | -0.14227 | 100.00% (10/10) | Non-Toxin | 1.35 | 0.38 | -1.06 | 1.73 | 0.67 | 11.5 |
| 366 | polymerase | LKSMSRCCFW | HLA-B*58:01 | 10 | -0.36895 | 90.00% (9/10) | Non-Toxin | 1.46 | 0.47 | -1.14 | 1.93 | 0.8 | 13.7 |
| 367 | polymerase | LLAEFFSFF | HLA-B*15:01 | 9 | 0.26643 | 100.00% (10/10) | Non-Toxin | 1.37 | 1.11 | -1.2 | 2.47 | 1.27 | 16 |
| 368 | polymerase | LLAEFFSFFR | HLA-A*68:01 | 10 | 0.35292 | 100.00% (10/10) | Non-Toxin | 1.02 | 0.65 | -0.99 | 1.67 | 0.68 | 9.7 |
| 369 | polymerase | LLLPTQGPY | HLA-B*15:01 | 9 | -0.06388 | 100.00% (10/10) | Non-Toxin | 0.86 | 1.28 | -1.47 | 2.13 | 0.67 | 29.2 |

| | | | | | | | | | | | | | |
|---|---|---|---|---|---|---|---|---|---|---|---|---|---|
| 370 | polymerase | LLVTSYMIY | HLA-B*15:01 | 9 | -0.18256 | 100.00% (10/10) | Non-Toxin | 1.33 | 1.38 | -1.32 | 2.71 | 1.38 | 21.1 |
| 371 | polymerase | LLVTSYMIYL | HLA-A*02:01 | 10 | -0.16738 | 100.00% (10/10) | Non-Toxin | 1.53 | 0.52 | -1.48 | 2.05 | 0.57 | 30.2 |
| 372 | polymerase | LPRAAHEIL | HLA-B*07:02 | 9 | 0.28698 | 100.00% (10/10) | Non-Toxin | 1.42 | 0.31 | -1.02 | 1.74 | 0.72 | 10.4 |
| 373 | polymerase | LPSGAQRLF | HLA-B*35:01 | 9 | -0.05334 | 100.00% (10/10) | Non-Toxin | 1.2 | 0.95 | -1.5 | 2.15 | 0.64 | 31.9 |
| 374 | polymerase | LSHPRVFKR | HLA-A*31:01 | 9 | 0.0614 | 100.00% (10/10) | Non-Toxin | 1.07 | 0.74 | -1.11 | 1.81 | 0.7 | 12.8 |
| 375 | polymerase | LSNREVKIWW | HLA-B*58:01 | 10 | 0.22642 | 100.00% (10/10) | Non-Toxin | 1.48 | 0.3 | -1.24 | 1.78 | 0.54 | 17.5 |
| 376 | polymerase | LSQNLLVTSY | HLA-B*15:01 | 10 | -0.09075 | 100.00% (10/10) | Non-Toxin | 1.6 | 1.32 | -1.52 | 2.91 | 1.4 | 32.8 |
| 377 | polymerase | LSQNLLVTSY | HLA-A*30:02 | 10 | -0.09075 | 100.00% (10/10) | Non-Toxin | 1.6 | 1.32 | -2.07 | 2.91 | 0.85 | 116.6 |
| 378 | polymerase | LTIDDCVKNW | HLA-B*58:01 | 10 | -0.1123 | 100.00% (10/10) | Non-Toxin | 1.5 | 0.43 | -1.42 | 1.93 | 0.51 | 26.1 |
| 379 | polymerase | LTPEMVLMY | HLA-A*01:01 | 9 | -0.14695 | 100.00% (10/10) | Non-Toxin | 1.52 | 1.23 | -1.97 | 2.75 | 0.78 | 93.8 |
| 380 | polymerase | LVLVCFPVY | HLA-B*35:01 | 9 | 0.1104 | 100.00% (10/10) | Non-Toxin | 1.54 | 1.33 | -2.25 | 2.86 | 0.61 | 176.8 |
| 381 | polymerase | LVSNTSKHTY | HLA-B*15:01 | 10 | -0.33653 | 100.00% (10/10) | Non-Toxin | 1.52 | 1.27 | -2.05 | 2.79 | 0.74 | 113.5 |
| 382 | polymerase | LVSNTSKHTY | HLA-A*30:02 | 10 | -0.33653 | 100.00% (10/10) | Non-Toxin | 1.52 | 1.27 | -2.11 | 2.79 | 0.68 | 129.8 |
| 383 | polymerase | LWGLIDPLF | HLA-A*23:01 | 9 | 0.13448 | 100.00% (10/10) | Non-Toxin | 1.16 | 1.2 | -1.35 | 2.37 | 1.02 | 22.4 |
| 384 | polymerase | LWGLIDPLF | HLA-A*24:02 | 9 | 0.13448 | 100.00% (10/10) | Non-Toxin | 1.16 | 1.2 | -1.59 | 2.37 | 0.77 | 39.2 |
| 385 | polymerase | MIYLMNWCDF | HLA-A*23:01 | 10 | -0.01369 | 100.00% (10/10) | Non-Toxin | 0.96 | 1.23 | -1.46 | 2.2 | 0.74 | 28.8 |
| 386 | polymerase | MIYLMNWCDF | HLA-A*24:02 | 10 | -0.01369 | 100.00% (10/10) | Non-Toxin | 0.96 | 1.23 | -1.66 | 2.2 | 0.54 | 45.6 |
| 387 | polymerase | MLADKVLEY | HLA-B*15:01 | 9 | -0.08698 | 100.00% (10/10) | Non-Toxin | 1.56 | 1.3 | -1.39 | 2.86 | 1.47 | 24.3 |
| 388 | polymerase | MLADKVLEY | HLA-B*35:01 | 9 | -0.08698 | 100.00% (10/10) | Non-Toxin | 1.56 | 1.3 | -1.45 | 2.86 | 1.4 | 28.4 |
| 389 | polymerase | MLADKVLEY | HLA-A*03:01 | 9 | -0.08698 | 100.00% (10/10) | Non-Toxin | 1.56 | 1.3 | -2.19 | 2.86 | 0.67 | 153.6 |
| 390 | polymerase | MLADKVLEY | HLA-A*30:02 | 9 | -0.08698 | 100.00% (10/10) | Non-Toxin | 1.56 | 1.3 | -2.26 | 2.86 | 0.6 | 180 |
| 391 | polymerase | MLADKVLEY | HLA-A*01:01 | 9 | -0.08698 | 100.00% (10/10) | Non-Toxin | 1.56 | 1.3 | -2.33 | 2.86 | 0.52 | 215.1 |
| 392 | polymerase | MLEYVLSGAY | HLA-B*15:01 | 10 | -0.04609 | 100.00% (10/10) | Non-Toxin | 1.04 | 1.3 | -1.37 | 2.33 | 0.97 | 23.3 |
| 393 | polymerase | MLEYVLSGAY | HLA-A*30:02 | 10 | -0.04609 | 100.00% (10/10) | Non-Toxin | 1.04 | 1.3 | -1.8 | 2.33 | 0.53 | 63.8 |
| 394 | polymerase | MLNEAMNYF | HLA-B*15:01 | 9 | -0.03617 | 100.00% (10/10) | Non-Toxin | 1.38 | 1.09 | -1.05 | 2.47 | 1.42 | 11.2 |
| 395 | polymerase | MLNEAMNYF | HLA-B*35:01 | 9 | -0.03617 | 100.00% (10/10) | Non-Toxin | 1.38 | 1.09 | -1.77 | 2.47 | 0.7 | 58.9 |
| 396 | polymerase | MMLLYQSTL | HLA-A*02:01 | 9 | -0.24434 | 100.00% (10/10) | Non-Toxin | 1.51 | 0.52 | -1.34 | 2.03 | 0.69 | 22 |
| 397 | polymerase | MMLLYQSTL | HLA-A*02:03 | 9 | -0.24434 | 100.00% (10/10) | Non-Toxin | 1.51 | 0.52 | -1.53 | 2.03 | 0.51 | 33.5 |
| 398 | polymerase | MYCDVLEGR | HLA-A*33:01 | 9 | 0.13888 | 100.00% (10/10) | Non-Toxin | 1.17 | 0.8 | -1.45 | 1.97 | 0.52 | 28.4 |
| 399 | polymerase | MYGWFFVPR | HLA-A*33:01 | 9 | 0.48645 | 100.00% (10/10) | Non-Toxin | 0.67 | 0.71 | -0.83 | 1.38 | 0.55 | 6.8 |
| 400 | polymerase | NIMKKSFKAY | HLA-B*15:01 | 10 | -0.69404 | 100.00% (10/10) | Non-Toxin | 1.59 | 1.31 | -2.05 | 2.9 | 0.86 | 111.2 |
| 401 | polymerase | NIRRKVLIL | HLA-B*08:01 | 9 | -0.03386 | 100.00% (10/10) | Non-Toxin | 1.44 | 0.57 | -1.19 | 2.01 | 0.82 | 15.5 |
| 402 | polymerase | NLHGKRKSL | HLA-B*08:01 | 9 | -0.39534 | 100.00% (10/10) | Non-Toxin | 1.84 | 0.48 | -1.7 | 2.31 | 0.61 | 50.3 |
| 403 | polymerase | NTCSVDLAR | HLA-A*68:01 | 9 | -0.10939 | 100.00% (10/10) | Non-Toxin | 1.14 | 0.68 | -0.96 | 1.82 | 0.86 | 9.2 |
| 404 | polymerase | NTEAAVLSRY | HLA-A*26:01 | 10 | 0.03035 | 100.00% (10/10) | Non-Toxin | 1.46 | 1.15 | -1.77 | 2.61 | 0.84 | 58.9 |
| 405 | polymerase | NTEAAVLSRY | HLA-A*01:01 | 10 | 0.03035 | 100.00% (10/10) | Non-Toxin | 1.46 | 1.15 | -1.91 | 2.61 | 0.7 | 80.7 |
| 406 | polymerase | NTMYGWFFV | HLA-A*68:02 | 9 | 0.34799 | 100.00% (10/10) | Non-Toxin | 0.84 | 0.16 | -0.41 | 0.99 | 0.58 | 2.6 |
| 407 | polymerase | NTSDLDFVIFY | HLA-A*01:01 | 11 | 0.313 | 100.00% (10/10) | Non-Toxin | 1.68 | 1.25 | -1.76 | 2.93 | 1.17 | 57.2 |
| 408 | polymerase | NTSKSFLDY | HLA-A*01:01 | 9 | -0.318 | 100.00% (10/10) | Non-Toxin | 1.17 | 1.3 | -1.73 | 2.47 | 0.74 | 54.1 |
| 409 | polymerase | NYLNLSRIF | HLA-A*23:01 | 9 | -0.0552 | 100.00% (10/10) | Non-Toxin | 1.29 | 1.19 | -1.74 | 2.48 | 0.74 | 55.1 |
| 410 | polymerase | QQLLISTEF | HLA-B*15:01 | 9 | 0.05037 | 100.00% (10/10) | Non-Toxin | 1.56 | 1.15 | -1.95 | 2.71 | 0.76 | 89.6 |
| 411 | polymerase | QSISIKWAF | HLA-B*58:01 | 9 | 0.01313 | 100.00% (10/10) | Non-Toxin | 1.4 | 1.14 | -1.51 | 2.54 | 1.03 | 32.4 |
| 412 | polymerase | QSISIKWAF | HLA-A*32:01 | 9 | 0.01313 | 100.00% (10/10) | Non-Toxin | 1.4 | 1.14 | -1.9 | 2.54 | 0.64 | 79.3 |
| 413 | polymerase | RALRSHMW | HLA-B*58:01 | 8 | -0.23037 | 100.00% (10/10) | Non-Toxin | 1.6 | 0.59 | -1.63 | 2.18 | 0.55 | 42.4 |
| 414 | polymerase | RALRSHMWR | HLA-A*31:01 | 9 | -0.10095 | 100.00% (10/10) | Non-Toxin | 0.93 | 0.85 | -0.83 | 1.78 | 0.95 | 6.8 |
| 415 | polymerase | REHMLADKVL | HLA-B*40:01 | 10 | -0.2759 | 100.00% (10/10) | Non-Toxin | 1.74 | 0.5 | -1.39 | 2.23 | 0.84 | 24.6 |
| 416 | polymerase | RELALGRVIY | HLA-B*44:03 | 10 | 0.21929 | 100.00% (10/10) | Non-Toxin | 1.61 | 1.3 | -1.8 | 2.91 | 1.1 | 63.4 |
| 417 | polymerase | RELALGRVIY | HLA-B*44:02 | 10 | 0.21929 | 100.00% (10/10) | Non-Toxin | 1.61 | 1.3 | -2.06 | 2.91 | 0.84 | 115.3 |

| | | | | | | | | | | | | | |
|---|---|---|---|---|---|---|---|---|---|---|---|---|---|
| 418 | polymerase | RELALGRVIY | HLA-B*15:01 | 10 | 0.21929 | 100.00% (10/10) | Non-Toxin | 1.61 | 1.3 | -2.13 | 2.91 | 0.78 | 133.8 |
| 419 | polymerase | RELKLFPSEY | HLA-B*44:03 | 10 | -0.20896 | 100.00% (10/10) | Non-Toxin | 1.48 | 1.3 | -1.66 | 2.78 | 1.13 | 45.3 |
| 420 | polymerase | RELKLFPSEY | HLA-B*44:02 | 10 | -0.20896 | 100.00% (10/10) | Non-Toxin | 1.48 | 1.3 | -1.88 | 2.78 | 0.91 | 75.5 |
| 421 | polymerase | RELKLFPSEY | HLA-A*30:02 | 10 | -0.20896 | 100.00% (10/10) | Non-Toxin | 1.48 | 1.3 | -2.14 | 2.78 | 0.65 | 136.5 |
| 422 | polymerase | RELKLFPSEY | HLA-B*15:01 | 10 | -0.20896 | 100.00% (10/10) | Non-Toxin | 1.48 | 1.3 | -2.16 | 2.78 | 0.63 | 143.4 |
| 423 | polymerase | RIATVYTW | HLA-B*58:01 | 8 | 0.12124 | 100.00% (10/10) | Non-Toxin | 1.73 | 0.56 | -1.7 | 2.29 | 0.59 | 50.4 |
| 424 | polymerase | RIATVYTWAY | HLA-A*30:02 | 10 | 0.3347 | 100.00% (10/10) | Non-Toxin | 1.56 | 1.44 | -1.54 | 3 | 1.46 | 35 |
| 425 | polymerase | RIATVYTWAY | HLA-B*15:01 | 10 | 0.3347 | 100.00% (10/10) | Non-Toxin | 1.56 | 1.44 | -1.69 | 3 | 1.31 | 49.3 |
| 426 | polymerase | RIATVYTWAY | HLA-B*58:01 | 10 | 0.3347 | 100.00% (10/10) | Non-Toxin | 1.56 | 1.44 | -2.19 | 3 | 0.81 | 156.3 |
| 427 | polymerase | RIATVYTWAY | HLA-A*32:01 | 10 | 0.3347 | 100.00% (10/10) | Non-Toxin | 1.56 | 1.44 | -2.31 | 3 | 0.69 | 203.2 |
| 428 | polymerase | RIATVYTWAY | HLA-A*03:01 | 10 | 0.3347 | 100.00% (10/10) | Non-Toxin | 1.56 | 1.44 | -2.49 | 3 | 0.51 | 309.1 |
| 429 | polymerase | RIGLNSSSCY | HLA-A*30:02 | 10 | -0.49441 | 100.00% (10/10) | Non-Toxin | 1.41 | 1.42 | -1.8 | 2.83 | 1.03 | 63.7 |
| 430 | polymerase | RLETDDYNGIY | HLA-A*01:01 | 11 | 0.21143 | 100.00% (10/10) | Non-Toxin | 1.41 | 1.34 | -1.76 | 2.75 | 0.99 | 57.9 |
| 431 | polymerase | RLFAKMTY | HLA-A*03:01 | 8 | -0.26517 | 100.00% (10/10) | Non-Toxin | 1.39 | 1.42 | -2.31 | 2.81 | 0.5 | 205 |
| 432 | polymerase | RLFAKMTYK | HLA-A*03:01 | 9 | -0.26733 | 100.00% (10/10) | Non-Toxin | 0.76 | 0.35 | -0.53 | 1.11 | 0.58 | 3.4 |
| 433 | polymerase | RLFAKMTYKMR | HLA-A*31:01 | 11 | -0.55391 | 100.00% (10/10) | Non-Toxin | 1.25 | 0.79 | -1.16 | 2.04 | 0.88 | 14.6 |
| 434 | polymerase | RLFNMYRSY | HLA-A*32:01 | 9 | -0.19597 | 100.00% (10/10) | Non-Toxin | 1.42 | 1.46 | -1.11 | 2.88 | 1.77 | 12.8 |
| 435 | polymerase | RLFNMYRSY | HLA-B*15:01 | 9 | -0.19597 | 100.00% (10/10) | Non-Toxin | 1.42 | 1.46 | -1.18 | 2.88 | 1.69 | 15.3 |
| 436 | polymerase | RLFNMYRSY | HLA-A*30:02 | 9 | -0.19597 | 100.00% (10/10) | Non-Toxin | 1.42 | 1.46 | -1.2 | 2.88 | 1.68 | 15.8 |
| 437 | polymerase | RLFNMYRSY | HLA-A*03:01 | 9 | -0.19597 | 100.00% (10/10) | Non-Toxin | 1.42 | 1.46 | -1.52 | 2.88 | 1.36 | 33.1 |
| 438 | polymerase | RLFNMYRSY | HLA-A*30:01 | 9 | -0.19597 | 100.00% (10/10) | Non-Toxin | 1.42 | 1.46 | -2.19 | 2.88 | 0.69 | 155 |
| 439 | polymerase | RLFNMYRSYF | HLA-A*32:01 | 10 | -0.23617 | 100.00% (10/10) | Non-Toxin | 1.16 | 1.29 | -1.2 | 2.45 | 1.25 | 15.7 |
| 440 | polymerase | RLFNMYRSYF | HLA-B*15:01 | 10 | -0.23617 | 100.00% (10/10) | Non-Toxin | 1.16 | 1.29 | -1.34 | 2.45 | 1.1 | 22 |
| 441 | polymerase | RLRDKSTQF | HLA-B*15:01 | 9 | -0.36153 | 100.00% (10/10) | Non-Toxin | 1.43 | 1.23 | -1.6 | 2.65 | 1.06 | 39.5 |
| 442 | polymerase | RLRLETDDY | HLA-A*30:02 | 9 | 0.17136 | 100.00% (10/10) | Non-Toxin | 1.19 | 1.36 | -1.98 | 2.55 | 0.57 | 96.4 |
| 443 | polymerase | RLRMNLRAL | HLA-B*07:02 | 9 | -0.1101 | 100.00% (10/10) | Non-Toxin | 1.6 | 0.48 | -1.32 | 2.08 | 0.76 | 20.9 |
| 444 | polymerase | RLSHHDYNQF | HLA-B*15:01 | 10 | -0.04467 | 100.00% (10/10) | Non-Toxin | 1.48 | 1.24 | -2.2 | 2.71 | 0.51 | 159.5 |
| 445 | polymerase | RPRGRHTM | HLA-B*07:02 | 8 | 0.16451 | 100.00% (10/10) | Non-Toxin | 1.31 | 0.18 | -0.89 | 1.5 | 0.61 | 7.8 |
| 446 | polymerase | RPRGRHTMV | HLA-B*07:02 | 9 | 0.06191 | 100.00% (10/10) | Non-Toxin | 1.1 | 0.19 | -0.64 | 1.29 | 0.64 | 4.4 |
| 447 | polymerase | RPTKALRSAI | HLA-B*07:02 | 10 | -0.24514 | 100.00% (10/10) | Non-Toxin | 1.24 | 0.23 | -0.68 | 1.48 | 0.8 | 4.8 |
| 448 | polymerase | RSAIRIATVY | HLA-B*58:01 | 10 | 0.42033 | 100.00% (10/10) | Non-Toxin | 1.43 | 1.42 | -1.36 | 2.85 | 1.49 | 22.9 |
| 449 | polymerase | RSAIRIATVY | HLA-A*30:02 | 10 | 0.42033 | 100.00% (10/10) | Non-Toxin | 1.43 | 1.42 | -1.5 | 2.85 | 1.36 | 31.4 |
| 450 | polymerase | RSAIRIATVY | HLA-B*15:01 | 10 | 0.42033 | 100.00% (10/10) | Non-Toxin | 1.43 | 1.42 | -1.77 | 2.85 | 1.09 | 58.3 |
| 451 | polymerase | RSAIRIATVY | HLA-B*57:01 | 10 | 0.42033 | 100.00% (10/10) | Non-Toxin | 1.43 | 1.42 | -1.9 | 2.85 | 0.95 | 80.2 |
| 452 | polymerase | RSMFIDDLL | HLA-B*58:01 | 9 | 0.22352 | 100.00% (10/10) | Non-Toxin | 1.55 | 0.57 | -1.16 | 2.12 | 0.96 | 14.5 |
| 453 | polymerase | RSPSHHLEPY | HLA-A*30:02 | 10 | -0.03949 | 100.00% (10/10) | Non-Toxin | 0.99 | 1.45 | -1.73 | 2.44 | 0.7 | 54.3 |
| 454 | polymerase | RSWNTSDLDF | HLA-B*58:01 | 10 | -0.03343 | 100.00% (10/10) | Non-Toxin | 1.21 | 1.32 | -1.41 | 2.53 | 1.12 | 26 |
| 455 | polymerase | RSWNTSDLDF | HLA-B*57:01 | 10 | -0.03343 | 100.00% (10/10) | Non-Toxin | 1.21 | 1.32 | -1.91 | 2.53 | 0.63 | 80.5 |
| 456 | polymerase | RSYFGLVLVCF | HLA-A*23:01 | 11 | 0.1719 | 100.00% (10/10) | Non-Toxin | 1.48 | 1.28 | -2.23 | 2.76 | 0.53 | 168.4 |
| 457 | polymerase | RYESMAIFAER | HLA-A*31:01 | 11 | 0.06445 | 100.00% (10/10) | Non-Toxin | 1 | 0.78 | -1.21 | 1.78 | 0.56 | 16.4 |
| 458 | polymerase | RYKCSNTSK | HLA-A*30:01 | 9 | -0.35534 | 100.00% (10/10) | Non-Toxin | 0.98 | 0.48 | -0.86 | 1.46 | 0.6 | 7.2 |
| 459 | polymerase | RYLPSGAQRLF | HLA-A*23:01 | 11 | -0.1766 | 100.00% (10/10) | Non-Toxin | 1.2 | 1.31 | -1.64 | 2.51 | 0.86 | 44.1 |
| 460 | polymerase | RYLPSGAQRLF | HLA-A*24:02 | 11 | -0.1766 | 100.00% (10/10) | Non-Toxin | 1.2 | 1.31 | -1.9 | 2.51 | 0.61 | 78.6 |
| 461 | polymerase | SAIEYAQLR | HLA-A*68:01 | 9 | 0.07294 | 100.00% (10/10) | Non-Toxin | 0.94 | 0.72 | -1.1 | 1.66 | 0.57 | 12.5 |
| 462 | polymerase | SAIRIATVY | HLA-B*35:01 | 9 | 0.31859 | 100.00% (10/10) | Non-Toxin | 1.43 | 1.4 | -1.41 | 2.84 | 1.42 | 25.8 |
| 463 | polymerase | SAIRIATVY | HLA-B*15:01 | 9 | 0.31859 | 100.00% (10/10) | Non-Toxin | 1.43 | 1.4 | -1.93 | 2.84 | 0.9 | 85.3 |
| 464 | polymerase | SAIRIATVY | HLA-A*30:02 | 9 | 0.31859 | 100.00% (10/10) | Non-Toxin | 1.43 | 1.4 | -2.2 | 2.84 | 0.63 | 158.7 |
| 465 | polymerase | SAIRIATVYTW | HLA-B*58:01 | 11 | 0.3592 | 100.00% (10/10) | Non-Toxin | 1.73 | 0.52 | -1.06 | 2.25 | 1.19 | 11.5 |

| | | | | | | | | | | | | | |
|---|---|---|---|---|---|---|---|---|---|---|---|---|---|
| 466 | polymerase | SAIRIATVYTW | HLA-B*57:01 | 11 | 0.3592 | 100.00% (10/10) | Non-Toxin | 1.73 | 0.52 | -1.54 | 2.25 | 0.71 | 34.7 |
| 467 | polymerase | SFCGIQFDCF | HLA-A*23:01 | 10 | 0.13082 | 100.00% (10/10) | Non-Toxin | 1.49 | 1.13 | -2.02 | 2.61 | 0.59 | 105.7 |
| 468 | polymerase | SFLDYHTEF | HLA-A*23:01 | 9 | 0.13683 | 100.00% (10/10) | Non-Toxin | 1.46 | 1.25 | -1.55 | 2.71 | 1.16 | 35.2 |
| 469 | polymerase | SFLDYHTEF | HLA-A*24:02 | 9 | 0.13683 | 100.00% (10/10) | Non-Toxin | 1.46 | 1.25 | -1.98 | 2.71 | 0.72 | 96.2 |
| 470 | polymerase | SMFIDDLLSI | HLA-A*02:03 | 10 | 0.09866 | 100.00% (10/10) | Non-Toxin | 1.16 | 0.41 | -0.81 | 1.58 | 0.76 | 6.5 |
| 471 | polymerase | SMIEPLVLAL | HLA-A*02:03 | 10 | 0.17471 | 100.00% (10/10) | Non-Toxin | 1.46 | 0.54 | -1.16 | 2 | 0.84 | 14.3 |
| 472 | polymerase | SMIEPLVLAL | HLA-A*02:01 | 10 | 0.17471 | 100.00% (10/10) | Non-Toxin | 1.46 | 0.54 | -1.28 | 2 | 0.72 | 19 |
| 473 | polymerase | SMIEPLVLAL | HLA-A*02:06 | 10 | 0.17471 | 100.00% (10/10) | Non-Toxin | 1.46 | 0.54 | -1.36 | 2 | 0.64 | 23 |
| 474 | polymerase | SPSHHLEPY | HLA-B*35:01 | 9 | 0.07793 | 100.00% (10/10) | Non-Toxin | 0.99 | 1.22 | -1.02 | 2.21 | 1.19 | 10.4 |
| 475 | polymerase | SQNLLVTSY | HLA-B*15:01 | 9 | -0.0491 | 100.00% (10/10) | Non-Toxin | 1.6 | 1.31 | -1.38 | 2.9 | 1.52 | 24.2 |
| 476 | polymerase | SQNLLVTSY | HLA-A*30:02 | 9 | -0.0491 | 100.00% (10/10) | Non-Toxin | 1.6 | 1.31 | -1.87 | 2.9 | 1.03 | 74.7 |
| 477 | polymerase | SQSKWYECF | HLA-B*15:01 | 9 | -0.00548 | 100.00% (10/10) | Non-Toxin | 1.23 | 1.17 | -1.85 | 2.4 | 0.55 | 70 |
| 478 | polymerase | SRLFNMYRSY | HLA-B*15:01 | 10 | -0.11956 | 100.00% (10/10) | Non-Toxin | 1.42 | 1.39 | -1.69 | 2.82 | 1.13 | 49 |
| 479 | polymerase | SRLFNMYRSY | HLA-A*30:02 | 10 | -0.11956 | 100.00% (10/10) | Non-Toxin | 1.42 | 1.39 | -1.94 | 2.82 | 0.87 | 87.5 |
| 480 | polymerase | SRLFNMYRSY | HLA-A*03:01 | 10 | -0.11956 | 100.00% (10/10) | Non-Toxin | 1.42 | 1.39 | -2.24 | 2.82 | 0.57 | 174.8 |
| 481 | polymerase | STHLFIYSK | HLA-A*11:01 | 9 | 0.13884 | 90.00% (9/10) | Non-Toxin | 1.13 | 0.23 | -0.72 | 1.36 | 0.64 | 5.3 |
| 482 | polymerase | STLGQSISF | HLA-A*32:01 | 9 | -0.22237 | 100.00% (10/10) | Non-Toxin | 1.44 | 1.1 | -1.63 | 2.54 | 0.91 | 42.7 |
| 483 | polymerase | STLGQSISFY | HLA-A*30:02 | 10 | -0.18934 | 100.00% (10/10) | Non-Toxin | 1.32 | 1.27 | -1.57 | 2.59 | 1.02 | 36.9 |
| 484 | polymerase | STLGQSISFY | HLA-A*11:01 | 10 | -0.18934 | 100.00% (10/10) | Non-Toxin | 1.32 | 1.27 | -1.65 | 2.59 | 0.94 | 44.4 |
| 485 | polymerase | SVLNRVSRY | HLA-A*30:02 | 9 | -0.03023 | 100.00% (10/10) | Non-Toxin | 1.26 | 1.35 | -1.63 | 2.61 | 0.97 | 43.1 |
| 486 | polymerase | SVYPREVLSY | HLA-A*03:01 | 10 | 0.06838 | 100.00% (10/10) | Non-Toxin | 1.31 | 1.37 | -1.52 | 2.68 | 1.16 | 32.8 |
| 487 | polymerase | SVYPREVLSY | HLA-A*11:01 | 10 | 0.06838 | 100.00% (10/10) | Non-Toxin | 1.31 | 1.37 | -1.53 | 2.68 | 1.15 | 34 |
| 488 | polymerase | SVYPREVLSY | HLA-B*15:01 | 10 | 0.06838 | 100.00% (10/10) | Non-Toxin | 1.31 | 1.37 | -1.68 | 2.68 | 1 | 48.2 |
| 489 | polymerase | SVYPREVLSY | HLA-A*30:02 | 10 | 0.06838 | 100.00% (10/10) | Non-Toxin | 1.31 | 1.37 | -1.87 | 2.68 | 0.81 | 73.5 |
| 490 | polymerase | SYFGLVLVCF | HLA-A*23:01 | 10 | 0.0944 | 100.00% (10/10) | Non-Toxin | 1.48 | 1.34 | -1.04 | 2.82 | 1.78 | 10.9 |
| 491 | polymerase | SYFGLVLVCF | HLA-A*24:02 | 10 | 0.0944 | 100.00% (10/10) | Non-Toxin | 1.48 | 1.34 | -1.51 | 2.82 | 1.31 | 32.6 |
| 492 | polymerase | SYMIYLMNW | HLA-A*23:01 | 9 | -0.0891 | 100.00% (10/10) | Non-Toxin | 1.53 | 0.55 | -1.07 | 2.09 | 1.02 | 11.7 |
| 493 | polymerase | SYMIYLMNW | HLA-A*24:02 | 9 | -0.0891 | 100.00% (10/10) | Non-Toxin | 1.53 | 0.55 | -1.31 | 2.09 | 0.78 | 20.2 |
| 494 | polymerase | TEAAVLSRY | HLA-B*44:03 | 9 | -0.02755 | 100.00% (10/10) | Non-Toxin | 1.46 | 1.27 | -1.57 | 2.73 | 1.16 | 37.2 |
| 495 | polymerase | TEAAVLSRY | HLA-B*44:02 | 9 | -0.02755 | 100.00% (10/10) | Non-Toxin | 1.46 | 1.27 | -1.82 | 2.73 | 0.91 | 66.5 |
| 496 | polymerase | TEFNPHNHY | HLA-B*44:03 | 9 | 0.06458 | 100.00% (10/10) | Non-Toxin | 1.36 | 1.24 | -2.09 | 2.6 | 0.51 | 123.6 |
| 497 | polymerase | TEFSINETL | HLA-B*40:01 | 9 | 0.10222 | 100.00% (10/10) | Non-Toxin | 1.6 | 0.42 | -1.14 | 2.02 | 0.88 | 13.7 |
| 498 | polymerase | TEVDNNHLIY | HLA-A*01:01 | 10 | 0.12197 | 100.00% (10/10) | Non-Toxin | 1.3 | 1.21 | -1.57 | 2.52 | 0.95 | 37.2 |
| 499 | polymerase | TIATIPFLF | HLA-A*23:01 | 9 | 0.26324 | 100.00% (10/10) | Non-Toxin | 1.08 | 1.13 | -1.61 | 2.21 | 0.6 | 40.9 |
| 500 | polymerase | TIISTHLFIY | HLA-A*11:01 | 10 | 0.11215 | 90.00% (9/10) | Non-Toxin | 1.26 | 1.27 | -1.98 | 2.53 | 0.55 | 95.7 |
| 501 | polymerase | TIPFLFLSAY | HLA-B*35:01 | 10 | 0.0902 | 100.00% (10/10) | Non-Toxin | 1.39 | 1.38 | -1.6 | 2.77 | 1.17 | 40 |
| 502 | polymerase | TMVDLLSDL | HLA-A*02:03 | 9 | -0.11218 | 100.00% (10/10) | Non-Toxin | 1.2 | 0.48 | -1.16 | 1.69 | 0.53 | 14.3 |
| 503 | polymerase | TMYGWFFVPR | HLA-A*31:01 | 10 | 0.50116 | 100.00% (10/10) | Non-Toxin | 0.67 | 0.83 | -0.72 | 1.5 | 0.78 | 5.2 |
| 504 | polymerase | TMYGWFFVPR | HLA-A*33:01 | 10 | 0.50116 | 100.00% (10/10) | Non-Toxin | 0.67 | 0.83 | -0.81 | 1.5 | 0.69 | 6.4 |
| 505 | polymerase | TQKVHPNLPY | HLA-B*15:01 | 10 | -0.02969 | 100.00% (10/10) | Non-Toxin | 1 | 1.3 | -1.43 | 2.3 | 0.87 | 26.8 |
| 506 | polymerase | TSDLDFVIFY | HLA-A*01:01 | 10 | 0.35122 | 100.00% (10/10) | Non-Toxin | 1.68 | 1.26 | -0.87 | 2.94 | 2.07 | 7.4 |
| 507 | polymerase | TSDLDFVIFY | HLA-A*11:01 | 10 | 0.35122 | 100.00% (10/10) | Non-Toxin | 1.68 | 1.26 | -2.31 | 2.94 | 0.63 | 203.7 |
| 508 | polymerase | TSYMIYLMNW | HLA-B*58:01 | 10 | -0.21432 | 100.00% (10/10) | Non-Toxin | 1.53 | 0.49 | -1.33 | 2.03 | 0.7 | 21.4 |
| 509 | polymerase | TTWIGNLDSY | HLA-A*26:01 | 10 | 0.14414 | 100.00% (10/10) | Non-Toxin | 1.29 | 1.25 | -1.93 | 2.54 | 0.61 | 85.6 |
| 510 | polymerase | TWTIATIPFLF | HLA-A*23:01 | 11 | 0.4339 | 100.00% (10/10) | Non-Toxin | 1.08 | 1.23 | -1.69 | 2.31 | 0.62 | 48.5 |
| 511 | polymerase | VIFYASLTY | HLA-B*15:01 | 9 | -0.07003 | 100.00% (10/10) | Non-Toxin | 1.22 | 1.36 | -1.45 | 2.58 | 1.13 | 28.2 |
| 512 | polymerase | VIFYASLTY | HLA-B*35:01 | 9 | -0.07003 | 100.00% (10/10) | Non-Toxin | 1.22 | 1.36 | -1.47 | 2.58 | 1.11 | 29.4 |
| 513 | polymerase | VIFYASLTY | HLA-A*30:02 | 9 | -0.07003 | 100.00% (10/10) | Non-Toxin | 1.22 | 1.36 | -1.65 | 2.58 | 0.92 | 45.1 |

| | | | | | | | | | | | | | |
|---|---|---|---|---|---|---|---|---|---|---|---|---|---|
| 514 | polymerase | VIFYASLTY | HLA-A*03:01 | 9 | -0.07003 | 100.00% (10/10) | Non-Toxin | 1.22 | 1.36 | -1.74 | 2.58 | 0.84 | 55.3 |
| 515 | polymerase | VIFYASLTY | HLA-A*11:01 | 9 | -0.07003 | 100.00% (10/10) | Non-Toxin | 1.22 | 1.36 | -1.85 | 2.58 | 0.73 | 70 |
| 516 | polymerase | VIFYASLTY | HLA-A*32:01 | 9 | -0.07003 | 100.00% (10/10) | Non-Toxin | 1.22 | 1.36 | -1.95 | 2.58 | 0.63 | 88.6 |
| 517 | polymerase | VMMEDGLLV | HLA-A*02:01 | 9 | 0.08141 | 100.00% (10/10) | Non-Toxin | 0.91 | 0.3 | -0.69 | 1.21 | 0.52 | 4.9 |
| 518 | polymerase | VSNTSKHTY | HLA-A*30:02 | 9 | -0.27716 | 100.00% (10/10) | Non-Toxin | 1.52 | 1.29 | -1.89 | 2.82 | 0.93 | 76.8 |
| 519 | polymerase | VSRLSHHDY | HLA-A*30:02 | 9 | -0.08475 | 100.00% (10/10) | Non-Toxin | 1.17 | 1.36 | -1.98 | 2.52 | 0.54 | 95.8 |
| 520 | polymerase | VTSYMIYLMNW | HLA-B*58:01 | 11 | -0.26484 | 100.00% (10/10) | Non-Toxin | 1.53 | 0.42 | -1.41 | 1.95 | 0.55 | 25.7 |
| 521 | polymerase | WAFEIHHRR | HLA-A*68:01 | 9 | 0.35634 | 100.00% (10/10) | Non-Toxin | 0.86 | 0.75 | -0.6 | 1.61 | 1.01 | 4 |
| 522 | polymerase | WAYGDNEECW | HLA-B*58:01 | 10 | 0.19545 | 100.00% (10/10) | Non-Toxin | 1.63 | 0.48 | -1.39 | 2.11 | 0.71 | 24.7 |
| 523 | polymerase | WESFCGIQF | HLA-B*40:01 | 9 | 0.08814 | 100.00% (10/10) | Non-Toxin | 1.27 | 1.09 | -1.77 | 2.36 | 0.59 | 58.3 |
| 524 | polymerase | WESFCGIQF | HLA-B*44:03 | 9 | 0.08814 | 100.00% (10/10) | Non-Toxin | 1.27 | 1.09 | -1.85 | 2.36 | 0.51 | 70.6 |
| 525 | polymerase | WTIATIPFL | HLA-A*02:06 | 9 | 0.30469 | 100.00% (10/10) | Non-Toxin | 1.53 | 0.43 | -0.76 | 1.96 | 1.2 | 5.8 |
| 526 | polymerase | WTIATIPFL | HLA-A*68:02 | 9 | 0.30469 | 100.00% (10/10) | Non-Toxin | 1.53 | 0.43 | -1.24 | 1.96 | 0.72 | 17.4 |
| 527 | polymerase | WTIATIPFL | HLA-A*02:01 | 9 | 0.30469 | 100.00% (10/10) | Non-Toxin | 1.53 | 0.43 | -1.37 | 1.96 | 0.59 | 23.5 |
| 528 | polymerase | WTIATIPFLF | HLA-B*58:01 | 10 | 0.33185 | 100.00% (10/10) | Non-Toxin | 1.08 | 1.11 | -1.25 | 2.19 | 0.94 | 17.8 |
| 529 | polymerase | YASLTYLRR | HLA-A*68:01 | 9 | -0.00966 | 100.00% (10/10) | Non-Toxin | 1.29 | 0.61 | -1.35 | 1.9 | 0.56 | 22.2 |
| 530 | polymerase | YESMAIFAER | HLA-A*68:01 | 10 | 0.13902 | 100.00% (10/10) | Non-Toxin | 1 | 0.63 | -0.76 | 1.63 | 0.87 | 5.8 |
| 531 | polymerase | YGLPGFFNW | HLA-B*58:01 | 9 | 0.22346 | 100.00% (10/10) | Non-Toxin | 1.52 | 0.27 | -1.12 | 1.8 | 0.67 | 13.2 |
| 532 | polymerase | YIINRTAGR | HLA-A*68:01 | 9 | 0.17645 | 100.00% (10/10) | Non-Toxin | 1.16 | 0.65 | -1.07 | 1.81 | 0.74 | 11.8 |
| 533 | polymerase | YLPSGAQRL | HLA-A*02:03 | 9 | -0.16776 | 100.00% (10/10) | Non-Toxin | 1.69 | 0.47 | -1.62 | 2.15 | 0.53 | 42.1 |
| 534 | polymerase | YLRRGIIKQL | HLA-A*02:03 | 10 | 0.10708 | 100.00% (10/10) | Non-Toxin | 1.81 | 0.44 | -1.51 | 2.25 | 0.74 | 32.3 |
| 535 | polymerase | YLTPEMVLMY | HLA-B*15:01 | 10 | -0.14516 | 100.00% (10/10) | Non-Toxin | 1.52 | 1.23 | -2.16 | 2.75 | 0.58 | 145.9 |
| 536 | polymerase | YMKDKALSPI | HLA-A*02:03 | 10 | -0.37612 | 100.00% (10/10) | Non-Toxin | 0.84 | 0.29 | -0.59 | 1.13 | 0.54 | 3.9 |
| 537 | polymerase | YPECNNILF | HLA-B*35:01 | 9 | 0.0717 | 90.00% (9/10) | Non-Toxin | 1.47 | 0.96 | -0.82 | 2.44 | 1.62 | 6.6 |
| 538 | polymerase | YPECNNILF | HLA-B*53:01 | 9 | 0.0717 | 90.00% (9/10) | Non-Toxin | 1.47 | 0.96 | -1.26 | 2.44 | 1.18 | 18.2 |
| 539 | polymerase | YPKGGIEGY | HLA-B*35:01 | 9 | 0.22668 | 100.00% (10/10) | Non-Toxin | 1.35 | 1.12 | -1.28 | 2.47 | 1.2 | 18.9 |
| 540 | polymerase | YPREVLSY | HLA-B*35:01 | 8 | 0.00769 | 100.00% (10/10) | Non-Toxin | 1.31 | 1.17 | -1.84 | 2.48 | 0.64 | 69.2 |
| 541 | polymerase | YQIDQPFFV | HLA-A*02:06 | 9 | 0.10948 | 100.00% (10/10) | Non-Toxin | 1.07 | 0.17 | -0.2 | 1.24 | 1.03 | 1.6 |
| 542 | polymerase | YQIDQPFFV | HLA-A*02:01 | 9 | 0.10948 | 100.00% (10/10) | Non-Toxin | 1.07 | 0.17 | -0.34 | 1.24 | 0.9 | 2.2 |
| 543 | polymerase | YQQAMLLGL | HLA-A*02:06 | 9 | -0.16923 | 100.00% (10/10) | Non-Toxin | 1.55 | 0.43 | -0.96 | 1.98 | 1.02 | 9.2 |
| 544 | polymerase | YQSTLGQSISF | HLA-B*15:01 | 11 | -0.24531 | 100.00% (10/10) | Non-Toxin | 1.44 | 1.14 | -1.99 | 2.58 | 0.59 | 98 |
| 545 | polymerase | YSNPDSTEVY | HLA-A*01:01 | 10 | -0.0076 | 100.00% (10/10) | Non-Toxin | 1.46 | 1.26 | -1.72 | 2.72 | 1 | 52.3 |
| 546 | polymerase | YSNPDSTEVY | HLA-B*15:01 | 10 | -0.0076 | 100.00% (10/10) | Non-Toxin | 1.46 | 1.26 | -1.95 | 2.72 | 0.77 | 88.9 |
| 547 | polymerase | YSNPDSTEVY | HLA-B*35:01 | 10 | -0.0076 | 100.00% (10/10) | Non-Toxin | 1.46 | 1.26 | -2.16 | 2.72 | 0.56 | 145 |
| 548 | polymerase | YTEVDNNHLIY | HLA-A*01:01 | 11 | 0.18189 | 100.00% (10/10) | Non-Toxin | 1.3 | 1.12 | -0.96 | 2.42 | 1.46 | 9.1 |
| 549 | polymerase | YTPGFPISR | HLA-A*68:01 | 9 | 0.14972 | 100.00% (10/10) | Non-Toxin | 1.08 | 0.69 | -1 | 1.76 | 0.76 | 10.1 |
| 550 | polymerase | YYQIDQPFF | HLA-A*23:01 | 9 | 0.06792 | 100.00% (10/10) | Non-Toxin | 1.33 | 1.24 | -1.43 | 2.57 | 1.14 | 26.8 |
| 551 | polymerase | YYQIDQPFF | HLA-A*24:02 | 9 | 0.06792 | 100.00% (10/10) | Non-Toxin | 1.33 | 1.24 | -1.54 | 2.57 | 1.02 | 35 |
| 552 | polymerase | YYQIDQPFFV | HLA-A*02:06 | 10 | 0.16188 | 100.00% (10/10) | Non-Toxin | 1.07 | 0.27 | -0.59 | 1.34 | 0.75 | 3.9 |
| | **Protein** | Peptide | **Allele** | **Peptide Length** | **Immunogenicity** | **Conservancy** | **Toxicity** | **Proteasome Score** | **TAP Score** | **MHC Score** | **Processing Score** | **Total Score** | **MHC IC50[nM]** |
| 553 | V protein | CLVSDAKMLSY | HLA-A*01:01 | 11 | -0.57469 | 19.30% (11/57) | Non-Toxin | 1.48 | 1.31 | -1.82 | 2.8 | 0.98 | 65.8 |
| 554 | V protein | DPVVTDVVY | HLA-B*35:01 | 9 | 0.17258 | 98.25% (56/57) | Non-Toxin | 1.74 | 1.05 | -1.4 | 2.8 | 1.4 | 25.1 |
| 555 | V protein | GLNPTAVPF | HLA-B*15:01 | 9 | 0.08973 | 19.30% (11/57) | Non-Toxin | 1.12 | 0.99 | -1.4 | 2.12 | 0.72 | 24.9 |
| 556 | V protein | KMLSYAPEI | HLA-A*02:01 | 9 | -0.0877 | 19.30% (11/57) | Non-Toxin | 1.24 | 0.4 | -0.68 | 1.65 | 0.97 | 4.8 |
| 557 | V protein | KMLSYAPEI | HLA-A*02:06 | 9 | -0.0877 | 19.30% (11/57) | Non-Toxin | 1.24 | 0.4 | -0.82 | 1.65 | 0.83 | 6.6 |
| 558 | V protein | KMLSYAPEI | HLA-A*32:01 | 9 | -0.0877 | 19.30% (11/57) | Non-Toxin | 1.24 | 0.4 | -1.13 | 1.65 | 0.52 | 13.4 |
| 559 | V protein | KSRGIPIKK | HLA-A*30:01 | 9 | 0.15638 | 100.00% (57/57) | Non-Toxin | 1.05 | 0.22 | -0.73 | 1.27 | 0.54 | 5.4 |

| | | | | | | | | | | | | | |
|---|---|---|---|---|---|---|---|---|---|---|---|---|---|
| 560 | V protein | LDPVVTDVVY | HLA-B*35:01 | 10 | 0.19578 | 98.25% (56/57) | Non-Toxin | 1.74 | 1.11 | -1.34 | 2.85 | 1.51 | 22.1 |
| 561 | V protein | LVSDAKMLSY | HLA-A*01:01 | 10 | -0.4746 | 19.30% (11/57) | Non-Toxin | 1.48 | 1.33 | -1.26 | 2.81 | 1.55 | 18 |
| 562 | V protein | LVSDAKMLSY | HLA-B*15:01 | 10 | -0.4746 | 19.30% (11/57) | Non-Toxin | 1.48 | 1.33 | -1.94 | 2.81 | 0.87 | 87.9 |
| 563 | V protein | NPTAVPFTLR | HLA-A*68:01 | 10 | 0.21785 | 19.30% (11/57) | Non-Toxin | 1.01 | 0.53 | -1.03 | 1.54 | 0.51 | 10.7 |
| 564 | V protein | SPVIAEHYY | HLA-B*35:01 | 9 | 0.30481 | 98.25% (56/57) | Non-Toxin | 1.36 | 1.13 | -1.31 | 2.5 | 1.18 | 20.6 |
| 565 | V protein | VSDAKMLSY | HLA-A*01:01 | 9 | -0.43475 | 19.30% (11/57) | Non-Toxin | 1.48 | 1.26 | -0.94 | 2.74 | 1.8 | 8.7 |
| | **Protein** | Peptide | **Allele** | **Peptide Length** | **Immunogenicity** | **Conservancy** | **Toxicity** | **Proteasome Score** | **TAP Score** | **MHC Score** | **Processing Score** | **Total Score** | **MHC IC50[nM]** |
| 566 | W protein | CLVSDAKMLSY | HLA-A*01:01 | 11 | -0.57469 | 20.51% (8/39) | Non-Toxin | 1.48 | 1.31 | -1.82 | 2.8 | 0.98 | 65.8 |
| 567 | W protein | DPVVTDVVY | HLA-B*35:01 | 9 | 0.17258 | 97.44% (38/39) | Non-Toxin | 1.74 | 1.05 | -1.4 | 2.8 | 1.4 | 25.1 |
| 568 | W protein | GLNPTAVPF | HLA-B*15:01 | 9 | 0.08973 | 20.51% (8/39) | Non-Toxin | 1.12 | 0.99 | -1.4 | 2.12 | 0.72 | 24.9 |
| 569 | W protein | KMLSYAPEI | HLA-A*02:01 | 9 | -0.0877 | 20.51% (8/39) | Non-Toxin | 1.24 | 0.4 | -0.68 | 1.65 | 0.97 | 4.8 |
| 570 | W protein | KMLSYAPEI | HLA-A*02:06 | 9 | -0.0877 | 20.51% (8/39) | Non-Toxin | 1.24 | 0.4 | -0.82 | 1.65 | 0.83 | 6.6 |
| 571 | W protein | KMLSYAPEI | HLA-A*32:01 | 9 | -0.0877 | 20.51% (8/39) | Non-Toxin | 1.24 | 0.4 | -1.13 | 1.65 | 0.52 | 13.4 |
| 572 | W protein | KSRGIPIKK | HLA-A*30:01 | 9 | 0.15638 | 100.00% (39/39) | Non-Toxin | 1.08 | 0.22 | -0.73 | 1.31 | 0.57 | 5.4 |
| 573 | W protein | LDPVVTDVVY | HLA-B*35:01 | 10 | 0.19578 | 97.44% (38/39) | Non-Toxin | 1.74 | 1.11 | -1.34 | 2.85 | 1.51 | 22.1 |
| 574 | W protein | LVSDAKMLSY | HLA-A*01:01 | 10 | -0.4746 | 20.51% (8/39) | Non-Toxin | 1.48 | 1.33 | -1.26 | 2.81 | 1.55 | 18 |
| 575 | W protein | LVSDAKMLSY | HLA-B*15:01 | 10 | -0.4746 | 20.51% (8/39) | Non-Toxin | 1.48 | 1.33 | -1.94 | 2.81 | 0.87 | 87.9 |
| 576 | W protein | NPTAVPFTLR | HLA-A*68:01 | 10 | 0.21785 | 20.51% (8/39) | Non-Toxin | 1.01 | 0.53 | -1.03 | 1.54 | 0.51 | 10.7 |
| 577 | W protein | QTRNIHLLGR | HLA-A*31:01 | 10 | 0.17139 | 97.44% (38/39) | Non-Toxin | 1.11 | 0.73 | -1.39 | 1.84 | 0.44 | 24.6 |
| 578 | W protein | SPVIAEHYY | HLA-B*35:01 | 9 | 0.30481 | 100.00% (39/39) | Non-Toxin | 1.36 | 1.13 | -1.31 | 2.5 | 1.18 | 20.6 |
| 579 | W protein | VSDAKMLSY | HLA-A*01:01 | 9 | -0.43475 | 20.51% (8/39) | Non-Toxin | 1.48 | 1.26 | -0.94 | 2.74 | 1.8 | 8.7 |

**Supplementary Table S2:** High Scoring CTL epitopes-HLA allele pairs screened from entire proteome of SARS-CoV-2 by the "MHC-I Binding Predictions" tool of IEDB. These epitopes were further utilized to identify the potentially immunogenic multiple epitope cluster based CTL Ag-Patches from the entire proteome of the SRAS-CoV-2. The screened epitopes are in consensus with the previous studies[38].

| S.No. | Protein | Peptide | Allele | Length | Score | Percentile rank | Immunogenicity | Conservancy | Toxicity | Method |
|---|---|---|---|---|---|---|---|---|---|---|
| 1 | C Protein | APVENLKKL | HLA-B*07:02 | 9 | 0.823222 | 0.07 | -0.21059 | 1.67% (1/60) | Non-Toxin | IEDB recommended 2020.09 (NetMHCpan EL 4.1) |
| 2 | C Protein | ASILLTLFR | HLA-A*31:01 | 9 | 0.728722 | 0.11 | 0.11682 | 100.00% (60/60) | Non-Toxin | IEDB recommended 2020.09 (NetMHCpan EL 4.1) |
| 3 | C Protein | ASILLTLFR | HLA-A*11:01 | 9 | 0.665698 | 0.16 | 0.11682 | 100.00% (60/60) | Non-Toxin | IEDB recommended 2020.09 (NetMHCpan EL 4.1) |
| 4 | C Protein | CPMLPKLQF | HLA-B*53:01 | 9 | 0.79741 | 0.04 | -0.359 | 100.00% (60/60) | Non-Toxin | IEDB recommended 2020.09 (NetMHCpan EL 4.1) |
| 5 | C Protein | CPMLPKLQF | HLA-B*35:01 | 9 | 0.787207 | 0.08 | -0.359 | 100.00% (60/60) | Non-Toxin | IEDB recommended 2020.09 (NetMHCpan EL 4.1) |
| 6 | C Protein | CPMLPKLQF | HLA-B*07:02 | 9 | 0.744382 | 0.11 | -0.359 | 100.00% (60/60) | Non-Toxin | IEDB recommended 2020.09 (NetMHCpan EL 4.1) |
| 7 | C Protein | EQTMGMYVLY | HLA-A*26:01 | 10 | 0.663978 | 0.06 | -0.27722 | 100.00% (60/60) | Non-Toxin | IEDB recommended 2020.09 (NetMHCpan EL 4.1) |
| 8 | C Protein | FLKKIGKL | HLA-B*08:01 | 8 | 0.710079 | 0.06 | -0.3075 | 98.33% (59/60) | Non-Toxin | IEDB recommended 2020.09 (NetMHCpan EL 4.1) |
| 9 | C Protein | GQKVQMTY | HLA-B*15:01 | 8 | 0.901547 | 0.02 | -0.2738 | 95.00% (57/60) | Non-Toxin | IEDB recommended 2020.09 (NetMHCpan EL 4.1) |
| 10 | C Protein | HQEQTMGMY | HLA-A*30:02 | 9 | 0.515293 | 0.16 | -0.28556 | 100.00% (60/60) | Non-Toxin | IEDB recommended 2020.09 (NetMHCpan EL 4.1) |
| 11 | C Protein | ITHQEQTMGMY | HLA-A*01:01 | 11 | 0.610291 | 0.14 | -0.32286 | 100.00% (60/60) | Non-Toxin | IEDB recommended 2020.09 (NetMHCpan EL 4.1) |
| 12 | C Protein | IYPVLLPQM | HLA-A*24:02 | 9 | 0.86784 | 0.04 | -0.06034 | 100.00% (60/60) | Non-Toxin | IEDB recommended 2020.09 (NetMHCpan EL 4.1) |
| 13 | C Protein | IYPVLLPQM | HLA-A*23:01 | 9 | 0.818931 | 0.04 | -0.06034 | 100.00% (60/60) | Non-Toxin | IEDB recommended 2020.09 (NetMHCpan EL 4.1) |
| 14 | C Protein | KLQFLKKIGK | HLA-A*03:01 | 10 | 0.719129 | 0.15 | -0.21148 | 98.33% (59/60) | Non-Toxin | IEDB recommended 2020.09 (NetMHCpan EL 4.1) |
| 15 | C Protein | KTEQKHGRTF | HLA-B*44:02 | 10 | 0.654226 | 0.12 | -0.1643 | 26.67% (16/60) | Non-Toxin | IEDB recommended 2020.09 (NetMHCpan EL 4.1) |
| 16 | C Protein | MASILLTLF | HLA-B*53:01 | 9 | 0.625818 | 0.07 | 0.08526 | 100.00% (60/60) | Non-Toxin | IEDB recommended 2020.09 (NetMHCpan EL 4.1) |

| | | | | | | | | | | |
|---|---|---|---|---|---|---|---|---|---|---|
| 17 | C Protein | MASILLTLF | HLA-B*58:01 | 9 | 0.816922 | 0.11 | 0.08526 | 100.00% (60/60) | Non-Toxin | IEDB recommended 2020.09 (NetMHCpan EL 4.1) |
| 18 | C Protein | MEVLKEEAW | HLA-B*44:02 | 9 | 0.912517 | 0.02 | -0.00615 | 78.33% (47/60) | Non-Toxin | IEDB recommended 2020.09 (NetMHCpan EL 4.1) |
| 19 | C Protein | MEVLKEEAW | HLA-B*44:03 | 9 | 0.918867 | 0.03 | -0.00615 | 78.33% (47/60) | Non-Toxin | IEDB recommended 2020.09 (NetMHCpan EL 4.1) |
| 20 | C Protein | MMASILLTL | HLA-A*32:01 | 9 | 0.557134 | 0.07 | -0.02129 | 100.00% (60/60) | Non-Toxin | IEDB recommended 2020.09 (NetMHCpan EL 4.1) |
| 21 | C Protein | MMASILLTL | HLA-A*02:01 | 9 | 0.771922 | 0.09 | -0.02129 | 100.00% (60/60) | Non-Toxin | IEDB recommended 2020.09 (NetMHCpan EL 4.1) |
| 22 | C Protein | MMASILLTL | HLA-A*02:03 | 9 | 0.727441 | 0.09 | -0.02129 | 100.00% (60/60) | Non-Toxin | IEDB recommended 2020.09 (NetMHCpan EL 4.1) |
| 23 | C Protein | MTYNWTQW | HLA-B*57:01 | 8 | 0.913336 | 0.09 | 0.14677 | 95.00% (57/60) | Non-Toxin | IEDB recommended 2020.09 (NetMHCpan EL 4.1) |
| 24 | C Protein | MTYNWTQW | HLA-B*58:01 | 8 | 0.808826 | 0.11 | 0.14677 | 95.00% (57/60) | Non-Toxin | IEDB recommended 2020.09 (NetMHCpan EL 4.1) |
| 25 | C Protein | MTYNWTQWL | HLA-A*68:02 | 9 | 0.575361 | 0.14 | 0.27619 | 95.00% (57/60) | Non-Toxin | IEDB recommended 2020.09 (NetMHCpan EL 4.1) |
| 26 | C Protein | MYVLYLMQR | HLA-A*33:01 | 9 | 0.734683 | 0.06 | -0.22768 | 100.00% (60/60) | Non-Toxin | IEDB recommended 2020.09 (NetMHCpan EL 4.1) |
| 27 | C Protein | QAFNNPASK | HLA-A*11:01 | 9 | 0.670372 | 0.16 | -0.04889 | 73.33% (44/60) | Non-Toxin | IEDB recommended 2020.09 (NetMHCpan EL 4.1) |
| 28 | C Protein | QAFNNPASK | HLA-A*30:01 | 9 | 0.476561 | 0.16 | -0.04889 | 73.33% (44/60) | Non-Toxin | IEDB recommended 2020.09 (NetMHCpan EL 4.1) |
| 29 | C Protein | QEQTMGMYVLY | HLA-B*44:03 | 11 | 0.693842 | 0.15 | -0.28266 | 100.00% (60/60) | Non-Toxin | IEDB recommended 2020.09 (NetMHCpan EL 4.1) |
| 30 | C Protein | QEQTMGMYVLY | HLA-B*44:02 | 11 | 0.57723 | 0.15 | -0.28266 | 100.00% (60/60) | Non-Toxin | IEDB recommended 2020.09 (NetMHCpan EL 4.1) |
| 31 | C Protein | QTMGMYVLY | HLA-A*26:01 | 9 | 0.775279 | 0.05 | -0.16902 | 100.00% (60/60) | Non-Toxin | IEDB recommended 2020.09 (NetMHCpan EL 4.1) |
| 32 | C Protein | QTMGMYVLY | HLA-A*30:02 | 9 | 0.715845 | 0.05 | -0.16902 | 100.00% (60/60) | Non-Toxin | IEDB recommended 2020.09 (NetMHCpan EL 4.1) |
| 33 | C Protein | QTMGMYVLY | HLA-A*32:01 | 9 | 0.43165 | 0.14 | -0.16902 | 100.00% (60/60) | Non-Toxin | IEDB recommended 2020.09 (NetMHCpan EL 4.1) |
| 34 | C Protein | QTMGMYVLY | HLA-A*01:01 | 9 | 0.576389 | 0.15 | -0.16902 | 100.00% (60/60) | Non-Toxin | IEDB recommended 2020.09 (NetMHCpan EL 4.1) |
| 35 | C Protein | RCCPMLPKLQF | HLA-B*07:02 | 11 | 0.814723 | 0.08 | -0.5013 | 100.00% (60/60) | Non-Toxin | IEDB recommended 2020.09 (NetMHCpan EL 4.1) |
| 36 | C Protein | RMMEVLKEEAW | HLA-A*32:01 | 11 | 0.668454 | 0.05 | 0.06476 | 78.33% (47/60) | Non-Toxin | IEDB recommended 2020.09 (NetMHCpan EL 4.1) |
| 37 | C Protein | SILLTLFRR | HLA-A*31:01 | 9 | 0.676568 | 0.15 | 0.14164 | 100.00% (60/60) | Non-Toxin | IEDB recommended 2020.09 (NetMHCpan EL 4.1) |
| 38 | C Protein | SKTEQKHGRTF | HLA-B*44:02 | 11 | 0.777859 | 0.06 | -0.07969 | 26.67% (16/60) | Non-Toxin | IEDB recommended 2020.09 (NetMHCpan EL 4.1) |
| 39 | C Protein | SKTEQKHGRTF | HLA-B*44:03 | 11 | 0.701106 | 0.14 | -0.07969 | 26.67% (16/60) | Non-Toxin | IEDB recommended 2020.09 (NetMHCpan EL 4.1) |
| 40 | C Protein | TEQKHGRTF | HLA-B*44:03 | 9 | 0.96597 | 0.01 | -0.12484 | 26.67% (16/60) | Non-Toxin | IEDB recommended 2020.09 (NetMHCpan EL 4.1) |
| 41 | C Protein | TEQKHGRTF | HLA-B*44:02 | 9 | 0.965601 | 0.01 | -0.12484 | 26.67% (16/60) | Non-Toxin | IEDB recommended 2020.09 (NetMHCpan EL 4.1) |
| 42 | C Protein | TGQKVQMTY | HLA-A*30:02 | 9 | 0.582272 | 0.11 | -0.44896 | 95.00% (57/60) | Non-Toxin | IEDB recommended 2020.09 (NetMHCpan EL 4.1) |
| 43 | C Protein | TLFRRTKKK | HLA-A*03:01 | 9 | 0.891743 | 0.03 | -0.13098 | 100.00% (60/60) | Non-Toxin | IEDB recommended 2020.09 (NetMHCpan EL 4.1) |
| 44 | C Protein | VLLPQMELL | HLA-A*02:01 | 9 | 0.870151 | 0.05 | -0.21484 | 100.00% (60/60) | Non-Toxin | IEDB recommended 2020.09 (NetMHCpan EL 4.1) |
| 45 | C Protein | VLLPQMELL | HLA-A*02:06 | 9 | 0.742716 | 0.09 | -0.21484 | 100.00% (60/60) | Non-Toxin | IEDB recommended 2020.09 (NetMHCpan EL 4.1) |
| 46 | C Protein | VPDMDLLQAL | HLA-B*07:02 | 10 | 0.69013 | 0.13 | -0.24404 | 100.00% (60/60) | Non-Toxin | IEDB recommended 2020.09 (NetMHCpan EL 4.1) |
| 47 | C Protein | VTGQKVQMTY | HLA-A*30:02 | 10 | 0.509179 | 0.16 | -0.50992 | 95.00% (57/60) | Non-Toxin | IEDB recommended 2020.09 (NetMHCpan EL 4.1) |
| 48 | C Protein | WTQWLQTLY | HLA-A*01:01 | 9 | 0.771517 | 0.07 | 0.09173 | 100.00% (60/60) | Non-Toxin | IEDB recommended 2020.09 (NetMHCpan EL 4.1) |
| 49 | C Protein | YPVLLPQM | HLA-B*51:01 | 8 | 0.69854 | 0.07 | -0.11676 | 100.00% (60/60) | Non-Toxin | IEDB recommended 2020.09 (NetMHCpan EL 4.1) |
| 50 | C Protein | YPVLLPQM | HLA-B*35:01 | 8 | 0.717162 | 0.12 | -0.11676 | 100.00% (60/60) | Non-Toxin | IEDB recommended 2020.09 (NetMHCpan EL 4.1) |
| **S.No.** | **Protein** | **Peptide** | **Allele** | **Length** | **Score** | **Percentile rank** | **Immunogenicity** | **Conservancy** | **Toxicity** | **Method** |
| 51 | Glycoprotein | AENPVFTVF | HLA-B*44:03 | 9 | 0.9927 | 0.01 | 0.19402 | 100.00% (66/66) | Non-Toxin | IEDB recommended 2020.09 (NetMHCpan EL 4.1) |
| 52 | Glycoprotein | AENPVFTVF | HLA-B*44:02 | 9 | 0.985092 | 0.01 | 0.19402 | 100.00% (66/66) | Non-Toxin | IEDB recommended 2020.09 (NetMHCpan EL 4.1) |
| 53 | Glycoprotein | AENPVFTVF | HLA-B*40:01 | 9 | 0.961615 | 0.02 | 0.19402 | 100.00% (66/66) | Non-Toxin | IEDB recommended 2020.09 (NetMHCpan EL 4.1) |
| 54 | Glycoprotein | ALQSIQQQI | HLA-A*02:03 | 9 | 0.75991 | 0.07 | -0.34895 | 75.76% (50/66) | Non-Toxin | IEDB recommended 2020.09 (NetMHCpan EL 4.1) |
| 55 | Glycoprotein | ALQSIQQQI | HLA-A*02:01 | 9 | 0.719158 | 0.12 | -0.34895 | 75.76% (50/66) | Non-Toxin | IEDB recommended 2020.09 (NetMHCpan EL 4.1) |
| 56 | Glycoprotein | AMDEGYFAY | HLA-A*01:01 | 9 | 0.938501 | 0.02 | 0.25913 | 90.91% (60/66) | Non-Toxin | IEDB recommended 2020.09 (NetMHCpan EL 4.1) |
| 57 | Glycoprotein | AMDEGYFAY | HLA-A*30:02 | 9 | 0.709552 | 0.05 | 0.25913 | 90.91% (60/66) | Non-Toxin | IEDB recommended 2020.09 (NetMHCpan EL 4.1) |
| 58 | Glycoprotein | AMIKDALQSI | HLA-A*02:03 | 10 | 0.69459 | 0.1 | -0.31896 | 71.21% (47/66) | Non-Toxin | IEDB recommended 2020.09 (NetMHCpan EL 4.1) |
| 59 | Glycoprotein | AQKTITNCF | HLA-B*15:01 | 9 | 0.969264 | 0.01 | 0.09824 | 100.00% (66/66) | Non-Toxin | IEDB recommended 2020.09 (NetMHCpan EL 4.1) |
| 60 | Glycoprotein | ASDKGKNPSK | HLA-A*30:01 | 10 | 0.58442 | 0.09 | -0.49891 | 75.76% (50/66) | Non-Toxin | IEDB recommended 2020.09 (NetMHCpan EL 4.1) |
| 61 | Glycoprotein | ASFSWDTMIK | HLA-A*11:01 | 10 | 0.809104 | 0.07 | 0.07493 | 100.00% (66/66) | Non-Toxin | IEDB recommended 2020.09 (NetMHCpan EL 4.1) |
| 62 | Glycoprotein | ASINENVNEK | HLA-A*11:01 | 10 | 0.759701 | 0.1 | 0.21979 | 100.00% (66/66) | Non-Toxin | IEDB recommended 2020.09 (NetMHCpan EL 4.1) |
| 63 | Glycoprotein | AVKPKNNGESY | HLA-B*15:01 | 11 | 0.821766 | 0.05 | -0.28402 | 71.21% (47/66) | Non-Toxin | IEDB recommended 2020.09 (NetMHCpan EL 4.1) |
| 64 | Glycoprotein | AVKPKNNGESY | HLA-A*30:02 | 11 | 0.566904 | 0.12 | -0.28402 | 71.21% (47/66) | Non-Toxin | IEDB recommended 2020.09 (NetMHCpan EL 4.1) |
| 65 | Glycoprotein | AVYNNEFYY | HLA-A*30:02 | 9 | 0.891164 | 0.01 | 0.17688 | 90.91% (60/66) | Non-Toxin | IEDB recommended 2020.09 (NetMHCpan EL 4.1) |
| 66 | Glycoprotein | AVYNNEFYY | HLA-A*11:01 | 9 | 0.880649 | 0.03 | 0.17688 | 90.91% (60/66) | Non-Toxin | IEDB recommended 2020.09 (NetMHCpan EL 4.1) |

| | | | | | | | | | | |
|---|---|---|---|---|---|---|---|---|---|---|
| 67 | Glycoprotein | CPNPLPFREY | HLA-B*53:01 | 10 | 0.52569 | 0.12 | 0.17752 | 100.00% (66/66) | Non-Toxin | IEDB recommended 2020.09 (NetMHCpan EL 4.1) |
| 68 | Glycoprotein | DAFLIDRI | HLA-B*51:01 | 8 | 0.928331 | 0.01 | 0.221 | 100.00% (66/66) | Non-Toxin | IEDB recommended 2020.09 (NetMHCpan EL 4.1) |
| 69 | Glycoprotein | DPVLNSTYW | HLA-B*53:01 | 9 | 0.922322 | 0.02 | -0.12919 | 1.52% (1/66) | Non-Toxin | IEDB recommended 2020.09 (NetMHCpan EL 4.1) |
| 70 | Glycoprotein | DSLGQPVFY | HLA-A*26:01 | 9 | 0.537647 | 0.1 | 0.0105 | 100.00% (66/66) | Non-Toxin | IEDB recommended 2020.09 (NetMHCpan EL 4.1) |
| 71 | Glycoprotein | DVQTVNPLV | HLA-A*68:02 | 9 | 0.604285 | 0.12 | 0.01973 | 78.79% (52/66) | Non-Toxin | IEDB recommended 2020.09 (NetMHCpan EL 4.1) |
| 72 | Glycoprotein | EENSKIVFI | HLA-B*44:02 | 9 | 0.814634 | 0.05 | -0.15005 | 66.67% (44/66) | Non-Toxin | IEDB recommended 2020.09 (NetMHCpan EL 4.1) |
| 73 | Glycoprotein | EENSKIVFI | HLA-B*44:03 | 9 | 0.806918 | 0.08 | -0.15005 | 66.67% (44/66) | Non-Toxin | IEDB recommended 2020.09 (NetMHCpan EL 4.1) |
| 74 | Glycoprotein | EIGPKVSL | HLA-B*08:01 | 8 | 0.686032 | 0.07 | -0.31092 | 100.00% (66/66) | Non-Toxin | IEDB recommended 2020.09 (NetMHCpan EL 4.1) |
| 75 | Glycoprotein | EISDQRLSI | HLA-A*68:02 | 9 | 0.605462 | 0.12 | -0.20148 | 98.48% (65/66) | Non-Toxin | IEDB recommended 2020.09 (NetMHCpan EL 4.1) |
| 76 | Glycoprotein | EIYDTGDNV | HLA-A*68:02 | 9 | 0.818284 | 0.05 | 0.10576 | 100.00% (66/66) | Non-Toxin | IEDB recommended 2020.09 (NetMHCpan EL 4.1) |
| 77 | Glycoprotein | EIYDTGDNVIR | HLA-A*68:01 | 11 | 0.922043 | 0.06 | 0.22003 | 100.00% (66/66) | Non-Toxin | IEDB recommended 2020.09 (NetMHCpan EL 4.1) |
| 78 | Glycoprotein | EVLDRGDEV | HLA-A*68:02 | 9 | 0.653933 | 0.1 | 0.17824 | 100.00% (66/66) | Non-Toxin | IEDB recommended 2020.09 (NetMHCpan EL 4.1) |
| 79 | Glycoprotein | EVPSLFMTNV | HLA-A*68:02 | 10 | 0.673201 | 0.09 | -0.20319 | 100.00% (66/66) | Non-Toxin | IEDB recommended 2020.09 (NetMHCpan EL 4.1) |
| 80 | Glycoprotein | FAYSHLEKI | HLA-B*51:01 | 9 | 0.875973 | 0.02 | -0.18811 | 74.24% (49/66) | Non-Toxin | IEDB recommended 2020.09 (NetMHCpan EL 4.1) |
| 81 | Glycoprotein | FAYSHLEKI | HLA-A*02:06 | 9 | 0.778243 | 0.08 | -0.18811 | 74.24% (49/66) | Non-Toxin | IEDB recommended 2020.09 (NetMHCpan EL 4.1) |
| 82 | Glycoprotein | FLIDRINWI | HLA-A*02:03 | 9 | 0.955217 | 0.02 | 0.36516 | 100.00% (66/66) | Non-Toxin | IEDB recommended 2020.09 (NetMHCpan EL 4.1) |
| 83 | Glycoprotein | FLIDRINWI | HLA-A*02:01 | 9 | 0.926293 | 0.03 | 0.36516 | 100.00% (66/66) | Non-Toxin | IEDB recommended 2020.09 (NetMHCpan EL 4.1) |
| 84 | Glycoprotein | FLIDRINWI | HLA-A*02:06 | 9 | 0.829268 | 0.07 | 0.36516 | 100.00% (66/66) | Non-Toxin | IEDB recommended 2020.09 (NetMHCpan EL 4.1) |
| 85 | Glycoprotein | FLIDRINWISA | HLA-A*02:03 | 11 | 0.833318 | 0.05 | 0.46339 | 100.00% (66/66) | Non-Toxin | IEDB recommended 2020.09 (NetMHCpan EL 4.1) |
| 86 | Glycoprotein | FLIDRINWISA | HLA-A*02:01 | 11 | 0.779718 | 0.09 | 0.46339 | 100.00% (66/66) | Non-Toxin | IEDB recommended 2020.09 (NetMHCpan EL 4.1) |
| 87 | Glycoprotein | FPAVGFLV | HLA-B*51:01 | 8 | 0.831141 | 0.03 | 0.18808 | 100.00% (66/66) | Non-Toxin | IEDB recommended 2020.09 (NetMHCpan EL 4.1) |
| 88 | Glycoprotein | FSWDTMIKF | HLA-B*58:01 | 9 | 0.824772 | 0.1 | -0.04696 | 100.00% (66/66) | Non-Toxin | IEDB recommended 2020.09 (NetMHCpan EL 4.1) |
| 89 | Glycoprotein | GESYNQHQF | HLA-B*44:02 | 9 | 0.933642 | 0.02 | -0.21314 | 75.76% (50/66) | Non-Toxin | IEDB recommended 2020.09 (NetMHCpan EL 4.1) |
| 90 | Glycoprotein | GESYNQHQF | HLA-B*44:03 | 9 | 0.94744 | 0.03 | -0.21314 | 75.76% (50/66) | Non-Toxin | IEDB recommended 2020.09 (NetMHCpan EL 4.1) |
| 91 | Glycoprotein | GESYNQHQF | HLA-B*40:01 | 9 | 0.864791 | 0.08 | -0.21314 | 75.76% (50/66) | Non-Toxin | IEDB recommended 2020.09 (NetMHCpan EL 4.1) |
| 92 | Glycoprotein | GIKQGDTLY | HLA-B*15:01 | 9 | 0.892776 | 0.03 | -0.1064 | 100.00% (66/66) | Non-Toxin | IEDB recommended 2020.09 (NetMHCpan EL 4.1) |
| 93 | Glycoprotein | GLADKIGTEI | HLA-A*02:03 | 10 | 0.861059 | 0.04 | 0.07778 | 100.00% (66/66) | Non-Toxin | IEDB recommended 2020.09 (NetMHCpan EL 4.1) |
| 94 | Glycoprotein | GLLDSKILSA | HLA-A*02:01 | 10 | 0.735282 | 0.11 | -0.33312 | 98.48% (65/66) | Non-Toxin | IEDB recommended 2020.09 (NetMHCpan EL 4.1) |
| 95 | Glycoprotein | GLLDSKILSA | HLA-A*02:03 | 10 | 0.65261 | 0.11 | -0.33312 | 98.48% (65/66) | Non-Toxin | IEDB recommended 2020.09 (NetMHCpan EL 4.1) |
| 96 | Glycoprotein | GTEIGPKVSL | HLA-B*40:01 | 10 | 0.970847 | 0.02 | -0.0762 | 100.00% (66/66) | Non-Toxin | IEDB recommended 2020.09 (NetMHCpan EL 4.1) |
| 97 | Glycoprotein | IGTEIGPKVSL | HLA-B*40:01 | 11 | 0.842277 | 0.09 | 0.00033 | 100.00% (66/66) | Non-Toxin | IEDB recommended 2020.09 (NetMHCpan EL 4.1) |
| 98 | Glycoprotein | ILKPKLISY | HLA-B*15:01 | 9 | 0.974856 | 0.01 | -0.28594 | 96.97% (64/66) | Non-Toxin | IEDB recommended 2020.09 (NetMHCpan EL 4.1) |
| 99 | Glycoprotein | ILKPKLISY | HLA-A*30:02 | 9 | 0.7787 | 0.03 | -0.28594 | 96.97% (64/66) | Non-Toxin | IEDB recommended 2020.09 (NetMHCpan EL 4.1) |
| 100 | Glycoprotein | ILKPKLISY | HLA-A*32:01 | 9 | 0.613876 | 0.06 | -0.28594 | 96.97% (64/66) | Non-Toxin | IEDB recommended 2020.09 (NetMHCpan EL 4.1) |
| 101 | Glycoprotein | ILKPKLISYTL | HLA-B*08:01 | 11 | 0.565422 | 0.11 | -0.30853 | 96.97% (64/66) | Non-Toxin | IEDB recommended 2020.09 (NetMHCpan EL 4.1) |
| 102 | Glycoprotein | ILRSGLLKY | HLA-B*15:01 | 9 | 0.915685 | 0.02 | -0.26247 | 98.48% (65/66) | Non-Toxin | IEDB recommended 2020.09 (NetMHCpan EL 4.1) |
| 103 | Glycoprotein | ILRSGLLKY | HLA-A*30:02 | 9 | 0.735218 | 0.04 | -0.26247 | 98.48% (65/66) | Non-Toxin | IEDB recommended 2020.09 (NetMHCpan EL 4.1) |
| 104 | Glycoprotein | IYDSLGQPVF | HLA-A*24:02 | 10 | 0.835396 | 0.05 | -0.23135 | 96.97% (64/66) | Non-Toxin | IEDB recommended 2020.09 (NetMHCpan EL 4.1) |
| 105 | Glycoprotein | IYDSLGQPVF | HLA-A*23:01 | 10 | 0.736497 | 0.07 | -0.23135 | 96.97% (64/66) | Non-Toxin | IEDB recommended 2020.09 (NetMHCpan EL 4.1) |
| 106 | Glycoprotein | KGKNPSKVIK | HLA-A*30:01 | 10 | 0.583524 | 0.09 | -0.33881 | 80.30% (53/66) | Non-Toxin | IEDB recommended 2020.09 (NetMHCpan EL 4.1) |
| 107 | Glycoprotein | KILSAFNTV | HLA-A*02:06 | 9 | 0.844148 | 0.06 | -0.00455 | 98.48% (65/66) | Non-Toxin | IEDB recommended 2020.09 (NetMHCpan EL 4.1) |
| 108 | Glycoprotein | KINEGLLDSK | HLA-A*03:01 | 10 | 0.944226 | 0.02 | 0.03247 | 98.48% (65/66) | Non-Toxin | IEDB recommended 2020.09 (NetMHCpan EL 4.1) |
| 109 | Glycoprotein | KINEGLLDSK | HLA-A*11:01 | 10 | 0.736511 | 0.11 | 0.03247 | 98.48% (65/66) | Non-Toxin | IEDB recommended 2020.09 (NetMHCpan EL 4.1) |
| 110 | Glycoprotein | KIYDSLGQPV | HLA-A*02:03 | 10 | 0.726573 | 0.09 | -0.22312 | 96.97% (64/66) | Non-Toxin | IEDB recommended 2020.09 (NetMHCpan EL 4.1) |
| 111 | Glycoprotein | KIYDSLGQPVF | HLA-A*32:01 | 11 | 0.530682 | 0.09 | -0.21206 | 96.97% (64/66) | Non-Toxin | IEDB recommended 2020.09 (NetMHCpan EL 4.1) |
| 112 | Glycoprotein | KLISYTLPV | HLA-A*02:01 | 9 | 0.871446 | 0.05 | -0.10617 | 95.45% (63/66) | Non-Toxin | IEDB recommended 2020.09 (NetMHCpan EL 4.1) |
| 113 | Glycoprotein | KLISYTLPV | HLA-A*02:03 | 9 | 0.830418 | 0.05 | -0.10617 | 95.45% (63/66) | Non-Toxin | IEDB recommended 2020.09 (NetMHCpan EL 4.1) |
| 114 | Glycoprotein | KLISYTLPV | HLA-A*02:06 | 9 | 0.757053 | 0.09 | -0.10617 | 95.45% (63/66) | Non-Toxin | IEDB recommended 2020.09 (NetMHCpan EL 4.1) |
| 115 | Glycoprotein | KPENCRLSM | HLA-B*07:02 | 9 | 0.924779 | 0.04 | -0.08381 | 98.48% (65/66) | Non-Toxin | IEDB recommended 2020.09 (NetMHCpan EL 4.1) |
| 116 | Glycoprotein | KPKLISYTL | HLA-B*07:02 | 9 | 0.981271 | 0.01 | -0.08773 | 96.97% (64/66) | Non-Toxin | IEDB recommended 2020.09 (NetMHCpan EL 4.1) |
| 117 | Glycoprotein | KPKLISYTL | HLA-B*08:01 | 9 | 0.794497 | 0.04 | -0.08773 | 96.97% (64/66) | Non-Toxin | IEDB recommended 2020.09 (NetMHCpan EL 4.1) |

| | | | | | | | | | | |
|---|---|---|---|---|---|---|---|---|---|---|
| 118 | Glycoprotein | KPKNNGESY | HLA-B*35:01 | 9 | 0.716665 | 0.12 | -0.06307 | 71.21% (47/66) | Non-Toxin | IEDB recommended 2020.09 (NetMHCpan EL 4.1) |
| 119 | Glycoprotein | KPQTEGVSNL | HLA-B*07:02 | 10 | 0.867951 | 0.05 | 0.02742 | 75.76% (50/66) | Non-Toxin | IEDB recommended 2020.09 (NetMHCpan EL 4.1) |
| 120 | Glycoprotein | KSYYGTMDIK | HLA-A*03:01 | 10 | 0.829379 | 0.06 | -0.00294 | 100.00% (66/66) | Non-Toxin | IEDB recommended 2020.09 (NetMHCpan EL 4.1) |
| 121 | Glycoprotein | KSYYGTMDIKK | HLA-A*03:01 | 11 | 0.871061 | 0.04 | -0.09792 | 100.00% (66/66) | Non-Toxin | IEDB recommended 2020.09 (NetMHCpan EL 4.1) |
| 122 | Glycoprotein | KTITNCFLLK | HLA-A*11:01 | 10 | 0.736348 | 0.11 | 0.11782 | 98.48% (65/66) | Non-Toxin | IEDB recommended 2020.09 (NetMHCpan EL 4.1) |
| 123 | Glycoprotein | KTSNQILKPK | HLA-A*03:01 | 10 | 0.876597 | 0.04 | -0.24233 | 98.48% (65/66) | Non-Toxin | IEDB recommended 2020.09 (NetMHCpan EL 4.1) |
| 124 | Glycoprotein | KTSNQILKPK | HLA-A*11:01 | 10 | 0.839489 | 0.06 | -0.24233 | 98.48% (65/66) | Non-Toxin | IEDB recommended 2020.09 (NetMHCpan EL 4.1) |
| 125 | Glycoprotein | KVMPYGPSGIK | HLA-A*03:01 | 11 | 0.94341 | 0.02 | -0.09893 | 100.00% (66/66) | Non-Toxin | IEDB recommended 2020.09 (NetMHCpan EL 4.1) |
| 126 | Glycoprotein | KVMPYGPSGIK | HLA-A*30:01 | 11 | 0.706511 | 0.04 | -0.09893 | 100.00% (66/66) | Non-Toxin | IEDB recommended 2020.09 (NetMHCpan EL 4.1) |
| 127 | Glycoprotein | KVMPYGPSGIK | HLA-A*11:01 | 11 | 0.79294 | 0.08 | -0.09893 | 100.00% (66/66) | Non-Toxin | IEDB recommended 2020.09 (NetMHCpan EL 4.1) |
| 128 | Glycoprotein | LAMDEGYFAY | HLA-A*01:01 | 10 | 0.829827 | 0.06 | 0.214 | 90.91% (60/66) | Non-Toxin | IEDB recommended 2020.09 (NetMHCpan EL 4.1) |
| 129 | Glycoprotein | LAMDEGYFAY | HLA-B*35:01 | 10 | 0.778031 | 0.09 | 0.214 | 90.91% (60/66) | Non-Toxin | IEDB recommended 2020.09 (NetMHCpan EL 4.1) |
| 130 | Glycoprotein | LLDSKILSA | HLA-A*02:01 | 9 | 0.802868 | 0.08 | -0.35001 | 98.48% (65/66) | Non-Toxin | IEDB recommended 2020.09 (NetMHCpan EL 4.1) |
| 131 | Glycoprotein | LLKNKIWCI | HLA-B*08:01 | 9 | 0.645816 | 0.08 | -0.00579 | 98.48% (65/66) | Non-Toxin | IEDB recommended 2020.09 (NetMHCpan EL 4.1) |
| 132 | Glycoprotein | LYFPAVGFL | HLA-A*23:01 | 9 | 0.673981 | 0.09 | 0.2008 | 100.00% (66/66) | Non-Toxin | IEDB recommended 2020.09 (NetMHCpan EL 4.1) |
| 133 | Glycoprotein | LYFPAVGFL | HLA-A*24:02 | 9 | 0.670528 | 0.12 | 0.2008 | 100.00% (66/66) | Non-Toxin | IEDB recommended 2020.09 (NetMHCpan EL 4.1) |
| 134 | Glycoprotein | MIKDALQSI | HLA-A*02:03 | 9 | 0.712158 | 0.1 | -0.21444 | 71.21% (47/66) | Non-Toxin | IEDB recommended 2020.09 (NetMHCpan EL 4.1) |
| 135 | Glycoprotein | MPTESKKV | HLA-B*51:01 | 8 | 0.612667 | 0.12 | -0.43275 | 74.24% (49/66) | Non-Toxin | IEDB recommended 2020.09 (NetMHCpan EL 4.1) |
| 136 | Glycoprotein | MPTESKKVRF | HLA-B*53:01 | 10 | 0.651627 | 0.07 | -0.39567 | 74.24% (49/66) | Non-Toxin | IEDB recommended 2020.09 (NetMHCpan EL 4.1) |
| 137 | Glycoprotein | MPYGPSGI | HLA-B*51:01 | 8 | 0.838764 | 0.03 | -0.10503 | 100.00% (66/66) | Non-Toxin | IEDB recommended 2020.09 (NetMHCpan EL 4.1) |
| 138 | Glycoprotein | MTRLAVKPK | HLA-A*30:01 | 9 | 0.74319 | 0.03 | -0.10588 | 100.00% (66/66) | Non-Toxin | IEDB recommended 2020.09 (NetMHCpan EL 4.1) |
| 139 | Glycoprotein | NPSKVIKSY | HLA-B*35:01 | 9 | 0.974763 | 0.01 | -0.38388 | 80.30% (53/66) | Non-Toxin | IEDB recommended 2020.09 (NetMHCpan EL 4.1) |
| 140 | Glycoprotein | NPSKVIKSY | HLA-B*53:01 | 9 | 0.754206 | 0.05 | -0.38388 | 80.30% (53/66) | Non-Toxin | IEDB recommended 2020.09 (NetMHCpan EL 4.1) |
| 141 | Glycoprotein | NQILKPKLISY | HLA-B*15:01 | 11 | 0.731875 | 0.12 | -0.39354 | 96.97% (64/66) | Non-Toxin | IEDB recommended 2020.09 (NetMHCpan EL 4.1) |
| 142 | Glycoprotein | QPVFYQASF | HLA-B*35:01 | 9 | 0.874428 | 0.05 | -0.04508 | 100.00% (66/66) | Non-Toxin | IEDB recommended 2020.09 (NetMHCpan EL 4.1) |
| 143 | Glycoprotein | QPVFYQASF | HLA-B*53:01 | 9 | 0.567699 | 0.09 | -0.04508 | 100.00% (66/66) | Non-Toxin | IEDB recommended 2020.09 (NetMHCpan EL 4.1) |
| 144 | Glycoprotein | QPVFYQASFSW | HLA-B*53:01 | 11 | 0.745175 | 0.05 | -0.10069 | 100.00% (66/66) | Non-Toxin | IEDB recommended 2020.09 (NetMHCpan EL 4.1) |
| 145 | Glycoprotein | QTAENPVFTV | HLA-A*68:02 | 10 | 0.896524 | 0.02 | 0.25669 | 100.00% (66/66) | Non-Toxin | IEDB recommended 2020.09 (NetMHCpan EL 4.1) |
| 146 | Glycoprotein | QTVNPLVVNW | HLA-B*57:01 | 10 | 0.989017 | 0.01 | 0.05521 | 72.73% (48/66) | Non-Toxin | IEDB recommended 2020.09 (NetMHCpan EL 4.1) |
| 147 | Glycoprotein | QTVNPLVVNW | HLA-B*58:01 | 10 | 0.940343 | 0.04 | 0.05521 | 72.73% (48/66) | Non-Toxin | IEDB recommended 2020.09 (NetMHCpan EL 4.1) |
| 148 | Glycoprotein | RLSIGSPSK | HLA-A*03:01 | 9 | 0.924551 | 0.02 | -0.14853 | 98.48% (65/66) | Non-Toxin | IEDB recommended 2020.09 (NetMHCpan EL 4.1) |
| 149 | Glycoprotein | RLSIGSPSK | HLA-A*30:01 | 9 | 0.611386 | 0.07 | -0.14853 | 98.48% (65/66) | Non-Toxin | IEDB recommended 2020.09 (NetMHCpan EL 4.1) |
| 150 | Glycoprotein | RPKLFAVKI | HLA-B*07:02 | 9 | 0.817556 | 0.07 | -0.02149 | 100.00% (66/66) | Non-Toxin | IEDB recommended 2020.09 (NetMHCpan EL 4.1) |
| 151 | Glycoprotein | RPNSHYIL | HLA-B*07:02 | 8 | 0.865362 | 0.06 | -0.02823 | 95.45% (63/66) | Non-Toxin | IEDB recommended 2020.09 (NetMHCpan EL 4.1) |
| 152 | Glycoprotein | SAVYNNEFYY | HLA-A*30:02 | 10 | 0.571065 | 0.11 | 0.18797 | 90.91% (60/66) | Non-Toxin | IEDB recommended 2020.09 (NetMHCpan EL 4.1) |
| 153 | Glycoprotein | SIGSPSKIY | HLA-A*30:02 | 9 | 0.638341 | 0.08 | -0.42624 | 95.45% (63/66) | Non-Toxin | IEDB recommended 2020.09 (NetMHCpan EL 4.1) |
| 154 | Glycoprotein | SINENVNEK | HLA-A*11:01 | 9 | 0.852696 | 0.05 | 0.18425 | 100.00% (66/66) | Non-Toxin | IEDB recommended 2020.09 (NetMHCpan EL 4.1) |
| 155 | Glycoprotein | SLIDTSSTI | HLA-A*02:03 | 9 | 0.85892 | 0.04 | -0.16935 | 100.00% (66/66) | Non-Toxin | IEDB recommended 2020.09 (NetMHCpan EL 4.1) |
| 156 | Glycoprotein | SLIDTSSTI | HLA-A*02:01 | 9 | 0.724287 | 0.12 | -0.16935 | 100.00% (66/66) | Non-Toxin | IEDB recommended 2020.09 (NetMHCpan EL 4.1) |
| 157 | Glycoprotein | SMGIRPNSHY | HLA-B*15:01 | 10 | 0.75637 | 0.1 | 0.05771 | 96.97% (64/66) | Non-Toxin | IEDB recommended 2020.09 (NetMHCpan EL 4.1) |
| 158 | Glycoprotein | SPSKIYDSL | HLA-B*07:02 | 9 | 0.936711 | 0.04 | -0.22252 | 95.45% (63/66) | Non-Toxin | IEDB recommended 2020.09 (NetMHCpan EL 4.1) |
| 159 | Glycoprotein | STASINENV | HLA-A*68:02 | 9 | 0.95749 | 0.01 | 0.05046 | 100.00% (66/66) | Non-Toxin | IEDB recommended 2020.09 (NetMHCpan EL 4.1) |
| 160 | Glycoprotein | STDNQAMIK | HLA-A*11:01 | 9 | 0.798004 | 0.08 | -0.14572 | 71.21% (47/66) | Non-Toxin | IEDB recommended 2020.09 (NetMHCpan EL 4.1) |
| 161 | Glycoprotein | STITIPANI | HLA-A*68:02 | 9 | 0.852249 | 0.04 | 0.23066 | 100.00% (66/66) | Non-Toxin | IEDB recommended 2020.09 (NetMHCpan EL 4.1) |
| 162 | Glycoprotein | SYNQHQFAL | HLA-A*24:02 | 9 | 0.75696 | 0.07 | -0.07454 | 75.76% (50/66) | Non-Toxin | IEDB recommended 2020.09 (NetMHCpan EL 4.1) |
| 163 | Glycoprotein | SYNQHQFAL | HLA-A*23:01 | 9 | 0.602283 | 0.11 | -0.07454 | 75.76% (50/66) | Non-Toxin | IEDB recommended 2020.09 (NetMHCpan EL 4.1) |
| 164 | Glycoprotein | TEIGPKVSL | HLA-B*40:01 | 9 | 0.993634 | 0.01 | -0.19832 | 100.00% (66/66) | Non-Toxin | IEDB recommended 2020.09 (NetMHCpan EL 4.1) |
| 165 | Glycoprotein | TEIGPKVSL | HLA-B*44:03 | 9 | 0.881783 | 0.04 | -0.19832 | 100.00% (66/66) | Non-Toxin | IEDB recommended 2020.09 (NetMHCpan EL 4.1) |
| 166 | Glycoprotein | TEIGPKVSL | HLA-B*44:02 | 9 | 0.809345 | 0.05 | -0.19832 | 100.00% (66/66) | Non-Toxin | IEDB recommended 2020.09 (NetMHCpan EL 4.1) |
| 167 | Glycoprotein | TPSNPNTVY | HLA-B*35:01 | 9 | 0.979377 | 0.01 | -0.02022 | 75.76% (50/66) | Non-Toxin | IEDB recommended 2020.09 (NetMHCpan EL 4.1) |
| 168 | Glycoprotein | TPSNPNTVY | HLA-B*53:01 | 9 | 0.663863 | 0.07 | -0.02022 | 75.76% (50/66) | Non-Toxin | IEDB recommended 2020.09 (NetMHCpan EL 4.1) |

| | | | | | | | | | | |
|---|---|---|---|---|---|---|---|---|---|---|
| 169 | Glycoprotein | TSNQILKPK | HLA-A*11:01 | 9 | 0.771223 | 0.09 | -0.18798 | 98.48% (65/66) | Non-Toxin | IEDB recommended 2020.09 (NetMHCpan EL 4.1) |
| 170 | Glycoprotein | TSNQILKPK | HLA-A*30:01 | 9 | 0.530047 | 0.11 | -0.18798 | 98.48% (65/66) | Non-Toxin | IEDB recommended 2020.09 (NetMHCpan EL 4.1) |
| 171 | Glycoprotein | TVFKDNEVLY | HLA-A*26:01 | 10 | 0.631707 | 0.07 | -0.04109 | 75.76% (50/66) | Non-Toxin | IEDB recommended 2020.09 (NetMHCpan EL 4.1) |
| 172 | Glycoprotein | TVFKDNEVLY | HLA-A*30:02 | 10 | 0.58975 | 0.11 | -0.04109 | 75.76% (50/66) | Non-Toxin | IEDB recommended 2020.09 (NetMHCpan EL 4.1) |
| 173 | Glycoprotein | TVFKDNEVLYR | HLA-A*68:01 | 11 | 0.883557 | 0.09 | -0.03886 | 75.76% (50/66) | Non-Toxin | IEDB recommended 2020.09 (NetMHCpan EL 4.1) |
| 174 | Glycoprotein | TVNPLVVNW | HLA-B*57:01 | 9 | 0.99016 | 0.01 | 0.04586 | 92.42% (61/66) | Non-Toxin | IEDB recommended 2020.09 (NetMHCpan EL 4.1) |
| 175 | Glycoprotein | TVNPLVVNW | HLA-B*58:01 | 9 | 0.98245 | 0.01 | 0.04586 | 92.42% (61/66) | Non-Toxin | IEDB recommended 2020.09 (NetMHCpan EL 4.1) |
| 176 | Glycoprotein | TVNPLVVNW | HLA-A*32:01 | 9 | 0.882614 | 0.01 | 0.04586 | 92.42% (61/66) | Non-Toxin | IEDB recommended 2020.09 (NetMHCpan EL 4.1) |
| 177 | Glycoprotein | TVNPLVVNW | HLA-B*53:01 | 9 | 0.853695 | 0.03 | 0.04586 | 92.42% (61/66) | Non-Toxin | IEDB recommended 2020.09 (NetMHCpan EL 4.1) |
| 178 | Glycoprotein | TVNPLVVNWR | HLA-A*68:01 | 10 | 0.9277 | 0.05 | 0.17896 | 92.42% (61/66) | Non-Toxin | IEDB recommended 2020.09 (NetMHCpan EL 4.1) |
| 179 | Glycoprotein | TVYHCSAVY | HLA-A*30:02 | 9 | 0.576146 | 0.11 | -0.11974 | 100.00% (66/66) | Non-Toxin | IEDB recommended 2020.09 (NetMHCpan EL 4.1) |
| 180 | Glycoprotein | TVYHCSAVY | HLA-A*26:01 | 9 | 0.502916 | 0.11 | -0.11974 | 100.00% (66/66) | Non-Toxin | IEDB recommended 2020.09 (NetMHCpan EL 4.1) |
| 181 | Glycoprotein | TYWSGSLMM | HLA-A*23:01 | 9 | 0.630795 | 0.1 | -0.32926 | 100.00% (66/66) | Non-Toxin | IEDB recommended 2020.09 (NetMHCpan EL 4.1) |
| 182 | Glycoprotein | TYWSGSLMMTR | HLA-A*33:01 | 11 | 0.584673 | 0.11 | -0.52429 | 100.00% (66/66) | Non-Toxin | IEDB recommended 2020.09 (NetMHCpan EL 4.1) |
| 183 | Glycoprotein | VFKDNEVLY | HLA-A*30:02 | 9 | 0.688437 | 0.06 | 0.06863 | 75.76% (50/66) | Non-Toxin | IEDB recommended 2020.09 (NetMHCpan EL 4.1) |
| 184 | Glycoprotein | VFYQASFSW | HLA-A*23:01 | 9 | 0.89986 | 0.02 | -0.23325 | 100.00% (66/66) | Non-Toxin | IEDB recommended 2020.09 (NetMHCpan EL 4.1) |
| 185 | Glycoprotein | VFYQASFSW | HLA-A*24:02 | 9 | 0.84855 | 0.04 | -0.23325 | 100.00% (66/66) | Non-Toxin | IEDB recommended 2020.09 (NetMHCpan EL 4.1) |
| 186 | Glycoprotein | VFYQASFSW | HLA-A*32:01 | 9 | 0.489801 | 0.11 | -0.23325 | 100.00% (66/66) | Non-Toxin | IEDB recommended 2020.09 (NetMHCpan EL 4.1) |
| 187 | Glycoprotein | VIRPKLFAV | HLA-B*08:01 | 9 | 0.764439 | 0.05 | -0.09314 | 100.00% (66/66) | Non-Toxin | IEDB recommended 2020.09 (NetMHCpan EL 4.1) |
| 188 | Glycoprotein | VIRPKLFAV | HLA-A*02:03 | 9 | 0.692449 | 0.1 | -0.09314 | 100.00% (66/66) | Non-Toxin | IEDB recommended 2020.09 (NetMHCpan EL 4.1) |
| 189 | Glycoprotein | VPSLFMTNV | HLA-B*51:01 | 9 | 0.820342 | 0.03 | -0.08718 | 100.00% (66/66) | Non-Toxin | IEDB recommended 2020.09 (NetMHCpan EL 4.1) |
| 190 | Glycoprotein | VPSLFMTNVW | HLA-B*53:01 | 10 | 0.730907 | 0.05 | -0.06666 | 100.00% (66/66) | Non-Toxin | IEDB recommended 2020.09 (NetMHCpan EL 4.1) |
| 191 | Glycoprotein | VWTPSNPNTVY | HLA-B*35:01 | 11 | 0.709721 | 0.12 | -0.12597 | 75.76% (50/66) | Non-Toxin | IEDB recommended 2020.09 (NetMHCpan EL 4.1) |
| 192 | Glycoprotein | VYNNEFYYV | HLA-A*24:02 | 9 | 0.773742 | 0.06 | 0.19381 | 90.91% (60/66) | Non-Toxin | IEDB recommended 2020.09 (NetMHCpan EL 4.1) |
| 193 | Glycoprotein | VYNNEFYYV | HLA-A*23:01 | 9 | 0.644897 | 0.1 | 0.19381 | 90.91% (60/66) | Non-Toxin | IEDB recommended 2020.09 (NetMHCpan EL 4.1) |
| 194 | Glycoprotein | WTPSNPNTVY | HLA-B*35:01 | 10 | 0.774703 | 0.09 | -0.13638 | 75.76% (50/66) | Non-Toxin | IEDB recommended 2020.09 (NetMHCpan EL 4.1) |
| 195 | Glycoprotein | WTPSNPNTVY | HLA-A*01:01 | 10 | 0.637408 | 0.12 | -0.13638 | 75.76% (50/66) | Non-Toxin | IEDB recommended 2020.09 (NetMHCpan EL 4.1) |
| 196 | Glycoprotein | YILRSGLLK | HLA-A*03:01 | 9 | 0.759821 | 0.12 | -0.09656 | 98.48% (65/66) | Non-Toxin | IEDB recommended 2020.09 (NetMHCpan EL 4.1) |
| **S.No.** | **Protein** | **Peptide** | **Allele** | **Length** | **Score** | **Percentile rank** | **Immunogenicity** | **Conservancy** | **Toxicity** | **Method** |
| 197 | Matrix Protein | AAYPLGVGK | HLA-A*11:01 | 9 | 0.958713 | 0.01 | 0.06338 | 97.44% (38/39) | Non-Toxin | IEDB recommended 2020.09 (NetMHCpan EL 4.1) |
| 198 | Matrix Protein | AAYPLGVGK | HLA-A*03:01 | 9 | 0.903419 | 0.03 | 0.06338 | 97.44% (38/39) | Non-Toxin | IEDB recommended 2020.09 (NetMHCpan EL 4.1) |
| 199 | Matrix Protein | AAYPLGVGK | HLA-A*30:01 | 9 | 0.724233 | 0.04 | 0.06338 | 97.44% (38/39) | Non-Toxin | IEDB recommended 2020.09 (NetMHCpan EL 4.1) |
| 200 | Matrix Protein | AVLQPSIPR | HLA-A*11:01 | 9 | 0.937807 | 0.01 | -0.18085 | 38.46% (15/39) | Non-Toxin | IEDB recommended 2020.09 (NetMHCpan EL 4.1) |
| 201 | Matrix Protein | AVLQPSIPR | HLA-A*31:01 | 9 | 0.877796 | 0.04 | -0.18085 | 38.46% (15/39) | Non-Toxin | IEDB recommended 2020.09 (NetMHCpan EL 4.1) |
| 202 | Matrix Protein | DENGSMIPKY | HLA-B*44:03 | 10 | 0.869933 | 0.05 | -0.31018 | 97.44% (38/39) | Non-Toxin | IEDB recommended 2020.09 (NetMHCpan EL 4.1) |
| 203 | Matrix Protein | DENGSMIPKY | HLA-B*44:02 | 10 | 0.705537 | 0.1 | -0.31018 | 97.44% (38/39) | Non-Toxin | IEDB recommended 2020.09 (NetMHCpan EL 4.1) |
| 204 | Matrix Protein | DVFIDNTGR | HLA-A*68:01 | 9 | 0.944623 | 0.04 | 0.23999 | 94.87% (37/39) | Non-Toxin | IEDB recommended 2020.09 (NetMHCpan EL 4.1) |
| 205 | Matrix Protein | DVFIDNTGR | HLA-A*33:01 | 9 | 0.566445 | 0.12 | 0.23999 | 94.87% (37/39) | Non-Toxin | IEDB recommended 2020.09 (NetMHCpan EL 4.1) |
| 206 | Matrix Protein | ESMEGVSDF | HLA-A*26:01 | 9 | 0.750935 | 0.05 | -0.01105 | 97.44% (38/39) | Non-Toxin | IEDB recommended 2020.09 (NetMHCpan EL 4.1) |
| 207 | Matrix Protein | FSLMDINPW | HLA-B*58:01 | 9 | 0.862849 | 0.08 | -0.04536 | 94.87% (37/39) | Non-Toxin | IEDB recommended 2020.09 (NetMHCpan EL 4.1) |
| 208 | Matrix Protein | GPLNHLVPW | HLA-B*53:01 | 9 | 0.536302 | 0.11 | 0.03931 | 97.44% (38/39) | Non-Toxin | IEDB recommended 2020.09 (NetMHCpan EL 4.1) |
| 209 | Matrix Protein | HPQDLLEEL | HLA-B*35:01 | 9 | 0.910201 | 0.04 | 0.10648 | 97.44% (38/39) | Non-Toxin | IEDB recommended 2020.09 (NetMHCpan EL 4.1) |
| 210 | Matrix Protein | HPQDLLEEL | HLA-B*53:01 | 9 | 0.770931 | 0.04 | 0.10648 | 97.44% (38/39) | Non-Toxin | IEDB recommended 2020.09 (NetMHCpan EL 4.1) |
| 211 | Matrix Protein | HPQDLLEEL | HLA-B*07:02 | 9 | 0.747457 | 0.11 | 0.10648 | 97.44% (38/39) | Non-Toxin | IEDB recommended 2020.09 (NetMHCpan EL 4.1) |
| 212 | Matrix Protein | IFNAVKVCR | HLA-A*33:01 | 9 | 0.719007 | 0.06 | -0.12219 | 97.44% (38/39) | Non-Toxin | IEDB recommended 2020.09 (NetMHCpan EL 4.1) |
| 213 | Matrix Protein | IFNAVKVCR | HLA-A*31:01 | 9 | 0.749649 | 0.09 | -0.12219 | 97.44% (38/39) | Non-Toxin | IEDB recommended 2020.09 (NetMHCpan EL 4.1) |
| 214 | Matrix Protein | INPWLNRLTW | HLA-B*53:01 | 10 | 0.540939 | 0.11 | 0.26423 | 94.87% (37/39) | Non-Toxin | IEDB recommended 2020.09 (NetMHCpan EL 4.1) |
| 215 | Matrix Protein | IQLDKHQAL | HLA-B*08:01 | 9 | 0.770985 | 0.04 | -0.23573 | 97.44% (38/39) | Non-Toxin | IEDB recommended 2020.09 (NetMHCpan EL 4.1) |
| 216 | Matrix Protein | IQLDKHQAL | HLA-A*02:06 | 9 | 0.72365 | 0.1 | -0.23573 | 97.44% (38/39) | Non-Toxin | IEDB recommended 2020.09 (NetMHCpan EL 4.1) |
| 217 | Matrix Protein | IYMIPRTML | HLA-A*24:02 | 9 | 0.77957 | 0.06 | 0.045 | 97.44% (38/39) | Non-Toxin | IEDB recommended 2020.09 (NetMHCpan EL 4.1) |
| 218 | Matrix Protein | IYMIPRTML | HLA-A*23:01 | 9 | 0.702654 | 0.08 | 0.045 | 97.44% (38/39) | Non-Toxin | IEDB recommended 2020.09 (NetMHCpan EL 4.1) |

| | | | | | | | | | | |
|---|---|---|---|---|---|---|---|---|---|---|
| 219 | Matrix Protein | IYMIPRTMLEF | HLA-A*24:02 | 11 | 0.779077 | 0.06 | 0.03816 | 97.44% (38/39) | Non-Toxin | IEDB recommended 2020.09 (NetMHCpan EL 4.1) |
| 220 | Matrix Protein | IYMIPRTMLEF | HLA-A*23:01 | 11 | 0.734098 | 0.07 | 0.03816 | 97.44% (38/39) | Non-Toxin | IEDB recommended 2020.09 (NetMHCpan EL 4.1) |
| 221 | Matrix Protein | KIDRMKLQF | HLA-A*32:01 | 9 | 0.565259 | 0.07 | -0.39176 | 94.87% (37/39) | Non-Toxin | IEDB recommended 2020.09 (NetMHCpan EL 4.1) |
| 222 | Matrix Protein | KINGVISKR | HLA-A*31:01 | 9 | 0.95937 | 0.01 | -0.06814 | 94.87% (37/39) | Non-Toxin | IEDB recommended 2020.09 (NetMHCpan EL 4.1) |
| 223 | Matrix Protein | KINGVISKR | HLA-A*03:01 | 9 | 0.828358 | 0.06 | -0.06814 | 94.87% (37/39) | Non-Toxin | IEDB recommended 2020.09 (NetMHCpan EL 4.1) |
| 224 | Matrix Protein | KIYTPGANER | HLA-A*31:01 | 10 | 0.798054 | 0.06 | 0.14993 | 97.44% (38/39) | Non-Toxin | IEDB recommended 2020.09 (NetMHCpan EL 4.1) |
| 225 | Matrix Protein | KIYTPGANER | HLA-A*03:01 | 10 | 0.82045 | 0.07 | 0.14993 | 97.44% (38/39) | Non-Toxin | IEDB recommended 2020.09 (NetMHCpan EL 4.1) |
| 226 | Matrix Protein | KIYTPGANERK | HLA-A*03:01 | 11 | 0.960329 | 0.01 | 0.20681 | 97.44% (38/39) | Non-Toxin | IEDB recommended 2020.09 (NetMHCpan EL 4.1) |
| 227 | Matrix Protein | KLNDSGIYM | HLA-A*02:01 | 9 | 0.766353 | 0.09 | 0.00118 | 97.44% (38/39) | Non-Toxin | IEDB recommended 2020.09 (NetMHCpan EL 4.1) |
| 228 | Matrix Protein | KLNDSGIYM | HLA-A*02:03 | 9 | 0.659643 | 0.11 | 0.00118 | 97.44% (38/39) | Non-Toxin | IEDB recommended 2020.09 (NetMHCpan EL 4.1) |
| 229 | Matrix Protein | KVASFMLHL | HLA-A*32:01 | 9 | 0.757709 | 0.02 | -0.19553 | 79.49% (31/39) | Non-Toxin | IEDB recommended 2020.09 (NetMHCpan EL 4.1) |
| 230 | Matrix Protein | KVASFMLHL | HLA-A*02:06 | 9 | 0.802942 | 0.07 | -0.19553 | 79.49% (31/39) | Non-Toxin | IEDB recommended 2020.09 (NetMHCpan EL 4.1) |
| 231 | Matrix Protein | KYNNYMYLI | HLA-A*24:02 | 9 | 0.902303 | 0.02 | -0.18711 | 97.44% (38/39) | Non-Toxin | IEDB recommended 2020.09 (NetMHCpan EL 4.1) |
| 232 | Matrix Protein | KYNNYMYLI | HLA-A*23:01 | 9 | 0.8084 | 0.05 | -0.18711 | 97.44% (38/39) | Non-Toxin | IEDB recommended 2020.09 (NetMHCpan EL 4.1) |
| 233 | Matrix Protein | KYYSVDYCR | HLA-A*31:01 | 9 | 0.860264 | 0.04 | -0.14121 | 97.44% (38/39) | Non-Toxin | IEDB recommended 2020.09 (NetMHCpan EL 4.1) |
| 234 | Matrix Protein | KYYSVDYCRR | HLA-A*31:01 | 10 | 0.735617 | 0.1 | -0.12461 | 97.44% (38/39) | Non-Toxin | IEDB recommended 2020.09 (NetMHCpan EL 4.1) |
| 235 | Matrix Protein | LQPSIPREF | HLA-B*15:01 | 9 | 0.769308 | 0.08 | 0.05127 | 38.46% (15/39) | Non-Toxin | IEDB recommended 2020.09 (NetMHCpan EL 4.1) |
| 236 | Matrix Protein | NAIAFNLLV | HLA-B*51:01 | 9 | 0.620195 | 0.12 | 0.17464 | 84.62% (33/39) | Non-Toxin | IEDB recommended 2020.09 (NetMHCpan EL 4.1) |
| 237 | Matrix Protein | NPWLNRLTW | HLA-B*53:01 | 9 | 0.949646 | 0.01 | 0.11648 | 94.87% (37/39) | Non-Toxin | IEDB recommended 2020.09 (NetMHCpan EL 4.1) |
| 238 | Matrix Protein | QLDKHQAL | HLA-B*08:01 | 8 | 0.588707 | 0.1 | -0.25432 | 97.44% (38/39) | Non-Toxin | IEDB recommended 2020.09 (NetMHCpan EL 4.1) |
| 239 | Matrix Protein | QPSIPREFM | HLA-B*07:02 | 9 | 0.861829 | 0.06 | 0.27104 | 38.46% (15/39) | Non-Toxin | IEDB recommended 2020.09 (NetMHCpan EL 4.1) |
| 240 | Matrix Protein | RIFFLSITK | HLA-A*03:01 | 9 | 0.949438 | 0.02 | 0.12427 | 97.44% (38/39) | Non-Toxin | IEDB recommended 2020.09 (NetMHCpan EL 4.1) |
| 241 | Matrix Protein | RIFFLSITK | HLA-A*11:01 | 9 | 0.88709 | 0.03 | 0.12427 | 97.44% (38/39) | Non-Toxin | IEDB recommended 2020.09 (NetMHCpan EL 4.1) |
| 242 | Matrix Protein | RIFFLSITK | HLA-A*30:01 | 9 | 0.736092 | 0.04 | 0.12427 | 97.44% (38/39) | Non-Toxin | IEDB recommended 2020.09 (NetMHCpan EL 4.1) |
| 243 | Matrix Protein | RLFAQMGFQK | HLA-A*03:01 | 10 | 0.946666 | 0.02 | -0.14341 | 94.87% (37/39) | Non-Toxin | IEDB recommended 2020.09 (NetMHCpan EL 4.1) |
| 244 | Matrix Protein | SESMEGVSDF | HLA-B*44:02 | 10 | 0.78351 | 0.06 | -0.1877 | 97.44% (38/39) | Non-Toxin | IEDB recommended 2020.09 (NetMHCpan EL 4.1) |
| 245 | Matrix Protein | SESMEGVSDF | HLA-B*44:03 | 10 | 0.816639 | 0.07 | -0.1877 | 97.44% (38/39) | Non-Toxin | IEDB recommended 2020.09 (NetMHCpan EL 4.1) |
| 246 | Matrix Protein | SLDKDGFKV | HLA-A*02:01 | 9 | 0.823165 | 0.07 | -0.1835 | 82.05% (32/39) | Non-Toxin | IEDB recommended 2020.09 (NetMHCpan EL 4.1) |
| 247 | Matrix Protein | SLHIKINGV | HLA-A*02:03 | 9 | 0.913363 | 0.03 | 0.07404 | 94.87% (37/39) | Non-Toxin | IEDB recommended 2020.09 (NetMHCpan EL 4.1) |
| 248 | Matrix Protein | SLMDINPWL | HLA-A*02:01 | 9 | 0.967916 | 0.02 | 0.20889 | 94.87% (37/39) | Non-Toxin | IEDB recommended 2020.09 (NetMHCpan EL 4.1) |
| 249 | Matrix Protein | SLMDINPWL | HLA-A*02:06 | 9 | 0.917932 | 0.03 | 0.20889 | 94.87% (37/39) | Non-Toxin | IEDB recommended 2020.09 (NetMHCpan EL 4.1) |
| 250 | Matrix Protein | SLMDINPWL | HLA-A*02:03 | 9 | 0.778072 | 0.06 | 0.20889 | 94.87% (37/39) | Non-Toxin | IEDB recommended 2020.09 (NetMHCpan EL 4.1) |
| 251 | Matrix Protein | SMIPKYKIY | HLA-B*15:01 | 9 | 0.884702 | 0.03 | -0.28568 | 97.44% (38/39) | Non-Toxin | IEDB recommended 2020.09 (NetMHCpan EL 4.1) |
| 252 | Matrix Protein | SMIPKYKIY | HLA-A*30:02 | 9 | 0.732656 | 0.04 | -0.28568 | 97.44% (38/39) | Non-Toxin | IEDB recommended 2020.09 (NetMHCpan EL 4.1) |
| 253 | Matrix Protein | SPSSWEHGGYL | HLA-B*07:02 | 11 | 0.719483 | 0.11 | 0.18287 | 43.59% (17/39) | Non-Toxin | IEDB recommended 2020.09 (NetMHCpan EL 4.1) |
| 254 | Matrix Protein | SSGPLNHLVPW | HLA-B*57:01 | 11 | 0.983765 | 0.02 | 0.03216 | 97.44% (38/39) | Non-Toxin | IEDB recommended 2020.09 (NetMHCpan EL 4.1) |
| 255 | Matrix Protein | SSGPLNHLVPW | HLA-B*58:01 | 11 | 0.964931 | 0.03 | 0.03216 | 97.44% (38/39) | Non-Toxin | IEDB recommended 2020.09 (NetMHCpan EL 4.1) |
| 256 | Matrix Protein | VAAVLQPSI | HLA-B*51:01 | 9 | 0.761215 | 0.05 | -0.17162 | 38.46% (15/39) | Non-Toxin | IEDB recommended 2020.09 (NetMHCpan EL 4.1) |
| 257 | Matrix Protein | VSDFSPSSW | HLA-B*58:01 | 9 | 0.979107 | 0.01 | -0.28282 | 97.44% (38/39) | Non-Toxin | IEDB recommended 2020.09 (NetMHCpan EL 4.1) |
| 258 | Matrix Protein | VSDFSPSSW | HLA-B*57:01 | 9 | 0.94504 | 0.06 | -0.28282 | 97.44% (38/39) | Non-Toxin | IEDB recommended 2020.09 (NetMHCpan EL 4.1) |
| 259 | Matrix Protein | YLDKVEPEI | HLA-A*02:01 | 9 | 0.970643 | 0.02 | -0.02621 | 97.44% (38/39) | Non-Toxin | IEDB recommended 2020.09 (NetMHCpan EL 4.1) |
| 260 | Matrix Protein | YLDKVEPEI | HLA-A*02:06 | 9 | 0.901827 | 0.04 | -0.02621 | 97.44% (38/39) | Non-Toxin | IEDB recommended 2020.09 (NetMHCpan EL 4.1) |
| 261 | Matrix Protein | YMIPRTML | HLA-B*08:01 | 8 | 0.840889 | 0.03 | -0.02922 | 97.44% (38/39) | Non-Toxin | IEDB recommended 2020.09 (NetMHCpan EL 4.1) |
| 262 | Matrix Protein | YMIPRTMLEF | HLA-B*15:01 | 10 | 0.718913 | 0.12 | 0.00408 | 97.44% (38/39) | Non-Toxin | IEDB recommended 2020.09 (NetMHCpan EL 4.1) |
| **S.No.** | **Protein** | **Peptide** | **Allele** | **Length** | **Score** | **Percentile rank** | **Immunogenicity** | **Conservancy** | **Toxicity** | **Method** |
| 263 | Nucleoprotein | AAFTLISMY | HLA-A*30:02 | 9 | 0.739162 | 0.04 | -0.05068 | 100.00% (13/13) | Non-Toxin | IEDB recommended 2020.09 (NetMHCpan EL 4.1) |
| 264 | Nucleoprotein | AAFTLISMY | HLA-B*35:01 | 9 | 0.710899 | 0.12 | -0.05068 | 100.00% (13/13) | Non-Toxin | IEDB recommended 2020.09 (NetMHCpan EL 4.1) |
| 265 | Nucleoprotein | AATATLTTK | HLA-A*11:01 | 9 | 0.746321 | 0.1 | 0.13477 | 100.00% (13/13) | Non-Toxin | IEDB recommended 2020.09 (NetMHCpan EL 4.1) |
| 266 | Nucleoprotein | AEESETRRW | HLA-B*44:02 | 9 | 0.995607 | 0.01 | 0.07399 | 100.00% (13/13) | Non-Toxin | IEDB recommended 2020.09 (NetMHCpan EL 4.1) |
| 267 | Nucleoprotein | AEESETRRW | HLA-B*44:03 | 9 | 0.995024 | 0.01 | 0.07399 | 100.00% (13/13) | Non-Toxin | IEDB recommended 2020.09 (NetMHCpan EL 4.1) |
| 268 | Nucleoprotein | AESMKVGAAF | HLA-B*44:02 | 10 | 0.870437 | 0.03 | -0.31242 | 100.00% (13/13) | Non-Toxin | IEDB recommended 2020.09 (NetMHCpan EL 4.1) |

| | | | | | | | | | | |
|---|---|---|---|---|---|---|---|---|---|---|
| 269 | Nucleoprotein | AESMKVGAAF | HLA-B*44:03 | 10 | 0.874851 | 0.05 | -0.31242 | 100.00% (13/13) | Non-Toxin | IEDB recommended 2020.09 (NetMHCpan EL 4.1) |
| 270 | Nucleoprotein | APGGYPLLW | HLA-B*53:01 | 9 | 0.84362 | 0.03 | 0.01522 | 100.00% (13/13) | Non-Toxin | IEDB recommended 2020.09 (NetMHCpan EL 4.1) |
| 271 | Nucleoprotein | ASFRSYQSK | HLA-A*30:01 | 9 | 0.868071 | 0.01 | -0.26892 | 100.00% (13/13) | Non-Toxin | IEDB recommended 2020.09 (NetMHCpan EL 4.1) |
| 272 | Nucleoprotein | ASFRSYQSK | HLA-A*11:01 | 9 | 0.913021 | 0.02 | -0.26892 | 100.00% (13/13) | Non-Toxin | IEDB recommended 2020.09 (NetMHCpan EL 4.1) |
| 273 | Nucleoprotein | ASFRSYQSK | HLA-A*03:01 | 9 | 0.837943 | 0.05 | -0.26892 | 100.00% (13/13) | Non-Toxin | IEDB recommended 2020.09 (NetMHCpan EL 4.1) |
| 274 | Nucleoprotein | ATDDPAISNK | HLA-A*11:01 | 10 | 0.910367 | 0.02 | 0.0387 | 84.62% (11/13) | Non-Toxin | IEDB recommended 2020.09 (NetMHCpan EL 4.1) |
| 275 | Nucleoprotein | ATNSPELRW | HLA-B*57:01 | 9 | 0.989609 | 0.01 | -0.06424 | 100.00% (13/13) | Non-Toxin | IEDB recommended 2020.09 (NetMHCpan EL 4.1) |
| 276 | Nucleoprotein | ATNSPELRW | HLA-B*58:01 | 9 | 0.989266 | 0.01 | -0.06424 | 100.00% (13/13) | Non-Toxin | IEDB recommended 2020.09 (NetMHCpan EL 4.1) |
| 277 | Nucleoprotein | ATNSPELRW | HLA-A*32:01 | 9 | 0.742075 | 0.03 | -0.06424 | 100.00% (13/13) | Non-Toxin | IEDB recommended 2020.09 (NetMHCpan EL 4.1) |
| 278 | Nucleoprotein | DIFEEAASF | HLA-A*26:01 | 9 | 0.844817 | 0.03 | 0.20944 | 100.00% (13/13) | Non-Toxin | IEDB recommended 2020.09 (NetMHCpan EL 4.1) |
| 279 | Nucleoprotein | DTAEESETR | HLA-A*68:01 | 9 | 0.938362 | 0.04 | 0.1624 | 100.00% (13/13) | Non-Toxin | IEDB recommended 2020.09 (NetMHCpan EL 4.1) |
| 280 | Nucleoprotein | DTAEESETRR | HLA-A*68:01 | 10 | 0.898259 | 0.07 | 0.2071 | 100.00% (13/13) | Non-Toxin | IEDB recommended 2020.09 (NetMHCpan EL 4.1) |
| 281 | Nucleoprotein | EEAASFRSY | HLA-B*44:03 | 9 | 0.995846 | 0.01 | -0.05181 | 100.00% (13/13) | Non-Toxin | IEDB recommended 2020.09 (NetMHCpan EL 4.1) |
| 282 | Nucleoprotein | EEAASFRSY | HLA-B*44:02 | 9 | 0.9927 | 0.01 | -0.05181 | 100.00% (13/13) | Non-Toxin | IEDB recommended 2020.09 (NetMHCpan EL 4.1) |
| 283 | Nucleoprotein | EEMEGLMRI | HLA-B*44:03 | 9 | 0.916195 | 0.03 | -0.05165 | 100.00% (13/13) | Non-Toxin | IEDB recommended 2020.09 (NetMHCpan EL 4.1) |
| 284 | Nucleoprotein | EEMEGLMRI | HLA-B*44:02 | 9 | 0.853212 | 0.03 | -0.05165 | 100.00% (13/13) | Non-Toxin | IEDB recommended 2020.09 (NetMHCpan EL 4.1) |
| 285 | Nucleoprotein | EEMEGLMRIL | HLA-B*44:03 | 10 | 0.731993 | 0.12 | 0.02209 | 100.00% (13/13) | Non-Toxin | IEDB recommended 2020.09 (NetMHCpan EL 4.1) |
| 286 | Nucleoprotein | EESETRRW | HLA-B*44:02 | 8 | 0.849991 | 0.03 | 0.17725 | 100.00% (13/13) | Non-Toxin | IEDB recommended 2020.09 (NetMHCpan EL 4.1) |
| 287 | Nucleoprotein | EESETRRW | HLA-B*44:03 | 8 | 0.859907 | 0.05 | 0.17725 | 100.00% (13/13) | Non-Toxin | IEDB recommended 2020.09 (NetMHCpan EL 4.1) |
| 288 | Nucleoprotein | EESETRRWAKY | HLA-B*44:02 | 11 | 0.914295 | 0.02 | 0.30118 | 100.00% (13/13) | Non-Toxin | IEDB recommended 2020.09 (NetMHCpan EL 4.1) |
| 289 | Nucleoprotein | EESETRRWAKY | HLA-B*44:03 | 11 | 0.925687 | 0.03 | 0.30118 | 100.00% (13/13) | Non-Toxin | IEDB recommended 2020.09 (NetMHCpan EL 4.1) |
| 290 | Nucleoprotein | EIISDIGNY | HLA-A*26:01 | 9 | 0.990437 | 0.01 | 0.04843 | 100.00% (13/13) | Non-Toxin | IEDB recommended 2020.09 (NetMHCpan EL 4.1) |
| 291 | Nucleoprotein | EIISDIGNYV | HLA-A*68:02 | 10 | 0.858595 | 0.04 | 0.05221 | 100.00% (13/13) | Non-Toxin | IEDB recommended 2020.09 (NetMHCpan EL 4.1) |
| 292 | Nucleoprotein | ESMKVGAAF | HLA-A*26:01 | 9 | 0.767075 | 0.05 | -0.14602 | 100.00% (13/13) | Non-Toxin | IEDB recommended 2020.09 (NetMHCpan EL 4.1) |
| 293 | Nucleoprotein | FAMGVATTI | HLA-B*51:01 | 9 | 0.899183 | 0.02 | 0.10957 | 100.00% (13/13) | Non-Toxin | IEDB recommended 2020.09 (NetMHCpan EL 4.1) |
| 294 | Nucleoprotein | FAPGGYPLLW | HLA-B*53:01 | 10 | 0.841903 | 0.03 | 0.03362 | 100.00% (13/13) | Non-Toxin | IEDB recommended 2020.09 (NetMHCpan EL 4.1) |
| 295 | Nucleoprotein | FMVEILIEV | HLA-A*02:06 | 9 | 0.969999 | 0.01 | 0.40413 | 100.00% (13/13) | Non-Toxin | IEDB recommended 2020.09 (NetMHCpan EL 4.1) |
| 296 | Nucleoprotein | FMVEILIEV | HLA-A*02:01 | 9 | 0.974244 | 0.02 | 0.40413 | 100.00% (13/13) | Non-Toxin | IEDB recommended 2020.09 (NetMHCpan EL 4.1) |
| 297 | Nucleoprotein | FMVEILIEV | HLA-A*02:03 | 9 | 0.947906 | 0.02 | 0.40413 | 100.00% (13/13) | Non-Toxin | IEDB recommended 2020.09 (NetMHCpan EL 4.1) |
| 298 | Nucleoprotein | FVDSRAYGL | HLA-A*02:06 | 9 | 0.675642 | 0.12 | -0.05536 | 100.00% (13/13) | Non-Toxin | IEDB recommended 2020.09 (NetMHCpan EL 4.1) |
| 299 | Nucleoprotein | GLMRILKTA | HLA-A*02:03 | 9 | 0.739614 | 0.08 | -0.04508 | 100.00% (13/13) | Non-Toxin | IEDB recommended 2020.09 (NetMHCpan EL 4.1) |
| 300 | Nucleoprotein | GMAGFFATI | HLA-A*02:03 | 9 | 0.694943 | 0.1 | 0.3267 | 100.00% (13/13) | Non-Toxin | IEDB recommended 2020.09 (NetMHCpan EL 4.1) |
| 301 | Nucleoprotein | GPRAPYMVL | HLA-B*07:02 | 9 | 0.994488 | 0.01 | -0.08219 | 100.00% (13/13) | Non-Toxin | IEDB recommended 2020.09 (NetMHCpan EL 4.1) |
| 302 | Nucleoprotein | GPRAPYMVLL | HLA-B*07:02 | 10 | 0.915242 | 0.04 | -0.09517 | 100.00% (13/13) | Non-Toxin | IEDB recommended 2020.09 (NetMHCpan EL 4.1) |
| 303 | Nucleoprotein | GYLEPMYFR | HLA-A*31:01 | 9 | 0.841577 | 0.05 | -0.01367 | 100.00% (13/13) | Non-Toxin | IEDB recommended 2020.09 (NetMHCpan EL 4.1) |
| 304 | Nucleoprotein | GYLEPMYFR | HLA-A*33:01 | 9 | 0.659681 | 0.08 | -0.01367 | 100.00% (13/13) | Non-Toxin | IEDB recommended 2020.09 (NetMHCpan EL 4.1) |
| 305 | Nucleoprotein | GYPLLWSF | HLA-A*23:01 | 8 | 0.576849 | 0.12 | 0.04333 | 100.00% (13/13) | Non-Toxin | IEDB recommended 2020.09 (NetMHCpan EL 4.1) |
| 306 | Nucleoprotein | IGPRAPYMVL | HLA-B*07:02 | 10 | 0.88781 | 0.05 | -0.05178 | 100.00% (13/13) | Non-Toxin | IEDB recommended 2020.09 (NetMHCpan EL 4.1) |
| 307 | Nucleoprotein | IISDIGNYV | HLA-A*02:03 | 9 | 0.750805 | 0.07 | 0.1225 | 100.00% (13/13) | Non-Toxin | IEDB recommended 2020.09 (NetMHCpan EL 4.1) |
| 308 | Nucleoprotein | IISDIGNYV | HLA-A*02:06 | 9 | 0.788065 | 0.08 | 0.1225 | 100.00% (13/13) | Non-Toxin | IEDB recommended 2020.09 (NetMHCpan EL 4.1) |
| 309 | Nucleoprotein | IISDIGNYV | HLA-A*68:02 | 9 | 0.64388 | 0.11 | 0.1225 | 100.00% (13/13) | Non-Toxin | IEDB recommended 2020.09 (NetMHCpan EL 4.1) |
| 310 | Nucleoprotein | ILIAKAVTA | HLA-A*02:03 | 9 | 0.664601 | 0.11 | -0.03308 | 100.00% (13/13) | Non-Toxin | IEDB recommended 2020.09 (NetMHCpan EL 4.1) |
| 311 | Nucleoprotein | ITIEAQIW | HLA-B*58:01 | 8 | 0.878098 | 0.07 | 0.18533 | 100.00% (13/13) | Non-Toxin | IEDB recommended 2020.09 (NetMHCpan EL 4.1) |
| 312 | Nucleoprotein | KFAPGGYPLLW | HLA-A*23:01 | 11 | 0.701594 | 0.08 | 0.03766 | 100.00% (13/13) | Non-Toxin | IEDB recommended 2020.09 (NetMHCpan EL 4.1) |
| 313 | Nucleoprotein | KFAPGGYPLLW | HLA-A*24:02 | 11 | 0.712723 | 0.09 | 0.03766 | 100.00% (13/13) | Non-Toxin | IEDB recommended 2020.09 (NetMHCpan EL 4.1) |
| 314 | Nucleoprotein | KGKTPFVDSR | HLA-A*31:01 | 10 | 0.752877 | 0.09 | 0.03318 | 100.00% (13/13) | Non-Toxin | IEDB recommended 2020.09 (NetMHCpan EL 4.1) |
| 315 | Nucleoprotein | KTARDSSKGK | HLA-A*30:01 | 10 | 0.603713 | 0.08 | -0.39265 | 100.00% (13/13) | Non-Toxin | IEDB recommended 2020.09 (NetMHCpan EL 4.1) |
| 316 | Nucleoprotein | KTPFVDSRAY | HLA-A*30:02 | 10 | 0.570315 | 0.11 | 0.08681 | 100.00% (13/13) | Non-Toxin | IEDB recommended 2020.09 (NetMHCpan EL 4.1) |
| 317 | Nucleoprotein | LEESIQTKF | HLA-B*44:03 | 9 | 0.848272 | 0.06 | -0.20665 | 100.00% (13/13) | Non-Toxin | IEDB recommended 2020.09 (NetMHCpan EL 4.1) |
| 318 | Nucleoprotein | LEESIQTKF | HLA-B*44:02 | 9 | 0.768804 | 0.07 | -0.20665 | 100.00% (13/13) | Non-Toxin | IEDB recommended 2020.09 (NetMHCpan EL 4.1) |
| 319 | Nucleoprotein | MVNGIPVMER | HLA-A*68:01 | 10 | 0.956768 | 0.03 | 0.09996 | 100.00% (13/13) | Non-Toxin | IEDB recommended 2020.09 (NetMHCpan EL 4.1) |

| | | | | | | | | | | |
|---|---|---|---|---|---|---|---|---|---|---|
| 320 | Nucleoprotein | MVNGIPVMER | HLA-A*31:01 | 10 | 0.742197 | 0.1 | 0.09996 | 100.00% (13/13) | Non-Toxin | IEDB recommended 2020.09 (NetMHCpan EL 4.1) |
| 321 | Nucleoprotein | MVNGIPVMER | HLA-A*33:01 | 10 | 0.619777 | 0.1 | 0.09996 | 100.00% (13/13) | Non-Toxin | IEDB recommended 2020.09 (NetMHCpan EL 4.1) |
| 322 | Nucleoprotein | MVNGIPVMERR | HLA-A*68:01 | 11 | 0.865518 | 0.12 | 0.14044 | 100.00% (13/13) | Non-Toxin | IEDB recommended 2020.09 (NetMHCpan EL 4.1) |
| 323 | Nucleoprotein | NLLSQSLSV | HLA-A*02:01 | 9 | 0.847652 | 0.06 | -0.54462 | 100.00% (13/13) | Non-Toxin | IEDB recommended 2020.09 (NetMHCpan EL 4.1) |
| 324 | Nucleoprotein | NPFFALTQQW | HLA-B*53:01 | 10 | 0.875745 | 0.03 | 0.0542 | 100.00% (13/13) | Non-Toxin | IEDB recommended 2020.09 (NetMHCpan EL 4.1) |
| 325 | Nucleoprotein | NRGYLEPMYFR | HLA-A*33:01 | 11 | 0.589758 | 0.11 | -0.01684 | 100.00% (13/13) | Non-Toxin | IEDB recommended 2020.09 (NetMHCpan EL 4.1) |
| 326 | Nucleoprotein | NTIKSLMLL | HLA-A*68:02 | 9 | 0.719062 | 0.08 | -0.50002 | 100.00% (13/13) | Non-Toxin | IEDB recommended 2020.09 (NetMHCpan EL 4.1) |
| 327 | Nucleoprotein | NTIKSLMLLY | HLA-A*26:01 | 10 | 0.560325 | 0.09 | -0.52684 | 100.00% (13/13) | Non-Toxin | IEDB recommended 2020.09 (NetMHCpan EL 4.1) |
| 328 | Nucleoprotein | PFFALTQQW | HLA-A*23:01 | 9 | 0.643026 | 0.1 | -0.06233 | 100.00% (13/13) | Non-Toxin | IEDB recommended 2020.09 (NetMHCpan EL 4.1) |
| 329 | Nucleoprotein | QQKRVNPFF | HLA-B*15:01 | 9 | 0.826914 | 0.05 | 0.07523 | 100.00% (13/13) | Non-Toxin | IEDB recommended 2020.09 (NetMHCpan EL 4.1) |
| 330 | Nucleoprotein | QTKFAPGGY | HLA-A*26:01 | 9 | 0.541353 | 0.1 | 0.12386 | 100.00% (13/13) | Non-Toxin | IEDB recommended 2020.09 (NetMHCpan EL 4.1) |
| 331 | Nucleoprotein | QVAELAAAV | HLA-A*68:02 | 9 | 0.920491 | 0.02 | 0.19536 | 100.00% (13/13) | Non-Toxin | IEDB recommended 2020.09 (NetMHCpan EL 4.1) |
| 332 | Nucleoprotein | REAKFAAGGVL | HLA-B*40:01 | 11 | 0.772258 | 0.12 | 0.07052 | 100.00% (13/13) | Non-Toxin | IEDB recommended 2020.09 (NetMHCpan EL 4.1) |
| 333 | Nucleoprotein | REIGPRAPY | HLA-B*44:02 | 9 | 0.837864 | 0.04 | 0.14176 | 100.00% (13/13) | Non-Toxin | IEDB recommended 2020.09 (NetMHCpan EL 4.1) |
| 334 | Nucleoprotein | REIGPRAPY | HLA-B*44:03 | 9 | 0.872258 | 0.05 | 0.14176 | 100.00% (13/13) | Non-Toxin | IEDB recommended 2020.09 (NetMHCpan EL 4.1) |
| 335 | Nucleoprotein | REMSISSL | HLA-B*40:01 | 8 | 0.822674 | 0.1 | -0.38922 | 100.00% (13/13) | Non-Toxin | IEDB recommended 2020.09 (NetMHCpan EL 4.1) |
| 336 | Nucleoprotein | RGYLEPMYFR | HLA-A*31:01 | 10 | 0.864046 | 0.04 | -0.02568 | 100.00% (13/13) | Non-Toxin | IEDB recommended 2020.09 (NetMHCpan EL 4.1) |
| 337 | Nucleoprotein | RLTNSLLNLR | HLA-A*31:01 | 10 | 0.77221 | 0.08 | -0.18819 | 100.00% (13/13) | Non-Toxin | IEDB recommended 2020.09 (NetMHCpan EL 4.1) |
| 338 | Nucleoprotein | RPGALIRSL | HLA-B*07:02 | 9 | 0.981392 | 0.01 | 0.11187 | 100.00% (13/13) | Non-Toxin | IEDB recommended 2020.09 (NetMHCpan EL 4.1) |
| 339 | Nucleoprotein | RPGALIRSLL | HLA-B*07:02 | 10 | 0.925615 | 0.04 | 0.07179 | 100.00% (13/13) | Non-Toxin | IEDB recommended 2020.09 (NetMHCpan EL 4.1) |
| 340 | Nucleoprotein | RSMGALNINR | HLA-A*31:01 | 10 | 0.841767 | 0.05 | 0.10685 | 100.00% (13/13) | Non-Toxin | IEDB recommended 2020.09 (NetMHCpan EL 4.1) |
| 341 | Nucleoprotein | RSRLAAKAAK | HLA-A*30:01 | 10 | 0.602552 | 0.08 | -0.06528 | 100.00% (13/13) | Non-Toxin | IEDB recommended 2020.09 (NetMHCpan EL 4.1) |
| 342 | Nucleoprotein | RSYQSKLGR | HLA-A*31:01 | 9 | 0.944357 | 0.01 | -0.47142 | 100.00% (13/13) | Non-Toxin | IEDB recommended 2020.09 (NetMHCpan EL 4.1) |
| 343 | Nucleoprotein | RSYQSKLGR | HLA-A*30:01 | 9 | 0.585278 | 0.09 | -0.47142 | 100.00% (13/13) | Non-Toxin | IEDB recommended 2020.09 (NetMHCpan EL 4.1) |
| 344 | Nucleoprotein | RSYQSKLGR | HLA-A*03:01 | 9 | 0.775679 | 0.11 | -0.47142 | 100.00% (13/13) | Non-Toxin | IEDB recommended 2020.09 (NetMHCpan EL 4.1) |
| 345 | Nucleoprotein | RYPALALNEF | HLA-A*24:02 | 10 | 0.937849 | 0.02 | 0.10567 | 100.00% (13/13) | Non-Toxin | IEDB recommended 2020.09 (NetMHCpan EL 4.1) |
| 346 | Nucleoprotein | RYPALALNEF | HLA-A*23:01 | 10 | 0.866903 | 0.03 | 0.10567 | 100.00% (13/13) | Non-Toxin | IEDB recommended 2020.09 (NetMHCpan EL 4.1) |
| 347 | Nucleoprotein | SAKGRAVEI | HLA-B*08:01 | 9 | 0.720362 | 0.06 | 0.14467 | 100.00% (13/13) | Non-Toxin | IEDB recommended 2020.09 (NetMHCpan EL 4.1) |
| 348 | Nucleoprotein | SETRRWAKY | HLA-B*44:02 | 9 | 0.874815 | 0.03 | 0.23061 | 100.00% (13/13) | Non-Toxin | IEDB recommended 2020.09 (NetMHCpan EL 4.1) |
| 349 | Nucleoprotein | SETRRWAKY | HLA-B*44:03 | 9 | 0.870731 | 0.05 | 0.23061 | 100.00% (13/13) | Non-Toxin | IEDB recommended 2020.09 (NetMHCpan EL 4.1) |
| 350 | Nucleoprotein | SGRQSVTFK | HLA-A*30:01 | 9 | 0.7602 | 0.02 | -0.12084 | 100.00% (13/13) | Non-Toxin | IEDB recommended 2020.09 (NetMHCpan EL 4.1) |
| 351 | Nucleoprotein | SLADSVPSSSV | HLA-A*02:03 | 11 | 0.912606 | 0.03 | -0.48869 | 100.00% (13/13) | Non-Toxin | IEDB recommended 2020.09 (NetMHCpan EL 4.1) |
| 352 | Nucleoprotein | SLADSVPSSSV | HLA-A*02:01 | 11 | 0.906387 | 0.03 | -0.48869 | 100.00% (13/13) | Non-Toxin | IEDB recommended 2020.09 (NetMHCpan EL 4.1) |
| 353 | Nucleoprotein | SLADSVPSSSV | HLA-A*02:06 | 11 | 0.725435 | 0.1 | -0.48869 | 100.00% (13/13) | Non-Toxin | IEDB recommended 2020.09 (NetMHCpan EL 4.1) |
| 354 | Nucleoprotein | SLLNDPDIEAV | HLA-A*02:01 | 11 | 0.787782 | 0.09 | 0.25493 | 100.00% (13/13) | Non-Toxin | IEDB recommended 2020.09 (NetMHCpan EL 4.1) |
| 355 | Nucleoprotein | SPELRWEL | HLA-B*08:01 | 8 | 0.548582 | 0.12 | 0.36475 | 100.00% (13/13) | Non-Toxin | IEDB recommended 2020.09 (NetMHCpan EL 4.1) |
| 356 | Nucleoprotein | SQSLSVRKF | HLA-B*15:01 | 9 | 0.888119 | 0.03 | -0.26942 | 100.00% (13/13) | Non-Toxin | IEDB recommended 2020.09 (NetMHCpan EL 4.1) |
| 357 | Nucleoprotein | SSKGKTPFV | HLA-A*30:01 | 9 | 0.651272 | 0.06 | -0.15032 | 100.00% (13/13) | Non-Toxin | IEDB recommended 2020.09 (NetMHCpan EL 4.1) |
| 358 | Nucleoprotein | SVRKFMVEI | HLA-A*30:01 | 9 | 0.518708 | 0.12 | -0.15816 | 100.00% (13/13) | Non-Toxin | IEDB recommended 2020.09 (NetMHCpan EL 4.1) |
| 359 | Nucleoprotein | TAEESETRRW | HLA-B*44:02 | 10 | 0.924372 | 0.02 | 0.18011 | 100.00% (13/13) | Non-Toxin | IEDB recommended 2020.09 (NetMHCpan EL 4.1) |
| 360 | Nucleoprotein | TAEESETRRW | HLA-B*44:03 | 10 | 0.914083 | 0.03 | 0.18011 | 100.00% (13/13) | Non-Toxin | IEDB recommended 2020.09 (NetMHCpan EL 4.1) |
| 361 | Nucleoprotein | TIKSLMLLY | HLA-A*30:02 | 9 | 0.747027 | 0.04 | -0.42841 | 100.00% (13/13) | Non-Toxin | IEDB recommended 2020.09 (NetMHCpan EL 4.1) |
| 362 | Nucleoprotein | TIKSLMLLY | HLA-A*26:01 | 9 | 0.708049 | 0.06 | -0.42841 | 100.00% (13/13) | Non-Toxin | IEDB recommended 2020.09 (NetMHCpan EL 4.1) |
| 363 | Nucleoprotein | TIRFGLETR | HLA-A*33:01 | 9 | 0.563756 | 0.12 | 0.26434 | 100.00% (13/13) | Non-Toxin | IEDB recommended 2020.09 (NetMHCpan EL 4.1) |
| 364 | Nucleoprotein | TPFVDSRAY | HLA-B*35:01 | 9 | 0.992487 | 0.01 | 0.01195 | 100.00% (13/13) | Non-Toxin | IEDB recommended 2020.09 (NetMHCpan EL 4.1) |
| 365 | Nucleoprotein | TPFVDSRAY | HLA-B*53:01 | 9 | 0.656965 | 0.07 | 0.01195 | 100.00% (13/13) | Non-Toxin | IEDB recommended 2020.09 (NetMHCpan EL 4.1) |
| 366 | Nucleoprotein | TTIDRSMGA | HLA-A*68:02 | 9 | 0.709706 | 0.08 | -0.16821 | 100.00% (13/13) | Non-Toxin | IEDB recommended 2020.09 (NetMHCpan EL 4.1) |
| 367 | Nucleoprotein | VPATNSPEL | HLA-B*07:02 | 9 | 0.931907 | 0.04 | -0.06113 | 100.00% (13/13) | Non-Toxin | IEDB recommended 2020.09 (NetMHCpan EL 4.1) |
| 368 | Nucleoprotein | VPATNSPEL | HLA-B*35:01 | 9 | 0.868157 | 0.05 | -0.06113 | 100.00% (13/13) | Non-Toxin | IEDB recommended 2020.09 (NetMHCpan EL 4.1) |
| 369 | Nucleoprotein | VPATNSPEL | HLA-B*53:01 | 9 | 0.568069 | 0.09 | -0.06113 | 100.00% (13/13) | Non-Toxin | IEDB recommended 2020.09 (NetMHCpan EL 4.1) |
| 370 | Nucleoprotein | VPATNSPEL | HLA-B*51:01 | 9 | 0.60984 | 0.12 | -0.06113 | 100.00% (13/13) | Non-Toxin | IEDB recommended 2020.09 (NetMHCpan EL 4.1) |

| | | | | | | | | | | |
|---|---|---|---|---|---|---|---|---|---|---|
| 371 | Nucleoprotein | VPATNSPELRW | HLA-B*53:01 | 11 | 0.826206 | 0.03 | -0.01131 | 100.00% (13/13) | Non-Toxin | IEDB recommended 2020.09 (NetMHCpan EL 4.1) |
| 372 | Nucleoprotein | VPATNSPELRW | HLA-B*57:01 | 11 | 0.973675 | 0.04 | -0.01131 | 100.00% (13/13) | Non-Toxin | IEDB recommended 2020.09 (NetMHCpan EL 4.1) |
| 373 | Nucleoprotein | VPATNSPELRW | HLA-B*58:01 | 11 | 0.936312 | 0.04 | -0.01131 | 100.00% (13/13) | Non-Toxin | IEDB recommended 2020.09 (NetMHCpan EL 4.1) |
| 374 | Nucleoprotein | VQQKRVNPF | HLA-B*15:01 | 9 | 0.83591 | 0.05 | -0.17728 | 100.00% (13/13) | Non-Toxin | IEDB recommended 2020.09 (NetMHCpan EL 4.1) |
| 375 | Nucleoprotein | VTFKREMSI | HLA-A*32:01 | 9 | 0.464414 | 0.12 | -0.27921 | 100.00% (13/13) | Non-Toxin | IEDB recommended 2020.09 (NetMHCpan EL 4.1) |
| 376 | Nucleoprotein | YLEPMYFRL | HLA-A*02:01 | 9 | 0.860397 | 0.05 | -0.0241 | 100.00% (13/13) | Non-Toxin | IEDB recommended 2020.09 (NetMHCpan EL 4.1) |
| 377 | Nucleoprotein | YLEPMYFRL | HLA-A*02:06 | 9 | 0.703857 | 0.11 | -0.0241 | 100.00% (13/13) | Non-Toxin | IEDB recommended 2020.09 (NetMHCpan EL 4.1) |
| 378 | Nucleoprotein | YPALALNEF | HLA-B*35:01 | 9 | 0.993417 | 0.01 | 0.08224 | 100.00% (13/13) | Non-Toxin | IEDB recommended 2020.09 (NetMHCpan EL 4.1) |
| 379 | Nucleoprotein | YPALALNEF | HLA-B*53:01 | 9 | 0.942342 | 0.01 | 0.08224 | 100.00% (13/13) | Non-Toxin | IEDB recommended 2020.09 (NetMHCpan EL 4.1) |
| 380 | Nucleoprotein | YPALALNEF | HLA-B*07:02 | 9 | 0.715817 | 0.12 | 0.08224 | 100.00% (13/13) | Non-Toxin | IEDB recommended 2020.09 (NetMHCpan EL 4.1) |
| 381 | Nucleoprotein | YPLLWSFAM | HLA-B*35:01 | 9 | 0.909678 | 0.04 | 0.16687 | 100.00% (13/13) | Non-Toxin | IEDB recommended 2020.09 (NetMHCpan EL 4.1) |
| **S.No.** | **Protein** | **Peptide** | **Allele** | **Length** | **Score** | **Percentile rank** | **Immunogenicity** | **Conservancy** | **Toxicity** | **Method** |
| 382 | Phosphoprotein | AEGSDDIQL | HLA-B*40:01 | 9 | 0.882909 | 0.07 | -0.06835 | 100.00% (27/27) | Non-Toxin | IEDB recommended 2020.09 (NetMHCpan EL 4.1) |
| 383 | Phosphoprotein | AENDEEIQEI | HLA-B*44:02 | 10 | 0.683639 | 0.11 | 0.30124 | 100.00% (27/27) | Non-Toxin | IEDB recommended 2020.09 (NetMHCpan EL 4.1) |
| 384 | Phosphoprotein | AQPPYHWSI | HLA-A*02:06 | 9 | 0.788192 | 0.08 | 0.10237 | 62.96% (17/27) | Non-Toxin | IEDB recommended 2020.09 (NetMHCpan EL 4.1) |
| 385 | Phosphoprotein | AVKEEPPQK | HLA-A*30:01 | 9 | 0.892611 | 0.01 | 0.04077 | 62.96% (17/27) | Non-Toxin | IEDB recommended 2020.09 (NetMHCpan EL 4.1) |
| 386 | Phosphoprotein | AVKEEPPQK | HLA-A*11:01 | 9 | 0.901199 | 0.02 | 0.04077 | 62.96% (17/27) | Non-Toxin | IEDB recommended 2020.09 (NetMHCpan EL 4.1) |
| 387 | Phosphoprotein | AVKEEPPQK | HLA-A*03:01 | 9 | 0.883607 | 0.04 | 0.04077 | 62.96% (17/27) | Non-Toxin | IEDB recommended 2020.09 (NetMHCpan EL 4.1) |
| 388 | Phosphoprotein | CLVSDAKMLSY | HLA-A*01:01 | 11 | 0.667959 | 0.1 | -0.57469 | 59.26% (16/27) | Non-Toxin | IEDB recommended 2020.09 (NetMHCpan EL 4.1) |
| 389 | Phosphoprotein | DAQPPYHW | HLA-B*53:01 | 8 | 0.601371 | 0.08 | -0.03574 | 62.96% (17/27) | Non-Toxin | IEDB recommended 2020.09 (NetMHCpan EL 4.1) |
| 390 | Phosphoprotein | DAQPPYHWSI | HLA-B*51:01 | 10 | 0.606259 | 0.12 | 0.05757 | 62.96% (17/27) | Non-Toxin | IEDB recommended 2020.09 (NetMHCpan EL 4.1) |
| 391 | Phosphoprotein | DILEQQSLF | HLA-A*26:01 | 9 | 0.609284 | 0.08 | -0.27079 | 62.96% (17/27) | Non-Toxin | IEDB recommended 2020.09 (NetMHCpan EL 4.1) |
| 392 | Phosphoprotein | DPAKDSPVI | HLA-B*51:01 | 9 | 0.73593 | 0.06 | -0.32367 | 55.56% (15/27) | Non-Toxin | IEDB recommended 2020.09 (NetMHCpan EL 4.1) |
| 393 | Phosphoprotein | DPVVTDVVY | HLA-B*35:01 | 9 | 0.932553 | 0.03 | 0.17258 | 100.00% (27/27) | Non-Toxin | IEDB recommended 2020.09 (NetMHCpan EL 4.1) |
| 394 | Phosphoprotein | EIPKIINKL | HLA-A*68:02 | 9 | 0.801002 | 0.05 | -0.09718 | 100.00% (27/27) | Non-Toxin | IEDB recommended 2020.09 (NetMHCpan EL 4.1) |
| 395 | Phosphoprotein | EIPKIINKL | HLA-A*26:01 | 9 | 0.591638 | 0.08 | -0.09718 | 100.00% (27/27) | Non-Toxin | IEDB recommended 2020.09 (NetMHCpan EL 4.1) |
| 396 | Phosphoprotein | EIQEIANTV | HLA-A*68:02 | 9 | 0.908477 | 0.02 | 0.2468 | 100.00% (27/27) | Non-Toxin | IEDB recommended 2020.09 (NetMHCpan EL 4.1) |
| 397 | Phosphoprotein | ELRSELIGY | HLA-A*26:01 | 9 | 0.770159 | 0.05 | 0.06951 | 96.30% (26/27) | Non-Toxin | IEDB recommended 2020.09 (NetMHCpan EL 4.1) |
| 398 | Phosphoprotein | EPYGAAVQL | HLA-B*51:01 | 9 | 0.763162 | 0.05 | 0.07499 | 100.00% (27/27) | Non-Toxin | IEDB recommended 2020.09 (NetMHCpan EL 4.1) |
| 399 | Phosphoprotein | EPYGAAVQL | HLA-B*35:01 | 9 | 0.763708 | 0.09 | 0.07499 | 100.00% (27/27) | Non-Toxin | IEDB recommended 2020.09 (NetMHCpan EL 4.1) |
| 400 | Phosphoprotein | EPYGAAVQL | HLA-B*53:01 | 9 | 0.526772 | 0.12 | 0.07499 | 100.00% (27/27) | Non-Toxin | IEDB recommended 2020.09 (NetMHCpan EL 4.1) |
| 401 | Phosphoprotein | ESIDRVLAK | HLA-A*68:01 | 9 | 0.96351 | 0.02 | 0.16828 | 100.00% (27/27) | Non-Toxin | IEDB recommended 2020.09 (NetMHCpan EL 4.1) |
| 402 | Phosphoprotein | FPHDTDRLNY | HLA-B*35:01 | 10 | 0.969478 | 0.02 | 0.1278 | 100.00% (27/27) | Non-Toxin | IEDB recommended 2020.09 (NetMHCpan EL 4.1) |
| 403 | Phosphoprotein | FPHDTDRLNY | HLA-B*53:01 | 10 | 0.768043 | 0.04 | 0.1278 | 100.00% (27/27) | Non-Toxin | IEDB recommended 2020.09 (NetMHCpan EL 4.1) |
| 404 | Phosphoprotein | FSFDNVKNF | HLA-A*26:01 | 9 | 0.498887 | 0.12 | -0.0929 | 100.00% (27/27) | Non-Toxin | IEDB recommended 2020.09 (NetMHCpan EL 4.1) |
| 405 | Phosphoprotein | FSFDNVKNFR | HLA-A*68:01 | 10 | 0.889376 | 0.08 | -0.04584 | 100.00% (27/27) | Non-Toxin | IEDB recommended 2020.09 (NetMHCpan EL 4.1) |
| 406 | Phosphoprotein | FTSSPERGW | HLA-B*58:01 | 9 | 0.966072 | 0.02 | -0.07324 | 100.00% (27/27) | Non-Toxin | IEDB recommended 2020.09 (NetMHCpan EL 4.1) |
| 407 | Phosphoprotein | FTSSPERGW | HLA-B*57:01 | 9 | 0.970202 | 0.04 | -0.07324 | 100.00% (27/27) | Non-Toxin | IEDB recommended 2020.09 (NetMHCpan EL 4.1) |
| 408 | Phosphoprotein | GLNPTAVPF | HLA-B*15:01 | 9 | 0.800904 | 0.07 | 0.08973 | 62.96% (17/27) | Non-Toxin | IEDB recommended 2020.09 (NetMHCpan EL 4.1) |
| 409 | Phosphoprotein | GPQTSRNVNL | HLA-B*07:02 | 10 | 0.865424 | 0.06 | -0.08427 | 62.96% (17/27) | Non-Toxin | IEDB recommended 2020.09 (NetMHCpan EL 4.1) |
| 410 | Phosphoprotein | GSEDPIIREL | HLA-B*40:01 | 10 | 0.90568 | 0.06 | 0.40108 | 62.96% (17/27) | Non-Toxin | IEDB recommended 2020.09 (NetMHCpan EL 4.1) |
| 411 | Phosphoprotein | GTIGKRVSNTR | HLA-A*31:01 | 11 | 0.734465 | 0.11 | -0.18061 | 100.00% (27/27) | Non-Toxin | IEDB recommended 2020.09 (NetMHCpan EL 4.1) |
| 412 | Phosphoprotein | IPIKKGTDAKY | HLA-B*35:01 | 11 | 0.715985 | 0.12 | -0.3851 | 55.56% (15/27) | Non-Toxin | IEDB recommended 2020.09 (NetMHCpan EL 4.1) |
| 413 | Phosphoprotein | IPKIINKL | HLA-B*51:01 | 8 | 0.687106 | 0.08 | 0.00543 | 100.00% (27/27) | Non-Toxin | IEDB recommended 2020.09 (NetMHCpan EL 4.1) |
| 414 | Phosphoprotein | IQLDPVVTDV | HLA-A*02:06 | 10 | 0.787286 | 0.08 | 0.1327 | 100.00% (27/27) | Non-Toxin | IEDB recommended 2020.09 (NetMHCpan EL 4.1) |
| 415 | Phosphoprotein | IQLDPVVTDVV | HLA-A*02:06 | 11 | 0.735367 | 0.1 | 0.1677 | 100.00% (27/27) | Non-Toxin | IEDB recommended 2020.09 (NetMHCpan EL 4.1) |
| 416 | Phosphoprotein | KDAQPPYHW | HLA-B*44:02 | 9 | 0.781823 | 0.06 | -0.10932 | 62.96% (17/27) | Non-Toxin | IEDB recommended 2020.09 (NetMHCpan EL 4.1) |
| 417 | Phosphoprotein | KEEPPQKRL | HLA-B*40:01 | 9 | 0.929366 | 0.05 | -0.25026 | 62.96% (17/27) | Non-Toxin | IEDB recommended 2020.09 (NetMHCpan EL 4.1) |
| 418 | Phosphoprotein | KEEPPQKRL | HLA-B*44:02 | 9 | 0.787822 | 0.05 | -0.25026 | 62.96% (17/27) | Non-Toxin | IEDB recommended 2020.09 (NetMHCpan EL 4.1) |
| 419 | Phosphoprotein | KEEPPQKRL | HLA-B*44:03 | 9 | 0.801772 | 0.08 | -0.25026 | 62.96% (17/27) | Non-Toxin | IEDB recommended 2020.09 (NetMHCpan EL 4.1) |
| 420 | Phosphoprotein | KEIPKIINKL | HLA-B*40:01 | 10 | 0.904097 | 0.06 | -0.05454 | 100.00% (27/27) | Non-Toxin | IEDB recommended 2020.09 (NetMHCpan EL 4.1) |

| | | | | | | | | | | |
|---|---|---|---|---|---|---|---|---|---|---|
| 421 | Phosphoprotein | KIINKLESI | HLA-A*02:03 | 9 | 0.806346 | 0.05 | -0.19591 | 100.00% (27/27) | Non-Toxin | IEDB recommended 2020.09 (NetMHCpan EL 4.1) |
| 422 | Phosphoprotein | KIINKLESI | HLA-A*32:01 | 9 | 0.600792 | 0.06 | -0.19591 | 100.00% (27/27) | Non-Toxin | IEDB recommended 2020.09 (NetMHCpan EL 4.1) |
| 423 | Phosphoprotein | KIINKLESI | HLA-A*02:06 | 9 | 0.814257 | 0.07 | -0.19591 | 100.00% (27/27) | Non-Toxin | IEDB recommended 2020.09 (NetMHCpan EL 4.1) |
| 424 | Phosphoprotein | KLESIDRVLAK | HLA-A*03:01 | 11 | 0.891681 | 0.03 | 0.11999 | 100.00% (27/27) | Non-Toxin | IEDB recommended 2020.09 (NetMHCpan EL 4.1) |
| 425 | Phosphoprotein | KLINLDMRL | HLA-A*02:01 | 9 | 0.902861 | 0.03 | -0.07119 | 100.00% (27/27) | Non-Toxin | IEDB recommended 2020.09 (NetMHCpan EL 4.1) |
| 426 | Phosphoprotein | KLINLDMRL | HLA-A*02:06 | 9 | 0.744941 | 0.09 | -0.07119 | 100.00% (27/27) | Non-Toxin | IEDB recommended 2020.09 (NetMHCpan EL 4.1) |
| 427 | Phosphoprotein | KMLSYAPEI | HLA-A*02:01 | 9 | 0.836092 | 0.06 | -0.0877 | 59.26% (16/27) | Non-Toxin | IEDB recommended 2020.09 (NetMHCpan EL 4.1) |
| 428 | Phosphoprotein | KMLSYAPEI | HLA-A*32:01 | 9 | 0.521775 | 0.09 | -0.0877 | 59.26% (16/27) | Non-Toxin | IEDB recommended 2020.09 (NetMHCpan EL 4.1) |
| 429 | Phosphoprotein | KNQKEIQKTY | HLA-B*15:01 | 10 | 0.948535 | 0.01 | -0.29586 | 100.00% (27/27) | Non-Toxin | IEDB recommended 2020.09 (NetMHCpan EL 4.1) |
| 430 | Phosphoprotein | KNQKEIQKTY | HLA-A*30:02 | 10 | 0.596421 | 0.11 | -0.29586 | 100.00% (27/27) | Non-Toxin | IEDB recommended 2020.09 (NetMHCpan EL 4.1) |
| 431 | Phosphoprotein | KPNESIGRTI | HLA-B*07:02 | 10 | 0.723959 | 0.11 | 0.16541 | 59.26% (16/27) | Non-Toxin | IEDB recommended 2020.09 (NetMHCpan EL 4.1) |
| 432 | Phosphoprotein | KPVIGRDIL | HLA-B*07:02 | 9 | 0.912854 | 0.04 | 0.32552 | 62.96% (17/27) | Non-Toxin | IEDB recommended 2020.09 (NetMHCpan EL 4.1) |
| 433 | Phosphoprotein | KSRGIPIKK | HLA-A*30:01 | 9 | 0.969386 | 0.01 | 0.15638 | 100.00% (27/27) | Non-Toxin | IEDB recommended 2020.09 (NetMHCpan EL 4.1) |
| 434 | Phosphoprotein | KSRGIPIKK | HLA-A*03:01 | 9 | 0.889371 | 0.04 | 0.15638 | 100.00% (27/27) | Non-Toxin | IEDB recommended 2020.09 (NetMHCpan EL 4.1) |
| 435 | Phosphoprotein | KSRGIPIKK | HLA-A*31:01 | 9 | 0.738898 | 0.1 | 0.15638 | 100.00% (27/27) | Non-Toxin | IEDB recommended 2020.09 (NetMHCpan EL 4.1) |
| 436 | Phosphoprotein | KTLIRTHIK | HLA-A*30:01 | 9 | 0.821307 | 0.02 | 0.32232 | 100.00% (27/27) | Non-Toxin | IEDB recommended 2020.09 (NetMHCpan EL 4.1) |
| 437 | Phosphoprotein | KTNTALSTI | HLA-A*32:01 | 9 | 0.51221 | 0.1 | -0.05232 | 100.00% (27/27) | Non-Toxin | IEDB recommended 2020.09 (NetMHCpan EL 4.1) |
| 438 | Phosphoprotein | KYIMPSDDF | HLA-A*24:02 | 9 | 0.820254 | 0.05 | -0.26835 | 96.30% (26/27) | Non-Toxin | IEDB recommended 2020.09 (NetMHCpan EL 4.1) |
| 439 | Phosphoprotein | KYIMPSDDF | HLA-A*23:01 | 9 | 0.684082 | 0.09 | -0.26835 | 96.30% (26/27) | Non-Toxin | IEDB recommended 2020.09 (NetMHCpan EL 4.1) |
| 440 | Phosphoprotein | LDPVVTDVVY | HLA-B*35:01 | 10 | 0.768381 | 0.09 | 0.19578 | 100.00% (27/27) | Non-Toxin | IEDB recommended 2020.09 (NetMHCpan EL 4.1) |
| 441 | Phosphoprotein | LEQQSLFSF | HLA-B*44:03 | 9 | 0.747962 | 0.1 | -0.32356 | 100.00% (27/27) | Non-Toxin | IEDB recommended 2020.09 (NetMHCpan EL 4.1) |
| 442 | Phosphoprotein | LEQQSLFSF | HLA-B*44:02 | 9 | 0.661797 | 0.12 | -0.32356 | 100.00% (27/27) | Non-Toxin | IEDB recommended 2020.09 (NetMHCpan EL 4.1) |
| 443 | Phosphoprotein | LVSDAKMLSY | HLA-A*01:01 | 10 | 0.928402 | 0.02 | -0.4746 | 59.26% (16/27) | Non-Toxin | IEDB recommended 2020.09 (NetMHCpan EL 4.1) |
| 444 | Phosphoprotein | MPKSRGIPI | HLA-B*08:01 | 9 | 0.812819 | 0.03 | -0.04833 | 100.00% (27/27) | Non-Toxin | IEDB recommended 2020.09 (NetMHCpan EL 4.1) |
| 445 | Phosphoprotein | MPKSRGIPI | HLA-B*07:02 | 9 | 0.939144 | 0.04 | -0.04833 | 100.00% (27/27) | Non-Toxin | IEDB recommended 2020.09 (NetMHCpan EL 4.1) |
| 446 | Phosphoprotein | MPSDDFSNTF | HLA-B*35:01 | 10 | 0.97685 | 0.01 | -0.03429 | 96.30% (26/27) | Non-Toxin | IEDB recommended 2020.09 (NetMHCpan EL 4.1) |
| 447 | Phosphoprotein | MPSDDFSNTF | HLA-B*53:01 | 10 | 0.926862 | 0.02 | -0.03429 | 96.30% (26/27) | Non-Toxin | IEDB recommended 2020.09 (NetMHCpan EL 4.1) |
| 448 | Phosphoprotein | MPSDDFSNTFF | HLA-B*53:01 | 11 | 0.829443 | 0.03 | 0.03819 | 96.30% (26/27) | Non-Toxin | IEDB recommended 2020.09 (NetMHCpan EL 4.1) |
| 449 | Phosphoprotein | MPSDDFSNTFF | HLA-B*35:01 | 11 | 0.811041 | 0.07 | 0.03819 | 96.30% (26/27) | Non-Toxin | IEDB recommended 2020.09 (NetMHCpan EL 4.1) |
| 450 | Phosphoprotein | NEPYGAAVQL | HLA-B*40:01 | 10 | 0.850427 | 0.08 | 0.06777 | 100.00% (27/27) | Non-Toxin | IEDB recommended 2020.09 (NetMHCpan EL 4.1) |
| 451 | Phosphoprotein | NPTAVPFTL | HLA-B*53:01 | 9 | 0.830012 | 0.03 | 0.20321 | 51.85% (14/27) | Non-Toxin | IEDB recommended 2020.09 (NetMHCpan EL 4.1) |
| 452 | Phosphoprotein | NPTAVPFTL | HLA-B*35:01 | 9 | 0.8582 | 0.06 | 0.20321 | 51.85% (14/27) | Non-Toxin | IEDB recommended 2020.09 (NetMHCpan EL 4.1) |
| 453 | Phosphoprotein | NPTAVPFTL | HLA-B*51:01 | 9 | 0.680366 | 0.08 | 0.20321 | 51.85% (14/27) | Non-Toxin | IEDB recommended 2020.09 (NetMHCpan EL 4.1) |
| 454 | Phosphoprotein | NPTAVPFTL | HLA-B*07:02 | 9 | 0.712601 | 0.12 | 0.20321 | 51.85% (14/27) | Non-Toxin | IEDB recommended 2020.09 (NetMHCpan EL 4.1) |
| 455 | Phosphoprotein | NPTAVPFTLR | HLA-A*68:01 | 10 | 0.878383 | 0.11 | 0.21785 | 51.85% (14/27) | Non-Toxin | IEDB recommended 2020.09 (NetMHCpan EL 4.1) |
| 456 | Phosphoprotein | NQKEIQKTY | HLA-B*15:01 | 9 | 0.96002 | 0.01 | -0.10801 | 100.00% (27/27) | Non-Toxin | IEDB recommended 2020.09 (NetMHCpan EL 4.1) |
| 457 | Phosphoprotein | NTFFPHDTDR | HLA-A*68:01 | 10 | 0.934647 | 0.04 | 0.2431 | 100.00% (27/27) | Non-Toxin | IEDB recommended 2020.09 (NetMHCpan EL 4.1) |
| 458 | Phosphoprotein | NVNLDSIKLY | HLA-A*26:01 | 10 | 0.542042 | 0.1 | -0.21596 | 100.00% (27/27) | Non-Toxin | IEDB recommended 2020.09 (NetMHCpan EL 4.1) |
| 459 | Phosphoprotein | QKNQKEIQKTY | HLA-B*15:01 | 11 | 0.794229 | 0.07 | -0.36992 | 100.00% (27/27) | Non-Toxin | IEDB recommended 2020.09 (NetMHCpan EL 4.1) |
| 460 | Phosphoprotein | QLDPVVTDV | HLA-A*02:01 | 9 | 0.870003 | 0.05 | 0.12082 | 100.00% (27/27) | Non-Toxin | IEDB recommended 2020.09 (NetMHCpan EL 4.1) |
| 461 | Phosphoprotein | QLDPVVTDV | HLA-A*02:06 | 9 | 0.714887 | 0.11 | 0.12082 | 100.00% (27/27) | Non-Toxin | IEDB recommended 2020.09 (NetMHCpan EL 4.1) |
| 462 | Phosphoprotein | QLDPVVTDVVY | HLA-A*01:01 | 11 | 0.83942 | 0.05 | 0.19408 | 100.00% (27/27) | Non-Toxin | IEDB recommended 2020.09 (NetMHCpan EL 4.1) |
| 463 | Phosphoprotein | QPPYHWSI | HLA-B*51:01 | 8 | 0.619756 | 0.12 | 0.09307 | 62.96% (17/27) | Non-Toxin | IEDB recommended 2020.09 (NetMHCpan EL 4.1) |
| 464 | Phosphoprotein | QPSIKDQTKAW | HLA-B*53:01 | 11 | 0.626939 | 0.07 | -0.34358 | 51.85% (14/27) | Non-Toxin | IEDB recommended 2020.09 (NetMHCpan EL 4.1) |
| 465 | Phosphoprotein | QQGKDAQPPY | HLA-B*15:01 | 10 | 0.871265 | 0.03 | -0.27118 | 59.26% (16/27) | Non-Toxin | IEDB recommended 2020.09 (NetMHCpan EL 4.1) |
| 466 | Phosphoprotein | REDLILPEL | HLA-B*40:01 | 9 | 0.980354 | 0.01 | 0.16434 | 62.96% (17/27) | Non-Toxin | IEDB recommended 2020.09 (NetMHCpan EL 4.1) |
| 467 | Phosphoprotein | RELLKENSL | HLA-B*40:01 | 9 | 0.956669 | 0.02 | -0.23263 | 62.96% (17/27) | Non-Toxin | IEDB recommended 2020.09 (NetMHCpan EL 4.1) |
| 468 | Phosphoprotein | RETDLVHL | HLA-B*40:01 | 8 | 0.804553 | 0.1 | 0.09028 | 100.00% (27/27) | Non-Toxin | IEDB recommended 2020.09 (NetMHCpan EL 4.1) |
| 469 | Phosphoprotein | RLNHIEEQV | HLA-A*02:03 | 9 | 0.755732 | 0.07 | 0.27112 | 100.00% (27/27) | Non-Toxin | IEDB recommended 2020.09 (NetMHCpan EL 4.1) |
| 470 | Phosphoprotein | RLNHIEEQV | HLA-A*02:01 | 9 | 0.811652 | 0.08 | 0.27112 | 100.00% (27/27) | Non-Toxin | IEDB recommended 2020.09 (NetMHCpan EL 4.1) |
| 471 | Phosphoprotein | RQRPGTPMPK | HLA-A*30:01 | 10 | 0.918589 | 0.01 | -0.08868 | 62.96% (17/27) | Non-Toxin | IEDB recommended 2020.09 (NetMHCpan EL 4.1) |

| | | | | | | | | | | |
|---|---|---|---|---|---|---|---|---|---|---|
| 472 | Phosphoprotein | RQRPGTPMPK | HLA-A*03:01 | 10 | 0.796456 | 0.09 | -0.08868 | 62.96% (17/27) | Non-Toxin | IEDB recommended 2020.09 (NetMHCpan EL 4.1) |
| 473 | Phosphoprotein | RTIEGQSIR | HLA-A*31:01 | 9 | 0.856405 | 0.04 | 0.00605 | 59.26% (16/27) | Non-Toxin | IEDB recommended 2020.09 (NetMHCpan EL 4.1) |
| 474 | Phosphoprotein | RVLAKTNTA | HLA-A*30:01 | 9 | 0.524121 | 0.12 | -0.12047 | 100.00% (27/27) | Non-Toxin | IEDB recommended 2020.09 (NetMHCpan EL 4.1) |
| 475 | Phosphoprotein | SDAKMLSY | HLA-A*01:01 | 8 | 0.630839 | 0.12 | -0.52536 | 59.26% (16/27) | Non-Toxin | IEDB recommended 2020.09 (NetMHCpan EL 4.1) |
| 476 | Phosphoprotein | SEDPIIREL | HLA-B*40:01 | 9 | 0.994468 | 0.01 | 0.3531 | 62.96% (17/27) | Non-Toxin | IEDB recommended 2020.09 (NetMHCpan EL 4.1) |
| 477 | Phosphoprotein | SEDPIIREL | HLA-B*44:03 | 9 | 0.974423 | 0.01 | 0.3531 | 62.96% (17/27) | Non-Toxin | IEDB recommended 2020.09 (NetMHCpan EL 4.1) |
| 478 | Phosphoprotein | SEDPIIREL | HLA-B*44:02 | 9 | 0.959815 | 0.01 | 0.3531 | 62.96% (17/27) | Non-Toxin | IEDB recommended 2020.09 (NetMHCpan EL 4.1) |
| 479 | Phosphoprotein | SEDPIIRELL | HLA-B*40:01 | 10 | 0.925972 | 0.05 | 0.38198 | 62.96% (17/27) | Non-Toxin | IEDB recommended 2020.09 (NetMHCpan EL 4.1) |
| 480 | Phosphoprotein | SEDPIIRELL | HLA-B*44:02 | 10 | 0.705619 | 0.1 | 0.38198 | 62.96% (17/27) | Non-Toxin | IEDB recommended 2020.09 (NetMHCpan EL 4.1) |
| 481 | Phosphoprotein | SEDPIIRELL | HLA-B*44:03 | 10 | 0.743206 | 0.11 | 0.38198 | 62.96% (17/27) | Non-Toxin | IEDB recommended 2020.09 (NetMHCpan EL 4.1) |
| 482 | Phosphoprotein | SFDNVKNFR | HLA-A*33:01 | 9 | 0.608098 | 0.11 | -0.09917 | 100.00% (27/27) | Non-Toxin | IEDB recommended 2020.09 (NetMHCpan EL 4.1) |
| 483 | Phosphoprotein | SIRDNLQAK | HLA-A*30:01 | 9 | 0.814193 | 0.02 | -0.05252 | 51.85% (14/27) | Non-Toxin | IEDB recommended 2020.09 (NetMHCpan EL 4.1) |
| 484 | Phosphoprotein | SPVIAEHYY | HLA-B*35:01 | 9 | 0.967986 | 0.02 | 0.30481 | 96.30% (26/27) | Non-Toxin | IEDB recommended 2020.09 (NetMHCpan EL 4.1) |
| 485 | Phosphoprotein | SPVIAEHYY | HLA-B*53:01 | 9 | 0.694604 | 0.06 | 0.30481 | 96.30% (26/27) | Non-Toxin | IEDB recommended 2020.09 (NetMHCpan EL 4.1) |
| 486 | Phosphoprotein | SSIQQPSIK | HLA-A*11:01 | 9 | 0.782261 | 0.09 | -0.25846 | 100.00% (27/27) | Non-Toxin | IEDB recommended 2020.09 (NetMHCpan EL 4.1) |
| 487 | Phosphoprotein | SSRDVIKTL | HLA-A*30:01 | 9 | 0.548679 | 0.1 | 0.04528 | 51.85% (14/27) | Non-Toxin | IEDB recommended 2020.09 (NetMHCpan EL 4.1) |
| 488 | Phosphoprotein | SSSEVIVGI | HLA-A*68:02 | 9 | 0.833703 | 0.04 | 0.26717 | 100.00% (27/27) | Non-Toxin | IEDB recommended 2020.09 (NetMHCpan EL 4.1) |
| 489 | Phosphoprotein | STIEGHLVSM | HLA-A*26:01 | 10 | 0.612154 | 0.08 | 0.13619 | 100.00% (27/27) | Non-Toxin | IEDB recommended 2020.09 (NetMHCpan EL 4.1) |
| 490 | Phosphoprotein | STSPTDGTIGK | HLA-A*11:01 | 11 | 0.936819 | 0.01 | 0.1962 | 62.96% (17/27) | Non-Toxin | IEDB recommended 2020.09 (NetMHCpan EL 4.1) |
| 491 | Phosphoprotein | TFFPHDTDR | HLA-A*33:01 | 9 | 0.623034 | 0.1 | 0.12494 | 100.00% (27/27) | Non-Toxin | IEDB recommended 2020.09 (NetMHCpan EL 4.1) |
| 492 | Phosphoprotein | TPMPKSRGI | HLA-B*07:02 | 9 | 0.892711 | 0.05 | -0.37041 | 100.00% (27/27) | Non-Toxin | IEDB recommended 2020.09 (NetMHCpan EL 4.1) |
| 493 | Phosphoprotein | TPMPKSRGI | HLA-B*51:01 | 9 | 0.709065 | 0.07 | -0.37041 | 100.00% (27/27) | Non-Toxin | IEDB recommended 2020.09 (NetMHCpan EL 4.1) |
| 494 | Phosphoprotein | TPMPKSRGI | HLA-B*08:01 | 9 | 0.547576 | 0.12 | -0.37041 | 100.00% (27/27) | Non-Toxin | IEDB recommended 2020.09 (NetMHCpan EL 4.1) |
| 495 | Phosphoprotein | VLMGVINSI | HLA-A*02:03 | 9 | 0.891655 | 0.03 | 0.04046 | 100.00% (27/27) | Non-Toxin | IEDB recommended 2020.09 (NetMHCpan EL 4.1) |
| 496 | Phosphoprotein | VLMGVINSI | HLA-A*02:01 | 9 | 0.889812 | 0.04 | 0.04046 | 100.00% (27/27) | Non-Toxin | IEDB recommended 2020.09 (NetMHCpan EL 4.1) |
| 497 | Phosphoprotein | VLMGVINSI | HLA-A*02:06 | 9 | 0.786218 | 0.08 | 0.04046 | 100.00% (27/27) | Non-Toxin | IEDB recommended 2020.09 (NetMHCpan EL 4.1) |
| 498 | Phosphoprotein | VLMGVINSI | HLA-A*32:01 | 9 | 0.499192 | 0.1 | 0.04046 | 100.00% (27/27) | Non-Toxin | IEDB recommended 2020.09 (NetMHCpan EL 4.1) |
| 499 | Phosphoprotein | VSDAKMLSY | HLA-A*01:01 | 9 | 0.994971 | 0.01 | -0.43475 | 59.26% (16/27) | Non-Toxin | IEDB recommended 2020.09 (NetMHCpan EL 4.1) |
| 500 | Phosphoprotein | VSDAKMLSY | HLA-A*30:02 | 9 | 0.699815 | 0.06 | -0.43475 | 59.26% (16/27) | Non-Toxin | IEDB recommended 2020.09 (NetMHCpan EL 4.1) |
| 501 | Phosphoprotein | YGFTSSPERGW | HLA-B*57:01 | 11 | 0.915052 | 0.09 | -0.09821 | 100.00% (27/27) | Non-Toxin | IEDB recommended 2020.09 (NetMHCpan EL 4.1) |
| 502 | Phosphoprotein | YPSAGTENV | HLA-B*51:01 | 9 | 0.630068 | 0.11 | 0.13593 | 92.59% (25/27) | Non-Toxin | IEDB recommended 2020.09 (NetMHCpan EL 4.1) |
| **S.No.** | **Protein** | **Peptide** | **Allele** | **Length** | **Score** | **Percentile rank** | **Immunogenicity** | **Conservancy** | **Toxicity** | **Method** |
| 503 | V Protein | AEGSDDIQL | HLA-B*40:01 | 9 | 0.882909 | 0.07 | -0.06835 | 94.74% (54/57) | Non-Toxin | IEDB recommended 2020.09 (NetMHCpan EL 4.1) |
| 504 | V Protein | AQPPYHWSI | HLA-A*02:06 | 9 | 0.788192 | 0.08 | 0.10237 | 19.30% (11/57) | Non-Toxin | IEDB recommended 2020.09 (NetMHCpan EL 4.1) |
| 505 | V Protein | AVKEEPPQK | HLA-A*30:01 | 9 | 0.892611 | 0.01 | 0.04077 | 19.30% (11/57) | Non-Toxin | IEDB recommended 2020.09 (NetMHCpan EL 4.1) |
| 506 | V Protein | AVKEEPPQK | HLA-A*11:01 | 9 | 0.901199 | 0.02 | 0.04077 | 19.30% (11/57) | Non-Toxin | IEDB recommended 2020.09 (NetMHCpan EL 4.1) |
| 507 | V Protein | AVKEEPPQK | HLA-A*03:01 | 9 | 0.883607 | 0.04 | 0.04077 | 19.30% (11/57) | Non-Toxin | IEDB recommended 2020.09 (NetMHCpan EL 4.1) |
| 508 | V Protein | CLVSDAKMLSY | HLA-A*01:01 | 11 | 0.667959 | 0.1 | -0.57469 | 19.30% (11/57) | Non-Toxin | IEDB recommended 2020.09 (NetMHCpan EL 4.1) |
| 509 | V Protein | DAQPPYHW | HLA-B*53:01 | 8 | 0.601371 | 0.08 | -0.03574 | 19.30% (11/57) | Non-Toxin | IEDB recommended 2020.09 (NetMHCpan EL 4.1) |
| 510 | V Protein | DAQPPYHWSI | HLA-B*51:01 | 10 | 0.606259 | 0.12 | 0.05757 | 19.30% (11/57) | Non-Toxin | IEDB recommended 2020.09 (NetMHCpan EL 4.1) |
| 511 | V Protein | DPAKDSPVI | HLA-B*51:01 | 9 | 0.73593 | 0.06 | -0.32367 | 17.54% (10/57) | Non-Toxin | IEDB recommended 2020.09 (NetMHCpan EL 4.1) |
| 512 | V Protein | DPVVTDVVY | HLA-B*35:01 | 9 | 0.932553 | 0.03 | 0.17258 | 98.25% (56/57) | Non-Toxin | IEDB recommended 2020.09 (NetMHCpan EL 4.1) |
| 513 | V Protein | FTSSPERGW | HLA-B*58:01 | 9 | 0.966072 | 0.02 | -0.07324 | 100.00% (57/57) | Non-Toxin | IEDB recommended 2020.09 (NetMHCpan EL 4.1) |
| 514 | V Protein | FTSSPERGW | HLA-B*57:01 | 9 | 0.970202 | 0.04 | -0.07324 | 100.00% (57/57) | Non-Toxin | IEDB recommended 2020.09 (NetMHCpan EL 4.1) |
| 515 | V Protein | GLNPTAVPF | HLA-B*15:01 | 9 | 0.800904 | 0.07 | 0.08973 | 19.30% (11/57) | Non-Toxin | IEDB recommended 2020.09 (NetMHCpan EL 4.1) |
| 516 | V Protein | GPQTSRNVNL | HLA-B*07:02 | 10 | 0.865424 | 0.06 | -0.08427 | 19.30% (11/57) | Non-Toxin | IEDB recommended 2020.09 (NetMHCpan EL 4.1) |
| 517 | V Protein | GSEDPIIREL | HLA-B*40:01 | 10 | 0.90568 | 0.06 | 0.40108 | 19.30% (11/57) | Non-Toxin | IEDB recommended 2020.09 (NetMHCpan EL 4.1) |
| 518 | V Protein | GTIGKRVSNTR | HLA-A*31:01 | 11 | 0.734465 | 0.11 | -0.18061 | 100.00% (57/57) | Non-Toxin | IEDB recommended 2020.09 (NetMHCpan EL 4.1) |
| 519 | V Protein | IQLDPVVTDV | HLA-A*02:06 | 10 | 0.787286 | 0.08 | 0.1327 | 94.74% (54/57) | Non-Toxin | IEDB recommended 2020.09 (NetMHCpan EL 4.1) |
| 520 | V Protein | IQLDPVVTDVV | HLA-A*02:06 | 11 | 0.735367 | 0.1 | 0.1677 | 94.74% (54/57) | Non-Toxin | IEDB recommended 2020.09 (NetMHCpan EL 4.1) |
| 521 | V Protein | ISICWDGKRAW | HLA-B*57:01 | 11 | 0.974216 | 0.04 | 0.12279 | 87.72% (50/57) | Non-Toxin | IEDB recommended 2020.09 (NetMHCpan EL 4.1) |

| | | | | | | | | | | |
|---|---|---|---|---|---|---|---|---|---|---|
| 522 | V Protein | ISICWDGKRAW | HLA-B*58:01 | 11 | 0.85944 | 0.08 | 0.12279 | 87.72% (50/57) | Non-Toxin | IEDB recommended 2020.09 (NetMHCpan EL 4.1) |
| 523 | V Protein | KDAQPPYHW | HLA-B*44:02 | 9 | 0.781823 | 0.06 | -0.10932 | 19.30% (11/57) | Non-Toxin | IEDB recommended 2020.09 (NetMHCpan EL 4.1) |
| 524 | V Protein | KEEPPQKRL | HLA-B*40:01 | 9 | 0.929366 | 0.05 | -0.25026 | 19.30% (11/57) | Non-Toxin | IEDB recommended 2020.09 (NetMHCpan EL 4.1) |
| 525 | V Protein | KEEPPQKRL | HLA-B*44:02 | 9 | 0.787822 | 0.05 | -0.25026 | 19.30% (11/57) | Non-Toxin | IEDB recommended 2020.09 (NetMHCpan EL 4.1) |
| 526 | V Protein | KEEPPQKRL | HLA-B*44:03 | 9 | 0.801772 | 0.08 | -0.25026 | 19.30% (11/57) | Non-Toxin | IEDB recommended 2020.09 (NetMHCpan EL 4.1) |
| 527 | V Protein | KMLSYAPEI | HLA-A*02:01 | 9 | 0.836092 | 0.06 | -0.0877 | 19.30% (11/57) | Non-Toxin | IEDB recommended 2020.09 (NetMHCpan EL 4.1) |
| 528 | V Protein | KMLSYAPEI | HLA-A*32:01 | 9 | 0.521775 | 0.09 | -0.0877 | 19.30% (11/57) | Non-Toxin | IEDB recommended 2020.09 (NetMHCpan EL 4.1) |
| 529 | V Protein | KNQKEIQKTY | HLA-B*15:01 | 10 | 0.948535 | 0.01 | -0.29586 | 100.00% (57/57) | Non-Toxin | IEDB recommended 2020.09 (NetMHCpan EL 4.1) |
| 530 | V Protein | KNQKEIQKTY | HLA-A*30:02 | 10 | 0.596421 | 0.11 | -0.29586 | 100.00% (57/57) | Non-Toxin | IEDB recommended 2020.09 (NetMHCpan EL 4.1) |
| 531 | V Protein | KPNESIGRTI | HLA-B*07:02 | 10 | 0.723959 | 0.11 | 0.16541 | 19.30% (11/57) | Non-Toxin | IEDB recommended 2020.09 (NetMHCpan EL 4.1) |
| 532 | V Protein | KSRGIPIKK | HLA-A*30:01 | 9 | 0.969386 | 0.01 | 0.15638 | 100.00% (57/57) | Non-Toxin | IEDB recommended 2020.09 (NetMHCpan EL 4.1) |
| 533 | V Protein | KSRGIPIKK | HLA-A*03:01 | 9 | 0.889371 | 0.04 | 0.15638 | 100.00% (57/57) | Non-Toxin | IEDB recommended 2020.09 (NetMHCpan EL 4.1) |
| 534 | V Protein | KSRGIPIKK | HLA-A*31:01 | 9 | 0.738898 | 0.1 | 0.15638 | 100.00% (57/57) | Non-Toxin | IEDB recommended 2020.09 (NetMHCpan EL 4.1) |
| 535 | V Protein | LDPVVTDVVY | HLA-B*35:01 | 10 | 0.768381 | 0.09 | 0.19578 | 98.25% (56/57) | Non-Toxin | IEDB recommended 2020.09 (NetMHCpan EL 4.1) |
| 536 | V Protein | LVSDAKMLSY | HLA-A*01:01 | 10 | 0.928402 | 0.02 | -0.4746 | 19.30% (11/57) | Non-Toxin | IEDB recommended 2020.09 (NetMHCpan EL 4.1) |
| 537 | V Protein | MPKSRGIPI | HLA-B*08:01 | 9 | 0.812819 | 0.03 | -0.04833 | 100.00% (57/57) | Non-Toxin | IEDB recommended 2020.09 (NetMHCpan EL 4.1) |
| 538 | V Protein | MPKSRGIPI | HLA-B*07:02 | 9 | 0.939144 | 0.04 | -0.04833 | 100.00% (57/57) | Non-Toxin | IEDB recommended 2020.09 (NetMHCpan EL 4.1) |
| 539 | V Protein | NPTAVPFTL | HLA-B*53:01 | 9 | 0.830012 | 0.03 | 0.20321 | 19.30% (11/57) | Non-Toxin | IEDB recommended 2020.09 (NetMHCpan EL 4.1) |
| 540 | V Protein | NPTAVPFTL | HLA-B*35:01 | 9 | 0.8582 | 0.06 | 0.20321 | 19.30% (11/57) | Non-Toxin | IEDB recommended 2020.09 (NetMHCpan EL 4.1) |
| 541 | V Protein | NPTAVPFTL | HLA-B*51:01 | 9 | 0.680366 | 0.08 | 0.20321 | 19.30% (11/57) | Non-Toxin | IEDB recommended 2020.09 (NetMHCpan EL 4.1) |
| 542 | V Protein | NPTAVPFTL | HLA-B*07:02 | 9 | 0.712601 | 0.12 | 0.20321 | 19.30% (11/57) | Non-Toxin | IEDB recommended 2020.09 (NetMHCpan EL 4.1) |
| 543 | V Protein | NPTAVPFTLR | HLA-A*68:01 | 10 | 0.878383 | 0.11 | 0.21785 | 19.30% (11/57) | Non-Toxin | IEDB recommended 2020.09 (NetMHCpan EL 4.1) |
| 544 | V Protein | NQKEIQKTY | HLA-B*15:01 | 9 | 0.96002 | 0.01 | -0.10801 | 100.00% (57/57) | Non-Toxin | IEDB recommended 2020.09 (NetMHCpan EL 4.1) |
| 545 | V Protein | NVNLDSIKLY | HLA-A*26:01 | 10 | 0.542042 | 0.1 | -0.21596 | 100.00% (57/57) | Non-Toxin | IEDB recommended 2020.09 (NetMHCpan EL 4.1) |
| 546 | V Protein | QKNQKEIQKTY | HLA-B*15:01 | 11 | 0.794229 | 0.07 | -0.36992 | 100.00% (57/57) | Non-Toxin | IEDB recommended 2020.09 (NetMHCpan EL 4.1) |
| 547 | V Protein | QLDPVVTDV | HLA-A*02:01 | 9 | 0.870003 | 0.05 | 0.12082 | 100.00% (57/57) | Non-Toxin | IEDB recommended 2020.09 (NetMHCpan EL 4.1) |
| 548 | V Protein | QLDPVVTDV | HLA-A*02:06 | 9 | 0.714887 | 0.11 | 0.12082 | 100.00% (57/57) | Non-Toxin | IEDB recommended 2020.09 (NetMHCpan EL 4.1) |
| 549 | V Protein | QLDPVVTDVVY | HLA-A*01:01 | 11 | 0.83942 | 0.05 | 0.19408 | 98.25% (56/57) | Non-Toxin | IEDB recommended 2020.09 (NetMHCpan EL 4.1) |
| 550 | V Protein | QPPYHWSI | HLA-B*51:01 | 8 | 0.619756 | 0.12 | 0.09307 | 19.30% (11/57) | Non-Toxin | IEDB recommended 2020.09 (NetMHCpan EL 4.1) |
| 551 | V Protein | QPSIKDQTKAW | HLA-B*53:01 | 11 | 0.626939 | 0.07 | -0.34358 | 19.30% (11/57) | Non-Toxin | IEDB recommended 2020.09 (NetMHCpan EL 4.1) |
| 552 | V Protein | QQGKDAQPPY | HLA-B*15:01 | 10 | 0.871265 | 0.03 | -0.27118 | 19.30% (11/57) | Non-Toxin | IEDB recommended 2020.09 (NetMHCpan EL 4.1) |
| 553 | V Protein | RELLKENSL | HLA-B*40:01 | 9 | 0.956669 | 0.02 | -0.23263 | 19.30% (11/57) | Non-Toxin | IEDB recommended 2020.09 (NetMHCpan EL 4.1) |
| 554 | V Protein | RETDLVHL | HLA-B*40:01 | 8 | 0.804553 | 0.1 | 0.09028 | 98.25% (56/57) | Non-Toxin | IEDB recommended 2020.09 (NetMHCpan EL 4.1) |
| 555 | V Protein | RQRPGTPMPK | HLA-A*30:01 | 10 | 0.918589 | 0.01 | -0.08868 | 19.30% (11/57) | Non-Toxin | IEDB recommended 2020.09 (NetMHCpan EL 4.1) |
| 556 | V Protein | RQRPGTPMPK | HLA-A*03:01 | 10 | 0.796456 | 0.09 | -0.08868 | 19.30% (11/57) | Non-Toxin | IEDB recommended 2020.09 (NetMHCpan EL 4.1) |
| 557 | V Protein | RTIEGQSIR | HLA-A*31:01 | 9 | 0.856405 | 0.04 | 0.00605 | 19.30% (11/57) | Non-Toxin | IEDB recommended 2020.09 (NetMHCpan EL 4.1) |
| 558 | V Protein | SDAKMLSY | HLA-A*01:01 | 8 | 0.630839 | 0.12 | -0.52536 | 19.30% (11/57) | Non-Toxin | IEDB recommended 2020.09 (NetMHCpan EL 4.1) |
| 559 | V Protein | SEDPIIREL | HLA-B*40:01 | 9 | 0.994468 | 0.01 | 0.3531 | 19.30% (11/57) | Non-Toxin | IEDB recommended 2020.09 (NetMHCpan EL 4.1) |
| 560 | V Protein | SEDPIIREL | HLA-B*44:03 | 9 | 0.974423 | 0.01 | 0.3531 | 19.30% (11/57) | Non-Toxin | IEDB recommended 2020.09 (NetMHCpan EL 4.1) |
| 561 | V Protein | SEDPIIREL | HLA-B*44:02 | 9 | 0.959815 | 0.01 | 0.3531 | 19.30% (11/57) | Non-Toxin | IEDB recommended 2020.09 (NetMHCpan EL 4.1) |
| 562 | V Protein | SEDPIIRELL | HLA-B*40:01 | 10 | 0.925972 | 0.05 | 0.38198 | 19.30% (11/57) | Non-Toxin | IEDB recommended 2020.09 (NetMHCpan EL 4.1) |
| 563 | V Protein | SEDPIIRELL | HLA-B*44:02 | 10 | 0.705619 | 0.1 | 0.38198 | 19.30% (11/57) | Non-Toxin | IEDB recommended 2020.09 (NetMHCpan EL 4.1) |
| 564 | V Protein | SEDPIIRELL | HLA-B*44:03 | 10 | 0.743206 | 0.11 | 0.38198 | 19.30% (11/57) | Non-Toxin | IEDB recommended 2020.09 (NetMHCpan EL 4.1) |
| 565 | V Protein | SIRDNLQAK | HLA-A*30:01 | 9 | 0.814193 | 0.02 | -0.05252 | 19.30% (11/57) | Non-Toxin | IEDB recommended 2020.09 (NetMHCpan EL 4.1) |
| 566 | V Protein | SPVIAEHYY | HLA-B*35:01 | 9 | 0.967986 | 0.02 | 0.30481 | 98.25% (56/57) | Non-Toxin | IEDB recommended 2020.09 (NetMHCpan EL 4.1) |
| 567 | V Protein | SPVIAEHYY | HLA-B*53:01 | 9 | 0.694604 | 0.06 | 0.30481 | 98.25% (56/57) | Non-Toxin | IEDB recommended 2020.09 (NetMHCpan EL 4.1) |
| 568 | V Protein | SSIQQPSIK | HLA-A*11:01 | 9 | 0.782261 | 0.09 | -0.25846 | 100.00% (57/57) | Non-Toxin | IEDB recommended 2020.09 (NetMHCpan EL 4.1) |
| 569 | V Protein | SSSEVIVGI | HLA-A*68:02 | 9 | 0.833703 | 0.04 | 0.26717 | 98.25% (56/57) | Non-Toxin | IEDB recommended 2020.09 (NetMHCpan EL 4.1) |
| 570 | V Protein | STSPTDGTIGK | HLA-A*11:01 | 11 | 0.936819 | 0.01 | 0.1962 | 100.00% (57/57) | Non-Toxin | IEDB recommended 2020.09 (NetMHCpan EL 4.1) |
| 571 | V Protein | TPMPKSRGI | HLA-B*07:02 | 9 | 0.892711 | 0.05 | -0.37041 | 100.00% (57/57) | Non-Toxin | IEDB recommended 2020.09 (NetMHCpan EL 4.1) |
| 572 | V Protein | TPMPKSRGI | HLA-B*51:01 | 9 | 0.709065 | 0.07 | -0.37041 | 100.00% (57/57) | Non-Toxin | IEDB recommended 2020.09 (NetMHCpan EL 4.1) |

| | | | | | | | | | | |
|---|---|---|---|---|---|---|---|---|---|---|
| 573 | V Protein | TPMPKSRGI | HLA-B*08:01 | 9 | 0.547576 | 0.12 | -0.37041 | 100.00% (57/57) | Non-Toxin | IEDB recommended 2020.09 (NetMHCpan EL 4.1) |
| 574 | V Protein | VSDAKMLSY | HLA-A*01:01 | 9 | 0.994971 | 0.01 | -0.43475 | 19.30% (11/57) | Non-Toxin | IEDB recommended 2020.09 (NetMHCpan EL 4.1) |
| 575 | V Protein | VSDAKMLSY | HLA-A*30:02 | 9 | 0.699815 | 0.06 | -0.43475 | 19.30% (11/57) | Non-Toxin | IEDB recommended 2020.09 (NetMHCpan EL 4.1) |
| 576 | V Protein | YGFTSSPERGW | HLA-B*57:01 | 11 | 0.915052 | 0.09 | -0.09821 | 100.00% (57/57) | Non-Toxin | IEDB recommended 2020.09 (NetMHCpan EL 4.1) |
| **S.No.** | **Protein** | **Peptide** | **Allele** | **Length** | **Score** | **Percentile rank** | **Immunogenicity** | **Conservancy** | **Toxicity** | **Method** |
| 577 | W Protein | AEGSDDIQL | HLA-B*40:01 | 9 | 0.882909 | 0.07 | -0.06835 | 100.00% (39/39) | Non-Toxin | IEDB recommended 2020.09 (NetMHCpan EL 4.1) |
| 578 | W Protein | AVKEKSPQK | HLA-A*30:01 | 9 | 0.945499 | 0.01 | -0.41202 | 79.49% (31/39) | Non-Toxin | IEDB recommended 2020.09 (NetMHCpan EL 4.1) |
| 579 | W Protein | AVKEKSPQK | HLA-A*03:01 | 9 | 0.928687 | 0.02 | -0.41202 | 79.49% (31/39) | Non-Toxin | IEDB recommended 2020.09 (NetMHCpan EL 4.1) |
| 580 | W Protein | AVKEKSPQK | HLA-A*11:01 | 9 | 0.835471 | 0.06 | -0.41202 | 79.49% (31/39) | Non-Toxin | IEDB recommended 2020.09 (NetMHCpan EL 4.1) |
| 581 | W Protein | CLVSDAKVLSY | HLA-A*01:01 | 11 | 0.72848 | 0.09 | -0.37053 | 76.92% (30/39) | Non-Toxin | IEDB recommended 2020.09 (NetMHCpan EL 4.1) |
| 582 | W Protein | DPVVTDVVY | HLA-B*35:01 | 9 | 0.932553 | 0.03 | 0.17258 | 97.44% (38/39) | Non-Toxin | IEDB recommended 2020.09 (NetMHCpan EL 4.1) |
| 583 | W Protein | FTSSPERGW | HLA-B*58:01 | 9 | 0.966072 | 0.02 | -0.07324 | 100.00% (39/39) | Non-Toxin | IEDB recommended 2020.09 (NetMHCpan EL 4.1) |
| 584 | W Protein | FTSSPERGW | HLA-B*57:01 | 9 | 0.970202 | 0.04 | -0.07324 | 100.00% (39/39) | Non-Toxin | IEDB recommended 2020.09 (NetMHCpan EL 4.1) |
| 585 | W Protein | GLNPTAIPF | HLA-B*15:01 | 9 | 0.761814 | 0.09 | 0.16721 | 79.49% (31/39) | Non-Toxin | IEDB recommended 2020.09 (NetMHCpan EL 4.1) |
| 586 | W Protein | GMFEDHPPTK | HLA-A*03:01 | 10 | 0.923007 | 0.02 | 0.19473 | 100.00% (39/39) | Non-Toxin | IEDB recommended 2020.09 (NetMHCpan EL 4.1) |
| 587 | W Protein | GMFEDHPPTKK | HLA-A*03:01 | 11 | 0.930908 | 0.02 | 0.07737 | 100.00% (39/39) | Non-Toxin | IEDB recommended 2020.09 (NetMHCpan EL 4.1) |
| 588 | W Protein | GSEDPIIQEL | HLA-B*40:01 | 10 | 0.955951 | 0.03 | 0.25964 | 79.49% (31/39) | Non-Toxin | IEDB recommended 2020.09 (NetMHCpan EL 4.1) |
| 589 | W Protein | GTIGKRVSNTR | HLA-A*31:01 | 11 | 0.734465 | 0.11 | -0.18061 | 100.00% (39/39) | Non-Toxin | IEDB recommended 2020.09 (NetMHCpan EL 4.1) |
| 590 | W Protein | HIIEGQSTR | HLA-A*68:01 | 9 | 0.94979 | 0.03 | -0.04903 | 33.33% (13/39) | Non-Toxin | IEDB recommended 2020.09 (NetMHCpan EL 4.1) |
| 591 | W Protein | HLLGRKTCL | HLA-B*08:01 | 9 | 0.892932 | 0.02 | -0.12084 | 92.31% (36/39) | Non-Toxin | IEDB recommended 2020.09 (NetMHCpan EL 4.1) |
| 592 | W Protein | HPPTKKARVSM | HLA-B*07:02 | 11 | 0.84177 | 0.07 | -0.35954 | 100.00% (39/39) | Non-Toxin | IEDB recommended 2020.09 (NetMHCpan EL 4.1) |
| 593 | W Protein | IPFTPKNL | HLA-B*51:01 | 8 | 0.707336 | 0.07 | -0.1422 | 79.49% (31/39) | Non-Toxin | IEDB recommended 2020.09 (NetMHCpan EL 4.1) |
| 594 | W Protein | IPFTPKNLSV | HLA-B*51:01 | 10 | 0.838223 | 0.03 | -0.25585 | 76.92% (30/39) | Non-Toxin | IEDB recommended 2020.09 (NetMHCpan EL 4.1) |
| 595 | W Protein | IQLDPVVTDV | HLA-A*02:06 | 10 | 0.787286 | 0.08 | 0.1327 | 100.00% (39/39) | Non-Toxin | IEDB recommended 2020.09 (NetMHCpan EL 4.1) |
| 596 | W Protein | IQLDPVVTDVV | HLA-A*02:06 | 11 | 0.735367 | 0.1 | 0.1677 | 100.00% (39/39) | Non-Toxin | IEDB recommended 2020.09 (NetMHCpan EL 4.1) |
| 597 | W Protein | KEKSPQKRL | HLA-B*44:02 | 9 | 0.697768 | 0.1 | -0.50807 | 79.49% (31/39) | Non-Toxin | IEDB recommended 2020.09 (NetMHCpan EL 4.1) |
| 598 | W Protein | KNQKEIQKTY | HLA-B*15:01 | 10 | 0.948535 | 0.01 | -0.29586 | 100.00% (39/39) | Non-Toxin | IEDB recommended 2020.09 (NetMHCpan EL 4.1) |
| 599 | W Protein | KNQKEIQKTY | HLA-A*30:02 | 10 | 0.596421 | 0.11 | -0.29586 | 100.00% (39/39) | Non-Toxin | IEDB recommended 2020.09 (NetMHCpan EL 4.1) |
| 600 | W Protein | KQRPGTPMPK | HLA-A*30:01 | 10 | 0.915351 | 0.01 | -0.08868 | 79.49% (31/39) | Non-Toxin | IEDB recommended 2020.09 (NetMHCpan EL 4.1) |
| 601 | W Protein | KQRPGTPMPK | HLA-A*03:01 | 10 | 0.799411 | 0.09 | -0.08868 | 79.49% (31/39) | Non-Toxin | IEDB recommended 2020.09 (NetMHCpan EL 4.1) |
| 602 | W Protein | KSRGIPIKK | HLA-A*30:01 | 9 | 0.969386 | 0.01 | 0.15638 | 100.00% (39/39) | Non-Toxin | IEDB recommended 2020.09 (NetMHCpan EL 4.1) |
| 603 | W Protein | KSRGIPIKK | HLA-A*03:01 | 9 | 0.889371 | 0.04 | 0.15638 | 100.00% (39/39) | Non-Toxin | IEDB recommended 2020.09 (NetMHCpan EL 4.1) |
| 604 | W Protein | KSRGIPIKK | HLA-A*31:01 | 9 | 0.738898 | 0.1 | 0.15638 | 100.00% (39/39) | Non-Toxin | IEDB recommended 2020.09 (NetMHCpan EL 4.1) |
| 605 | W Protein | KVLSYAPEI | HLA-A*32:01 | 9 | 0.600689 | 0.06 | -0.0877 | 76.92% (30/39) | Non-Toxin | IEDB recommended 2020.09 (NetMHCpan EL 4.1) |
| 606 | W Protein | KVLSYAPEI | HLA-A*02:06 | 9 | 0.752378 | 0.09 | -0.0877 | 76.92% (30/39) | Non-Toxin | IEDB recommended 2020.09 (NetMHCpan EL 4.1) |
| 607 | W Protein | LDPVVTDVVY | HLA-B*35:01 | 10 | 0.768381 | 0.09 | 0.19578 | 97.44% (38/39) | Non-Toxin | IEDB recommended 2020.09 (NetMHCpan EL 4.1) |
| 608 | W Protein | LVSDAKVLSY | HLA-A*01:01 | 10 | 0.949801 | 0.02 | -0.27044 | 76.92% (30/39) | Non-Toxin | IEDB recommended 2020.09 (NetMHCpan EL 4.1) |
| 609 | W Protein | MPKSRGIPI | HLA-B*08:01 | 9 | 0.812819 | 0.03 | -0.04833 | 100.00% (39/39) | Non-Toxin | IEDB recommended 2020.09 (NetMHCpan EL 4.1) |
| 610 | W Protein | MPKSRGIPI | HLA-B*07:02 | 9 | 0.939144 | 0.04 | -0.04833 | 100.00% (39/39) | Non-Toxin | IEDB recommended 2020.09 (NetMHCpan EL 4.1) |
| 611 | W Protein | NQKEIQKTY | HLA-B*15:01 | 9 | 0.96002 | 0.01 | -0.10801 | 100.00% (39/39) | Non-Toxin | IEDB recommended 2020.09 (NetMHCpan EL 4.1) |
| 612 | W Protein | NVNLDSIKLY | HLA-A*26:01 | 10 | 0.542042 | 0.1 | -0.21596 | 100.00% (39/39) | Non-Toxin | IEDB recommended 2020.09 (NetMHCpan EL 4.1) |
| 613 | W Protein | QELLKENSF | HLA-B*44:02 | 9 | 0.85788 | 0.03 | -0.23263 | 79.49% (31/39) | Non-Toxin | IEDB recommended 2020.09 (NetMHCpan EL 4.1) |
| 614 | W Protein | QELLKENSF | HLA-B*44:03 | 9 | 0.877268 | 0.04 | -0.23263 | 79.49% (31/39) | Non-Toxin | IEDB recommended 2020.09 (NetMHCpan EL 4.1) |
| 615 | W Protein | QGKDAQPLYY | HLA-A*01:01 | 10 | 0.671592 | 0.1 | -0.14434 | 79.49% (31/39) | Non-Toxin | IEDB recommended 2020.09 (NetMHCpan EL 4.1) |
| 616 | W Protein | QKNQKEIQKTY | HLA-B*15:01 | 11 | 0.794229 | 0.07 | -0.36992 | 100.00% (39/39) | Non-Toxin | IEDB recommended 2020.09 (NetMHCpan EL 4.1) |
| 617 | W Protein | QLDPVVTDV | HLA-A*02:01 | 9 | 0.870003 | 0.05 | 0.12082 | 100.00% (39/39) | Non-Toxin | IEDB recommended 2020.09 (NetMHCpan EL 4.1) |
| 618 | W Protein | QLDPVVTDV | HLA-A*02:06 | 9 | 0.714887 | 0.11 | 0.12082 | 100.00% (39/39) | Non-Toxin | IEDB recommended 2020.09 (NetMHCpan EL 4.1) |
| 619 | W Protein | QLDPVVTDVVY | HLA-A*01:01 | 11 | 0.83942 | 0.05 | 0.19408 | 97.44% (38/39) | Non-Toxin | IEDB recommended 2020.09 (NetMHCpan EL 4.1) |
| 620 | W Protein | QQGKDAQPLY | HLA-B*15:01 | 10 | 0.888736 | 0.03 | -0.27118 | 79.49% (31/39) | Non-Toxin | IEDB recommended 2020.09 (NetMHCpan EL 4.1) |
| 621 | W Protein | RETDLVHL | HLA-B*40:01 | 8 | 0.804553 | 0.1 | 0.09028 | 100.00% (39/39) | Non-Toxin | IEDB recommended 2020.09 (NetMHCpan EL 4.1) |
| 622 | W Protein | SAVKEKSPQK | HLA-A*30:01 | 10 | 0.597659 | 0.08 | -0.54887 | 76.92% (30/39) | Non-Toxin | IEDB recommended 2020.09 (NetMHCpan EL 4.1) |

| | | | | | | | | | | |
|---|---|---|---|---|---|---|---|---|---|---|
| 623 | W Protein | SDAKVLSY | HLA-A*01:01 | 8 | 0.693711 | 0.1 | -0.31416 | 76.92% (30/39) | Non-Toxin | IEDB recommended 2020.09 (NetMHCpan EL 4.1) |
| 624 | W Protein | SEDPIIQEL | HLA-B*40:01 | 9 | 0.996375 | 0.01 | 0.21166 | 79.49% (31/39) | Non-Toxin | IEDB recommended 2020.09 (NetMHCpan EL 4.1) |
| 625 | W Protein | SEDPIIQEL | HLA-B*44:03 | 9 | 0.963793 | 0.01 | 0.21166 | 79.49% (31/39) | Non-Toxin | IEDB recommended 2020.09 (NetMHCpan EL 4.1) |
| 626 | W Protein | SEDPIIQEL | HLA-B*44:02 | 9 | 0.940116 | 0.01 | 0.21166 | 79.49% (31/39) | Non-Toxin | IEDB recommended 2020.09 (NetMHCpan EL 4.1) |
| 627 | W Protein | SEDPIIQELL | HLA-B*40:01 | 10 | 0.95565 | 0.03 | 0.22422 | 79.49% (31/39) | Non-Toxin | IEDB recommended 2020.09 (NetMHCpan EL 4.1) |
| 628 | W Protein | SGSEDPIIQEL | HLA-B*40:01 | 11 | 0.854104 | 0.08 | 0.27347 | 79.49% (31/39) | Non-Toxin | IEDB recommended 2020.09 (NetMHCpan EL 4.1) |
| 629 | W Protein | SPQKRLPM | HLA-B*07:02 | 8 | 0.831128 | 0.07 | -0.224 | 74.36% (29/39) | Non-Toxin | IEDB recommended 2020.09 (NetMHCpan EL 4.1) |
| 630 | W Protein | SPQKRLPM | HLA-B*08:01 | 8 | 0.612801 | 0.09 | -0.224 | 74.36% (29/39) | Non-Toxin | IEDB recommended 2020.09 (NetMHCpan EL 4.1) |
| 631 | W Protein | SPQKRLPML | HLA-B*07:02 | 9 | 0.972555 | 0.02 | -0.3266 | 74.36% (29/39) | Non-Toxin | IEDB recommended 2020.09 (NetMHCpan EL 4.1) |
| 632 | W Protein | SPQKRLPML | HLA-B*08:01 | 9 | 0.897326 | 0.02 | -0.3266 | 74.36% (29/39) | Non-Toxin | IEDB recommended 2020.09 (NetMHCpan EL 4.1) |
| 633 | W Protein | SPVIAEHYY | HLA-B*35:01 | 9 | 0.967986 | 0.02 | 0.30481 | 100.00% (39/39) | Non-Toxin | IEDB recommended 2020.09 (NetMHCpan EL 4.1) |
| 634 | W Protein | SPVIAEHYY | HLA-B*53:01 | 9 | 0.694604 | 0.06 | 0.30481 | 100.00% (39/39) | Non-Toxin | IEDB recommended 2020.09 (NetMHCpan EL 4.1) |
| 635 | W Protein | SQQGKDAQPLY | HLA-B*15:01 | 11 | 0.87658 | 0.03 | -0.27868 | 79.49% (31/39) | Non-Toxin | IEDB recommended 2020.09 (NetMHCpan EL 4.1) |
| 636 | W Protein | SQQGKDAQPLY | HLA-A*30:02 | 11 | 0.650536 | 0.08 | -0.27868 | 79.49% (31/39) | Non-Toxin | IEDB recommended 2020.09 (NetMHCpan EL 4.1) |
| 637 | W Protein | SSIQQPSIK | HLA-A*11:01 | 9 | 0.782261 | 0.09 | -0.25846 | 100.00% (39/39) | Non-Toxin | IEDB recommended 2020.09 (NetMHCpan EL 4.1) |
| 638 | W Protein | SSSEVIVGI | HLA-A*68:02 | 9 | 0.833703 | 0.04 | 0.26717 | 97.44% (38/39) | Non-Toxin | IEDB recommended 2020.09 (NetMHCpan EL 4.1) |
| 639 | W Protein | STRDSLQIK | HLA-A*30:01 | 9 | 0.935075 | 0.01 | -0.15242 | 71.79% (28/39) | Non-Toxin | IEDB recommended 2020.09 (NetMHCpan EL 4.1) |
| 640 | W Protein | STRDSLQIK | HLA-A*11:01 | 9 | 0.789457 | 0.08 | -0.15242 | 71.79% (28/39) | Non-Toxin | IEDB recommended 2020.09 (NetMHCpan EL 4.1) |
| 641 | W Protein | STSPTDGTIGK | HLA-A*11:01 | 11 | 0.936819 | 0.01 | 0.1962 | 100.00% (39/39) | Non-Toxin | IEDB recommended 2020.09 (NetMHCpan EL 4.1) |
| 642 | W Protein | TPMPKSRGI | HLA-B*07:02 | 9 | 0.892711 | 0.05 | -0.37041 | 100.00% (39/39) | Non-Toxin | IEDB recommended 2020.09 (NetMHCpan EL 4.1) |
| 643 | W Protein | TPMPKSRGI | HLA-B*51:01 | 9 | 0.709065 | 0.07 | -0.37041 | 100.00% (39/39) | Non-Toxin | IEDB recommended 2020.09 (NetMHCpan EL 4.1) |
| 644 | W Protein | VPAKDSPVI | HLA-B*51:01 | 9 | 0.838774 | 0.03 | -0.32367 | 74.36% (29/39) | Non-Toxin | IEDB recommended 2020.09 (NetMHCpan EL 4.1) |
| 645 | W Protein | VSDAKVLSY | HLA-A*01:01 | 9 | 0.996544 | 0.01 | -0.23059 | 76.92% (30/39) | Non-Toxin | IEDB recommended 2020.09 (NetMHCpan EL 4.1) |
| 646 | W Protein | VSDAKVLSY | HLA-A*30:02 | 9 | 0.751839 | 0.04 | -0.23059 | 76.92% (30/39) | Non-Toxin | IEDB recommended 2020.09 (NetMHCpan EL 4.1) |
| 647 | W Protein | YGFTSSPERGW | HLA-B*57:01 | 11 | 0.915052 | 0.09 | -0.09821 | 100.00% (39/39) | Non-Toxin | IEDB recommended 2020.09 (NetMHCpan EL 4.1) |
| **S.No.** | **Protein** | **Peptide** | **Allele** | **Length** | **Score** | **Percentile rank** | **Immunogenicity** | **Conservancy** | **Toxicity** | **Method** |
| 648 | Fusion Protein | AIGPPVFTDK | HLA-A*11:01 | 10 | 0.883408 | 0.03 | 0.18516 | 97.30% (36/37) | Non-Toxin | IEDB recommended 2020.09 (NetMHCpan EL 4.1) |
| 649 | Fusion Protein | AIGPPVFTDK | HLA-A*03:01 | 10 | 0.860799 | 0.05 | 0.18516 | 97.30% (36/37) | Non-Toxin | IEDB recommended 2020.09 (NetMHCpan EL 4.1) |
| 650 | Fusion Protein | ALYEAMKNA | HLA-A*02:03 | 9 | 0.941496 | 0.02 | -0.21343 | 100.00% (37/37) | Non-Toxin | IEDB recommended 2020.09 (NetMHCpan EL 4.1) |
| 651 | Fusion Protein | ALYEAMKNA | HLA-A*02:01 | 9 | 0.790189 | 0.09 | -0.21343 | 100.00% (37/37) | Non-Toxin | IEDB recommended 2020.09 (NetMHCpan EL 4.1) |
| 652 | Fusion Protein | ALYEAMKNA | HLA-A*02:06 | 9 | 0.713829 | 0.11 | -0.21343 | 100.00% (37/37) | Non-Toxin | IEDB recommended 2020.09 (NetMHCpan EL 4.1) |
| 653 | Fusion Protein | AQITAGVAL | HLA-B*15:01 | 9 | 0.809091 | 0.06 | 0.20996 | 97.30% (36/37) | Non-Toxin | IEDB recommended 2020.09 (NetMHCpan EL 4.1) |
| 654 | Fusion Protein | AQITAGVALY | HLA-B*15:01 | 10 | 0.964206 | 0.01 | 0.21876 | 97.30% (36/37) | Non-Toxin | IEDB recommended 2020.09 (NetMHCpan EL 4.1) |
| 655 | Fusion Protein | AQITAGVALY | HLA-A*30:02 | 10 | 0.598327 | 0.11 | 0.21876 | 97.30% (36/37) | Non-Toxin | IEDB recommended 2020.09 (NetMHCpan EL 4.1) |
| 656 | Fusion Protein | AYIQELLPVSF | HLA-A*23:01 | 11 | 0.869392 | 0.03 | -0.06972 | 100.00% (37/37) | Non-Toxin | IEDB recommended 2020.09 (NetMHCpan EL 4.1) |
| 657 | Fusion Protein | AYIQELLPVSF | HLA-A*24:02 | 11 | 0.86702 | 0.04 | -0.06972 | 100.00% (37/37) | Non-Toxin | IEDB recommended 2020.09 (NetMHCpan EL 4.1) |
| 658 | Fusion Protein | DISSQISSM | HLA-A*26:01 | 9 | 0.75497 | 0.05 | -0.44397 | 97.30% (36/37) | Non-Toxin | IEDB recommended 2020.09 (NetMHCpan EL 4.1) |
| 659 | Fusion Protein | DPVSNSMTI | HLA-B*51:01 | 9 | 0.836921 | 0.03 | -0.44062 | 97.30% (36/37) | Non-Toxin | IEDB recommended 2020.09 (NetMHCpan EL 4.1) |
| 660 | Fusion Protein | DSITGQIIY | HLA-A*26:01 | 9 | 0.644882 | 0.07 | 0.1963 | 100.00% (37/37) | Non-Toxin | IEDB recommended 2020.09 (NetMHCpan EL 4.1) |
| 661 | Fusion Protein | DSITGQIIY | HLA-B*35:01 | 9 | 0.728264 | 0.11 | 0.1963 | 100.00% (37/37) | Non-Toxin | IEDB recommended 2020.09 (NetMHCpan EL 4.1) |
| 662 | Fusion Protein | DSITGQIIYV | HLA-A*68:02 | 10 | 0.642853 | 0.11 | 0.2379 | 100.00% (37/37) | Non-Toxin | IEDB recommended 2020.09 (NetMHCpan EL 4.1) |
| 663 | Fusion Protein | DTVNPSLISM | HLA-A*26:01 | 10 | 0.635248 | 0.07 | -0.15979 | 100.00% (37/37) | Non-Toxin | IEDB recommended 2020.09 (NetMHCpan EL 4.1) |
| 664 | Fusion Protein | DYATPMTNNMR | HLA-A*33:01 | 11 | 0.638953 | 0.09 | -0.20639 | 97.30% (36/37) | Non-Toxin | IEDB recommended 2020.09 (NetMHCpan EL 4.1) |
| 665 | Fusion Protein | ELVVSSHVPR | HLA-A*68:01 | 10 | 0.920002 | 0.06 | -0.20845 | 100.00% (37/37) | Non-Toxin | IEDB recommended 2020.09 (NetMHCpan EL 4.1) |
| 666 | Fusion Protein | ESDSITGQIIY | HLA-A*01:01 | 11 | 0.964183 | 0.01 | 0.12217 | 100.00% (37/37) | Non-Toxin | IEDB recommended 2020.09 (NetMHCpan EL 4.1) |
| 667 | Fusion Protein | ESTNEAVVKL | HLA-A*68:02 | 10 | 0.71031 | 0.08 | 0.08939 | 97.30% (36/37) | Non-Toxin | IEDB recommended 2020.09 (NetMHCpan EL 4.1) |
| 668 | Fusion Protein | ETAEKTVY | HLA-A*26:01 | 8 | 0.556129 | 0.09 | -0.02517 | 100.00% (37/37) | Non-Toxin | IEDB recommended 2020.09 (NetMHCpan EL 4.1) |
| 669 | Fusion Protein | ETAEKTVYV | HLA-A*68:02 | 9 | 0.994144 | 0.01 | -0.02733 | 100.00% (37/37) | Non-Toxin | IEDB recommended 2020.09 (NetMHCpan EL 4.1) |
| 670 | Fusion Protein | ETAEKTVYV | HLA-A*26:01 | 9 | 0.644069 | 0.07 | -0.02733 | 100.00% (37/37) | Non-Toxin | IEDB recommended 2020.09 (NetMHCpan EL 4.1) |
| 671 | Fusion Protein | ETAEKTVYVL | HLA-A*68:02 | 10 | 0.902807 | 0.02 | 0.00111 | 100.00% (37/37) | Non-Toxin | IEDB recommended 2020.09 (NetMHCpan EL 4.1) |
| 672 | Fusion Protein | ETLLRTLGY | HLA-A*26:01 | 9 | 0.85457 | 0.03 | 0.08262 | 97.30% (36/37) | Non-Toxin | IEDB recommended 2020.09 (NetMHCpan EL 4.1) |

| | | | | | | | | | | |
|---|---|---|---|---|---|---|---|---|---|---|
| 673 | Fusion Protein | EWISIVPNF | HLA-A*23:01 | 9 | 0.872759 | 0.03 | 0.03205 | 97.30% (36/37) | Non-Toxin | IEDB recommended 2020.09 (NetMHCpan EL 4.1) |
| 674 | Fusion Protein | EWISIVPNF | HLA-A*24:02 | 9 | 0.836855 | 0.04 | 0.03205 | 97.30% (36/37) | Non-Toxin | IEDB recommended 2020.09 (NetMHCpan EL 4.1) |
| 675 | Fusion Protein | FALSNGVLF | HLA-B*35:01 | 9 | 0.806076 | 0.08 | -0.11611 | 100.00% (37/37) | Non-Toxin | IEDB recommended 2020.09 (NetMHCpan EL 4.1) |
| 676 | Fusion Protein | FALSNGVLF | HLA-B*53:01 | 9 | 0.587275 | 0.09 | -0.11611 | 100.00% (37/37) | Non-Toxin | IEDB recommended 2020.09 (NetMHCpan EL 4.1) |
| 677 | Fusion Protein | FILVRNTL | HLA-B*08:01 | 8 | 0.715157 | 0.06 | 0.11501 | 100.00% (37/37) | Non-Toxin | IEDB recommended 2020.09 (NetMHCpan EL 4.1) |
| 678 | Fusion Protein | FPILTEIQQAY | HLA-B*35:01 | 11 | 0.988425 | 0.01 | 0.113 | 100.00% (37/37) | Non-Toxin | IEDB recommended 2020.09 (NetMHCpan EL 4.1) |
| 679 | Fusion Protein | FPILTEIQQAY | HLA-B*53:01 | 11 | 0.738909 | 0.05 | 0.113 | 100.00% (37/37) | Non-Toxin | IEDB recommended 2020.09 (NetMHCpan EL 4.1) |
| 680 | Fusion Protein | GLVKGVTRK | HLA-A*03:01 | 9 | 0.811953 | 0.08 | -0.06874 | 43.24% (16/37) | Non-Toxin | IEDB recommended 2020.09 (NetMHCpan EL 4.1) |
| 681 | Fusion Protein | GQIIYVDLSSY | HLA-B*15:01 | 11 | 0.850268 | 0.04 | -0.0114 | 45.95% (17/37) | Non-Toxin | IEDB recommended 2020.09 (NetMHCpan EL 4.1) |
| 682 | Fusion Protein | IIYVDLSSY | HLA-B*15:01 | 9 | 0.841065 | 0.05 | -0.18478 | 45.95% (17/37) | Non-Toxin | IEDB recommended 2020.09 (NetMHCpan EL 4.1) |
| 683 | Fusion Protein | IIYVDLSSY | HLA-A*30:02 | 9 | 0.6073 | 0.1 | -0.18478 | 45.95% (17/37) | Non-Toxin | IEDB recommended 2020.09 (NetMHCpan EL 4.1) |
| 684 | Fusion Protein | IIYVDLSSYY | HLA-A*30:02 | 10 | 0.577055 | 0.11 | -0.24637 | 45.95% (17/37) | Non-Toxin | IEDB recommended 2020.09 (NetMHCpan EL 4.1) |
| 685 | Fusion Protein | ILTEIQQAY | HLA-B*15:01 | 9 | 0.920345 | 0.02 | 0.05901 | 100.00% (37/37) | Non-Toxin | IEDB recommended 2020.09 (NetMHCpan EL 4.1) |
| 686 | Fusion Protein | ILTEIQQAY | HLA-A*30:02 | 9 | 0.696212 | 0.06 | 0.05901 | 100.00% (37/37) | Non-Toxin | IEDB recommended 2020.09 (NetMHCpan EL 4.1) |
| 687 | Fusion Protein | ILTEIQQAY | HLA-B*35:01 | 9 | 0.721227 | 0.11 | 0.05901 | 100.00% (37/37) | Non-Toxin | IEDB recommended 2020.09 (NetMHCpan EL 4.1) |
| 688 | Fusion Protein | IQQAYIQEL | HLA-A*02:06 | 9 | 0.780163 | 0.08 | 0.08419 | 100.00% (37/37) | Non-Toxin | IEDB recommended 2020.09 (NetMHCpan EL 4.1) |
| 689 | Fusion Protein | ISECSVGILHY | HLA-A*01:01 | 11 | 0.655221 | 0.11 | 0.02517 | 100.00% (37/37) | Non-Toxin | IEDB recommended 2020.09 (NetMHCpan EL 4.1) |
| 690 | Fusion Protein | IVIKMIPNV | HLA-A*02:06 | 9 | 0.865559 | 0.05 | -0.23266 | 97.30% (36/37) | Non-Toxin | IEDB recommended 2020.09 (NetMHCpan EL 4.1) |
| 691 | Fusion Protein | IYKNNTHDL | HLA-A*24:02 | 9 | 0.727965 | 0.08 | -0.00601 | 100.00% (37/37) | Non-Toxin | IEDB recommended 2020.09 (NetMHCpan EL 4.1) |
| 692 | Fusion Protein | IYVDLSSYY | HLA-A*30:02 | 9 | 0.662808 | 0.07 | -0.27259 | 45.95% (17/37) | Non-Toxin | IEDB recommended 2020.09 (NetMHCpan EL 4.1) |
| 693 | Fusion Protein | KIKSNPLTK | HLA-A*03:01 | 9 | 0.968066 | 0.01 | -0.23989 | 100.00% (37/37) | Non-Toxin | IEDB recommended 2020.09 (NetMHCpan EL 4.1) |
| 694 | Fusion Protein | KIKSNPLTK | HLA-A*30:01 | 9 | 0.960464 | 0.01 | -0.23989 | 100.00% (37/37) | Non-Toxin | IEDB recommended 2020.09 (NetMHCpan EL 4.1) |
| 695 | Fusion Protein | KIKSNPLTK | HLA-A*11:01 | 9 | 0.792886 | 0.08 | -0.23989 | 100.00% (37/37) | Non-Toxin | IEDB recommended 2020.09 (NetMHCpan EL 4.1) |
| 696 | Fusion Protein | KLSKIGLVK | HLA-A*03:01 | 9 | 0.949033 | 0.02 | -0.09444 | 100.00% (37/37) | Non-Toxin | IEDB recommended 2020.09 (NetMHCpan EL 4.1) |
| 697 | Fusion Protein | KLSKIGLVK | HLA-A*30:01 | 9 | 0.540892 | 0.11 | -0.09444 | 100.00% (37/37) | Non-Toxin | IEDB recommended 2020.09 (NetMHCpan EL 4.1) |
| 698 | Fusion Protein | KMIPNVSNM | HLA-B*15:01 | 9 | 0.760931 | 0.09 | -0.0788 | 97.30% (36/37) | Non-Toxin | IEDB recommended 2020.09 (NetMHCpan EL 4.1) |
| 699 | Fusion Protein | KMIPNVSNM | HLA-A*02:03 | 9 | 0.704273 | 0.1 | -0.0788 | 97.30% (36/37) | Non-Toxin | IEDB recommended 2020.09 (NetMHCpan EL 4.1) |
| 700 | Fusion Protein | KMIPNVSNM | HLA-A*32:01 | 9 | 0.498504 | 0.1 | -0.0788 | 97.30% (36/37) | Non-Toxin | IEDB recommended 2020.09 (NetMHCpan EL 4.1) |
| 701 | Fusion Protein | KYKIKSNPLTK | HLA-A*30:01 | 11 | 0.710571 | 0.04 | -0.3106 | 100.00% (37/37) | Non-Toxin | IEDB recommended 2020.09 (NetMHCpan EL 4.1) |
| 702 | Fusion Protein | KYKIKSNPLTK | HLA-A*03:01 | 11 | 0.785606 | 0.1 | -0.3106 | 100.00% (37/37) | Non-Toxin | IEDB recommended 2020.09 (NetMHCpan EL 4.1) |
| 703 | Fusion Protein | KYLSDLLF | HLA-A*23:01 | 8 | 0.641251 | 0.1 | -0.16827 | 97.30% (36/37) | Non-Toxin | IEDB recommended 2020.09 (NetMHCpan EL 4.1) |
| 704 | Fusion Protein | KYLSDLLFVF | HLA-A*23:01 | 10 | 0.918982 | 0.02 | -0.04679 | 97.30% (36/37) | Non-Toxin | IEDB recommended 2020.09 (NetMHCpan EL 4.1) |
| 705 | Fusion Protein | KYLSDLLFVF | HLA-A*24:02 | 10 | 0.905914 | 0.02 | -0.04679 | 97.30% (36/37) | Non-Toxin | IEDB recommended 2020.09 (NetMHCpan EL 4.1) |
| 706 | Fusion Protein | LLDTVNPSL | HLA-A*02:01 | 9 | 0.952173 | 0.02 | -0.02565 | 100.00% (37/37) | Non-Toxin | IEDB recommended 2020.09 (NetMHCpan EL 4.1) |
| 707 | Fusion Protein | LLDTVNPSL | HLA-A*02:06 | 9 | 0.841892 | 0.06 | -0.02565 | 100.00% (37/37) | Non-Toxin | IEDB recommended 2020.09 (NetMHCpan EL 4.1) |
| 708 | Fusion Protein | LQDPVSNSM | HLA-A*02:06 | 9 | 0.736395 | 0.1 | -0.22161 | 97.30% (36/37) | Non-Toxin | IEDB recommended 2020.09 (NetMHCpan EL 4.1) |
| 709 | Fusion Protein | LSLDLALSK | HLA-A*11:01 | 9 | 0.76376 | 0.1 | -0.06127 | 97.30% (36/37) | Non-Toxin | IEDB recommended 2020.09 (NetMHCpan EL 4.1) |
| 710 | Fusion Protein | LSLDLALSKY | HLA-A*01:01 | 10 | 0.912099 | 0.03 | -0.23004 | 97.30% (36/37) | Non-Toxin | IEDB recommended 2020.09 (NetMHCpan EL 4.1) |
| 711 | Fusion Protein | LTEIQQAY | HLA-A*01:01 | 8 | 0.72202 | 0.09 | -0.0224 | 100.00% (37/37) | Non-Toxin | IEDB recommended 2020.09 (NetMHCpan EL 4.1) |
| 712 | Fusion Protein | LTKDIVIKM | HLA-A*26:01 | 9 | 0.60023 | 0.08 | 0.1071 | 100.00% (37/37) | Non-Toxin | IEDB recommended 2020.09 (NetMHCpan EL 4.1) |
| 713 | Fusion Protein | LVKGVTRKY | HLA-A*26:01 | 9 | 0.677122 | 0.06 | -0.04148 | 43.24% (16/37) | Non-Toxin | IEDB recommended 2020.09 (NetMHCpan EL 4.1) |
| 714 | Fusion Protein | LVKGVTRKY | HLA-B*15:01 | 9 | 0.784744 | 0.08 | -0.04148 | 43.24% (16/37) | Non-Toxin | IEDB recommended 2020.09 (NetMHCpan EL 4.1) |
| 715 | Fusion Protein | LVKGVTRKY | HLA-A*30:02 | 9 | 0.647528 | 0.08 | -0.04148 | 43.24% (16/37) | Non-Toxin | IEDB recommended 2020.09 (NetMHCpan EL 4.1) |
| 716 | Fusion Protein | MTIQAISQA | HLA-A*68:02 | 9 | 0.811518 | 0.05 | -0.11728 | 97.30% (36/37) | Non-Toxin | IEDB recommended 2020.09 (NetMHCpan EL 4.1) |
| 717 | Fusion Protein | MTIQAISQAF | HLA-A*26:01 | 10 | 0.525524 | 0.11 | -0.13629 | 97.30% (36/37) | Non-Toxin | IEDB recommended 2020.09 (NetMHCpan EL 4.1) |
| 718 | Fusion Protein | NPLTKDIVI | HLA-B*51:01 | 9 | 0.709989 | 0.07 | -0.01722 | 100.00% (37/37) | Non-Toxin | IEDB recommended 2020.09 (NetMHCpan EL 4.1) |
| 719 | Fusion Protein | NTYSRLEDR | HLA-A*33:01 | 9 | 0.686379 | 0.08 | -0.03025 | 100.00% (37/37) | Non-Toxin | IEDB recommended 2020.09 (NetMHCpan EL 4.1) |
| 720 | Fusion Protein | NVIISLGKY | HLA-A*26:01 | 9 | 0.85858 | 0.03 | -0.09182 | 100.00% (37/37) | Non-Toxin | IEDB recommended 2020.09 (NetMHCpan EL 4.1) |
| 721 | Fusion Protein | PTSSGDLYY | HLA-A*01:01 | 9 | 0.90589 | 0.03 | -0.17781 | 100.00% (37/37) | Non-Toxin | IEDB recommended 2020.09 (NetMHCpan EL 4.1) |
| 722 | Fusion Protein | QETAEKTVY | HLA-B*44:02 | 9 | 0.911123 | 0.02 | 0.00335 | 100.00% (37/37) | Non-Toxin | IEDB recommended 2020.09 (NetMHCpan EL 4.1) |
| 723 | Fusion Protein | QETAEKTVY | HLA-B*44:03 | 9 | 0.944413 | 0.03 | 0.00335 | 100.00% (37/37) | Non-Toxin | IEDB recommended 2020.09 (NetMHCpan EL 4.1) |

| | | | | | | | | | | |
|---|---|---|---|---|---|---|---|---|---|---|
| 724 | Fusion Protein | QETAEKTVYV | HLA-A*68:02 | 10 | 0.760356 | 0.06 | 0.00869 | 100.00% (37/37) | Non-Toxin | IEDB recommended 2020.09 (NetMHCpan EL 4.1) |
| 725 | Fusion Protein | QITAGVALY | HLA-A*26:01 | 9 | 0.636128 | 0.07 | 0.15037 | 97.30% (36/37) | Non-Toxin | IEDB recommended 2020.09 (NetMHCpan EL 4.1) |
| 726 | Fusion Protein | RFALSNGVLF | HLA-A*23:01 | 10 | 0.697575 | 0.08 | -0.1056 | 100.00% (37/37) | Non-Toxin | IEDB recommended 2020.09 (NetMHCpan EL 4.1) |
| 727 | Fusion Protein | RFALSNGVLF | HLA-A*24:02 | 10 | 0.723486 | 0.09 | -0.1056 | 100.00% (37/37) | Non-Toxin | IEDB recommended 2020.09 (NetMHCpan EL 4.1) |
| 728 | Fusion Protein | RLAGVIMAGV | HLA-A*02:03 | 10 | 0.694341 | 0.1 | 0.10412 | 100.00% (37/37) | Non-Toxin | IEDB recommended 2020.09 (NetMHCpan EL 4.1) |
| 729 | Fusion Protein | RLLDTVNPSL | HLA-A*02:01 | 10 | 0.879786 | 0.04 | -0.01539 | 100.00% (37/37) | Non-Toxin | IEDB recommended 2020.09 (NetMHCpan EL 4.1) |
| 730 | Fusion Protein | RLNGILTPI | HLA-A*02:03 | 9 | 0.781955 | 0.06 | 0.17744 | 100.00% (37/37) | Non-Toxin | IEDB recommended 2020.09 (NetMHCpan EL 4.1) |
| 731 | Fusion Protein | RLNGILTPI | HLA-A*32:01 | 9 | 0.53261 | 0.09 | 0.17744 | 100.00% (37/37) | Non-Toxin | IEDB recommended 2020.09 (NetMHCpan EL 4.1) |
| 732 | Fusion Protein | RLNGILTPIK | HLA-A*03:01 | 10 | 0.909081 | 0.03 | 0.25574 | 100.00% (37/37) | Non-Toxin | IEDB recommended 2020.09 (NetMHCpan EL 4.1) |
| 733 | Fusion Protein | RPTSSGDLYY | HLA-A*01:01 | 10 | 0.759856 | 0.08 | -0.27261 | 100.00% (37/37) | Non-Toxin | IEDB recommended 2020.09 (NetMHCpan EL 4.1) |
| 734 | Fusion Protein | RVRPTSSGDLY | HLA-A*30:02 | 11 | 0.778439 | 0.03 | -0.23462 | 100.00% (37/37) | Non-Toxin | IEDB recommended 2020.09 (NetMHCpan EL 4.1) |
| 735 | Fusion Protein | RVRPTSSGDLY | HLA-A*30:01 | 11 | 0.531437 | 0.11 | -0.23462 | 100.00% (37/37) | Non-Toxin | IEDB recommended 2020.09 (NetMHCpan EL 4.1) |
| 736 | Fusion Protein | SECSVGILHY | HLA-B*44:03 | 10 | 0.909429 | 0.04 | 0.02405 | 100.00% (37/37) | Non-Toxin | IEDB recommended 2020.09 (NetMHCpan EL 4.1) |
| 737 | Fusion Protein | SECSVGILHY | HLA-B*44:02 | 10 | 0.824313 | 0.05 | 0.02405 | 100.00% (37/37) | Non-Toxin | IEDB recommended 2020.09 (NetMHCpan EL 4.1) |
| 738 | Fusion Protein | SEWISIVPNF | HLA-B*44:03 | 10 | 0.779422 | 0.09 | 0.20004 | 97.30% (36/37) | Non-Toxin | IEDB recommended 2020.09 (NetMHCpan EL 4.1) |
| 739 | Fusion Protein | SITGQIIYV | HLA-A*02:06 | 9 | 0.884539 | 0.05 | 0.16934 | 100.00% (37/37) | Non-Toxin | IEDB recommended 2020.09 (NetMHCpan EL 4.1) |
| 740 | Fusion Protein | SITGQIIYV | HLA-A*02:01 | 9 | 0.817011 | 0.07 | 0.16934 | 100.00% (37/37) | Non-Toxin | IEDB recommended 2020.09 (NetMHCpan EL 4.1) |
| 741 | Fusion Protein | SITGQIIYV | HLA-A*02:03 | 9 | 0.743961 | 0.08 | 0.16934 | 100.00% (37/37) | Non-Toxin | IEDB recommended 2020.09 (NetMHCpan EL 4.1) |
| 742 | Fusion Protein | SITGQIIYV | HLA-A*68:02 | 9 | 0.703353 | 0.08 | 0.16934 | 100.00% (37/37) | Non-Toxin | IEDB recommended 2020.09 (NetMHCpan EL 4.1) |
| 743 | Fusion Protein | SIVPNFILV | HLA-A*02:06 | 9 | 0.911247 | 0.03 | 0.21198 | 100.00% (37/37) | Non-Toxin | IEDB recommended 2020.09 (NetMHCpan EL 4.1) |
| 744 | Fusion Protein | SIVPNFILV | HLA-A*68:02 | 9 | 0.769957 | 0.06 | 0.21198 | 100.00% (37/37) | Non-Toxin | IEDB recommended 2020.09 (NetMHCpan EL 4.1) |
| 745 | Fusion Protein | SIVPNFILV | HLA-A*02:03 | 9 | 0.672035 | 0.11 | 0.21198 | 100.00% (37/37) | Non-Toxin | IEDB recommended 2020.09 (NetMHCpan EL 4.1) |
| 746 | Fusion Protein | SLDLALSKY | HLA-A*01:01 | 9 | 0.938569 | 0.02 | -0.24192 | 97.30% (36/37) | Non-Toxin | IEDB recommended 2020.09 (NetMHCpan EL 4.1) |
| 747 | Fusion Protein | SLGKYLGSV | HLA-A*02:03 | 9 | 0.747405 | 0.07 | -0.2881 | 100.00% (37/37) | Non-Toxin | IEDB recommended 2020.09 (NetMHCpan EL 4.1) |
| 748 | Fusion Protein | SMLSMIILY | HLA-A*30:02 | 9 | 0.674773 | 0.06 | -0.10995 | 100.00% (37/37) | Non-Toxin | IEDB recommended 2020.09 (NetMHCpan EL 4.1) |
| 749 | Fusion Protein | SSMNQSLQQSK | HLA-A*11:01 | 11 | 0.844997 | 0.05 | -0.65167 | 100.00% (37/37) | Non-Toxin | IEDB recommended 2020.09 (NetMHCpan EL 4.1) |
| 750 | Fusion Protein | SSYYIIVRV | HLA-A*68:02 | 9 | 0.837314 | 0.04 | 0.31504 | 45.95% (17/37) | Non-Toxin | IEDB recommended 2020.09 (NetMHCpan EL 4.1) |
| 751 | Fusion Protein | STNEAVVKL | HLA-A*68:02 | 9 | 0.826345 | 0.04 | 0.08445 | 97.30% (36/37) | Non-Toxin | IEDB recommended 2020.09 (NetMHCpan EL 4.1) |
| 752 | Fusion Protein | SVMENYKTR | HLA-A*68:01 | 9 | 0.905726 | 0.07 | -0.12535 | 100.00% (37/37) | Non-Toxin | IEDB recommended 2020.09 (NetMHCpan EL 4.1) |
| 753 | Fusion Protein | SVMENYKTR | HLA-A*31:01 | 9 | 0.769175 | 0.08 | -0.12535 | 100.00% (37/37) | Non-Toxin | IEDB recommended 2020.09 (NetMHCpan EL 4.1) |
| 754 | Fusion Protein | SVMENYKTR | HLA-A*33:01 | 9 | 0.610846 | 0.1 | -0.12535 | 100.00% (37/37) | Non-Toxin | IEDB recommended 2020.09 (NetMHCpan EL 4.1) |
| 755 | Fusion Protein | SYYIIVRVY | HLA-A*30:02 | 9 | 0.709825 | 0.05 | 0.36898 | 45.95% (17/37) | Non-Toxin | IEDB recommended 2020.09 (NetMHCpan EL 4.1) |
| 756 | Fusion Protein | SYYIIVRVYF | HLA-A*23:01 | 10 | 0.623632 | 0.1 | 0.38392 | 45.95% (17/37) | Non-Toxin | IEDB recommended 2020.09 (NetMHCpan EL 4.1) |
| 757 | Fusion Protein | TAAQITAGV | HLA-A*68:02 | 9 | 0.962136 | 0.01 | 0.1151 | 97.30% (36/37) | Non-Toxin | IEDB recommended 2020.09 (NetMHCpan EL 4.1) |
| 758 | Fusion Protein | TELSLDLAL | HLA-B*40:01 | 9 | 0.9368 | 0.05 | -0.14649 | 97.30% (36/37) | Non-Toxin | IEDB recommended 2020.09 (NetMHCpan EL 4.1) |
| 759 | Fusion Protein | TPIKGALEI | HLA-B*51:01 | 9 | 0.83588 | 0.03 | -0.05483 | 100.00% (37/37) | Non-Toxin | IEDB recommended 2020.09 (NetMHCpan EL 4.1) |
| 760 | Fusion Protein | TPIKGALEI | HLA-B*53:01 | 9 | 0.61083 | 0.08 | -0.05483 | 100.00% (37/37) | Non-Toxin | IEDB recommended 2020.09 (NetMHCpan EL 4.1) |
| 761 | Fusion Protein | TPIKGALEIY | HLA-B*35:01 | 10 | 0.920784 | 0.03 | 0.04912 | 100.00% (37/37) | Non-Toxin | IEDB recommended 2020.09 (NetMHCpan EL 4.1) |
| 762 | Fusion Protein | TVNPSLISM | HLA-A*26:01 | 9 | 0.712661 | 0.06 | -0.16914 | 100.00% (37/37) | Non-Toxin | IEDB recommended 2020.09 (NetMHCpan EL 4.1) |
| 763 | Fusion Protein | TVNPSLISM | HLA-A*68:02 | 9 | 0.633604 | 0.11 | -0.16914 | 100.00% (37/37) | Non-Toxin | IEDB recommended 2020.09 (NetMHCpan EL 4.1) |
| 764 | Fusion Protein | VLGNVIISL | HLA-A*02:01 | 9 | 0.879121 | 0.04 | 0.18563 | 100.00% (37/37) | Non-Toxin | IEDB recommended 2020.09 (NetMHCpan EL 4.1) |
| 765 | Fusion Protein | VPNFILVRNTL | HLA-B*07:02 | 11 | 0.74277 | 0.11 | 0.34064 | 100.00% (37/37) | Non-Toxin | IEDB recommended 2020.09 (NetMHCpan EL 4.1) |
| 766 | Fusion Protein | VPRFALSNGVL | HLA-B*07:02 | 11 | 0.834371 | 0.07 | 0.04743 | 100.00% (37/37) | Non-Toxin | IEDB recommended 2020.09 (NetMHCpan EL 4.1) |
| 767 | Fusion Protein | VRPTSSGDLYY | HLA-A*01:01 | 11 | 0.811377 | 0.06 | -0.24438 | 100.00% (37/37) | Non-Toxin | IEDB recommended 2020.09 (NetMHCpan EL 4.1) |
| 768 | Fusion Protein | VSFNNDNSEW | HLA-B*58:01 | 10 | 0.944867 | 0.04 | -0.04042 | 97.30% (36/37) | Non-Toxin | IEDB recommended 2020.09 (NetMHCpan EL 4.1) |
| 769 | Fusion Protein | VSFNNDNSEW | HLA-B*57:01 | 10 | 0.946468 | 0.06 | -0.04042 | 97.30% (36/37) | Non-Toxin | IEDB recommended 2020.09 (NetMHCpan EL 4.1) |
| 770 | Fusion Protein | VVSSHVPRF | HLA-A*32:01 | 9 | 0.603745 | 0.06 | -0.12893 | 100.00% (37/37) | Non-Toxin | IEDB recommended 2020.09 (NetMHCpan EL 4.1) |
| 771 | Fusion Protein | VYFPILTEI | HLA-A*24:02 | 9 | 0.978204 | 0.01 | 0.23726 | 100.00% (37/37) | Non-Toxin | IEDB recommended 2020.09 (NetMHCpan EL 4.1) |
| 772 | Fusion Protein | VYFPILTEI | HLA-A*23:01 | 9 | 0.978018 | 0.01 | 0.23726 | 100.00% (37/37) | Non-Toxin | IEDB recommended 2020.09 (NetMHCpan EL 4.1) |
| 773 | Fusion Protein | YATPMTNNM | HLA-B*35:01 | 9 | 0.776901 | 0.09 | -0.14226 | 97.30% (36/37) | Non-Toxin | IEDB recommended 2020.09 (NetMHCpan EL 4.1) |
| 774 | Fusion Protein | YEKLSKIGL | HLA-B*40:01 | 9 | 0.858409 | 0.08 | -0.31314 | 100.00% (37/37) | Non-Toxin | IEDB recommended 2020.09 (NetMHCpan EL 4.1) |

| | | | | | | | | | | |
|---|---|---|---|---|---|---|---|---|---|---|
| 775 | Fusion Protein | YVDLSSYY | HLA-A*01:01 | 8 | 0.865485 | 0.05 | -0.32391 | 45.95% (17/37) | Non-Toxin | IEDB recommended 2020.09 (NetMHCpan EL 4.1) |
| 776 | Fusion Protein | YYIIVRVYF | HLA-A*23:01 | 9 | 0.888305 | 0.03 | 0.29872 | 100.00% (37/37) | Non-Toxin | IEDB recommended 2020.09 (NetMHCpan EL 4.1) |
| 777 | Fusion Protein | YYIIVRVYF | HLA-A*24:02 | 9 | 0.856049 | 0.04 | 0.29872 | 100.00% (37/37) | Non-Toxin | IEDB recommended 2020.09 (NetMHCpan EL 4.1) |
| **S.No.** | **Protein** | **Peptide** | **Allele** | **Length** | **Score** | **Percentile rank** | **Immunogenicity** | **Conservancy** | **Toxicity** | **Method** |
| 778 | Polymerase | AEFFSFFRTF | HLA-B*44:03 | 10 | 0.823382 | 0.07 | 0.28526 | 100.00% (10/10) | Non-Toxin | IEDB recommended 2020.09 (NetMHCpan EL 4.1) |
| 779 | Polymerase | AEFFSFFRTF | HLA-B*44:02 | 10 | 0.727453 | 0.09 | 0.28526 | 100.00% (10/10) | Non-Toxin | IEDB recommended 2020.09 (NetMHCpan EL 4.1) |
| 780 | Polymerase | AELPIPEY | HLA-B*44:03 | 8 | 0.874582 | 0.05 | 0.1889 | 100.00% (10/10) | Non-Toxin | IEDB recommended 2020.09 (NetMHCpan EL 4.1) |
| 781 | Polymerase | AELPIPEY | HLA-B*44:02 | 8 | 0.731635 | 0.08 | 0.1889 | 100.00% (10/10) | Non-Toxin | IEDB recommended 2020.09 (NetMHCpan EL 4.1) |
| 782 | Polymerase | AEQDETVISL | HLA-B*40:01 | 10 | 0.981355 | 0.01 | 0.17454 | 100.00% (10/10) | Non-Toxin | IEDB recommended 2020.09 (NetMHCpan EL 4.1) |
| 783 | Polymerase | AGRLFAKMTYK | HLA-A*03:01 | 11 | 0.811549 | 0.08 | -0.18696 | 100.00% (10/10) | Non-Toxin | IEDB recommended 2020.09 (NetMHCpan EL 4.1) |
| 784 | Polymerase | ALIASGVGK | HLA-A*03:01 | 9 | 0.883642 | 0.04 | 0.00801 | 100.00% (10/10) | Non-Toxin | IEDB recommended 2020.09 (NetMHCpan EL 4.1) |
| 785 | Polymerase | ALIASGVGKY | HLA-B*15:01 | 10 | 0.778333 | 0.08 | -0.10407 | 100.00% (10/10) | Non-Toxin | IEDB recommended 2020.09 (NetMHCpan EL 4.1) |
| 786 | Polymerase | ALSHPRVFK | HLA-A*03:01 | 9 | 0.971332 | 0.01 | 0.12001 | 100.00% (10/10) | Non-Toxin | IEDB recommended 2020.09 (NetMHCpan EL 4.1) |
| 787 | Polymerase | ALSHPRVFK | HLA-A*11:01 | 9 | 0.875731 | 0.04 | 0.12001 | 100.00% (10/10) | Non-Toxin | IEDB recommended 2020.09 (NetMHCpan EL 4.1) |
| 788 | Polymerase | ALSHPRVFK | HLA-A*30:01 | 9 | 0.666133 | 0.06 | 0.12001 | 100.00% (10/10) | Non-Toxin | IEDB recommended 2020.09 (NetMHCpan EL 4.1) |
| 789 | Polymerase | ALSHPRVFKR | HLA-A*31:01 | 10 | 0.787573 | 0.07 | 0.03011 | 100.00% (10/10) | Non-Toxin | IEDB recommended 2020.09 (NetMHCpan EL 4.1) |
| 790 | Polymerase | ALSPIKDEW | HLA-A*32:01 | 9 | 0.662995 | 0.05 | -0.06104 | 100.00% (10/10) | Non-Toxin | IEDB recommended 2020.09 (NetMHCpan EL 4.1) |
| 791 | Polymerase | ALSPIKDEW | HLA-B*58:01 | 9 | 0.871421 | 0.08 | -0.06104 | 100.00% (10/10) | Non-Toxin | IEDB recommended 2020.09 (NetMHCpan EL 4.1) |
| 792 | Polymerase | ALYLGQSISIK | HLA-A*03:01 | 11 | 0.813961 | 0.07 | -0.18984 | 100.00% (10/10) | Non-Toxin | IEDB recommended 2020.09 (NetMHCpan EL 4.1) |
| 793 | Polymerase | APIGGFNYL | HLA-B*07:02 | 9 | 0.907381 | 0.04 | 0.21288 | 100.00% (10/10) | Non-Toxin | IEDB recommended 2020.09 (NetMHCpan EL 4.1) |
| 794 | Polymerase | APIMKAHAI | HLA-B*08:01 | 9 | 0.854209 | 0.03 | -0.25651 | 100.00% (10/10) | Non-Toxin | IEDB recommended 2020.09 (NetMHCpan EL 4.1) |
| 795 | Polymerase | APIMKAHAI | HLA-B*07:02 | 9 | 0.928737 | 0.04 | -0.25651 | 100.00% (10/10) | Non-Toxin | IEDB recommended 2020.09 (NetMHCpan EL 4.1) |
| 796 | Polymerase | ASKHIIRLK | HLA-A*30:01 | 9 | 0.884537 | 0.01 | 0.25463 | 100.00% (10/10) | Non-Toxin | IEDB recommended 2020.09 (NetMHCpan EL 4.1) |
| 797 | Polymerase | AYPECNNILF | HLA-A*24:02 | 10 | 0.872057 | 0.04 | 0.1381 | 90.00% (9/10) | Non-Toxin | IEDB recommended 2020.09 (NetMHCpan EL 4.1) |
| 798 | Polymerase | AYPECNNILF | HLA-A*23:01 | 10 | 0.758397 | 0.06 | 0.1381 | 90.00% (9/10) | Non-Toxin | IEDB recommended 2020.09 (NetMHCpan EL 4.1) |
| 799 | Polymerase | DAELPIPEY | HLA-B*35:01 | 9 | 0.904508 | 0.04 | 0.18496 | 100.00% (10/10) | Non-Toxin | IEDB recommended 2020.09 (NetMHCpan EL 4.1) |
| 800 | Polymerase | DALEAMVGR | HLA-A*68:01 | 9 | 0.924229 | 0.06 | 0.02459 | 100.00% (10/10) | Non-Toxin | IEDB recommended 2020.09 (NetMHCpan EL 4.1) |
| 801 | Polymerase | DALEAMVGR | HLA-A*33:01 | 9 | 0.726337 | 0.06 | 0.02459 | 100.00% (10/10) | Non-Toxin | IEDB recommended 2020.09 (NetMHCpan EL 4.1) |
| 802 | Polymerase | DEHELLKTL | HLA-B*44:03 | 9 | 0.820297 | 0.07 | -0.06931 | 100.00% (10/10) | Non-Toxin | IEDB recommended 2020.09 (NetMHCpan EL 4.1) |
| 803 | Polymerase | DEHELLKTLF | HLA-B*44:03 | 10 | 0.767646 | 0.1 | -0.08707 | 100.00% (10/10) | Non-Toxin | IEDB recommended 2020.09 (NetMHCpan EL 4.1) |
| 804 | Polymerase | DGLLVSKI | HLA-B*51:01 | 8 | 0.700667 | 0.07 | -0.31229 | 100.00% (10/10) | Non-Toxin | IEDB recommended 2020.09 (NetMHCpan EL 4.1) |
| 805 | Polymerase | DIGSDINTLR | HLA-A*68:01 | 10 | 0.911368 | 0.07 | 0.01592 | 100.00% (10/10) | Non-Toxin | IEDB recommended 2020.09 (NetMHCpan EL 4.1) |
| 806 | Polymerase | DIKLGNVKR | HLA-A*33:01 | 9 | 0.843853 | 0.02 | -0.14541 | 100.00% (10/10) | Non-Toxin | IEDB recommended 2020.09 (NetMHCpan EL 4.1) |
| 807 | Polymerase | DIKYQPLISR | HLA-A*33:01 | 10 | 0.812091 | 0.03 | -0.1921 | 100.00% (10/10) | Non-Toxin | IEDB recommended 2020.09 (NetMHCpan EL 4.1) |
| 808 | Polymerase | DIKYQPLISR | HLA-A*68:01 | 10 | 0.902947 | 0.07 | -0.1921 | 100.00% (10/10) | Non-Toxin | IEDB recommended 2020.09 (NetMHCpan EL 4.1) |
| 809 | Polymerase | DLKKYYQI | HLA-B*08:01 | 8 | 0.579563 | 0.1 | -0.39184 | 100.00% (10/10) | Non-Toxin | IEDB recommended 2020.09 (NetMHCpan EL 4.1) |
| 810 | Polymerase | DPYNMLEYV | HLA-B*51:01 | 9 | 0.882661 | 0.02 | -0.10681 | 100.00% (10/10) | Non-Toxin | IEDB recommended 2020.09 (NetMHCpan EL 4.1) |
| 811 | Polymerase | DSQSITKTI | HLA-B*51:01 | 9 | 0.758219 | 0.05 | -0.19725 | 100.00% (10/10) | Non-Toxin | IEDB recommended 2020.09 (NetMHCpan EL 4.1) |
| 812 | Polymerase | DSYEYIINR | HLA-A*68:01 | 9 | 0.978937 | 0.01 | 0.32977 | 100.00% (10/10) | Non-Toxin | IEDB recommended 2020.09 (NetMHCpan EL 4.1) |
| 813 | Polymerase | DSYEYIINR | HLA-A*33:01 | 9 | 0.915796 | 0.01 | 0.32977 | 100.00% (10/10) | Non-Toxin | IEDB recommended 2020.09 (NetMHCpan EL 4.1) |
| 814 | Polymerase | DVLKAITPV | HLA-A*68:02 | 9 | 0.681954 | 0.09 | -0.03094 | 100.00% (10/10) | Non-Toxin | IEDB recommended 2020.09 (NetMHCpan EL 4.1) |
| 815 | Polymerase | EALIASGVGKY | HLA-A*26:01 | 11 | 0.749702 | 0.05 | -0.01822 | 100.00% (10/10) | Non-Toxin | IEDB recommended 2020.09 (NetMHCpan EL 4.1) |
| 816 | Polymerase | EAMVGRYI | HLA-B*51:01 | 8 | 0.641085 | 0.1 | 0.06314 | 100.00% (10/10) | Non-Toxin | IEDB recommended 2020.09 (NetMHCpan EL 4.1) |
| 817 | Polymerase | EAWYLASQR | HLA-A*68:01 | 9 | 0.946822 | 0.04 | -0.11309 | 100.00% (10/10) | Non-Toxin | IEDB recommended 2020.09 (NetMHCpan EL 4.1) |
| 818 | Polymerase | EAWYLASQR | HLA-A*33:01 | 9 | 0.761082 | 0.05 | -0.11309 | 100.00% (10/10) | Non-Toxin | IEDB recommended 2020.09 (NetMHCpan EL 4.1) |
| 819 | Polymerase | EFFSFFRTF | HLA-A*23:01 | 9 | 0.831873 | 0.04 | 0.16209 | 100.00% (10/10) | Non-Toxin | IEDB recommended 2020.09 (NetMHCpan EL 4.1) |
| 820 | Polymerase | EFFSFFRTF | HLA-A*24:02 | 9 | 0.747448 | 0.07 | 0.16209 | 100.00% (10/10) | Non-Toxin | IEDB recommended 2020.09 (NetMHCpan EL 4.1) |
| 821 | Polymerase | EFNPHNHYK | HLA-A*33:01 | 9 | 0.712964 | 0.06 | 0.03729 | 100.00% (10/10) | Non-Toxin | IEDB recommended 2020.09 (NetMHCpan EL 4.1) |
| 822 | Polymerase | EICAKQAQLY | HLA-A*26:01 | 10 | 0.622105 | 0.07 | -0.36834 | 100.00% (10/10) | Non-Toxin | IEDB recommended 2020.09 (NetMHCpan EL 4.1) |
| 823 | Polymerase | EIINIHECR | HLA-A*68:01 | 9 | 0.954696 | 0.03 | 0.24974 | 100.00% (10/10) | Non-Toxin | IEDB recommended 2020.09 (NetMHCpan EL 4.1) |
| 824 | Polymerase | EIINIHECR | HLA-A*33:01 | 9 | 0.785381 | 0.03 | 0.24974 | 100.00% (10/10) | Non-Toxin | IEDB recommended 2020.09 (NetMHCpan EL 4.1) |

| | | | | | | | | | | |
|---|---|---|---|---|---|---|---|---|---|---|
| 825 | Polymerase | EIYGLPGFF | HLA-A*26:01 | 9 | 0.832767 | 0.04 | 0.10866 | 100.00% (10/10) | Non-Toxin | IEDB recommended 2020.09 (NetMHCpan EL 4.1) |
| 826 | Polymerase | ELKLDSDLSMY | HLA-A*01:01 | 11 | 0.729152 | 0.09 | -0.45172 | 100.00% (10/10) | Non-Toxin | IEDB recommended 2020.09 (NetMHCpan EL 4.1) |
| 827 | Polymerase | ELKLFPSEY | HLA-A*26:01 | 9 | 0.712676 | 0.06 | -0.05872 | 100.00% (10/10) | Non-Toxin | IEDB recommended 2020.09 (NetMHCpan EL 4.1) |
| 828 | Polymerase | ESMAIFAER | HLA-A*68:01 | 9 | 0.986103 | 0.01 | 0.31369 | 100.00% (10/10) | Non-Toxin | IEDB recommended 2020.09 (NetMHCpan EL 4.1) |
| 829 | Polymerase | ESMAIFAER | HLA-A*33:01 | 9 | 0.920339 | 0.01 | 0.31369 | 100.00% (10/10) | Non-Toxin | IEDB recommended 2020.09 (NetMHCpan EL 4.1) |
| 830 | Polymerase | ETDDYNGIY | HLA-A*01:01 | 9 | 0.979774 | 0.01 | 0.12619 | 100.00% (10/10) | Non-Toxin | IEDB recommended 2020.09 (NetMHCpan EL 4.1) |
| 831 | Polymerase | ETDDYNGIY | HLA-A*26:01 | 9 | 0.534209 | 0.1 | 0.12619 | 100.00% (10/10) | Non-Toxin | IEDB recommended 2020.09 (NetMHCpan EL 4.1) |
| 832 | Polymerase | ETDDYNGIYHL | HLA-A*68:02 | 11 | 0.683156 | 0.09 | 0.19368 | 100.00% (10/10) | Non-Toxin | IEDB recommended 2020.09 (NetMHCpan EL 4.1) |
| 833 | Polymerase | ETIISTHLF | HLA-A*26:01 | 9 | 0.944687 | 0.01 | 0.07338 | 100.00% (10/10) | Non-Toxin | IEDB recommended 2020.09 (NetMHCpan EL 4.1) |
| 834 | Polymerase | ETIISTHLFIY | HLA-A*26:01 | 11 | 0.612948 | 0.08 | 0.25144 | 90.00% (9/10) | Non-Toxin | IEDB recommended 2020.09 (NetMHCpan EL 4.1) |
| 835 | Polymerase | ETKQAGRLF | HLA-A*26:01 | 9 | 0.698682 | 0.06 | -0.07936 | 100.00% (10/10) | Non-Toxin | IEDB recommended 2020.09 (NetMHCpan EL 4.1) |
| 836 | Polymerase | ETNTRIAAI | HLA-A*68:02 | 9 | 0.762928 | 0.06 | 0.26852 | 100.00% (10/10) | Non-Toxin | IEDB recommended 2020.09 (NetMHCpan EL 4.1) |
| 837 | Polymerase | ETTVKSDIKY | HLA-A*26:01 | 10 | 0.619616 | 0.07 | -0.30976 | 100.00% (10/10) | Non-Toxin | IEDB recommended 2020.09 (NetMHCpan EL 4.1) |
| 838 | Polymerase | ETTWIGNLDSY | HLA-A*26:01 | 11 | 0.797646 | 0.04 | 0.30341 | 100.00% (10/10) | Non-Toxin | IEDB recommended 2020.09 (NetMHCpan EL 4.1) |
| 839 | Polymerase | EVDNNHLIY | HLA-A*01:01 | 9 | 0.982598 | 0.01 | 0.09324 | 100.00% (10/10) | Non-Toxin | IEDB recommended 2020.09 (NetMHCpan EL 4.1) |
| 840 | Polymerase | EVDNNHLIY | HLA-A*26:01 | 9 | 0.562828 | 0.09 | 0.09324 | 100.00% (10/10) | Non-Toxin | IEDB recommended 2020.09 (NetMHCpan EL 4.1) |
| 841 | Polymerase | EVNDQGITSV | HLA-A*68:02 | 10 | 0.871511 | 0.03 | 0.0018 | 100.00% (10/10) | Non-Toxin | IEDB recommended 2020.09 (NetMHCpan EL 4.1) |
| 842 | Polymerase | EVPDALEAM | HLA-A*26:01 | 9 | 0.823839 | 0.04 | 0.15374 | 100.00% (10/10) | Non-Toxin | IEDB recommended 2020.09 (NetMHCpan EL 4.1) |
| 843 | Polymerase | EVPDALEAMV | HLA-A*68:02 | 10 | 0.680838 | 0.09 | 0.07069 | 100.00% (10/10) | Non-Toxin | IEDB recommended 2020.09 (NetMHCpan EL 4.1) |
| 844 | Polymerase | EYIINRTAGR | HLA-A*33:01 | 10 | 0.678265 | 0.08 | 0.31058 | 100.00% (10/10) | Non-Toxin | IEDB recommended 2020.09 (NetMHCpan EL 4.1) |
| 845 | Polymerase | FAERLDEIY | HLA-A*01:01 | 9 | 0.732737 | 0.09 | 0.25692 | 100.00% (10/10) | Non-Toxin | IEDB recommended 2020.09 (NetMHCpan EL 4.1) |
| 846 | Polymerase | FALYLGQSI | HLA-B*51:01 | 9 | 0.699321 | 0.07 | -0.18064 | 100.00% (10/10) | Non-Toxin | IEDB recommended 2020.09 (NetMHCpan EL 4.1) |
| 847 | Polymerase | FEKLSQNLL | HLA-B*40:01 | 9 | 0.885711 | 0.07 | -0.36324 | 100.00% (10/10) | Non-Toxin | IEDB recommended 2020.09 (NetMHCpan EL 4.1) |
| 848 | Polymerase | FIDDLLSIL | HLA-A*02:06 | 9 | 0.820214 | 0.07 | -0.05358 | 100.00% (10/10) | Non-Toxin | IEDB recommended 2020.09 (NetMHCpan EL 4.1) |
| 849 | Polymerase | FIDDLLSIL | HLA-A*02:01 | 9 | 0.753755 | 0.1 | -0.05358 | 100.00% (10/10) | Non-Toxin | IEDB recommended 2020.09 (NetMHCpan EL 4.1) |
| 850 | Polymerase | FIYSKKIHY | HLA-B*15:01 | 9 | 0.878613 | 0.03 | -0.44945 | 90.00% (9/10) | Non-Toxin | IEDB recommended 2020.09 (NetMHCpan EL 4.1) |
| 851 | Polymerase | FIYSKKIHY | HLA-A*26:01 | 9 | 0.676405 | 0.06 | -0.44945 | 90.00% (9/10) | Non-Toxin | IEDB recommended 2020.09 (NetMHCpan EL 4.1) |
| 852 | Polymerase | FIYSKKIHY | HLA-A*30:02 | 9 | 0.647328 | 0.08 | -0.44945 | 90.00% (9/10) | Non-Toxin | IEDB recommended 2020.09 (NetMHCpan EL 4.1) |
| 853 | Polymerase | FLMDRRVIL | HLA-B*08:01 | 9 | 0.946245 | 0.01 | 0.17704 | 100.00% (10/10) | Non-Toxin | IEDB recommended 2020.09 (NetMHCpan EL 4.1) |
| 854 | Polymerase | FLMDRRVIL | HLA-A*02:03 | 9 | 0.817851 | 0.05 | 0.17704 | 100.00% (10/10) | Non-Toxin | IEDB recommended 2020.09 (NetMHCpan EL 4.1) |
| 855 | Polymerase | FLMDRRVIL | HLA-A*02:01 | 9 | 0.851636 | 0.06 | 0.17704 | 100.00% (10/10) | Non-Toxin | IEDB recommended 2020.09 (NetMHCpan EL 4.1) |
| 856 | Polymerase | FNKVKSAL | HLA-B*08:01 | 8 | 0.82196 | 0.03 | -0.36117 | 100.00% (10/10) | Non-Toxin | IEDB recommended 2020.09 (NetMHCpan EL 4.1) |
| 857 | Polymerase | FPISRLFNM | HLA-B*35:01 | 9 | 0.92836 | 0.03 | 0.01171 | 100.00% (10/10) | Non-Toxin | IEDB recommended 2020.09 (NetMHCpan EL 4.1) |
| 858 | Polymerase | FPISRLFNM | HLA-B*51:01 | 9 | 0.769648 | 0.05 | 0.01171 | 100.00% (10/10) | Non-Toxin | IEDB recommended 2020.09 (NetMHCpan EL 4.1) |
| 859 | Polymerase | FPISRLFNM | HLA-B*08:01 | 9 | 0.744343 | 0.05 | 0.01171 | 100.00% (10/10) | Non-Toxin | IEDB recommended 2020.09 (NetMHCpan EL 4.1) |
| 860 | Polymerase | FPISRLFNM | HLA-B*53:01 | 9 | 0.716075 | 0.05 | 0.01171 | 100.00% (10/10) | Non-Toxin | IEDB recommended 2020.09 (NetMHCpan EL 4.1) |
| 861 | Polymerase | FPISRLFNMY | HLA-B*35:01 | 10 | 0.81414 | 0.07 | -0.08153 | 100.00% (10/10) | Non-Toxin | IEDB recommended 2020.09 (NetMHCpan EL 4.1) |
| 862 | Polymerase | FPLWSTEEL | HLA-B*35:01 | 9 | 0.919156 | 0.03 | 0.23773 | 100.00% (10/10) | Non-Toxin | IEDB recommended 2020.09 (NetMHCpan EL 4.1) |
| 863 | Polymerase | FPLWSTEEL | HLA-B*53:01 | 9 | 0.742916 | 0.05 | 0.23773 | 100.00% (10/10) | Non-Toxin | IEDB recommended 2020.09 (NetMHCpan EL 4.1) |
| 864 | Polymerase | FPLWSTEEL | HLA-B*51:01 | 9 | 0.677867 | 0.08 | 0.23773 | 100.00% (10/10) | Non-Toxin | IEDB recommended 2020.09 (NetMHCpan EL 4.1) |
| 865 | Polymerase | FPVMGNRI | HLA-B*51:01 | 8 | 0.813447 | 0.04 | -0.09271 | 100.00% (10/10) | Non-Toxin | IEDB recommended 2020.09 (NetMHCpan EL 4.1) |
| 866 | Polymerase | FPVMGNRIY | HLA-B*35:01 | 9 | 0.988335 | 0.01 | -0.01495 | 100.00% (10/10) | Non-Toxin | IEDB recommended 2020.09 (NetMHCpan EL 4.1) |
| 867 | Polymerase | FPVMGNRIY | HLA-B*53:01 | 9 | 0.709672 | 0.06 | -0.01495 | 100.00% (10/10) | Non-Toxin | IEDB recommended 2020.09 (NetMHCpan EL 4.1) |
| 868 | Polymerase | FTDQKIRSM | HLA-B*08:01 | 9 | 0.590356 | 0.1 | -0.24706 | 100.00% (10/10) | Non-Toxin | IEDB recommended 2020.09 (NetMHCpan EL 4.1) |
| 869 | Polymerase | FYASLTYLR | HLA-A*33:01 | 9 | 0.814289 | 0.03 | -0.13763 | 100.00% (10/10) | Non-Toxin | IEDB recommended 2020.09 (NetMHCpan EL 4.1) |
| 870 | Polymerase | GEGSGSMML | HLA-B*40:01 | 9 | 0.943271 | 0.04 | -0.529 | 100.00% (10/10) | Non-Toxin | IEDB recommended 2020.09 (NetMHCpan EL 4.1) |
| 871 | Polymerase | GEKVDTNLIY | HLA-B*44:03 | 10 | 0.761614 | 0.1 | 0.09325 | 100.00% (10/10) | Non-Toxin | IEDB recommended 2020.09 (NetMHCpan EL 4.1) |
| 872 | Polymerase | GFFNWMHKR | HLA-A*31:01 | 9 | 0.864797 | 0.04 | -0.01681 | 100.00% (10/10) | Non-Toxin | IEDB recommended 2020.09 (NetMHCpan EL 4.1) |
| 873 | Polymerase | GFFNWMHKR | HLA-A*33:01 | 9 | 0.652583 | 0.09 | -0.01681 | 100.00% (10/10) | Non-Toxin | IEDB recommended 2020.09 (NetMHCpan EL 4.1) |
| 874 | Polymerase | GIEGYSQKTW | HLA-B*44:02 | 10 | 0.795517 | 0.05 | -0.36646 | 100.00% (10/10) | Non-Toxin | IEDB recommended 2020.09 (NetMHCpan EL 4.1) |
| 875 | Polymerase | GLIDPLFPV | HLA-A*02:01 | 9 | 0.976885 | 0.01 | 0.1366 | 100.00% (10/10) | Non-Toxin | IEDB recommended 2020.09 (NetMHCpan EL 4.1) |

| | | | | | | | | | | |
|---|---|---|---|---|---|---|---|---|---|---|
| 876 | Polymerase | GLIDPLFPV | HLA-A*02:06 | 9 | 0.971103 | 0.01 | 0.1366 | 100.00% (10/10) | Non-Toxin | IEDB recommended 2020.09 (NetMHCpan EL 4.1) |
| 877 | Polymerase | GLIDPLFPV | HLA-A*02:03 | 9 | 0.942868 | 0.02 | 0.1366 | 100.00% (10/10) | Non-Toxin | IEDB recommended 2020.09 (NetMHCpan EL 4.1) |
| 878 | Polymerase | GLIRSGLRK | HLA-A*03:01 | 9 | 0.863636 | 0.05 | -0.01304 | 100.00% (10/10) | Non-Toxin | IEDB recommended 2020.09 (NetMHCpan EL 4.1) |
| 879 | Polymerase | GLLDTTKGL | HLA-A*02:01 | 9 | 0.791648 | 0.08 | -0.06914 | 100.00% (10/10) | Non-Toxin | IEDB recommended 2020.09 (NetMHCpan EL 4.1) |
| 880 | Polymerase | GLLDTTKGL | HLA-A*02:03 | 9 | 0.689945 | 0.1 | -0.06914 | 100.00% (10/10) | Non-Toxin | IEDB recommended 2020.09 (NetMHCpan EL 4.1) |
| 881 | Polymerase | GLQPKLVSR | HLA-A*31:01 | 9 | 0.875448 | 0.04 | -0.33102 | 90.00% (9/10) | Non-Toxin | IEDB recommended 2020.09 (NetMHCpan EL 4.1) |
| 882 | Polymerase | GRLFAKMTYK | HLA-A*03:01 | 10 | 0.832314 | 0.06 | -0.1924 | 100.00% (10/10) | Non-Toxin | IEDB recommended 2020.09 (NetMHCpan EL 4.1) |
| 883 | Polymerase | GSGSMMLLY | HLA-A*30:02 | 9 | 0.658464 | 0.07 | -0.50761 | 100.00% (10/10) | Non-Toxin | IEDB recommended 2020.09 (NetMHCpan EL 4.1) |
| 884 | Polymerase | GYSQKTWTI | HLA-A*24:02 | 9 | 0.876993 | 0.04 | -0.1341 | 100.00% (10/10) | Non-Toxin | IEDB recommended 2020.09 (NetMHCpan EL 4.1) |
| 885 | Polymerase | GYSQKTWTI | HLA-A*23:01 | 9 | 0.718316 | 0.08 | -0.1341 | 100.00% (10/10) | Non-Toxin | IEDB recommended 2020.09 (NetMHCpan EL 4.1) |
| 886 | Polymerase | HIKNNIRRK | HLA-A*30:01 | 9 | 0.60974 | 0.07 | 0.11639 | 100.00% (10/10) | Non-Toxin | IEDB recommended 2020.09 (NetMHCpan EL 4.1) |
| 887 | Polymerase | HLDSPIVSGK | HLA-A*03:01 | 10 | 0.800942 | 0.08 | -0.12143 | 100.00% (10/10) | Non-Toxin | IEDB recommended 2020.09 (NetMHCpan EL 4.1) |
| 888 | Polymerase | HLLAEFFSF | HLA-A*32:01 | 9 | 0.699547 | 0.04 | 0.24561 | 100.00% (10/10) | Non-Toxin | IEDB recommended 2020.09 (NetMHCpan EL 4.1) |
| 889 | Polymerase | HMLADKVLEY | HLA-B*15:01 | 10 | 0.796862 | 0.07 | -0.06463 | 100.00% (10/10) | Non-Toxin | IEDB recommended 2020.09 (NetMHCpan EL 4.1) |
| 890 | Polymerase | HMRDKSTEI | HLA-B*08:01 | 9 | 0.893986 | 0.02 | -0.23535 | 100.00% (10/10) | Non-Toxin | IEDB recommended 2020.09 (NetMHCpan EL 4.1) |
| 891 | Polymerase | HPNLPYKV | HLA-B*51:01 | 8 | 0.661363 | 0.09 | -0.20954 | 100.00% (10/10) | Non-Toxin | IEDB recommended 2020.09 (NetMHCpan EL 4.1) |
| 892 | Polymerase | HTEFNPHNHY | HLA-A*01:01 | 10 | 0.963169 | 0.01 | 0.17709 | 100.00% (10/10) | Non-Toxin | IEDB recommended 2020.09 (NetMHCpan EL 4.1) |
| 893 | Polymerase | HTEFNPHNHY | HLA-B*44:03 | 10 | 0.915897 | 0.03 | 0.17709 | 100.00% (10/10) | Non-Toxin | IEDB recommended 2020.09 (NetMHCpan EL 4.1) |
| 894 | Polymerase | HTEFNPHNHY | HLA-A*30:02 | 10 | 0.754132 | 0.04 | 0.17709 | 100.00% (10/10) | Non-Toxin | IEDB recommended 2020.09 (NetMHCpan EL 4.1) |
| 895 | Polymerase | HTEFNPHNHY | HLA-B*44:02 | 10 | 0.820447 | 0.05 | 0.17709 | 100.00% (10/10) | Non-Toxin | IEDB recommended 2020.09 (NetMHCpan EL 4.1) |
| 896 | Polymerase | HTEFNPHNHY | HLA-A*26:01 | 10 | 0.67573 | 0.06 | 0.17709 | 100.00% (10/10) | Non-Toxin | IEDB recommended 2020.09 (NetMHCpan EL 4.1) |
| 897 | Polymerase | HTYKVLSNA | HLA-A*68:02 | 9 | 0.768275 | 0.06 | -0.33184 | 100.00% (10/10) | Non-Toxin | IEDB recommended 2020.09 (NetMHCpan EL 4.1) |
| 898 | Polymerase | HVDTSSRSW | HLA-B*58:01 | 9 | 0.910284 | 0.06 | -0.32355 | 100.00% (10/10) | Non-Toxin | IEDB recommended 2020.09 (NetMHCpan EL 4.1) |
| 899 | Polymerase | HVDTSSRSW | HLA-B*53:01 | 9 | 0.689189 | 0.06 | -0.32355 | 100.00% (10/10) | Non-Toxin | IEDB recommended 2020.09 (NetMHCpan EL 4.1) |
| 900 | Polymerase | IATVYTWAY | HLA-B*35:01 | 9 | 0.771632 | 0.09 | 0.29688 | 100.00% (10/10) | Non-Toxin | IEDB recommended 2020.09 (NetMHCpan EL 4.1) |
| 901 | Polymerase | IAYPECNNI | HLA-B*51:01 | 9 | 0.742431 | 0.06 | 0.02515 | 90.00% (9/10) | Non-Toxin | IEDB recommended 2020.09 (NetMHCpan EL 4.1) |
| 902 | Polymerase | IEGYSQKTW | HLA-B*44:02 | 9 | 0.930451 | 0.02 | -0.42218 | 100.00% (10/10) | Non-Toxin | IEDB recommended 2020.09 (NetMHCpan EL 4.1) |
| 903 | Polymerase | IEGYSQKTW | HLA-B*44:03 | 9 | 0.934111 | 0.03 | -0.42218 | 100.00% (10/10) | Non-Toxin | IEDB recommended 2020.09 (NetMHCpan EL 4.1) |
| 904 | Polymerase | ILNIDNIHLL | HLA-A*02:01 | 10 | 0.814141 | 0.07 | 0.29322 | 100.00% (10/10) | Non-Toxin | IEDB recommended 2020.09 (NetMHCpan EL 4.1) |
| 905 | Polymerase | ILNPNLICI | HLA-A*02:01 | 9 | 0.764226 | 0.1 | 0.05082 | 100.00% (10/10) | Non-Toxin | IEDB recommended 2020.09 (NetMHCpan EL 4.1) |
| 906 | Polymerase | ILRNSPNPMLK | HLA-A*03:01 | 11 | 0.833016 | 0.06 | -0.33303 | 100.00% (10/10) | Non-Toxin | IEDB recommended 2020.09 (NetMHCpan EL 4.1) |
| 907 | Polymerase | ILVEHSHLI | HLA-A*02:03 | 9 | 0.917118 | 0.03 | 0.01074 | 100.00% (10/10) | Non-Toxin | IEDB recommended 2020.09 (NetMHCpan EL 4.1) |
| 908 | Polymerase | ILVEHSHLI | HLA-A*02:01 | 9 | 0.908684 | 0.03 | 0.01074 | 100.00% (10/10) | Non-Toxin | IEDB recommended 2020.09 (NetMHCpan EL 4.1) |
| 909 | Polymerase | ILVEHSHLI | HLA-A*02:06 | 9 | 0.784133 | 0.08 | 0.01074 | 100.00% (10/10) | Non-Toxin | IEDB recommended 2020.09 (NetMHCpan EL 4.1) |
| 910 | Polymerase | IMKKSFKAY | HLA-B*15:01 | 9 | 0.934486 | 0.01 | -0.49704 | 100.00% (10/10) | Non-Toxin | IEDB recommended 2020.09 (NetMHCpan EL 4.1) |
| 911 | Polymerase | IMLNEAMNY | HLA-A*30:02 | 9 | 0.656456 | 0.08 | -0.02776 | 100.00% (10/10) | Non-Toxin | IEDB recommended 2020.09 (NetMHCpan EL 4.1) |
| 912 | Polymerase | IMTESVLQK | HLA-A*03:01 | 9 | 0.902445 | 0.03 | -0.08593 | 100.00% (10/10) | Non-Toxin | IEDB recommended 2020.09 (NetMHCpan EL 4.1) |
| 913 | Polymerase | IPAPIGGFNY | HLA-B*35:01 | 10 | 0.916029 | 0.04 | 0.29106 | 100.00% (10/10) | Non-Toxin | IEDB recommended 2020.09 (NetMHCpan EL 4.1) |
| 914 | Polymerase | IPAPIGGFNY | HLA-B*53:01 | 10 | 0.72893 | 0.05 | 0.29106 | 100.00% (10/10) | Non-Toxin | IEDB recommended 2020.09 (NetMHCpan EL 4.1) |
| 915 | Polymerase | IPFLFLSAY | HLA-B*35:01 | 9 | 0.962059 | 0.02 | 0.01364 | 100.00% (10/10) | Non-Toxin | IEDB recommended 2020.09 (NetMHCpan EL 4.1) |
| 916 | Polymerase | IPGQRELKL | HLA-B*08:01 | 9 | 0.699375 | 0.07 | -0.09627 | 100.00% (10/10) | Non-Toxin | IEDB recommended 2020.09 (NetMHCpan EL 4.1) |
| 917 | Polymerase | IPGQRELKL | HLA-B*07:02 | 9 | 0.780531 | 0.09 | -0.09627 | 100.00% (10/10) | Non-Toxin | IEDB recommended 2020.09 (NetMHCpan EL 4.1) |
| 918 | Polymerase | IQFDCFMEL | HLA-A*02:06 | 9 | 0.819612 | 0.07 | 0.02832 | 100.00% (10/10) | Non-Toxin | IEDB recommended 2020.09 (NetMHCpan EL 4.1) |
| 919 | Polymerase | ISRLFNMYR | HLA-A*31:01 | 9 | 0.796194 | 0.07 | -0.03681 | 100.00% (10/10) | Non-Toxin | IEDB recommended 2020.09 (NetMHCpan EL 4.1) |
| 920 | Polymerase | ITSQGMTSK | HLA-A*11:01 | 9 | 0.858372 | 0.04 | -0.36646 | 100.00% (10/10) | Non-Toxin | IEDB recommended 2020.09 (NetMHCpan EL 4.1) |
| 921 | Polymerase | ITSQGMTSK | HLA-A*03:01 | 9 | 0.790546 | 0.09 | -0.36646 | 100.00% (10/10) | Non-Toxin | IEDB recommended 2020.09 (NetMHCpan EL 4.1) |
| 922 | Polymerase | IYGLPGFFNW | HLA-A*23:01 | 10 | 0.894838 | 0.02 | 0.22726 | 100.00% (10/10) | Non-Toxin | IEDB recommended 2020.09 (NetMHCpan EL 4.1) |
| 923 | Polymerase | IYGLPGFFNW | HLA-A*24:02 | 10 | 0.873025 | 0.04 | 0.22726 | 100.00% (10/10) | Non-Toxin | IEDB recommended 2020.09 (NetMHCpan EL 4.1) |
| 924 | Polymerase | KALNLSPLIQR | HLA-A*31:01 | 11 | 0.846067 | 0.05 | -0.15861 | 100.00% (10/10) | Non-Toxin | IEDB recommended 2020.09 (NetMHCpan EL 4.1) |
| 925 | Polymerase | KALSPIKDEW | HLA-B*57:01 | 10 | 0.991766 | 0.01 | -0.17705 | 100.00% (10/10) | Non-Toxin | IEDB recommended 2020.09 (NetMHCpan EL 4.1) |
| 926 | Polymerase | KALSPIKDEW | HLA-B*58:01 | 10 | 0.983327 | 0.01 | -0.17705 | 100.00% (10/10) | Non-Toxin | IEDB recommended 2020.09 (NetMHCpan EL 4.1) |

| | | | | | | | | | | |
|---|---|---|---|---|---|---|---|---|---|---|
| 927 | Polymerase | KAYNIISKK | HLA-A*03:01 | 9 | 0.975367 | 0.01 | -0.01845 | 100.00% (10/10) | Non-Toxin | IEDB recommended 2020.09 (NetMHCpan EL 4.1) |
| 928 | Polymerase | KAYNIISKK | HLA-A*11:01 | 9 | 0.94852 | 0.01 | -0.01845 | 100.00% (10/10) | Non-Toxin | IEDB recommended 2020.09 (NetMHCpan EL 4.1) |
| 929 | Polymerase | KAYNIISKK | HLA-A*30:01 | 9 | 0.878783 | 0.01 | -0.01845 | 100.00% (10/10) | Non-Toxin | IEDB recommended 2020.09 (NetMHCpan EL 4.1) |
| 930 | Polymerase | KEICAKQAQLY | HLA-B*44:03 | 11 | 0.847747 | 0.06 | -0.36316 | 100.00% (10/10) | Non-Toxin | IEDB recommended 2020.09 (NetMHCpan EL 4.1) |
| 931 | Polymerase | KEICAKQAQLY | HLA-B*44:02 | 11 | 0.771193 | 0.07 | -0.36316 | 100.00% (10/10) | Non-Toxin | IEDB recommended 2020.09 (NetMHCpan EL 4.1) |
| 932 | Polymerase | KETKQAGRLF | HLA-B*44:02 | 10 | 0.698587 | 0.1 | -0.21 | 100.00% (10/10) | Non-Toxin | IEDB recommended 2020.09 (NetMHCpan EL 4.1) |
| 933 | Polymerase | KFAGSVLNR | HLA-A*31:01 | 9 | 0.861168 | 0.04 | -0.08858 | 100.00% (10/10) | Non-Toxin | IEDB recommended 2020.09 (NetMHCpan EL 4.1) |
| 934 | Polymerase | KFKDTKSSELY | HLA-A*30:02 | 11 | 0.653727 | 0.08 | -0.45869 | 100.00% (10/10) | Non-Toxin | IEDB recommended 2020.09 (NetMHCpan EL 4.1) |
| 935 | Polymerase | KGLIRSGLRK | HLA-A*03:01 | 10 | 0.833249 | 0.06 | 0.0724 | 100.00% (10/10) | Non-Toxin | IEDB recommended 2020.09 (NetMHCpan EL 4.1) |
| 936 | Polymerase | KIWWKIIGY | HLA-A*32:01 | 9 | 0.723173 | 0.03 | 0.34219 | 100.00% (10/10) | Non-Toxin | IEDB recommended 2020.09 (NetMHCpan EL 4.1) |
| 937 | Polymerase | KIWWKIIGY | HLA-A*30:02 | 9 | 0.6134 | 0.1 | 0.34219 | 100.00% (10/10) | Non-Toxin | IEDB recommended 2020.09 (NetMHCpan EL 4.1) |
| 938 | Polymerase | KKYYQIDQPF | HLA-A*23:01 | 10 | 0.636779 | 0.1 | -0.07148 | 100.00% (10/10) | Non-Toxin | IEDB recommended 2020.09 (NetMHCpan EL 4.1) |
| 939 | Polymerase | KLDSDLSMY | HLA-A*01:01 | 9 | 0.923022 | 0.03 | -0.39033 | 100.00% (10/10) | Non-Toxin | IEDB recommended 2020.09 (NetMHCpan EL 4.1) |
| 940 | Polymerase | KLDSDLSMY | HLA-A*30:02 | 9 | 0.715847 | 0.05 | -0.39033 | 100.00% (10/10) | Non-Toxin | IEDB recommended 2020.09 (NetMHCpan EL 4.1) |
| 941 | Polymerase | KLFPSEYSI | HLA-A*32:01 | 9 | 0.910285 | 0.01 | -0.13979 | 100.00% (10/10) | Non-Toxin | IEDB recommended 2020.09 (NetMHCpan EL 4.1) |
| 942 | Polymerase | KLFPSEYSI | HLA-A*02:01 | 9 | 0.956367 | 0.02 | -0.13979 | 100.00% (10/10) | Non-Toxin | IEDB recommended 2020.09 (NetMHCpan EL 4.1) |
| 943 | Polymerase | KLFPSEYSI | HLA-A*02:06 | 9 | 0.881674 | 0.05 | -0.13979 | 100.00% (10/10) | Non-Toxin | IEDB recommended 2020.09 (NetMHCpan EL 4.1) |
| 944 | Polymerase | KLFPSEYSI | HLA-A*02:03 | 9 | 0.832089 | 0.05 | -0.13979 | 100.00% (10/10) | Non-Toxin | IEDB recommended 2020.09 (NetMHCpan EL 4.1) |
| 945 | Polymerase | KLISAIEYA | HLA-A*02:01 | 9 | 0.820528 | 0.07 | 0.12245 | 100.00% (10/10) | Non-Toxin | IEDB recommended 2020.09 (NetMHCpan EL 4.1) |
| 946 | Polymerase | KLISAIEYA | HLA-A*02:06 | 9 | 0.80208 | 0.07 | 0.12245 | 100.00% (10/10) | Non-Toxin | IEDB recommended 2020.09 (NetMHCpan EL 4.1) |
| 947 | Polymerase | KLISAIEYA | HLA-A*02:03 | 9 | 0.775657 | 0.07 | 0.12245 | 100.00% (10/10) | Non-Toxin | IEDB recommended 2020.09 (NetMHCpan EL 4.1) |
| 948 | Polymerase | KLKGLVVPL | HLA-A*02:03 | 9 | 0.740366 | 0.08 | 0.02052 | 100.00% (10/10) | Non-Toxin | IEDB recommended 2020.09 (NetMHCpan EL 4.1) |
| 949 | Polymerase | KLMTKTLPK | HLA-A*03:01 | 9 | 0.974443 | 0.01 | -0.20724 | 100.00% (10/10) | Non-Toxin | IEDB recommended 2020.09 (NetMHCpan EL 4.1) |
| 950 | Polymerase | KLMTKTLPK | HLA-A*30:01 | 9 | 0.769015 | 0.02 | -0.20724 | 100.00% (10/10) | Non-Toxin | IEDB recommended 2020.09 (NetMHCpan EL 4.1) |
| 951 | Polymerase | KQAGRLFAK | HLA-A*30:01 | 9 | 0.662612 | 0.06 | 0.20842 | 100.00% (10/10) | Non-Toxin | IEDB recommended 2020.09 (NetMHCpan EL 4.1) |
| 952 | Polymerase | KQAGRLFAK | HLA-A*03:01 | 9 | 0.817215 | 0.07 | 0.20842 | 100.00% (10/10) | Non-Toxin | IEDB recommended 2020.09 (NetMHCpan EL 4.1) |
| 953 | Polymerase | KQAQLYFER | HLA-A*31:01 | 9 | 0.888244 | 0.03 | 0.03916 | 100.00% (10/10) | Non-Toxin | IEDB recommended 2020.09 (NetMHCpan EL 4.1) |
| 954 | Polymerase | KQFHDDLKKY | HLA-B*15:01 | 10 | 0.880014 | 0.03 | -0.20469 | 100.00% (10/10) | Non-Toxin | IEDB recommended 2020.09 (NetMHCpan EL 4.1) |
| 955 | Polymerase | KQFHDDLKKY | HLA-A*30:02 | 10 | 0.743434 | 0.04 | -0.20469 | 100.00% (10/10) | Non-Toxin | IEDB recommended 2020.09 (NetMHCpan EL 4.1) |
| 956 | Polymerase | KQFHDDLKKYY | HLA-A*30:02 | 11 | 0.732901 | 0.04 | -0.28421 | 100.00% (10/10) | Non-Toxin | IEDB recommended 2020.09 (NetMHCpan EL 4.1) |
| 957 | Polymerase | KQFHDDLKKYY | HLA-B*15:01 | 11 | 0.752747 | 0.1 | -0.28421 | 100.00% (10/10) | Non-Toxin | IEDB recommended 2020.09 (NetMHCpan EL 4.1) |
| 958 | Polymerase | KSDKTGKKCY | HLA-A*01:01 | 10 | 0.673985 | 0.1 | -0.5555 | 70.00% (7/10) | Non-Toxin | IEDB recommended 2020.09 (NetMHCpan EL 4.1) |
| 959 | Polymerase | KSDKTGKKCYY | HLA-A*01:01 | 11 | 0.759146 | 0.08 | -0.59966 | 70.00% (7/10) | Non-Toxin | IEDB recommended 2020.09 (NetMHCpan EL 4.1) |
| 960 | Polymerase | KSFLDYHTEF | HLA-B*57:01 | 10 | 0.939633 | 0.07 | 0.16655 | 100.00% (10/10) | Non-Toxin | IEDB recommended 2020.09 (NetMHCpan EL 4.1) |
| 961 | Polymerase | KSFLDYHTEF | HLA-B*58:01 | 10 | 0.836987 | 0.09 | 0.16655 | 100.00% (10/10) | Non-Toxin | IEDB recommended 2020.09 (NetMHCpan EL 4.1) |
| 962 | Polymerase | KSGLQPKLVSR | HLA-A*31:01 | 11 | 0.908371 | 0.02 | -0.40602 | 90.00% (9/10) | Non-Toxin | IEDB recommended 2020.09 (NetMHCpan EL 4.1) |
| 963 | Polymerase | KSRELDFPLW | HLA-B*57:01 | 10 | 0.994996 | 0.01 | 0.22271 | 100.00% (10/10) | Non-Toxin | IEDB recommended 2020.09 (NetMHCpan EL 4.1) |
| 964 | Polymerase | KSRELDFPLW | HLA-B*58:01 | 10 | 0.968245 | 0.02 | 0.22271 | 100.00% (10/10) | Non-Toxin | IEDB recommended 2020.09 (NetMHCpan EL 4.1) |
| 965 | Polymerase | KSSELYHIK | HLA-A*30:01 | 9 | 0.729676 | 0.04 | 0.13783 | 100.00% (10/10) | Non-Toxin | IEDB recommended 2020.09 (NetMHCpan EL 4.1) |
| 966 | Polymerase | KSSELYHIK | HLA-A*11:01 | 9 | 0.805859 | 0.08 | 0.13783 | 100.00% (10/10) | Non-Toxin | IEDB recommended 2020.09 (NetMHCpan EL 4.1) |
| 967 | Polymerase | KTIKNITAR | HLA-A*31:01 | 9 | 0.968599 | 0.01 | 0.0008 | 100.00% (10/10) | Non-Toxin | IEDB recommended 2020.09 (NetMHCpan EL 4.1) |
| 968 | Polymerase | KTIKNITAR | HLA-A*11:01 | 9 | 0.822354 | 0.06 | 0.0008 | 100.00% (10/10) | Non-Toxin | IEDB recommended 2020.09 (NetMHCpan EL 4.1) |
| 969 | Polymerase | KTLPKGMQER | HLA-A*31:01 | 10 | 0.929844 | 0.01 | -0.39632 | 100.00% (10/10) | Non-Toxin | IEDB recommended 2020.09 (NetMHCpan EL 4.1) |
| 970 | Polymerase | KTLPKGMQERR | HLA-A*31:01 | 11 | 0.908707 | 0.02 | -0.35706 | 100.00% (10/10) | Non-Toxin | IEDB recommended 2020.09 (NetMHCpan EL 4.1) |
| 971 | Polymerase | KTPEDDIFIHY | HLA-A*01:01 | 11 | 0.915235 | 0.03 | 0.51137 | 90.00% (9/10) | Non-Toxin | IEDB recommended 2020.09 (NetMHCpan EL 4.1) |
| 972 | Polymerase | KVHPNLPYK | HLA-A*03:01 | 9 | 0.983067 | 0.01 | -0.02892 | 100.00% (10/10) | Non-Toxin | IEDB recommended 2020.09 (NetMHCpan EL 4.1) |
| 973 | Polymerase | KVHPNLPYK | HLA-A*30:01 | 9 | 0.9816 | 0.01 | -0.02892 | 100.00% (10/10) | Non-Toxin | IEDB recommended 2020.09 (NetMHCpan EL 4.1) |
| 974 | Polymerase | KVHPNLPYK | HLA-A*11:01 | 9 | 0.975148 | 0.01 | -0.02892 | 100.00% (10/10) | Non-Toxin | IEDB recommended 2020.09 (NetMHCpan EL 4.1) |
| 975 | Polymerase | KVHPNLPYK | HLA-A*31:01 | 9 | 0.810218 | 0.06 | -0.02892 | 100.00% (10/10) | Non-Toxin | IEDB recommended 2020.09 (NetMHCpan EL 4.1) |
| 976 | Polymerase | KVHPNLPYKVK | HLA-A*30:01 | 11 | 0.797072 | 0.02 | -0.18992 | 100.00% (10/10) | Non-Toxin | IEDB recommended 2020.09 (NetMHCpan EL 4.1) |
| 977 | Polymerase | KVHPNLPYKVK | HLA-A*03:01 | 11 | 0.786951 | 0.1 | -0.18992 | 100.00% (10/10) | Non-Toxin | IEDB recommended 2020.09 (NetMHCpan EL 4.1) |

| | | | | | | | | | | |
|---|---|---|---|---|---|---|---|---|---|---|
| 978 | Polymerase | KVIGMLQNI | HLA-A*32:01 | 9 | 0.655429 | 0.05 | -0.20568 | 100.00% (10/10) | Non-Toxin | IEDB recommended 2020.09 (NetMHCpan EL 4.1) |
| 979 | Polymerase | KVIGMLQNI | HLA-A*02:06 | 9 | 0.858934 | 0.06 | -0.20568 | 100.00% (10/10) | Non-Toxin | IEDB recommended 2020.09 (NetMHCpan EL 4.1) |
| 980 | Polymerase | KVIGMLQNITR | HLA-A*31:01 | 11 | 0.800345 | 0.06 | -0.08839 | 100.00% (10/10) | Non-Toxin | IEDB recommended 2020.09 (NetMHCpan EL 4.1) |
| 981 | Polymerase | KVKKEICAK | HLA-A*30:01 | 9 | 0.803901 | 0.02 | -0.08686 | 100.00% (10/10) | Non-Toxin | IEDB recommended 2020.09 (NetMHCpan EL 4.1) |
| 982 | Polymerase | KVKKEICAK | HLA-A*03:01 | 9 | 0.783982 | 0.1 | -0.08686 | 100.00% (10/10) | Non-Toxin | IEDB recommended 2020.09 (NetMHCpan EL 4.1) |
| 983 | Polymerase | KVLEYAPIMK | HLA-A*03:01 | 10 | 0.933888 | 0.02 | 0.13093 | 100.00% (10/10) | Non-Toxin | IEDB recommended 2020.09 (NetMHCpan EL 4.1) |
| 984 | Polymerase | KVLEYAPIMK | HLA-A*11:01 | 10 | 0.908142 | 0.02 | 0.13093 | 100.00% (10/10) | Non-Toxin | IEDB recommended 2020.09 (NetMHCpan EL 4.1) |
| 985 | Polymerase | KVREHMLADK | HLA-A*30:01 | 10 | 0.709284 | 0.04 | 0.01359 | 100.00% (10/10) | Non-Toxin | IEDB recommended 2020.09 (NetMHCpan EL 4.1) |
| 986 | Polymerase | KVREHMLADK | HLA-A*03:01 | 10 | 0.789649 | 0.09 | 0.01359 | 100.00% (10/10) | Non-Toxin | IEDB recommended 2020.09 (NetMHCpan EL 4.1) |
| 987 | Polymerase | KWAFEIHHRR | HLA-A*31:01 | 10 | 0.833449 | 0.05 | 0.44559 | 100.00% (10/10) | Non-Toxin | IEDB recommended 2020.09 (NetMHCpan EL 4.1) |
| 988 | Polymerase | KWYECFLFW | HLA-A*23:01 | 9 | 0.643792 | 0.1 | 0.21629 | 100.00% (10/10) | Non-Toxin | IEDB recommended 2020.09 (NetMHCpan EL 4.1) |
| 989 | Polymerase | KYFKENGMVK | HLA-A*30:01 | 10 | 0.576802 | 0.09 | -0.17998 | 100.00% (10/10) | Non-Toxin | IEDB recommended 2020.09 (NetMHCpan EL 4.1) |
| 990 | Polymerase | KYYQIDQPF | HLA-A*23:01 | 9 | 0.949346 | 0.01 | -0.07152 | 100.00% (10/10) | Non-Toxin | IEDB recommended 2020.09 (NetMHCpan EL 4.1) |
| 991 | Polymerase | KYYQIDQPF | HLA-A*24:02 | 9 | 0.942987 | 0.01 | -0.07152 | 100.00% (10/10) | Non-Toxin | IEDB recommended 2020.09 (NetMHCpan EL 4.1) |
| 992 | Polymerase | KYYQIDQPFF | HLA-A*23:01 | 10 | 0.941388 | 0.01 | -0.01656 | 100.00% (10/10) | Non-Toxin | IEDB recommended 2020.09 (NetMHCpan EL 4.1) |
| 993 | Polymerase | KYYQIDQPFF | HLA-A*24:02 | 10 | 0.930055 | 0.02 | -0.01656 | 100.00% (10/10) | Non-Toxin | IEDB recommended 2020.09 (NetMHCpan EL 4.1) |
| 994 | Polymerase | KYYQIDQPFFV | HLA-A*02:06 | 11 | 0.743572 | 0.09 | 0.0774 | 100.00% (10/10) | Non-Toxin | IEDB recommended 2020.09 (NetMHCpan EL 4.1) |
| 995 | Polymerase | LADKVLEY | HLA-A*01:01 | 8 | 0.722731 | 0.09 | -0.09554 | 100.00% (10/10) | Non-Toxin | IEDB recommended 2020.09 (NetMHCpan EL 4.1) |
| 996 | Polymerase | LAKTVAQTV | HLA-B*51:01 | 9 | 0.795097 | 0.04 | -0.02899 | 100.00% (10/10) | Non-Toxin | IEDB recommended 2020.09 (NetMHCpan EL 4.1) |
| 997 | Polymerase | LARALRSHMW | HLA-B*57:01 | 10 | 0.923896 | 0.09 | -0.13526 | 100.00% (10/10) | Non-Toxin | IEDB recommended 2020.09 (NetMHCpan EL 4.1) |
| 998 | Polymerase | LETDDYNGIY | HLA-A*01:01 | 10 | 0.931807 | 0.02 | 0.15319 | 100.00% (10/10) | Non-Toxin | IEDB recommended 2020.09 (NetMHCpan EL 4.1) |
| 999 | Polymerase | LGQSISIKW | HLA-B*58:01 | 9 | 0.932578 | 0.04 | -0.24388 | 100.00% (10/10) | Non-Toxin | IEDB recommended 2020.09 (NetMHCpan EL 4.1) |
| 1000 | Polymerase | LGQSISIKW | HLA-B*57:01 | 9 | 0.94388 | 0.07 | -0.24388 | 100.00% (10/10) | Non-Toxin | IEDB recommended 2020.09 (NetMHCpan EL 4.1) |
| 1001 | Polymerase | LIASGVGKY | HLA-A*26:01 | 9 | 0.58391 | 0.09 | -0.17931 | 100.00% (10/10) | Non-Toxin | IEDB recommended 2020.09 (NetMHCpan EL 4.1) |
| 1002 | Polymerase | LISRSNALW | HLA-B*58:01 | 9 | 0.894877 | 0.06 | -0.14227 | 100.00% (10/10) | Non-Toxin | IEDB recommended 2020.09 (NetMHCpan EL 4.1) |
| 1003 | Polymerase | LKLDSDLSMY | HLA-A*01:01 | 10 | 0.84994 | 0.05 | -0.37344 | 100.00% (10/10) | Non-Toxin | IEDB recommended 2020.09 (NetMHCpan EL 4.1) |
| 1004 | Polymerase | LPAHASKHI | HLA-B*51:01 | 9 | 0.765138 | 0.05 | -0.23548 | 90.00% (9/10) | Non-Toxin | IEDB recommended 2020.09 (NetMHCpan EL 4.1) |
| 1005 | Polymerase | LPIPEYTEV | HLA-B*51:01 | 9 | 0.976165 | 0.01 | 0.21732 | 100.00% (10/10) | Non-Toxin | IEDB recommended 2020.09 (NetMHCpan EL 4.1) |
| 1006 | Polymerase | LPIPEYTEV | HLA-B*53:01 | 9 | 0.570883 | 0.09 | 0.21732 | 100.00% (10/10) | Non-Toxin | IEDB recommended 2020.09 (NetMHCpan EL 4.1) |
| 1007 | Polymerase | LPSGAQRLF | HLA-B*53:01 | 9 | 0.875556 | 0.03 | -0.05334 | 100.00% (10/10) | Non-Toxin | IEDB recommended 2020.09 (NetMHCpan EL 4.1) |
| 1008 | Polymerase | LPSGAQRLF | HLA-B*35:01 | 9 | 0.893121 | 0.05 | -0.05334 | 100.00% (10/10) | Non-Toxin | IEDB recommended 2020.09 (NetMHCpan EL 4.1) |
| 1009 | Polymerase | LPYKVKKEI | HLA-B*51:01 | 9 | 0.950014 | 0.01 | -0.5045 | 100.00% (10/10) | Non-Toxin | IEDB recommended 2020.09 (NetMHCpan EL 4.1) |
| 1010 | Polymerase | LPYKVKKEI | HLA-B*07:02 | 9 | 0.776658 | 0.09 | -0.5045 | 100.00% (10/10) | Non-Toxin | IEDB recommended 2020.09 (NetMHCpan EL 4.1) |
| 1011 | Polymerase | LSHPRVFKR | HLA-A*31:01 | 9 | 0.915212 | 0.02 | 0.0614 | 100.00% (10/10) | Non-Toxin | IEDB recommended 2020.09 (NetMHCpan EL 4.1) |
| 1012 | Polymerase | LSHPRVFKR | HLA-A*33:01 | 9 | 0.727337 | 0.06 | 0.0614 | 100.00% (10/10) | Non-Toxin | IEDB recommended 2020.09 (NetMHCpan EL 4.1) |
| 1013 | Polymerase | LSNREVKIW | HLA-B*57:01 | 9 | 0.991806 | 0.01 | 0.0821 | 100.00% (10/10) | Non-Toxin | IEDB recommended 2020.09 (NetMHCpan EL 4.1) |
| 1014 | Polymerase | LSNREVKIW | HLA-B*58:01 | 9 | 0.980314 | 0.01 | 0.0821 | 100.00% (10/10) | Non-Toxin | IEDB recommended 2020.09 (NetMHCpan EL 4.1) |
| 1015 | Polymerase | LSNREVKIWW | HLA-B*57:01 | 10 | 0.95387 | 0.06 | 0.22642 | 100.00% (10/10) | Non-Toxin | IEDB recommended 2020.09 (NetMHCpan EL 4.1) |
| 1016 | Polymerase | LTIDDCVKNW | HLA-B*57:01 | 10 | 0.965981 | 0.05 | -0.1123 | 100.00% (10/10) | Non-Toxin | IEDB recommended 2020.09 (NetMHCpan EL 4.1) |
| 1017 | Polymerase | LTIDDCVKNW | HLA-B*58:01 | 10 | 0.914447 | 0.05 | -0.1123 | 100.00% (10/10) | Non-Toxin | IEDB recommended 2020.09 (NetMHCpan EL 4.1) |
| 1018 | Polymerase | LTPEMVLMY | HLA-A*01:01 | 9 | 0.786954 | 0.07 | -0.14695 | 100.00% (10/10) | Non-Toxin | IEDB recommended 2020.09 (NetMHCpan EL 4.1) |
| 1019 | Polymerase | LTPEMVLMY | HLA-A*26:01 | 9 | 0.633998 | 0.07 | -0.14695 | 100.00% (10/10) | Non-Toxin | IEDB recommended 2020.09 (NetMHCpan EL 4.1) |
| 1020 | Polymerase | LWGLIDPLF | HLA-A*23:01 | 9 | 0.652191 | 0.1 | 0.13448 | 100.00% (10/10) | Non-Toxin | IEDB recommended 2020.09 (NetMHCpan EL 4.1) |
| 1021 | Polymerase | LYLGQSISI | HLA-A*24:02 | 9 | 0.716974 | 0.09 | -0.22237 | 100.00% (10/10) | Non-Toxin | IEDB recommended 2020.09 (NetMHCpan EL 4.1) |
| 1022 | Polymerase | MLADKVLEY | HLA-B*15:01 | 9 | 0.949274 | 0.01 | -0.08698 | 100.00% (10/10) | Non-Toxin | IEDB recommended 2020.09 (NetMHCpan EL 4.1) |
| 1023 | Polymerase | MLADKVLEY | HLA-A*30:02 | 9 | 0.808511 | 0.02 | -0.08698 | 100.00% (10/10) | Non-Toxin | IEDB recommended 2020.09 (NetMHCpan EL 4.1) |
| 1024 | Polymerase | MLADKVLEY | HLA-A*01:01 | 9 | 0.844751 | 0.05 | -0.08698 | 100.00% (10/10) | Non-Toxin | IEDB recommended 2020.09 (NetMHCpan EL 4.1) |
| 1025 | Polymerase | MLADKVLEY | HLA-A*26:01 | 9 | 0.754185 | 0.05 | -0.08698 | 100.00% (10/10) | Non-Toxin | IEDB recommended 2020.09 (NetMHCpan EL 4.1) |
| 1026 | Polymerase | MLADKVLEY | HLA-B*35:01 | 9 | 0.83305 | 0.07 | -0.08698 | 100.00% (10/10) | Non-Toxin | IEDB recommended 2020.09 (NetMHCpan EL 4.1) |
| 1027 | Polymerase | MLADKVLEYA | HLA-A*02:03 | 10 | 0.694179 | 0.1 | -0.06288 | 100.00% (10/10) | Non-Toxin | IEDB recommended 2020.09 (NetMHCpan EL 4.1) |
| 1028 | Polymerase | MTESVLQKV | HLA-A*68:02 | 9 | 0.716639 | 0.08 | -0.32797 | 100.00% (10/10) | Non-Toxin | IEDB recommended 2020.09 (NetMHCpan EL 4.1) |

| | | | | | | | | | | |
|---|---|---|---|---|---|---|---|---|---|---|
| 1029 | Polymerase | MYCDVLEGR | HLA-A*33:01 | 9 | 0.659634 | 0.08 | 0.13888 | 100.00% (10/10) | Non-Toxin | IEDB recommended 2020.09 (NetMHCpan EL 4.1) |
| 1030 | Polymerase | NIRRKVLIL | HLA-B*08:01 | 9 | 0.94942 | 0.01 | -0.03386 | 100.00% (10/10) | Non-Toxin | IEDB recommended 2020.09 (NetMHCpan EL 4.1) |
| 1031 | Polymerase | NITARTILR | HLA-A*33:01 | 9 | 0.688272 | 0.08 | 0.24475 | 100.00% (10/10) | Non-Toxin | IEDB recommended 2020.09 (NetMHCpan EL 4.1) |
| 1032 | Polymerase | NLHGKRKSL | HLA-B*08:01 | 9 | 0.949676 | 0.01 | -0.39534 | 100.00% (10/10) | Non-Toxin | IEDB recommended 2020.09 (NetMHCpan EL 4.1) |
| 1033 | Polymerase | NLSPLIQRY | HLA-A*30:02 | 9 | 0.779952 | 0.03 | -0.0179 | 100.00% (10/10) | Non-Toxin | IEDB recommended 2020.09 (NetMHCpan EL 4.1) |
| 1034 | Polymerase | NLSPLIQRY | HLA-A*26:01 | 9 | 0.642122 | 0.07 | -0.0179 | 100.00% (10/10) | Non-Toxin | IEDB recommended 2020.09 (NetMHCpan EL 4.1) |
| 1035 | Polymerase | NPDSTEVYL | HLA-B*53:01 | 9 | 0.584975 | 0.09 | 0.00546 | 100.00% (10/10) | Non-Toxin | IEDB recommended 2020.09 (NetMHCpan EL 4.1) |
| 1036 | Polymerase | NPPIKTGVL | HLA-B*08:01 | 9 | 0.592548 | 0.1 | 0.00958 | 100.00% (10/10) | Non-Toxin | IEDB recommended 2020.09 (NetMHCpan EL 4.1) |
| 1037 | Polymerase | NPQEKICVL | HLA-B*08:01 | 9 | 0.884446 | 0.02 | -0.04295 | 100.00% (10/10) | Non-Toxin | IEDB recommended 2020.09 (NetMHCpan EL 4.1) |
| 1038 | Polymerase | NTMYGWFFV | HLA-A*68:02 | 9 | 0.656525 | 0.1 | 0.34799 | 100.00% (10/10) | Non-Toxin | IEDB recommended 2020.09 (NetMHCpan EL 4.1) |
| 1039 | Polymerase | NTSDLDFVIFY | HLA-A*01:01 | 11 | 0.704239 | 0.1 | 0.313 | 100.00% (10/10) | Non-Toxin | IEDB recommended 2020.09 (NetMHCpan EL 4.1) |
| 1040 | Polymerase | NTSKHTYKV | HLA-A*68:02 | 9 | 0.869929 | 0.03 | -0.33178 | 100.00% (10/10) | Non-Toxin | IEDB recommended 2020.09 (NetMHCpan EL 4.1) |
| 1041 | Polymerase | NTSKSFLDY | HLA-A*01:01 | 9 | 0.836734 | 0.05 | -0.318 | 100.00% (10/10) | Non-Toxin | IEDB recommended 2020.09 (NetMHCpan EL 4.1) |
| 1042 | Polymerase | NVKRPTKAL | HLA-B*08:01 | 9 | 0.746773 | 0.05 | -0.15132 | 100.00% (10/10) | Non-Toxin | IEDB recommended 2020.09 (NetMHCpan EL 4.1) |
| 1043 | Polymerase | NVKRPTKALR | HLA-A*33:01 | 10 | 0.704769 | 0.07 | -0.16738 | 100.00% (10/10) | Non-Toxin | IEDB recommended 2020.09 (NetMHCpan EL 4.1) |
| 1044 | Polymerase | NYLNLSRIF | HLA-A*23:01 | 9 | 0.864851 | 0.03 | -0.0552 | 100.00% (10/10) | Non-Toxin | IEDB recommended 2020.09 (NetMHCpan EL 4.1) |
| 1045 | Polymerase | NYLNLSRIF | HLA-A*24:02 | 9 | 0.871082 | 0.04 | -0.0552 | 100.00% (10/10) | Non-Toxin | IEDB recommended 2020.09 (NetMHCpan EL 4.1) |
| 1046 | Polymerase | QESKSRELDF | HLA-B*44:02 | 10 | 0.695373 | 0.1 | -0.28355 | 100.00% (10/10) | Non-Toxin | IEDB recommended 2020.09 (NetMHCpan EL 4.1) |
| 1047 | Polymerase | QPFFVPTKI | HLA-B*51:01 | 9 | 0.952773 | 0.01 | 0.09232 | 100.00% (10/10) | Non-Toxin | IEDB recommended 2020.09 (NetMHCpan EL 4.1) |
| 1048 | Polymerase | QPFFVPTKI | HLA-B*53:01 | 9 | 0.59665 | 0.08 | 0.09232 | 100.00% (10/10) | Non-Toxin | IEDB recommended 2020.09 (NetMHCpan EL 4.1) |
| 1049 | Polymerase | QPLISRSNAL | HLA-B*07:02 | 10 | 0.91172 | 0.04 | -0.11871 | 100.00% (10/10) | Non-Toxin | IEDB recommended 2020.09 (NetMHCpan EL 4.1) |
| 1050 | Polymerase | QQLLISTEF | HLA-B*15:01 | 9 | 0.881393 | 0.03 | 0.05037 | 100.00% (10/10) | Non-Toxin | IEDB recommended 2020.09 (NetMHCpan EL 4.1) |
| 1051 | Polymerase | QSISIKWAF | HLA-A*32:01 | 9 | 0.558811 | 0.07 | 0.01313 | 100.00% (10/10) | Non-Toxin | IEDB recommended 2020.09 (NetMHCpan EL 4.1) |
| 1052 | Polymerase | QSISIKWAF | HLA-B*57:01 | 9 | 0.919057 | 0.09 | 0.01313 | 100.00% (10/10) | Non-Toxin | IEDB recommended 2020.09 (NetMHCpan EL 4.1) |
| 1053 | Polymerase | QSISIKWAF | HLA-B*58:01 | 9 | 0.849445 | 0.09 | 0.01313 | 100.00% (10/10) | Non-Toxin | IEDB recommended 2020.09 (NetMHCpan EL 4.1) |
| 1054 | Polymerase | QTVLEIITK | HLA-A*11:01 | 9 | 0.789251 | 0.08 | 0.36002 | 100.00% (10/10) | Non-Toxin | IEDB recommended 2020.09 (NetMHCpan EL 4.1) |
| 1055 | Polymerase | QVDAELPIPEY | HLA-A*01:01 | 11 | 0.688061 | 0.1 | 0.29689 | 100.00% (10/10) | Non-Toxin | IEDB recommended 2020.09 (NetMHCpan EL 4.1) |
| 1056 | Polymerase | RALRSHMW | HLA-B*57:01 | 8 | 0.913389 | 0.09 | -0.23037 | 100.00% (10/10) | Non-Toxin | IEDB recommended 2020.09 (NetMHCpan EL 4.1) |
| 1057 | Polymerase | RALRSHMW | HLA-B*58:01 | 8 | 0.836902 | 0.09 | -0.23037 | 100.00% (10/10) | Non-Toxin | IEDB recommended 2020.09 (NetMHCpan EL 4.1) |
| 1058 | Polymerase | RALRSHMWR | HLA-A*31:01 | 9 | 0.753547 | 0.08 | -0.10095 | 100.00% (10/10) | Non-Toxin | IEDB recommended 2020.09 (NetMHCpan EL 4.1) |
| 1059 | Polymerase | RAVIKNSQK | HLA-A*30:01 | 9 | 0.591082 | 0.08 | -0.27607 | 100.00% (10/10) | Non-Toxin | IEDB recommended 2020.09 (NetMHCpan EL 4.1) |
| 1060 | Polymerase | REHSSIRVPY | HLA-B*44:02 | 10 | 0.791683 | 0.05 | -0.11039 | 100.00% (10/10) | Non-Toxin | IEDB recommended 2020.09 (NetMHCpan EL 4.1) |
| 1061 | Polymerase | REHSSIRVPY | HLA-B*44:03 | 10 | 0.82941 | 0.06 | -0.11039 | 100.00% (10/10) | Non-Toxin | IEDB recommended 2020.09 (NetMHCpan EL 4.1) |
| 1062 | Polymerase | REKNGFKEVW | HLA-B*44:02 | 10 | 0.893812 | 0.03 | -0.02389 | 90.00% (9/10) | Non-Toxin | IEDB recommended 2020.09 (NetMHCpan EL 4.1) |
| 1063 | Polymerase | REKNGFKEVW | HLA-B*44:03 | 10 | 0.877629 | 0.04 | -0.02389 | 90.00% (9/10) | Non-Toxin | IEDB recommended 2020.09 (NetMHCpan EL 4.1) |
| 1064 | Polymerase | RELALGRVI | HLA-B*40:01 | 9 | 0.80287 | 0.1 | 0.12467 | 100.00% (10/10) | Non-Toxin | IEDB recommended 2020.09 (NetMHCpan EL 4.1) |
| 1065 | Polymerase | RELKLFPSEY | HLA-B*44:03 | 10 | 0.80047 | 0.08 | -0.20896 | 100.00% (10/10) | Non-Toxin | IEDB recommended 2020.09 (NetMHCpan EL 4.1) |
| 1066 | Polymerase | RELKLFPSEY | HLA-B*44:02 | 10 | 0.701365 | 0.1 | -0.20896 | 100.00% (10/10) | Non-Toxin | IEDB recommended 2020.09 (NetMHCpan EL 4.1) |
| 1067 | Polymerase | RIATVYTW | HLA-A*32:01 | 8 | 0.515644 | 0.1 | 0.12124 | 100.00% (10/10) | Non-Toxin | IEDB recommended 2020.09 (NetMHCpan EL 4.1) |
| 1068 | Polymerase | RIYNIVSMI | HLA-A*32:01 | 9 | 0.63658 | 0.05 | -0.08147 | 100.00% (10/10) | Non-Toxin | IEDB recommended 2020.09 (NetMHCpan EL 4.1) |
| 1069 | Polymerase | RLETDDYNGIY | HLA-A*01:01 | 11 | 0.841886 | 0.05 | 0.21143 | 100.00% (10/10) | Non-Toxin | IEDB recommended 2020.09 (NetMHCpan EL 4.1) |
| 1070 | Polymerase | RLFAKMTYK | HLA-A*03:01 | 9 | 0.992137 | 0.01 | -0.26733 | 100.00% (10/10) | Non-Toxin | IEDB recommended 2020.09 (NetMHCpan EL 4.1) |
| 1071 | Polymerase | RLFAKMTYK | HLA-A*30:01 | 9 | 0.857769 | 0.01 | -0.26733 | 100.00% (10/10) | Non-Toxin | IEDB recommended 2020.09 (NetMHCpan EL 4.1) |
| 1072 | Polymerase | RLFAKMTYK | HLA-A*11:01 | 9 | 0.869853 | 0.04 | -0.26733 | 100.00% (10/10) | Non-Toxin | IEDB recommended 2020.09 (NetMHCpan EL 4.1) |
| 1073 | Polymerase | RLFAKMTYK | HLA-A*31:01 | 9 | 0.82979 | 0.05 | -0.26733 | 100.00% (10/10) | Non-Toxin | IEDB recommended 2020.09 (NetMHCpan EL 4.1) |
| 1074 | Polymerase | RLFNMYRSY | HLA-A*32:01 | 9 | 0.853832 | 0.01 | -0.19597 | 100.00% (10/10) | Non-Toxin | IEDB recommended 2020.09 (NetMHCpan EL 4.1) |
| 1075 | Polymerase | RLFNMYRSY | HLA-A*30:02 | 9 | 0.835929 | 0.01 | -0.19597 | 100.00% (10/10) | Non-Toxin | IEDB recommended 2020.09 (NetMHCpan EL 4.1) |
| 1076 | Polymerase | RLFNMYRSY | HLA-B*15:01 | 9 | 0.902758 | 0.02 | -0.19597 | 100.00% (10/10) | Non-Toxin | IEDB recommended 2020.09 (NetMHCpan EL 4.1) |
| 1077 | Polymerase | RLFNMYRSYF | HLA-A*32:01 | 10 | 0.538636 | 0.08 | -0.23617 | 100.00% (10/10) | Non-Toxin | IEDB recommended 2020.09 (NetMHCpan EL 4.1) |
| 1078 | Polymerase | RLNLHGKRK | HLA-A*03:01 | 9 | 0.840472 | 0.05 | -0.10162 | 100.00% (10/10) | Non-Toxin | IEDB recommended 2020.09 (NetMHCpan EL 4.1) |
| 1079 | Polymerase | RLRDKSTQF | HLA-B*15:01 | 9 | 0.964699 | 0.01 | -0.36153 | 100.00% (10/10) | Non-Toxin | IEDB recommended 2020.09 (NetMHCpan EL 4.1) |

| | | | | | | | | | | |
|---|---|---|---|---|---|---|---|---|---|---|
| 1080 | Polymerase | RLRDKSTQF | HLA-A*32:01 | 9 | 0.781152 | 0.02 | -0.36153 | 100.00% (10/10) | Non-Toxin | IEDB recommended 2020.09 (NetMHCpan EL 4.1) |
| 1081 | Polymerase | RLRDKSTQF | HLA-B*08:01 | 9 | 0.714697 | 0.06 | -0.36153 | 100.00% (10/10) | Non-Toxin | IEDB recommended 2020.09 (NetMHCpan EL 4.1) |
| 1082 | Polymerase | RLRDKSTQFK | HLA-A*03:01 | 10 | 0.951989 | 0.01 | -0.3248 | 100.00% (10/10) | Non-Toxin | IEDB recommended 2020.09 (NetMHCpan EL 4.1) |
| 1083 | Polymerase | RLRDKSTQFK | HLA-A*30:01 | 10 | 0.799443 | 0.02 | -0.3248 | 100.00% (10/10) | Non-Toxin | IEDB recommended 2020.09 (NetMHCpan EL 4.1) |
| 1084 | Polymerase | RMMMETTVK | HLA-A*03:01 | 9 | 0.794141 | 0.09 | -0.04278 | 100.00% (10/10) | Non-Toxin | IEDB recommended 2020.09 (NetMHCpan EL 4.1) |
| 1085 | Polymerase | RPRGRHTM | HLA-B*07:02 | 8 | 0.977882 | 0.02 | 0.16451 | 100.00% (10/10) | Non-Toxin | IEDB recommended 2020.09 (NetMHCpan EL 4.1) |
| 1086 | Polymerase | RPRGRHTM | HLA-B*08:01 | 8 | 0.652974 | 0.08 | 0.16451 | 100.00% (10/10) | Non-Toxin | IEDB recommended 2020.09 (NetMHCpan EL 4.1) |
| 1087 | Polymerase | RPRGRHTMV | HLA-B*07:02 | 9 | 0.977509 | 0.02 | 0.06191 | 100.00% (10/10) | Non-Toxin | IEDB recommended 2020.09 (NetMHCpan EL 4.1) |
| 1088 | Polymerase | RPRGRHTMV | HLA-B*08:01 | 9 | 0.613908 | 0.09 | 0.06191 | 100.00% (10/10) | Non-Toxin | IEDB recommended 2020.09 (NetMHCpan EL 4.1) |
| 1089 | Polymerase | RPRGRHTMVDL | HLA-B*07:02 | 11 | 0.946034 | 0.03 | 0.0531 | 100.00% (10/10) | Non-Toxin | IEDB recommended 2020.09 (NetMHCpan EL 4.1) |
| 1090 | Polymerase | RPSVENHKY | HLA-B*35:01 | 9 | 0.814186 | 0.07 | -0.01945 | 100.00% (10/10) | Non-Toxin | IEDB recommended 2020.09 (NetMHCpan EL 4.1) |
| 1091 | Polymerase | RPTKALRSA | HLA-B*07:02 | 9 | 0.751886 | 0.1 | -0.22972 | 100.00% (10/10) | Non-Toxin | IEDB recommended 2020.09 (NetMHCpan EL 4.1) |
| 1092 | Polymerase | RPTKALRSAI | HLA-B*07:02 | 10 | 0.927416 | 0.04 | -0.24514 | 100.00% (10/10) | Non-Toxin | IEDB recommended 2020.09 (NetMHCpan EL 4.1) |
| 1093 | Polymerase | RSKLMTKTLPK | HLA-A*30:01 | 11 | 0.584701 | 0.09 | -0.40366 | 100.00% (10/10) | Non-Toxin | IEDB recommended 2020.09 (NetMHCpan EL 4.1) |
| 1094 | Polymerase | RTFGHPILEAK | HLA-A*03:01 | 11 | 0.977686 | 0.01 | 0.31932 | 100.00% (10/10) | Non-Toxin | IEDB recommended 2020.09 (NetMHCpan EL 4.1) |
| 1095 | Polymerase | RTFGHPILEAK | HLA-A*11:01 | 11 | 0.962112 | 0.01 | 0.31932 | 100.00% (10/10) | Non-Toxin | IEDB recommended 2020.09 (NetMHCpan EL 4.1) |
| 1096 | Polymerase | RTFGHPILEAK | HLA-A*30:01 | 11 | 0.834546 | 0.01 | 0.31932 | 100.00% (10/10) | Non-Toxin | IEDB recommended 2020.09 (NetMHCpan EL 4.1) |
| 1097 | Polymerase | RYEDNTGTKF | HLA-A*24:02 | 10 | 0.74196 | 0.07 | 0.02498 | 100.00% (10/10) | Non-Toxin | IEDB recommended 2020.09 (NetMHCpan EL 4.1) |
| 1098 | Polymerase | RYKCSNTSK | HLA-A*30:01 | 9 | 0.731337 | 0.04 | -0.35534 | 100.00% (10/10) | Non-Toxin | IEDB recommended 2020.09 (NetMHCpan EL 4.1) |
| 1099 | Polymerase | RYLPSGAQR | HLA-A*31:01 | 9 | 0.915958 | 0.02 | -0.17862 | 100.00% (10/10) | Non-Toxin | IEDB recommended 2020.09 (NetMHCpan EL 4.1) |
| 1100 | Polymerase | RYLPSGAQRL | HLA-A*24:02 | 10 | 0.864861 | 0.04 | -0.17355 | 100.00% (10/10) | Non-Toxin | IEDB recommended 2020.09 (NetMHCpan EL 4.1) |
| 1101 | Polymerase | RYLPSGAQRL | HLA-A*23:01 | 10 | 0.793876 | 0.05 | -0.17355 | 100.00% (10/10) | Non-Toxin | IEDB recommended 2020.09 (NetMHCpan EL 4.1) |
| 1102 | Polymerase | RYLPSGAQRLF | HLA-A*24:02 | 11 | 0.963175 | 0.01 | -0.1766 | 100.00% (10/10) | Non-Toxin | IEDB recommended 2020.09 (NetMHCpan EL 4.1) |
| 1103 | Polymerase | RYLPSGAQRLF | HLA-A*23:01 | 11 | 0.950483 | 0.01 | -0.1766 | 100.00% (10/10) | Non-Toxin | IEDB recommended 2020.09 (NetMHCpan EL 4.1) |
| 1104 | Polymerase | SAIEYAQLR | HLA-A*68:01 | 9 | 0.939067 | 0.04 | 0.07294 | 100.00% (10/10) | Non-Toxin | IEDB recommended 2020.09 (NetMHCpan EL 4.1) |
| 1105 | Polymerase | SAIRIATV | HLA-B*51:01 | 8 | 0.646464 | 0.1 | 0.29447 | 100.00% (10/10) | Non-Toxin | IEDB recommended 2020.09 (NetMHCpan EL 4.1) |
| 1106 | Polymerase | SAIRIATVY | HLA-B*35:01 | 9 | 0.96342 | 0.02 | 0.31859 | 100.00% (10/10) | Non-Toxin | IEDB recommended 2020.09 (NetMHCpan EL 4.1) |
| 1107 | Polymerase | SAIRIATVY | HLA-B*15:01 | 9 | 0.78797 | 0.08 | 0.31859 | 100.00% (10/10) | Non-Toxin | IEDB recommended 2020.09 (NetMHCpan EL 4.1) |
| 1108 | Polymerase | SAIRIATVY | HLA-A*30:02 | 9 | 0.60604 | 0.1 | 0.31859 | 100.00% (10/10) | Non-Toxin | IEDB recommended 2020.09 (NetMHCpan EL 4.1) |
| 1109 | Polymerase | SAIRIATVYTW | HLA-B*57:01 | 11 | 0.950289 | 0.06 | 0.3592 | 100.00% (10/10) | Non-Toxin | IEDB recommended 2020.09 (NetMHCpan EL 4.1) |
| 1110 | Polymerase | SAIRIATVYTW | HLA-B*58:01 | 11 | 0.850536 | 0.09 | 0.3592 | 100.00% (10/10) | Non-Toxin | IEDB recommended 2020.09 (NetMHCpan EL 4.1) |
| 1111 | Polymerase | SASKKIFNL | HLA-B*08:01 | 9 | 0.713945 | 0.06 | -0.2604 | 100.00% (10/10) | Non-Toxin | IEDB recommended 2020.09 (NetMHCpan EL 4.1) |
| 1112 | Polymerase | SAYETNTRI | HLA-B*51:01 | 9 | 0.755332 | 0.05 | 0.19426 | 100.00% (10/10) | Non-Toxin | IEDB recommended 2020.09 (NetMHCpan EL 4.1) |
| 1113 | Polymerase | SFLDYHTEF | HLA-A*23:01 | 9 | 0.914152 | 0.02 | 0.13683 | 100.00% (10/10) | Non-Toxin | IEDB recommended 2020.09 (NetMHCpan EL 4.1) |
| 1114 | Polymerase | SFLDYHTEF | HLA-A*24:02 | 9 | 0.908613 | 0.02 | 0.13683 | 100.00% (10/10) | Non-Toxin | IEDB recommended 2020.09 (NetMHCpan EL 4.1) |
| 1115 | Polymerase | SGLQPKLVSR | HLA-A*31:01 | 10 | 0.771077 | 0.08 | -0.41322 | 90.00% (9/10) | Non-Toxin | IEDB recommended 2020.09 (NetMHCpan EL 4.1) |
| 1116 | Polymerase | SIMTESVLQK | HLA-A*11:01 | 10 | 0.90894 | 0.02 | -0.11972 | 100.00% (10/10) | Non-Toxin | IEDB recommended 2020.09 (NetMHCpan EL 4.1) |
| 1117 | Polymerase | SIMTESVLQK | HLA-A*03:01 | 10 | 0.896155 | 0.03 | -0.11972 | 100.00% (10/10) | Non-Toxin | IEDB recommended 2020.09 (NetMHCpan EL 4.1) |
| 1118 | Polymerase | SITSQGMTSK | HLA-A*11:01 | 10 | 0.826196 | 0.06 | -0.46287 | 100.00% (10/10) | Non-Toxin | IEDB recommended 2020.09 (NetMHCpan EL 4.1) |
| 1119 | Polymerase | SLREDIITSK | HLA-A*03:01 | 10 | 0.900281 | 0.03 | 0.33013 | 100.00% (10/10) | Non-Toxin | IEDB recommended 2020.09 (NetMHCpan EL 4.1) |
| 1120 | Polymerase | SLYILRQSK | HLA-A*03:01 | 9 | 0.893911 | 0.03 | -0.02378 | 100.00% (10/10) | Non-Toxin | IEDB recommended 2020.09 (NetMHCpan EL 4.1) |
| 1121 | Polymerase | SLYILRQSK | HLA-A*30:01 | 9 | 0.6114 | 0.07 | -0.02378 | 100.00% (10/10) | Non-Toxin | IEDB recommended 2020.09 (NetMHCpan EL 4.1) |
| 1122 | Polymerase | SMFIDDLLSI | HLA-A*02:03 | 10 | 0.689384 | 0.1 | 0.09866 | 100.00% (10/10) | Non-Toxin | IEDB recommended 2020.09 (NetMHCpan EL 4.1) |
| 1123 | Polymerase | SMIEPLVLA | HLA-A*02:03 | 9 | 0.895288 | 0.03 | 0.15107 | 100.00% (10/10) | Non-Toxin | IEDB recommended 2020.09 (NetMHCpan EL 4.1) |
| 1124 | Polymerase | SMIEPLVLA | HLA-A*02:06 | 9 | 0.898162 | 0.04 | 0.15107 | 100.00% (10/10) | Non-Toxin | IEDB recommended 2020.09 (NetMHCpan EL 4.1) |
| 1125 | Polymerase | SMIEPLVLA | HLA-A*02:01 | 9 | 0.886276 | 0.04 | 0.15107 | 100.00% (10/10) | Non-Toxin | IEDB recommended 2020.09 (NetMHCpan EL 4.1) |
| 1126 | Polymerase | SMNTKSASK | HLA-A*03:01 | 9 | 0.886178 | 0.04 | -0.39241 | 100.00% (10/10) | Non-Toxin | IEDB recommended 2020.09 (NetMHCpan EL 4.1) |
| 1127 | Polymerase | SMNTKSASK | HLA-A*30:01 | 9 | 0.577742 | 0.09 | -0.39241 | 100.00% (10/10) | Non-Toxin | IEDB recommended 2020.09 (NetMHCpan EL 4.1) |
| 1128 | Polymerase | SMYMKDKAL | HLA-B*08:01 | 9 | 0.83085 | 0.03 | -0.52616 | 100.00% (10/10) | Non-Toxin | IEDB recommended 2020.09 (NetMHCpan EL 4.1) |
| 1129 | Polymerase | SPIKDEWDSVY | HLA-B*35:01 | 11 | 0.891279 | 0.05 | 0.06638 | 100.00% (10/10) | Non-Toxin | IEDB recommended 2020.09 (NetMHCpan EL 4.1) |
| 1130 | Polymerase | SPIVSGKLI | HLA-B*51:01 | 9 | 0.650402 | 0.1 | -0.23294 | 100.00% (10/10) | Non-Toxin | IEDB recommended 2020.09 (NetMHCpan EL 4.1) |

| | | | | | | | | | | |
|---|---|---|---|---|---|---|---|---|---|---|
| 1131 | Polymerase | SPNPMLKGL | HLA-B*07:02 | 9 | 0.96201 | 0.03 | -0.3569 | 100.00% (10/10) | Non-Toxin | IEDB recommended 2020.09 (NetMHCpan EL 4.1) |
| 1132 | Polymerase | SPSHHLEPY | HLA-B*35:01 | 9 | 0.926763 | 0.03 | 0.07793 | 100.00% (10/10) | Non-Toxin | IEDB recommended 2020.09 (NetMHCpan EL 4.1) |
| 1133 | Polymerase | SQNLLVTSY | HLA-B*15:01 | 9 | 0.988794 | 0.01 | -0.0491 | 100.00% (10/10) | Non-Toxin | IEDB recommended 2020.09 (NetMHCpan EL 4.1) |
| 1134 | Polymerase | SQNLLVTSY | HLA-A*30:02 | 9 | 0.843801 | 0.01 | -0.0491 | 100.00% (10/10) | Non-Toxin | IEDB recommended 2020.09 (NetMHCpan EL 4.1) |
| 1135 | Polymerase | STEFSINETL | HLA-B*40:01 | 10 | 0.956627 | 0.02 | 0.21989 | 100.00% (10/10) | Non-Toxin | IEDB recommended 2020.09 (NetMHCpan EL 4.1) |
| 1136 | Polymerase | STHLFIYSK | HLA-A*11:01 | 9 | 0.957062 | 0.01 | 0.13884 | 90.00% (9/10) | Non-Toxin | IEDB recommended 2020.09 (NetMHCpan EL 4.1) |
| 1137 | Polymerase | STHLFIYSK | HLA-A*30:01 | 9 | 0.710044 | 0.04 | 0.13884 | 90.00% (9/10) | Non-Toxin | IEDB recommended 2020.09 (NetMHCpan EL 4.1) |
| 1138 | Polymerase | STLGQSISF | HLA-A*32:01 | 9 | 0.807368 | 0.02 | -0.22237 | 100.00% (10/10) | Non-Toxin | IEDB recommended 2020.09 (NetMHCpan EL 4.1) |
| 1139 | Polymerase | STLGQSISFY | HLA-A*30:02 | 10 | 0.639465 | 0.08 | -0.18934 | 100.00% (10/10) | Non-Toxin | IEDB recommended 2020.09 (NetMHCpan EL 4.1) |
| 1140 | Polymerase | STSNNLSHR | HLA-A*68:01 | 9 | 0.922277 | 0.06 | -0.19767 | 100.00% (10/10) | Non-Toxin | IEDB recommended 2020.09 (NetMHCpan EL 4.1) |
| 1141 | Polymerase | SVLEGKFRL | HLA-A*02:06 | 9 | 0.833349 | 0.06 | 0.05619 | 100.00% (10/10) | Non-Toxin | IEDB recommended 2020.09 (NetMHCpan EL 4.1) |
| 1142 | Polymerase | SVLEGKFRLR | HLA-A*31:01 | 10 | 0.797535 | 0.06 | 0.06755 | 100.00% (10/10) | Non-Toxin | IEDB recommended 2020.09 (NetMHCpan EL 4.1) |
| 1143 | Polymerase | SVLNRVSRY | HLA-A*30:02 | 9 | 0.832181 | 0.02 | -0.03023 | 100.00% (10/10) | Non-Toxin | IEDB recommended 2020.09 (NetMHCpan EL 4.1) |
| 1144 | Polymerase | SVLNRVSRY | HLA-A*26:01 | 9 | 0.790896 | 0.04 | -0.03023 | 100.00% (10/10) | Non-Toxin | IEDB recommended 2020.09 (NetMHCpan EL 4.1) |
| 1145 | Polymerase | SVYPREVLSY | HLA-A*30:02 | 10 | 0.87422 | 0.01 | 0.06838 | 100.00% (10/10) | Non-Toxin | IEDB recommended 2020.09 (NetMHCpan EL 4.1) |
| 1146 | Polymerase | SVYPREVLSY | HLA-B*15:01 | 10 | 0.905934 | 0.02 | 0.06838 | 100.00% (10/10) | Non-Toxin | IEDB recommended 2020.09 (NetMHCpan EL 4.1) |
| 1147 | Polymerase | SVYPREVLSY | HLA-A*03:01 | 10 | 0.881268 | 0.04 | 0.06838 | 100.00% (10/10) | Non-Toxin | IEDB recommended 2020.09 (NetMHCpan EL 4.1) |
| 1148 | Polymerase | SVYPREVLSY | HLA-A*26:01 | 10 | 0.811638 | 0.04 | 0.06838 | 100.00% (10/10) | Non-Toxin | IEDB recommended 2020.09 (NetMHCpan EL 4.1) |
| 1149 | Polymerase | SVYPREVLSY | HLA-A*11:01 | 10 | 0.800093 | 0.08 | 0.06838 | 100.00% (10/10) | Non-Toxin | IEDB recommended 2020.09 (NetMHCpan EL 4.1) |
| 1150 | Polymerase | SVYPREVLSY | HLA-A*32:01 | 10 | 0.539727 | 0.08 | 0.06838 | 100.00% (10/10) | Non-Toxin | IEDB recommended 2020.09 (NetMHCpan EL 4.1) |
| 1151 | Polymerase | SYFGLVLVCF | HLA-A*23:01 | 10 | 0.744381 | 0.07 | 0.0944 | 100.00% (10/10) | Non-Toxin | IEDB recommended 2020.09 (NetMHCpan EL 4.1) |
| 1152 | Polymerase | SYMIYLMNW | HLA-A*23:01 | 9 | 0.822194 | 0.04 | -0.0891 | 100.00% (10/10) | Non-Toxin | IEDB recommended 2020.09 (NetMHCpan EL 4.1) |
| 1153 | Polymerase | SYMIYLMNW | HLA-A*24:02 | 9 | 0.804882 | 0.06 | -0.0891 | 100.00% (10/10) | Non-Toxin | IEDB recommended 2020.09 (NetMHCpan EL 4.1) |
| 1154 | Polymerase | TEFNPHNHY | HLA-B*44:03 | 9 | 0.982412 | 0.01 | 0.06458 | 100.00% (10/10) | Non-Toxin | IEDB recommended 2020.09 (NetMHCpan EL 4.1) |
| 1155 | Polymerase | TEFNPHNHY | HLA-B*44:02 | 9 | 0.965017 | 0.01 | 0.06458 | 100.00% (10/10) | Non-Toxin | IEDB recommended 2020.09 (NetMHCpan EL 4.1) |
| 1156 | Polymerase | TEFSINETL | HLA-B*40:01 | 9 | 0.992205 | 0.01 | 0.10222 | 100.00% (10/10) | Non-Toxin | IEDB recommended 2020.09 (NetMHCpan EL 4.1) |
| 1157 | Polymerase | TEFSINETL | HLA-B*44:03 | 9 | 0.774084 | 0.09 | 0.10222 | 100.00% (10/10) | Non-Toxin | IEDB recommended 2020.09 (NetMHCpan EL 4.1) |
| 1158 | Polymerase | TEFSINETLTL | HLA-B*40:01 | 11 | 0.922328 | 0.06 | 0.14219 | 100.00% (10/10) | Non-Toxin | IEDB recommended 2020.09 (NetMHCpan EL 4.1) |
| 1159 | Polymerase | TEIILNPNL | HLA-B*40:01 | 9 | 0.912165 | 0.06 | 0.14709 | 100.00% (10/10) | Non-Toxin | IEDB recommended 2020.09 (NetMHCpan EL 4.1) |
| 1160 | Polymerase | TESVLQKVM | HLA-B*44:03 | 9 | 0.799231 | 0.08 | -0.28988 | 100.00% (10/10) | Non-Toxin | IEDB recommended 2020.09 (NetMHCpan EL 4.1) |
| 1161 | Polymerase | TESVLQKVM | HLA-B*44:02 | 9 | 0.720928 | 0.09 | -0.28988 | 100.00% (10/10) | Non-Toxin | IEDB recommended 2020.09 (NetMHCpan EL 4.1) |
| 1162 | Polymerase | TETIISTHL | HLA-B*40:01 | 9 | 0.944836 | 0.04 | 0.17205 | 100.00% (10/10) | Non-Toxin | IEDB recommended 2020.09 (NetMHCpan EL 4.1) |
| 1163 | Polymerase | TETIISTHLF | HLA-B*44:03 | 10 | 0.750623 | 0.1 | 0.17238 | 100.00% (10/10) | Non-Toxin | IEDB recommended 2020.09 (NetMHCpan EL 4.1) |
| 1164 | Polymerase | TETIISTHLF | HLA-B*44:02 | 10 | 0.696101 | 0.1 | 0.17238 | 100.00% (10/10) | Non-Toxin | IEDB recommended 2020.09 (NetMHCpan EL 4.1) |
| 1165 | Polymerase | TEVDNNHLI | HLA-B*40:01 | 9 | 0.848758 | 0.09 | 0.04415 | 100.00% (10/10) | Non-Toxin | IEDB recommended 2020.09 (NetMHCpan EL 4.1) |
| 1166 | Polymerase | TEVDNNHLI | HLA-B*44:03 | 9 | 0.773184 | 0.09 | 0.04415 | 100.00% (10/10) | Non-Toxin | IEDB recommended 2020.09 (NetMHCpan EL 4.1) |
| 1167 | Polymerase | TEVDNNHLIY | HLA-B*44:03 | 10 | 0.921608 | 0.03 | 0.12197 | 100.00% (10/10) | Non-Toxin | IEDB recommended 2020.09 (NetMHCpan EL 4.1) |
| 1168 | Polymerase | TEVDNNHLIY | HLA-A*01:01 | 10 | 0.912782 | 0.03 | 0.12197 | 100.00% (10/10) | Non-Toxin | IEDB recommended 2020.09 (NetMHCpan EL 4.1) |
| 1169 | Polymerase | TEVDNNHLIY | HLA-B*44:02 | 10 | 0.812143 | 0.05 | 0.12197 | 100.00% (10/10) | Non-Toxin | IEDB recommended 2020.09 (NetMHCpan EL 4.1) |
| 1170 | Polymerase | TGKLKGLVV | HLA-B*08:01 | 9 | 0.616068 | 0.09 | -0.2445 | 100.00% (10/10) | Non-Toxin | IEDB recommended 2020.09 (NetMHCpan EL 4.1) |
| 1171 | Polymerase | TIATIPFLF | HLA-A*32:01 | 9 | 0.530401 | 0.09 | 0.26324 | 100.00% (10/10) | Non-Toxin | IEDB recommended 2020.09 (NetMHCpan EL 4.1) |
| 1172 | Polymerase | TIMNCVTKK | HLA-A*11:01 | 9 | 0.771159 | 0.09 | -0.17039 | 90.00% (9/10) | Non-Toxin | IEDB recommended 2020.09 (NetMHCpan EL 4.1) |
| 1173 | Polymerase | TKTIKNITAR | HLA-A*68:01 | 10 | 0.894531 | 0.08 | 0.11112 | 100.00% (10/10) | Non-Toxin | IEDB recommended 2020.09 (NetMHCpan EL 4.1) |
| 1174 | Polymerase | TLGQSISFY | HLA-A*30:02 | 9 | 0.614879 | 0.1 | -0.2126 | 100.00% (10/10) | Non-Toxin | IEDB recommended 2020.09 (NetMHCpan EL 4.1) |
| 1175 | Polymerase | TLVDETRSA | HLA-A*02:03 | 9 | 0.6851 | 0.1 | 0.11678 | 100.00% (10/10) | Non-Toxin | IEDB recommended 2020.09 (NetMHCpan EL 4.1) |
| 1176 | Polymerase | TPEDDIFIHY | HLA-B*35:01 | 10 | 0.834987 | 0.06 | 0.44744 | 100.00% (10/10) | Non-Toxin | IEDB recommended 2020.09 (NetMHCpan EL 4.1) |
| 1177 | Polymerase | TPEDDIFIHY | HLA-B*53:01 | 10 | 0.63585 | 0.07 | 0.44744 | 100.00% (10/10) | Non-Toxin | IEDB recommended 2020.09 (NetMHCpan EL 4.1) |
| 1178 | Polymerase | TPEDDIFIHY | HLA-B*44:03 | 10 | 0.796685 | 0.08 | 0.44744 | 100.00% (10/10) | Non-Toxin | IEDB recommended 2020.09 (NetMHCpan EL 4.1) |
| 1179 | Polymerase | TPGFPISRL | HLA-B*07:02 | 9 | 0.83853 | 0.07 | 0.1339 | 100.00% (10/10) | Non-Toxin | IEDB recommended 2020.09 (NetMHCpan EL 4.1) |
| 1180 | Polymerase | TPGFPISRLF | HLA-B*53:01 | 10 | 0.721765 | 0.05 | 0.12907 | 100.00% (10/10) | Non-Toxin | IEDB recommended 2020.09 (NetMHCpan EL 4.1) |
| 1181 | Polymerase | TPVSTSNNL | HLA-B*07:02 | 9 | 0.815954 | 0.08 | -0.28024 | 100.00% (10/10) | Non-Toxin | IEDB recommended 2020.09 (NetMHCpan EL 4.1) |

| | | | | | | | | | | |
|---|---|---|---|---|---|---|---|---|---|---|
| 1182 | Polymerase | TSDLDFVIFY | HLA-A*01:01 | 10 | 0.973912 | 0.01 | 0.35122 | 100.00% (10/10) | Non-Toxin | IEDB recommended 2020.09 (NetMHCpan EL 4.1) |
| 1183 | Polymerase | TTVKSDIKY | HLA-A*26:01 | 9 | 0.64814 | 0.07 | -0.3575 | 100.00% (10/10) | Non-Toxin | IEDB recommended 2020.09 (NetMHCpan EL 4.1) |
| 1184 | Polymerase | TTVKSDIKY | HLA-A*01:01 | 9 | 0.69926 | 0.1 | -0.3575 | 100.00% (10/10) | Non-Toxin | IEDB recommended 2020.09 (NetMHCpan EL 4.1) |
| 1185 | Polymerase | TTWIGNLDSY | HLA-A*26:01 | 10 | 0.554422 | 0.09 | 0.14414 | 100.00% (10/10) | Non-Toxin | IEDB recommended 2020.09 (NetMHCpan EL 4.1) |
| 1186 | Polymerase | TVAQTVLEI | HLA-A*68:02 | 9 | 0.671157 | 0.09 | 0.02194 | 100.00% (10/10) | Non-Toxin | IEDB recommended 2020.09 (NetMHCpan EL 4.1) |
| 1187 | Polymerase | TVLEIITKA | HLA-A*02:06 | 9 | 0.880375 | 0.05 | 0.25879 | 100.00% (10/10) | Non-Toxin | IEDB recommended 2020.09 (NetMHCpan EL 4.1) |
| 1188 | Polymerase | TVLEIITKA | HLA-A*68:02 | 9 | 0.740554 | 0.07 | 0.25879 | 100.00% (10/10) | Non-Toxin | IEDB recommended 2020.09 (NetMHCpan EL 4.1) |
| 1189 | Polymerase | VIFYASLTY | HLA-A*30:02 | 9 | 0.722156 | 0.04 | -0.07003 | 100.00% (10/10) | Non-Toxin | IEDB recommended 2020.09 (NetMHCpan EL 4.1) |
| 1190 | Polymerase | VIFYASLTY | HLA-B*15:01 | 9 | 0.829588 | 0.05 | -0.07003 | 100.00% (10/10) | Non-Toxin | IEDB recommended 2020.09 (NetMHCpan EL 4.1) |
| 1191 | Polymerase | VIKNSQKPK | HLA-A*30:01 | 9 | 0.606311 | 0.08 | -0.53513 | 100.00% (10/10) | Non-Toxin | IEDB recommended 2020.09 (NetMHCpan EL 4.1) |
| 1192 | Polymerase | VMMEDGLLV | HLA-A*02:01 | 9 | 0.871134 | 0.05 | 0.08141 | 100.00% (10/10) | Non-Toxin | IEDB recommended 2020.09 (NetMHCpan EL 4.1) |
| 1193 | Polymerase | VMMEDGLLV | HLA-A*02:06 | 9 | 0.746373 | 0.09 | 0.08141 | 100.00% (10/10) | Non-Toxin | IEDB recommended 2020.09 (NetMHCpan EL 4.1) |
| 1194 | Polymerase | VMMEDGLLV | HLA-A*02:03 | 9 | 0.702884 | 0.1 | 0.08141 | 100.00% (10/10) | Non-Toxin | IEDB recommended 2020.09 (NetMHCpan EL 4.1) |
| 1195 | Polymerase | VSNTSKHTY | HLA-A*30:02 | 9 | 0.859883 | 0.01 | -0.27716 | 100.00% (10/10) | Non-Toxin | IEDB recommended 2020.09 (NetMHCpan EL 4.1) |
| 1196 | Polymerase | VSNTSKHTY | HLA-A*01:01 | 9 | 0.736085 | 0.09 | -0.27716 | 100.00% (10/10) | Non-Toxin | IEDB recommended 2020.09 (NetMHCpan EL 4.1) |
| 1197 | Polymerase | VYPREVLSY | HLA-A*24:02 | 9 | 0.720043 | 0.09 | 0.07882 | 100.00% (10/10) | Non-Toxin | IEDB recommended 2020.09 (NetMHCpan EL 4.1) |
| 1198 | Polymerase | VYPREVLSY | HLA-A*23:01 | 9 | 0.647236 | 0.1 | 0.07882 | 100.00% (10/10) | Non-Toxin | IEDB recommended 2020.09 (NetMHCpan EL 4.1) |
| 1199 | Polymerase | WAFEIHHRR | HLA-A*33:01 | 9 | 0.829405 | 0.02 | 0.35634 | 100.00% (10/10) | Non-Toxin | IEDB recommended 2020.09 (NetMHCpan EL 4.1) |
| 1200 | Polymerase | WAFEIHHRR | HLA-A*68:01 | 9 | 0.939272 | 0.04 | 0.35634 | 100.00% (10/10) | Non-Toxin | IEDB recommended 2020.09 (NetMHCpan EL 4.1) |
| 1201 | Polymerase | YEDNTGTKF | HLA-B*44:02 | 9 | 0.709644 | 0.09 | -0.02285 | 100.00% (10/10) | Non-Toxin | IEDB recommended 2020.09 (NetMHCpan EL 4.1) |
| 1202 | Polymerase | YEDNTGTKF | HLA-B*40:01 | 9 | 0.805676 | 0.1 | -0.02285 | 100.00% (10/10) | Non-Toxin | IEDB recommended 2020.09 (NetMHCpan EL 4.1) |
| 1203 | Polymerase | YEDNTGTKF | HLA-B*44:03 | 9 | 0.769024 | 0.1 | -0.02285 | 100.00% (10/10) | Non-Toxin | IEDB recommended 2020.09 (NetMHCpan EL 4.1) |
| 1204 | Polymerase | YGLPGFFNW | HLA-B*58:01 | 9 | 0.949328 | 0.04 | 0.22346 | 100.00% (10/10) | Non-Toxin | IEDB recommended 2020.09 (NetMHCpan EL 4.1) |
| 1205 | Polymerase | YGLPGFFNW | HLA-B*57:01 | 9 | 0.963054 | 0.05 | 0.22346 | 100.00% (10/10) | Non-Toxin | IEDB recommended 2020.09 (NetMHCpan EL 4.1) |
| 1206 | Polymerase | YHTEFNPHNHY | HLA-B*44:03 | 11 | 0.790711 | 0.08 | 0.25414 | 100.00% (10/10) | Non-Toxin | IEDB recommended 2020.09 (NetMHCpan EL 4.1) |
| 1207 | Polymerase | YLASQRVNI | HLA-A*02:03 | 9 | 0.772641 | 0.07 | -0.18679 | 100.00% (10/10) | Non-Toxin | IEDB recommended 2020.09 (NetMHCpan EL 4.1) |
| 1208 | Polymerase | YLEDEQFNV | HLA-A*02:01 | 9 | 0.965747 | 0.02 | 0.1383 | 100.00% (10/10) | Non-Toxin | IEDB recommended 2020.09 (NetMHCpan EL 4.1) |
| 1209 | Polymerase | YLEDEQFNV | HLA-A*02:06 | 9 | 0.902384 | 0.04 | 0.1383 | 100.00% (10/10) | Non-Toxin | IEDB recommended 2020.09 (NetMHCpan EL 4.1) |
| 1210 | Polymerase | YLEDEQFNVSY | HLA-A*01:01 | 11 | 0.869685 | 0.05 | 0.08561 | 100.00% (10/10) | Non-Toxin | IEDB recommended 2020.09 (NetMHCpan EL 4.1) |
| 1211 | Polymerase | YLPSGAQRL | HLA-A*02:03 | 9 | 0.722541 | 0.09 | -0.16776 | 100.00% (10/10) | Non-Toxin | IEDB recommended 2020.09 (NetMHCpan EL 4.1) |
| 1212 | Polymerase | YLPSGAQRLF | HLA-B*53:01 | 10 | 0.636284 | 0.07 | -0.17081 | 100.00% (10/10) | Non-Toxin | IEDB recommended 2020.09 (NetMHCpan EL 4.1) |
| 1213 | Polymerase | YLTPEMVLM | HLA-A*02:01 | 9 | 0.749817 | 0.1 | -0.038 | 100.00% (10/10) | Non-Toxin | IEDB recommended 2020.09 (NetMHCpan EL 4.1) |
| 1214 | Polymerase | YPECNNILF | HLA-B*53:01 | 9 | 0.766098 | 0.04 | 0.0717 | 90.00% (9/10) | Non-Toxin | IEDB recommended 2020.09 (NetMHCpan EL 4.1) |
| 1215 | Polymerase | YPECNNILF | HLA-B*35:01 | 9 | 0.85598 | 0.06 | 0.0717 | 90.00% (9/10) | Non-Toxin | IEDB recommended 2020.09 (NetMHCpan EL 4.1) |
| 1216 | Polymerase | YPKGGIEGY | HLA-B*35:01 | 9 | 0.981469 | 0.01 | 0.22668 | 100.00% (10/10) | Non-Toxin | IEDB recommended 2020.09 (NetMHCpan EL 4.1) |
| 1217 | Polymerase | YPKGGIEGY | HLA-B*53:01 | 9 | 0.663676 | 0.07 | 0.22668 | 100.00% (10/10) | Non-Toxin | IEDB recommended 2020.09 (NetMHCpan EL 4.1) |
| 1218 | Polymerase | YPKGGIEGY | HLA-A*26:01 | 9 | 0.537597 | 0.1 | 0.22668 | 100.00% (10/10) | Non-Toxin | IEDB recommended 2020.09 (NetMHCpan EL 4.1) |
| 1219 | Polymerase | YPREVLSY | HLA-B*35:01 | 8 | 0.924899 | 0.03 | 0.00769 | 100.00% (10/10) | Non-Toxin | IEDB recommended 2020.09 (NetMHCpan EL 4.1) |
| 1220 | Polymerase | YPVLERTRI | HLA-B*51:01 | 9 | 0.790006 | 0.04 | 0.21146 | 100.00% (10/10) | Non-Toxin | IEDB recommended 2020.09 (NetMHCpan EL 4.1) |
| 1221 | Polymerase | YQIDQPFFV | HLA-A*02:06 | 9 | 0.99605 | 0.01 | 0.10948 | 100.00% (10/10) | Non-Toxin | IEDB recommended 2020.09 (NetMHCpan EL 4.1) |
| 1222 | Polymerase | YQIDQPFFV | HLA-A*02:01 | 9 | 0.9837 | 0.01 | 0.10948 | 100.00% (10/10) | Non-Toxin | IEDB recommended 2020.09 (NetMHCpan EL 4.1) |
| 1223 | Polymerase | YQIDQPFFV | HLA-A*02:03 | 9 | 0.894139 | 0.03 | 0.10948 | 100.00% (10/10) | Non-Toxin | IEDB recommended 2020.09 (NetMHCpan EL 4.1) |
| 1224 | Polymerase | YSNPDSTEVY | HLA-A*01:01 | 10 | 0.945 | 0.02 | -0.0076 | 100.00% (10/10) | Non-Toxin | IEDB recommended 2020.09 (NetMHCpan EL 4.1) |
| 1225 | Polymerase | YTEVDNNHLIY | HLA-A*01:01 | 11 | 0.989212 | 0.01 | 0.18189 | 100.00% (10/10) | Non-Toxin | IEDB recommended 2020.09 (NetMHCpan EL 4.1) |
| 1226 | Polymerase | YTPGFPISR | HLA-A*68:01 | 9 | 0.941523 | 0.04 | 0.14972 | 100.00% (10/10) | Non-Toxin | IEDB recommended 2020.09 (NetMHCpan EL 4.1) |
| 1227 | Polymerase | YTPGFPISRLF | HLA-B*57:01 | 11 | 0.914005 | 0.09 | 0.14477 | 100.00% (10/10) | Non-Toxin | IEDB recommended 2020.09 (NetMHCpan EL 4.1) |
| 1228 | Polymerase | YYLTPEMVL | HLA-A*24:02 | 9 | 0.743055 | 0.07 | -0.00517 | 100.00% (10/10) | Non-Toxin | IEDB recommended 2020.09 (NetMHCpan EL 4.1) |
| 1229 | Polymerase | YYLTPEMVL | HLA-A*23:01 | 9 | 0.688171 | 0.09 | -0.00517 | 100.00% (10/10) | Non-Toxin | IEDB recommended 2020.09 (NetMHCpan EL 4.1) |
| 1230 | Polymerase | YYQIDQPFF | HLA-A*24:02 | 9 | 0.973241 | 0.01 | 0.06792 | 100.00% (10/10) | Non-Toxin | IEDB recommended 2020.09 (NetMHCpan EL 4.1) |
| 1231 | Polymerase | YYQIDQPFF | HLA-A*23:01 | 9 | 0.944604 | 0.01 | 0.06792 | 100.00% (10/10) | Non-Toxin | IEDB recommended 2020.09 (NetMHCpan EL 4.1) |
| 1232 | Polymerase | YYQIDQPFFV | HLA-A*02:06 | 10 | 0.811423 | 0.07 | 0.16188 | 100.00% (10/10) | Non-Toxin | IEDB recommended 2020.09 (NetMHCpan EL 4.1) |

**Supplementary Table S3:** High Percentile Ranking HTL epitopes-HLA allele pairs screened from entire proteome of SARS-CoV-2 by the "MHC-II Binding Predictions" tool of IEDB. These epitopes were further utilized to identify the potentially immunogenic multiple epitope cluster based HTL Ag-Patches from the entire proteome of the SRAS-CoV-2. The screened epitopes are in consensus with the previous studies[38].

| S.No. | Protein | Peptide | Allele | Conservancy | Toxicity | Percentile rank | Method used |
|---|---|---|---|---|---|---|---|
| 1 | c protein | ASILLTLFRRTKKKY | HLA-DPA1*03:01/DPB1*04:02 | 100.00% (60/60) | Non-Toxin | 0.62 | Consensus (comb.lib./smm/nn) |
| 2 | c protein | ASILLTLFRRTKKKY | HLA-DRB1*11:01 | 100.00% (60/60) | Non-Toxin | 0.63 | Consensus (smm/nn/sturniolo) |
| 3 | c protein | CLRMMEMLKEETWRI | HLA-DRB3*01:01 | 21.67% (13/60) | Non-Toxin | 0.27 | Consensus (comb.lib./smm/nn) |
| 4 | c protein | CPMLPKLQFLKKIGK | HLA-DRB1*11:01 | 98.33% (59/60) | Non-Toxin | 0.19 | Consensus (smm/nn/sturniolo) |
| 5 | c protein | CPMLPKLQFLKKIGK | HLA-DRB5*01:01 | 98.33% (59/60) | Non-Toxin | 0.7 | Consensus (smm/nn/sturniolo) |
| 6 | c protein | DDQVFNNPASKIKQK | HLA-DRB1*13:02 | 21.67% (13/60) | Non-Toxin | 0.27 | Consensus (smm/nn/sturniolo) |
| 7 | c protein | DQVFNNPASKIKQKP | HLA-DRB1*13:02 | 21.67% (13/60) | Non-Toxin | 0.4 | Consensus (smm/nn/sturniolo) |
| 8 | c protein | EETWRIYPVLLPQME | HLA-DRB1*08:02 | 21.67% (13/60) | Non-Toxin | 0.41 | Consensus (smm/nn/sturniolo) |
| 9 | c protein | EMLKEETWRIYPVLL | HLA-DRB3*01:01 | 21.67% (13/60) | Non-Toxin | 0.72 | Consensus (comb.lib./smm/nn) |
| 10 | c protein | ETWRIYPVLLPQMEL | HLA-DRB1*08:02 | 21.67% (13/60) | Non-Toxin | 0.35 | Consensus (smm/nn/sturniolo) |
| 11 | c protein | FRRTKKKYRRHTDDQ | HLA-DRB5*01:01 | 95.00% (57/60) | Non-Toxin | 0.17 | Consensus (smm/nn/sturniolo) |
| 12 | c protein | HTDDQVFNNPASKIK | HLA-DRB1*13:02 | 21.67% (13/60) | Non-Toxin | 0.41 | Consensus (smm/nn/sturniolo) |
| 13 | c protein | ILLTLFRRTKKKYRR | HLA-DRB5*01:01 | 100.00% (60/60) | Non-Toxin | 0.17 | Consensus (smm/nn/sturniolo) |
| 14 | c protein | ILLTLFRRTKKKYRR | HLA-DRB1*11:01 | 100.00% (60/60) | Non-Toxin | 0.38 | Consensus (smm/nn/sturniolo) |
| 15 | c protein | IYPVLLPQMELLERE | HLA-DPA1*03:01/DPB1*04:02 | 21.67% (13/60) | Non-Toxin | 1.01 | Consensus (comb.lib./smm/nn) |
| 16 | c protein | KEETWRIYPVLLPQM | HLA-DRB1*08:02 | 21.67% (13/60) | Non-Toxin | 0.63 | Consensus (smm/nn/sturniolo) |
| 17 | c protein | KVQMTYNWTQWLQTL | HLA-DPA1*01:03/DPB1*02:01 | 95.00% (57/60) | Non-Toxin | 0.09 | Consensus (comb.lib./smm/nn) |
| 18 | c protein | KVQMTYNWTQWLQTL | HLA-DPA1*01/DPB1*04:01 | 95.00% (57/60) | Non-Toxin | 0.87 | Consensus (comb.lib./smm/nn) |
| 19 | c protein | LFRRTKKKYRRHTDD | HLA-DRB5*01:01 | 95.00% (57/60) | Non-Toxin | 0.17 | Consensus (smm/nn/sturniolo) |
| 20 | c protein | LLTLFRRTKKKYRRH | HLA-DRB5*01:01 | 100.00% (60/60) | Non-Toxin | 0.17 | Consensus (smm/nn/sturniolo) |
| 21 | c protein | LLTLFRRTKKKYRRH | HLA-DRB1*11:01 | 100.00% (60/60) | Non-Toxin | 0.34 | Consensus (smm/nn/sturniolo) |
| 22 | c protein | LQTLYTMIMEENVPD | HLA-DQA1*04:01/DQB1*04:02 | 100.00% (60/60) | Non-Toxin | 0.22 | Consensus (comb.lib./smm/nn) |
| 23 | c protein | LRMMEMLKEETWRIY | HLA-DRB3*01:01 | 21.67% (13/60) | Non-Toxin | 0.18 | Consensus (comb.lib./smm/nn) |
| 24 | c protein | LTLFRRTKKKYRRHT | HLA-DRB5*01:01 | 100.00% (60/60) | Non-Toxin | 0.17 | Consensus (smm/nn/sturniolo) |
| 25 | c protein | LTLFRRTKKKYRRHT | HLA-DRB1*11:01 | 100.00% (60/60) | Non-Toxin | 0.52 | Consensus (smm/nn/sturniolo) |
| 26 | c protein | MASILLTLFRRTKKK | HLA-DPA1*03:01/DPB1*04:02 | 100.00% (60/60) | Non-Toxin | 0.39 | Consensus (comb.lib./smm/nn) |
| 27 | c protein | MASILLTLFRRTKKK | HLA-DPA1*01:03/DPB1*02:01 | 100.00% (60/60) | Non-Toxin | 0.82 | Consensus (comb.lib./smm/nn) |
| 28 | c protein | MEMLKEETWRIYPVL | HLA-DRB3*01:01 | 21.67% (13/60) | Non-Toxin | 0.18 | Consensus (comb.lib./smm/nn) |
| 29 | c protein | MLKEETWRIYPVLLP | HLA-DRB3*01:01 | 21.67% (13/60) | Non-Toxin | 0.16 | Consensus (comb.lib./smm/nn) |
| 30 | c protein | MLPKLQFLKKIGKLI | HLA-DRB1*11:01 | 98.33% (59/60) | Non-Toxin | 0.19 | Consensus (smm/nn/sturniolo) |
| 31 | c protein | MLPKLQFLKKIGKLI | HLA-DRB5*01:01 | 98.33% (59/60) | Non-Toxin | 0.7 | Consensus (smm/nn/sturniolo) |
| 32 | c protein | MMASILLTLFRRTKK | HLA-DPA1*03:01/DPB1*04:02 | 100.00% (60/60) | Non-Toxin | 0.35 | Consensus (comb.lib./smm/nn) |
| 33 | c protein | MMASILLTLFRRTKK | HLA-DPA1*01:03/DPB1*02:01 | 100.00% (60/60) | Non-Toxin | 0.73 | Consensus (comb.lib./smm/nn) |
| 34 | c protein | MMEMLKEETWRIYPV | HLA-DRB3*01:01 | 21.67% (13/60) | Non-Toxin | 0.18 | Consensus (comb.lib./smm/nn) |
| 35 | c protein | MTYNWTQWLQTLYTM | HLA-DPA1*01:03/DPB1*02:01 | 95.00% (57/60) | Non-Toxin | 0.23 | Consensus (comb.lib./smm/nn) |
| 36 | c protein | PMLPKLQFLKKIGKL | HLA-DRB1*11:01 | 98.33% (59/60) | Non-Toxin | 0.19 | Consensus (smm/nn/sturniolo) |
| 37 | c protein | PMLPKLQFLKKIGKL | HLA-DRB5*01:01 | 98.33% (59/60) | Non-Toxin | 0.7 | Consensus (smm/nn/sturniolo) |
| 38 | c protein | QKVQMTYNWTQWLQT | HLA-DPA1*01:03/DPB1*02:01 | 95.00% (57/60) | Non-Toxin | 0.14 | Consensus (comb.lib./smm/nn) |
| 39 | c protein | QMTYNWTQWLQTLYT | HLA-DPA1*01:03/DPB1*02:01 | 95.00% (57/60) | Non-Toxin | 0.1 | Consensus (comb.lib./smm/nn) |
| 40 | c protein | QMTYNWTQWLQTLYT | HLA-DPA1*01/DPB1*04:01 | 95.00% (57/60) | Non-Toxin | 0.77 | Consensus (comb.lib./smm/nn) |
| 41 | c protein | QMTYNWTQWLQTLYT | HLA-DPA1*02:01/DPB1*01:01 | 95.00% (57/60) | Non-Toxin | 0.79 | Consensus (comb.lib./smm/nn) |
| 42 | c protein | QMTYNWTQWLQTLYT | HLA-DPA1*03:01/DPB1*04:02 | 95.00% (57/60) | Non-Toxin | 0.86 | Consensus (comb.lib./smm/nn) |

| | | | | | | | |
|---|---|---|---|---|---|---|---|
| 43 | c protein | QTLYTMIMEENVPDM | HLA-DQA1*04:01/DQB1*04:02 | 100.00% (60/60) | Non-Toxin | 0.43 | Consensus (comb.lib./smm/nn) |
| 44 | c protein | QWLQTLYTMIMEENV | HLA-DQA1*04:01/DQB1*04:02 | 100.00% (60/60) | Non-Toxin | 0.09 | Consensus (comb.lib./smm/nn) |
| 45 | c protein | RHTDDQVFNNPASKI | HLA-DRB1*13:02 | 21.67% (13/60) | Non-Toxin | 0.61 | Consensus (smm/nn/sturniolo) |
| 46 | c protein | RMMEMLKEETWRIYP | HLA-DRB3*01:01 | 21.67% (13/60) | Non-Toxin | 0.19 | Consensus (comb.lib./smm/nn) |
| 47 | c protein | SILLTLFRRTKKKYR | HLA-DRB5*01:01 | 100.00% (60/60) | Non-Toxin | 0.17 | Consensus (smm/nn/sturniolo) |
| 48 | c protein | SILLTLFRRTKKKYR | HLA-DRB1*11:01 | 100.00% (60/60) | Non-Toxin | 0.4 | Consensus (smm/nn/sturniolo) |
| 49 | c protein | TDDQVFNNPASKIKQ | HLA-DRB1*13:02 | 21.67% (13/60) | Non-Toxin | 0.3 | Consensus (smm/nn/sturniolo) |
| 50 | c protein | TLFRRTKKKYRRHTD | HLA-DRB5*01:01 | 95.00% (57/60) | Non-Toxin | 0.17 | Consensus (smm/nn/sturniolo) |
| 51 | c protein | TLFRRTKKKYRRHTD | HLA-DRB1*11:01 | 95.00% (57/60) | Non-Toxin | 0.95 | Consensus (smm/nn/sturniolo) |
| 52 | c protein | TLYTMIMEENVPDMD | HLA-DQA1*04:01/DQB1*04:02 | 100.00% (60/60) | Non-Toxin | 0.72 | Consensus (comb.lib./smm/nn) |
| 53 | c protein | TQWLQTLYTMIMEEN | HLA-DQA1*04:01/DQB1*04:02 | 100.00% (60/60) | Non-Toxin | 0.08 | Consensus (comb.lib./smm/nn) |
| 54 | c protein | TWRIYPVLLPQMELL | HLA-DRB1*08:02 | 21.67% (13/60) | Non-Toxin | 0.36 | Consensus (smm/nn/sturniolo) |
| 55 | c protein | TYNWTQWLQTLYTMI | HLA-DPA1*01:03/DPB1*02:01 | 95.00% (57/60) | Non-Toxin | 0.52 | Consensus (comb.lib./smm/nn) |
| 56 | c protein | TYNWTQWLQTLYTMI | HLA-DPA1*02:01/DPB1*01:01 | 95.00% (57/60) | Non-Toxin | 0.96 | Consensus (comb.lib./smm/nn) |
| 57 | c protein | VQMTYNWTQWLQTLY | HLA-DPA1*01:03/DPB1*02:01 | 95.00% (57/60) | Non-Toxin | 0.08 | Consensus (comb.lib./smm/nn) |
| 58 | c protein | VQMTYNWTQWLQTLY | HLA-DPA1*01/DPB1*04:01 | 95.00% (57/60) | Non-Toxin | 0.7 | Consensus (comb.lib./smm/nn) |
| 59 | c protein | VQMTYNWTQWLQTLY | HLA-DPA1*02:01/DPB1*01:01 | 95.00% (57/60) | Non-Toxin | 0.8 | Consensus (comb.lib./smm/nn) |
| 60 | c protein | WLQTLYTMIMEENVP | HLA-DQA1*04:01/DQB1*04:02 | 100.00% (60/60) | Non-Toxin | 0.1 | Consensus (comb.lib./smm/nn) |
| 61 | c protein | WRIYPVLLPQMELLE | HLA-DRB1*08:02 | 100.00% (60/60) | Non-Toxin | 0.66 | Consensus (smm/nn/sturniolo) |
| 62 | c protein | WTQWLQTLYTMIMEE | HLA-DQA1*04:01/DQB1*04:02 | 100.00% (60/60) | Non-Toxin | 0.1 | Consensus (comb.lib./smm/nn) |
| **S.No.** | **Protein** | **Peptide** | **Allele** | **Conservancy** | **Toxicity** | **Percentile rank** | **Method used** |
| 63 | fusion protein | AEKTVYVLTALQDYI | HLA-DQA1*05:01/DQB1*02:01 | 100.00% (37/37) | Non-Toxin | 0.46 | Consensus (comb.lib./smm/nn) |
| 64 | fusion protein | AFGGNYETLLRTLGY | HLA-DRB1*11:01 | 97.30% (36/37) | Non-Toxin | 0.44 | Consensus (smm/nn/sturniolo) |
| 65 | fusion protein | AGVAIGIATAAQITA | HLA-DRB1*08:02 | 97.30% (36/37) | Non-Toxin | 0.31 | Consensus (smm/nn/sturniolo) |
| 66 | fusion protein | AGVIMAGVAIGIATA | HLA-DQA1*05:01/DQB1*03:01 | 97.30% (36/37) | Non-Toxin | 0.27 | Consensus (comb.lib./smm/nn) |
| 67 | fusion protein | ALSKYLSDLLFVFGP | HLA-DRB3*01:01 | 97.30% (36/37) | Non-Toxin | 0.01 | Consensus (comb.lib./smm/nn) |
| 68 | fusion protein | ASLCIGLITFISFII | HLA-DRB1*15:01 | 100.00% (37/37) | Non-Toxin | 0.11 | Consensus (smm/nn/sturniolo) |
| 69 | fusion protein | ASLCIGLITFISFII | HLA-DPA1*01:03/DPB1*02:01 | 100.00% (37/37) | Non-Toxin | 0.37 | Consensus (comb.lib./smm/nn) |
| 70 | fusion protein | CIGLITFISFIIVEK | HLA-DPA1*01:03/DPB1*02:01 | 100.00% (37/37) | Non-Toxin | 0.05 | Consensus (comb.lib./smm/nn) |
| 71 | fusion protein | CIGLITFISFIIVEK | HLA-DRB1*15:01 | 100.00% (37/37) | Non-Toxin | 0.1 | Consensus (smm/nn/sturniolo) |
| 72 | fusion protein | CNLLILILMISECSV | HLA-DRB1*11:01 | 40.54% (15/37) | Non-Toxin | 0.12 | Consensus (smm/nn/sturniolo) |
| 73 | fusion protein | CNLLILILMISECSV | HLA-DRB1*04:05 | 40.54% (15/37) | Non-Toxin | 0.47 | Consensus (smm/nn/sturniolo) |
| 74 | fusion protein | CPRELVVSSHVPRFA | HLA-DRB1*03:01 | 100.00% (37/37) | Non-Toxin | 0.06 | Consensus (smm/nn/sturniolo) |
| 75 | fusion protein | CYCNLLILILMISEC | HLA-DPA1*03:01/DPB1*04:02 | 40.54% (15/37) | Non-Toxin | 0.12 | Consensus (comb.lib./smm/nn) |
| 76 | fusion protein | CYCNLLILILMISEC | HLA-DRB1*11:01 | 40.54% (15/37) | Non-Toxin | 0.28 | Consensus (smm/nn/sturniolo) |
| 77 | fusion protein | DIVIKMIPNVSNMSQ | HLA-DRB1*08:02 | 97.30% (36/37) | Non-Toxin | 0.1 | Consensus (smm/nn/sturniolo) |
| 78 | fusion protein | DKRCYCNLLILILMI | HLA-DPA1*03:01/DPB1*04:02 | 40.54% (15/37) | Non-Toxin | 0.05 | Consensus (comb.lib./smm/nn) |
| 79 | fusion protein | DLALSKYLSDLLFVF | HLA-DRB3*01:01 | 97.30% (36/37) | Non-Toxin | 0.01 | Consensus (comb.lib./smm/nn) |
| 80 | fusion protein | DLSSYYIIVRVYFPI | HLA-DRB3*01:01 | 45.95% (17/37) | Non-Toxin | 0.05 | Consensus (comb.lib./smm/nn) |
| 81 | fusion protein | DLSSYYIIVRVYFPI | HLA-DPA1*02:01/DPB1*01:01 | 45.95% (17/37) | Non-Toxin | 0.43 | Consensus (comb.lib./smm/nn) |
| 82 | fusion protein | DLVGDVRLAGVIMAG | HLA-DRB1*03:01 | 100.00% (37/37) | Non-Toxin | 0.43 | Consensus (smm/nn/sturniolo) |
| 83 | fusion protein | DNSEWISIVPNFILV | HLA-DRB5*01:01 | 97.30% (36/37) | Non-Toxin | 0.38 | Consensus (smm/nn/sturniolo) |
| 84 | fusion protein | DPVSNSMTIQAISQA | HLA-DQA1*01:02/DQB1*06:02 | 97.30% (36/37) | Non-Toxin | 0.06 | Consensus (comb.lib./smm/nn) |
| 85 | fusion protein | DTVNPSLISMLSMII | HLA-DRB1*15:01 | 100.00% (37/37) | Non-Toxin | 0.3 | Consensus (smm/nn/sturniolo) |
| 86 | fusion protein | EIGFCLITKRSVICN | HLA-DRB1*11:01 | 100.00% (37/37) | Non-Toxin | 0.42 | Consensus (smm/nn/sturniolo) |
| 87 | fusion protein | EKKRNTYSRLEDRRV | HLA-DRB1*09:01 | 100.00% (37/37) | Non-Toxin | 0.25 | Consensus (comb.lib./smm/nn) |
| 88 | fusion protein | EKTVYVLTALQDYIN | HLA-DQA1*05:01/DQB1*02:01 | 100.00% (37/37) | Non-Toxin | 0.49 | Consensus (comb.lib./smm/nn) |
| 89 | fusion protein | ELVVSSHVPRFALSN | HLA-DRB1*03:01 | 100.00% (37/37) | Non-Toxin | 0.06 | Consensus (smm/nn/sturniolo) |

| | | | | | | | |
|---|---|---|---|---|---|---|---|
| 90 | fusion protein | FGGNYETLLRTLGYA | HLA-DRB1*11:01 | 97.30% (36/37) | Non-Toxin | 0.18 | Consensus (smm/nn/sturniolo) |
| 91 | fusion protein | FISFIIVEKKRNTYS | HLA-DRB1*11:01 | 100.00% (37/37) | Non-Toxin | 0.34 | Consensus (smm/nn/sturniolo) |
| 92 | fusion protein | GGNYETLLRTLGYAT | HLA-DRB1*11:01 | 94.59% (35/37) | Non-Toxin | 0.09 | Consensus (smm/nn/sturniolo) |
| 93 | fusion protein | GLITFISFIIVEKKR | HLA-DPA1*01:03/DPB1*02:01 | 100.00% (37/37) | Non-Toxin | 0.05 | Consensus (comb.lib./smm/nn) |
| 94 | fusion protein | GNYETLLRTLGYATE | HLA-DRB1*11:01 | 94.59% (35/37) | Non-Toxin | 0.22 | Consensus (smm/nn/sturniolo) |
| 95 | fusion protein | GQIIYVDLSSYYIIV | HLA-DRB1*15:01 | 45.95% (17/37) | Non-Toxin | 0.28 | Consensus (smm/nn/sturniolo) |
| 96 | fusion protein | GQIIYVDLSSYYIIV | HLA-DRB3*01:01 | 45.95% (17/37) | Non-Toxin | 0.4 | Consensus (comb.lib./smm/nn) |
| 97 | fusion protein | GVAIGIATAAQITAG | HLA-DRB1*08:02 | 97.30% (36/37) | Non-Toxin | 0.32 | Consensus (smm/nn/sturniolo) |
| 98 | fusion protein | GVIMAGVAIGIATAA | HLA-DQA1*05:01/DQB1*03:01 | 97.30% (36/37) | Non-Toxin | 0.37 | Consensus (comb.lib./smm/nn) |
| 99 | fusion protein | HDLVGDVRLAGVIMA | HLA-DRB1*03:01 | 100.00% (37/37) | Non-Toxin | 0.05 | Consensus (smm/nn/sturniolo) |
| 100 | fusion protein | IGLITFISFIIVEKK | HLA-DPA1*01:03/DPB1*02:01 | 100.00% (37/37) | Non-Toxin | 0.05 | Consensus (comb.lib./smm/nn) |
| 101 | fusion protein | IGLITFISFIIVEKK | HLA-DRB1*15:01 | 100.00% (37/37) | Non-Toxin | 0.1 | Consensus (smm/nn/sturniolo) |
| 102 | fusion protein | IIVRVYFPILTEIQQ | HLA-DPA1*02:01/DPB1*01:01 | 100.00% (37/37) | Non-Toxin | 0.27 | Consensus (comb.lib./smm/nn) |
| 103 | fusion protein | IIVRVYFPILTEIQQ | HLA-DPA1*01:03/DPB1*02:01 | 100.00% (37/37) | Non-Toxin | 0.28 | Consensus (comb.lib./smm/nn) |
| 104 | fusion protein | IIVRVYFPILTEIQQ | HLA-DQA1*01:01/DQB1*05:01 | 100.00% (37/37) | Non-Toxin | 0.38 | Consensus (comb.lib./smm/nn) |
| 105 | fusion protein | IKMIPNVSNMSQCTG | HLA-DRB1*08:02 | 97.30% (36/37) | Non-Toxin | 0.3 | Consensus (smm/nn/sturniolo) |
| 106 | fusion protein | IKSNPLTKDIVIKMI | HLA-DRB1*03:01 | 100.00% (37/37) | Non-Toxin | 0.25 | Consensus (smm/nn/sturniolo) |
| 107 | fusion protein | ILDKRCYCNLLILIL | HLA-DPA1*03:01/DPB1*04:02 | 40.54% (15/37) | Non-Toxin | 0.06 | Consensus (comb.lib./smm/nn) |
| 108 | fusion protein | ISFIIVEKKRNTYSR | HLA-DRB1*11:01 | 100.00% (37/37) | Non-Toxin | 0.44 | Consensus (smm/nn/sturniolo) |
| 109 | fusion protein | ISIVPNFILVRNTLI | HLA-DRB1*07:01 | 100.00% (37/37) | Non-Toxin | 0.07 | Consensus (comb.lib./smm/nn) |
| 110 | fusion protein | ISIVPNFILVRNTLI | HLA-DRB1*01:01 | 100.00% (37/37) | Non-Toxin | 0.28 | Consensus (comb.lib./smm/nn) |
| 111 | fusion protein | ISIVPNFILVRNTLI | HLA-DRB1*04:05 | 100.00% (37/37) | Non-Toxin | 0.34 | Consensus (smm/nn/sturniolo) |
| 112 | fusion protein | ISMLSMIILYVLSIA | HLA-DRB4*01:01 | 100.00% (37/37) | Non-Toxin | 0.16 | Consensus (comb.lib./smm/nn) |
| 113 | fusion protein | ISMLSMIILYVLSIA | HLA-DRB1*11:01 | 100.00% (37/37) | Non-Toxin | 0.39 | Consensus (smm/nn/sturniolo) |
| 114 | fusion protein | ITGQIIYVDLSSYYI | HLA-DRB3*01:01 | 45.95% (17/37) | Non-Toxin | 0.42 | Consensus (comb.lib./smm/nn) |
| 115 | fusion protein | IVIKMIPNVSNMSQC | HLA-DRB1*08:02 | 97.30% (36/37) | Non-Toxin | 0.11 | Consensus (smm/nn/sturniolo) |
| 116 | fusion protein | IVPNFILVRNTLISN | HLA-DRB1*01:01 | 100.00% (37/37) | Non-Toxin | 0.28 | Consensus (comb.lib./smm/nn) |
| 117 | fusion protein | IVPNFILVRNTLISN | HLA-DRB1*07:01 | 100.00% (37/37) | Non-Toxin | 0.29 | Consensus (comb.lib./smm/nn) |
| 118 | fusion protein | IVPNFILVRNTLISN | HLA-DRB1*04:05 | 100.00% (37/37) | Non-Toxin | 0.34 | Consensus (smm/nn/sturniolo) |
| 119 | fusion protein | IVPNFILVRNTLISN | HLA-DRB1*08:02 | 100.00% (37/37) | Non-Toxin | 0.42 | Consensus (smm/nn/sturniolo) |
| 120 | fusion protein | IVRVYFPILTEIQQA | HLA-DPA1*02:01/DPB1*01:01 | 100.00% (37/37) | Non-Toxin | 0.27 | Consensus (comb.lib./smm/nn) |
| 121 | fusion protein | IVRVYFPILTEIQQA | HLA-DQA1*01:01/DQB1*05:01 | 100.00% (37/37) | Non-Toxin | 0.43 | Consensus (comb.lib./smm/nn) |
| 122 | fusion protein | IVRVYFPILTEIQQA | HLA-DPA1*01:03/DPB1*02:01 | 100.00% (37/37) | Non-Toxin | 0.47 | Consensus (comb.lib./smm/nn) |
| 123 | fusion protein | KCPRELVVSSHVPRF | HLA-DRB1*03:01 | 100.00% (37/37) | Non-Toxin | 0.06 | Consensus (smm/nn/sturniolo) |
| 124 | fusion protein | KDIVIKMIPNVSNMS | HLA-DRB1*08:02 | 97.30% (36/37) | Non-Toxin | 0.16 | Consensus (smm/nn/sturniolo) |
| 125 | fusion protein | KIKSNPLTKDIVIKM | HLA-DRB1*03:01 | 100.00% (37/37) | Non-Toxin | 0.25 | Consensus (smm/nn/sturniolo) |
| 126 | fusion protein | KNNTHDLVGDVRLAG | HLA-DRB1*03:01 | 100.00% (37/37) | Non-Toxin | 0.08 | Consensus (smm/nn/sturniolo) |
| 127 | fusion protein | KQTELSLDLALSKYL | HLA-DRB1*03:01 | 97.30% (36/37) | Non-Toxin | 0.37 | Consensus (smm/nn/sturniolo) |
| 128 | fusion protein | KRCYCNLLILILMIS | HLA-DPA1*03:01/DPB1*04:02 | 40.54% (15/37) | Non-Toxin | 0.05 | Consensus (comb.lib./smm/nn) |
| 129 | fusion protein | KRCYCNLLILILMIS | HLA-DRB1*11:01 | 40.54% (15/37) | Non-Toxin | 0.19 | Consensus (smm/nn/sturniolo) |
| 130 | fusion protein | KSNPLTKDIVIKMIP | HLA-DRB1*03:01 | 97.30% (36/37) | Non-Toxin | 0.25 | Consensus (smm/nn/sturniolo) |
| 131 | fusion protein | KTVYVLTALQDYINT | HLA-DQA1*05:01/DQB1*02:01 | 100.00% (37/37) | Non-Toxin | 0.49 | Consensus (comb.lib./smm/nn) |
| 132 | fusion protein | KYLSDLLFVFGPNLQ | HLA-DRB3*01:01 | 97.30% (36/37) | Non-Toxin | 0.01 | Consensus (comb.lib./smm/nn) |
| 133 | fusion protein | LAGVIMAGVAIGIAT | HLA-DQA1*05:01/DQB1*03:01 | 97.30% (36/37) | Non-Toxin | 0.32 | Consensus (comb.lib./smm/nn) |
| 134 | fusion protein | LALSKYLSDLLFVFG | HLA-DRB3*01:01 | 97.30% (36/37) | Non-Toxin | 0.01 | Consensus (comb.lib./smm/nn) |
| 135 | fusion protein | LCIGLITFISFIIVE | HLA-DPA1*01:03/DPB1*02:01 | 100.00% (37/37) | Non-Toxin | 0.04 | Consensus (comb.lib./smm/nn) |
| 136 | fusion protein | LCIGLITFISFIIVE | HLA-DRB1*15:01 | 100.00% (37/37) | Non-Toxin | 0.11 | Consensus (smm/nn/sturniolo) |
| 137 | fusion protein | LDKRCYCNLLILILM | HLA-DPA1*03:01/DPB1*04:02 | 40.54% (15/37) | Non-Toxin | 0.06 | Consensus (comb.lib./smm/nn) |

| | | | | | | | |
|---|---|---|---|---|---|---|---|
| 138 | fusion protein | LISMLSMIILYVLSI | HLA-DRB4*01:01 | 100.00% (37/37) | Non-Toxin | 0.16 | Consensus (comb.lib./smm/nn) |
| 139 | fusion protein | LISMLSMIILYVLSI | HLA-DRB1*11:01 | 100.00% (37/37) | Non-Toxin | 0.3 | Consensus (smm/nn/sturniolo) |
| 140 | fusion protein | LITFISFIIVEKKRN | HLA-DPA1*01:03/DPB1*02:01 | 100.00% (37/37) | Non-Toxin | 0.37 | Consensus (comb.lib./smm/nn) |
| 141 | fusion protein | LLILILMISECSVGI | HLA-DRB1*11:01 | 72.97% (27/37) | Non-Toxin | 0.3 | Consensus (smm/nn/sturniolo) |
| 142 | fusion protein | LLPVSFNNDNSEWIS | HLA-DRB3*01:01 | 97.30% (36/37) | Non-Toxin | 0.5 | Consensus (comb.lib./smm/nn) |
| 143 | fusion protein | LPVSFNNDNSEWISI | HLA-DRB3*01:01 | 97.30% (36/37) | Non-Toxin | 0.46 | Consensus (comb.lib./smm/nn) |
| 144 | fusion protein | LQDPVSNSMTIQAIS | HLA-DQA1*01:02/DQB1*06:02 | 97.30% (36/37) | Non-Toxin | 0.06 | Consensus (comb.lib./smm/nn) |
| 145 | fusion protein | LRTLGYATEDFDDLL | HLA-DQA1*05:01/DQB1*02:01 | 97.30% (36/37) | Non-Toxin | 0.46 | Consensus (comb.lib./smm/nn) |
| 146 | fusion protein | LSKYLSDLLFVFGPN | HLA-DRB3*01:01 | 97.30% (36/37) | Non-Toxin | 0.01 | Consensus (comb.lib./smm/nn) |
| 147 | fusion protein | LSMIILYVLSIASLC | HLA-DRB1*11:01 | 100.00% (37/37) | Non-Toxin | 0.05 | Consensus (smm/nn/sturniolo) |
| 148 | fusion protein | LSMIILYVLSIASLC | HLA-DRB1*15:01 | 100.00% (37/37) | Non-Toxin | 0.23 | Consensus (smm/nn/sturniolo) |
| 149 | fusion protein | LSMIILYVLSIASLC | HLA-DRB1*04:01 | 100.00% (37/37) | Non-Toxin | 0.42 | Consensus (smm/nn/sturniolo) |
| 150 | fusion protein | LSSYYIIVRVYFPIL | HLA-DPA1*01:03/DPB1*02:01 | 45.95% (17/37) | Non-Toxin | 0.04 | Consensus (comb.lib./smm/nn) |
| 151 | fusion protein | LSSYYIIVRVYFPIL | HLA-DRB3*01:01 | 45.95% (17/37) | Non-Toxin | 0.05 | Consensus (comb.lib./smm/nn) |
| 152 | fusion protein | LSSYYIIVRVYFPIL | HLA-DPA1*01/DPB1*04:01 | 45.95% (17/37) | Non-Toxin | 0.15 | Consensus (comb.lib./smm/nn) |
| 153 | fusion protein | LSSYYIIVRVYFPIL | HLA-DPA1*02:01/DPB1*01:01 | 45.95% (17/37) | Non-Toxin | 0.27 | Consensus (comb.lib./smm/nn) |
| 154 | fusion protein | LTEIQQAYIQELLPV | HLA-DQA1*05:01/DQB1*02:01 | 100.00% (37/37) | Non-Toxin | 0.3 | Consensus (comb.lib./smm/nn) |
| 155 | fusion protein | LTKDIVIKMIPNVSN | HLA-DRB1*08:02 | 97.30% (36/37) | Non-Toxin | 0.22 | Consensus (smm/nn/sturniolo) |
| 156 | fusion protein | LTKDIVIKMIPNVSN | HLA-DRB1*03:01 | 97.30% (36/37) | Non-Toxin | 0.25 | Consensus (smm/nn/sturniolo) |
| 157 | fusion protein | LVGDVRLAGVIMAGV | HLA-DRB1*03:01 | 100.00% (37/37) | Non-Toxin | 0.41 | Consensus (smm/nn/sturniolo) |
| 158 | fusion protein | LVVSSHVPRFALSNG | HLA-DRB1*03:01 | 100.00% (37/37) | Non-Toxin | 0.4 | Consensus (smm/nn/sturniolo) |
| 159 | fusion protein | MIILYVLSIASLCIG | HLA-DRB1*11:01 | 100.00% (37/37) | Non-Toxin | 0.05 | Consensus (smm/nn/sturniolo) |
| 160 | fusion protein | MIILYVLSIASLCIG | HLA-DRB1*15:01 | 100.00% (37/37) | Non-Toxin | 0.28 | Consensus (smm/nn/sturniolo) |
| 161 | fusion protein | MIILYVLSIASLCIG | HLA-DRB1*04:01 | 100.00% (37/37) | Non-Toxin | 0.4 | Consensus (smm/nn/sturniolo) |
| 162 | fusion protein | MLSMIILYVLSIASL | HLA-DRB1*11:01 | 100.00% (37/37) | Non-Toxin | 0.05 | Consensus (smm/nn/sturniolo) |
| 163 | fusion protein | MLSMIILYVLSIASL | HLA-DRB1*15:01 | 100.00% (37/37) | Non-Toxin | 0.23 | Consensus (smm/nn/sturniolo) |
| 164 | fusion protein | MLSMIILYVLSIASL | HLA-DRB1*04:01 | 100.00% (37/37) | Non-Toxin | 0.46 | Consensus (smm/nn/sturniolo) |
| 165 | fusion protein | MLSMIILYVLSIASL | HLA-DPA1*02:01/DPB1*01:01 | 100.00% (37/37) | Non-Toxin | 0.51 | Consensus (comb.lib./smm/nn) |
| 166 | fusion protein | MVVILDKRCYCNLLI | HLA-DRB1*03:01 | 40.54% (15/37) | Non-Toxin | 0.15 | Consensus (smm/nn/sturniolo) |
| 167 | fusion protein | MVVILDKRCYCNLLI | HLA-DRB1*11:01 | 40.54% (15/37) | Non-Toxin | 0.48 | Consensus (smm/nn/sturniolo) |
| 168 | fusion protein | NDNSEWISIVPNFIL | HLA-DRB5*01:01 | 97.30% (36/37) | Non-Toxin | 0.38 | Consensus (smm/nn/sturniolo) |
| 169 | fusion protein | NFILVRNTLISNIEI | HLA-DRB1*08:02 | 100.00% (37/37) | Non-Toxin | 0.4 | Consensus (smm/nn/sturniolo) |
| 170 | fusion protein | NLLILILMISECSVG | HLA-DRB1*11:01 | 72.97% (27/37) | Non-Toxin | 0.26 | Consensus (smm/nn/sturniolo) |
| 171 | fusion protein | NLLILILMISECSVG | HLA-DRB1*04:05 | 72.97% (27/37) | Non-Toxin | 0.47 | Consensus (smm/nn/sturniolo) |
| 172 | fusion protein | NLQDPVSNSMTIQAI | HLA-DQA1*01:02/DQB1*06:02 | 97.30% (36/37) | Non-Toxin | 0.18 | Consensus (comb.lib./smm/nn) |
| 173 | fusion protein | NNDNSEWISIVPNFI | HLA-DRB5*01:01 | 97.30% (36/37) | Non-Toxin | 0.38 | Consensus (smm/nn/sturniolo) |
| 174 | fusion protein | NNTHDLVGDVRLAGV | HLA-DRB1*03:01 | 100.00% (37/37) | Non-Toxin | 0.06 | Consensus (smm/nn/sturniolo) |
| 175 | fusion protein | NPLTKDIVIKMIPNV | HLA-DRB1*03:01 | 97.30% (36/37) | Non-Toxin | 0.25 | Consensus (smm/nn/sturniolo) |
| 176 | fusion protein | NPSLISMLSMIILYV | HLA-DRB4*01:01 | 100.00% (37/37) | Non-Toxin | 0.17 | Consensus (comb.lib./smm/nn) |
| 177 | fusion protein | NPSLISMLSMIILYV | HLA-DRB1*15:01 | 100.00% (37/37) | Non-Toxin | 0.27 | Consensus (smm/nn/sturniolo) |
| 178 | fusion protein | NSEWISIVPNFILVR | HLA-DRB5*01:01 | 97.30% (36/37) | Non-Toxin | 0.38 | Consensus (smm/nn/sturniolo) |
| 179 | fusion protein | NSMTIQAISQAFGGN | HLA-DRB1*08:02 | 97.30% (36/37) | Non-Toxin | 0.35 | Consensus (smm/nn/sturniolo) |
| 180 | fusion protein | NTHDLVGDVRLAGVI | HLA-DRB1*03:01 | 100.00% (37/37) | Non-Toxin | 0.05 | Consensus (smm/nn/sturniolo) |
| 181 | fusion protein | PLTKDIVIKMIPNVS | HLA-DRB1*03:01 | 97.30% (36/37) | Non-Toxin | 0.25 | Consensus (smm/nn/sturniolo) |
| 182 | fusion protein | PNFILVRNTLISNIE | HLA-DRB1*01:01 | 100.00% (37/37) | Non-Toxin | 0.28 | Consensus (comb.lib./smm/nn) |
| 183 | fusion protein | PNFILVRNTLISNIE | HLA-DRB1*08:02 | 100.00% (37/37) | Non-Toxin | 0.34 | Consensus (smm/nn/sturniolo) |
| 184 | fusion protein | PNFILVRNTLISNIE | HLA-DRB1*04:05 | 100.00% (37/37) | Non-Toxin | 0.34 | Consensus (smm/nn/sturniolo) |
| 185 | fusion protein | PRELVVSSHVPRFAL | HLA-DRB1*03:01 | 100.00% (37/37) | Non-Toxin | 0.06 | Consensus (smm/nn/sturniolo) |

| | | | | | | | |
|---|---|---|---|---|---|---|---|
| 186 | fusion protein | PSLISMLSMIILYVL | HLA-DRB4*01:01 | 100.00% (37/37) | Non-Toxin | 0.15 | Consensus (comb.lib./smm/nn) |
| 187 | fusion protein | PSLISMLSMIILYVL | HLA-DRB1*15:01 | 100.00% (37/37) | Non-Toxin | 0.27 | Consensus (smm/nn/sturniolo) |
| 188 | fusion protein | PVSFNNDNSEWISIV | HLA-DRB3*01:01 | 97.30% (36/37) | Non-Toxin | 0.48 | Consensus (comb.lib./smm/nn) |
| 189 | fusion protein | PVSNSMTIQAISQAF | HLA-DQA1*01:02/DQB1*06:02 | 97.30% (36/37) | Non-Toxin | 0.07 | Consensus (comb.lib./smm/nn) |
| 190 | fusion protein | QDPVSNSMTIQAISQ | HLA-DQA1*01:02/DQB1*06:02 | 97.30% (36/37) | Non-Toxin | 0.06 | Consensus (comb.lib./smm/nn) |
| 191 | fusion protein | QIIYVDLSSYYIIVR | HLA-DRB1*15:01 | 45.95% (17/37) | Non-Toxin | 0.33 | Consensus (smm/nn/sturniolo) |
| 192 | fusion protein | QTELSLDLALSKYLS | HLA-DRB1*03:01 | 97.30% (36/37) | Non-Toxin | 0.28 | Consensus (smm/nn/sturniolo) |
| 193 | fusion protein | RCYCNLLILILMISE | HLA-DPA1*03:01/DPB1*04:02 | 40.54% (15/37) | Non-Toxin | 0.05 | Consensus (comb.lib./smm/nn) |
| 194 | fusion protein | RCYCNLLILILMISE | HLA-DRB1*11:01 | 40.54% (15/37) | Non-Toxin | 0.17 | Consensus (smm/nn/sturniolo) |
| 195 | fusion protein | RELVVSSHVPRFALS | HLA-DRB1*03:01 | 100.00% (37/37) | Non-Toxin | 0.06 | Consensus (smm/nn/sturniolo) |
| 196 | fusion protein | RLAGVIMAGVAIGIA | HLA-DQA1*05:01/DQB1*03:01 | 100.00% (37/37) | Non-Toxin | 0.46 | Consensus (comb.lib./smm/nn) |
| 197 | fusion protein | RTLGYATEDFDDLLE | HLA-DQA1*05:01/DQB1*02:01 | 97.30% (36/37) | Non-Toxin | 0.5 | Consensus (comb.lib./smm/nn) |
| 198 | fusion protein | SEWISIVPNFILVRN | HLA-DRB5*01:01 | 97.30% (36/37) | Non-Toxin | 0.38 | Consensus (smm/nn/sturniolo) |
| 199 | fusion protein | SITGQIIYVDLSSYY | HLA-DRB3*01:01 | 45.95% (17/37) | Non-Toxin | 0.49 | Consensus (comb.lib./smm/nn) |
| 200 | fusion protein | SIVPNFILVRNTLIS | HLA-DRB1*07:01 | 100.00% (37/37) | Non-Toxin | 0.13 | Consensus (comb.lib./smm/nn) |
| 201 | fusion protein | SIVPNFILVRNTLIS | HLA-DRB1*01:01 | 100.00% (37/37) | Non-Toxin | 0.28 | Consensus (comb.lib./smm/nn) |
| 202 | fusion protein | SIVPNFILVRNTLIS | HLA-DRB1*04:05 | 100.00% (37/37) | Non-Toxin | 0.34 | Consensus (smm/nn/sturniolo) |
| 203 | fusion protein | SKYLSDLLFVFGPNL | HLA-DRB3*01:01 | 97.30% (36/37) | Non-Toxin | 0.01 | Consensus (comb.lib./smm/nn) |
| 204 | fusion protein | SLCIGLITFISFIIV | HLA-DPA1*01:03/DPB1*02:01 | 100.00% (37/37) | Non-Toxin | 0.04 | Consensus (comb.lib./smm/nn) |
| 205 | fusion protein | SLCIGLITFISFIIV | HLA-DRB1*15:01 | 100.00% (37/37) | Non-Toxin | 0.11 | Consensus (smm/nn/sturniolo) |
| 206 | fusion protein | SLISMLSMIILYVLS | HLA-DRB4*01:01 | 100.00% (37/37) | Non-Toxin | 0.15 | Consensus (comb.lib./smm/nn) |
| 207 | fusion protein | SMIILYVLSIASLCI | HLA-DRB1*11:01 | 100.00% (37/37) | Non-Toxin | 0.05 | Consensus (smm/nn/sturniolo) |
| 208 | fusion protein | SMIILYVLSIASLCI | HLA-DRB1*15:01 | 100.00% (37/37) | Non-Toxin | 0.23 | Consensus (smm/nn/sturniolo) |
| 209 | fusion protein | SMIILYVLSIASLCI | HLA-DRB1*04:01 | 100.00% (37/37) | Non-Toxin | 0.38 | Consensus (smm/nn/sturniolo) |
| 210 | fusion protein | SMLSMIILYVLSIAS | HLA-DRB1*11:01 | 100.00% (37/37) | Non-Toxin | 0.05 | Consensus (smm/nn/sturniolo) |
| 211 | fusion protein | SMLSMIILYVLSIAS | HLA-DRB1*15:01 | 100.00% (37/37) | Non-Toxin | 0.23 | Consensus (smm/nn/sturniolo) |
| 212 | fusion protein | SMTIQAISQAFGGNY | HLA-DRB1*08:02 | 97.30% (36/37) | Non-Toxin | 0.37 | Consensus (smm/nn/sturniolo) |
| 213 | fusion protein | SNPLTKDIVIKMIPN | HLA-DRB1*03:01 | 97.30% (36/37) | Non-Toxin | 0.25 | Consensus (smm/nn/sturniolo) |
| 214 | fusion protein | SSYYIIVRVYFPILT | HLA-DPA1*01:03/DPB1*02:01 | 45.95% (17/37) | Non-Toxin | 0.04 | Consensus (comb.lib./smm/nn) |
| 215 | fusion protein | SSYYIIVRVYFPILT | HLA-DRB3*01:01 | 45.95% (17/37) | Non-Toxin | 0.05 | Consensus (comb.lib./smm/nn) |
| 216 | fusion protein | SSYYIIVRVYFPILT | HLA-DPA1*01/DPB1*04:01 | 45.95% (17/37) | Non-Toxin | 0.15 | Consensus (comb.lib./smm/nn) |
| 217 | fusion protein | SSYYIIVRVYFPILT | HLA-DPA1*02:01/DPB1*01:01 | 45.95% (17/37) | Non-Toxin | 0.27 | Consensus (comb.lib./smm/nn) |
| 218 | fusion protein | SYYIIVRVYFPILTE | HLA-DPA1*01:03/DPB1*02:01 | 45.95% (17/37) | Non-Toxin | 0.05 | Consensus (comb.lib./smm/nn) |
| 219 | fusion protein | SYYIIVRVYFPILTE | HLA-DPA1*01/DPB1*04:01 | 45.95% (17/37) | Non-Toxin | 0.16 | Consensus (comb.lib./smm/nn) |
| 220 | fusion protein | SYYIIVRVYFPILTE | HLA-DRB3*01:01 | 45.95% (17/37) | Non-Toxin | 0.24 | Consensus (comb.lib./smm/nn) |
| 221 | fusion protein | SYYIIVRVYFPILTE | HLA-DPA1*02:01/DPB1*01:01 | 45.95% (17/37) | Non-Toxin | 0.27 | Consensus (comb.lib./smm/nn) |
| 222 | fusion protein | SYYIIVRVYFPILTE | HLA-DQA1*01:01/DQB1*05:01 | 45.95% (17/37) | Non-Toxin | 0.3 | Consensus (comb.lib./smm/nn) |
| 223 | fusion protein | TGQIIYVDLSSYYII | HLA-DRB1*15:01 | 45.95% (17/37) | Non-Toxin | 0.38 | Consensus (smm/nn/sturniolo) |
| 224 | fusion protein | TGQIIYVDLSSYYII | HLA-DRB3*01:01 | 45.95% (17/37) | Non-Toxin | 0.39 | Consensus (comb.lib./smm/nn) |
| 225 | fusion protein | THDLVGDVRLAGVIM | HLA-DRB1*03:01 | 100.00% (37/37) | Non-Toxin | 0.05 | Consensus (smm/nn/sturniolo) |
| 226 | fusion protein | TKDIVIKMIPNVSNM | HLA-DRB1*08:02 | 97.30% (36/37) | Non-Toxin | 0.22 | Consensus (smm/nn/sturniolo) |
| 227 | fusion protein | TVNPSLISMLSMIIL | HLA-DRB4*01:01 | 100.00% (37/37) | Non-Toxin | 0.22 | Consensus (comb.lib./smm/nn) |
| 228 | fusion protein | TVNPSLISMLSMIIL | HLA-DRB1*15:01 | 100.00% (37/37) | Non-Toxin | 0.27 | Consensus (smm/nn/sturniolo) |
| 229 | fusion protein | VIKMIPNVSNMSQCT | HLA-DRB1*08:02 | 97.30% (36/37) | Non-Toxin | 0.21 | Consensus (smm/nn/sturniolo) |
| 230 | fusion protein | VNPSLISMLSMIILY | HLA-DRB4*01:01 | 100.00% (37/37) | Non-Toxin | 0.22 | Consensus (comb.lib./smm/nn) |
| 231 | fusion protein | VNPSLISMLSMIILY | HLA-DRB1*15:01 | 100.00% (37/37) | Non-Toxin | 0.27 | Consensus (smm/nn/sturniolo) |
| 232 | fusion protein | VPNFILVRNTLISNI | HLA-DRB1*01:01 | 100.00% (37/37) | Non-Toxin | 0.28 | Consensus (comb.lib./smm/nn) |
| 233 | fusion protein | VPNFILVRNTLISNI | HLA-DRB1*08:02 | 100.00% (37/37) | Non-Toxin | 0.33 | Consensus (smm/nn/sturniolo) |

| | | | | | | | |
|---|---|---|---|---|---|---|---|
| 234 | fusion protein | VPNFILVRNTLISNI | HLA-DRB1*04:05 | 100.00% (37/37) | Non-Toxin | 0.34 | Consensus (smm/nn/sturniolo) |
| 235 | fusion protein | VPNFILVRNTLISNI | HLA-DRB1*07:01 | 100.00% (37/37) | Non-Toxin | 0.43 | Consensus (comb.lib./smm/nn) |
| 236 | fusion protein | VRLAGVIMAGVAIGI | HLA-DQA1*05:01/DQB1*03:01 | 100.00% (37/37) | Non-Toxin | 0.48 | Consensus (comb.lib./smm/nn) |
| 237 | fusion protein | VSNSMTIQAISQAFG | HLA-DQA1*01:02/DQB1*06:02 | 97.30% (36/37) | Non-Toxin | 0.46 | Consensus (comb.lib./smm/nn) |
| 238 | fusion protein | YCNLLILILMISECS | HLA-DPA1*03:01/DPB1*04:02 | 40.54% (15/37) | Non-Toxin | 0.12 | Consensus (comb.lib./smm/nn) |
| 239 | fusion protein | YCNLLILILMISECS | HLA-DRB1*11:01 | 40.54% (15/37) | Non-Toxin | 0.12 | Consensus (smm/nn/sturniolo) |
| 240 | fusion protein | YCNLLILILMISECS | HLA-DRB1*04:05 | 40.54% (15/37) | Non-Toxin | 0.47 | Consensus (smm/nn/sturniolo) |
| 241 | fusion protein | YIIVRVYFPILTEIQ | HLA-DPA1*01:03/DPB1*02:01 | 100.00% (37/37) | Non-Toxin | 0.08 | Consensus (comb.lib./smm/nn) |
| 242 | fusion protein | YIIVRVYFPILTEIQ | HLA-DPA1*01/DPB1*04:01 | 100.00% (37/37) | Non-Toxin | 0.15 | Consensus (comb.lib./smm/nn) |
| 243 | fusion protein | YIIVRVYFPILTEIQ | HLA-DPA1*02:01/DPB1*01:01 | 100.00% (37/37) | Non-Toxin | 0.27 | Consensus (comb.lib./smm/nn) |
| 244 | fusion protein | YIIVRVYFPILTEIQ | HLA-DQA1*01:01/DQB1*05:01 | 100.00% (37/37) | Non-Toxin | 0.28 | Consensus (comb.lib./smm/nn) |
| 245 | fusion protein | YLSDLLFVFGPNLQD | HLA-DRB3*01:01 | 97.30% (36/37) | Non-Toxin | 0.01 | Consensus (comb.lib./smm/nn) |
| 246 | fusion protein | YYIIVRVYFPILTEI | HLA-DPA1*01:03/DPB1*02:01 | 100.00% (37/37) | Non-Toxin | 0.04 | Consensus (comb.lib./smm/nn) |
| 247 | fusion protein | YYIIVRVYFPILTEI | HLA-DPA1*01/DPB1*04:01 | 100.00% (37/37) | Non-Toxin | 0.14 | Consensus (comb.lib./smm/nn) |
| 248 | fusion protein | YYIIVRVYFPILTEI | HLA-DPA1*02:01/DPB1*01:01 | 100.00% (37/37) | Non-Toxin | 0.27 | Consensus (comb.lib./smm/nn) |
| 249 | fusion protein | YYIIVRVYFPILTEI | HLA-DQA1*01:01/DQB1*05:01 | 100.00% (37/37) | Non-Toxin | 0.28 | Consensus (comb.lib./smm/nn) |
| 250 | fusion protein | YYIIVRVYFPILTEI | HLA-DRB3*01:01 | 100.00% (37/37) | Non-Toxin | 0.36 | Consensus (comb.lib./smm/nn) |
| **S.No.** | **Protein** | **Peptide** | **Allele** | **Conservancy** | **Toxicity** | **Percentile rank** | **Method used** |
| 251 | glycoprotein | AFLIDRINWISAGVF | HLA-DRB3*01:01 | 100.00% (66/66) | Non-Toxin | 0.01 | Consensus (comb.lib./smm/nn) |
| 252 | glycoprotein | ALLGSIVIIVMNIMI | HLA-DRB4*01:01 | 98.48% (65/66) | Non-Toxin | 0.2 | Consensus (comb.lib./smm/nn) |
| 253 | glycoprotein | ALLGSIVIIVMNIMI | HLA-DRB1*15:01 | 98.48% (65/66) | Non-Toxin | 0.34 | Consensus (smm/nn/sturniolo) |
| 254 | glycoprotein | AMDEGYFAYSHLERI | HLA-DPA1*03:01/DPB1*04:02 | 24.24% (16/66) | Non-Toxin | 0.08 | Consensus (comb.lib./smm/nn) |
| 255 | glycoprotein | AMDEGYFAYSHLERI | HLA-DPA1*02:01/DPB1*01:01 | 24.24% (16/66) | Non-Toxin | 0.25 | Consensus (comb.lib./smm/nn) |
| 256 | glycoprotein | AMDEGYFAYSHLERI | HLA-DPA1*01:03/DPB1*02:01 | 24.24% (16/66) | Non-Toxin | 0.31 | Consensus (comb.lib./smm/nn) |
| 257 | glycoprotein | ASFSWDTMIKFGDVL | HLA-DRB3*01:01 | 19.70% (13/66) | Non-Toxin | 0.01 | Consensus (comb.lib./smm/nn) |
| 258 | glycoprotein | AVIKDALQGIQQQIK | HLA-DRB4*01:01 | 24.24% (16/66) | Non-Toxin | 0.35 | Consensus (comb.lib./smm/nn) |
| 259 | glycoprotein | AYSHLERIGSCSRGV | HLA-DRB1*11:01 | 24.24% (16/66) | Non-Toxin | 0.38 | Consensus (smm/nn/sturniolo) |
| 260 | glycoprotein | CSAVYNNEFYYVLCA | HLA-DPA1*01/DPB1*04:01 | 90.91% (60/66) | Non-Toxin | 0.37 | Consensus (comb.lib./smm/nn) |
| 261 | glycoprotein | CWEGVYNDAFLIDRI | HLA-DRB3*01:01 | 100.00% (66/66) | Non-Toxin | 0.18 | Consensus (comb.lib./smm/nn) |
| 262 | glycoprotein | CWEGVYNDAFLIDRI | HLA-DPA1*01:03/DPB1*02:01 | 100.00% (66/66) | Non-Toxin | 0.34 | Consensus (comb.lib./smm/nn) |
| 263 | glycoprotein | DAFLIDRINWISAGV | HLA-DRB3*01:01 | 100.00% (66/66) | Non-Toxin | 0.01 | Consensus (comb.lib./smm/nn) |
| 264 | glycoprotein | DAFLIDRINWISAGV | HLA-DRB1*03:01 | 100.00% (66/66) | Non-Toxin | 0.22 | Consensus (smm/nn/sturniolo) |
| 265 | glycoprotein | DALQGIQQQIKGLAD | HLA-DRB4*01:01 | 24.24% (16/66) | Non-Toxin | 0.35 | Consensus (comb.lib./smm/nn) |
| 266 | glycoprotein | DEGYFAYSHLERIGS | HLA-DPA1*03:01/DPB1*04:02 | 24.24% (16/66) | Non-Toxin | 0.03 | Consensus (comb.lib./smm/nn) |
| 267 | glycoprotein | DEGYFAYSHLERIGS | HLA-DPA1*01:03/DPB1*02:01 | 24.24% (16/66) | Non-Toxin | 0.22 | Consensus (comb.lib./smm/nn) |
| 268 | glycoprotein | DEGYFAYSHLERIGS | HLA-DPA1*02:01/DPB1*01:01 | 24.24% (16/66) | Non-Toxin | 0.29 | Consensus (comb.lib./smm/nn) |
| 269 | glycoprotein | DEGYFAYSHLERIGS | HLA-DRB1*11:01 | 24.24% (16/66) | Non-Toxin | 0.38 | Consensus (smm/nn/sturniolo) |
| 270 | glycoprotein | DEGYFAYSHLERIGS | HLA-DPA1*02:01/DPB1*14:01 | 24.24% (16/66) | Non-Toxin | 0.48 | NetMHCIIpan |
| 271 | glycoprotein | DEGYFAYSHLERIGS | HLA-DPA1*01/DPB1*04:01 | 24.24% (16/66) | Non-Toxin | 0.51 | Consensus (comb.lib./smm/nn) |
| 272 | glycoprotein | DSKILSAFNTVIALL | HLA-DPA1*03:01/DPB1*04:02 | 98.48% (65/66) | Non-Toxin | 0.44 | Consensus (comb.lib./smm/nn) |
| 273 | glycoprotein | EGVYNDAFLIDRINW | HLA-DPA1*01:03/DPB1*02:01 | 100.00% (66/66) | Non-Toxin | 0.35 | Consensus (comb.lib./smm/nn) |
| 274 | glycoprotein | EGYFAYSHLERIGSC | HLA-DPA1*03:01/DPB1*04:02 | 24.24% (16/66) | Non-Toxin | 0.02 | Consensus (comb.lib./smm/nn) |
| 275 | glycoprotein | EGYFAYSHLERIGSC | HLA-DPA1*01:03/DPB1*02:01 | 24.24% (16/66) | Non-Toxin | 0.21 | Consensus (comb.lib./smm/nn) |
| 276 | glycoprotein | EGYFAYSHLERIGSC | HLA-DRB1*11:01 | 24.24% (16/66) | Non-Toxin | 0.38 | Consensus (smm/nn/sturniolo) |
| 277 | glycoprotein | EGYFAYSHLERIGSC | HLA-DPA1*02:01/DPB1*01:01 | 24.24% (16/66) | Non-Toxin | 0.38 | Consensus (comb.lib./smm/nn) |
| 278 | glycoprotein | EGYFAYSHLERIGSC | HLA-DPA1*01/DPB1*04:01 | 24.24% (16/66) | Non-Toxin | 0.45 | Consensus (comb.lib./smm/nn) |
| 279 | glycoprotein | EGYFAYSHLERIGSC | HLA-DPA1*02:01/DPB1*14:01 | 24.24% (16/66) | Non-Toxin | 0.5 | NetMHCIIpan |
| 280 | glycoprotein | EICWEGVYNDAFLID | HLA-DRB3*01:01 | 25.76% (17/66) | Non-Toxin | 0.17 | Consensus (comb.lib./smm/nn) |

| | | | | | | | |
|---|---|---|---|---|---|---|---|
| 281 | glycoprotein | EICWEGVYNDAFLID | HLA-DPA1*01:03/DPB1*02:01 | 25.76% (17/66) | Non-Toxin | 0.37 | Consensus (comb.lib./smm/nn) |
| 282 | glycoprotein | FAYSHLERIGSCSRG | HLA-DRB1*11:01 | 24.24% (16/66) | Non-Toxin | 0.38 | Consensus (smm/nn/sturniolo) |
| 283 | glycoprotein | FLIDRINWISAGVFL | HLA-DRB3*01:01 | 100.00% (66/66) | Non-Toxin | 0.01 | Consensus (comb.lib./smm/nn) |
| 284 | glycoprotein | FSWDTMIKFGDVLTV | HLA-DRB3*01:01 | 19.70% (13/66) | Non-Toxin | 0.01 | Consensus (comb.lib./smm/nn) |
| 285 | glycoprotein | FYQASFSWDTMIKFG | HLA-DRB3*01:01 | 100.00% (66/66) | Non-Toxin | 0.26 | Consensus (comb.lib./smm/nn) |
| 286 | glycoprotein | GSIVIIVMNIMIIQN | HLA-DRB4*01:01 | 98.48% (65/66) | Non-Toxin | 0.15 | Consensus (comb.lib./smm/nn) |
| 287 | glycoprotein | GSIVIIVMNIMIIQN | HLA-DRB1*15:01 | 98.48% (65/66) | Non-Toxin | 0.31 | Consensus (smm/nn/sturniolo) |
| 288 | glycoprotein | GVYNDAFLIDRINWI | HLA-DRB3*01:01 | 100.00% (66/66) | Non-Toxin | 0.01 | Consensus (comb.lib./smm/nn) |
| 289 | glycoprotein | GVYNDAFLIDRINWI | HLA-DRB1*03:01 | 100.00% (66/66) | Non-Toxin | 0.22 | Consensus (smm/nn/sturniolo) |
| 290 | glycoprotein | GYFAYSHLERIGSCS | HLA-DPA1*03:01/DPB1*04:02 | 24.24% (16/66) | Non-Toxin | 0.03 | Consensus (comb.lib./smm/nn) |
| 291 | glycoprotein | GYFAYSHLERIGSCS | HLA-DPA1*01:03/DPB1*02:01 | 24.24% (16/66) | Non-Toxin | 0.26 | Consensus (comb.lib./smm/nn) |
| 292 | glycoprotein | GYFAYSHLERIGSCS | HLA-DRB1*11:01 | 24.24% (16/66) | Non-Toxin | 0.38 | Consensus (smm/nn/sturniolo) |
| 293 | glycoprotein | HCSAVYNNEFYYVLC | HLA-DPA1*01/DPB1*04:01 | 90.91% (60/66) | Non-Toxin | 0.38 | Consensus (comb.lib./smm/nn) |
| 294 | glycoprotein | IALLGSIVIIVMNIM | HLA-DRB4*01:01 | 98.48% (65/66) | Non-Toxin | 0.21 | Consensus (comb.lib./smm/nn) |
| 295 | glycoprotein | IALLGSIVIIVMNIM | HLA-DRB1*15:01 | 98.48% (65/66) | Non-Toxin | 0.46 | Consensus (smm/nn/sturniolo) |
| 296 | glycoprotein | ICWEGVYNDAFLIDR | HLA-DRB3*01:01 | 25.76% (17/66) | Non-Toxin | 0.17 | Consensus (comb.lib./smm/nn) |
| 297 | glycoprotein | ICWEGVYNDAFLIDR | HLA-DPA1*01:03/DPB1*02:01 | 25.76% (17/66) | Non-Toxin | 0.36 | Consensus (comb.lib./smm/nn) |
| 298 | glycoprotein | IIVMNIMIIQNYTRS | HLA-DRB4*01:01 | 98.48% (65/66) | Non-Toxin | 0.1 | Consensus (comb.lib./smm/nn) |
| 299 | glycoprotein | IIVMNIMIIQNYTRS | HLA-DRB5*01:01 | 98.48% (65/66) | Non-Toxin | 0.12 | Consensus (smm/nn/sturniolo) |
| 300 | glycoprotein | IIVMNIMIIQNYTRS | HLA-DRB1*15:01 | 98.48% (65/66) | Non-Toxin | 0.39 | Consensus (smm/nn/sturniolo) |
| 301 | glycoprotein | IKDALQGIQQQIKGL | HLA-DRB4*01:01 | 24.24% (16/66) | Non-Toxin | 0.3 | Consensus (comb.lib./smm/nn) |
| 302 | glycoprotein | ILSAFNTVIALLGSI | HLA-DRB1*07:01 | 98.48% (65/66) | Non-Toxin | 0.44 | Consensus (comb.lib./smm/nn) |
| 303 | glycoprotein | ILSAFNTVIALLGSI | HLA-DPA1*03:01/DPB1*04:02 | 98.48% (65/66) | Non-Toxin | 0.46 | Consensus (comb.lib./smm/nn) |
| 304 | glycoprotein | ITNCFLLKNKIWCIS | HLA-DRB1*11:01 | 98.48% (65/66) | Non-Toxin | 0.43 | Consensus (smm/nn/sturniolo) |
| 305 | glycoprotein | IVIIVMNIMIIQNYT | HLA-DRB4*01:01 | 98.48% (65/66) | Non-Toxin | 0.08 | Consensus (comb.lib./smm/nn) |
| 306 | glycoprotein | IVMNIMIIQNYTRST | HLA-DRB4*01:01 | 98.48% (65/66) | Non-Toxin | 0.1 | Consensus (comb.lib./smm/nn) |
| 307 | glycoprotein | IVMNIMIIQNYTRST | HLA-DRB5*01:01 | 98.48% (65/66) | Non-Toxin | 0.12 | Consensus (smm/nn/sturniolo) |
| 308 | glycoprotein | IVMNIMIIQNYTRST | HLA-DRB1*15:01 | 98.48% (65/66) | Non-Toxin | 0.44 | Consensus (smm/nn/sturniolo) |
| 309 | glycoprotein | KDALQGIQQQIKGLA | HLA-DRB4*01:01 | 24.24% (16/66) | Non-Toxin | 0.34 | Consensus (comb.lib./smm/nn) |
| 310 | glycoprotein | KILSAFNTVIALLGS | HLA-DPA1*03:01/DPB1*04:02 | 98.48% (65/66) | Non-Toxin | 0.46 | Consensus (comb.lib./smm/nn) |
| 311 | glycoprotein | KILSAFNTVIALLGS | HLA-DRB1*07:01 | 98.48% (65/66) | Non-Toxin | 0.47 | Consensus (comb.lib./smm/nn) |
| 312 | glycoprotein | KTITNCFLLKNKIWC | HLA-DRB1*11:01 | 98.48% (65/66) | Non-Toxin | 0.45 | Consensus (smm/nn/sturniolo) |
| 313 | glycoprotein | LGSIVIIVMNIMIIQ | HLA-DRB4*01:01 | 98.48% (65/66) | Non-Toxin | 0.14 | Consensus (comb.lib./smm/nn) |
| 314 | glycoprotein | LGSIVIIVMNIMIIQ | HLA-DRB1*15:01 | 98.48% (65/66) | Non-Toxin | 0.3 | Consensus (smm/nn/sturniolo) |
| 315 | glycoprotein | LLGSIVIIVMNIMII | HLA-DRB4*01:01 | 98.48% (65/66) | Non-Toxin | 0.14 | Consensus (comb.lib./smm/nn) |
| 316 | glycoprotein | LLGSIVIIVMNIMII | HLA-DRB1*15:01 | 98.48% (65/66) | Non-Toxin | 0.3 | Consensus (smm/nn/sturniolo) |
| 317 | glycoprotein | LPNNICLQKTSNQIL | HLA-DRB1*07:01 | 96.97% (64/66) | Non-Toxin | 0.34 | Consensus (comb.lib./smm/nn) |
| 318 | glycoprotein | LSAFNTVIALLGSIV | HLA-DRB1*07:01 | 98.48% (65/66) | Non-Toxin | 0.41 | Consensus (comb.lib./smm/nn) |
| 319 | glycoprotein | LSAFNTVIALLGSIV | HLA-DPA1*03:01/DPB1*04:02 | 98.48% (65/66) | Non-Toxin | 0.48 | Consensus (comb.lib./smm/nn) |
| 320 | glycoprotein | MDEGYFAYSHLERIG | HLA-DPA1*03:01/DPB1*04:02 | 24.24% (16/66) | Non-Toxin | 0.05 | Consensus (comb.lib./smm/nn) |
| 321 | glycoprotein | MDEGYFAYSHLERIG | HLA-DPA1*01:03/DPB1*02:01 | 24.24% (16/66) | Non-Toxin | 0.23 | Consensus (comb.lib./smm/nn) |
| 322 | glycoprotein | MDEGYFAYSHLERIG | HLA-DPA1*02:01/DPB1*01:01 | 24.24% (16/66) | Non-Toxin | 0.27 | Consensus (comb.lib./smm/nn) |
| 323 | glycoprotein | MNIMIIQNYTRSTDN | HLA-DRB5*01:01 | 98.48% (65/66) | Non-Toxin | 0.13 | Consensus (smm/nn/sturniolo) |
| 324 | glycoprotein | NDAFLIDRINWISAG | HLA-DRB3*01:01 | 100.00% (66/66) | Non-Toxin | 0.01 | Consensus (comb.lib./smm/nn) |
| 325 | glycoprotein | NDAFLIDRINWISAG | HLA-DRB1*03:01 | 100.00% (66/66) | Non-Toxin | 0.22 | Consensus (smm/nn/sturniolo) |
| 326 | glycoprotein | PEICWEGVYNDAFLI | HLA-DRB3*01:01 | 25.76% (17/66) | Non-Toxin | 0.19 | Consensus (comb.lib./smm/nn) |
| 327 | glycoprotein | QASFSWDTMIKFGDV | HLA-DRB3*01:01 | 100.00% (66/66) | Non-Toxin | 0.01 | Consensus (comb.lib./smm/nn) |
| 328 | glycoprotein | SAFNTVIALLGSIVI | HLA-DRB1*07:01 | 98.48% (65/66) | Non-Toxin | 0.38 | Consensus (comb.lib./smm/nn) |

| | | | | | | | |
|---|---|---|---|---|---|---|---|
| 329 | glycoprotein | SAVYNNEFYYVLCAV | HLA-DPA1*01/DPB1*04:01 | 89.39% (59/66) | Non-Toxin | 0.32 | Consensus (comb.lib./smm/nn) |
| 330 | glycoprotein | SFSWDTMIKFGDVLT | HLA-DRB3*01:01 | 19.70% (13/66) | Non-Toxin | 0.01 | Consensus (comb.lib./smm/nn) |
| 331 | glycoprotein | SIVIIVMNIMIIQNY | HLA-DRB4*01:01 | 98.48% (65/66) | Non-Toxin | 0.08 | Consensus (comb.lib./smm/nn) |
| 332 | glycoprotein | SIVIIVMNIMIIQNY | HLA-DRB1*15:01 | 98.48% (65/66) | Non-Toxin | 0.48 | Consensus (smm/nn/sturniolo) |
| 333 | glycoprotein | SKILSAFNTVIALLG | HLA-DRB1*07:01 | 98.48% (65/66) | Non-Toxin | 0.32 | Consensus (comb.lib./smm/nn) |
| 334 | glycoprotein | SKILSAFNTVIALLG | HLA-DPA1*03:01/DPB1*04:02 | 98.48% (65/66) | Non-Toxin | 0.48 | Consensus (comb.lib./smm/nn) |
| 335 | glycoprotein | TITNCFLLKNKIWCI | HLA-DRB1*11:01 | 98.48% (65/66) | Non-Toxin | 0.44 | Consensus (smm/nn/sturniolo) |
| 336 | glycoprotein | VFYQASFSWDTMIKF | HLA-DRB3*01:01 | 100.00% (66/66) | Non-Toxin | 0.23 | Consensus (comb.lib./smm/nn) |
| 337 | glycoprotein | VIIVMNIMIIQNYTR | HLA-DRB4*01:01 | 98.48% (65/66) | Non-Toxin | 0.1 | Consensus (comb.lib./smm/nn) |
| 338 | glycoprotein | VIIVMNIMIIQNYTR | HLA-DRB5*01:01 | 98.48% (65/66) | Non-Toxin | 0.12 | Consensus (smm/nn/sturniolo) |
| 339 | glycoprotein | VIIVMNIMIIQNYTR | HLA-DRB1*15:01 | 98.48% (65/66) | Non-Toxin | 0.31 | Consensus (smm/nn/sturniolo) |
| 340 | glycoprotein | VIKDALQGIQQQIKG | HLA-DRB4*01:01 | 24.24% (16/66) | Non-Toxin | 0.35 | Consensus (comb.lib./smm/nn) |
| 341 | glycoprotein | VMNIMIIQNYTRSTD | HLA-DRB5*01:01 | 98.48% (65/66) | Non-Toxin | 0.12 | Consensus (smm/nn/sturniolo) |
| 342 | glycoprotein | VYNDAFLIDRINWIS | HLA-DRB3*01:01 | 100.00% (66/66) | Non-Toxin | 0.01 | Consensus (comb.lib./smm/nn) |
| 343 | glycoprotein | VYNDAFLIDRINWIS | HLA-DRB1*03:01 | 100.00% (66/66) | Non-Toxin | 0.22 | Consensus (smm/nn/sturniolo) |
| 344 | glycoprotein | WEGVYNDAFLIDRIN | HLA-DRB3*01:01 | 100.00% (66/66) | Non-Toxin | 0.17 | Consensus (comb.lib./smm/nn) |
| 345 | glycoprotein | WEGVYNDAFLIDRIN | HLA-DPA1*01:03/DPB1*02:01 | 100.00% (66/66) | Non-Toxin | 0.34 | Consensus (comb.lib./smm/nn) |
| 346 | glycoprotein | YFAYSHLERIGSCSR | HLA-DPA1*03:01/DPB1*04:02 | 24.24% (16/66) | Non-Toxin | 0.1 | Consensus (comb.lib./smm/nn) |
| 347 | glycoprotein | YFAYSHLERIGSCSR | HLA-DRB1*11:01 | 24.24% (16/66) | Non-Toxin | 0.38 | Consensus (smm/nn/sturniolo) |
| 348 | glycoprotein | YHCSAVYNNEFYYVL | HLA-DPA1*01/DPB1*04:01 | 90.91% (60/66) | Non-Toxin | 0.36 | Consensus (comb.lib./smm/nn) |
| 349 | glycoprotein | YNDAFLIDRINWISA | HLA-DRB3*01:01 | 100.00% (66/66) | Non-Toxin | 0.01 | Consensus (comb.lib./smm/nn) |
| 350 | glycoprotein | YNDAFLIDRINWISA | HLA-DRB1*03:01 | 100.00% (66/66) | Non-Toxin | 0.22 | Consensus (smm/nn/sturniolo) |
| 351 | glycoprotein | YQASFSWDTMIKFGD | HLA-DRB3*01:01 | 100.00% (66/66) | Non-Toxin | 0.01 | Consensus (comb.lib./smm/nn) |
| 352 | glycoprotein | YSHLERIGSCSRGVS | HLA-DRB1*11:01 | 24.24% (16/66) | Non-Toxin | 0.38 | Consensus (smm/nn/sturniolo) |
| **S.No.** | **Protein** | **Peptide** | **Allele** | **Conservancy** | **Toxicity** | **Percentile rank** | **Method used** |
| 353 | Matrix Protein | ALRIFFLSITKLNDS | HLA-DPA1*01/DPB1*04:01 | 97.44% (38/39) | Non-Toxin | 0.21 | Consensus (comb.lib./smm/nn) |
| 354 | Matrix Protein | ALRIFFLSITKLNDS | HLA-DPA1*02:01/DPB1*14:01 | 97.44% (38/39) | Non-Toxin | 0.3 | NetMHCIIpan |
| 355 | Matrix Protein | CFSLMDINPWLNRLT | HLA-DRB3*01:01 | 94.87% (37/39) | Non-Toxin | 0.07 | Consensus (comb.lib./smm/nn) |
| 356 | Matrix Protein | DKHQALRIFFLSITK | HLA-DPA1*01/DPB1*04:01 | 97.44% (38/39) | Non-Toxin | 0.05 | Consensus (comb.lib./smm/nn) |
| 357 | Matrix Protein | DKHQALRIFFLSITK | HLA-DPA1*01:03/DPB1*02:01 | 97.44% (38/39) | Non-Toxin | 0.08 | Consensus (comb.lib./smm/nn) |
| 358 | Matrix Protein | DSGIYMIPRTMLEFR | HLA-DRB1*11:01 | 97.44% (38/39) | Non-Toxin | 0.44 | Consensus (smm/nn/sturniolo) |
| 359 | Matrix Protein | EFMIYDDVFIDNTGR | HLA-DRB3*01:01 | 94.87% (37/39) | Non-Toxin | 0.02 | Consensus (comb.lib./smm/nn) |
| 360 | Matrix Protein | EFRRNNAIAFNLLVY | HLA-DRB3*02:02 | 84.62% (33/39) | Non-Toxin | 0.35 | NetMHCIIpan |
| 361 | Matrix Protein | FMIYDDVFIDNTGRI | HLA-DRB3*01:01 | 94.87% (37/39) | Non-Toxin | 0.01 | Consensus (comb.lib./smm/nn) |
| 362 | Matrix Protein | FMIYDDVFIDNTGRI | HLA-DRB1*03:01 | 94.87% (37/39) | Non-Toxin | 0.08 | Consensus (smm/nn/sturniolo) |
| 363 | Matrix Protein | GKRKKIRTIAAYPLG | HLA-DRB1*08:02 | 97.44% (38/39) | Non-Toxin | 0.32 | Consensus (smm/nn/sturniolo) |
| 364 | Matrix Protein | GNFVRRAGKYYSVDY | HLA-DRB1*11:01 | 79.49% (31/39) | Non-Toxin | 0.38 | Consensus (smm/nn/sturniolo) |
| 365 | Matrix Protein | HIKINGVISKRLFAQ | HLA-DRB1*11:01 | 94.87% (37/39) | Non-Toxin | 0.48 | Consensus (smm/nn/sturniolo) |
| 366 | Matrix Protein | HLGNFVRRAGKYYSV | HLA-DRB1*11:01 | 79.49% (31/39) | Non-Toxin | 0.38 | Consensus (smm/nn/sturniolo) |
| 367 | Matrix Protein | HLVPWKKVLTSGSIF | HLA-DRB1*08:02 | 35.90% (14/39) | Non-Toxin | 0.36 | Consensus (smm/nn/sturniolo) |
| 368 | Matrix Protein | HQALRIFFLSITKLN | HLA-DPA1*01/DPB1*04:01 | 97.44% (38/39) | Non-Toxin | 0.05 | Consensus (comb.lib./smm/nn) |
| 369 | Matrix Protein | HQALRIFFLSITKLN | HLA-DPA1*01:03/DPB1*02:01 | 97.44% (38/39) | Non-Toxin | 0.07 | Consensus (comb.lib./smm/nn) |
| 370 | Matrix Protein | HQALRIFFLSITKLN | HLA-DPA1*02:01/DPB1*14:01 | 97.44% (38/39) | Non-Toxin | 0.5 | NetMHCIIpan |
| 371 | Matrix Protein | IKINGVISKRLFAQM | HLA-DRB1*11:01 | 94.87% (37/39) | Non-Toxin | 0.48 | Consensus (smm/nn/sturniolo) |
| 372 | Matrix Protein | IPREFMIYDDVFIDN | HLA-DRB3*01:01 | 38.46% (15/39) | Non-Toxin | 0.01 | Consensus (comb.lib./smm/nn) |
| 373 | Matrix Protein | IPREFMIYDDVFIDN | HLA-DQA1*05:01/DQB1*02:01 | 38.46% (15/39) | Non-Toxin | 0.34 | Consensus (comb.lib./smm/nn) |
| 374 | Matrix Protein | IPREFMIYDDVFIDN | HLA-DQA1*01:01/DQB1*05:01 | 38.46% (15/39) | Non-Toxin | 0.36 | Consensus (comb.lib./smm/nn) |
| 375 | Matrix Protein | IYDDVFIDNTGRILK | HLA-DRB3*01:01 | 94.87% (37/39) | Non-Toxin | 0.01 | Consensus (comb.lib./smm/nn) |

| | | | | | | | |
|---|---|---|---|---|---|---|---|
| 376 | Matrix Protein | IYDDVFIDNTGRILK | HLA-DRB1*03:01 | 94.87% (37/39) | Non-Toxin | 0.05 | Consensus (smm/nn/sturniolo) |
| 377 | Matrix Protein | KHQALRIFFLSITKL | HLA-DPA1*01/DPB1*04:01 | 97.44% (38/39) | Non-Toxin | 0.04 | Consensus (comb.lib./smm/nn) |
| 378 | Matrix Protein | KHQALRIFFLSITKL | HLA-DPA1*01:03/DPB1*02:01 | 97.44% (38/39) | Non-Toxin | 0.06 | Consensus (comb.lib./smm/nn) |
| 379 | Matrix Protein | KKIRTIAAYPLGVGK | HLA-DRB1*08:02 | 97.44% (38/39) | Non-Toxin | 0.42 | Consensus (smm/nn/sturniolo) |
| 380 | Matrix Protein | KNLCFSLMDINPWLN | HLA-DRB3*01:01 | 94.87% (37/39) | Non-Toxin | 0.06 | Consensus (comb.lib./smm/nn) |
| 381 | Matrix Protein | KRKKIRTIAAYPLGV | HLA-DRB1*08:02 | 97.44% (38/39) | Non-Toxin | 0.36 | Consensus (smm/nn/sturniolo) |
| 382 | Matrix Protein | KRKKIRTIAAYPLGV | HLA-DRB5*01:01 | 97.44% (38/39) | Non-Toxin | 0.5 | Consensus (smm/nn/sturniolo) |
| 383 | Matrix Protein | LCFSLMDINPWLNRL | HLA-DRB3*01:01 | 94.87% (37/39) | Non-Toxin | 0.06 | Consensus (comb.lib./smm/nn) |
| 384 | Matrix Protein | LDKHQALRIFFLSIT | HLA-DPA1*01/DPB1*04:01 | 97.44% (38/39) | Non-Toxin | 0.06 | Consensus (comb.lib./smm/nn) |
| 385 | Matrix Protein | LDKHQALRIFFLSIT | HLA-DPA1*01:03/DPB1*02:01 | 97.44% (38/39) | Non-Toxin | 0.08 | Consensus (comb.lib./smm/nn) |
| 386 | Matrix Protein | LEFRRNNAIAFNLLV | HLA-DRB3*02:02 | 84.62% (33/39) | Non-Toxin | 0.08 | NetMHCIIpan |
| 387 | Matrix Protein | LEFRRNNAIAFNLLV | HLA-DRB3*01:01 | 84.62% (33/39) | Non-Toxin | 0.46 | Consensus (comb.lib./smm/nn) |
| 388 | Matrix Protein | LEFRRNNAIAFNLLV | HLA-DRB1*13:02 | 84.62% (33/39) | Non-Toxin | 0.48 | Consensus (smm/nn/sturniolo) |
| 389 | Matrix Protein | LGNFVRRAGKYYSVD | HLA-DRB1*11:01 | 79.49% (31/39) | Non-Toxin | 0.38 | Consensus (smm/nn/sturniolo) |
| 390 | Matrix Protein | LHIKINGVISKRLFA | HLA-DRB1*07:01 | 94.87% (37/39) | Non-Toxin | 0.4 | Consensus (comb.lib./smm/nn) |
| 391 | Matrix Protein | LHIKINGVISKRLFA | HLA-DRB1*11:01 | 94.87% (37/39) | Non-Toxin | 0.48 | Consensus (smm/nn/sturniolo) |
| 392 | Matrix Protein | LHIKINGVISKRLFA | HLA-DRB3*02:02 | 94.87% (37/39) | Non-Toxin | 0.5 | NetMHCIIpan |
| 393 | Matrix Protein | LHLGNFVRRAGKYYS | HLA-DRB1*11:01 | 79.49% (31/39) | Non-Toxin | 0.38 | Consensus (smm/nn/sturniolo) |
| 394 | Matrix Protein | LNHLVPWKKVLTSGS | HLA-DRB1*08:02 | 35.90% (14/39) | Non-Toxin | 0.45 | Consensus (smm/nn/sturniolo) |
| 395 | Matrix Protein | LRIFFLSITKLNDSG | HLA-DPA1*02:01/DPB1*14:01 | 97.44% (38/39) | Non-Toxin | 0.48 | NetMHCIIpan |
| 396 | Matrix Protein | LSLHIKINGVISKRL | HLA-DRB1*07:01 | 94.87% (37/39) | Non-Toxin | 0.4 | Consensus (comb.lib./smm/nn) |
| 397 | Matrix Protein | MIYDDVFIDNTGRIL | HLA-DRB3*01:01 | 94.87% (37/39) | Non-Toxin | 0.01 | Consensus (comb.lib./smm/nn) |
| 398 | Matrix Protein | MIYDDVFIDNTGRIL | HLA-DRB1*03:01 | 94.87% (37/39) | Non-Toxin | 0.06 | Consensus (smm/nn/sturniolo) |
| 399 | Matrix Protein | MLEFRRNNAIAFNLL | HLA-DRB3*02:02 | 97.44% (38/39) | Non-Toxin | 0.06 | NetMHCIIpan |
| 400 | Matrix Protein | MLEFRRNNAIAFNLL | HLA-DRB3*01:01 | 97.44% (38/39) | Non-Toxin | 0.22 | Consensus (comb.lib./smm/nn) |
| 401 | Matrix Protein | MLEFRRNNAIAFNLL | HLA-DRB1*13:02 | 97.44% (38/39) | Non-Toxin | 0.48 | Consensus (smm/nn/sturniolo) |
| 402 | Matrix Protein | MYLICYGFVEDVERT | HLA-DQA1*01:01/DQB1*05:01 | 97.44% (38/39) | Non-Toxin | 0.28 | Consensus (comb.lib./smm/nn) |
| 403 | Matrix Protein | NFVRRAGKYYSVDYC | HLA-DRB1*11:01 | 79.49% (31/39) | Non-Toxin | 0.38 | Consensus (smm/nn/sturniolo) |
| 404 | Matrix Protein | NHLVPWKKVLTSGSI | HLA-DRB1*08:02 | 35.90% (14/39) | Non-Toxin | 0.29 | Consensus (smm/nn/sturniolo) |
| 405 | Matrix Protein | NLCFSLMDINPWLNR | HLA-DRB3*01:01 | 94.87% (37/39) | Non-Toxin | 0.06 | Consensus (comb.lib./smm/nn) |
| 406 | Matrix Protein | NYMYLICYGFVEDVE | HLA-DQA1*01:01/DQB1*05:01 | 97.44% (38/39) | Non-Toxin | 0.39 | Consensus (comb.lib./smm/nn) |
| 407 | Matrix Protein | PREFMIYDDVFIDNT | HLA-DRB3*01:01 | 94.87% (37/39) | Non-Toxin | 0.01 | Consensus (comb.lib./smm/nn) |
| 408 | Matrix Protein | PREFMIYDDVFIDNT | HLA-DQA1*01:01/DQB1*05:01 | 94.87% (37/39) | Non-Toxin | 0.35 | Consensus (comb.lib./smm/nn) |
| 409 | Matrix Protein | PREFMIYDDVFIDNT | HLA-DQA1*05:01/DQB1*02:01 | 94.87% (37/39) | Non-Toxin | 0.39 | Consensus (comb.lib./smm/nn) |
| 410 | Matrix Protein | PRTMLEFRRNNAIAF | HLA-DRB3*01:01 | 97.44% (38/39) | Non-Toxin | 0.3 | Consensus (comb.lib./smm/nn) |
| 411 | Matrix Protein | PRTMLEFRRNNAIAF | HLA-DRB1*13:02 | 97.44% (38/39) | Non-Toxin | 0.51 | Consensus (smm/nn/sturniolo) |
| 412 | Matrix Protein | PSIPREFMIYDDVFI | HLA-DRB3*01:01 | 38.46% (15/39) | Non-Toxin | 0.11 | Consensus (comb.lib./smm/nn) |
| 413 | Matrix Protein | PSIPREFMIYDDVFI | HLA-DQA1*05:01/DQB1*02:01 | 38.46% (15/39) | Non-Toxin | 0.41 | Consensus (comb.lib./smm/nn) |
| 414 | Matrix Protein | QALRIFFLSITKLND | HLA-DPA1*01/DPB1*04:01 | 97.44% (38/39) | Non-Toxin | 0.19 | Consensus (comb.lib./smm/nn) |
| 415 | Matrix Protein | QALRIFFLSITKLND | HLA-DPA1*02:01/DPB1*14:01 | 97.44% (38/39) | Non-Toxin | 0.33 | NetMHCIIpan |
| 416 | Matrix Protein | QKNLCFSLMDINPWL | HLA-DRB3*01:01 | 94.87% (37/39) | Non-Toxin | 0.06 | Consensus (comb.lib./smm/nn) |
| 417 | Matrix Protein | QLDKHQALRIFFLSI | HLA-DPA1*01/DPB1*04:01 | 97.44% (38/39) | Non-Toxin | 0.06 | Consensus (comb.lib./smm/nn) |
| 418 | Matrix Protein | QLDKHQALRIFFLSI | HLA-DPA1*01:03/DPB1*02:01 | 97.44% (38/39) | Non-Toxin | 0.09 | Consensus (comb.lib./smm/nn) |
| 419 | Matrix Protein | REFMIYDDVFIDNTG | HLA-DRB3*01:01 | 94.87% (37/39) | Non-Toxin | 0.01 | Consensus (comb.lib./smm/nn) |
| 420 | Matrix Protein | REFMIYDDVFIDNTG | HLA-DQA1*01:01/DQB1*05:01 | 94.87% (37/39) | Non-Toxin | 0.35 | Consensus (comb.lib./smm/nn) |
| 421 | Matrix Protein | RKKIRTIAAYPLGVG | HLA-DRB1*08:02 | 97.44% (38/39) | Non-Toxin | 0.36 | Consensus (smm/nn/sturniolo) |
| 422 | Matrix Protein | RTMLEFRRNNAIAFN | HLA-DRB3*02:02 | 97.44% (38/39) | Non-Toxin | 0.1 | NetMHCIIpan |
| 423 | Matrix Protein | RTMLEFRRNNAIAFN | HLA-DRB3*01:01 | 97.44% (38/39) | Non-Toxin | 0.27 | Consensus (comb.lib./smm/nn) |

| | | | | | | | |
|---|---|---|---|---|---|---|---|
| 424 | Matrix Protein | SIPREFMIYDDVFID | HLA-DRB3*01:01 | 38.46% (15/39) | Non-Toxin | 0.01 | Consensus (comb.lib./smm/nn) |
| 425 | Matrix Protein | SIPREFMIYDDVFID | HLA-DQA1*05:01/DQB1*02:01 | 38.46% (15/39) | Non-Toxin | 0.34 | Consensus (comb.lib./smm/nn) |
| 426 | Matrix Protein | SIPREFMIYDDVFID | HLA-DQA1*01:01/DQB1*05:01 | 38.46% (15/39) | Non-Toxin | 0.43 | Consensus (comb.lib./smm/nn) |
| 427 | Matrix Protein | SLHIKINGVISKRLF | HLA-DRB1*07:01 | 94.87% (37/39) | Non-Toxin | 0.34 | Consensus (comb.lib./smm/nn) |
| 428 | Matrix Protein | TMLEFRRNNAIAFNL | HLA-DRB3*02:02 | 97.44% (38/39) | Non-Toxin | 0.06 | NetMHCIIpan |
| 429 | Matrix Protein | TMLEFRRNNAIAFNL | HLA-DRB3*01:01 | 97.44% (38/39) | Non-Toxin | 0.23 | Consensus (comb.lib./smm/nn) |
| 430 | Matrix Protein | TMLEFRRNNAIAFNL | HLA-DRB1*13:02 | 97.44% (38/39) | Non-Toxin | 0.48 | Consensus (smm/nn/sturniolo) |
| 431 | Matrix Protein | YDDVFIDNTGRILKG | HLA-DRB3*01:01 | 94.87% (37/39) | Non-Toxin | 0.01 | Consensus (comb.lib./smm/nn) |
| 432 | Matrix Protein | YDDVFIDNTGRILKG | HLA-DRB1*03:01 | 94.87% (37/39) | Non-Toxin | 0.06 | Consensus (smm/nn/sturniolo) |
| 433 | Matrix Protein | YLICYGFVEDVERTP | HLA-DQA1*01:01/DQB1*05:01 | 97.44% (38/39) | Non-Toxin | 0.27 | Consensus (comb.lib./smm/nn) |
| 434 | Matrix Protein | YMYLICYGFVEDVER | HLA-DQA1*01:01/DQB1*05:01 | 97.44% (38/39) | Non-Toxin | 0.35 | Consensus (comb.lib./smm/nn) |
| | **Protein** | **Peptide** | **Allele** | **Conservancy** | **Toxicity** | **Percentile rank** | **Method used** |
| 435 | nucleocapsid | AELAAAVQETSAGRQ | HLA-DQA1*04:01/DQB1*04:02 | 100.00% (13/13) | Non-Toxin | 0.27 | Consensus (comb.lib./smm/nn) |
| 436 | nucleocapsid | AGGIDQNMANRLGLS | HLA-DRB1*13:02 | 100.00% (13/13) | Non-Toxin | 0.15 | Consensus (smm/nn/sturniolo) |
| 437 | nucleocapsid | ALIRSLLNDPDIEAV | HLA-DRB3*01:01 | 100.00% (13/13) | Non-Toxin | 0.07 | Consensus (comb.lib./smm/nn) |
| 438 | nucleocapsid | ALIRSLLNDPDIEAV | HLA-DRB1*04:05 | 100.00% (13/13) | Non-Toxin | 0.19 | Consensus (smm/nn/sturniolo) |
| 439 | nucleocapsid | AVEIISDIGNYVEET | HLA-DRB3*01:01 | 100.00% (13/13) | Non-Toxin | 0.26 | Consensus (comb.lib./smm/nn) |
| 440 | nucleocapsid | AVEIISDIGNYVEET | HLA-DRB1*13:02 | 100.00% (13/13) | Non-Toxin | 0.27 | Consensus (smm/nn/sturniolo) |
| 441 | nucleocapsid | AVIIDVGSMVNGIPV | HLA-DRB1*03:01 | 100.00% (13/13) | Non-Toxin | 0.45 | Consensus (smm/nn/sturniolo) |
| 442 | nucleocapsid | DIEAVIIDVGSMVNG | HLA-DRB1*03:01 | 100.00% (13/13) | Non-Toxin | 0.45 | Consensus (smm/nn/sturniolo) |
| 443 | nucleocapsid | DMSTLVSAVITIEAQ | HLA-DQA1*04:01/DQB1*04:02 | 100.00% (13/13) | Non-Toxin | 0.03 | Consensus (comb.lib./smm/nn) |
| 444 | nucleocapsid | DPDIEAVIIDVGSMV | HLA-DRB1*03:01 | 100.00% (13/13) | Non-Toxin | 0.45 | Consensus (smm/nn/sturniolo) |
| 445 | nucleocapsid | DQVAELAAAVQETSA | HLA-DQA1*04:01/DQB1*04:02 | 15.38% (2/13) | Non-Toxin | 0.01 | Consensus (comb.lib./smm/nn) |
| 446 | nucleocapsid | DQVAELAAAVQETSA | HLA-DQA1*03:01/DQB1*03:02 | 15.38% (2/13) | Non-Toxin | 0.03 | Consensus (comb.lib./smm/nn) |
| 447 | nucleocapsid | EAVIIDVGSMVNGIP | HLA-DRB1*03:01 | 100.00% (13/13) | Non-Toxin | 0.45 | Consensus (smm/nn/sturniolo) |
| 448 | nucleocapsid | EFQSDLNTIKSLMLL | HLA-DRB1*07:01 | 100.00% (13/13) | Non-Toxin | 0.11 | Consensus (comb.lib./smm/nn) |
| 449 | nucleocapsid | ELRWELTLFALDVIR | HLA-DQA1*01:01/DQB1*05:01 | 100.00% (13/13) | Non-Toxin | 0.02 | Consensus (comb.lib./smm/nn) |
| 450 | nucleocapsid | ELRWELTLFALDVIR | HLA-DPA1*01/DPB1*04:01 | 100.00% (13/13) | Non-Toxin | 0.18 | Consensus (comb.lib./smm/nn) |
| 451 | nucleocapsid | ELTLFALDVIRSPSA | HLA-DQA1*01:01/DQB1*05:01 | 100.00% (13/13) | Non-Toxin | 0.2 | Consensus (comb.lib./smm/nn) |
| 452 | nucleocapsid | ELTLFALDVIRSPSA | HLA-DRB1*11:01 | 100.00% (13/13) | Non-Toxin | 0.44 | Consensus (smm/nn/sturniolo) |
| 453 | nucleocapsid | ERPGALIRSLLNDPD | HLA-DRB1*04:05 | 100.00% (13/13) | Non-Toxin | 0.22 | Consensus (smm/nn/sturniolo) |
| 454 | nucleocapsid | FQSDLNTIKSLMLLY | HLA-DRB1*07:01 | 100.00% (13/13) | Non-Toxin | 0.24 | Consensus (comb.lib./smm/nn) |
| 455 | nucleocapsid | GALIRSLLNDPDIEA | HLA-DRB3*01:01 | 100.00% (13/13) | Non-Toxin | 0.07 | Consensus (comb.lib./smm/nn) |
| 456 | nucleocapsid | GALIRSLLNDPDIEA | HLA-DRB1*04:05 | 100.00% (13/13) | Non-Toxin | 0.19 | Consensus (smm/nn/sturniolo) |
| 457 | nucleocapsid | GGIDQNMANRLGLSS | HLA-DRB1*13:02 | 100.00% (13/13) | Non-Toxin | 0.15 | Consensus (smm/nn/sturniolo) |
| 458 | nucleocapsid | GGYPLLWSFAMGVAT | HLA-DRB1*09:01 | 100.00% (13/13) | Non-Toxin | 0.44 | Consensus (comb.lib./smm/nn) |
| 459 | nucleocapsid | GLRITDMSTLVSAVI | HLA-DQA1*04:01/DQB1*04:02 | 100.00% (13/13) | Non-Toxin | 0.45 | Consensus (comb.lib./smm/nn) |
| 460 | nucleocapsid | GLSSDQVAELAAAVQ | HLA-DQA1*04:01/DQB1*04:02 | 15.38% (2/13) | Non-Toxin | 0.17 | Consensus (comb.lib./smm/nn) |
| 461 | nucleocapsid | GMAGFFATIRFGLET | HLA-DPA1*01/DPB1*04:01 | 100.00% (13/13) | Non-Toxin | 0.43 | Consensus (comb.lib./smm/nn) |
| 462 | nucleocapsid | GRAVEIISDIGNYVE | HLA-DRB3*01:01 | 100.00% (13/13) | Non-Toxin | 0.19 | Consensus (comb.lib./smm/nn) |
| 463 | nucleocapsid | GYPLLWSFAMGVATT | HLA-DRB1*09:01 | 100.00% (13/13) | Non-Toxin | 0.46 | Consensus (comb.lib./smm/nn) |
| 464 | nucleocapsid | HAGGIDQNMANRLGL | HLA-DRB1*13:02 | 100.00% (13/13) | Non-Toxin | 0.15 | Consensus (smm/nn/sturniolo) |
| 465 | nucleocapsid | HHAGGIDQNMANRLG | HLA-DRB1*13:02 | 100.00% (13/13) | Non-Toxin | 0.24 | Consensus (smm/nn/sturniolo) |
| 466 | nucleocapsid | IEAVIIDVGSMVNGI | HLA-DRB1*03:01 | 100.00% (13/13) | Non-Toxin | 0.45 | Consensus (smm/nn/sturniolo) |
| 467 | nucleocapsid | IRIFVPATNSPELRW | HLA-DRB1*04:05 | 100.00% (13/13) | Non-Toxin | 0.11 | Consensus (smm/nn/sturniolo) |
| 468 | nucleocapsid | IRSLLNDPDIEAVII | HLA-DRB3*01:01 | 100.00% (13/13) | Non-Toxin | 0.05 | Consensus (comb.lib./smm/nn) |
| 469 | nucleocapsid | ITDMSTLVSAVITIE | HLA-DQA1*04:01/DQB1*04:02 | 100.00% (13/13) | Non-Toxin | 0.05 | Consensus (comb.lib./smm/nn) |
| 470 | nucleocapsid | KGRAVEIISDIGNYV | HLA-DRB3*01:01 | 100.00% (13/13) | Non-Toxin | 0.19 | Consensus (comb.lib./smm/nn) |

| | | | | | | | |
|---|---|---|---|---|---|---|---|
| 471 | nucleocapsid | KIRIFVPATNSPELR | HLA-DRB1*04:05 | 100.00% (13/13) | Non-Toxin | 0.12 | Consensus (smm/nn/sturniolo) |
| 472 | nucleocapsid | LGLSSDQVAELAAAV | HLA-DQA1*04:01/DQB1*04:02 | 15.38% (2/13) | Non-Toxin | 0.16 | Consensus (comb.lib./smm/nn) |
| 473 | nucleocapsid | LIRSLLNDPDIEAVI | HLA-DRB3*01:01 | 100.00% (13/13) | Non-Toxin | 0.06 | Consensus (comb.lib./smm/nn) |
| 474 | nucleocapsid | LLWSFAMGVATTIDR | HLA-DRB1*09:01 | 100.00% (13/13) | Non-Toxin | 0.01 | Consensus (comb.lib./smm/nn) |
| 475 | nucleocapsid | LRITDMSTLVSAVIT | HLA-DQA1*04:01/DQB1*04:02 | 100.00% (13/13) | Non-Toxin | 0.43 | Consensus (comb.lib./smm/nn) |
| 476 | nucleocapsid | LRWELTLFALDVIRS | HLA-DQA1*01:01/DQB1*05:01 | 100.00% (13/13) | Non-Toxin | 0.02 | Consensus (comb.lib./smm/nn) |
| 477 | nucleocapsid | LRWELTLFALDVIRS | HLA-DPA1*01/DPB1*04:01 | 100.00% (13/13) | Non-Toxin | 0.21 | Consensus (comb.lib./smm/nn) |
| 478 | nucleocapsid | LSSDQVAELAAAVQE | HLA-DQA1*04:01/DQB1*04:02 | 15.38% (2/13) | Non-Toxin | 0.01 | Consensus (comb.lib./smm/nn) |
| 479 | nucleocapsid | LSSDQVAELAAAVQE | HLA-DQA1*03:01/DQB1*03:02 | 15.38% (2/13) | Non-Toxin | 0.02 | Consensus (comb.lib./smm/nn) |
| 480 | nucleocapsid | LTEMRNLLSQSLSVR | HLA-DRB4*01:01 | 100.00% (13/13) | Non-Toxin | 0.39 | Consensus (comb.lib./smm/nn) |
| 481 | nucleocapsid | LTLFALDVIRSPSAA | HLA-DQA1*01:01/DQB1*05:01 | 100.00% (13/13) | Non-Toxin | 0.22 | Consensus (comb.lib./smm/nn) |
| 482 | nucleocapsid | LTLFALDVIRSPSAA | HLA-DRB1*11:01 | 100.00% (13/13) | Non-Toxin | 0.42 | Consensus (smm/nn/sturniolo) |
| 483 | nucleocapsid | LTNSLLNLRSRLAAK | HLA-DRB1*15:01 | 100.00% (13/13) | Non-Toxin | 0.49 | Consensus (smm/nn/sturniolo) |
| 484 | nucleocapsid | LTTKIRIFVPATNSP | HLA-DRB1*08:02 | 100.00% (13/13) | Non-Toxin | 0.37 | Consensus (smm/nn/sturniolo) |
| 485 | nucleocapsid | LVSAVITIEAQIWIL | HLA-DQA1*03:01/DQB1*03:02 | 100.00% (13/13) | Non-Toxin | 0.18 | Consensus (comb.lib./smm/nn) |
| 486 | nucleocapsid | LWSFAMGVATTIDRS | HLA-DRB1*09:01 | 100.00% (13/13) | Non-Toxin | 0.01 | Consensus (comb.lib./smm/nn) |
| 487 | nucleocapsid | MAGFFATIRFGLETR | HLA-DPA1*01/DPB1*04:01 | 100.00% (13/13) | Non-Toxin | 0.43 | Consensus (comb.lib./smm/nn) |
| 488 | nucleocapsid | MSTLVSAVITIEAQI | HLA-DQA1*03:01/DQB1*03:02 | 100.00% (13/13) | Non-Toxin | 0.21 | Consensus (comb.lib./smm/nn) |
| 489 | nucleocapsid | NEFQSDLNTIKSLML | HLA-DRB1*07:01 | 100.00% (13/13) | Non-Toxin | 0.1 | Consensus (comb.lib./smm/nn) |
| 490 | nucleocapsid | NEFQSDLNTIKSLML | HLA-DRB1*04:01 | 100.00% (13/13) | Non-Toxin | 0.38 | Consensus (smm/nn/sturniolo) |
| 491 | nucleocapsid | NSLLNLRSRLAAKAA | HLA-DRB1*15:01 | 100.00% (13/13) | Non-Toxin | 0.51 | Consensus (smm/nn/sturniolo) |
| 492 | nucleocapsid | PDIEAVIIDVGSMVN | HLA-DRB1*03:01 | 100.00% (13/13) | Non-Toxin | 0.45 | Consensus (smm/nn/sturniolo) |
| 493 | nucleocapsid | PELRWELTLFALDVI | HLA-DQA1*01:01/DQB1*05:01 | 100.00% (13/13) | Non-Toxin | 0.02 | Consensus (comb.lib./smm/nn) |
| 494 | nucleocapsid | PELRWELTLFALDVI | HLA-DPA1*01/DPB1*04:01 | 100.00% (13/13) | Non-Toxin | 0.16 | Consensus (comb.lib./smm/nn) |
| 495 | nucleocapsid | PGALIRSLLNDPDIE | HLA-DRB1*04:05 | 100.00% (13/13) | Non-Toxin | 0.19 | Consensus (smm/nn/sturniolo) |
| 496 | nucleocapsid | PLLWSFAMGVATTID | HLA-DRB1*09:01 | 100.00% (13/13) | Non-Toxin | 0.01 | Consensus (comb.lib./smm/nn) |
| 497 | nucleocapsid | QVAELAAAVQETSAG | HLA-DQA1*04:01/DQB1*04:02 | 100.00% (13/13) | Non-Toxin | 0.01 | Consensus (comb.lib./smm/nn) |
| 498 | nucleocapsid | QVAELAAAVQETSAG | HLA-DQA1*03:01/DQB1*03:02 | 100.00% (13/13) | Non-Toxin | 0.07 | Consensus (comb.lib./smm/nn) |
| 499 | nucleocapsid | RAVEIISDIGNYVEE | HLA-DRB3*01:01 | 100.00% (13/13) | Non-Toxin | 0.22 | Consensus (comb.lib./smm/nn) |
| 500 | nucleocapsid | RAVEIISDIGNYVEE | HLA-DRB1*13:02 | 100.00% (13/13) | Non-Toxin | 0.33 | Consensus (smm/nn/sturniolo) |
| 501 | nucleocapsid | RHHAGGIDQNMANRL | HLA-DRB1*13:02 | 100.00% (13/13) | Non-Toxin | 0.33 | Consensus (smm/nn/sturniolo) |
| 502 | nucleocapsid | RIFVPATNSPELRWE | HLA-DRB1*04:05 | 100.00% (13/13) | Non-Toxin | 0.27 | Consensus (smm/nn/sturniolo) |
| 503 | nucleocapsid | RITDMSTLVSAVITI | HLA-DQA1*04:01/DQB1*04:02 | 100.00% (13/13) | Non-Toxin | 0.4 | Consensus (comb.lib./smm/nn) |
| 504 | nucleocapsid | RLTNSLLNLRSRLAA | HLA-DRB1*15:01 | 100.00% (13/13) | Non-Toxin | 0.49 | Consensus (smm/nn/sturniolo) |
| 505 | nucleocapsid | RPGALIRSLLNDPDI | HLA-DRB1*04:05 | 100.00% (13/13) | Non-Toxin | 0.19 | Consensus (smm/nn/sturniolo) |
| 506 | nucleocapsid | RWELTLFALDVIRSP | HLA-DQA1*01:01/DQB1*05:01 | 100.00% (13/13) | Non-Toxin | 0.02 | Consensus (comb.lib./smm/nn) |
| 507 | nucleocapsid | SDQVAELAAAVQETS | HLA-DQA1*04:01/DQB1*04:02 | 15.38% (2/13) | Non-Toxin | 0.01 | Consensus (comb.lib./smm/nn) |
| 508 | nucleocapsid | SDQVAELAAAVQETS | HLA-DQA1*03:01/DQB1*03:02 | 15.38% (2/13) | Non-Toxin | 0.03 | Consensus (comb.lib./smm/nn) |
| 509 | nucleocapsid | SKGKTPFVDSRAYGL | HLA-DRB1*07:01 | 100.00% (13/13) | Non-Toxin | 0.38 | Consensus (comb.lib./smm/nn) |
| 510 | nucleocapsid | SSDQVAELAAAVQET | HLA-DQA1*04:01/DQB1*04:02 | 15.38% (2/13) | Non-Toxin | 0.01 | Consensus (comb.lib./smm/nn) |
| 511 | nucleocapsid | SSDQVAELAAAVQET | HLA-DQA1*03:01/DQB1*03:02 | 15.38% (2/13) | Non-Toxin | 0.02 | Consensus (comb.lib./smm/nn) |
| 512 | nucleocapsid | STLVSAVITIEAQIW | HLA-DQA1*03:01/DQB1*03:02 | 100.00% (13/13) | Non-Toxin | 0.18 | Consensus (comb.lib./smm/nn) |
| 513 | nucleocapsid | SVTFKREMSISSLAN | HLA-DRB1*04:01 | 0.00% (0/13) | Non-Toxin | 0.49 | Consensus (smm/nn/sturniolo) |
| 514 | nucleocapsid | TDMSTLVSAVITIEA | HLA-DQA1*04:01/DQB1*04:02 | 100.00% (13/13) | Non-Toxin | 0.07 | Consensus (comb.lib./smm/nn) |
| 515 | nucleocapsid | TEMRNLLSQSLSVRK | HLA-DRB4*01:01 | 100.00% (13/13) | Non-Toxin | 0.32 | Consensus (comb.lib./smm/nn) |
| 516 | nucleocapsid | TKIRIFVPATNSPEL | HLA-DRB1*04:05 | 100.00% (13/13) | Non-Toxin | 0.16 | Consensus (smm/nn/sturniolo) |
| 517 | nucleocapsid | TKIRIFVPATNSPEL | HLA-DRB1*08:02 | 100.00% (13/13) | Non-Toxin | 0.35 | Consensus (smm/nn/sturniolo) |
| 518 | nucleocapsid | TLFALDVIRSPSAAE | HLA-DRB1*11:01 | 100.00% (13/13) | Non-Toxin | 0.45 | Consensus (smm/nn/sturniolo) |

| | | | | | | | |
|---|---|---|---|---|---|---|---|
| 519 | nucleocapsid | TLTTKIRIFVPATNS | HLA-DRB1*08:02 | 100.00% (13/13) | Non-Toxin | 0.37 | Consensus (smm/nn/sturniolo) |
| 520 | nucleocapsid | TLVSAVITIEAQIWI | HLA-DQA1*03:01/DQB1*03:02 | 100.00% (13/13) | Non-Toxin | 0.19 | Consensus (comb.lib./smm/nn) |
| 521 | nucleocapsid | TNSLLNLRSRLAAKA | HLA-DRB1*15:01 | 100.00% (13/13) | Non-Toxin | 0.49 | Consensus (smm/nn/sturniolo) |
| 522 | nucleocapsid | TTKIRIFVPATNSPE | HLA-DRB1*08:02 | 100.00% (13/13) | Non-Toxin | 0.29 | Consensus (smm/nn/sturniolo) |
| 523 | nucleocapsid | TTKIRIFVPATNSPE | HLA-DRB1*04:05 | 100.00% (13/13) | Non-Toxin | 0.34 | Consensus (smm/nn/sturniolo) |
| 524 | nucleocapsid | VAELAAAVQETSAGR | HLA-DQA1*04:01/DQB1*04:02 | 100.00% (13/13) | Non-Toxin | 0.12 | Consensus (comb.lib./smm/nn) |
| 525 | nucleocapsid | VEIISDIGNYVEETG | HLA-DRB1*13:02 | 100.00% (13/13) | Non-Toxin | 0.34 | Consensus (smm/nn/sturniolo) |
| 526 | nucleocapsid | VIIDVGSMVNGIPVM | HLA-DRB1*03:01 | 100.00% (13/13) | Non-Toxin | 0.45 | Consensus (smm/nn/sturniolo) |
| 527 | nucleocapsid | WELTLFALDVIRSPS | HLA-DQA1*01:01/DQB1*05:01 | 100.00% (13/13) | Non-Toxin | 0.02 | Consensus (comb.lib./smm/nn) |
| 528 | nucleocapsid | WSFAMGVATTIDRSM | HLA-DRB1*09:01 | 100.00% (13/13) | Non-Toxin | 0.01 | Consensus (comb.lib./smm/nn) |
| 529 | nucleocapsid | YPLLWSFAMGVATTI | HLA-DRB1*09:01 | 100.00% (13/13) | Non-Toxin | 0.02 | Consensus (comb.lib./smm/nn) |
| **S.No.** | **Protein** | **Peptide** | **Allele** | **Conservancy** | **Toxicity** | **Percentile rank** | **Method used** |
| 530 | Phosphoprotein | AEGSDDIQLDPVVTD | HLA-DRB1*03:01 | 100.00% (27/27) | Non-Toxin | 0.13 | Consensus (smm/nn/sturniolo) |
| 531 | Phosphoprotein | CLVSDAKMLSYAPEI | HLA-DRB1*03:01 | 59.26% (16/27) | Non-Toxin | 0.02 | Consensus (smm/nn/sturniolo) |
| 532 | Phosphoprotein | DDIQLDPVVTDVVYH | HLA-DRB1*03:01 | 100.00% (27/27) | Non-Toxin | 0.13 | Consensus (smm/nn/sturniolo) |
| 533 | Phosphoprotein | DIQLDPVVTDVVYHD | HLA-DRB3*01:01 | 100.00% (27/27) | Non-Toxin | 0.02 | Consensus (comb.lib./smm/nn) |
| 534 | Phosphoprotein | DIQLDPVVTDVVYHD | HLA-DRB1*03:01 | 100.00% (27/27) | Non-Toxin | 0.27 | Consensus (smm/nn/sturniolo) |
| 535 | Phosphoprotein | DPVVTDVVYHDHGGE | HLA-DRB3*01:01 | 100.00% (27/27) | Non-Toxin | 0.02 | Consensus (comb.lib./smm/nn) |
| 536 | Phosphoprotein | DSIKLYTSDDEEADQ | HLA-DRB3*01:01 | 100.00% (27/27) | Non-Toxin | 0.28 | Consensus (comb.lib./smm/nn) |
| 537 | Phosphoprotein | EGSDDIQLDPVVTDV | HLA-DRB1*03:01 | 100.00% (27/27) | Non-Toxin | 0.13 | Consensus (smm/nn/sturniolo) |
| 538 | Phosphoprotein | EQQSLFSFDNVKNFR | HLA-DRB3*01:01 | 100.00% (27/27) | Non-Toxin | 0.14 | Consensus (comb.lib./smm/nn) |
| 539 | Phosphoprotein | FSFDNVKNFRDGSLT | HLA-DRB3*01:01 | 100.00% (27/27) | Non-Toxin | 0.01 | Consensus (comb.lib./smm/nn) |
| 540 | Phosphoprotein | GHLVSMMIMIPGKGK | HLA-DRB5*01:01 | 100.00% (27/27) | Non-Toxin | 0.27 | Consensus (smm/nn/sturniolo) |
| 541 | Phosphoprotein | GNVCLVSDAKMLSYA | HLA-DRB1*03:01 | 59.26% (16/27) | Non-Toxin | 0.01 | Consensus (smm/nn/sturniolo) |
| 542 | Phosphoprotein | GSDDIQLDPVVTDVV | HLA-DRB1*03:01 | 100.00% (27/27) | Non-Toxin | 0.13 | Consensus (smm/nn/sturniolo) |
| 543 | Phosphoprotein | GVINSIKLINLDMRL | HLA-DRB4*01:01 | 100.00% (27/27) | Non-Toxin | 0.44 | Consensus (comb.lib./smm/nn) |
| 544 | Phosphoprotein | HLVSMMIMIPGKGKG | HLA-DRB5*01:01 | 100.00% (27/27) | Non-Toxin | 0.28 | Consensus (smm/nn/sturniolo) |
| 545 | Phosphoprotein | IEGHLVSMMIMIPGK | HLA-DRB4*01:01 | 100.00% (27/27) | Non-Toxin | 0.47 | Consensus (comb.lib./smm/nn) |
| 546 | Phosphoprotein | IKLINLDMRLNHIEE | HLA-DRB1*03:01 | 100.00% (27/27) | Non-Toxin | 0.1 | Consensus (smm/nn/sturniolo) |
| 547 | Phosphoprotein | IKLYTSDDEEADQLE | HLA-DRB3*01:01 | 100.00% (27/27) | Non-Toxin | 0.27 | Consensus (comb.lib./smm/nn) |
| 548 | Phosphoprotein | INLDMRLNHIEEQVK | HLA-DRB1*03:01 | 100.00% (27/27) | Non-Toxin | 0.35 | Consensus (smm/nn/sturniolo) |
| 549 | Phosphoprotein | INSIKLINLDMRLNH | HLA-DRB1*03:01 | 100.00% (27/27) | Non-Toxin | 0.33 | Consensus (smm/nn/sturniolo) |
| 550 | Phosphoprotein | IQLDPVVTDVVYHDH | HLA-DRB3*01:01 | 100.00% (27/27) | Non-Toxin | 0.02 | Consensus (comb.lib./smm/nn) |
| 551 | Phosphoprotein | IQLDPVVTDVVYHDH | HLA-DRB1*03:01 | 100.00% (27/27) | Non-Toxin | 0.26 | Consensus (smm/nn/sturniolo) |
| 552 | Phosphoprotein | KLINLDMRLNHIEEQ | HLA-DRB1*03:01 | 100.00% (27/27) | Non-Toxin | 0.31 | Consensus (smm/nn/sturniolo) |
| 553 | Phosphoprotein | KLYTSDDEEADQLEF | HLA-DRB3*01:01 | 100.00% (27/27) | Non-Toxin | 0.28 | Consensus (comb.lib./smm/nn) |
| 554 | Phosphoprotein | LDPVVTDVVYHDHGG | HLA-DRB3*01:01 | 100.00% (27/27) | Non-Toxin | 0.02 | Consensus (comb.lib./smm/nn) |
| 555 | Phosphoprotein | LDSIKLYTSDDEEAD | HLA-DRB3*01:01 | 100.00% (27/27) | Non-Toxin | 0.27 | Consensus (comb.lib./smm/nn) |
| 556 | Phosphoprotein | LEFEDEFAGSSSEVI | HLA-DRB1*07:01 | 100.00% (27/27) | Non-Toxin | 0.45 | Consensus (comb.lib./smm/nn) |
| 557 | Phosphoprotein | LEQQSLFSFDNVKNF | HLA-DRB3*01:01 | 100.00% (27/27) | Non-Toxin | 0.15 | Consensus (comb.lib./smm/nn) |
| 558 | Phosphoprotein | LFSFDNVKNFRDGSL | HLA-DRB3*01:01 | 100.00% (27/27) | Non-Toxin | 0.01 | Consensus (comb.lib./smm/nn) |
| 559 | Phosphoprotein | LINLDMRLNHIEEQV | HLA-DRB1*03:01 | 100.00% (27/27) | Non-Toxin | 0.35 | Consensus (smm/nn/sturniolo) |
| 560 | Phosphoprotein | LTNEPYGAAVQLRED | HLA-DQA1*04:01/DQB1*04:02 | 62.96% (17/27) | Non-Toxin | 0.21 | Consensus (comb.lib./smm/nn) |
| 561 | Phosphoprotein | LVSDAKMLSYAPEIA | HLA-DRB1*03:01 | 59.26% (16/27) | Non-Toxin | 0.02 | Consensus (smm/nn/sturniolo) |
| 562 | Phosphoprotein | LVSMMIMIPGKGKGE | HLA-DRB5*01:01 | 100.00% (27/27) | Non-Toxin | 0.28 | Consensus (smm/nn/sturniolo) |
| 563 | Phosphoprotein | MPSDDFSNTFFPHDT | HLA-DPA1*01/DPB1*04:01 | 96.30% (26/27) | Non-Toxin | 0.49 | Consensus (comb.lib./smm/nn) |
| 564 | Phosphoprotein | NEPYGAAVQLREDLI | HLA-DQA1*04:01/DQB1*04:02 | 62.96% (17/27) | Non-Toxin | 0.39 | Consensus (comb.lib./smm/nn) |
| 565 | Phosphoprotein | NGNVCLVSDAKMLSY | HLA-DRB1*03:01 | 59.26% (16/27) | Non-Toxin | 0.01 | Consensus (smm/nn/sturniolo) |

| | | | | | | | |
|---|---|---|---|---|---|---|---|
| 566 | Phosphoprotein | NNGNVCLVSDAKMLS | HLA-DRB1*03:01 | 62.96% (17/27) | Non-Toxin | 0.01 | Consensus (smm/nn/sturniolo) |
| 567 | Phosphoprotein | NSIKLINLDMRLNHI | HLA-DRB1*03:01 | 100.00% (27/27) | Non-Toxin | 0.16 | Consensus (smm/nn/sturniolo) |
| 568 | Phosphoprotein | NVCLVSDAKMLSYAP | HLA-DRB1*03:01 | 59.26% (16/27) | Non-Toxin | 0.01 | Consensus (smm/nn/sturniolo) |
| 569 | Phosphoprotein | PSDDFSNTFFPHDTD | HLA-DPA1*01/DPB1*04:01 | 96.30% (26/27) | Non-Toxin | 0.51 | Consensus (comb.lib./smm/nn) |
| 570 | Phosphoprotein | PVVTDVVYHDHGGEC | HLA-DRB3*01:01 | 100.00% (27/27) | Non-Toxin | 0.2 | Consensus (comb.lib./smm/nn) |
| 571 | Phosphoprotein | QLDPVVTDVVYHDHG | HLA-DRB3*01:01 | 100.00% (27/27) | Non-Toxin | 0.02 | Consensus (comb.lib./smm/nn) |
| 572 | Phosphoprotein | QQSLFSFDNVKNFRD | HLA-DRB3*01:01 | 100.00% (27/27) | Non-Toxin | 0.01 | Consensus (comb.lib./smm/nn) |
| 573 | Phosphoprotein | QSLFSFDNVKNFRDG | HLA-DRB3*01:01 | 100.00% (27/27) | Non-Toxin | 0.01 | Consensus (comb.lib./smm/nn) |
| 574 | Phosphoprotein | SDDFSNTFFPHDTDR | HLA-DPA1*01/DPB1*04:01 | 96.30% (26/27) | Non-Toxin | 0.5 | Consensus (comb.lib./smm/nn) |
| 575 | Phosphoprotein | SDDIQLDPVVTDVVY | HLA-DRB1*03:01 | 100.00% (27/27) | Non-Toxin | 0.13 | Consensus (smm/nn/sturniolo) |
| 576 | Phosphoprotein | SIKLINLDMRLNHIE | HLA-DRB1*03:01 | 100.00% (27/27) | Non-Toxin | 0.11 | Consensus (smm/nn/sturniolo) |
| 577 | Phosphoprotein | SIKLYTSDDEEADQL | HLA-DRB3*01:01 | 100.00% (27/27) | Non-Toxin | 0.27 | Consensus (comb.lib./smm/nn) |
| 578 | Phosphoprotein | SLFSFDNVKNFRDGS | HLA-DRB3*01:01 | 100.00% (27/27) | Non-Toxin | 0.01 | Consensus (comb.lib./smm/nn) |
| 579 | Phosphoprotein | SLTNEPYGAAVQLRE | HLA-DQA1*04:01/DQB1*04:02 | 62.96% (17/27) | Non-Toxin | 0.19 | Consensus (comb.lib./smm/nn) |
| 580 | Phosphoprotein | SMMIMIPGKGKGERK | HLA-DRB5*01:01 | 100.00% (27/27) | Non-Toxin | 0.3 | Consensus (smm/nn/sturniolo) |
| 581 | Phosphoprotein | SSEVIVGISPEDEEP | HLA-DQA1*04:01/DQB1*04:02 | 62.96% (17/27) | Non-Toxin | 0.39 | Consensus (comb.lib./smm/nn) |
| 582 | Phosphoprotein | SSSEVIVGISPEDEE | HLA-DQA1*04:01/DQB1*04:02 | 62.96% (17/27) | Non-Toxin | 0.29 | Consensus (comb.lib./smm/nn) |
| 583 | Phosphoprotein | TIEGHLVSMMIMIPG | HLA-DRB4*01:01 | 100.00% (27/27) | Non-Toxin | 0.48 | Consensus (comb.lib./smm/nn) |
| 584 | Phosphoprotein | TNEPYGAAVQLREDL | HLA-DQA1*04:01/DQB1*04:02 | 62.96% (17/27) | Non-Toxin | 0.25 | Consensus (comb.lib./smm/nn) |
| 585 | Phosphoprotein | VCLVSDAKMLSYAPE | HLA-DRB1*03:01 | 59.26% (16/27) | Non-Toxin | 0.01 | Consensus (smm/nn/sturniolo) |
| 586 | Phosphoprotein | VSMMIMIPGKGKGER | HLA-DRB5*01:01 | 100.00% (27/27) | Non-Toxin | 0.27 | Consensus (smm/nn/sturniolo) |
| 587 | Phosphoprotein | VVTDVVYHDHGGECT | HLA-DRB3*01:01 | 100.00% (27/27) | Non-Toxin | 0.21 | Consensus (comb.lib./smm/nn) |
| **S.No.** | **Protein** | **Peptide** | **Allele** | **Conservancy** | **Toxicity** | **Percentile rank** | **Method used** |
| 588 | Polymerase | ASFLMDRRVILPRAA | HLA-DRB3*01:01 | 100.00% (20/20) | Non-Toxin | 0.01 | Consensus (comb.lib./smm/nn) |
| 589 | Polymerase | ASFLMDRRVILPRAA | HLA-DRB1*03:01 | 100.00% (20/20) | Non-Toxin | 0.01 | Consensus (smm/nn/sturniolo) |
| 590 | Polymerase | ASQRVNIDLDVLKAI | HLA-DRB3*01:01 | 100.00% (20/20) | Non-Toxin | 0.01 | Consensus (comb.lib./smm/nn) |
| 591 | Polymerase | CINILKVIQQLLIST | HLA-DRB4*01:01 | 100.00% (20/20) | Non-Toxin | 0.01 | Consensus (comb.lib./smm/nn) |
| 592 | Polymerase | CLNWRYESMAIFAER | HLA-DRB3*01:01 | 100.00% (20/20) | Non-Toxin | 0.01 | Consensus (comb.lib./smm/nn) |
| 593 | Polymerase | DDLKKYYQIDQPFFV | HLA-DRB3*01:01 | 100.00% (20/20) | Non-Toxin | 0.01 | Consensus (comb.lib./smm/nn) |
| 594 | Polymerase | DLELASFLMDRRVIL | HLA-DRB3*01:01 | 100.00% (20/20) | Non-Toxin | 0.01 | Consensus (comb.lib./smm/nn) |
| 595 | Polymerase | DLELASFLMDRRVIL | HLA-DRB1*03:01 | 100.00% (20/20) | Non-Toxin | 0.02 | Consensus (smm/nn/sturniolo) |
| 596 | Polymerase | DLKKYYQIDQPFFVP | HLA-DRB3*01:01 | 100.00% (20/20) | Non-Toxin | 0.01 | Consensus (comb.lib./smm/nn) |
| 597 | Polymerase | DLSMYMKDKALSPIK | HLA-DRB3*01:01 | 100.00% (20/20) | Non-Toxin | 0.01 | Consensus (comb.lib./smm/nn) |
| 598 | Polymerase | DNIHLLAEFFSFFRT | HLA-DPA1*01:03/DPB1*02:01 | 100.00% (20/20) | Non-Toxin | 0.01 | Consensus (comb.lib./smm/nn) |
| 599 | Polymerase | DQKIRSMFIDDLLSI | HLA-DRB3*01:01 | 100.00% (20/20) | Non-Toxin | 0.01 | Consensus (comb.lib./smm/nn) |
| 600 | Polymerase | DSDLSMYMKDKALSP | HLA-DRB3*01:01 | 100.00% (20/20) | Non-Toxin | 0.01 | Consensus (comb.lib./smm/nn) |
| 601 | Polymerase | DTIIIMLNEAMNYFD | HLA-DRB3*01:01 | 100.00% (20/20) | Non-Toxin | 0.01 | Consensus (comb.lib./smm/nn) |
| 602 | Polymerase | DYNQFLILNKLLSNR | HLA-DRB1*11:01 | 100.00% (20/20) | Non-Toxin | 0.01 | Consensus (smm/nn/sturniolo) |
| 603 | Polymerase | ELASFLMDRRVILPR | HLA-DRB3*01:01 | 100.00% (20/20) | Non-Toxin | 0.01 | Consensus (comb.lib./smm/nn) |
| 604 | Polymerase | ELASFLMDRRVILPR | HLA-DRB1*03:01 | 100.00% (20/20) | Non-Toxin | 0.01 | Consensus (smm/nn/sturniolo) |
| 605 | Polymerase | ETLTLDVTSPISNNL | HLA-DRB3*01:01 | 100.00% (20/20) | Non-Toxin | 0.01 | Consensus (comb.lib./smm/nn) |
| 606 | Polymerase | FCLNWRYESMAIFAE | HLA-DRB3*01:01 | 100.00% (20/20) | Non-Toxin | 0.01 | Consensus (comb.lib./smm/nn) |
| 607 | Polymerase | FLILNKLLSNRRQND | HLA-DRB1*11:01 | 100.00% (20/20) | Non-Toxin | 0.01 | Consensus (smm/nn/sturniolo) |
| 608 | Polymerase | FSINETLTLDVTSPI | HLA-DRB3*01:01 | 100.00% (20/20) | Non-Toxin | 0.01 | Consensus (comb.lib./smm/nn) |
| 609 | Polymerase | FVIFYASLTYLRRGI | HLA-DPA1*02:01/DPB1*14:01 | 100.00% (20/20) | Non-Toxin | 0.01 | NetMHCIIpan |
| 610 | Polymerase | GYCINILKVIQQLLI | HLA-DRB4*01:01 | 100.00% (20/20) | Non-Toxin | 0.02 | Consensus (comb.lib./smm/nn) |
| 611 | Polymerase | HDYNQFLILNKLLSN | HLA-DRB1*11:01 | 100.00% (20/20) | Non-Toxin | 0.01 | Consensus (smm/nn/sturniolo) |
| 612 | Polymerase | HHDYNQFLILNKLLS | HLA-DRB1*11:01 | 100.00% (20/20) | Non-Toxin | 0.01 | Consensus (smm/nn/sturniolo) |

| | | | | | | | |
|---|---|---|---|---|---|---|---|
| 613 | Polymerase | HHKPPWIIDLNPQEK | HLA-DRB3*01:01 | 100.00% (20/20) | Non-Toxin | 0.02 | Consensus (comb.lib./smm/nn) |
| 614 | Polymerase | HKPPWIIDLNPQEKI | HLA-DRB3*01:01 | 100.00% (20/20) | Non-Toxin | 0.02 | Consensus (comb.lib./smm/nn) |
| 615 | Polymerase | IDNIHLLAEFFSFFR | HLA-DPA1*01:03/DPB1*02:01 | 100.00% (20/20) | Non-Toxin | 0.01 | Consensus (comb.lib./smm/nn) |
| 616 | Polymerase | IHLLAEFFSFFRTFG | HLA-DPA1*01:03/DPB1*02:01 | 100.00% (20/20) | Non-Toxin | 0.01 | Consensus (comb.lib./smm/nn) |
| 617 | Polymerase | IIIMLNEAMNYFDDN | HLA-DRB3*01:01 | 100.00% (20/20) | Non-Toxin | 0.01 | Consensus (comb.lib./smm/nn) |
| 618 | Polymerase | ILKVIQQLLISTEFS | HLA-DRB4*01:01 | 100.00% (20/20) | Non-Toxin | 0.01 | Consensus (comb.lib./smm/nn) |
| 619 | Polymerase | INETLTLDVTSPISN | HLA-DRB3*01:01 | 100.00% (20/20) | Non-Toxin | 0.01 | Consensus (comb.lib./smm/nn) |
| 620 | Polymerase | INILKVIQQLLISTE | HLA-DRB4*01:01 | 100.00% (20/20) | Non-Toxin | 0.01 | Consensus (comb.lib./smm/nn) |
| 621 | Polymerase | IRSMFIDDLLSILNI | HLA-DRB3*01:01 | 100.00% (20/20) | Non-Toxin | 0.01 | Consensus (comb.lib./smm/nn) |
| 622 | Polymerase | KIRSMFIDDLLSILN | HLA-DRB3*01:01 | 100.00% (20/20) | Non-Toxin | 0.01 | Consensus (comb.lib./smm/nn) |
| 623 | Polymerase | KKYYQIDQPFFVPTK | HLA-DRB3*01:01 | 100.00% (20/20) | Non-Toxin | 0.01 | Consensus (comb.lib./smm/nn) |
| 624 | Polymerase | KPPWIIDLNPQEKIC | HLA-DRB3*01:01 | 100.00% (20/20) | Non-Toxin | 0.02 | Consensus (comb.lib./smm/nn) |
| 625 | Polymerase | KYYQIDQPFFVPTKI | HLA-DRB3*01:01 | 100.00% (20/20) | Non-Toxin | 0.01 | Consensus (comb.lib./smm/nn) |
| 626 | Polymerase | LASFLMDRRVILPRA | HLA-DRB3*01:01 | 100.00% (20/20) | Non-Toxin | 0.01 | Consensus (comb.lib./smm/nn) |
| 627 | Polymerase | LASFLMDRRVILPRA | HLA-DRB1*03:01 | 100.00% (20/20) | Non-Toxin | 0.01 | Consensus (smm/nn/sturniolo) |
| 628 | Polymerase | LDFVIFYASLTYLRR | HLA-DPA1*02:01/DPB1*14:01 | 100.00% (20/20) | Non-Toxin | 0.01 | NetMHCIIpan |
| 629 | Polymerase | LELASFLMDRRVILP | HLA-DRB3*01:01 | 100.00% (20/20) | Non-Toxin | 0.01 | Consensus (comb.lib./smm/nn) |
| 630 | Polymerase | LELASFLMDRRVILP | HLA-DRB1*03:01 | 100.00% (20/20) | Non-Toxin | 0.01 | Consensus (smm/nn/sturniolo) |
| 631 | Polymerase | LKKYYQIDQPFFVPT | HLA-DRB3*01:01 | 100.00% (20/20) | Non-Toxin | 0.01 | Consensus (comb.lib./smm/nn) |
| 632 | Polymerase | LNWRYESMAIFAERL | HLA-DRB3*01:01 | 100.00% (20/20) | Non-Toxin | 0.01 | Consensus (comb.lib./smm/nn) |
| 633 | Polymerase | LSMYMKDKALSPIKD | HLA-DRB3*01:01 | 100.00% (20/20) | Non-Toxin | 0.01 | Consensus (comb.lib./smm/nn) |
| 634 | Polymerase | MFIDDLLSILNIDNI | HLA-DRB3*01:01 | 100.00% (20/20) | Non-Toxin | 0.01 | Consensus (comb.lib./smm/nn) |
| 635 | Polymerase | NETLTLDVTSPISNN | HLA-DRB3*01:01 | 100.00% (20/20) | Non-Toxin | 0.01 | Consensus (comb.lib./smm/nn) |
| 636 | Polymerase | NHHKPPWIIDLNPQE | HLA-DRB3*01:01 | 100.00% (20/20) | Non-Toxin | 0.02 | Consensus (comb.lib./smm/nn) |
| 637 | Polymerase | NIDNIHLLAEFFSFF | HLA-DPA1*01:03/DPB1*02:01 | 100.00% (20/20) | Non-Toxin | 0.01 | Consensus (comb.lib./smm/nn) |
| 638 | Polymerase | NIHLLAEFFSFFRTF | HLA-DPA1*01:03/DPB1*02:01 | 100.00% (20/20) | Non-Toxin | 0.01 | Consensus (comb.lib./smm/nn) |
| 639 | Polymerase | NILKVIQQLLISTEF | HLA-DRB4*01:01 | 100.00% (20/20) | Non-Toxin | 0.01 | Consensus (comb.lib./smm/nn) |
| 640 | Polymerase | NQFLILNKLLSNRRQ | HLA-DRB1*11:01 | 100.00% (20/20) | Non-Toxin | 0.01 | Consensus (smm/nn/sturniolo) |
| 641 | Polymerase | NWRYESMAIFAERLD | HLA-DRB3*01:01 | 100.00% (20/20) | Non-Toxin | 0.01 | Consensus (comb.lib./smm/nn) |
| 642 | Polymerase | PPWIIDLNPQEKICV | HLA-DRB3*01:01 | 100.00% (20/20) | Non-Toxin | 0.02 | Consensus (comb.lib./smm/nn) |
| 643 | Polymerase | QFLILNKLLSNRRQN | HLA-DRB1*11:01 | 100.00% (20/20) | Non-Toxin | 0.01 | Consensus (smm/nn/sturniolo) |
| 644 | Polymerase | QKIRSMFIDDLLSIL | HLA-DRB3*01:01 | 100.00% (20/20) | Non-Toxin | 0.01 | Consensus (comb.lib./smm/nn) |
| 645 | Polymerase | QRVNIDLDVLKAITP | HLA-DRB3*01:01 | 100.00% (20/20) | Non-Toxin | 0.01 | Consensus (comb.lib./smm/nn) |
| 646 | Polymerase | RDTIIIMLNEAMNYF | HLA-DRB3*01:01 | 100.00% (20/20) | Non-Toxin | 0.01 | Consensus (comb.lib./smm/nn) |
| 647 | Polymerase | RKVLILDFRSKLMTK | HLA-DRB1*03:01 | 100.00% (20/20) | Non-Toxin | 0.01 | Consensus (smm/nn/sturniolo) |
| 648 | Polymerase | RSMFIDDLLSILNID | HLA-DRB3*01:01 | 100.00% (20/20) | Non-Toxin | 0.01 | Consensus (comb.lib./smm/nn) |
| 649 | Polymerase | RVNIDLDVLKAITPV | HLA-DRB3*01:01 | 100.00% (20/20) | Non-Toxin | 0.01 | Consensus (comb.lib./smm/nn) |
| 650 | Polymerase | SDLSMYMKDKALSPI | HLA-DRB3*01:01 | 100.00% (20/20) | Non-Toxin | 0.01 | Consensus (comb.lib./smm/nn) |
| 651 | Polymerase | SFLMDRRVILPRAAH | HLA-DRB3*01:01 | 100.00% (20/20) | Non-Toxin | 0.01 | Consensus (comb.lib./smm/nn) |
| 652 | Polymerase | SINETLTLDVTSPIS | HLA-DRB3*01:01 | 100.00% (20/20) | Non-Toxin | 0.01 | Consensus (comb.lib./smm/nn) |
| 653 | Polymerase | SMFIDDLLSILNIDN | HLA-DRB3*01:01 | 100.00% (20/20) | Non-Toxin | 0.01 | Consensus (comb.lib./smm/nn) |
| 654 | Polymerase | SMYMKDKALSPIKDE | HLA-DRB3*01:01 | 100.00% (20/20) | Non-Toxin | 0.01 | Consensus (comb.lib./smm/nn) |
| 655 | Polymerase | SQRVNIDLDVLKAIT | HLA-DRB3*01:01 | 100.00% (20/20) | Non-Toxin | 0.01 | Consensus (comb.lib./smm/nn) |
| 656 | Polymerase | TIIIMLNEAMNYFDD | HLA-DRB3*01:01 | 100.00% (20/20) | Non-Toxin | 0.01 | Consensus (comb.lib./smm/nn) |
| 657 | Polymerase | VNIDLDVLKAITPVS | HLA-DRB3*01:01 | 100.00% (20/20) | Non-Toxin | 0.01 | Consensus (comb.lib./smm/nn) |
| 658 | Polymerase | WRYESMAIFAERLDE | HLA-DRB3*01:01 | 100.00% (20/20) | Non-Toxin | 0.01 | Consensus (comb.lib./smm/nn) |
| 659 | Polymerase | YCINILKVIQQLLIS | HLA-DRB4*01:01 | 100.00% (20/20) | Non-Toxin | 0.01 | Consensus (comb.lib./smm/nn) |
| 660 | Polymerase | YNQFLILNKLLSNRR | HLA-DRB1*11:01 | 100.00% (20/20) | Non-Toxin | 0.01 | Consensus (smm/nn/sturniolo) |

| | | | | | | | |
|---|---|---|---|---|---|---|---|
| 661 | Polymerase | YQIDQPFFVPTKITS | HLA-DRB3*01:01 | 100.00% (20/20) | Non-Toxin | 0.01 | Consensus (comb.lib./smm/nn) |
| 662 | Polymerase | YYQIDQPFFVPTKIT | HLA-DRB3*01:01 | 100.00% (20/20) | Non-Toxin | 0.01 | Consensus (comb.lib./smm/nn) |
| **S.No.** | **Protein** | **Peptide** | **Allele** | **Conservancy** | **Toxicity** | **Percentile rank** | **Method used** |
| 663 | V Protein | AEGSDDIQLDPVVTD | HLA-DRB1*03:01 | 94.74% (54/57) | Non-Toxin | 0.13 | Consensus (smm/nn/sturniolo) |
| 664 | V Protein | AKMLSYAPEIAVSKE | HLA-DRB1*09:01 | 19.30% (11/57) | Non-Toxin | 0.55 | Consensus (comb.lib./smm/nn) |
| 665 | V Protein | AQPPYHWSIERSISP | HLA-DRB1*09:01 | 19.30% (11/57) | Non-Toxin | 0.85 | Consensus (comb.lib./smm/nn) |
| 666 | V Protein | CLVSDAKMLSYAPEI | HLA-DRB1*03:01 | 19.30% (11/57) | Non-Toxin | 0.02 | Consensus (smm/nn/sturniolo) |
| 667 | V Protein | DAKMLSYAPEIAVSK | HLA-DRB1*09:01 | 19.30% (11/57) | Non-Toxin | 0.54 | Consensus (comb.lib./smm/nn) |
| 668 | V Protein | DDIQLDPVVTDVVYH | HLA-DRB1*03:01 | 92.98% (53/57) | Non-Toxin | 0.13 | Consensus (smm/nn/sturniolo) |
| 669 | V Protein | DGKRAWVEEWCNPAC | HLA-DQA1*01:01/DQB1*05:01 | 87.72% (50/57) | Non-Toxin | 0.82 | Consensus (comb.lib./smm/nn) |
| 670 | V Protein | DIQLDPVVTDVVYHD | HLA-DRB3*01:01 | 92.98% (53/57) | Non-Toxin | 0.02 | Consensus (comb.lib./smm/nn) |
| 671 | V Protein | DIQLDPVVTDVVYHD | HLA-DRB1*03:01 | 92.98% (53/57) | Non-Toxin | 0.27 | Consensus (smm/nn/sturniolo) |
| 672 | V Protein | DKLELVNDGLNIIDF | HLA-DRB1*03:01 | 100.00% (57/57) | Non-Toxin | 0.57 | Consensus (smm/nn/sturniolo) |
| 673 | V Protein | DKLELVNDGLNIIDF | HLA-DRB3*01:01 | 100.00% (57/57) | Non-Toxin | 1.02 | Consensus (comb.lib./smm/nn) |
| 674 | V Protein | DPVVTDVVYHDHGGE | HLA-DRB3*01:01 | 98.25% (56/57) | Non-Toxin | 0.02 | Consensus (comb.lib./smm/nn) |
| 675 | V Protein | DPVVTDVVYHDHGGE | HLA-DRB1*03:01 | 98.25% (56/57) | Non-Toxin | 0.6 | Consensus (smm/nn/sturniolo) |
| 676 | V Protein | DSIKLYTSDDEEADQ | HLA-DRB3*01:01 | 100.00% (57/57) | Non-Toxin | 0.28 | Consensus (comb.lib./smm/nn) |
| 677 | V Protein | EFEDEFAGSSSEVIV | HLA-DRB1*07:01 | 98.25% (56/57) | Non-Toxin | 0.56 | Consensus (comb.lib./smm/nn) |
| 678 | V Protein | EGSDDIQLDPVVTDV | HLA-DRB1*03:01 | 94.74% (54/57) | Non-Toxin | 0.13 | Consensus (smm/nn/sturniolo) |
| 679 | V Protein | GNVCLVSDAKMLSYA | HLA-DRB1*03:01 | 19.30% (11/57) | Non-Toxin | 0.01 | Consensus (smm/nn/sturniolo) |
| 680 | V Protein | GNVCLVSDAKMLSYA | HLA-DRB1*11:01 | 19.30% (11/57) | Non-Toxin | 0.7 | Consensus (smm/nn/sturniolo) |
| 681 | V Protein | GSDDIQLDPVVTDVV | HLA-DRB1*03:01 | 94.74% (54/57) | Non-Toxin | 0.13 | Consensus (smm/nn/sturniolo) |
| 682 | V Protein | GSSSEVIVGISPEDE | HLA-DQA1*04:01/DQB1*04:02 | 19.30% (11/57) | Non-Toxin | 0.83 | Consensus (comb.lib./smm/nn) |
| 683 | V Protein | IKLYTSDDEEADQLE | HLA-DRB3*01:01 | 100.00% (57/57) | Non-Toxin | 0.27 | Consensus (comb.lib./smm/nn) |
| 684 | V Protein | IQLDPVVTDVVYHDH | HLA-DRB3*01:01 | 92.98% (53/57) | Non-Toxin | 0.02 | Consensus (comb.lib./smm/nn) |
| 685 | V Protein | IQLDPVVTDVVYHDH | HLA-DRB1*03:01 | 92.98% (53/57) | Non-Toxin | 0.26 | Consensus (smm/nn/sturniolo) |
| 686 | V Protein | KLELVNDGLNIIDFI | HLA-DRB1*03:01 | 100.00% (57/57) | Non-Toxin | 0.53 | Consensus (smm/nn/sturniolo) |
| 687 | V Protein | KLELVNDGLNIIDFI | HLA-DRB3*01:01 | 100.00% (57/57) | Non-Toxin | 1 | Consensus (comb.lib./smm/nn) |
| 688 | V Protein | KLYTSDDEEADQLEF | HLA-DRB3*01:01 | 100.00% (57/57) | Non-Toxin | 0.28 | Consensus (comb.lib./smm/nn) |
| 689 | V Protein | KMLSYAPEIAVSKED | HLA-DRB1*09:01 | 19.30% (11/57) | Non-Toxin | 0.67 | Consensus (comb.lib./smm/nn) |
| 690 | V Protein | LDPVVTDVVYHDHGG | HLA-DRB3*01:01 | 98.25% (56/57) | Non-Toxin | 0.02 | Consensus (comb.lib./smm/nn) |
| 691 | V Protein | LDPVVTDVVYHDHGG | HLA-DRB1*03:01 | 98.25% (56/57) | Non-Toxin | 0.56 | Consensus (smm/nn/sturniolo) |
| 692 | V Protein | LDSIKLYTSDDEEAD | HLA-DRB3*01:01 | 100.00% (57/57) | Non-Toxin | 0.27 | Consensus (comb.lib./smm/nn) |
| 693 | V Protein | LEFEDEFAGSSSEVI | HLA-DRB1*07:01 | 100.00% (57/57) | Non-Toxin | 0.45 | Consensus (comb.lib./smm/nn) |
| 694 | V Protein | LELVNDGLNIIDFIQ | HLA-DRB1*03:01 | 100.00% (57/57) | Non-Toxin | 0.55 | Consensus (smm/nn/sturniolo) |
| 695 | V Protein | LVSDAKMLSYAPEIA | HLA-DRB1*03:01 | 19.30% (11/57) | Non-Toxin | 0.02 | Consensus (smm/nn/sturniolo) |
| 696 | V Protein | LYTSDDEEADQLEFE | HLA-DQA1*03:01/DQB1*03:02 | 100.00% (57/57) | Non-Toxin | 0.79 | Consensus (comb.lib./smm/nn) |
| 697 | V Protein | MDKLELVNDGLNIID | HLA-DRB1*03:01 | 100.00% (57/57) | Non-Toxin | 0.59 | Consensus (smm/nn/sturniolo) |
| 698 | V Protein | NGNVCLVSDAKMLSY | HLA-DRB1*03:01 | 19.30% (11/57) | Non-Toxin | 0.01 | Consensus (smm/nn/sturniolo) |
| 699 | V Protein | NGNVCLVSDAKMLSY | HLA-DRB1*11:01 | 19.30% (11/57) | Non-Toxin | 0.7 | Consensus (smm/nn/sturniolo) |
| 700 | V Protein | NNGNVCLVSDAKMLS | HLA-DRB1*03:01 | 19.30% (11/57) | Non-Toxin | 0.01 | Consensus (smm/nn/sturniolo) |
| 701 | V Protein | NNGNVCLVSDAKMLS | HLA-DRB1*11:01 | 19.30% (11/57) | Non-Toxin | 0.7 | Consensus (smm/nn/sturniolo) |
| 702 | V Protein | NVCLVSDAKMLSYAP | HLA-DRB1*03:01 | 19.30% (11/57) | Non-Toxin | 0.01 | Consensus (smm/nn/sturniolo) |
| 703 | V Protein | NVCLVSDAKMLSYAP | HLA-DRB1*11:01 | 19.30% (11/57) | Non-Toxin | 0.7 | Consensus (smm/nn/sturniolo) |
| 704 | V Protein | NVNLDSIKLYTSDDE | HLA-DRB1*03:01 | 100.00% (57/57) | Non-Toxin | 0.76 | Consensus (smm/nn/sturniolo) |
| 705 | V Protein | PPYHWSIERSISPDK | HLA-DRB1*09:01 | 19.30% (11/57) | Non-Toxin | 0.99 | Consensus (comb.lib./smm/nn) |
| 706 | V Protein | PQTSRNVNLDSIKLY | HLA-DRB1*03:01 | 19.30% (11/57) | Non-Toxin | 0.76 | Consensus (smm/nn/sturniolo) |
| 707 | V Protein | PVVTDVVYHDHGGEC | HLA-DRB3*01:01 | 98.25% (56/57) | Non-Toxin | 0.2 | Consensus (comb.lib./smm/nn) |

| | | | | | | | |
|---|---|---|---|---|---|---|---|
| 708 | V Protein | QLDPVVTDVVYHDHG | HLA-DRB3*01:01 | 98.25% (56/57) | Non-Toxin | 0.02 | Consensus (comb.lib./smm/nn) |
| 709 | V Protein | QLDPVVTDVVYHDHG | HLA-DRB1*03:01 | 98.25% (56/57) | Non-Toxin | 0.52 | Consensus (smm/nn/sturniolo) |
| 710 | V Protein | QPPYHWSIERSISPD | HLA-DRB1*09:01 | 19.30% (11/57) | Non-Toxin | 0.56 | Consensus (comb.lib./smm/nn) |
| 711 | V Protein | QTSRNVNLDSIKLYT | HLA-DRB1*03:01 | 19.30% (11/57) | Non-Toxin | 0.76 | Consensus (smm/nn/sturniolo) |
| 712 | V Protein | RNVNLDSIKLYTSDD | HLA-DRB1*03:01 | 78.95% (45/57) | Non-Toxin | 0.76 | Consensus (smm/nn/sturniolo) |
| 713 | V Protein | SDAKMLSYAPEIAVS | HLA-DRB1*09:01 | 19.30% (11/57) | Non-Toxin | 0.55 | Consensus (comb.lib./smm/nn) |
| 714 | V Protein | SDDIQLDPVVTDVVY | HLA-DRB1*03:01 | 92.98% (53/57) | Non-Toxin | 0.13 | Consensus (smm/nn/sturniolo) |
| 715 | V Protein | SEVIVGISPEDEEPS | HLA-DQA1*04:01/DQB1*04:02 | 19.30% (11/57) | Non-Toxin | 0.65 | Consensus (comb.lib./smm/nn) |
| 716 | V Protein | SIKLYTSDDEEADQL | HLA-DRB3*01:01 | 100.00% (57/57) | Non-Toxin | 0.27 | Consensus (comb.lib./smm/nn) |
| 717 | V Protein | SRNVNLDSIKLYTSD | HLA-DRB1*03:01 | 19.30% (11/57) | Non-Toxin | 0.76 | Consensus (smm/nn/sturniolo) |
| 718 | V Protein | SSEVIVGISPEDEEP | HLA-DQA1*04:01/DQB1*04:02 | 19.30% (11/57) | Non-Toxin | 0.39 | Consensus (comb.lib./smm/nn) |
| 719 | V Protein | SSEVIVGISPEDEEP | HLA-DQA1*03:01/DQB1*03:02 | 19.30% (11/57) | Non-Toxin | 0.98 | Consensus (comb.lib./smm/nn) |
| 720 | V Protein | SSSEVIVGISPEDEE | HLA-DQA1*04:01/DQB1*04:02 | 19.30% (11/57) | Non-Toxin | 0.29 | Consensus (comb.lib./smm/nn) |
| 721 | V Protein | SSSEVIVGISPEDEE | HLA-DQA1*03:01/DQB1*03:02 | 19.30% (11/57) | Non-Toxin | 0.82 | Consensus (comb.lib./smm/nn) |
| 722 | V Protein | TSRNVNLDSIKLYTS | HLA-DRB1*03:01 | 19.30% (11/57) | Non-Toxin | 0.76 | Consensus (smm/nn/sturniolo) |
| 723 | V Protein | VCLVSDAKMLSYAPE | HLA-DRB1*03:01 | 19.30% (11/57) | Non-Toxin | 0.01 | Consensus (smm/nn/sturniolo) |
| 724 | V Protein | VCLVSDAKMLSYAPE | HLA-DRB1*11:01 | 19.30% (11/57) | Non-Toxin | 0.7 | Consensus (smm/nn/sturniolo) |
| 725 | V Protein | VNLDSIKLYTSDDEE | HLA-DRB1*03:01 | 100.00% (57/57) | Non-Toxin | 0.76 | Consensus (smm/nn/sturniolo) |
| 726 | V Protein | VSDAKMLSYAPEIAV | HLA-DRB1*09:01 | 19.30% (11/57) | Non-Toxin | 0.55 | Consensus (comb.lib./smm/nn) |
| 727 | V Protein | VVTDVVYHDHGGECT | HLA-DRB3*01:01 | 98.25% (56/57) | Non-Toxin | 0.21 | Consensus (comb.lib./smm/nn) |
| 728 | V Protein | WDGKRAWVEEWCNPA | HLA-DQA1*01:01/DQB1*05:01 | 87.72% (50/57) | Non-Toxin | 0.86 | Consensus (comb.lib./smm/nn) |
| 729 | V Protein | YTSDDEEADQLEFED | HLA-DQA1*03:01/DQB1*03:02 | 100.00% (57/57) | Non-Toxin | 0.72 | Consensus (comb.lib./smm/nn) |
| **S.No.** | **Protein** | **Peptide** | **Allele** | **Conservancy** | **Toxicity** | **Percentile rank** | **Method used** |
| 730 | W Protein | AEGSDDIQLDPVVTD | HLA-DRB1*03:01 | 100.00% (39/39) | Non-Toxin | 0.13 | Consensus (smm/nn/sturniolo) |
| 731 | W Protein | AKMLSYAPEIAVSKE | HLA-DRB1*09:01 | 20.51% (8/39) | Non-Toxin | 0.55 | Consensus (comb.lib./smm/nn) |
| 732 | W Protein | CLVSDAKMLSYAPEI | HLA-DRB1*03:01 | 20.51% (8/39) | Non-Toxin | 0.02 | Consensus (smm/nn/sturniolo) |
| 733 | W Protein | DAKMLSYAPEIAVSK | HLA-DRB1*09:01 | 20.51% (8/39) | Non-Toxin | 0.54 | Consensus (comb.lib./smm/nn) |
| 734 | W Protein | DDIQLDPVVTDVVYH | HLA-DRB1*03:01 | 97.44% (38/39) | Non-Toxin | 0.13 | Consensus (smm/nn/sturniolo) |
| 735 | W Protein | DIQLDPVVTDVVYHD | HLA-DRB3*01:01 | 97.44% (38/39) | Non-Toxin | 0.02 | Consensus (comb.lib./smm/nn) |
| 736 | W Protein | DIQLDPVVTDVVYHD | HLA-DRB1*03:01 | 97.44% (38/39) | Non-Toxin | 0.27 | Consensus (smm/nn/sturniolo) |
| 737 | W Protein | DKLELVNDGLNIIDF | HLA-DRB1*03:01 | 100.00% (39/39) | Non-Toxin | 0.57 | Consensus (smm/nn/sturniolo) |
| 738 | W Protein | DPVVTDVVYHDHGGE | HLA-DRB3*01:01 | 97.44% (38/39) | Non-Toxin | 0.02 | Consensus (comb.lib./smm/nn) |
| 739 | W Protein | DPVVTDVVYHDHGGE | HLA-DRB1*03:01 | 97.44% (38/39) | Non-Toxin | 0.6 | Consensus (smm/nn/sturniolo) |
| 740 | W Protein | DSIKLYTSDDEEADQ | HLA-DRB3*01:01 | 100.00% (39/39) | Non-Toxin | 0.28 | Consensus (comb.lib./smm/nn) |
| 741 | W Protein | EFEDEFAGSSSEVIV | HLA-DRB1*07:01 | 97.44% (38/39) | Non-Toxin | 0.56 | Consensus (comb.lib./smm/nn) |
| 742 | W Protein | EGSDDIQLDPVVTDV | HLA-DRB1*03:01 | 100.00% (39/39) | Non-Toxin | 0.13 | Consensus (smm/nn/sturniolo) |
| 743 | W Protein | GIPIKKAQTRNIHLL | HLA-DRB1*07:01 | 5.13% (2/39) | Non-Toxin | 0.53 | Consensus (comb.lib./smm/nn) |
| 744 | W Protein | GNVCLVSDAKMLSYA | HLA-DRB1*03:01 | 20.51% (8/39) | Non-Toxin | 0.01 | Consensus (smm/nn/sturniolo) |
| 745 | W Protein | GSDDIQLDPVVTDVV | HLA-DRB1*03:01 | 100.00% (39/39) | Non-Toxin | 0.13 | Consensus (smm/nn/sturniolo) |
| 746 | W Protein | IKLYTSDDEEADQLE | HLA-DRB3*01:01 | 100.00% (39/39) | Non-Toxin | 0.27 | Consensus (comb.lib./smm/nn) |
| 747 | W Protein | IQLDPVVTDVVYHDH | HLA-DRB3*01:01 | 97.44% (38/39) | Non-Toxin | 0.02 | Consensus (comb.lib./smm/nn) |
| 748 | W Protein | IQLDPVVTDVVYHDH | HLA-DRB1*03:01 | 97.44% (38/39) | Non-Toxin | 0.26 | Consensus (smm/nn/sturniolo) |
| 749 | W Protein | KLELVNDGLNIIDFI | HLA-DRB1*03:01 | 100.00% (39/39) | Non-Toxin | 0.53 | Consensus (smm/nn/sturniolo) |
| 750 | W Protein | KLYTSDDEEADQLEF | HLA-DRB3*01:01 | 100.00% (39/39) | Non-Toxin | 0.28 | Consensus (comb.lib./smm/nn) |
| 751 | W Protein | LDPVVTDVVYHDHGG | HLA-DRB3*01:01 | 97.44% (38/39) | Non-Toxin | 0.02 | Consensus (comb.lib./smm/nn) |
| 752 | W Protein | LDPVVTDVVYHDHGG | HLA-DRB1*03:01 | 97.44% (38/39) | Non-Toxin | 0.56 | Consensus (smm/nn/sturniolo) |
| 753 | W Protein | LDSIKLYTSDDEEAD | HLA-DRB3*01:01 | 100.00% (39/39) | Non-Toxin | 0.27 | Consensus (comb.lib./smm/nn) |
| 754 | W Protein | LEFEDEFAGSSSEVI | HLA-DRB1*07:01 | 100.00% (39/39) | Non-Toxin | 0.45 | Consensus (comb.lib./smm/nn) |

| | | | | | | | |
|---|---|---|---|---|---|---|---|
| 755 | W Protein | LELVNDGLNIIDFIQ | HLA-DRB1*03:01 | 100.00% (39/39) | Non-Toxin | 0.55 | Consensus (smm/nn/sturniolo) |
| 756 | W Protein | LVSDAKMLSYAPEIA | HLA-DRB1*03:01 | 20.51% (8/39) | Non-Toxin | 0.02 | Consensus (smm/nn/sturniolo) |
| 757 | W Protein | MDKLELVNDGLNIID | HLA-DRB1*03:01 | 100.00% (39/39) | Non-Toxin | 0.59 | Consensus (smm/nn/sturniolo) |
| 758 | W Protein | NGNVCLVSDAKMLSY | HLA-DRB1*03:01 | 20.51% (8/39) | Non-Toxin | 0.01 | Consensus (smm/nn/sturniolo) |
| 759 | W Protein | NNGNVCLVSDAKMLS | HLA-DRB1*03:01 | 20.51% (8/39) | Non-Toxin | 0.01 | Consensus (smm/nn/sturniolo) |
| 760 | W Protein | NVCLVSDAKMLSYAP | HLA-DRB1*03:01 | 20.51% (8/39) | Non-Toxin | 0.01 | Consensus (smm/nn/sturniolo) |
| 761 | W Protein | PVVTDVVYHDHGGEC | HLA-DRB3*01:01 | 97.44% (38/39) | Non-Toxin | 0.2 | Consensus (comb.lib./smm/nn) |
| 762 | W Protein | QLDPVVTDVVYHDHG | HLA-DRB3*01:01 | 97.44% (38/39) | Non-Toxin | 0.02 | Consensus (comb.lib./smm/nn) |
| 763 | W Protein | QLDPVVTDVVYHDHG | HLA-DRB1*03:01 | 97.44% (38/39) | Non-Toxin | 0.52 | Consensus (smm/nn/sturniolo) |
| 764 | W Protein | QPPYHWSIERSISPD | HLA-DRB1*09:01 | 20.51% (8/39) | Non-Toxin | 0.56 | Consensus (comb.lib./smm/nn) |
| 765 | W Protein | SDAKMLSYAPEIAVS | HLA-DRB1*09:01 | 20.51% (8/39) | Non-Toxin | 0.55 | Consensus (comb.lib./smm/nn) |
| 766 | W Protein | SDDIQLDPVVTDVVY | HLA-DRB1*03:01 | 97.44% (38/39) | Non-Toxin | 0.13 | Consensus (smm/nn/sturniolo) |
| 767 | W Protein | SIKLYTSDDEEADQL | HLA-DRB3*01:01 | 100.00% (39/39) | Non-Toxin | 0.27 | Consensus (comb.lib./smm/nn) |
| 768 | W Protein | SSEVIVGISPEDEEP | HLA-DQA1*04:01/DQB1*04:02 | 20.51% (8/39) | Non-Toxin | 0.39 | Consensus (comb.lib./smm/nn) |
| 769 | W Protein | SSSEVIVGISPEDEE | HLA-DQA1*04:01/DQB1*04:02 | 20.51% (8/39) | Non-Toxin | 0.29 | Consensus (comb.lib./smm/nn) |
| 770 | W Protein | TRNIHLLGRKTCLGR | HLA-DRB1*11:01 | 92.31% (36/39) | Non-Toxin | 0.44 | Consensus (smm/nn/sturniolo) |
| 771 | W Protein | VCLVSDAKMLSYAPE | HLA-DRB1*03:01 | 20.51% (8/39) | Non-Toxin | 0.01 | Consensus (smm/nn/sturniolo) |
| 772 | W Protein | VSDAKMLSYAPEIAV | HLA-DRB1*09:01 | 20.51% (8/39) | Non-Toxin | 0.55 | Consensus (comb.lib./smm/nn) |
| 773 | W Protein | VVTDVVYHDHGGECT | HLA-DRB3*01:01 | 97.44% (38/39) | Non-Toxin | 0.21 | Consensus (comb.lib./smm/nn) |

**Supplementary Table S4. Homology models of full length protein tertiary structure of NIPAH virus.** Nipah protein sequences were retrieved from NCBI. The homology modeling tools SwissModel and I-Tasser were used.

| S.No | NIPAH-CoV Protein | Templet used | Protein length | Tools used | QMEAN | C-Score | Estimated TM-score | Estimated RMSD | Iden1 |
|---|---|---|---|---|---|---|---|---|---|
| 1 | C Protein | 3dyjA | Full length | I-TASSER | - | -2.88 | 0.39±0.13 | 11.5±4.5Å | 0.14 |
| 2 | Fusion Protein | 5evmA | Full length | I-TASSER | - | -0.53 | 0.65±0.13 | 8.7±4.6Å | 1 |
| 3 | Glycoprotein | 3d11A | Full length | I-TASSER | - | -0.83 | 0.61±0.14 | 9.7±4.6Å | 0.95 |
| 4 | Matrix Protein | 4g1gA | Full length | I-TASSER | - | 0.18 | 0.74±0.11 | 6.1±3.8Å | 0.2 |
| 5 | Nucleocapsid | 6jc3A | Full length | I-TASSER | - | -0.88 | 0.60±0.14 | 9.5±4.6Å | 0.31 |
| 6 | Phosphoprotein | 4n5bA | Full length | I-TASSER | - | -1.29 | 0.55±0.15 | 11.2±4.6Å | 1 |
| 7 | Polymerase | 6v85.1.A | Full length | SwissModel | -6.06 | - | - | - | - |
| 8 | V protein | 2b5lA | Full length | I-TASSER | - | -1.09 | 0.58±0.14 | 9.6±4.6Å | 0.26 |
| 9 | W protein | 4co6A | Full length | I-TASSER | - | -0.75 | 0.62±0.14 | 8.8±4.6Å | 1 |

**Supplementary Table S5:** HLA alleles covered by the overlapping CTL and HTL epitopes.

| S.No. | HLA alleles | Class of HLA alleles |
|---|---|---|
| 1 | HLA-A*01:01 | HLA Class I alleles |
| 2 | HLA-A*02:01 | |
| 3 | HLA-A*02:03 | |
| 4 | HLA-A*02:06 | |
| 5 | HLA-A*03:01 | |
| 6 | HLA-A*11:01 | |

| | | |
|---|---|---|
| 7 | HLA-A*23:01 | |
| 8 | HLA-A*24:02 | |
| 9 | HLA-A*26:01 | |
| 10 | HLA-A*30:01 | |
| 11 | HLA-A*30:02 | |
| 12 | HLA-A*31:01 | |
| 13 | HLA-A*32:01 | |
| 14 | HLA-A*33:01 | |
| 15 | HLA-A*68:01 | |
| 16 | HLA-A*68:02 | |
| 17 | HLA-B*07:02 | |
| 18 | HLA-B*08:01 | |
| 19 | HLA-B*15:01 | |
| 20 | HLA-B*35:01 | |
| 21 | HLA-B*40:01 | |
| 22 | HLA-B*44:02 | |
| 23 | HLA-B*44:03 | |
| 24 | HLA-B*51:01 | |
| 25 | HLA-B*53:01 | |
| 26 | HLA-B*57:01 | |
| 27 | HLA-B*58:01 | |
| 28 | HLA-DPA1*01/DPB1*04:01 | HLA Class II alleles |
| 29 | HLA-DPA1*01:03/DPB1*02:01 | |
| 30 | HLA-DPA1*02:01/DPB1*01:01 | |
| 31 | HLA-DPA1*02:01/DPB1*14:01 | |
| 32 | HLA-DPA1*03:01/DPB1*04:02 | |
| 33 | HLA-DQA1*01:01/DQB1*05:01 | |
| 34 | HLA-DQA1*01:02/DQB1*06:02 | |
| 35 | HLA-DQA1*03:01/DQB1*03:02 | |
| 36 | HLA-DQA1*04:01/DQB1*04:02 | |
| 37 | HLA-DQA1*05:01/DQB1*02:01 | |
| 38 | HLA-DQA1*05:01/DQB1*03:01 | |
| 39 | HLA-DRB1*01:01 | |
| 40 | HLA-DRB1*03:01 | |
| 41 | HLA-DRB1*04:01 | |
| 42 | HLA-DRB1*04:05 | |
| 43 | HLA-DRB1*07:01 | |

| | |
|---|---|
| 44 | HLA-DRB1*08:02 |
| 45 | HLA-DRB1*09:01 |
| 46 | HLA-DRB1*11:01 |
| 47 | HLA-DRB1*13:02 |
| 48 | HLA-DRB1*15:01 |
| 49 | HLA-DRB3*01:01 |
| 50 | HLA-DRB3*02:02 |
| 51 | HLA-DRB4*01:01 |
| 52 | HLA-DRB5*01:01 |

**Supplementary Table S6:** Population coverage by all the overlapping CTL and HTL epitopes forming epitope clusters.

| Population/area | Class combined | | |
|---|---|---|---|
| | coveragea | average_hitb | pc90c |
| Algeria | 75.83% | 49.09 | 2.07 |
| American Samoa | 98.75% | 123.14 | 58.71 |
| Argentina | 99.04% | 148.28 | 63.93 |
| Australia | 95.84% | 131.68 | 58.36 |
| Austria | 99.99% | 249.29 | 148.97 |
| Belarus | 43.81% | 19.06 | 1.07 |
| Belgium | 99.87% | 197.29 | 103.64 |
| Bolivia | 77.82% | 27.34 | 7.66 |
| Borneo | 42.09% | 16.89 | 3.11 |
| Brazil | 98.55% | 164.55 | 73.95 |
| Bulgaria | 99.75% | 193.33 | 104.4 |
| Burkina Faso | 67.18% | 58.94 | 14.32 |
| Cameroon | 93.95% | 149.53 | 54.78 |
| Canada | 27.22% | 12.32 | 0.69 |
| Cape Verde | 99.83% | 232.79 | 125.86 |
| Central Africa | 94.14% | 140.31 | 34.19 |
| Central African Republic | 77.51% | 57.41 | 2.22 |
| Central America | 53.29% | 24.9 | 1.07 |
| Chile | 98.57% | 158.04 | 67.81 |
| China | 97.61% | 146.36 | 59.33 |
| Colombia | 56.35% | 28.97 | 1.15 |
| Congo | 63.54% | 31.88 | 1.37 |
| Cook Islands | 62.91% | 30 | 1.35 |
| Costa Rica | 24.31% | 7.86 | 2.38 |
| Croatia | 99.93% | 224.31 | 126 |
| Cuba | 99.84% | 207.24 | 108.55 |
| Czech Republic | 99.84% | 218.88 | 117.13 |
| Denmark | 87.68% | 38.39 | 4.06 |
| East Africa | 96.51% | 156.14 | 63.27 |
| East Asia | 99.61% | 186.19 | 81.45 |
| Ecuador | 89.16% | 67.36 | 19.38 |
| England | 100.00% | 259.61 | 159.85 |
| Equatorial Guinea | 43.90% | 24.85 | 0.89 |

| | | | |
|---|---|---|---|
| Ethiopia | 82.77% | 40.17 | 3.48 |
| Europe | 99.95% | 234.55 | 135.3 |
| Fiji | 75.74% | 45.68 | 2.06 |
| Finland | 100.00% | 245.18 | 142.03 |
| France | 99.97% | 239.02 | 139.04 |
| Gabon | 41.78% | 20.76 | 0.86 |
| Georgia | 99.57% | 206.5 | 104.32 |
| Germany | 99.99% | 258.01 | 156.1 |
| Greece | 65.84% | 36.83 | 1.46 |
| Guatemala | 52.58% | 25.98 | 1.05 |
| Guinea-Bissau | 98.93% | 188.56 | 82.62 |
| Hong Kong | 96.05% | 120.67 | 58.65 |
| India | 97.17% | 146.09 | 58.74 |
| Indonesia | 92.82% | 111.45 | 25.26 |
| Iran | 99.10% | 186.13 | 78.38 |
| Ireland Northern | 100.00% | 264.94 | 165.96 |
| Ireland South | 99.99% | 261.39 | 160.81 |
| Israel | 96.51% | 145.07 | 58.8 |
| Italy | 99.85% | 257.55 | 141.32 |
| Ivory Coast | 67.75% | 55.21 | 13.33 |
| Jamaica | 27.41% | 10.06 | 0.83 |
| Japan | 99.61% | 180.01 | 79.85 |
| Jordan | 95.62% | 117.51 | 50.03 |
| Kenya | 89.56% | 111.28 | 33.52 |
| Korea; South | 99.68% | 193.37 | 89.99 |
| Lebanon | 69.40% | 40.22 | 1.63 |
| Macedonia | 74.69% | 53.35 | 1.98 |
| Malaysia | 91.87% | 105.12 | 19.88 |
| Mali | 96.02% | 147.68 | 63.78 |
| Martinique | 77.11% | 54.36 | 2.18 |
| Mexico | 99.08% | 171.17 | 75.24 |
| Mongolia | 98.98% | 141.56 | 64.56 |
| Morocco | 99.77% | 212.65 | 108.95 |
| Nauru | 37.40% | 14.64 | 0.96 |
| Netherlands | 82.53% | 48.92 | 2.86 |
| New Caledonia | 99.66% | 145.08 | 64.83 |
| New Zealand | 63.88% | 32.43 | 1.38 |
| Niue | 40.56% | 17.37 | 3.53 |
| North Africa | 98.98% | 182.84 | 80.96 |
| North America | 99.87% | 218.51 | 116.1 |
| Northeast Asia | 97.67% | 146.82 | 59.36 |
| Norway | 93.75% | 54.36 | 5.81 |
| Oceania | 97.51% | 128.14 | 58.58 |
| Oman | 99.69% | 171.64 | 102.28 |
| Pakistan | 97.13% | 105.39 | 59.06 |
| Papua New Guinea | 99.30% | 117.19 | 58.78 |
| Peru | 100.00% | 161.92 | 75.69 |
| Philippines | 96.36% | 110.68 | 58.28 |
| Poland | 99.96% | 231.67 | 135.44 |
| Portugal | 99.70% | 204.4 | 102.59 |
| Romania | 99.67% | 178 | 98.67 |

| | | | |
|---|---|---|---|
| Russia | 99.79% | 188.78 | 96.93 |
| Rwanda | 70.09% | 49.73 | 1.67 |
| Samoa | 71.25% | 28.82 | 1.74 |
| Sao Tome and Principe | 98.89% | 194.11 | 82.59 |
| Saudi Arabia | 99.65% | 186.61 | 95.11 |
| Scotland | 96.52% | 95.32 | 23.08 |
| Senegal | 96.92% | 163.73 | 65.82 |
| Serbia | 73.37% | 61.35 | 13.15 |
| Singapore | 97.24% | 140.39 | 58.99 |
| Slovenia | 83.68% | 47.94 | 3.06 |
| South Africa | 95.27% | 151.25 | 58.34 |
| South America | 95.02% | 123.47 | 43.49 |
| South Asia | 98.61% | 162.24 | 66.72 |
| Southeast Asia | 97.51% | 140.12 | 58.91 |
| Southwest Asia | 95.73% | 138.47 | 58.57 |
| Spain | 97.47% | 141.95 | 48.85 |
| Sri Lanka | 52.39% | 43.15 | 12.39 |
| Sudan | 97.47% | 166.24 | 63.94 |
| Sweden | 100.00% | 260.91 | 157.95 |
| Taiwan | 99.22% | 163.27 | 75.28 |
| Thailand | 96.75% | 137.1 | 59.11 |
| Tokelau | 20.79% | 4.08 | 2.27 |
| Tonga | 62.79% | 28.86 | 4.84 |
| Tunisia | 99.33% | 181.42 | 82.85 |
| Turkey | 85.87% | 78.2 | 3.54 |
| Uganda | 94.87% | 135.67 | 58.56 |
| Ukraine | 50.64% | 23.43 | 1.22 |
| United Arab Emirates | 35.19% | 41.36 | 7.56 |
| United States | 99.88% | 219.7 | 117.86 |
| Venezuela | 90.04% | 87.35 | 47.01 |
| Vietnam | 96.12% | 133.53 | 58.42 |
| West Africa | 98.37% | 177.28 | 75.18 |
| West Indies | 99.65% | 201.68 | 105.88 |
| Zambia | 98.10% | 159.01 | 73.61 |
| Zimbabwe | 97.65% | 168.43 | 66.45 |
| **World** | **99.71%** | **201.5** | **98.56** |
| **Average** | **85.79** | **127.56** | **56.91** |
| **Standard deviation** | **20.73** | **74.82** | **47.14** |

Note: Following allele(s) were not available at IEDB "Population coverage" tool database, and therefore not included in the calculation, please note that allele names are case sensitive:DPA1*01, DRB5*01:01, DRB3*02:02, DRB3*01:01, DRB4*01:01.
**a:** projected population coverage; **b:** average number of epitope hits / HLA combinations recognized by the population; **c:** minimum number of epitope hits / HLA combinations recognized by 90% of the population

**Supplementary Table S7:** Construct of CTL-MPV-1, CTL-MPV-2, CTL-MPV-3, HTL-MPV-1, and HTL-MPV-2. Physicochemical property analysis based on the amino acid sequences of all the designed three CTL and two HTL Multi-Patch Vaccines.

| S.No. | Vaccine | Vaccine constructs |
|---|---|---|
| 1 | CTL-MPV-1 construct<br><br>(Comprising of identified CTL Ag-PATCHES from c Protein, Fusion Protein, Glycoprotein, Matrix Protein, Nucleoprotein, Phosphoprotein of NiV): | GIGDPVTCLKSGAICHPVFCPRRYKQIGTCGLPGTKCCKKPEAAAKRMMEVLKEEAWRIYPVLLPQMELLGGGGSITHQEQTMGMYVLYLMQRCCPMLPKLQFLKKIGKLGGGGSISECSVGILHYEKLSKIGLVKGVTRKYKIKSNPLTKDIVIKMIPNVSNMGGGGSESDSITGQIIYVDLSSYYIIVRVYFPILTEIQQAYIQELLPVSFNNDNSEWISIVPNFILVRNTLGGGGSASDKGKNPSKVIKSYYGTMDIKKINEGLLDSKILSAFNTVGGGGSEVLDRGDEVPSLFMTNVWTPSNPNTVYHCSAVYNNEFYYVGGGGSEENSKIVFIEISDQRLSIGSPSKIYDSLGQPVFYQASFSWDTMIKFGDVQTVNPLVVNWRGGGGSSESMEGVSDFSPSSWEHGGYLDKVEPEIDENGSMIPKYKIYTPGANERKYNNYMYLICYGFGGGGSIQLDKHQALRIFFLSITKLNDSGIYMIPRTMLEFGGGGSAESMKVGAAFTLISMYSERPGALIRSLLNDPDIEAVGGGGSEEMEGLMRILKTARDSSKGKTPFVDSRAYGLGGGGSITIEAQIWILIAKAVTAPDTAEESETRRWAKYVQQKRVNPFFALTQQWGGGGSNTIKSLMLLYREIGPRAPYMVLLEESIQTKFAPGGYPLLWSFAMGVATTIDRSMGALNINRGYLEPMYFRLGGGGSKYIMPSDDFSNTFFPHDTDRLNYGGGGSSVLMGVINSIKLINLDMRLNHIEEQVKEIPKIINKLESIDRVLAKTNTALSTIEGHLVSMEAAAKGIINTLQKYYCRVRGGRCAVLSCLPKEEQIGKCSTRGRKCCRRKKHHHHHH |
| 2 | CTL-MPV-2 construct<br><br>(Comprising of identified CTL Ag-PATCHES from Polymerase, V Protein,W Protein of NiV): | GIGDPVTCLKSGAICHPVFCPRRYKQIGTCGLPGTKCCKKPEAAAKKSDKTGKKCYYLTPEMVLMYCDVLEGRMMMETTVKSDIKYQPLISRSNALWGLIDPLFPVMGNRIYNIVSMIEPLVLALGGGGSFTDQKIRSMFIDDLLSILNIDNIHLLAEFFSFFRTFGHPILEAKGGGGSLTIDDCVKNWESFCGIQFDCFMELKLDSDLSMYMKDKALSPIKDEWDSVYPREVLSYGGGGSEALIASGVGKYFKENGMVKDEHELLKTLFGGGGSKTPEDDIFIHYPKGGIEGYSQKTWTIATIPFLFLSAYETNTRIAAIGGGGSTQKVHPNLPYKVKKEICAKQAQLYFERLRMNLRALGGGGSDIKLGNVKRPTKALRSAIRIATVYTWAYGDNEECWGGGGSFLIVDPELFALYLGQSISIKWAFEIHHRRPRGRHTMVDLLSDLVSNTSKHTYKVLSNALSHPRVFKRGGGGSFEKLSQNLLVTSYMIYLMNWCDFGGGGSHVDTSSRSWNTSDLDFVIFYASLTYLRRGIIKQLGGGGSRPSVENHKYRRIGLNSSSCYKALNLSPLIQRYLPSGAQRLFIGEGSGSMMLLYQSTLGQSISFYGGGGSVMMEDGLLVSKIAYTPGFPISRLFNMYRSYFGLVLVCFPVYSNPDSTEVYLGGGGSAEGSDDIQLDPVVTDVVYGGGGSCLVSDAKVLSYAPEIGGGGSQQGKDAQPPYHWSIEAAAKGIINTLQKYYCRVRGGRCAVLSCLPKEEQIGKCSTRGRKCCRRKKHHHHHH |
| 3 | HTL-MPV-1 construct | GIGDPVTCLKSGAICHPVFCPRRYKQIGTCGLPGTKCCKKPEAAAKMMASILLTLFRRTKKKY |

| | | |
|---|---|---|
| | (Comprising of identified HTL Ag-PATCHES from C Protein, Fusion Protein, Glycoprotein of NiV): | RRHTDDQAFNNPASKTEQKHGGGGSCLRMMEVLKEEAWRIYPVLLPQMELLEKEGGGGS QKVQMTYNWTQWLQTLYTMIMEENVPDMDGGGGSMVVILDKRCYCNLLILILMISECSVGIG GGGSKIKSNPLTKDIVIKMIPNVSNMSQCTGGGGGSKNNTHDLVGDVRLAGVIMAGVAIGIAT AAQITAGGGGGSKQTELSLDLALSKYLSDLLFVFGPNLQDPVSNSMTIQAISQAFGGNYETLL RTLGYATEDFDDLLEGGGGSSITGQIIYVDLSSYYIIVRVYFPILTEIQQAYIQELLPVSFNNDNS EWISIVPNFILVRNTLISNIEIGFCLITKRSVICNGGGGSDTVNPSLISMLSMIILYVLSIASLCIGLI TFISFIIVEKKRNTYSRLEDRRVGGGGSDSKILSAFNTVIALLGSIVIIVMNIMIIQNYTRSTDNQA MIKDALQSIQQQIKGLADGGGGSAMDEGYFAYSHLEKIGSCSRGVSGGGGSPEVCWEGVY NDAFLIDRINWISAGVFLEAAAKGIINTLQKYYCRVRGGRCAVLSCLPKEEQIGKCSTRGRKC CRRKKHHHHHH |
| 4 | HTL-MPV-2 construct<br><br>(Comprising of identified HTL Ag-PATCHES from Matrix protein, Nucleoprotein, Phosphoprotein, Polymerase, V Protein of NiV): | GIGDPVTCLKSGAICHPVFCPRRYKQIGTCGLPGTKCCKKPEAAAKQLDKHQALRIFFLSITKL NDSGIYMIPRTMLEFRRNNAIAFNLLVYGGGGSPSIPREFMIYDDVFIDNTGRILKGGGGGST LTTKIRIFVPATNSPELRWELTLFALDVIRSPSAAEGGGGSERPGALIRSLLNDPDIEAVIIDVG SMVNGIPVMGGGGSSKGKTPFVDSRAYGLRITDMSTLVSAVITIEAQIWILGGGGSRHHAGG IDQNMANRLGLSSNQVAELAAVQETSAGRQGGGGSAEGSDDIQLDPVVTDVVYHDHGGE CTGGGGSLEQQSLFSFDNVKNFRDGSLTNEPYGAAVQLREDLIGGGGSDQKIRSMFIDDLL SILNIDNIHLLAEFFSFFRTFGGGGGSGYCINILKVIQQLLISTEFSINETLTLDVTSPISNNLGGG SHHDYNQFLILNKLLSNRRQNDGGGGSDDLKKYYQIDQPFFVPTKITSGGGGSAEGSDDIQL DPVVTDVVYHDHGGECTGGGGSNNGNVCLVSDAKMLSYAPEIAVSKEDGGGGSPQTSRN VNLDSIKLYTSDDEEADQLEFEDEFAGSSSEVIVGISPEDEEPSEAAAKGIINTLQKYYCRVRG GRCAVLSCLPKEEQIGKCSTRGRKCCRRKKHHHHHH |

**Supplementary Table S8:** INF-γ inducing POSITIVE epitopes with a score of 1 or more than 1, screened from the CTL MPVs.

| # | CTL-MPV | Sequence | Method | Result | Score |
|---|---|---|---|---|---|
| 1 | CTL-MPV-1 | QQQSYLDSGIHFGAT | SVM | POSITIVE | 0.014601 |
| 2 | CTL-MPV-1 | HWQQQSYLDSGIHFG | SVM | POSITIVE | 0.021451 |
| 3 | CTL-MPV-1 | HFGATTTAPSLSGKG | SVM | POSITIVE | 0.006054 |
| 4 | CTL-MPV-1 | EVDSAVVGIFMPAFT | SVM | POSITIVE | 0.363695 |
| 5 | CTL-MPV-1 | FARAFVEVYLFFYFW | SVM | POSITIVE | 0.349729 |
| 6 | CTL-MPV-1 | GSSVLFVTNPFYQSV | SVM | POSITIVE | 0.3411 |
| 7 | CTL-MPV-1 | GYTGYIDWFQPNISG | SVM | POSITIVE | 0.085113 |
| 8 | CTL-MPV-1 | HEEHVLQTVIPAPWY | SVM | POSITIVE | 0.559065 |
| 9 | CTL-MPV-1 | KISSLILCNFQFQCC | SVM | POSITIVE | 0.408967 |

| | | | | | |
|---|---|---|---|---|---|
| 10 | CTL-MPV-1 | RWNRSLFERVLSVAL | SVM | POSITIVE | 0.444948 |
| 11 | CTL-MPV-1 | CGSYDYRDYYENESF | SVM | POSITIVE | 0.243738 |
| 12 | CTL-MPV-1 | TVYSYIRTPVDSYLK | SVM | POSITIVE | 0.314022 |
| 13 | CTL-MPV-1 | VFKDELFTLAVNRMS | SVM | POSITIVE | 0.012104 |
| 14 | CTL-MPV-1 | VYSYIRTPVDSYLKF | SVM | POSITIVE | 0.061491 |
| 15 | CTL-MPV-1 | DSGIHFGATTTAPSL | SVM | NEGATIVE | -0.0848 |
| 16 | CTL-MPV-1 | GIHFGATTTAPSLSG | SVM | NEGATIVE | -0.06577 |
| 17 | CTL-MPV-1 | IHFGATTTAPSLSGK | SVM | NEGATIVE | -0.04903 |
| 18 | CTL-MPV-1 | LDSGIHFGATTTAPS | SVM | NEGATIVE | -0.13812 |
| 19 | CTL-MPV-1 | SGIHFGATTTAPSLS | SVM | NEGATIVE | -0.18675 |
| 20 | CTL-MPV-1 | DSGIHFGATTTAPSL | SVM | NEGATIVE | -0.0848 |
| 21 | CTL-MPV-1 | LDSGIHFGATTTAPS | SVM | NEGATIVE | -0.13812 |
| 22 | CTL-MPV-1 | WQQQSYLDSGIHFGA | SVM | NEGATIVE | -0.03573 |
| 23 | CTL-MPV-1 | SGIHFGATTTAPSLS | SVM | NEGATIVE | -0.18675 |
| 24 | CTL-MPV-1 | FGATTTAPSLSGKGN | SVM | NEGATIVE | -0.22921 |
| 25 | CTL-MPV-1 | GIHFGATTTAPSLSG | SVM | NEGATIVE | -0.06577 |
| 26 | CTL-MPV-1 | QQSYLDSGIHFGATT | SVM | NEGATIVE | -0.12639 |
| 27 | CTL-MPV-1 | SHWQQQSYLDSGIHF | SVM | NEGATIVE | -0.16041 |
| 28 | CTL-MPV-1 | IHFGATTTAPSLSGK | SVM | NEGATIVE | -0.04903 |
| 29 | CTL-MPV-1 | YLDSGIHFGATTTAP | SVM | NEGATIVE | -0.24225 |
| 30 | CTL-MPV-1 | SGIHFGATTTAPSLS | SVM | NEGATIVE | -0.18675 |
| 31 | CTL-MPV-1 | QSYLDSGIHFGATTT | SVM | NEGATIVE | -0.03631 |
| 32 | CTL-MPV-1 | SYLDSGIHFGATTTA | SVM | NEGATIVE | -0.22194 |
| 33 | CTL-MPV-1 | EDQKYSEFLGNMVNE | SVM | NEGATIVE | -0.88771 |
| **#** | **CTL-MPV** | **Sequence** | **Method** | **Result** | **Score** |
| 34 | CTL-MPV-2 | SYLGSGIHSGATTTA | SVM | POSITIVE | 0.037787 |
| 35 | CTL-MPV-2 | QQQSYLGSGIHSGAT | SVM | POSITIVE | 0.145343 |
| 36 | CTL-MPV-2 | FWSAGVTLYNITTGL | SVM | POSITIVE | 0.465067 |
| 37 | CTL-MPV-2 | ANGLDTFSGFKVDFW | SVM | POSITIVE | 0.260875 |
| 38 | CTL-MPV-2 | YLGSGIHSGATTTAP | SVM | NEGATIVE | -0.25657 |
| 39 | CTL-MPV-2 | LGSGIHSGATTTAPS | SVM | NEGATIVE | -0.03696 |
| 40 | CTL-MPV-2 | GSGIHSGATTTAPSL | SVM | NEGATIVE | -0.20969 |
| 41 | CTL-MPV-2 | QSYLGSGIHSGATTT | SVM | NEGATIVE | -0.00588 |
| 42 | CTL-MPV-2 | QQSYLGSGIHSGATT | SVM | NEGATIVE | -0.0339 |
| 43 | CTL-MPV-2 | WQQQSYLGSGIHSGA | SVM | NEGATIVE | -0.06562 |
| 44 | CTL-MPV-2 | NGLDTFSGFKVDFWS | SVM | NEGATIVE | -0.4901 |

**Supplementary Table S9:** INF-γ inducing POSITIVE epitopes with a score of 1 or more than 1, screened from the HTL MPVs.

| # | HTL-MPV | Sequence | Method | Result | Score |
|---|---|---|---|---|---|
| 1 | HTL-MPV-1 | FGLARYIKNDSNYVVK | SVM | POSITIVE | 0.152406 |
| 2 | HTL-MPV-1 | AAIAFNVYVNTKPEILDF | SVM | POSITIVE | 0.038756 |
| 3 | HTL-MPV-1 | AIAFNVYVNTKPEILDFD | SVM | POSITIVE | 0.024703 |
| 4 | HTL-MPV-1 | GLARYIKNDSNYVVKG | SVM | NEGATIVE | -0.04135 |
| 5 | HTL-MPV-1 | LARYIKNDSNYVVKGN | SVM | NEGATIVE | -0.05923 |
| 6 | HTL-MPV-1 | ARYIKNDSNYVVKGNA | SVM | NEGATIVE | -0.0928 |
| 7 | HTL-MPV-1 | DFGLARYIKNDSNYVV | SVM | NEGATIVE | -0.08624 |
| 8 | HTL-MPV-1 | AFNVYVNTKPEILDF | SVM | NEGATIVE | -0.78488 |
| 9 | HTL-MPV-1 | FNVYVNTKPEILDFD | SVM | NEGATIVE | -0.70084 |
| 10 | HTL-MPV-1 | ILDFDRLVNGMLQCV | SVM | NEGATIVE | -0.502 |
| 11 | HTL-MPV-1 | KPEILDFDRLVNGML | SVM | NEGATIVE | -0.44773 |
| 12 | HTL-MPV-1 | AFNVYVNTKPEILDFD | SVM | NEGATIVE | -0.40182 |
| 13 | HTL-MPV-1 | IAFNVYVNTKPEILDF | SVM | NEGATIVE | -0.5766 |
| 14 | HTL-MPV-1 | AIAFNVYVNTKPEILDF | SVM | NEGATIVE | -0.41184 |
| 15 | HTL-MPV-1 | IAFNVYVNTKPEILDFD | SVM | NEGATIVE | -0.17287 |
| # | HTL-MPV | Sequence | Method | Result | Score |
| 16 | HTL-MPV-2 | GIVLKDIKLGNLLLT | SVM | POSITIVE | 0.006928 |
| 17 | HTL-MPV-2 | QGIVLKDIKLGNLLL | SVM | POSITIVE | 0.076582 |
| 18 | HTL-MPV-2 | SQGIVLKDIKLGNLL | SVM | POSITIVE | 0.10677 |
| 19 | HTL-MPV-2 | HSQGIVLKDIKLGNL | SVM | POSITIVE | 0.042983 |
| 20 | HTL-MPV-2 | LHSQGIVLKDIKLGN | SVM | POSITIVE | 0.011449 |
| 21 | HTL-MPV-2 | YLHSQGIVLKDIKLG | SVM | POSITIVE | 2.047511 |
| 22 | HTL-MPV-2 | FKVYFWSAGVTLYNI | SVM | POSITIVE | 0.175602 |
| 23 | HTL-MPV-2 | VYFWSAGVTLYNITT | SVM | POSITIVE | 0.215957 |
| 24 | HTL-MPV-2 | KVYFWSAGVTLYNIT | SVM | POSITIVE | 0.131178 |
| 25 | HTL-MPV-2 | FSGFKVYFWSAGVTL | SVM | POSITIVE | 0.037046 |
| 26 | HTL-MPV-2 | SGFKVYFWSAGVTLY | SVM | POSITIVE | 1.969099 |
| 27 | HTL-MPV-2 | IVLKDIKLGNLLLTT | SVM | NEGATIVE | -0.37097 |
| 28 | HTL-MPV-2 | EYLHSQGIVLKDIKL | SVM | NEGATIVE | -0.23876 |
| 29 | HTL-MPV-2 | VLKDIKLGNLLLTTG | SVM | NEGATIVE | -0.61579 |
| 30 | HTL-MPV-2 | ANGLDTFSGFKVYFW | SVM | NEGATIVE | -0.55794 |
| 31 | HTL-MPV-2 | NGLDTFSGFKVYFWS | SVM | NEGATIVE | -0.51605 |

**Supplementary Table S10:** AllergenFP and Vaxijen analysis of MPVs. For the Vaxijen the default threshold is 0.4, and here all the MPVs have scored above 0.4, indicating potential antigenic nature.

| S.No. | MPVs | AllergenFP v.1.0 | Vaxijen | |
|---|---|---|---|---|
| 1 | CTL-MPV-1 | NON-ALLERGEN | ANTIGENS | 0.5571 |
| 2 | CTL-MPV-2 | NON-ALLERGEN | ANTIGENS | 0.5360 |
| 3 | HTL-MPV-1 | NON-ALLERGEN | ANTIGENS | 0.5455 |
| 4 | HTL-MPV-2 | NON-ALLERGEN | ANTIGENS | 0.5201 |

**Supplementary Table S11:** Parameters for the tertiary structure homology modeling of all the CTL and HTL MPVs by the I-TASSER tool.

| S.No. | MPVs | PDB hit | C-Score | TM-Score | RMSD (Å) |
|---|---|---|---|---|---|
| 1 | CTL-MPV-1 | 7abhu | -0.42 | 0.66 ± 0.13 | 9.4 ± 4.6 |
| 2 | CTL-MPV-2 | 5a22A | -2.01 | 0.47 ± 0.15 | 13.3 ± 4.1 |
| 3 | HTL-MPV-1 | 2b9bB | -3.41 | 0.34 ± 0.11 | 16.6 ±2.9 |
| 4 | HTL-MPV-2 | 7wooF | -2.00 | 0.48 ± 0.15 | 12.8 ± 4.2 |

**Supplementary Table S12:** Refinement parameter values for CTL and HTL MPV models after refinement by GalaxyRefine tool. RMSD value in Å indicated deviation from initial model. GDT-HA (global distance test-High Accuracy): backbone structure accuracy measured by GDT-HA.

| S.No. | MPV Models | GDT-HA | RMSD | MolProbity | Clash score | Poor rotamers | Rama favored |
|---|---|---|---|---|---|---|---|
| 1 | CTL-MPV-1 | 0.9461 | 0.429 | 2.235 | 17.6 | 0.7 | 91.9 |
| 2 | CTL-MPV-2 | 0.9668 | 0.370 | 2.274 | 15.4 | 0.6 | 88.7 |
| 3 | HTL-MPV-1 | 0.9555 | 0.398 | 2.709 | 33.2 | 1.1 | 84.5 |
| 4 | HTL-MPV-2 | 0.9834 | 0.330 | 2.465 | 18.4 | 0.4 | 82.3 |

**Supplementary Table S13:** B cell linear epitopes screened from CTL & HTL MPVs.

| No. | CTL-MPV-1 | Chain | Start | End | Peptide | Number of residues | Score |
|---|---|---|---|---|---|---|---|
| 1 | CTL-MPV-1 | A | 795 | 830 | GRCAVLSCLPKEEQIGKCSTRGRKCCRRKKHHHHHH | 36 | 0.909 |
| 2 | CTL-MPV-1 | A | 1 | 41 | GIGDPVTCLKSGAICHPVFCPRRYKQIGTCGLPGTKCCKKP | 41 | 0.849 |
| 3 | CTL-MPV-1 | A | 226 | 249 | KNPSKVIKSYYGTMDIKKINEGLL | 24 | 0.762 |
| 4 | CTL-MPV-1 | A | 761 | 786 | NTALSTIEGHLVSMEAAAKGIINTLQ | 26 | 0.76 |
| 5 | CTL-MPV-1 | A | 572 | 585 | VTAPDTAEESETRR | 14 | 0.752 |
| 6 | CTL-MPV-1 | A | 158 | 170 | PGANERKYNNYMY | 13 | 0.749 |
| 7 | CTL-MPV-1 | A | 262 | 287 | GGGSEVLDRGDEVPSLFMTNVWTPSN | 26 | 0.747 |
| 8 | CTL-MPV-1 | A | 478 | 498 | GGSAESMKVGAAFTLISMYSE | 21 | 0.701 |

| | | | | | | | |
|---|---|---|---|---|---|---|---|
| 9 | CTL-MPV-1 | A | 719 | 738 | GVINSIKLINLDMRLNHIEE | 20 | 0.694 |
| 10 | CTL-MPV-1 | A | 658 | 675 | TTIDRSMGALNINRGYLE | 18 | 0.69 |
| 11 | CTL-MPV-1 | A | 190 | 211 | LRIFFLSITKLNDSGIYMIPRT | 22 | 0.673 |
| 12 | CTL-MPV-1 | A | 437 | 460 | GGGGSIQLDKHQALRIFFLSITKL | 24 | 0.67 |
| 13 | CTL-MPV-1 | A | 309 | 329 | GSEENSKIVFIEISDQRLSIG | 21 | 0.669 |
| 14 | CTL-MPV-1 | A | 694 | 711 | DFSNTFFPHDTDRLNYGG | 18 | 0.664 |
| 15 | CTL-MPV-1 | A | 510 | 529 | DPDIEAVGGGGSEEMEGLMR | 20 | 0.66 |
| 16 | CTL-MPV-1 | A | 396 | 410 | LDKVEPEIDENGSMI | 15 | 0.643 |
| 17 | CTL-MPV-1 | A | 55 | 75 | EAWRIYPVLLPQMELLGGGGS | 21 | 0.633 |
| 18 | CTL-MPV-1 | A | 135 | 150 | YLDKVEPEIDENGSMI | 16 | 0.624 |
| 19 | CTL-MPV-1 | A | 622 | 637 | EIGPRAPYMVLLEESI | 16 | 0.574 |
| 20 | CTL-MPV-1 | A | 367 | 374 | VNWRGGGG | 8 | 0.534 |
| 21 | CTL-MPV-1 | A | 95 | 102 | CPMLPKLQ | 8 | 0.531 |
| **No.** | CTL-MPV-2 | **Chain** | **Start** | **End** | **Peptide** | **Number of residues** | **Score** |
| 1 | CTL-MPV-2 | A | 5 | 59 | PVTCLKSGAICHPVFCPRRYKQIGTCGLPGTKCCKKPEAAAKKSDKTGKKCYYLT | 55 | 0.801 |
| 2 | CTL-MPV-2 | A | 722 | 752 | PYHWSIEAAAKGIINTLQKYYCRVRGGRCAV | 31 | 0.78 |
| 3 | CTL-MPV-2 | A | 332 | 366 | PNLPYKVKKEICAKQAQLYFERLRMNLRALGGGGS | 35 | 0.773 |
| 4 | CTL-MPV-2 | A | 228 | 241 | VYPREVLSYGGGGS | 14 | 0.77 |
| 5 | CTL-MPV-2 | A | 488 | 534 | VTSYMIYLMNWCDFGGGGSHVDTSSRSWNTSDLDFVIFYASLTYLRR | 47 | 0.729 |
| 6 | CTL-MPV-2 | A | 259 | 289 | VKDEHELLKTLFGGGGSKTPEDDIFIHYPKG | 31 | 0.714 |
| 7 | CTL-MPV-2 | A | 754 | 767 | SCLPKEEQIGKCST | 14 | 0.713 |
| 8 | CTL-MPV-2 | A | 574 | 597 | IQRYLPSGAQRLFIGEGSGSMMLL | 24 | 0.71 |
| 9 | CTL-MPV-2 | A | 309 | 317 | LSAYETNTR | 9 | 0.691 |
| 10 | CTL-MPV-2 | A | 185 | 201 | CVKNWESFCGIQFDCFM | 17 | 0.682 |
| 11 | CTL-MPV-2 | A | 655 | 704 | YSNPDSTEVYLGGGGSAEGSDDIQLDPVVTDVVYGGGGSCLVSDAKVLSY | 50 | 0.678 |
| 12 | CTL-MPV-2 | A | 127 | 152 | GGGSFTDQKIRSMFIDDLLSILNIDN | 26 | 0.654 |
| 13 | CTL-MPV-2 | A | 600 | 609 | STLGQSISFY | 10 | 0.649 |
| 14 | CTL-MPV-2 | A | 220 | 226 | PIKDEWD | 7 | 0.618 |
| 15 | CTL-MPV-2 | A | 83 | 95 | DIKYQPLISRSNA | 13 | 0.604 |
| 16 | CTL-MPV-2 | A | 434 | 437 | RRPR | 4 | 0.603 |
| 17 | CTL-MPV-2 | A | 630 | 642 | PGFPISRLFNMYR | 13 | 0.599 |
| 18 | CTL-MPV-2 | A | 473 | 478 | RGGGGS | 6 | 0.597 |
| **No.** | HTL-MPV-1 | **Chain** | **Start** | **End** | **Peptide** | **Number of residues** | **Score** |
| 1 | HTL-MPV-1 | A | 505 | 587 | TRSTDNQAMIKDALQSIQQQIKGLADGGGGSAMDEGYFAYSHLEKIGSCSRGVSGGGGSPEVCWEGVYNDAFLIDRINWISAG | 83 | 0.839 |
| 2 | HTL-MPV-1 | A | 426 | 489 | MLSMIILYVLSIASLCIGLITFISFIIVEKKRNTYSRLEDRRVGGGGSDSKILSAFNTVIALLG | 64 | 0.815 |
| 3 | HTL-MPV-1 | A | 241 | 272 | VAIGIATAAQITAGGGGGSKQTELSLDLALSK | 32 | 0.722 |
| 4 | HTL-MPV-1 | A | 53 | 102 | LTLFRRTKKKYRRHTDDQAFNNPASKTEQKHGGGGSCLRMMEVLKEEAWR | 50 | 0.71 |
| 5 | HTL-MPV-1 | A | 369 | 379 | VSFNNDNSEWI | 11 | 0.641 |
| 6 | HTL-MPV-1 | A | 115 | 152 | EKEGGGGSQKVQMTYNWTQWLQTLYTMIMEENVPDMDG | 38 | 0.638 |
| 7 | HTL-MPV-1 | A | 173 | 185 | ILMISECSVGIGG | 13 | 0.612 |
| 8 | HTL-MPV-1 | A | 225 | 233 | HDLVGDVRL | 9 | 0.607 |
| 9 | HTL-MPV-1 | A | 212 | 222 | QCTGGGGGSKN | 11 | 0.538 |
| 10 | HTL-MPV-1 | A | 20 | 32 | CPRRYKQIGTCGL | 13 | 0.521 |
| **No.** | HTL-MPV-1 | **Chain** | **Start** | **End** | **Peptide** | **Number of residues** | **Score** |
| 1 | HTL-MPV-2 | A | 623 | 663 | CRVRGGRCAVLSCLPKEEQIGKCSTRGRKCCRRKKHHHHHH | 41 | 0.832 |
| 2 | HTL-MPV-2 | A | 1 | 43 | GIGDPVTCLKSGAICHPVFCPRRYKQIGTCGLPGTKCCKKPEA | 43 | 0.808 |
| 3 | HTL-MPV-2 | A | 542 | 578 | YAPEIAVSKEDGGGGSPQTSRNVNLDSIKLYTSDDEE | 37 | 0.755 |
| 4 | HTL-MPV-2 | A | 582 | 607 | LEFEDEFAGSSSEVIVGISPEDEEPS | 26 | 0.705 |

| | | | | | | | |
|---|---|---|---|---|---|---|---|
| 5 | HTL-MPV-2 | A | 154 | 176 | DVIRSPSAAEGGGGSERPGALIR | 23 | 0.697 |
| 6 | HTL-MPV-2 | A | 85 | 116 | IAFNLLVYGGGGSPSIPREFMIYDDVFIDNTG | 32 | 0.695 |
| 7 | HTL-MPV-2 | A | 306 | 329 | VVYHDHGGECTGGGGSLEQQSLFS | 24 | 0.674 |
| 8 | HTL-MPV-2 | A | 416 | 438 | LISTEFSINETLTLDVTSPISNN | 23 | 0.667 |
| 9 | HTL-MPV-2 | A | 463 | 481 | NDGGGGSDDLKKYYQIDQP | 19 | 0.638 |
| 10 | HTL-MPV-2 | A | 485 | 502 | PTKITSGGGGSAEGSDDI | 18 | 0.611 |
| 11 | HTL-MPV-2 | A | 137 | 144 | PATNSPEL | 8 | 0.61 |
| 12 | HTL-MPV-2 | A | 179 | 209 | LNDPDIEAVIIDVGSMVNGIPVMGGGGSSKG | 31 | 0.602 |
| 13 | HTL-MPV-2 | A | 281 | 287 | SAGRQGG | 7 | 0.578 |
| 14 | HTL-MPV-2 | A | 290 | 303 | SAEGSDDIQLDPVV | 14 | 0.578 |
| 15 | HTL-MPV-2 | A | 119 | 126 | LKGGGGGS | 8 | 0.562 |
| 16 | HTL-MPV-2 | A | 336 | 343 | FRDGSLTN | 8 | 0.508 |

**Supplementary Table S14:** B cell Discontinuous epitopes screened from CTL MPVs.

| No. | | Residues | Number of residues | Score |
|---|---|---|---|---|
| 1 | CTL-MPV-1 | A:G1, A:I2, A:G3, A:D4, A:P5, A:V6, A:T7, A:C8, A:L9, A:K10, A:S11, A:G12, A:A13, A:I14, A:C15, A:H16, A:P17, A:V18, A:F19, A:C20, A:P21, A:R22, A:R23, A:K25, A:Q26, A:I27, A:G28, A:T29, A:C30, A:G31, A:L32, A:P33, A:G34, A:T35, A:K36, A:C37, A:K39, A:K40, A:P41, A:V51, A:E55, A:A56, A:W57, A:R58, A:I59, A:Y60, A:P61, A:V62, A:L63, A:L64, A:P65, A:M67, A:E68, A:L69, A:C94, A:C95, A:P96, A:M97, A:L98, A:P99, A:L101, A:Q102, A:L136, A:D137, A:K138, A:V139, A:E140, A:P141, A:E142, A:I143, A:D144, A:E145, A:N146, A:G147, A:S148, A:I150, A:K154, A:P158, A:G159, A:A160, A:N161, A:E162, A:K164, A:Y165, A:N166, A:N167, A:Y168, A:M169, A:Y170, A:R191, A:F193, A:L195, A:S196, A:I197, A:T198, A:K199, A:L200, A:N201, A:D202, A:S203, A:G204, A:I205, A:Y206, A:M207, A:I208, A:P209, A:T211, A:F215, A:G225, A:K226, A:N227, A:P228, A:S229, A:K230, A:V231, A:I232, A:K233, A:S234, A:Y235, A:Y236, A:G237, A:T238, A:M239, A:D240, A:I241, A:K242, A:K243, A:I244, A:N245, A:E246, A:G247, A:L248, A:L249, A:K252, A:G261, A:G262, A:G263, A:G264, A:S265, A:E266, A:V267, A:L268, A:D269, A:R270, A:G271, A:D272, A:E273, A:V274, A:P275, A:S276, A:L277, A:F278, A:M279, A:T280, A:N281, A:V282, A:W283, A:T284, A:P285, A:S286, A:N287, A:E301, A:F302, A:Y303, A:G309, A:S310, A:E311, A:E312, A:N313, A:S314, A:K315, A:I316, A:V317, A:F318, A:I319, A:E320, A:I321, A:S322, A:D323, A:Q324, A:L326, A:S327, A:I328, A:G329, A:S330, A:F356, A:G357, A:V359, A:Q360, A:N363, A:P364, A:R370, A:G394, A:L396, A:D397, A:K398, A:V399, A:E400, A:P401, A:E402, A:I403, A:D404, A:E405, A:N406, A:G407, A:M409, A:I410, A:Y413, A:K414, A:T417, A:P418, A:G437, A:G438, A:G439, A:G440, A:S441, A:I442, A:Q443, A:L444, A:D445, A:H447, A:Q448, A:A449, A:L450, A:R451, A:I452, A:F453, A:F454, A:L455, A:S456, A:I457, A:T458, A:K459, A:L460, A:N461, A:D462, A:S463, A:G478, A:G479, A:S480, A:A481, A:E482, A:S483, A:M484, A:K485, A:V486, A:G487, A:A488, A:A489, A:F490, A:T491, A:L492, A:I493, A:S494, A:M495, A:Y496, A:S497, A:E498, A:P500, A:D510, A:P511, A:D512, A:I513, A:E514, A:A515, A:V516, A:G517, A:G518, A:G519, A:G520, A:S521, A:E522, A:E523, A:M524, A:E525, A:G526, A:M528, A:R529, A:V572, A:T573, A:A574, A:P575, A:D576, A:T577, A:A578, A:E579, A:E580, A:S581, A:E582, A:T583, A:R584, A:R585, A:K588, A:Y589, A:Q591, A:Q592, A:K593, A:R594, A:E622, A:I623, A:G624, A:P625, A:R626, A:A627, A:P628, A:M630, A:V631, A:L633, A:E634, A:E635, A:S636, A:I637, A:Q638, A:T658, A:T659, A:I660, A:D661, A:R662, A:S663, A:M664, A:G665, A:A666, A:L667, A:N668, A:I669, A:N670, A:R671, A:G672, A:Y673, A:L674, A:E675, A:Y678, A:D694, A:F695, A:S696, A:N697, A:T698, A:F699, A:F700, A:P701, A:H702, A:D703, A:T704, A:D705, A:R706, A:L707, A:N708, A:Y709, A:G710, A:G711, A:G712, A:G713, A:G719, A:V720, A:I721, A:N722, A:S723, A:I724, A:K725, A:L726, A:I727, A:N728, A:L729, A:D730, A:M731, A:R732, A:L733, A:N734, A:E737, A:N761, A:T762, A:A763, A:L764, A:S765, A:T766, A:I767, A:E768, A:G769, A:H770, A:L771, A:V772, A:S773, A:M774, A:E775, A:A776, A:A777, A:A778, A:K779, A:G780, A:I781, A:I782, A:N783, A:T784, A:L785, A:Q786, A:G795, A:R796, A:C797, A:A798, A:V799, A:L800, A:S801, A:C802, A:L803, A:P804, A:K805, A:E806, A:Q808, A:I809, A:G810, A:K811, A:C812, A:S813, A:T814, A:R815, A:G816, A:R817, A:K818, A:C819, A:C820, A:R821, A:K823, A:K824, A:H825, A:H826, A:H827, A:H828, A:H829, A:H830 | 429 | 0.71 |
| 2 | CTL-MPV-1 | A:G371, A:G372, A:G373, A:G374, A:S375 | 5 | 0.511 |
| 3 | CTL-MPV-1 | A:G71, A:G72, A:G73, A:G74, A:S75, A:I76 | 6 | 0.51 |
| **No.** | | **Residues** | **Number of residues** | **Score** |
| 1 | CTL-MPV-2 | A:Y723, A:H724, A:E728, A:A729, A:A730, A:A731, A:K732, A:G733, A:I734, A:I735, A:N736, A:T737, A:L738, A:Q739, A:K740, A:Y741, A:Y742, A:C743, A:R744, A:V745, A:R746, A:G747, A:G748, A:R749, A:C750, A:A751, A:V752 | 27 | 0.824 |
| 2 | CTL-MPV-2 | A:G1, A:I2, A:G3, A:P5, A:V6, A:T7, A:C8, A:L9, A:K10, A:S11, A:G12, A:A13, A:I14, A:C15, A:H16, A:P17, A:V18, A:F19, A:C20, A:P21, A:R22, A:R23, A:Y24, A:K25, A:Q26, A:I27, A:G28, A:T29, A:C30, A:G31, A:L32, A:P33, A:G34, A:T35, A:K36, A:C37, A:C38, A:K39, A:K40, A:P41, A:E42, A:A43, A:A44, A:A45, A:K46, A:K47, A:S48, A:D49, A:K50, A:T51, A:G52, A:K53, A:K54, A:C55, A:Y56, A:Y57, A:L58, A:T59 | 58 | 0.784 |
| 3 | CTL-MPV-2 | A:Y491, A:Y494, A:L495, A:M496 | 4 | 0.763 |
| 4 | CTL-MPV-2 | A:E172, A:K174, A:G178, A:T181, A:I182, A:C185, A:V186, A:K187, A:N188, A:W189, A:E190, A:S191, A:F192, A:C193, A:G194, A:I195, A:Q196, A:P220, A:I221, A:K222, A:D223, A:E224, A:W225, A:D226, A:V228, A:Y229, A:P230, A:R231, A:E232, A:V233, A:L234, A:S235, A:Y236, A:G237, A:G238, A:G239, A:G240, A:S241, A:E242, A:D261, A:H263, A:E264, A:L265, A:L266, A:K267, A:T268, A:L269, A:F270, A:G271, A:G272, A:G273, A:G274, A:S275, A:K276, A:T277, A:P278, A:E279, A:D280, A:D281, A:I282, A:F283, A:I284, A:H285, A:Y286, A:P287, A:K288, A:L309, A:S310, A:A311, A:Y312, A:E313, A:T314, A:N315, A:T316, A:H331, A:P332, A:N333, A:L334, A:P335, A:Y336, A:K337, A:V338, A:K339, A:K340, A:E341, A:I342, A:C343, A:A344, A:K345, A:Q346, A:A347, A:Q348, A:L349, A:Y350, A:F351, A:E352, A:R353, A:L354, A:R355, A:M356, A:L358, A:R359, A:A360, A:G362, A:G363, A:G364, A:G365, A:S366, A:D367, A:I368 | 110 | 0.733 |

| | | | | |
|---|---|---|---|---|
| 5 | CTL-MPV-2 | A:R434, A:R435, A:P436, A:R437, A:G438, A:W498, A:C499, A:D500, A:F501, A:G502, A:G503, A:G504, A:G505, A:S506, A:H507, A:V508, A:D509, A:T510, A:S511, A:S512, A:R513, A:S514, A:W515, A:N516, A:T517, A:S518, A:D519, A:L520, A:D521, A:F522, A:V523, A:I524, A:Y526, A:A527, A:S528, A:L529, A:T530, A:Y531, A:L532, A:R533, A:I536, A:I574, A:Q575, A:L578, A:P579, A:S580, A:G581, A:A582, A:Q583, A:R584, A:L585, A:F586, A:I587, A:G588, A:E589, A:G590, A:S591, A:G592, A:S593, A:M594, A:L596, A:L597, A:S600, A:T601, A:L602, A:G603, A:Q604, A:S605, A:I606, A:S607, A:F608, A:Y609, A:Y628, A:T629, A:P630, A:G631, A:F632, A:P633, A:I634, A:S635, A:R636, A:L637, A:F638, A:N639, A:M640, A:Y641, A:R642, A:S643, A:F652, A:S656, A:N657, A:P658, A:D659, A:S660, A:T661, A:E662, A:V663, A:Y664, A:L665, A:G666, A:G667, A:G668, A:G669, A:S670, A:A671, A:E672, A:G673, A:S674, A:D675, A:D676, A:I677, A:Q678, A:L679, A:D680, A:P681, A:V682, A:V683, A:T684, A:D685, A:V686, A:V687, A:Y688, A:G689, A:G690, A:G691, A:G692, A:S693, A:C694, A:L695, A:V696, A:S697, A:D698, A:A699, A:K700, A:V701, A:L702, A:S703, A:Y704, A:L753, A:S754, A:C755, A:L756, A:P757, A:K758, A:E759, A:E760, A:Q761, A:I762, A:G763, A:K764, A:C765, A:S766, A:T767, A:R768, A:G769, A:R770, A:C772, A:H778, A:H779 | 159 | 0.679 |
| 6 | CTL-MPV-2 | A:Q484, A:N485, A:V488, A:T489, A:M492, A:I493 | 6 | 0.657 |
| 7 | CTL-MPV-2 | A:D83, A:I84, A:K85, A:Y86, A:Q87, A:P88, A:L89, A:I90, A:S91, A:R92, A:N94, A:A95, A:G98, A:L99, A:P102, A:L103, A:P105, A:V106, A:M107, A:G108, A:R110, A:L125, A:G127, A:G128, A:G129, A:S130, A:F131, A:T132, A:D133, A:Q134, A:K135, A:I136, A:R137, A:S138, A:M139, A:F140, A:I141, A:D142, A:D143, A:L144, A:L145, A:S146, A:I147, A:L148, A:N149, A:I150, A:D151, A:N152, A:K369, A:L370, A:G371 | 51 | 0.628 |
| 8 | CTL-MPV-2 | A:G474, A:G475, A:G476, A:G477, A:S478, A:K481 | 6 | 0.626 |
| 9 | CTL-MPV-2 | A:F197, A:D198, A:C199, A:F200 | 4 | 0.614 |
| 10 | CTL-MPV-2 | A:S726, A:I727, A:K771 | 3 | 0.509 |
| 11 | CTL-MPV-2 | A:E255, A:N256, A:G257, A:V259, A:K260 | 5 | 0.507 |
| **No.** | | **Residues** | **Number of residues** | **Score** |
| 1 | HTL-MPV-1 | A:T505, A:T508, A:D509, A:N510, A:Q511, A:A512, A:M513, A:I514, A:K515, A:D516, A:A517, A:L518, A:Q519, A:S520, A:I521, A:Q522, A:Q523, A:Q524, A:I525, A:K526, A:G527, A:L528, A:A529, A:D530, A:G531, A:G532, A:G533, A:G534, A:S535, A:A536, A:M537, A:D538, A:E539, A:G540, A:Y541, A:F542, A:A543, A:Y544, A:S545, A:H546, A:L547, A:E548, A:K549, A:I550, A:G551, A:S552, A:C553, A:S554, A:R555, A:G556, A:V557, A:S558, A:G559, A:G560, A:G561, A:G562, A:S563, A:P564, A:E565, A:V566, A:C567, A:W568 | 62 | 0.878 |
| 2 | HTL-MPV-1 | A:C20, A:P21, A:R22, A:R23, A:K25, A:Q26, A:G28, A:C30, A:L32, A:V369, A:S370, A:F371, A:N372, A:N373, A:D374, A:N375, A:S376, A:E377, A:W378, A:I379, A:S425, A:M426, A:L427, A:S428, A:M429, A:I430, A:I431, A:L432, A:Y433, A:V434, A:L435, A:S436, A:I437, A:A438, A:S439, A:L440, A:C441, A:I442, A:G443, A:L444, A:I445, A:T446, A:F447, A:I448, A:S449, A:F450, A:I451, A:I452, A:V453, A:E454, A:K455, A:K456, A:R457, A:N458, A:T459, A:Y460, A:S461, A:R462, A:L463, A:E464, A:D465, A:R466, A:R467, A:V468, A:G469, A:G470, A:G471, A:G472, A:S473, A:D474, A:S475, A:K476, A:I477, A:L478, A:S479, A:A480, A:F481, A:N482, A:T483, A:V484, A:I485, A:A486, A:L487, A:L488, A:G489, A:V492 | 86 | 0.758 |
| 3 | HTL-MPV-1 | A:E569, A:G570, A:V571, A:Y572, A:N573, A:D574, A:A575, A:F576, A:L577, A:I578, A:D579, A:R580, A:I581, A:N582, A:W583, A:I584, A:S585, A:A586, A:G587, A:V588 | 20 | 0.727 |
| 4 | HTL-MPV-1 | A:L53, A:T54, A:F56, A:R57, A:R58, A:T59, A:K60, A:K61, A:K62, A:Y63, A:R64, A:R65, A:H66, A:T67, A:D68, A:D69, A:Q70, A:A71, A:F72, A:N73, A:N74, A:P75, A:A76, A:S77, A:K78, A:T79, A:E80, A:Q81, A:K82, A:H83, A:G84, A:G85, A:G86, A:G87, A:S88, A:C89, A:L90, A:R91, A:M92, A:M93, A:E94, A:V95, A:L96, A:K97, A:E98, A:E99, A:W101, A:R102, A:G215, A:G216, A:G218, A:G219, A:S220, A:K221, A:N222, A:N223, A:G240, A:V241, A:A242, A:I243, A:G244, A:I245, A:A246, A:T247, A:A248, A:A249, A:Q250, A:I251, A:T252, A:A253, A:G254, A:G255, A:G256, A:G257, A:G258, A:S259, A:K260, A:Q261, A:T262, A:E263, A:L264, A:S265, A:L266, A:D267, A:L268, A:A269, A:L270, A:S271, A:K272, A:L341, A:S342, A:S343, A:Y344 | 93 | 0.69 |
| 5 | HTL-MPV-1 | A:H225, A:D226, A:L227, A:V228, A:G229, A:D230, A:V231, A:R232 | 8 | 0.609 |
| 6 | HTL-MPV-1 | A:P109, A:E112, A:L113, A:E115, A:K116, A:E117, A:G118, A:G119, A:G120, A:G121, A:S122, A:Q123, A:K124, A:V125, A:Q126, A:M127, A:N130, A:W131, A:T132, A:Q133, A:W134, A:L135, A:Q136, A:T137, A:L138, A:Y139, A:T140, A:M141, A:I142, A:M143, A:E144, A:E145, A:N146, A:V147, A:P148, A:D149, A:M150, A:D151, A:G152, A:G155, A:M157, A:L170, A:I171, A:I173, A:L174, A:I176, A:S177, A:E178, A:C179, A:S180, A:V181, A:G182, A:I183, A:G184, A:G185, A:G187 | 56 | 0.608 |
| 7 | HTL-MPV-1 | A:P205, A:N206, A:S208, A:N209, A:Q212, A:C213 | 6 | 0.556 |
| **No.** | | **Residues** | **Number of residues** | **Score** |
| 1 | HTL-MPV-2 | A:A589, A:G590, A:S591, A:S592, A:S593, A:E594, A:V595, A:I596, A:V597, A:G598, A:I599, A:S600, A:P601, A:E602, A:D603, A:E604 | 16 | 0.799 |
| 2 | HTL-MPV-2 | A:N463, A:D464, A:G465, A:G466, A:G467, A:G468, A:S469, A:D470, A:D471, A:L472, A:K473, A:K474, A:Y475, A:Y476, A:Q477, A:I478, A:D479, A:Q480, A:P481, A:F482, A:L540, A:S541, A:Y542, A:A543, A:P544, A:E545, A:I546, A:A547, A:V548, A:S549, A:K550, A:D552, A:G553, A:G554, A:G555, A:G556, A:S557, A:P558, A:Q559, A:T560, A:S561, A:R562, A:N563, A:V564, A:N565, A:L566, A:D567, A:S568, A:I569, A:L571, A:T573, A:S574, A:D575, A:D576, A:E577, A:E578, A:C623, A:V625, A:R626, A:G627, A:G628, A:R629, A:C630, A:A631, A:V632, A:L633, A:S634, A:C635, A:L636, A:P637, A:K638, A:E639, A:E640, A:Q641, A:I642, A:G643, A:K644, A:C645, A:S646, A:T647, A:R648 | 81 | 0.761 |
| 3 | HTL-MPV-2 | A:L416, A:I417, A:S418, A:T419, A:F421, A:S422, A:I423, A:N424, A:E425, A:T426, A:L427, A:T428, A:L429, A:D430, A:V431, A:T432, A:S433, A:P434, A:S436, A:N437, A:N438, A:T489, A:S490, A:G491, A:G492, A:G493, A:G494, A:S495, A:A496, A:E497, A:G498, A:S499, A:D500, A:D501, A:I502, A:Q503, A:L504, A:G649, A:R650, A:K651, A:C652, A:C653, A:R654, A:R655, A:K656, A:K657, A:H658, A:H659, A:H660, A:H661, A:H662, A:H663 | 52 | 0.682 |
| 4 | HTL- | A:G1, A:I2, A:G3, A:D4, A:P5, A:V6, A:T7, A:C8, A:L9, A:S11, A:G12, A:A13, A:I14, A:C15, A:H16, A:P17, A:V18, A:F19, A:C20, A:P21, A:R22, A:Y24, A:K25, A:Q26, A:I27, A:G28, A:T29, A:C30, A:G31, A:L32, | 157 | 0.666 |

| | | | | |
|---|---|---|---|---|
| | MPV-2 | A:P33, A:G34, A:T35, A:K36, A:C37, A:C38, A:K39, A:K40, A:P41, A:E42, A:A43, A:K46, A:Q47, A:L48, A:D49, A:L150, A:F151, A:D154, A:V155, A:I156, A:S158, A:P159, A:S160, A:A161, A:A162, A:E163, A:G164, A:G165, A:G166, A:G167, A:S168, A:E169, A:R170, A:P171, A:G172, A:A173, A:L174, A:R176, A:L179, A:N180, A:D181, A:P182, A:D183, A:I184, A:E185, A:A186, A:V187, A:I188, A:I189, A:D190, A:V191, A:G192, A:S193, A:M194, A:V195, A:N196, A:G197, A:I198, A:P199, A:V200, A:M201, A:G202, A:G203, A:G204, A:G205, A:S206, A:S207, A:K208, A:G209, A:K210, A:V214, A:D215, A:G244, A:G245, A:G246, A:R249, A:H250, A:A252, A:G253, A:S281, A:A282, A:G283, A:R284, A:Q285, A:G286, A:G287, A:G288, A:G289, A:S290, A:A291, A:E292, A:G293, A:S294, A:D295, A:D296, A:I297, A:Q298, A:L299, A:D300, A:P301, A:V302, A:V303, A:V306, A:V307, A:Y308, A:H309, A:D310, A:H311, A:G312, A:G313, A:E314, A:C315, A:T316, A:G317, A:G318, A:G319, A:G320, A:S321, A:L322, A:E323, A:Q324, A:G339, A:S340, A:L341, A:T342, A:N343, A:D372 | | |
| 5 | HTL-MPV-2 | A:P137, A:A138, A:T139, A:N140, A:S141, A:E143, A:L144, A:E147 | 8 | 0.639 |
| 6 | HTL-MPV-2 | A:R80, A:N83, A:I85, A:A86, A:F87, A:N88, A:L89, A:L90, A:V91, A:Y92, A:G93, A:G94, A:G95, A:G96, A:S97, A:P98, A:S99, A:I100, A:P101, A:R102, A:E103, A:F104, A:M105, A:I106, A:Y107, A:D108, A:D109, A:V110, A:F111, A:I112, A:D113, A:N114, A:T115, A:G116, A:R117, A:L119, A:K120, A:G121, A:G122, A:G123, A:G124, A:G125, A:S126, A:T127, A:V136, A:N335, A:R337 | 47 | 0.62 |
| 7 | HTL-MPV-2 | A:A579, A:L582, A:E583, A:E585, A:N616, A:Q619 | 6 | 0.558 |
| 8 | HTL-MPV-2 | A:E605, A:P606, A:S607 | 3 | 0.55 |

**Supplementary Table S15:** Analysis of codon-optimized cDNA of all the MPVs.

| S.No. | MPVs | GC content | CAI (Codon Adaptation Index) score | Tandem rare codons |
|---|---|---|---|---|
| 1 | CTL-MPV-1 | 64.74% | 1 | 0% |
| 2 | CTL-MPV-2 | 66.06% | 1 | 0% |
| 3 | HTL-MPV-1 | 64.86% | 1 | 0% |
| 4 | HTL-MPV-2 | 67.96% | 1 | 0% |
| | Ideal values | 30-70% | 0.8-1.0 | <30% |
